# The landscape of high-affinity human antibodies against intratumoral antigens

**DOI:** 10.1101/2021.02.06.430058

**Authors:** Goran Rakocevic, Irina Glotova, Ines de Santiago, Berke Cagkan Toptas, Milena Popovic, Milos Popovic, Dario Leone, Andrew L. Stachyra, Raphael Rozenfeld, Deniz Kural, Daniele Biasci

**Affiliations:** Totient Inc. 1 Alewife Center Suite 120, Cambridge, MA 02140

## Abstract

High expression of immunoglobulin transcripts is often detected in human tumours and correlates with favourable clinical outcomes across different cancer types. However, the antigens recognized by such immunoglobulins (Ig) and their biological significance remain largely unknown. We computationally inferred the paired sequence of thousands of clonally expanded Ig using bulk RNA sequencing data from solid tumors in The Cancer Genome Atlas (TCGA). After expressing 283 Ig as recombinant antibodies in mammalian cells, we individually tested them against a library of twenty thousands full-length human proteins and an additional library of six thousand structurally intact membrane proteins, thus identifying their candidate target antigens. Using surface plasmon resonance we then confirmed 16 high-affinity antibodies that bind to their targets with *K*_D_ in the nanomolar range. Our work provides insights into the antigens that drive B cell responses in human cancers, while also obtaining fully human, high-affinity recombinant antibodies against them.

## Main

Transcripts encoding immunoglobulin light and heavy chains are often detected in solid tumors across different cancer types^1^, but their functional relevance remains unclear. Certain characteristics of the intratumoral Ig repertoire (e.g., transcripts abundance, clonality and number of detectable somatic mutations) have been associated with favourable clinical outcomes, such as longer overall survival and response to immune checkpoint inhibitors^1–3^. Moreover, the presence of intratumoral plasma cells and ectopic germinal centers, which are key components of the antibody selection and production machinery^4^, has been associated with longer overall survival and immunotherapy response^5–8^. Despite these observations, the contribution of intratumoral Ig to immune responses against cancer remains largely unknown.

The main obstacle to the functional characterization of intratumoral Ig is our limited understanding of their target antigens. Previous studies have demonstrated that sequences of individual Ig chains can be reconstructed using bioinformatics methods from bulk RNA sequencing data (RNA-Seq) generated by large-scale cancer genomics efforts^1,3^. Compared to specialized B-cell receptor (BCR) sequencing or single-cell sequencing, bulk RNA-seq has the significant advantage of being readily available for thousands of clinically annotated tumor samples^9^. However, previous studies have been limited to in-silico analysis: they did not attempt to pair heavy and light Ig chains, nor to express the resulting sequences as complete antibodies, two key steps required to experimentally identify their target antigens.

In this work we assembled and paired thousands of intratumoral Ig chains using legacy bulk RNA-Seq data from TCGA, one of the most comprehensive genomic studies of human cancer to date^9^ (Figure 1). We individually performed gene synthesis, mammalian expression, and purification for 283 pairs of Ig chains, obtaining high quality, fully human recombinant antibodies in most cases. We then individually screened each antibody against two large collections of recombinant proteins, covering the vast majority of the human proteome, in order to obtain the most likely binding targets. In selected cases, we confirmed binding using surface plasmon resonance (SPR) in order to characterize the binding kinetics. Our results show that fully functional antibodies can be obtained from legacy bulk tumour RNA-Seq, without the need for specialized BCR or single-cell sequencing. Using this approach, we were able to identify the target antigens of several high-affinity Ig expressed in human tumors.

**Fig. 1.**
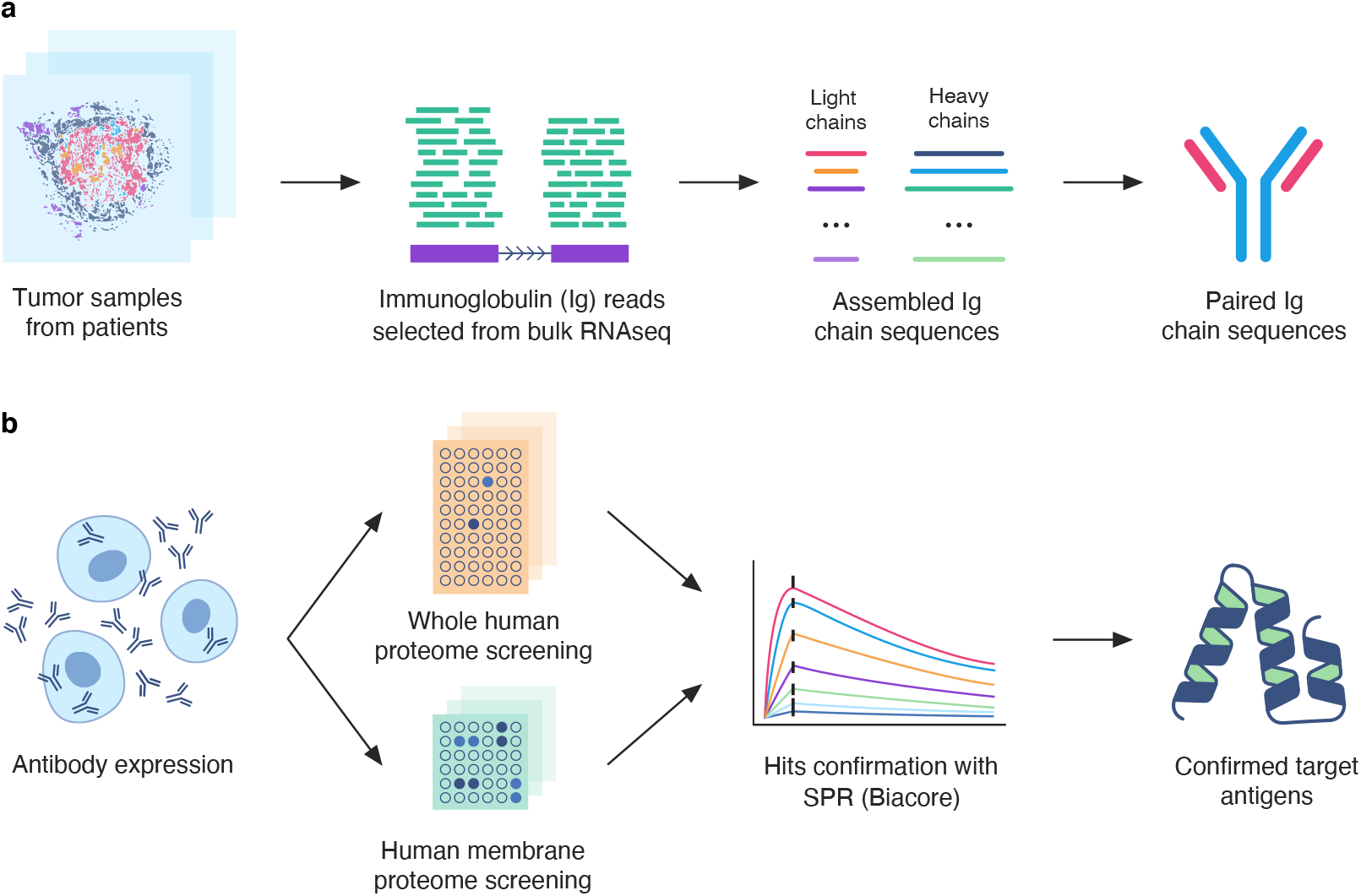
Ig reconstruction and characterization protocol. **a**. Steps of the computational workflow which starts from raw RNA sequencing data of tumor samples as the input, removes the reads mapped to non-Ig transcripts, reconstructs the Ig chain sequences, and outputs the paired sequences for which both chains satisfy the dominance threshold. **b**. Steps of the experimental workflow aimed at expressing the reconstructed Ig as recombinant antibodies, screening for their target antigens using two different human protein libraries, and confirming the result using surface plasmon resonance (SPR).

## Results

### Identification of dominant Ig sequences in the TCGA dataset

It has been observed that in some tumor samples the Ig repertoire is particularly clonal: a small set of Ig transcripts are expressed at high levels and account for the vast majority of all the reads attributable to Ig genes in that sample^1,3^. We speculated that this phenomenon, which probably results from selected B cell clones winning the competition for the limited access to T follicular helper (Tfh) cells in the germinal center (GC) reaction^4^, could be used to correctly pair heavy and light Ig chains from bulk RNA-Seq data, at least in some cases. Accordingly, we computationally assembled Ig chain sequences expressed in each TCGA tumor sample and we calculated the associated Berger–Parker index^10^ in order to identify samples with particularly dominant sequences. We observed that a small, but not insignificant, proportion of the analyzed samples expressed highly dominant Ig sequences (Supplementary Figure 2).

### Dominant Ig sequences can be correctly paired from bulk RNA-Seq in-silico

As a next step, we wanted to assess whether the dominance information captured by the Berger–Parker index could be used to correctly pair heavy and light chains from bulk RNA-Seq data. In order to do so, we used RNA-Seq data from single B cells^11^ to simulate bulk RNA-Seq samples for which the expression level, sequence, and correct pairing of all Ig chains was known. Using this approach, we established that for samples with dominance scores larger than 0.382 we could identify and correctly pair the dominant Ig chains with high accuracy (true positive rate 0.46, false positive rate 0.1, AUC=0.757; Figure 2). We used this result to select 191 paired intratumoral Ig sequences from our TCGA analysis for expression in mammalian cells and further characterization (Figure 3). As a control, we also selected for expression and characterization 92 sequences below our chosen dominance score cutoff.

**Fig. 2.**
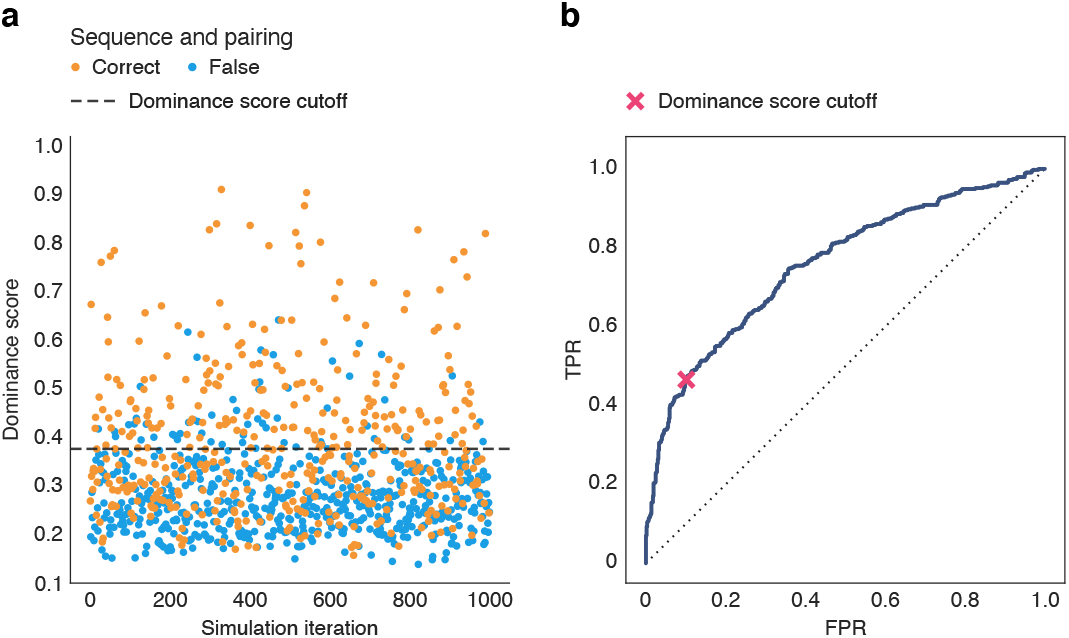
Evaluation of reconstruction performance on synthetic data. **a**. Distribution of correctly and incorrectly reconstructed samples depending on the dominance score (geometric mean of the Berger-Parker indices for heavy and light Ig chains) estimated by the workflow. Above the threshold of 0.382 the top Ig for 90% of the synthetic samples were correctly reconstructed. **b**. ROC curve for the evaluation at different values of the dominance score. The red cross marks the dominance score of 0.382 selected as the point with the highest true positive rate (0.46) where the false positive rate is below 0.1.

**Fig. 3.**
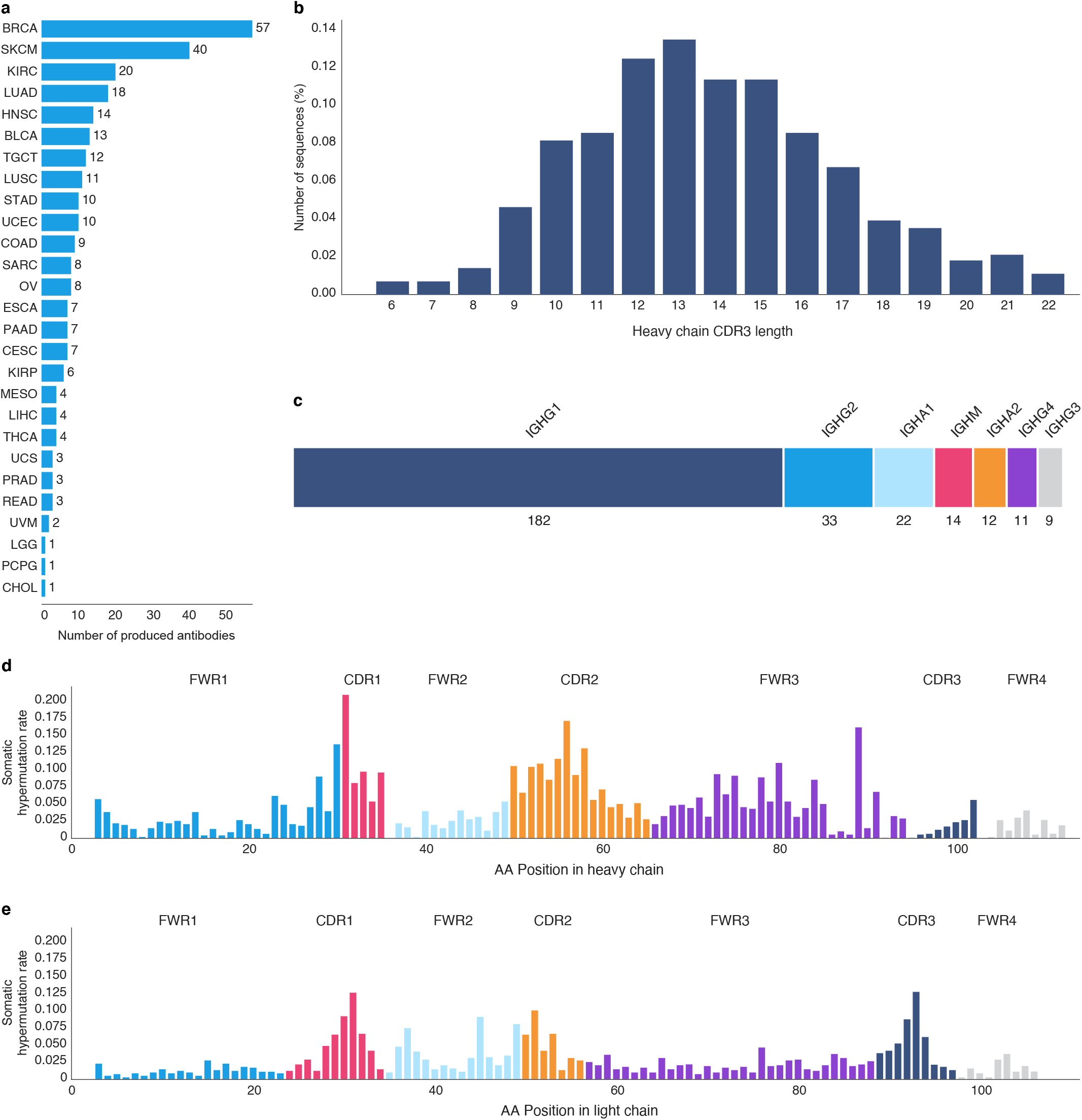
Immunoglobulin isotypes, SHM, and cancer types. **a**. Distribution of the reconstructed Ig across TCGA cancer types. **b**. Distribution of amino acid lengths of IgH CDR3 regions. **c**. Number of selected Ig by putative isotype. **d**. and **e**. Mean SHM rate for each position in the Martin numbering scheme across the heavy and light chains in the selected set of Ig. For each chain, per amino-acid mutation rates were estimated by mapping the chains to germline sequence constructs by IgBlast and numbered following the Martin numbering scheme. For chains where multiple amino acids map to the same Martin number we used the mean value across those amino acids.

### In-silico paired Ig sequences can be expressed at high levels in mammalian cells

We performed gene synthesis for 283 paired Ig sequences and attempted expression of the corresponding antibody proteins in mammalian cells (HEK293). For each candidate, we replaced the heavy constant region with a standard human IgG1 sequence in order to facilitate subsequent detection and screening. We successfully expressed 275 recombinant antibodies out of 283 with high yield (mean yield 2.35 mg/150mL; Supplementary Table S1). After expression, each antibody was purified using protein A, stored in PBS at a standard concentration of 1ug/uL and used for subsequent target antigen screening.

### Identification of target antigens using high-throughput proteomics

Having successfully expressed the intratumoral Ig we paired in-silico, we decided to use the resulting recombinant antibodies to attempt identification of their target antigens. To this aim, we used high throughput proteomics to screen individual antibodies against a large collection of wild-type human proteins. We screened 173 antibodies against a collection of about twenty thousand human proteins, each represented in duplicate on the surface of a protein array, covering about 80% of the known human proteome^12^. While protein arrays are a cost effective way to test antibodies against a large number of potential targets, this technology is not designed to display membrane proteins in their correct conformation. Complex membrane proteins are unlikely to fold correctly when immobilized on the surface of the array, outside or their native membrane context. For this reason, we used high-throughput fluorescence-activated cell sorting (HT-FACS) to screen 92 antibodies against a library of six thousand structurally-intact membrane proteins expressed in human HEK0293T cells^13^. As a result of these screenings, we were able to identify high confidence hits for 84 of our antibody candidates screened using protein arrays (48%) and 21 of candidates screened using HT-FACS (23%). Targets recognized by our in-silico paired intratumoral Ig included well known cancer-specific antigens^14^ (NY-ESO-1, MAGEA3, GAGE2A, DLL3) as well as immunomodulatory molecules expressed in the tumor microenvironment (ANXA1, TGFBI, C4BPB; Figure 4 and Supplementary tables 2, 4 and 5).

**Fig. 4.**
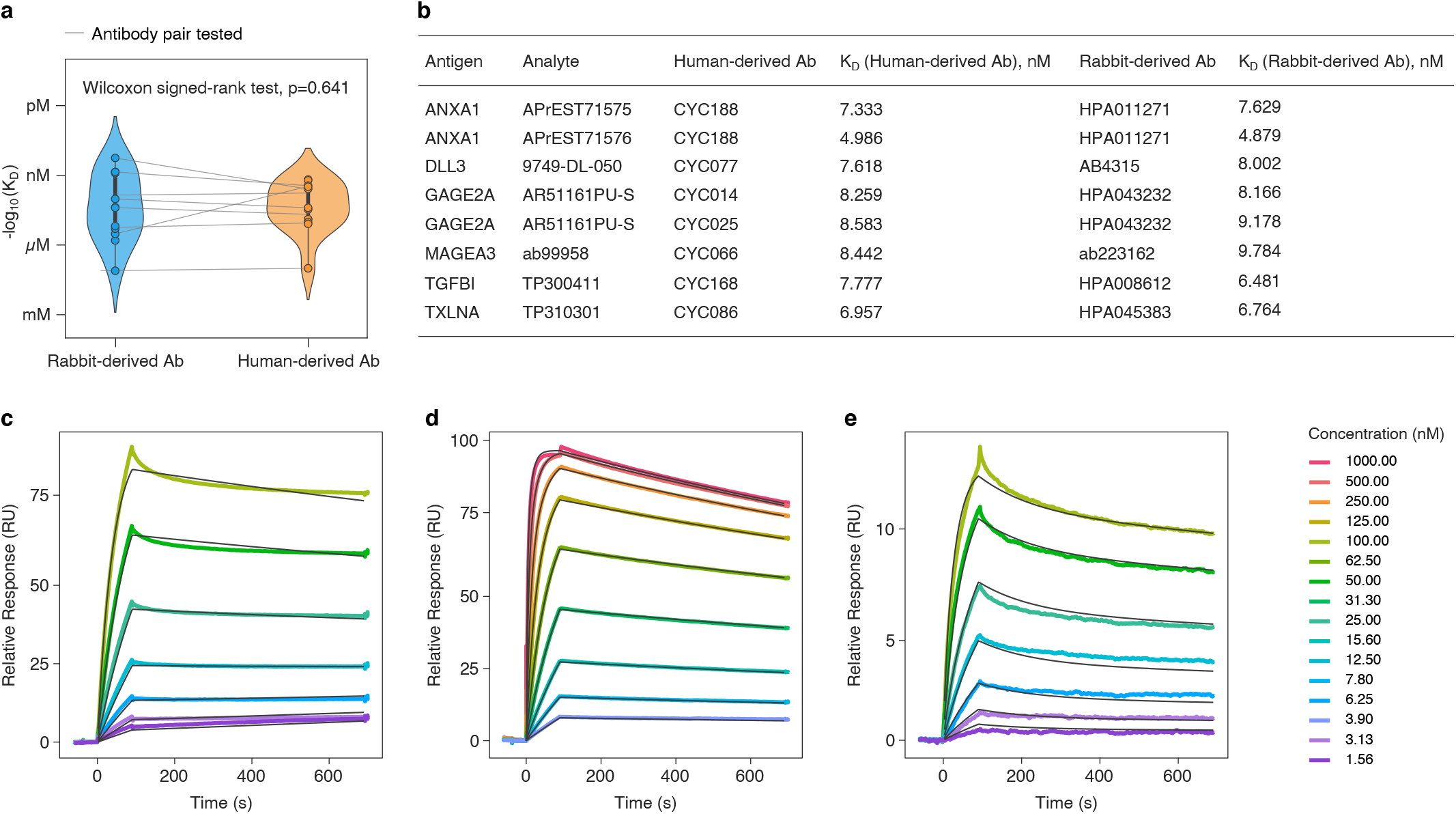
Distribution of *K*_D_ values for antibodies derived from intratumoral Ig and commercially available antibodies against the same antigens. **a**. Distribution of empirically determined *K*_D_ values for human-derived antibodies (Abs) and their rabbit-derived counterparts for the same antigens, showing no statistically significant difference using a paired t-test. As Abs have been tested in multiple experiments, to calculate the p-value, we first averaged the −log10(*K*_D_) across the experiments with a specific analyte (different source of antigen), then paired each of these averages from human-derived Abs with averages from rabbit-derived Abs for the same analyte and applied paired t-test. **b**. *K*_D_ values for human and rabbit-derived antibodies presented in a. When multiple experiments have been performed for a human or rabbit Ab, we calculated the average *K*_D_ from these experiments. For detailed information about each experiment consult Supplementary Table S2. **c**. Representative sensorgrams of SPR-determined antibody-antigen interaction for CYC214 (anti-C4BPB antibody), for **d**. CYC066 (anti-MAGEA3 antibody) and for **e.** CYC168 (ant-TGFBI antibody). Solid colored lines indicate raw data observed by Biacore 8K instrument, and overlaid solid black lines indicate the fit result estimated using the model. For additional experimental details and sensograms, see Supplementary Figure S3.

### Intratumoral Ig bind their target antigens with high-affinity

As a next step, we decided to independently confirm the interaction between selected intratumoral Ig and their putative target antigens by using surface plasmon resonance (SPR)^15^. For each intratumoral Ig sequence, we characterized the binding affinity of the corresponding recombinant antibody to its putative target antigen, having sourced the antigen independently of the vendor used for the high-throughput proteomics screening. We confirmed 19 antibody-antigen interactions (Supplementary Table S2). We observed that the recombinant antibodies derived from intratumoral Ig sequences bind to their target antigen with very high-affinity, with *K*_D_ in the low nanomolar range (Figure 4 and Supplementary Table S2). When we compared our fully human antibodies against commercial antibodies obtained from rabbits (a model organism known to generate very high-affinity antibodies^16,17^) after immunisation with the same antigen, we found no significant differences in their binding affinity (Figure 4).

### Epitope mapping for recombinant antibodies derived from intratumoral Ig

Having demonstrated that our in-silico paired intratumoral Ig bind to their target antigens with high-affinity, we decided to assess whether it would be possible to identify their epitope, at least in principle. We selected one corresponding recombinant antibody and performed epitope mapping using hydrogen/deuterium exchange mass spectrometry (HDX-MS)^18^. Using this technique, we were able to identify the putative binding region of the antibody on its designated target (C4BPB; Figure 5). We observed that the most likely epitope overlaps with the C4BPB binding site for protein S^19^, which is crucial for its biological function^20^. Although additional functional studies would be required to clarify whether or not this particular antibody is able to interfere with the binding between C4BPB and protein S, our workflow shows that in principle it is possible to identify the target antigen and the corresponding epitope for in-silico paired intratumoral Ig, thus gaining further insights into their biological function.

**Fig. 5.**
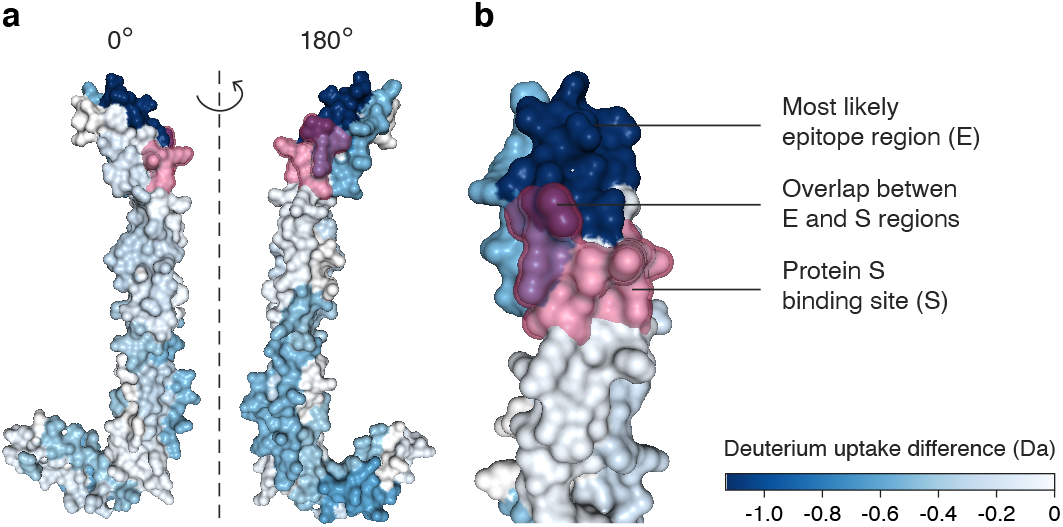
Epitope mapping results for C4BPB show overlap with protein S binding site. HDX-MS was used to measure the level of deuterium (D) uptake by C4BPB alone or in presence of CYC214 antibody. **a.** Relative D uptake difference per residue (blue shading) across the entire protein surface. **b**. Details of the protein region containing the known binding site^20^ for protein S. High D uptake difference (dark blue) is detected in the region containing the binding site for protein S, thus suggesting that CYC214 might be disrupting the interaction between C4BPB and protein S. Each C4BPB fragment is measured up to three times and each residue can be covered by one or more overlapping fragments: the uptake difference per residue was calculated using the mean of the uptake differences of the fragments covering the residue.

## Discussion

Despite the striking correlation between expression of Ig transcripts and favourable clinical outcomes in human tumors, the functional role of intratumoral Ig remains substantially unknown. In this study, we demonstrated for the first time that in-silico pairing of intratumoral Ig can be used to obtain fully functional antibodies from legacy tumor RNA sequencing data. Moreover, we showed that it is possible to use high-throughput proteomics techniques to identify their target antigens, characterize their binding kinetics, and map the corresponding epitopes. These steps are crucial to enable further functional characterization studies. Interestingly, we have demonstrated that highly clonal intratumoral Ig are selected to bind not only cancer-specific antigens (NY-ESO-1, MAGEA3, GAGE2A, DLL3), but also wild-type proteins expressed in the tumor microenvironment (ANXA1, TGFBI, C4BPB). Despite being directed against non-mutated self-antigens, these Ig bind to their target with very high-affinity, similarly to antibodies obtained by immunizing a different species with the same antigen. The importance of this observation is twofold: on one hand, it highlights the extent to which peripheral tolerance might be compromised during immune responses in cancer-affected tissues; on the other hand, it suggests that high-affinity fully human antibodies against human proteins can be obtained by sequencing Ig transcripts expressed in tissues affected by chronic inflammation. While we have not identified the target antigens for many of the antibodies produced in this study, it should be noted that our antigen screening only considered non-mutated proteins. The remaining orphan antibodies could be explained by errors in our in-silico pairing method. Alternatively, it is possible that some of these orphan antibodies bind to antigens for which we could not screen in this study, including neoantigens specific for particular patients, or non-protein antigens such as glycans^20,21^. Advancements in antigen screening methods might in the future allow the deorphaning of additional candidates. Despite these limitations, this study generated the largest collection of individually screened fully human intratumoral Ig generated so far, paves the way to improve our understanding of their functional role during anti-tumor immune responses, and suggests a novel way to extract immunological insights from legacy RNA sequencing data.

## Methods

### Ig reconstruction

For the purpose of reconstruction of Ig chain sequences from RNA sequencing data we implemented a computational workflow following the same basic idea underlying previously described approaches ^11,22^. We reduced the number of sequencing reads not originating from Ig transcripts through three filtering steps. First, we selected only the reads that either map to the Ig genes, or fail to map using Kallisto (version 0.44) ^23^ and Gencode ^24^ (version 22) protein coding sequences as the reference. Second, we mapped the remaining reads to the full human genome (version GRCh38, ^24,25^), and retained only the reads that mapped to the Ig loci, or failed to map at all. In order to extract the selected reads and revert to FASTQ format we used Sambamba ^26^ view (version 0.5.9) and Samtools ^27^ Fastq (version 1.8) commands. Third, we depleted the reads originating from viral and bacterial sources by running kraken2^28^ and retained the unclassified reads. We assembled the retained reads using Trinity RNA Seq assembler (version 2.8.4)^29^ with custom parameters “--no_normalize_reads”, “--max_chrysalis_cluster_size 100”, and “--max_reads_per_graph 5000000”. We then mapped the assembled sequences germline V, D, and J gene regions using IgBLAST^28,30^ (version 1.13). Sequences with high V gene match scores (cutoff = 100) were kept as putative Ig chains.

### Ig reconstruction performance evaluation

To evaluate the performance of this workflow and evaluate the possibility of identifying which heavy and light chain form a pair in cases where there is a strongly expressed clone we generated a synthetic benchmark using single cell sequencing data for sample PW2 from Lindman et.al.^11^. We estimated the distribution of Ig reads in TCGA data and found it to closely follow a lognormal model (Supplementary Figure 1). We then generated a set of 1000 synthetic samples. We generated each sample by sampling 25 numbers from this distribution and assigning them as the total number of Ig reads to 25 random B cells from the PW2 data. We then randomly selected the corresponding number of RNA sequencing reads from those mapped to Ig chains in each of the 25 B cells. To this mix we added 10 million reads randomly selected from a bulk RNA sequencing sample (TCGA-04-1348-01A), to act as background. In evaluation, we considered the reconstruction to be a success if all of the CDR regions (CDR1, CDR2, and CDR3) for both the heavy and light chain of the top clone were correct and the pair was correctly selected as the most abundant Ig.

### Ig quantification, pairing, and somatic hypermutation analysis

For each processed TCGA sample we used Kallisto to quantify the amount of RNA sequencing reads originating from each of the putative Ig chain transcripts. We separately calculated the *Berger-Parker dominance index*^10^ for both the heavy and light Ig chains, as the proportion of reads attributed to the most common chain of the corresponding type. We then calculated the dominance score for each sample as the geometric mean of the heavy and light chain Berger-Parker indices. Using the results of the synthetic benchmark we determined the threshold on this score (>0.382) above which we expect the sequence reconstruction to be accurate, and the top (most abundant) heavy and light chain to form a paired Ig. For each such Ig we assigned the putative isotypes by mapping the constant portions of the heavy chains (sections of the assembled sequences post the annotated J regions) to the Gencode v22 reference immunoglobulin C sequences. We used the IgBlast mappings to the germline segment to identify the mutated positions in each chain. For every position as defined by the Martin numbering scheme^31^ we calculated the mutation frequency across all of the reconstructed immunoglobulin chains.

### Processing of TCGA RNA sequencing data

TCGA BAM files were retrieved and processed in Cancer Genomics Cloud^32^ (CGC). We applied sambamba-sort to sort the BAM files and Samtools fastq to convert them to FASTQ files, followed by Salmon quant in mapping-based mode (GENOME v27 (GRCh38.p10) transcript assembly was used) with GC bias correction. We obtained TPM values for 10,533 tumor samples and 730 normal samples across 33 TCGA cancer types (Supplementary Table S3).

### Statistical analysis of antigen arrays

Raw signal intensities (mean F635 values) were correlated for background signal (mean B635 values) using the function background from R package limma^33^, with the correction method set to ‘normexpr’ with ‘mle’ parameter estimation strategy. The replicate spots for the same target clone were summarized by calculating the geometric mean followed by logarithmic transformation. We noticed that a number of spots reported high signal intensities across multiple arrays likely due to non-specific binding. To counter this issue we subtracted the mean signal value across all of the 172 arrays for each target clone. We then centered and scaled the signal intensities and performed multiple testing corrections by calculating the false discovery rate according to the Benjamini-Hochberg method^33,34^. We used a stringent q-value cutoff of 0.01 in order to control the number of false positive hits.

### Antibody synthesis

The variable region of the antibodies were recombined with a constant region of human IgG class I using AbAb’s recombinant platform (Absolute Antibody Ltd, Oxford, UK). The antibodies were expressed into HEK293 cells using the Absolute Antibody transient expression system and purified by one-step affinity chromatography. We performed Quality Control (QC) analysis to assess concentration, total amount, level of endotoxin, and sodium dodecyl sulfate polyacrylamide gel electrophoresis (SDS-PAGE) (Supplementary Table S1).

### HuProt™ Protein arrays

We tested each candidate antibody using the human proteome microarray HuProt^™^ 4.0, a comprehensive library of GST-tagged recombinant human protein expressed in yeast. Briefly, the HuProt arrays were blocked with 5% BSA/1×TBS-T at room temperature for 1 h. Then antibodies were incubated at 4 °C overnight, washed and labelled with secondary antibodies prior detection.

### Membrane Proteome Array (MPA) specificity testing

Membrane Proteome Array (MPA) is Integral Molecular’s cell-based array of ~6,000 human membrane proteins, each expressed in live unfixed cells in separate wells of a 384-well plate^13^. In this study, the MPA was expressed in HEK-293T cells 36h prior to testing. Each MAb was fluorescently labeled and added to the MPA at a concentration optimized for the best signal-to-background ratio for target detection using an independent immunofluorescence titration curve against membrane-tethered protein A. Binding was measured by Intellicyt iQue3. Each 384-well plate contained positive (Fc-binding) and negative (empty vector) controls to ensure plate-by-plate data validity. Hits were validated by flow cytometry with serial dilutions of antibody, and the target identity was confirmed by sequencing.

### Surface Plasmon Resonance (SPR) assay

**A** Biacore 8K instrument, exploiting the SPR principle, was used to calculate the equilibrium dissociation constant (*K*_D_). Selected antibodies (ligand) were individually immobilized to a sensor chip coated with Protein A to ensure the correct orientation of the antibody while the antigens (analyte) were injected within the flow stream at 30 μ1/min. We used 90s association time for each experiment with 600s dissociation time. Data analysis was performed using the R package *pbm^35^*. Sensorgram raw data time series measurements were downloaded from the Biacore 8K instrument, and fitted to an appropriate observation model from the pbm package using non-linear least squares parameter estimation techniques. Models were selected parsimoniously; those which could adequately explain the data using fewer parameters were generally preferred. In a small number cases, selected concentration curves were excluded from the fitting procedure due to either instrumental measurement anomalies, or apparent statistical outliers. Examples of qualifying anomalies included refractive index bulk shift discontinuities^36^ at the transition between the association and dissociation measurement phases.

### Epitope mapping

Hydrogen deuterium exchange mass spectrometry (HDX-MS) was used to determine the epitope recognized by the antibodies. Linear peptides spanning the target antigen length were incubated in a deuterium containing buffer in presence or absence of the relevant antibody to observe differential exchange with hydrogen at the putative binding sites.

## Data availability

The raw RNA sequencing reads used in this study are generated by the TCGA Research Network: https://www.cancer.gov/tcga, and available through the TCGA data portal. All relevant processed data are included as Source Data linked to this paper.

## Code availability

The code for generation of the synthetic data used in the benchmarks, statistical analyses of the arrays, and plots is from the authors available upon a reasonable request.

## Authors contributions

GR, IDS, and BCT developed the sequence reconstruction workflow and generated the Ig chain sequences. GR performed the performance evaluation of the workflow and analysed the protein array data. ALS prepared supplementary figures depicting sensorgram plots for SPR assay experiments. Milena P. performed statistical analysis comparing *K*_D_ values of human vs. commercially available rabbit-derived antibodies. IG prepared the main figures. Milos P. and IG processed the epitope mapping and generated the corresponding visualization. DB and DK designed research and DB oversaw the implementation. GR, DB, DAL and DK prepared the manuscript. All authors reviewed the final manuscript.

## Supplementary figures

**Figure S1 | Distribution of Ig reads mapping to different chains in TCGA data follows a lognormal model.**

The plot represents the distribution of the read counts mapped to each reconstructed Ig chain in the TCGA dataset. The data closely match a lognormal distribution with the estimated parameters mean=5.78, standard deviation=1.52.

**Figure S2 | Distribution of Berger-Parker dominance score across different cancer types in the TCGA data**

The plot represents the distribution of Berger-Parker dominance scores of the heavy and the light chains reconstructed from tumour samples in the TCGA dataset.

**Figure S3 | Surface plasmon resonance assay sensorgram plots providing antibody-antigen equilibrium dissociation constants**

Sensorgrams plots depict the SPR response of the Biacore 8K instrument in relative units (RU) vs. time, as commercially sourced antigens are flowed across surface-captured antibodies during an initial 90 s measurement interval, and then permitted to dissociate without antigen flowing, during a subsequent 600 s measurement interval. Antigens are prepared across a range of two-fold serially diluted concentrations, and within a given set of concentration curves comprising a single measurement attempt, data from all concentrations are fitted simultaneously using non-linear least squares regression to arrive at a single unifying *K*_D_ estimate (or a pair of *K*_D_ estimates, when a biphasic binding model provides a better fit) valid for that measurement attempt. Solid colored lines indicate raw data observed by the apparatus, and overlaid solid black lines indicate the fit result estimated using the model; a closer match between the two indicates a better fit, with correspondingly superior explanatory power. In some cases, indicated by the absence of an overlaid solid black regression curve, data corresponding to one or more concentration curves may be discarded from the input to the regression model, if the result leads to a significantly better fit across the remaining concentration curves. Parameter estimates provided by the fit model are listed in the table at the top of each page, while lettered panels enumerate cases in which multiple replicate measurements were attempted for a given antibody-antigen pairing.

**Figure S4 | Binding scores for top 5 hits from protein array screening.**

Plots depict the scaled binding scores from top 5 hits from each of the protein arrays. Binding scores were calculated by fitting random effects models to the signal intestines, weighting each target by its mean rank across the arrays (to eliminate common promiscuous binders) and scaled by taking the z scores (Methods section). Asterisks mark hits with Benjamini and Hochberg false discovery rate < 0.01.

**Figure S5 | Membrane proteome-wide specificity testing of recombinant antibodies.** Each antibody was screened for binding against Integral Molecular’s Membrane Proteome Array, consisting of ~6,000 human membrane proteins in their native state in unfixed cells. Antibody binding was detected by flow cytometry, and hits were defined as resulting in binding signals more than 3 standard deviations higher than background. Validated targets are shown in blue.

## Supplementary tables

**Table S1 |** Antibody expression and antigen screening summary.

**Table S2 |** Biacore results.

**Table S3 |** Gene expression table TCGA (Salmon, TPM, no other normalization).

**Table S4 |** Protein array results.

**Table S5 |** Membrane proteome-wide screening results

