## Supplementary Figures for "The landscape of high-affinity human antibodies against intratumoral antigens": Supplementary Figure 1.pdf

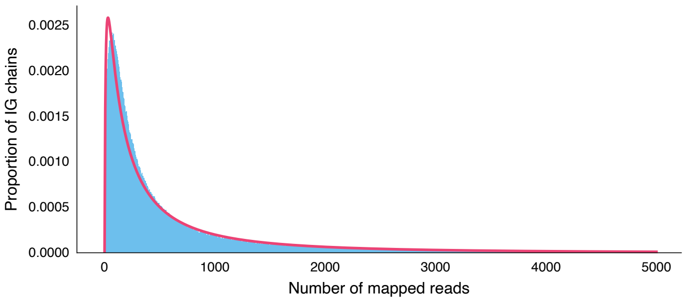


**Supplementary Figure 1 | Distribution of IG reads mapping to different chains in TCGA data follows a lognormal model.**
