## Supplementary Figures for "The landscape of high-affinity human antibodies against intratumoral antigens": Supplementary Figure 2.pdf

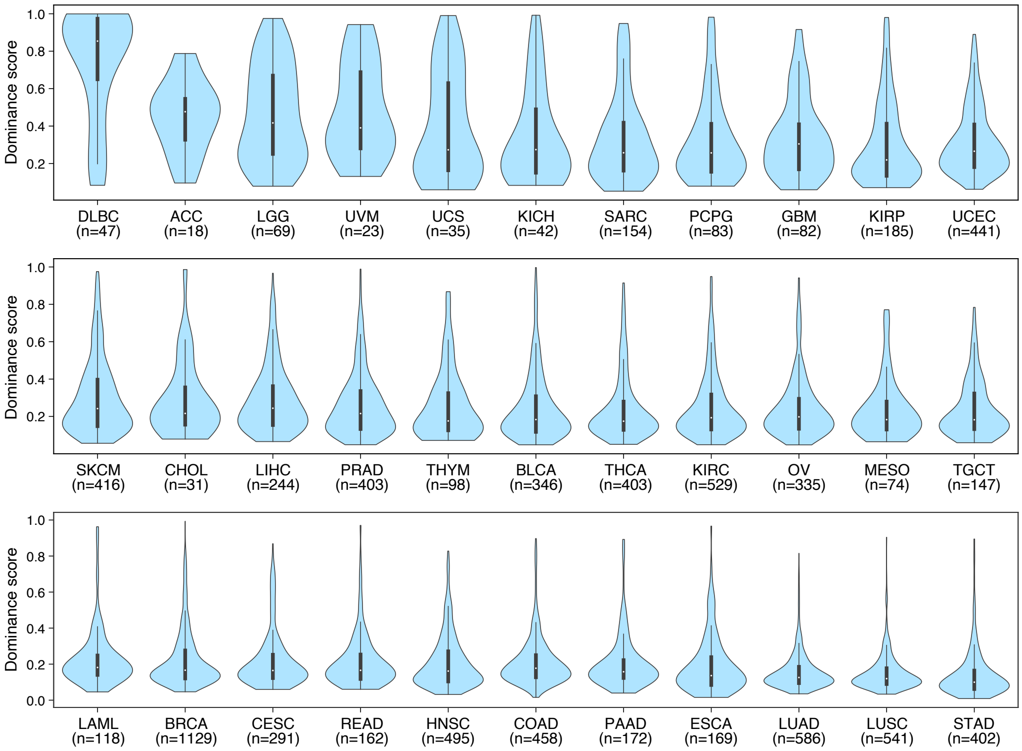


**Supplementary Figure 2 | Distribution of Berger-Parker index across different cancer types in the TCGA data.**
