## Supplementary Figures for "The landscape of high-affinity human antibodies against intratumoral antigens": Supplementary Figure 3.pdf

### ACTN4 (TP302873) Surface Plasmon Resonance Sensorgrams

| Panel | Antibody Name | Antibody Type | KD1 (nM) | kon1 (1/nM·s) | koff1 (1/s) | rmax1 (RU) | KD2 (nM) | kon2 (1/nM·s) | koff2 (1/s) | rmax2 (RU) | drift (RU/s) | offset (RU) |
|---|---|---|---|---|---|---|---|---|---|---|---|---|
| a | CYC088 | Human | 1.718 | 4.26e−04 | 7.31e−04 | 12.1 | 79.665 | 8.17e−04 | 6.51e−02 | 14.7 | | 0.9 |

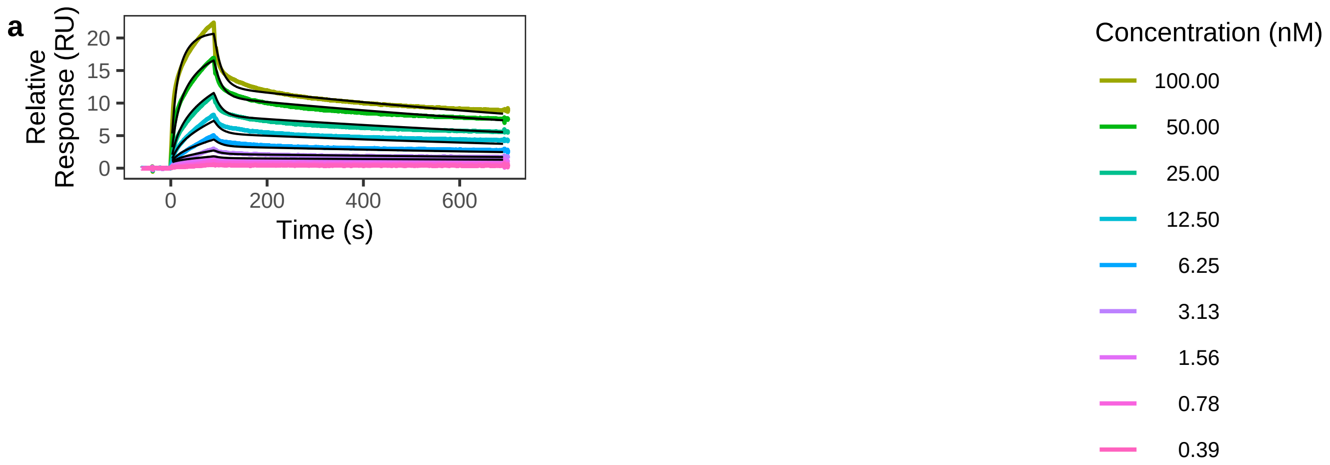

### ANXA1 (APrEST71575) Surface Plasmon Resonance Sensorgrams

| Panel | Antibody Name | Antibody Type | KD1 (nM) | kon1 (1/nM·s) | koff1 (1/s) | rmax1 (RU) | KD2 (nM) | kon2 (1/nM·s) | koff2 (1/s) | rmax2 (RU) | drift (RU/s) | offset (RU) |
|---|---|---|---|---|---|---|---|---|---|---|---|---|
| a | CYC188 | Human | 3.740 | 2.01e−04 | 7.50e−04 | 22.8 | 52.977 | 3.17e−04 | 1.68e−02 | 9.5 | | |
| b | CYC188 | Human | 3.441 | 1.60e−04 | 5.52e−04 | 13.8 | 148.687 | 2.11e−04 | 3.14e−02 | 10.3 | | |
| c | CYC188 | Human | 0.853 | 2.35e−04 | 2.00e−04 | 17.9 | 6.084 | 9.75e−04 | 5.93e−03 | 8.5 | −4.06e−03 | |
| d | CYC188 | Human | 0.771 | 4.44e−04 | 3.42e−04 | 68.8 | 498.887 | 2.73e−05 | 1.36e−02 | 166.9 | | |
| e | HPA011271 | Rabbit | 1.657 | 3.15e−04 | 5.22e−04 | 47.7 | 44.440 | 3.51e−04 | 1.56e−02 | 15.5 | | 1.6 |
| f | HPA011271 | Rabbit | 1.407 | 2.41e−04 | 3.39e−04 | 30.8 | 46.684 | 7.12e−04 | 3.32e−02 | 14.4 | | |

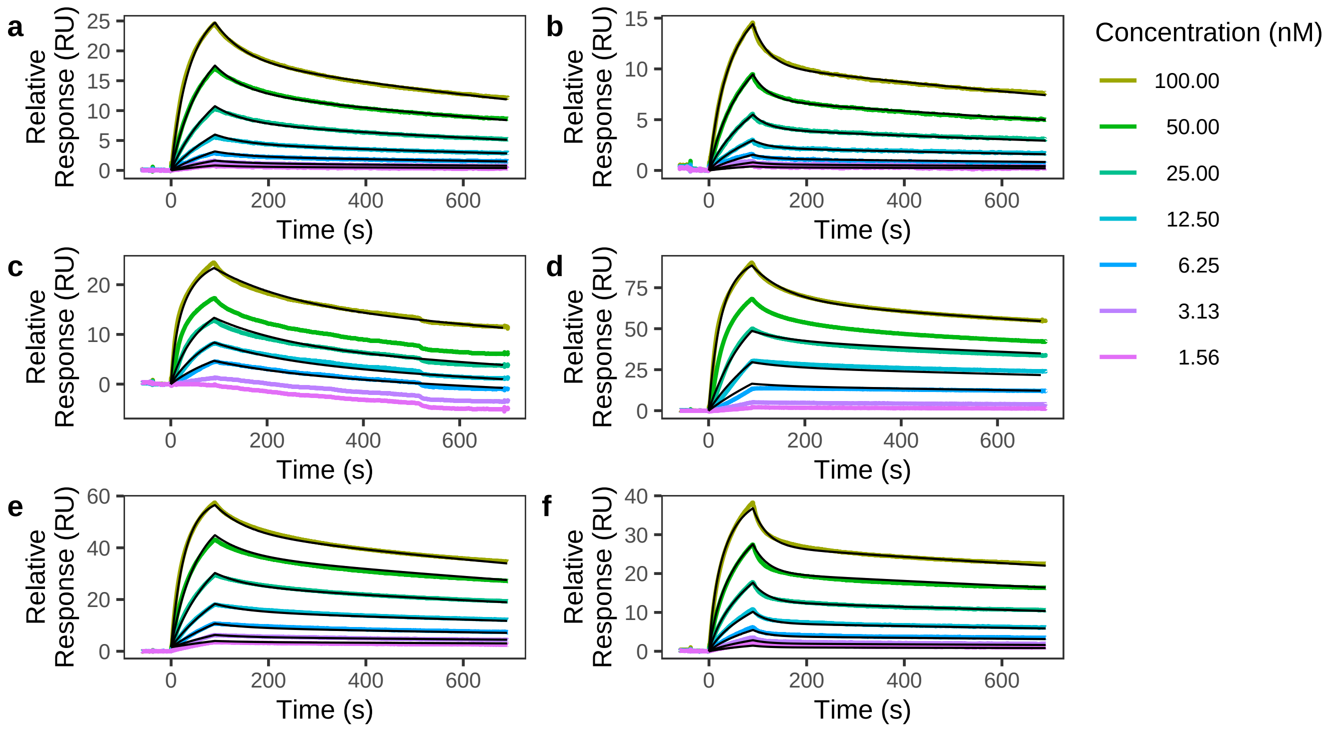

### ANXA1 (APrEST71576) Surface Plasmon Resonance Sensorgrams

| Panel | Antibody Name | Antibody Type | KD1 (nM) | kon1 (1/nM·s) | koff1 (1/s) | rmax1 (RU) | KD2 (nM) | kon2 (1/nM·s) | koff2 (1/s) | rmax2 (RU) | drift (RU/s) | offset (RU) |
|---|---|---|---|---|---|---|---|---|---|---|---|---|
| a | CYC188 | Human | 1154.396 | 7.19e−07 | 8.30e−04 | 427.2 | 19461.426 | 9.45e−07 | 1.84e−02 | 283.4 | | |
| b | HPA011271 | Rabbit | 7.049 | 1.52e−04 | 1.07e−03 | 3.3 | 26380.678 | 1.15e−06 | 3.05e−02 | 394.3 | | |
| c | HPA011272 | Rabbit | 1.344 | 2.47e−04 | 3.32e−04 | 29.3 | 2939.835 | 2.77e−06 | 8.15e−03 | 366.9 | | |
| d | HPA011272 | Rabbit | 1.408 | 2.95e−04 | 4.15e−04 | 21.7 | 111.843 | 2.63e−04 | 2.94e−02 | 11.3 | | |

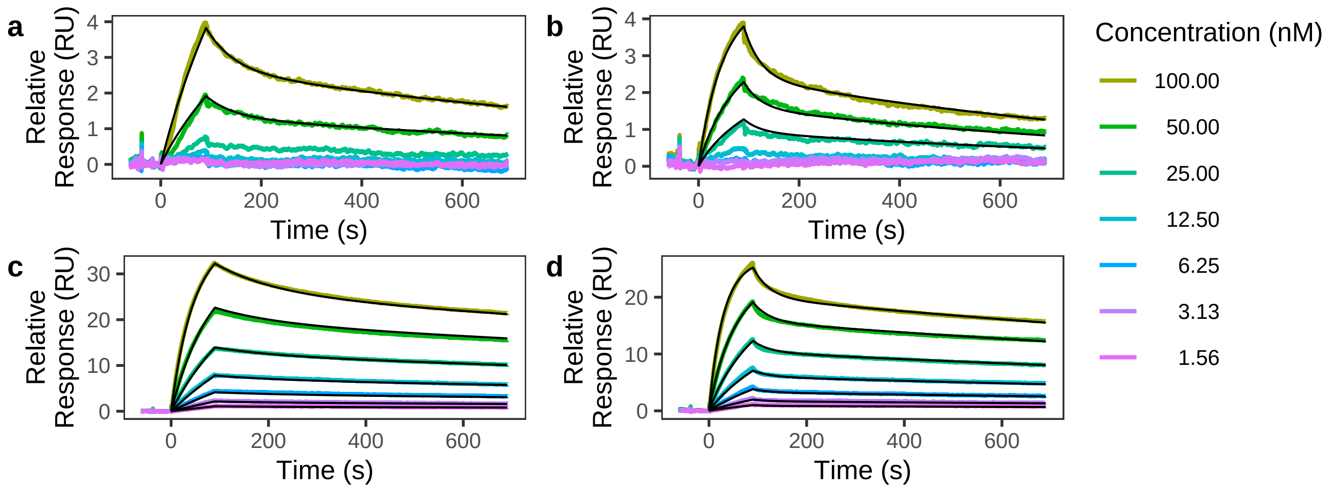

BIRC7 (ab87179) Surface Plasmon Resonance Sensorgrams

| Panel | Antibody Name | Antibody Type | KD1 (nM) | kon1 (1/nM·s) | koff1 (1/s) | rmax1 (RU) | KD2 (nM) | kon2 (1/nM·s) | koff2 (1/s) | rmax2 (RU) | drift (RU/s) | offset (RU) |
|---|---|---|---|---|---|---|---|---|---|---|---|---|
| a | CYC278 | Human | 5.276 | 2.45e−05 | 1.29e−04 | 210.4 | | | | | 1.21e−03 | |

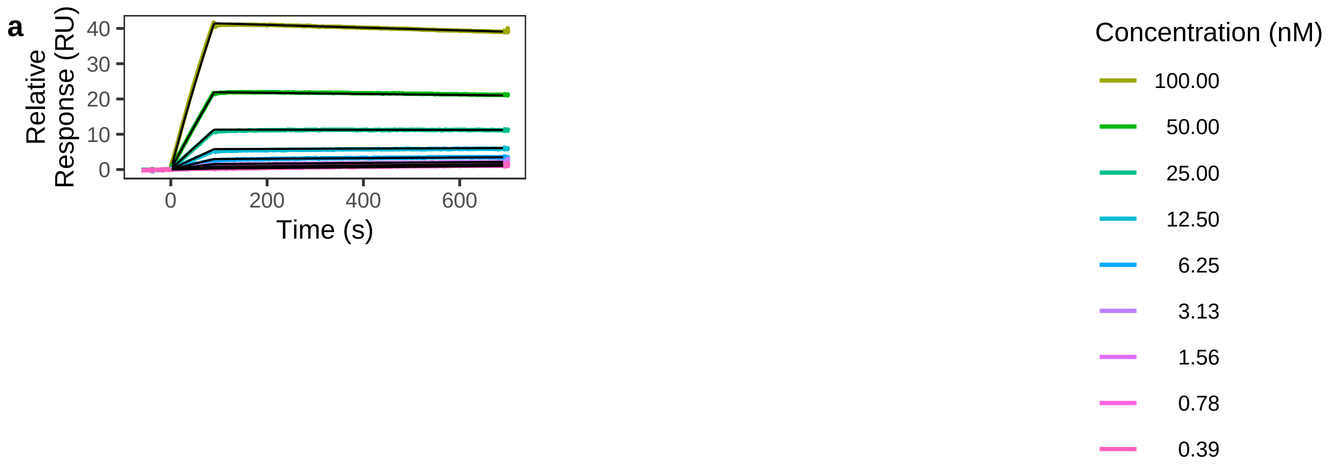

### C4BPB (ab130028) Surface Plasmon Resonance Sensorgrams

| Panel | Antibody Name | Antibody Type | KD1 (nM) | kon1 (1/nM·s) | koff1 (1/s) | rmax1 (RU) | KD2 (nM) | kon2 (1/nM·s) | koff2 (1/s) | rmax2 (RU) | drift (RU/s) | offset (RU) |
|---|---|---|---|---|---|---|---|---|---|---|---|---|
| a | CYC214 | Human | 0.356 | 3.89e−04 | 1.38e−04 | 71.9 | | | | | | |
| b | CYC214 | Human | 0.686 | 1.91e−04 | 1.31e−04 | 114.0 | | | | | | |
| c | CYC214 | Human | 1.093 | 2.56e−04 | 2.80e−04 | 96.8 | | | | | 5.60e−03 | |
| d | CYC214 | Human | 0.418 | 2.46e−04 | 1.03e−04 | 209.3 | | | | | | |

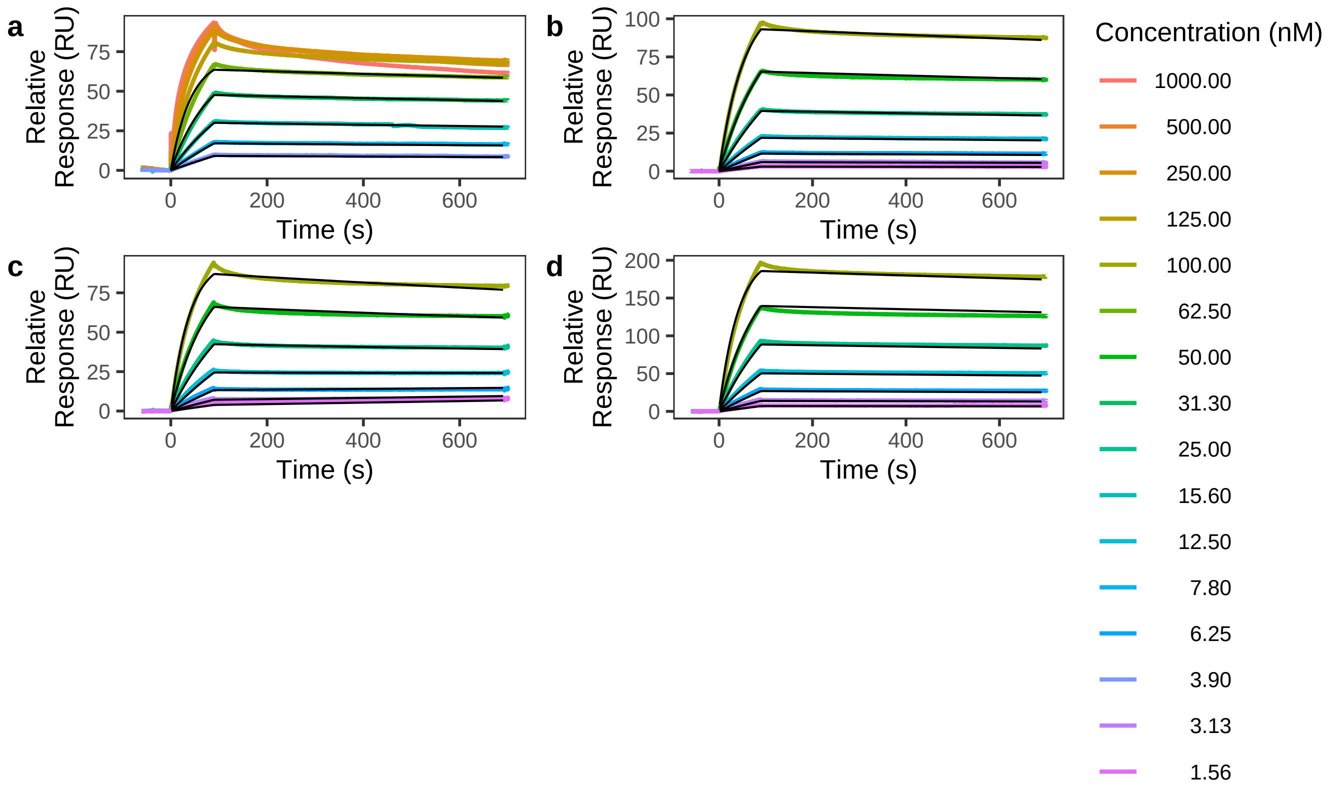

CTAG1A (TP316285) Surface Plasmon Resonance Sensorgrams

| Panel | Antibody Name | Antibody Type | KD1 (nM) | kon1 (1/nM·s) | koff1 (1/s) | rmax1 (RU) | KD2 (nM) | kon2 (1/nM·s) | koff2 (1/s) | rmax2 (RU) | drift (RU/s) | offset (RU) |
|---|---|---|---|---|---|---|---|---|---|---|---|---|
| a | CYC002 | Human | 0.236 | 4.52e−04 | 1.07e−04 | 26.1 | 7.960 | 2.20e−05 | 1.75e−04 | 79.9 | | |
| b | CYC002 | Human | 0.305 | 3.99e−04 | 1.22e−04 | 25.6 | 7.499 | 1.73e−05 | 1.30e−04 | 81.5 | | |
| c | CYC002 | Human | 0.222 | 4.72e−04 | 1.05e−04 | 25.5 | 3.888 | 4.38e−05 | 1.70e−04 | 56.3 | | |

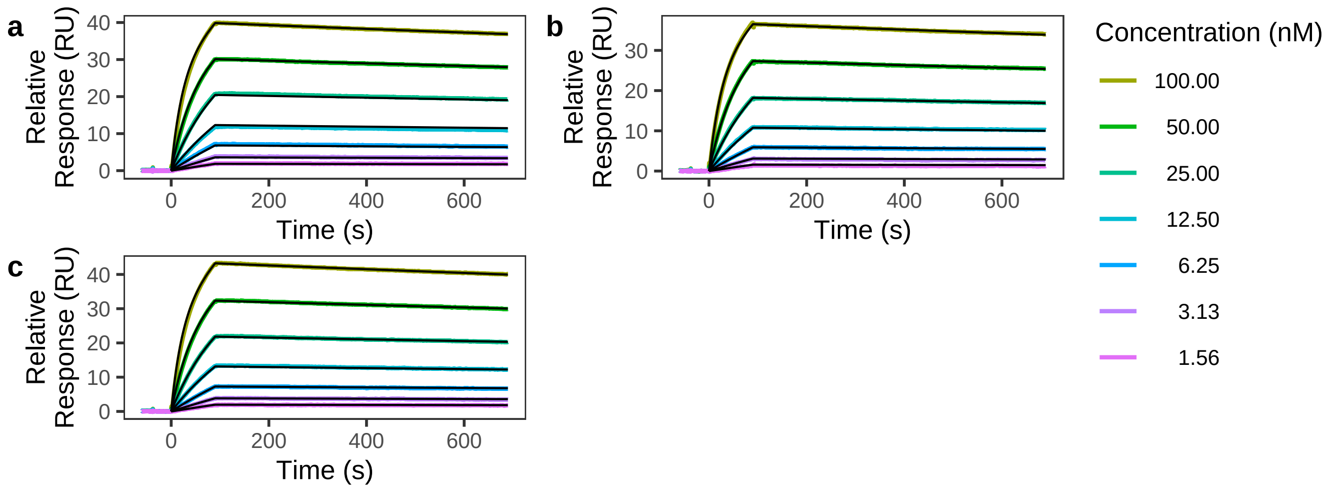

### CTAG1A (TP316285) Surface Plasmon Resonance Sensorgrams

| Panel | Antibody Name | Antibody Type | KD1 (nM) | kon1 (1/nM·s) | koff1 (1/s) | rmax1 (RU) | KD2 (nM) | kon2 (1/nM·s) | koff2 (1/s) | rmax2 (RU) | drift (RU/s) | offset (RU) |
|---|---|---|---|---|---|---|---|---|---|---|---|---|
| a | CYC156 | Human | 0.461 | 8.06e−05 | 3.71e−05 | 45.7 | | | | | 8.69e−04 | |

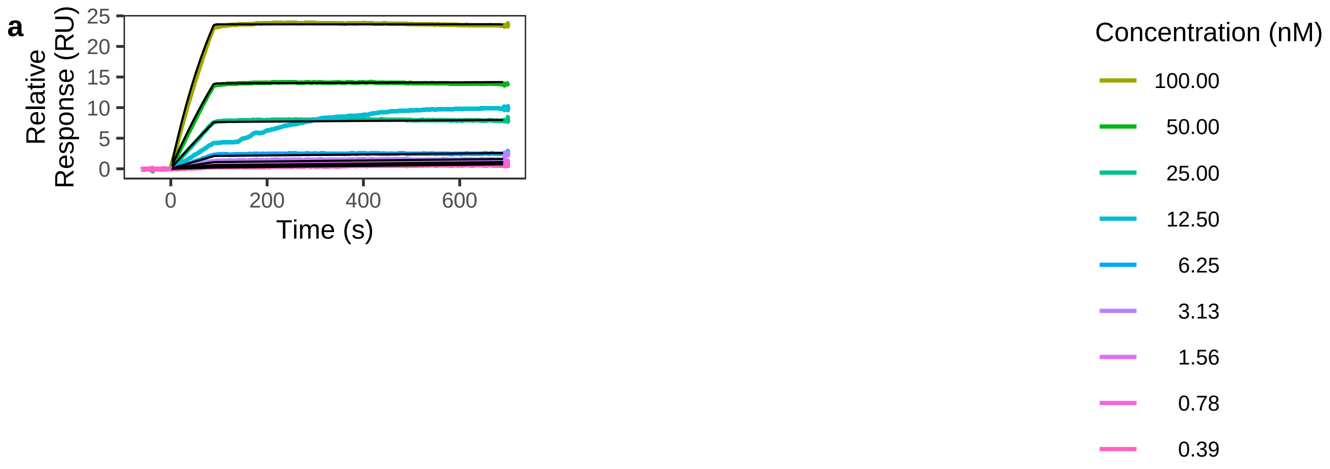

### CTAG1A (TP316285) Surface Plasmon Resonance Sensorgrams

| Panel | Antibody Name | Antibody Type | KD1 (nM) | kon1 (1/nM·s) | koff1 (1/s) | rmax1 (RU) | KD2 (nM) | kon2 (1/nM·s) | koff2 (1/s) | rmax2 (RU) | drift (RU/s) | offset (RU) |
|---|---|---|---|---|---|---|---|---|---|---|---|---|
| a | CYC187 | Human | 0.491 | 2.43e−04 | 1.20e−04 | 82.7 | | | | | | |

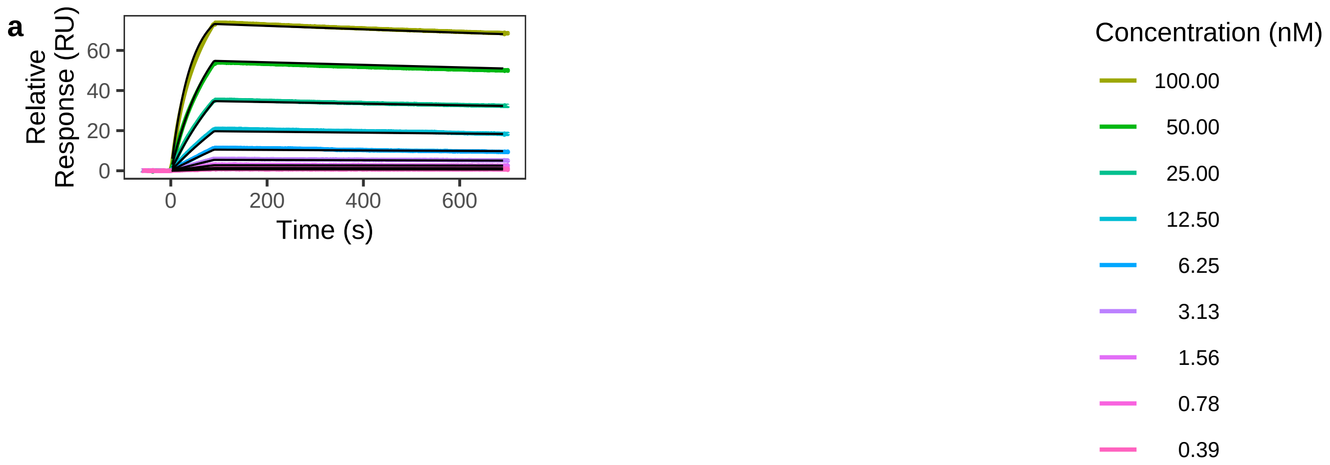

DLL3 (9749−DL−050) Surface Plasmon Resonance Sensorgrams

| Panel | Antibody Name | Antibody Type | KD1 (nM) | kon1 (1/nM·s) | koff1 (1/s) | rmax1 (RU) | KD2 (nM) | kon2 (1/nM·s) | koff2 (1/s) | rmax2 (RU) | drift (RU/s) | offset (RU) |
|---|---|---|---|---|---|---|---|---|---|---|---|---|
| a | CYC077 | Human | 11.202 | 1.41e−04 | 1.58e−03 | 11.2 | 37.005 | 1.03e−03 | 3.82e−02 | 10.9 | | |

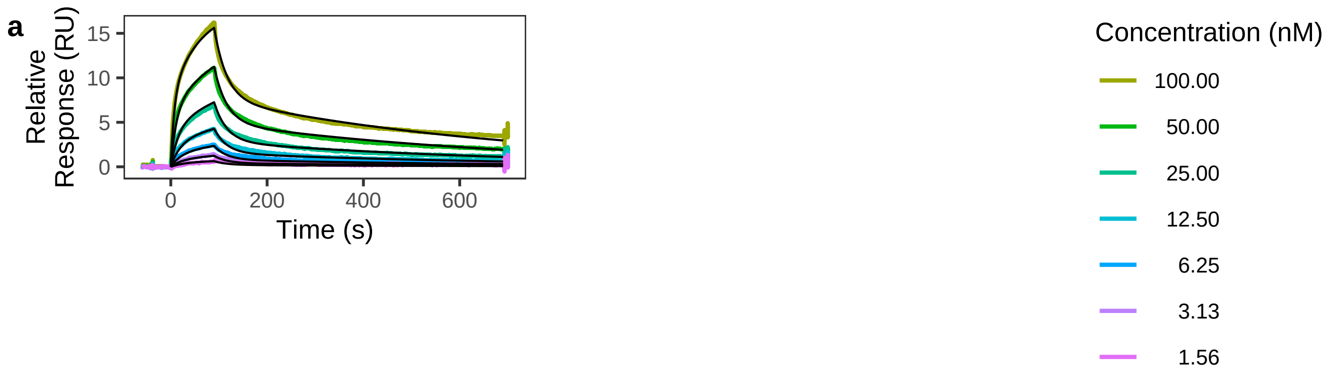

GAGE1 (TP761867) Surface Plasmon Resonance Sensorgrams

| Panel | Antibody Name | Antibody Type | KD1 (nM) | kon1 (1/nM·s) | koff1 (1/s) | rmax1 (RU) | KD2 (nM) | kon2 (1/nM·s) | koff2 (1/s) | rmax2 (RU) | drift (RU/s) | offset (RU) |
|---|---|---|---|---|---|---|---|---|---|---|---|---|
| a | CYC014 | Human | 2.073 | 2.61e−04 | 5.40e−04 | 3.5 | | | | | | |

a

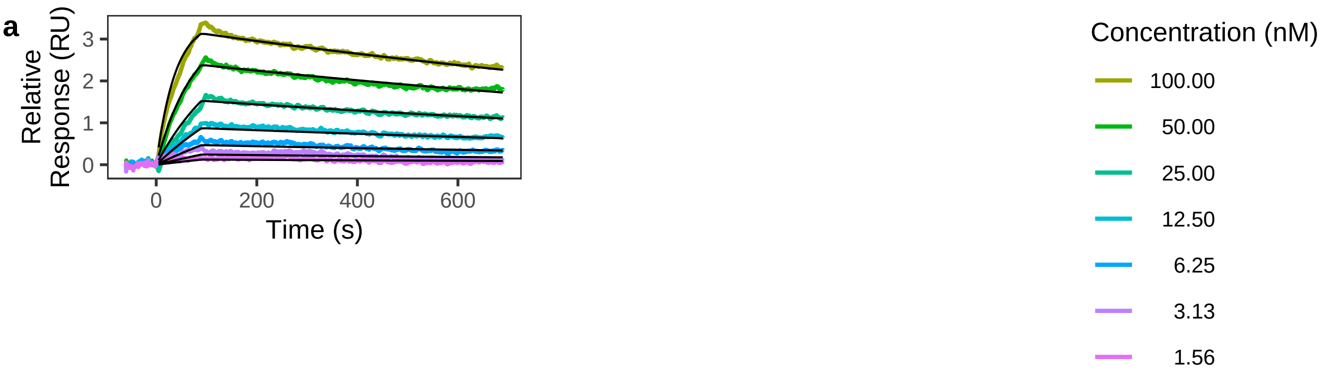

GAGE1 (TP761867) Surface Plasmon Resonance Sensorgrams

| Panel | Antibody Name | Antibody Type | KD1 (nM) | kon1 (1/nM·s) | koff1 (1/s) | rmax1 (RU) | KD2 (nM) | kon2 (1/nM·s) | koff2 (1/s) | rmax2 (RU) | drift (RU/s) | offset (RU) |
|---|---|---|---|---|---|---|---|---|---|---|---|---|
| a | CYC025 | Human | 1.265 | 1.98e−04 | 2.50e−04 | 2.1 | | | | | | |

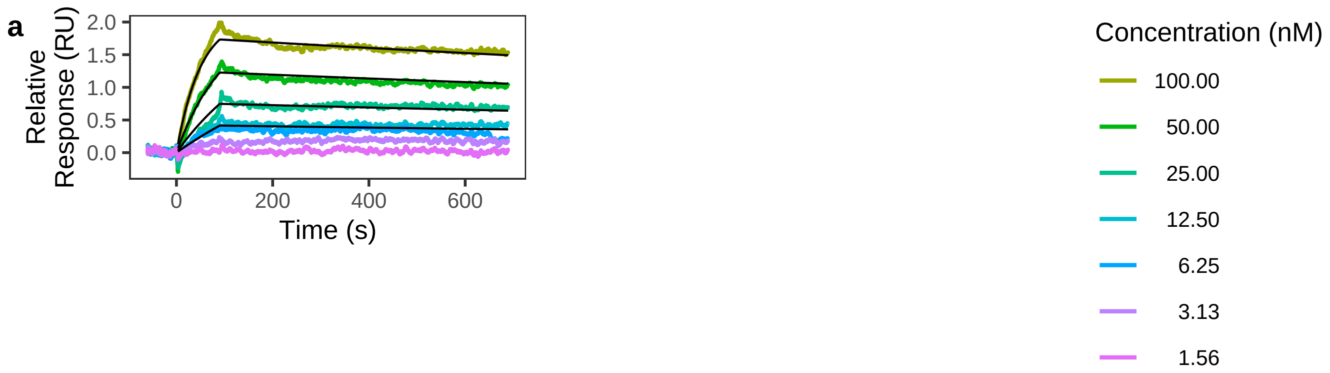

### GAGE2A (AR51161PU−S) Surface Plasmon Resonance Sensorgrams

| Panel | Antibody Name | Antibody Type | KD1 (nM) | kon1 (1/nM·s) | koff1 (1/s) | rmax1 (RU) | KD2 (nM) | kon2 (1/nM·s) | koff2 (1/s) | rmax2 (RU) | drift (RU/s) | offset (RU) |
|---|---|---|---|---|---|---|---|---|---|---|---|---|
| a | CYC014 | Human | 0.616 | 7.58e−04 | 4.67e−04 | 10.6 | 150.920 | 8.97e−05 | 1.35e−02 | 20.9 | | −1.0 |
| b | CYC014 | Human | 0.340 | 8.92e−04 | 3.03e−04 | 5.1 | 2.515 | 9.09e−03 | 2.29e−02 | 1.2 | | 0.3 |
| c | CYC014 | Human | 0.874 | 6.31e−04 | 5.51e−04 | 6.6 | 2.207 | 7.02e−03 | 1.55e−02 | 1.6 | | |
| d | CYC014 | Human | 0.476 | 5.47e−04 | 2.60e−04 | 12.2 | 2.124 | 7.89e−03 | 1.68e−02 | 2.3 | 2.24e−03 | |
| e | HPA043232 | Rabbit | 0.060 | 6.48e−03 | 3.86e−04 | 11.7 | 1.444 | 5.44e−04 | 7.86e−04 | 29.1 | | |
| f | HPA043232 | Rabbit | 0.061 | 4.16e−03 | 2.56e−04 | 14.7 | 1.270 | 4.62e−04 | 5.87e−04 | 26.7 | | |
| g | HPA043232 | Rabbit | 0.122 | 1.33e−03 | 1.62e−04 | 31.5 | 936.404 | 6.89e−06 | 6.45e−03 | 772.5 | | |
| h | HPA043232 | Rabbit | 0.164 | 1.33e−03 | 2.17e−04 | 33.2 | 1232.109 | 6.16e−06 | 7.59e−03 | 986.4 | | |
| i | HPA043232 | Rabbit | 0.235 | 1.22e−03 | 2.88e−04 | 36.4 | 1265.428 | 5.17e−06 | 6.54e−03 | 1045.5 | | 1.9 |
| j | HPA043232 | Rabbit | 0.127 | 1.50e−03 | 1.90e−04 | 22.7 | 347.642 | 1.09e−05 | 3.79e−03 | 345.9 | | 1.5 |

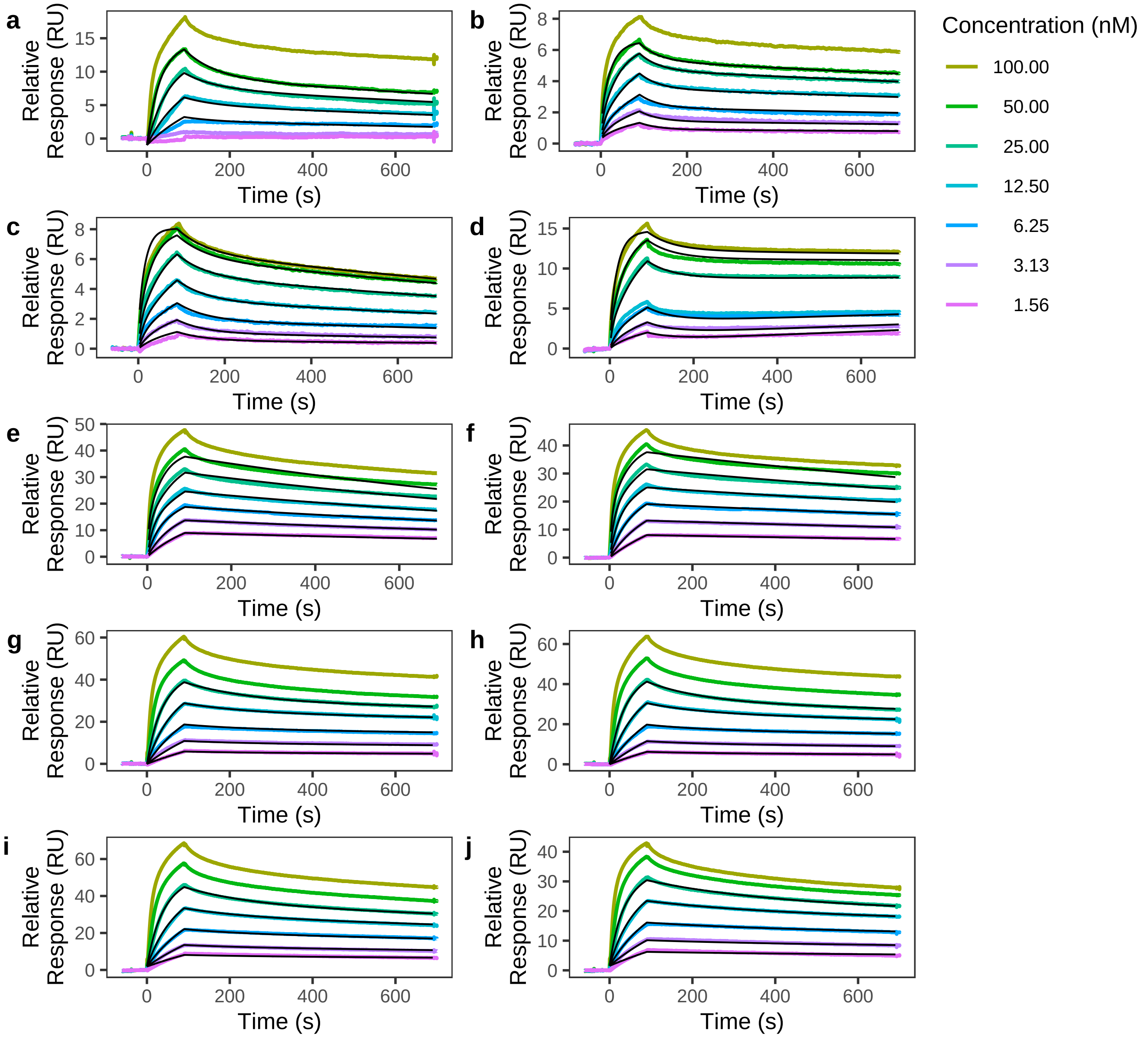

### GAGE2A (AR51161PU−S) Surface Plasmon Resonance Sensorgrams

| Panel | Antibody Name | Antibody Type | KD1 (nM) | kon1 (1/nM·s) | koff1 (1/s) | rmax1 (RU) | KD2 (nM) | kon2 (1/nM·s) | koff2 (1/s) | rmax2 (RU) | drift (RU/s) | offset (RU) |
|---|---|---|---|---|---|---|---|---|---|---|---|---|
| a | CYC025 | Human | 0.470 | 6.96e−04 | 3.27e−04 | 2.4 | 4.761 | 5.24e−03 | 2.50e−02 | 0.7 | | |
| b | CYC025 | Human | 1.076 | 5.42e−04 | 5.84e−04 | 26.2 | 26.721 | 7.93e−04 | 2.12e−02 | 10.6 | | −1.3 |
| c | CYC025 | Human | 0.991 | 4.94e−04 | 4.90e−04 | 30.1 | 13.067 | 1.10e−03 | 1.44e−02 | 10.3 | | −0.9 |
| d | HPA043232 | Rabbit | 0.060 | 6.48e−03 | 3.86e−04 | 11.7 | 1.444 | 5.44e−04 | 7.86e−04 | 29.1 | | |
| e | HPA043232 | Rabbit | 0.061 | 4.16e−03 | 2.56e−04 | 14.7 | 1.270 | 4.62e−04 | 5.87e−04 | 26.7 | | |
| f | HPA043232 | Rabbit | 0.122 | 1.33e−03 | 1.62e−04 | 31.5 | 936.404 | 6.89e−06 | 6.45e−03 | 772.5 | | |
| g | HPA043232 | Rabbit | 0.164 | 1.33e−03 | 2.17e−04 | 33.2 | 1232.109 | 6.16e−06 | 7.59e−03 | 986.4 | | |
| h | HPA043232 | Rabbit | 0.235 | 1.22e−03 | 2.88e−04 | 36.4 | 1265.428 | 5.17e−06 | 6.54e−03 | 1045.5 | | 1.9 |
| i | HPA043232 | Rabbit | 0.127 | 1.50e−03 | 1.90e−04 | 22.7 | 347.642 | 1.09e−05 | 3.79e−03 | 345.9 | | 1.5 |

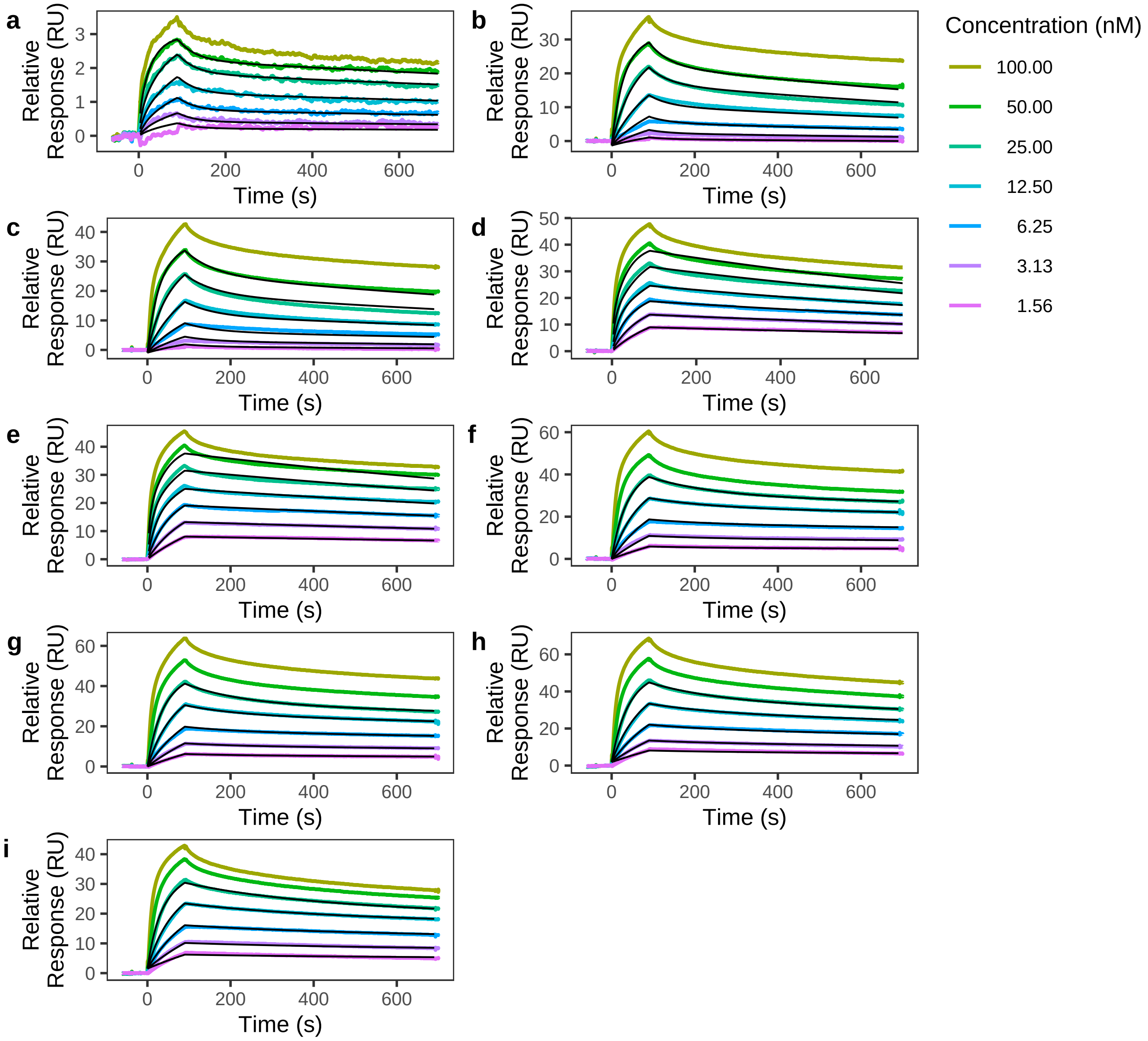

### GPR83 (H00010888−G01) Surface Plasmon Resonance Sensorgrams

| Panel | Antibody Name | Antibody Type | KD1 (nM) | kon1 (1/nM·s) | koff1 (1/s) | rmax1 (RU) | KD2 (nM) | kon2 (1/nM·s) | koff2 (1/s) | rmax2 (RU) | drift (RU/s) | offset (RU) |
|---|---|---|---|---|---|---|---|---|---|---|---|---|
| a | CYC181 | Human | | | | | | | | | | |
| b | CYC181 | Human | | | | | | | | | | |
| c | ab140767 | Rabbit | | | | | | | | | | |
| d | ab228351 | Rabbit | | | | | | | | | | |

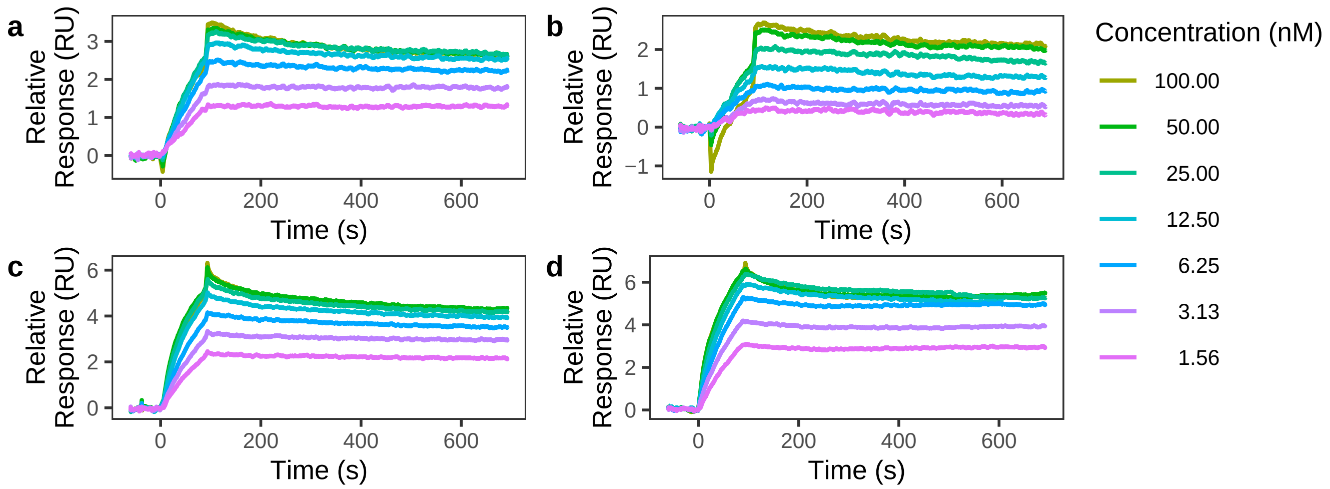

### HCLS1 (TP300329) Surface Plasmon Resonance Sensorgrams

| Panel | Antibody Name | Antibody Type | KD1 (nM) | kon1 (1/nM·s) | koff1 (1/s) | rmax1 (RU) | KD2 (nM) | kon2 (1/nM·s) | koff2 (1/s) | rmax2 (RU) | drift (RU/s) | offset (RU) |
|---|---|---|---|---|---|---|---|---|---|---|---|---|
| a | CYC101 | Human | 3.156 | 6.65e−05 | 2.10e−04 | 36.8 | | | | | | 0.7 |

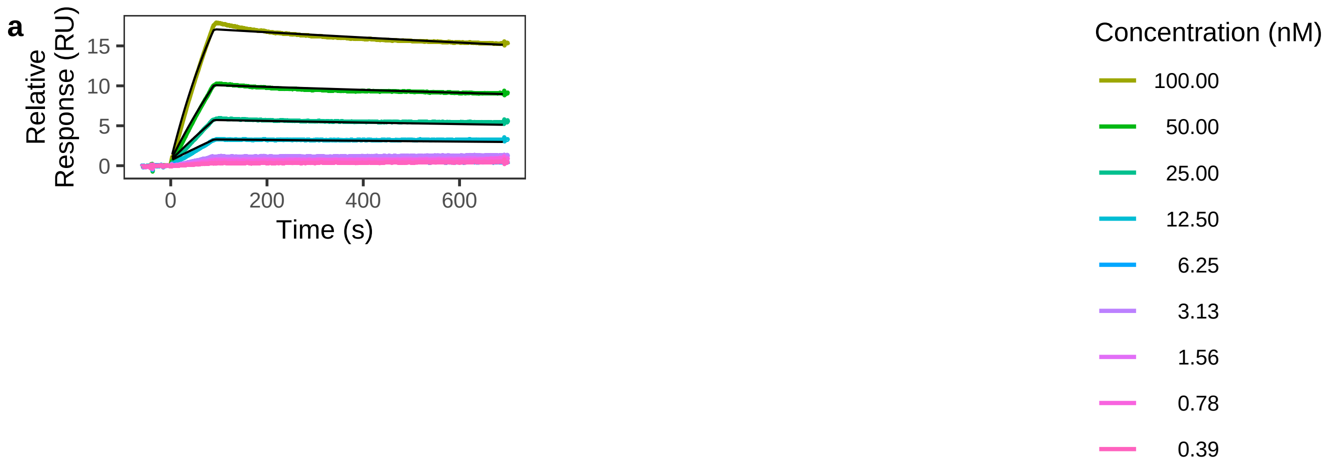

KLRC1 (13905−H07H) Surface Plasmon Resonance Sensorgrams

| Panel | Antibody Name | Antibody Type | KD1 (nM) | kon1 (1/nM·s) | koff1 (1/s) | rmax1 (RU) | KD2 (nM) | kon2 (1/nM·s) | koff2 (1/s) | rmax2 (RU) | drift (RU/s) | offset (RU) |
|---|---|---|---|---|---|---|---|---|---|---|---|---|
| a | CYC038 | Human | 9.894 | 4.20e−05 | 4.15e−04 | 11.7 | | | | | −9.69e−04 | |

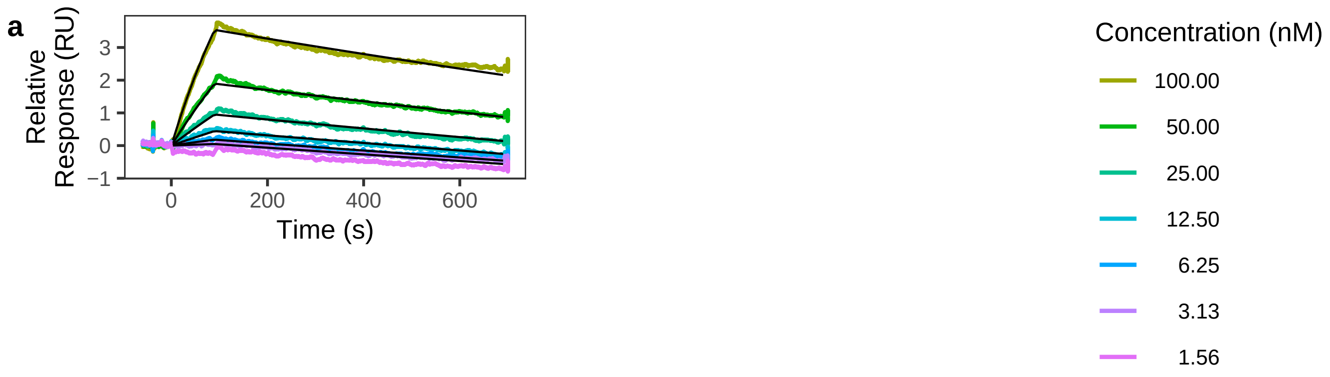

### MAGEA3 (ab99958) Surface Plasmon Resonance Sensorgrams

| Panel | Antibody Name | Antibody Type | KD1 (nM) | kon1 (1/nM·s) | koff1 (1/s) | rmax1 (RU) | KD2 (nM) | kon2 (1/nM·s) | koff2 (1/s) | rmax2 (RU) | drift (RU/s) | offset (RU) |
|---|---|---|---|---|---|---|---|---|---|---|---|---|
| a | CYC066 | Human | 0.514 | 3.44e−04 | 1.77e−04 | 59.4 | 6.725 | 8.83e−05 | 5.94e−04 | 41.4 | | |
| b | ab223162 | Rabbit | 0.148 | 7.40e−04 | 1.10e−04 | 18.1 | 0.181 | 1.92e−03 | 3.48e−04 | 18.4 | | |

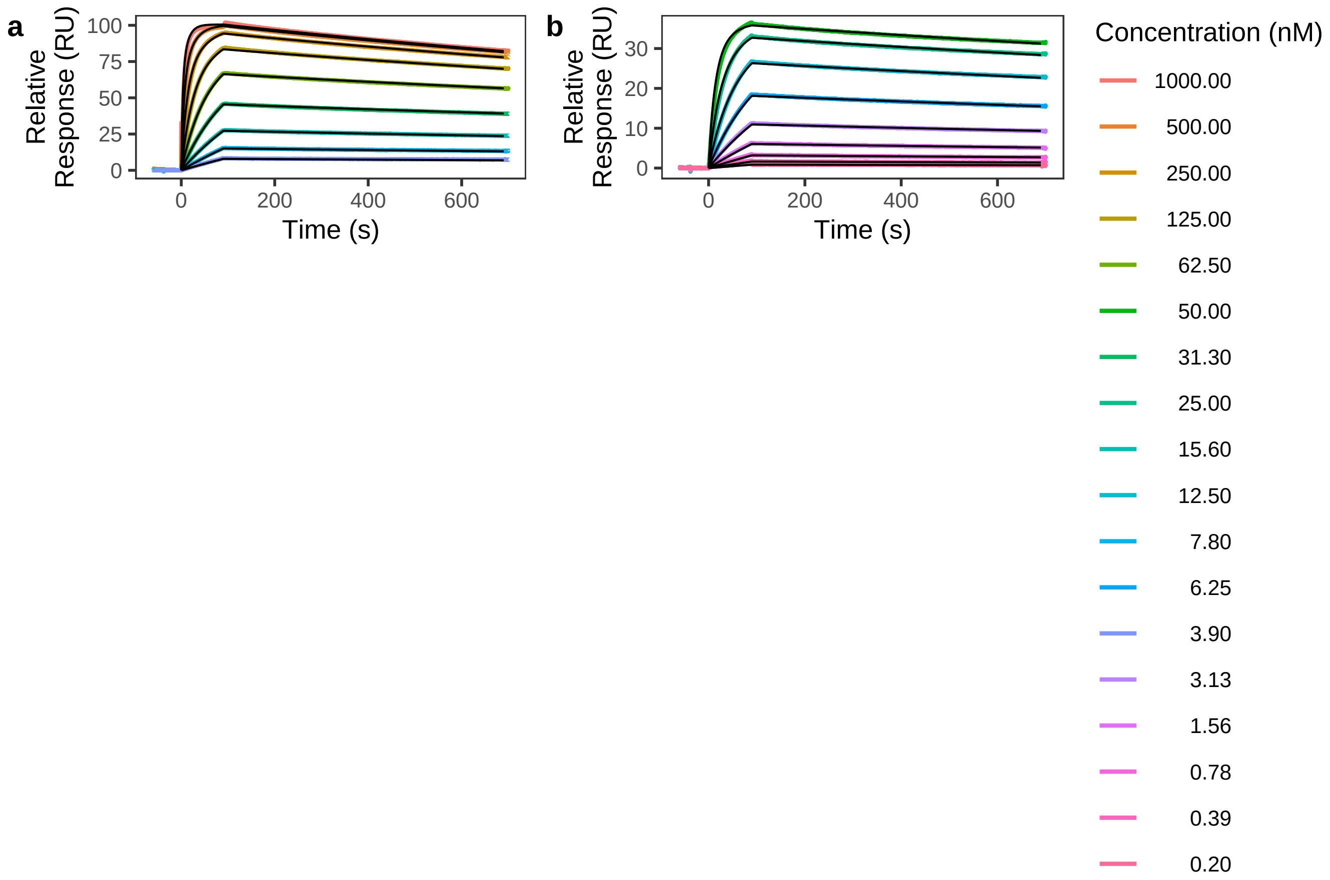

### TGFBI (TP300411) Surface Plasmon Resonance Sensorgrams

| Panel | Antibody Name | Antibody Type | KD1 (nM) | kon1 (1/nM·s) | koff1 (1/s) | rmax1 (RU) | KD2 (nM) | kon2 (1/nM·s) | koff2 (1/s) | rmax2 (RU) | drift (RU/s) | offset (RU) |
|---|---|---|---|---|---|---|---|---|---|---|---|---|
| a | CYC168 | Human | 0.502 | 3.41e−04 | 1.71e−04 | 12.1 | 5.340 | 1.39e−03 | 7.40e−03 | 1.9 | | |
| b | CYC168 | Human | 0.476 | 3.40e−04 | 1.62e−04 | 13.8 | 191.130 | 2.72e−03 | 5.19e−01 | 36.4 | | −1.7 |
| c | HPA008612 | Rabbit | 0.037 | 6.46e−03 | 2.38e−04 | 2.7 | 0.485 | 8.75e−04 | 4.25e−04 | 5.4 | | |
| d | HPA008612 | Rabbit | 1.098 | 3.45e−04 | 3.79e−04 | 8.2 | 659.384 | 3.96e−04 | 2.61e−01 | 20.3 | | −0.1 |
| e | HPA017019 | Rabbit | 0.041 | 2.84e−03 | 1.16e−04 | 30.6 | 10.465 | 5.87e−04 | 6.15e−03 | 6.0 | | −2.2 |
| f | HPA017019 | Rabbit | 0.261 | 5.16e−04 | 1.35e−04 | 23.1 | 357.089 | 1.88e−03 | 6.71e−01 | 130.2 | | −0.7 |

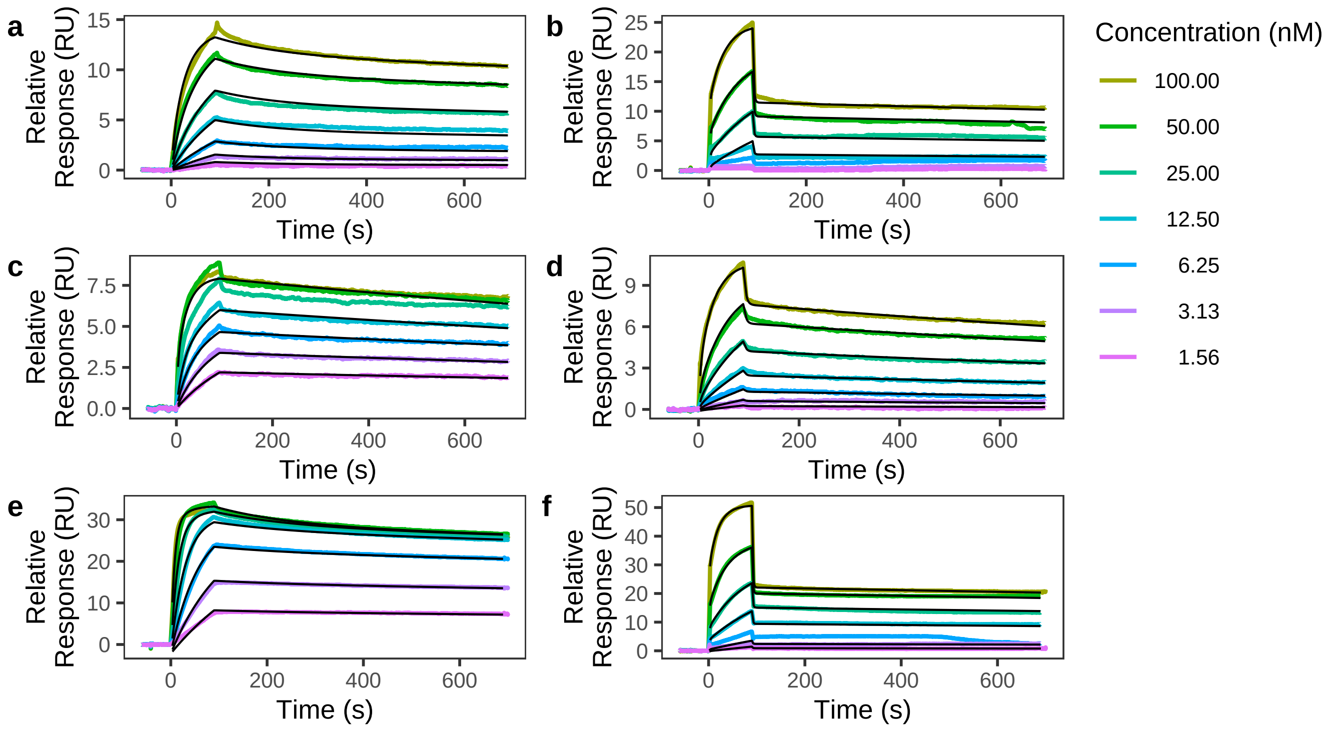

### TXLNA (TP310301) Surface Plasmon Resonance Sensorgrams

| Panel | Antibody Name | Antibody Type | KD1 (nM) | kon1 (1/nM·s) | koff1 (1/s) | rmax1 (RU) | KD2 (nM) | kon2 (1/nM·s) | koff2 (1/s) | rmax2 (RU) | drift (RU/s) | offset (RU) |
|---|---|---|---|---|---|---|---|---|---|---|---|---|
| a | CYC086 | Human | 10.860 | 8.91e−05 | 9.67e−04 | 9.3 | 191.602 | 1.54e−03 | 2.96e−01 | 14.5 | | |
| b | CYC086 | Human | 28.412 | 2.05e−05 | 5.82e−04 | 14.3 | 212.243 | 2.17e−03 | 4.60e−01 | 11.6 | | |
| c | CYC086 | Human | 0.716 | 2.12e−04 | 1.52e−04 | 6.2 | 167.228 | 1.12e−04 | 1.88e−02 | 2.8 | | |
| d | HPA045383 | Rabbit | 1.273 | 2.03e−04 | 2.59e−04 | 28.2 | 389.790 | 1.15e−03 | 4.48e−01 | 32.3 | | 0.8 |
| e | HPA045383 | Rabbit | 1.090 | 1.82e−04 | 1.99e−04 | 19.5 | 301.971 | 2.13e−03 | 6.44e−01 | 27.0 | | |
| f | HPA072477 | Rabbit | 0.140 | 2.23e−03 | 3.12e−04 | 4.2 | 0.732 | 5.03e−04 | 3.68e−04 | 27.8 | | |

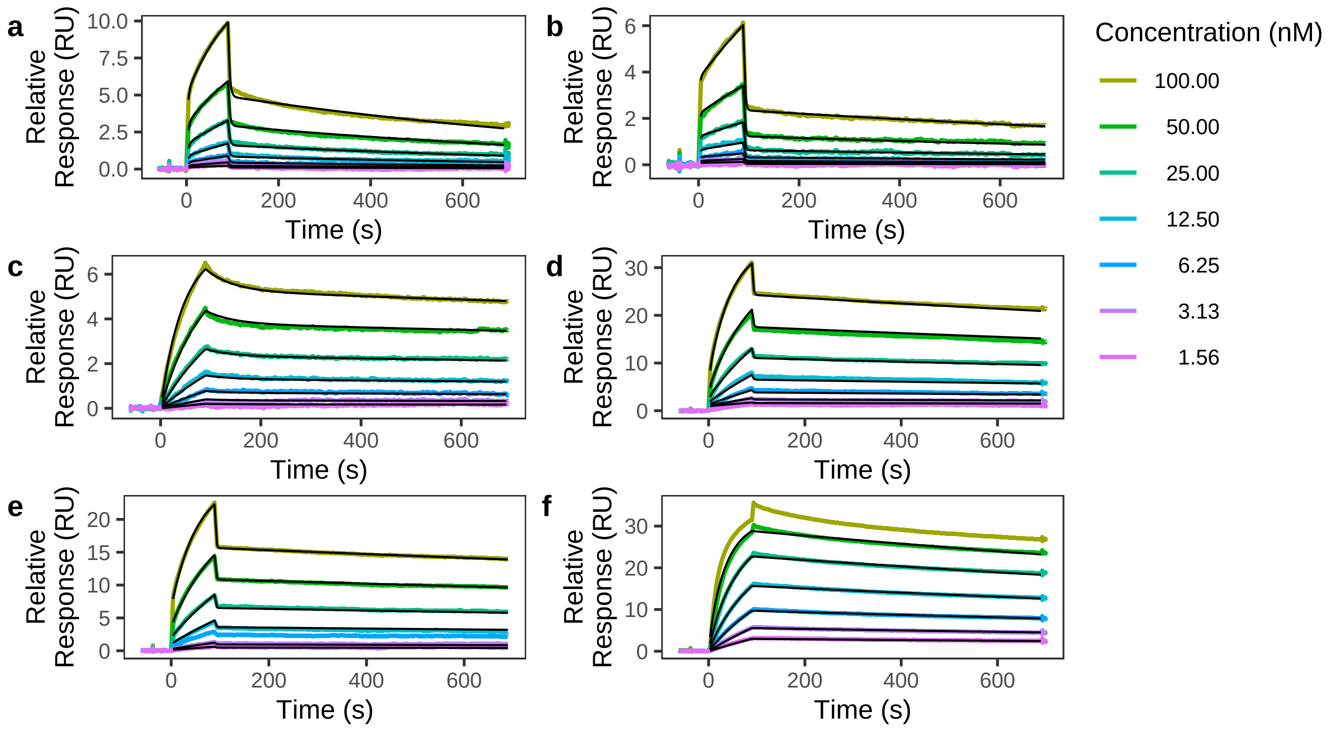

### UBQLN2 (ab207109) Surface Plasmon Resonance Sensorgrams

| Panel | Antibody Name | Antibody Type | KD1 (nM) | kon1 (1/nM·s) | koff1 (1/s) | rmax1 (RU) | KD2 (nM) | kon2 (1/nM·s) | koff2 (1/s) | rmax2 (RU) | drift (RU/s) | offset (RU) |
|---|---|---|---|---|---|---|---|---|---|---|---|---|
| a | CYC049 | Human | 0.991 | 3.22e−03 | 3.19e−03 | 7.7 | 66.438 | 8.09e−04 | 5.37e−02 | 73.1 | | |
| b | CYC049 | Human | 0.711 | 3.46e−03 | 2.46e−03 | 6.0 | 40.364 | 8.66e−04 | 3.50e−02 | 52.4 | | |

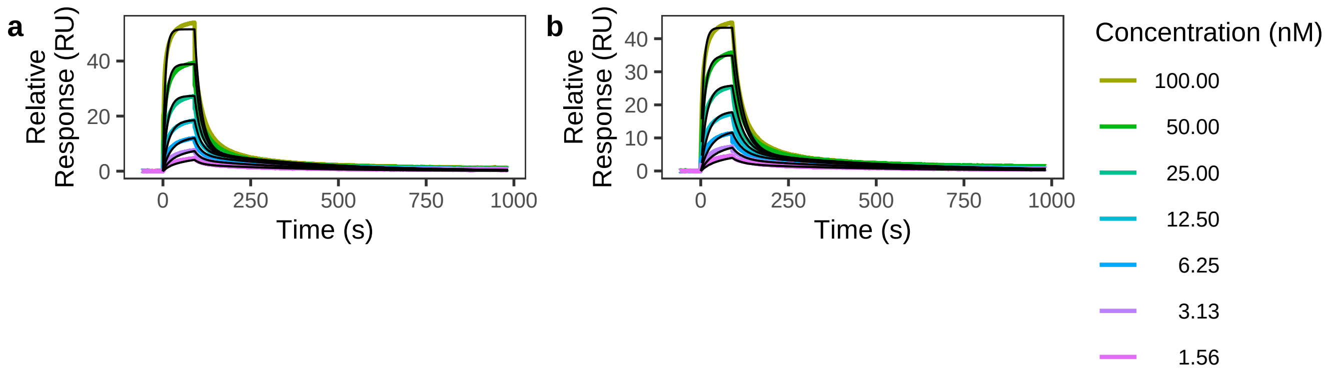

### UBQLN2 (ab207109) Surface Plasmon Resonance Sensorgrams

| Panel | Antibody Name | Antibody Type | KD1 (nM) | kon1 (1/nM·s) | koff1 (1/s) | rmax1 (RU) | KD2 (nM) | kon2 (1/nM·s) | koff2 (1/s) | rmax2 (RU) | drift (RU/s) | offset (RU) |
|---|---|---|---|---|---|---|---|---|---|---|---|---|
| a | CYC146 | Human | 13.102 | 1.09e−03 | 1.43e−02 | 3.1 | 604.712 | 4.03e−04 | 2.44e−01 | 146.8 | −6.35e−04 | |

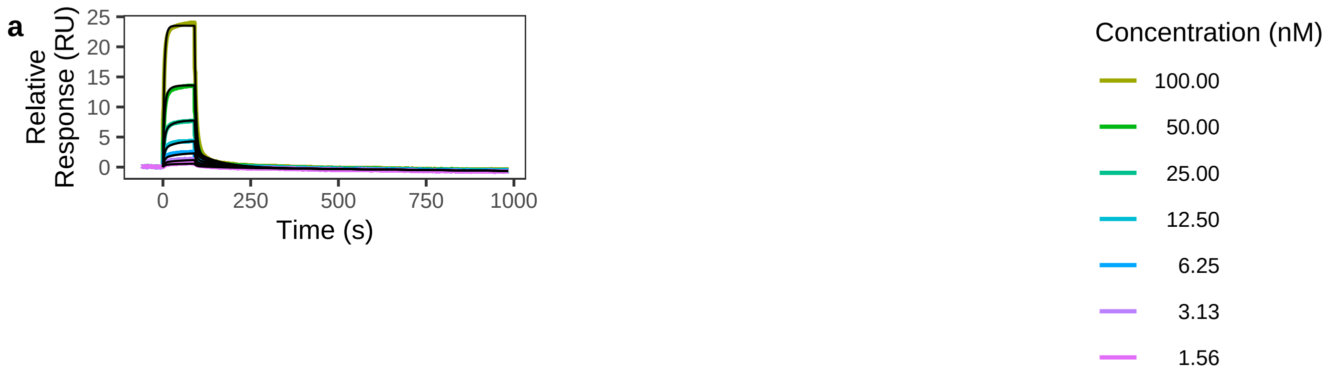
