## Supplementary Figures for "The landscape of high-affinity human antibodies against intratumoral antigens": Supplementary Figure 4.pdf

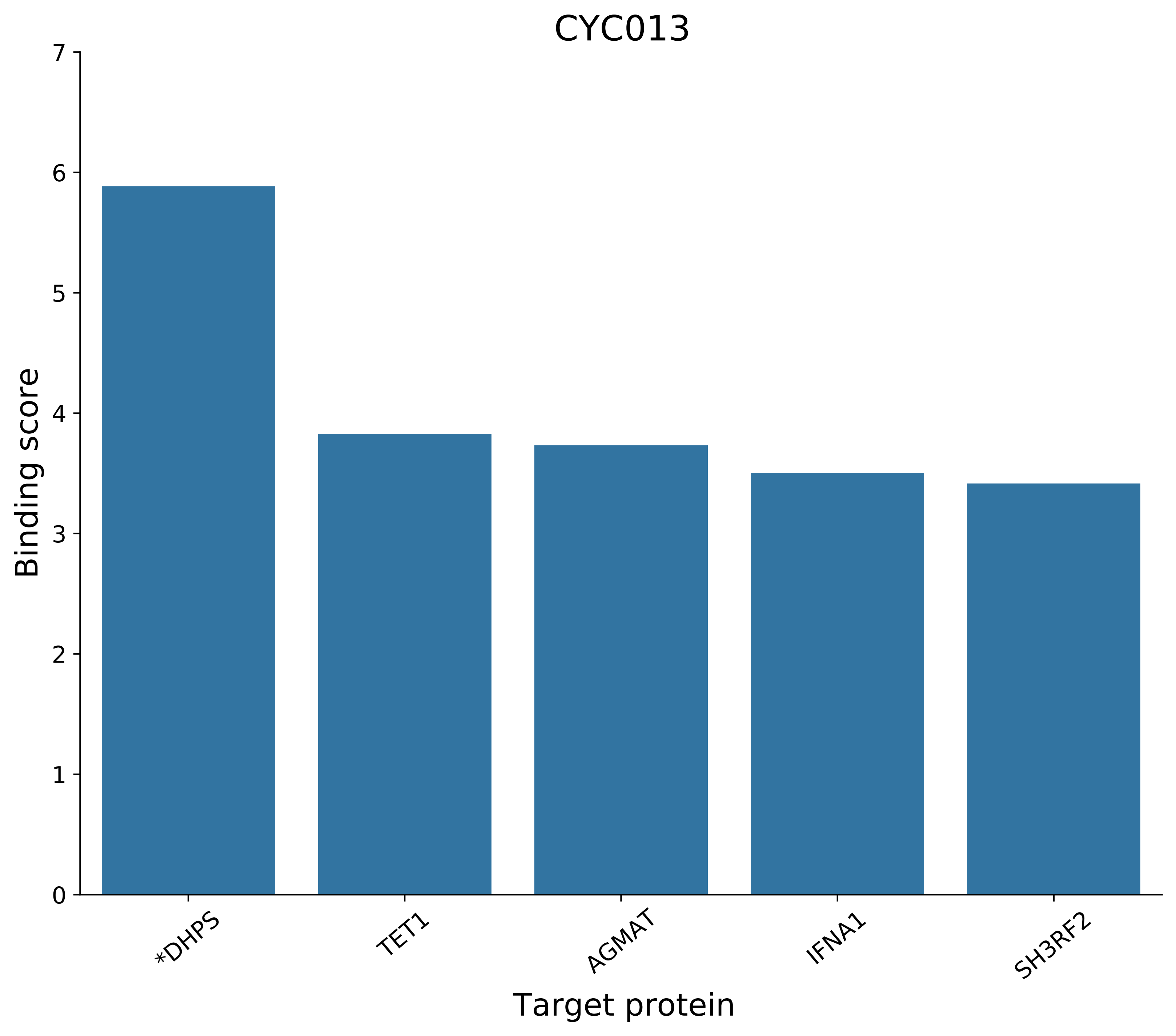
CYC013
Binding score
0
1
2
3
4
5
6
7
*DHPS
TET1
AGMAT
IFNA1
SH3RF2
Target protein

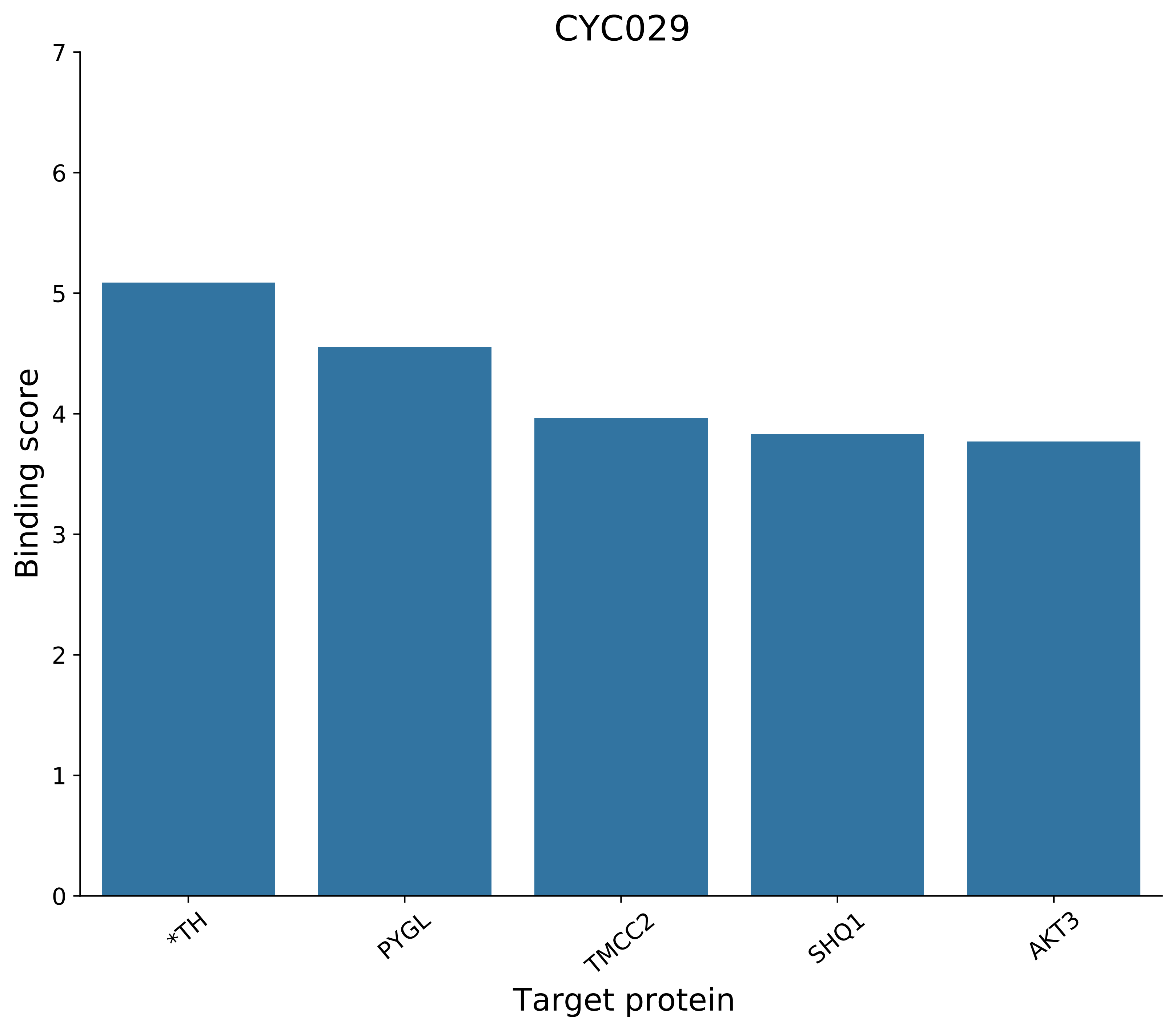

CYC029
7
6
5
4
3
2
1
0
Binding score
*TH
PYGL
TMCC2
SHQ1
AKT3
Target protein

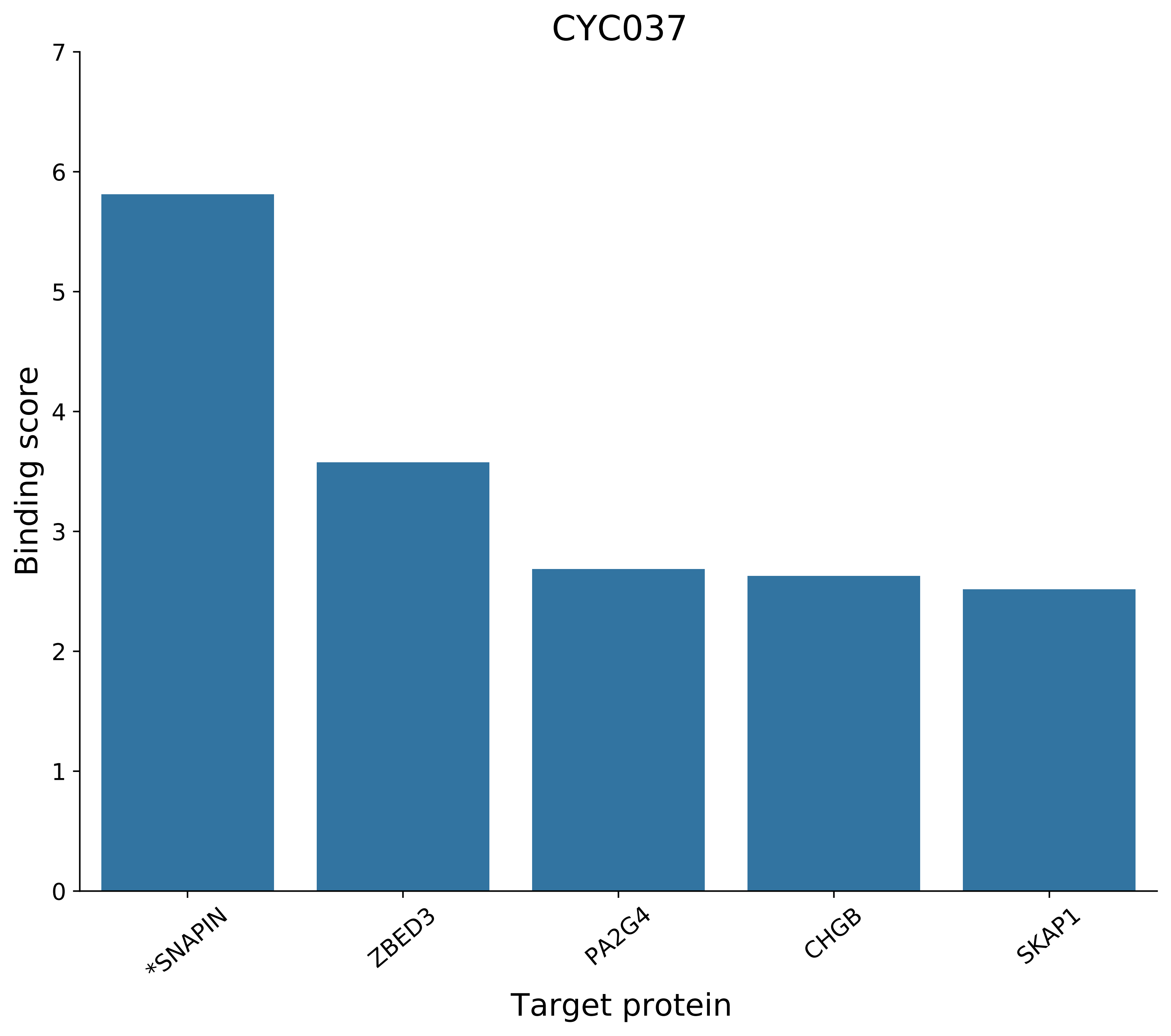
CYC037
Binding score
0
1
2
3
4
5
6
7
*SNAPIN
ZBED3
PA2G4
CHGB
SKAP1
Target protein

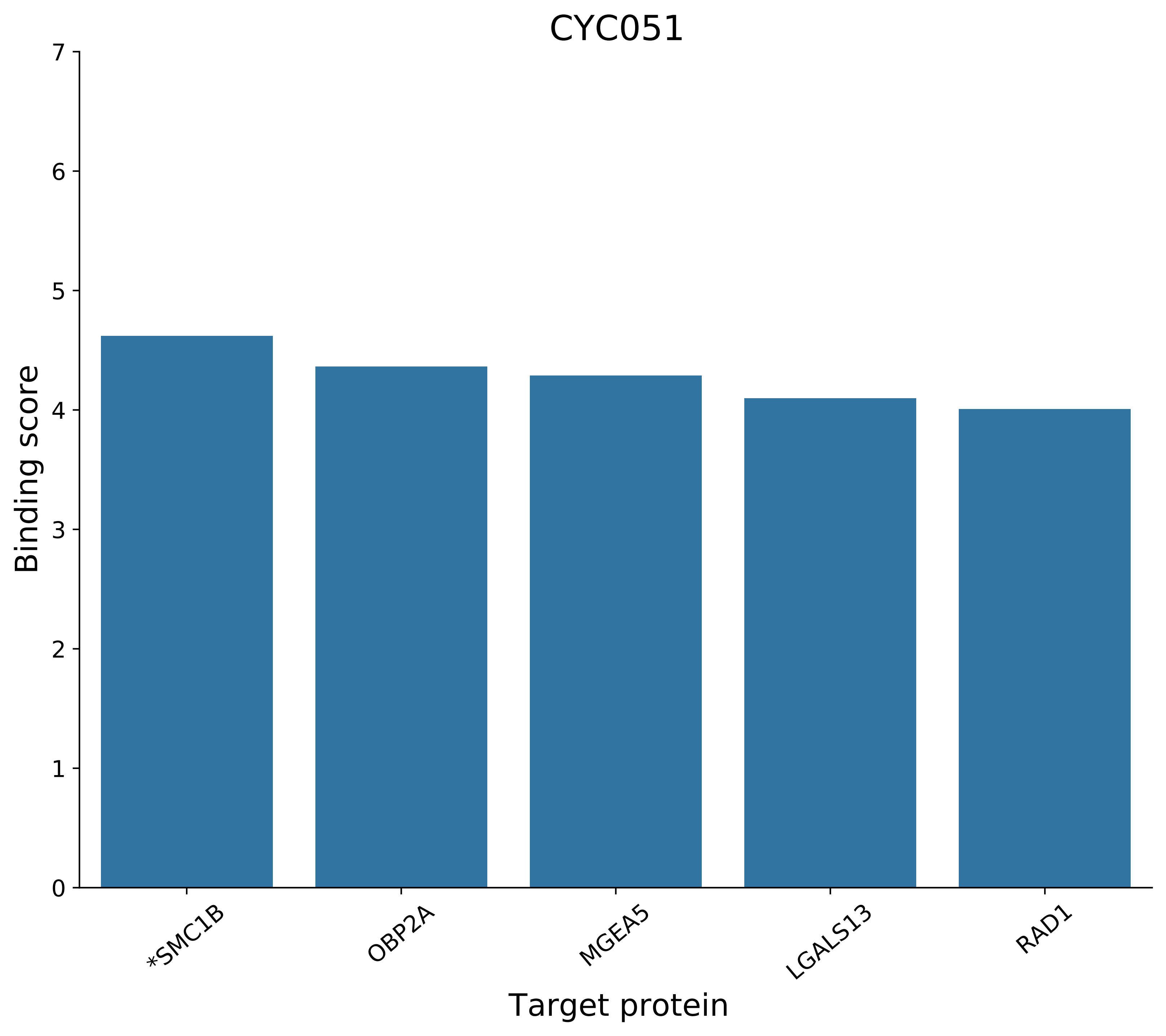
CYC051
Binding score
0
1
2
3
4
5
6
7
*SMC1B
OBP2A
MGEA5
LGALS13
RAD1
Target protein

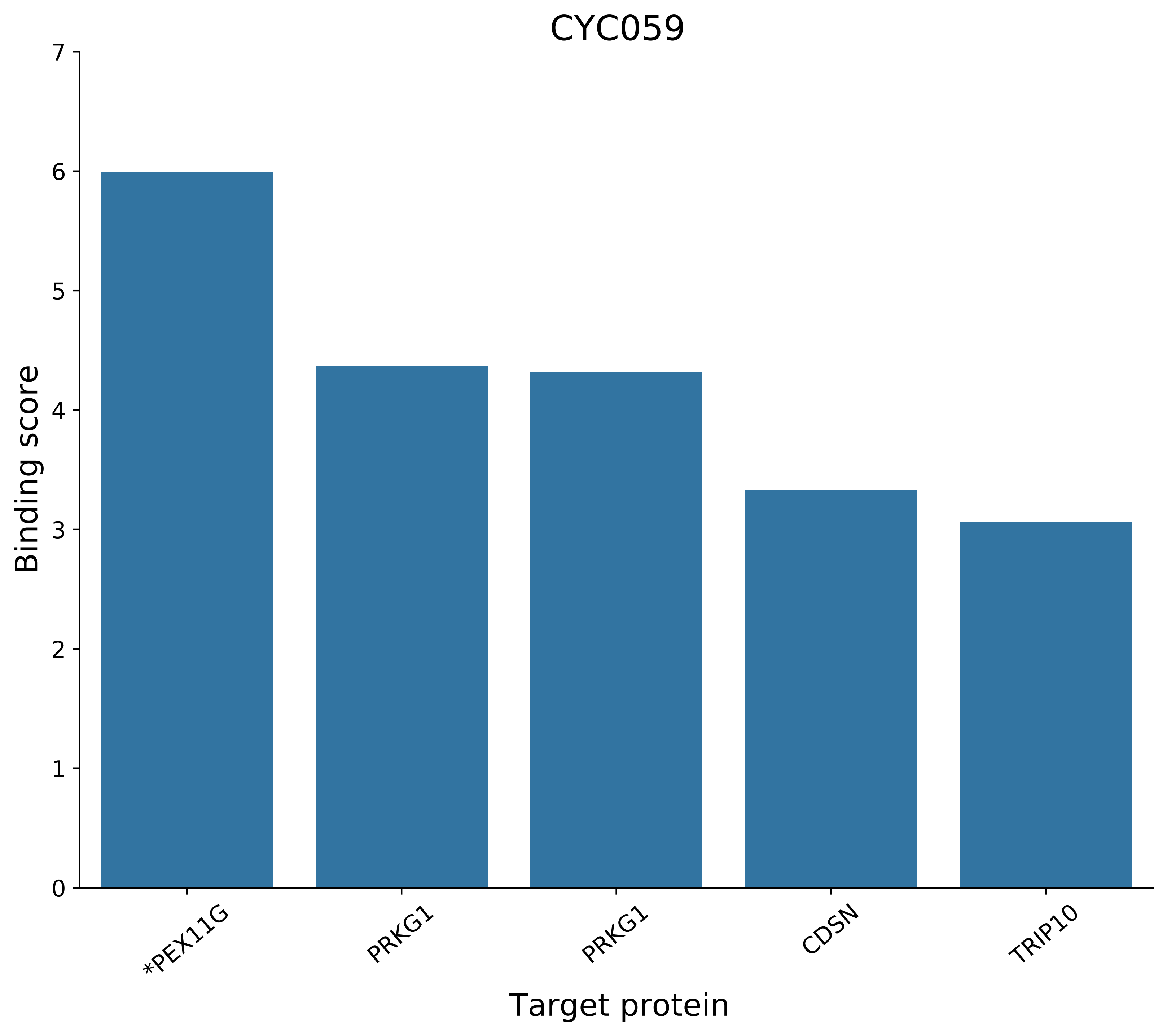
CYC059
Binding score
0
1
2
3
4
5
6
7
*PEX11G
PRKG1
PRKG1
CDSN
TRIP10
Target protein

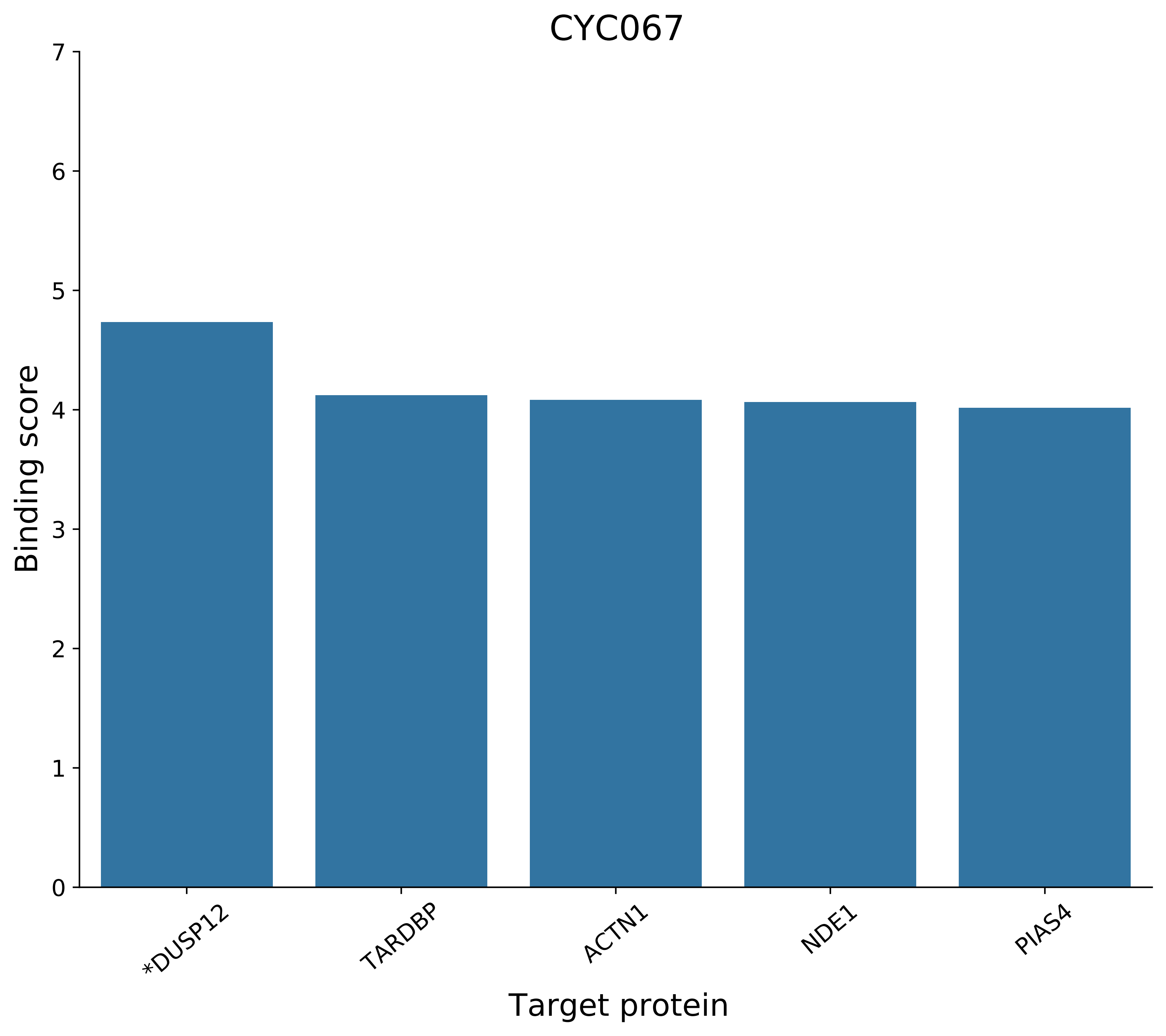

CYC067
Binding score
Target protein
0
1
2
3
4
5
6
7
*DUSP12
TARDBP
ACTN1
NDE1
PIAS4

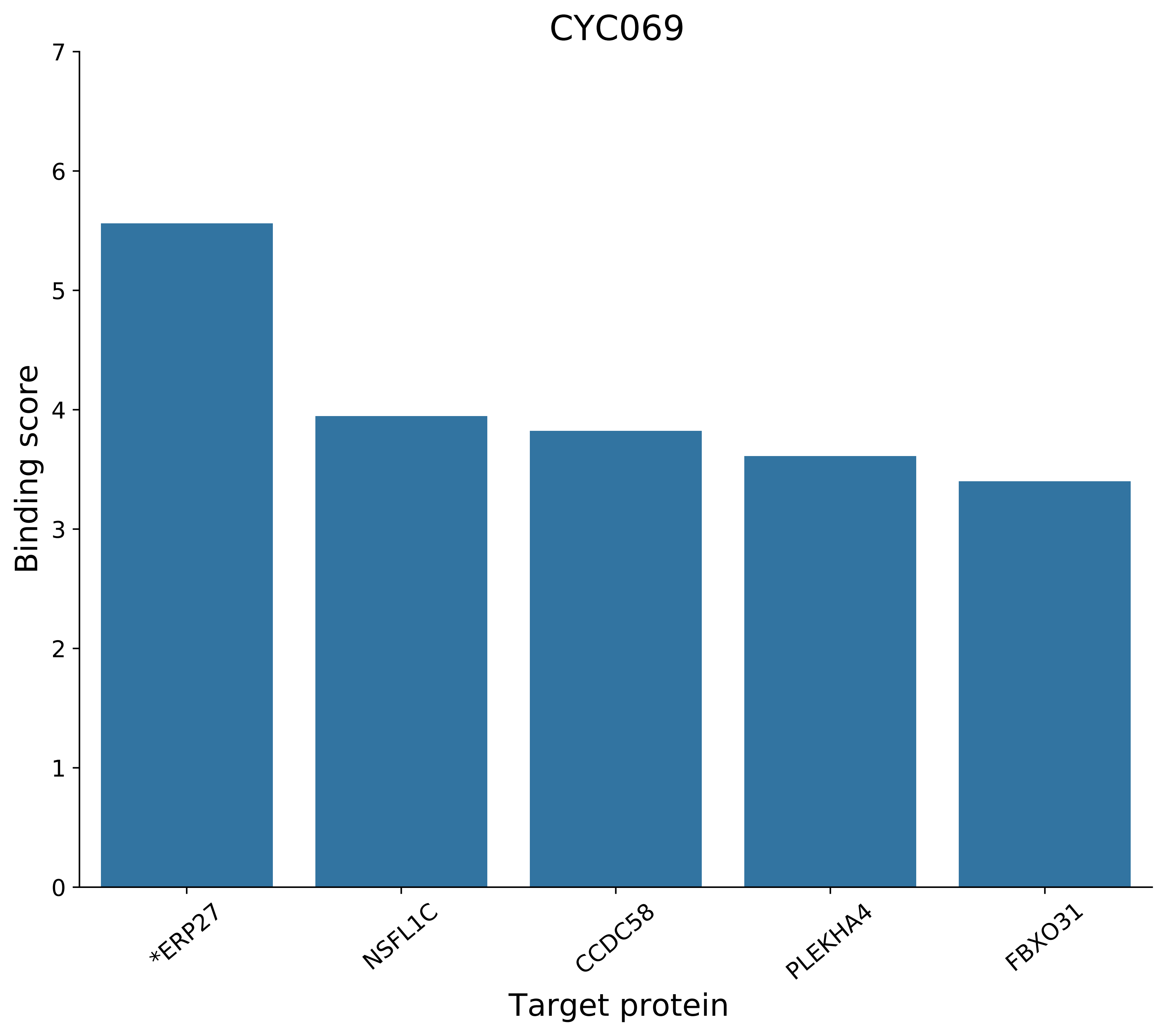
CYC069
Binding score
0
1
2
3
4
5
6
7
*ERP27
NSFL1C
CCDC58
PLEKHA4
FBXO31
Target protein

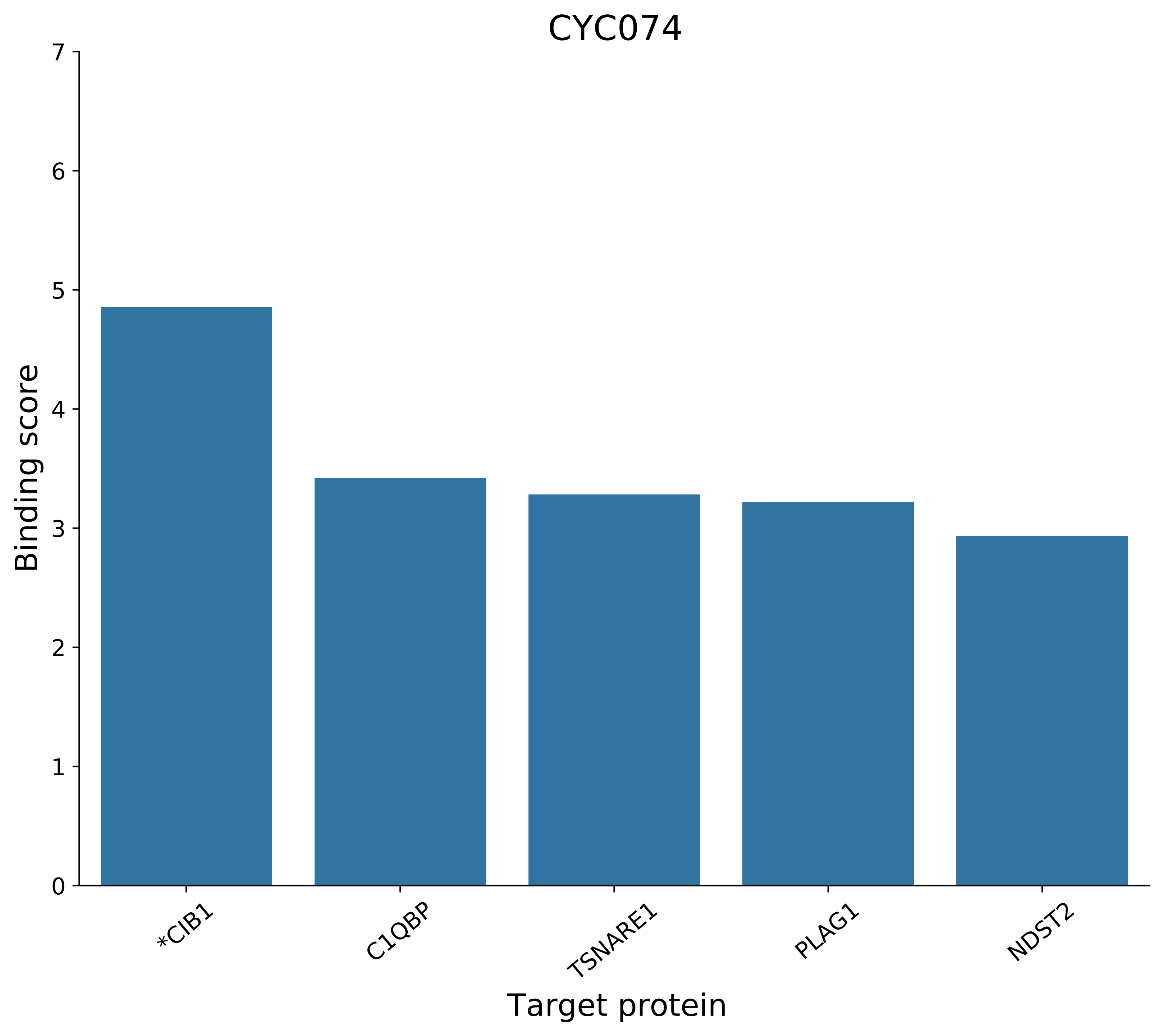
CYC074
Binding score
Target protein
0
1
2
3
4
5
6
7
*CIB1
C1QBP
TSNARE1
PLAG1
NDST2

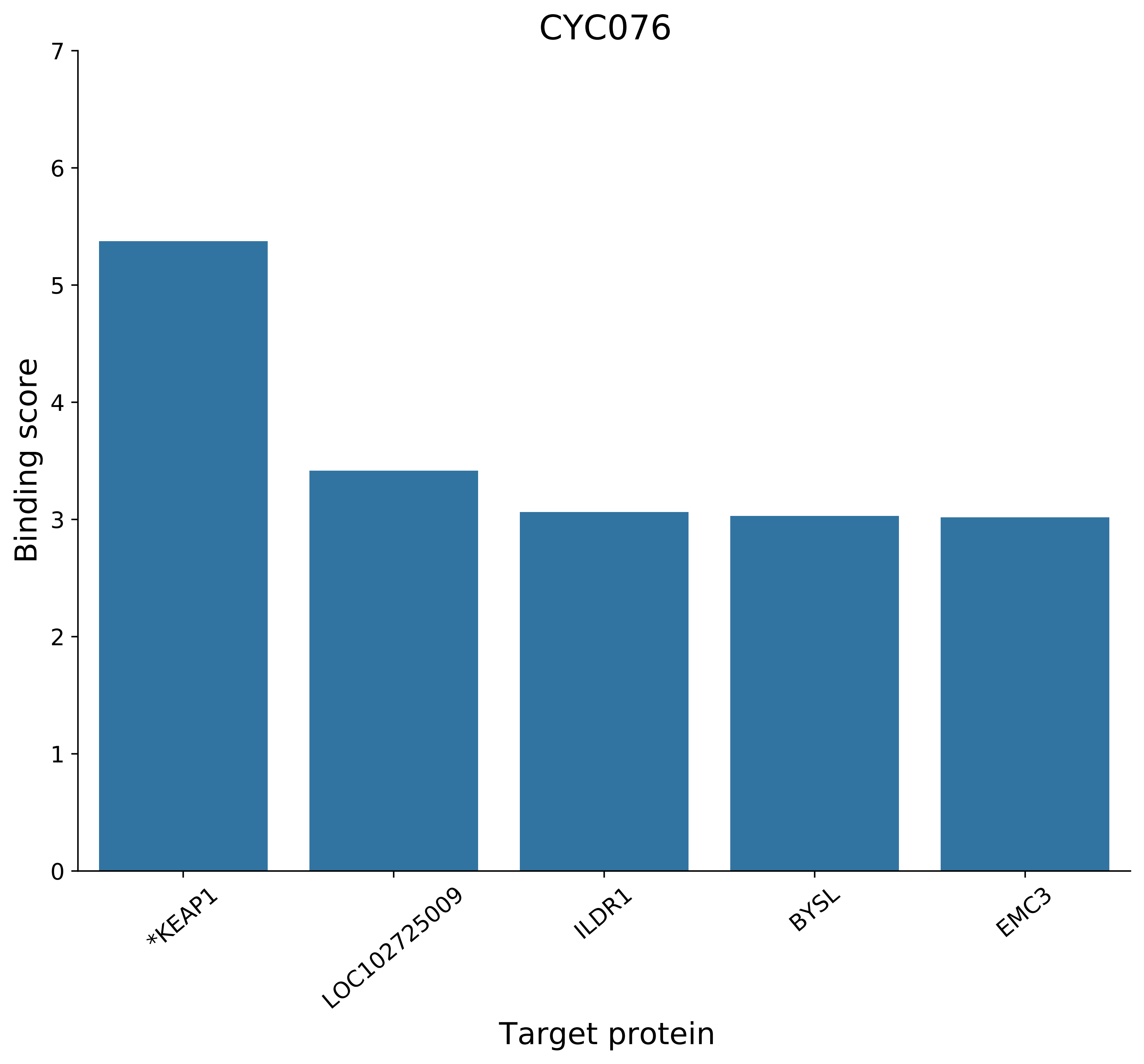
CYC076
Binding score
0
1
2
3
4
5
6
7
*KEAP1
LOC102725009
ILDR1
BYSL
EMC3
Target protein

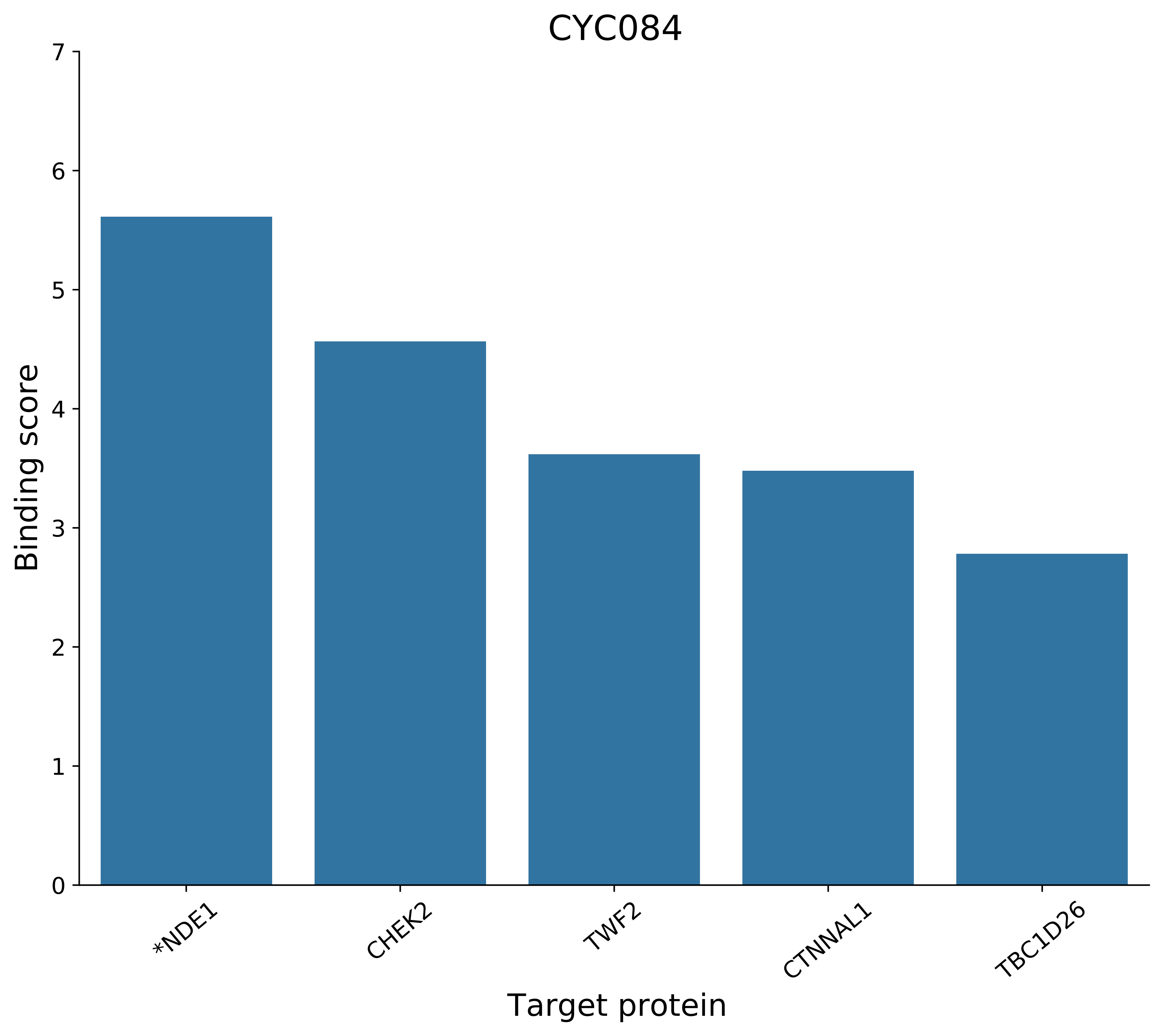
CYC084
Binding score
Target protein
*NDE1
CHEK2
TWF2
CTNNAL1
TBC1D26
0
1
2
3
4
5
6
7

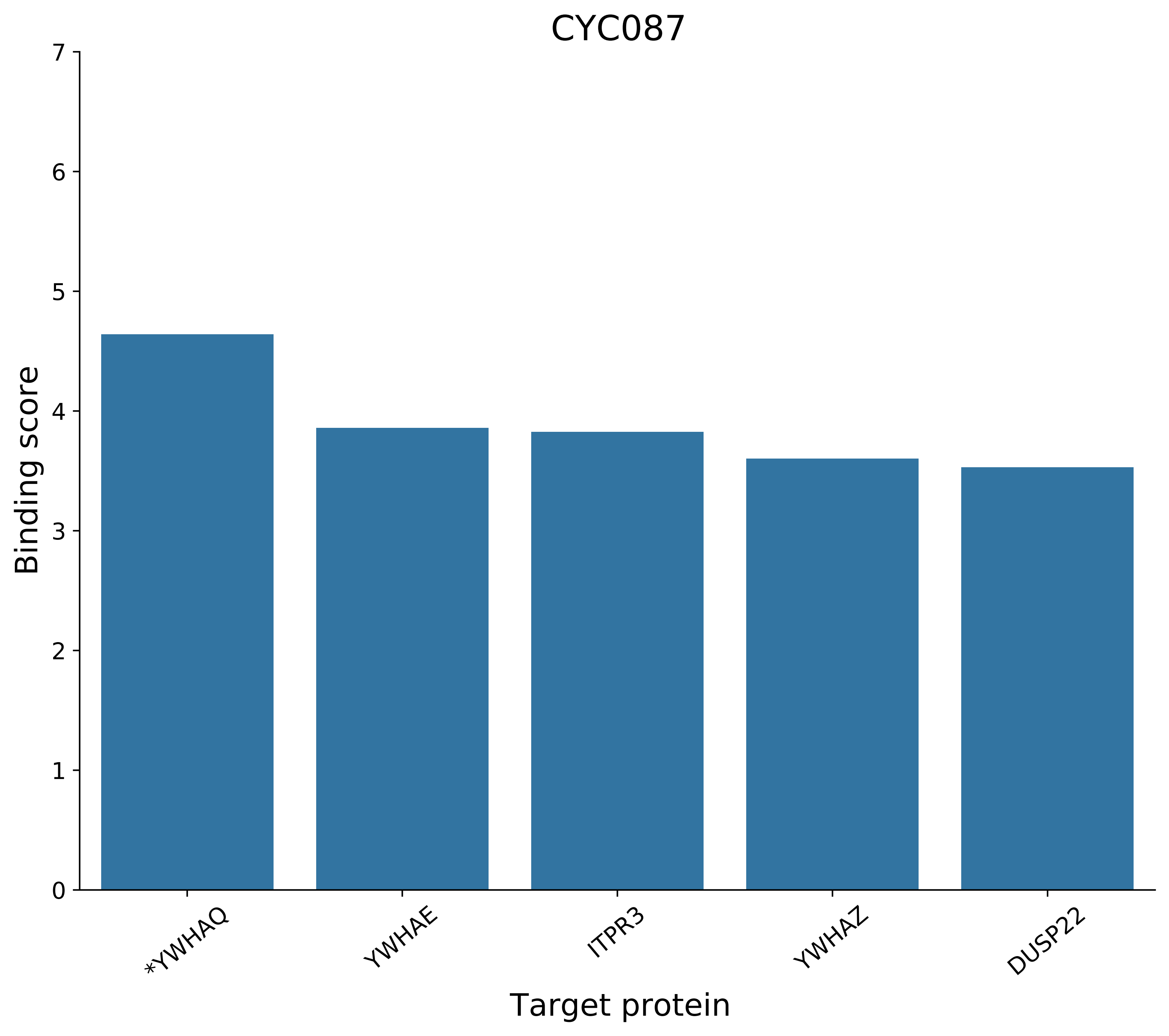
CYC087
Binding score
Target protein
0
1
2
3
4
5
6
7
*YWHAQ
YWHAE
ITPR3
YWHAZ
DUSP22

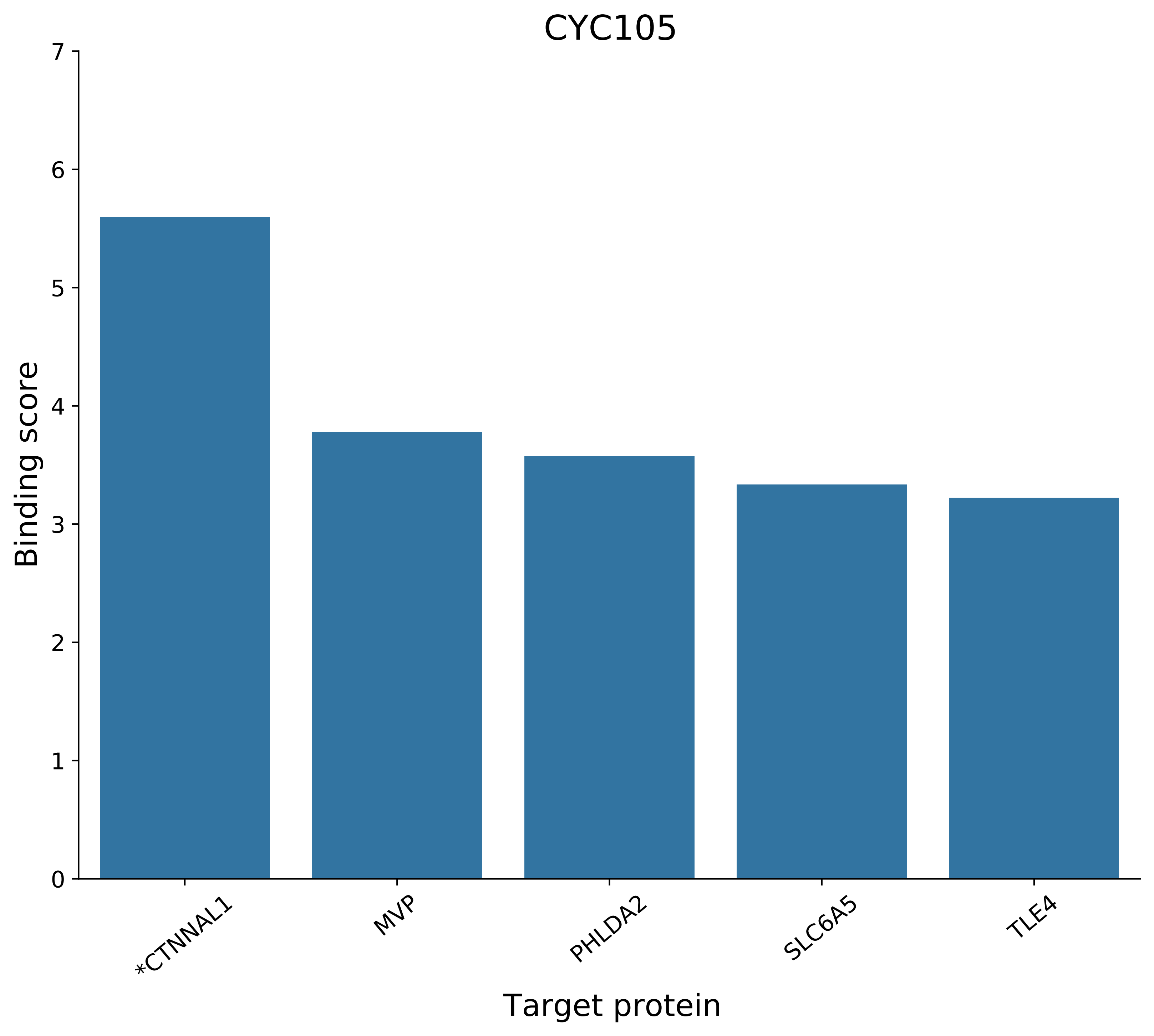
CYC105
Binding score
7
6
5
4
3
2
1
0
*CTNNAL1
MVP
PHLDA2
SLC6A5
TLE4
Target protein

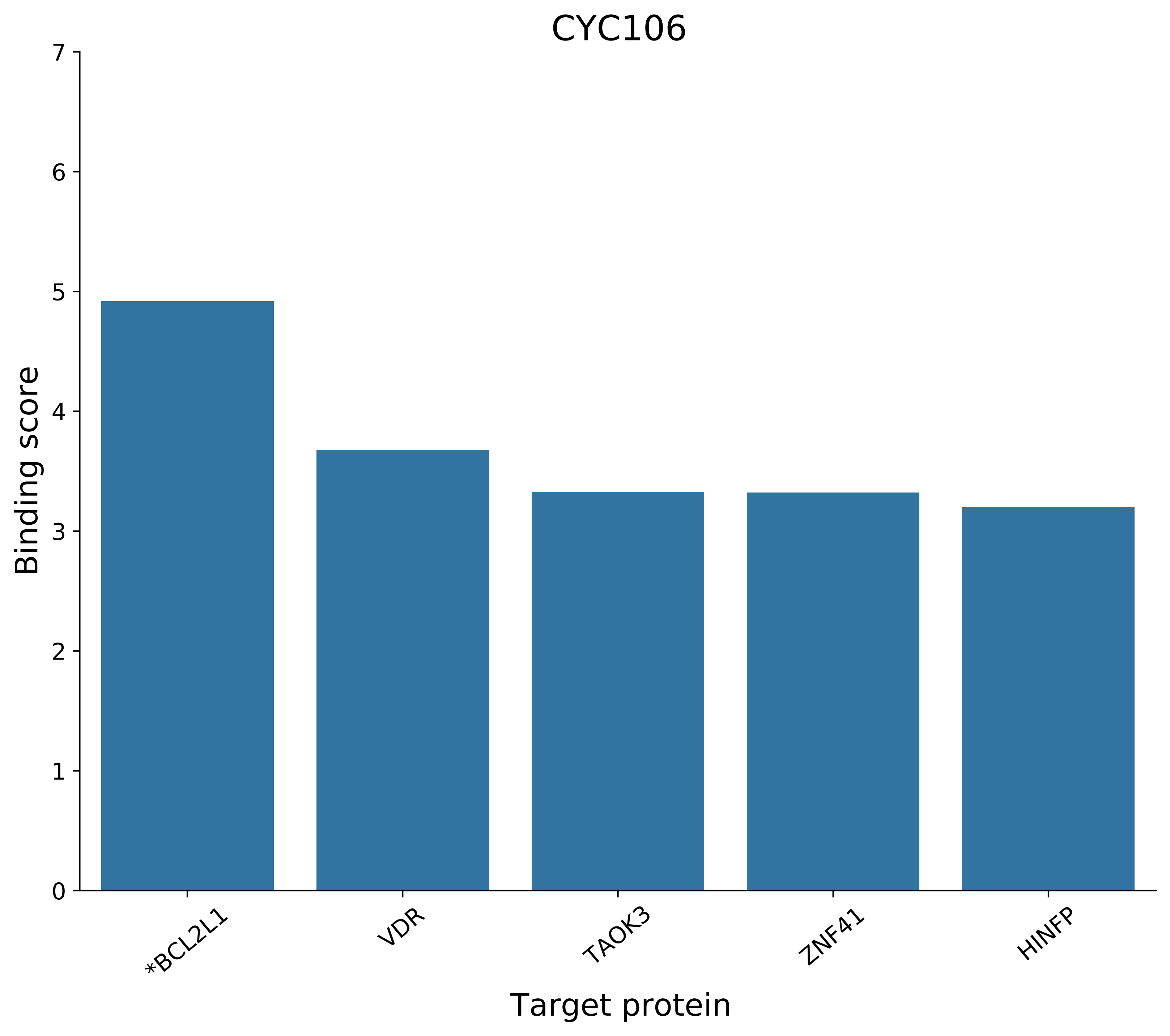
CYC106
Binding score
7
6
5
4
3
2
1
0
*BCL2L1
VDR
TAOK3
ZNF41
HINFP
Target protein

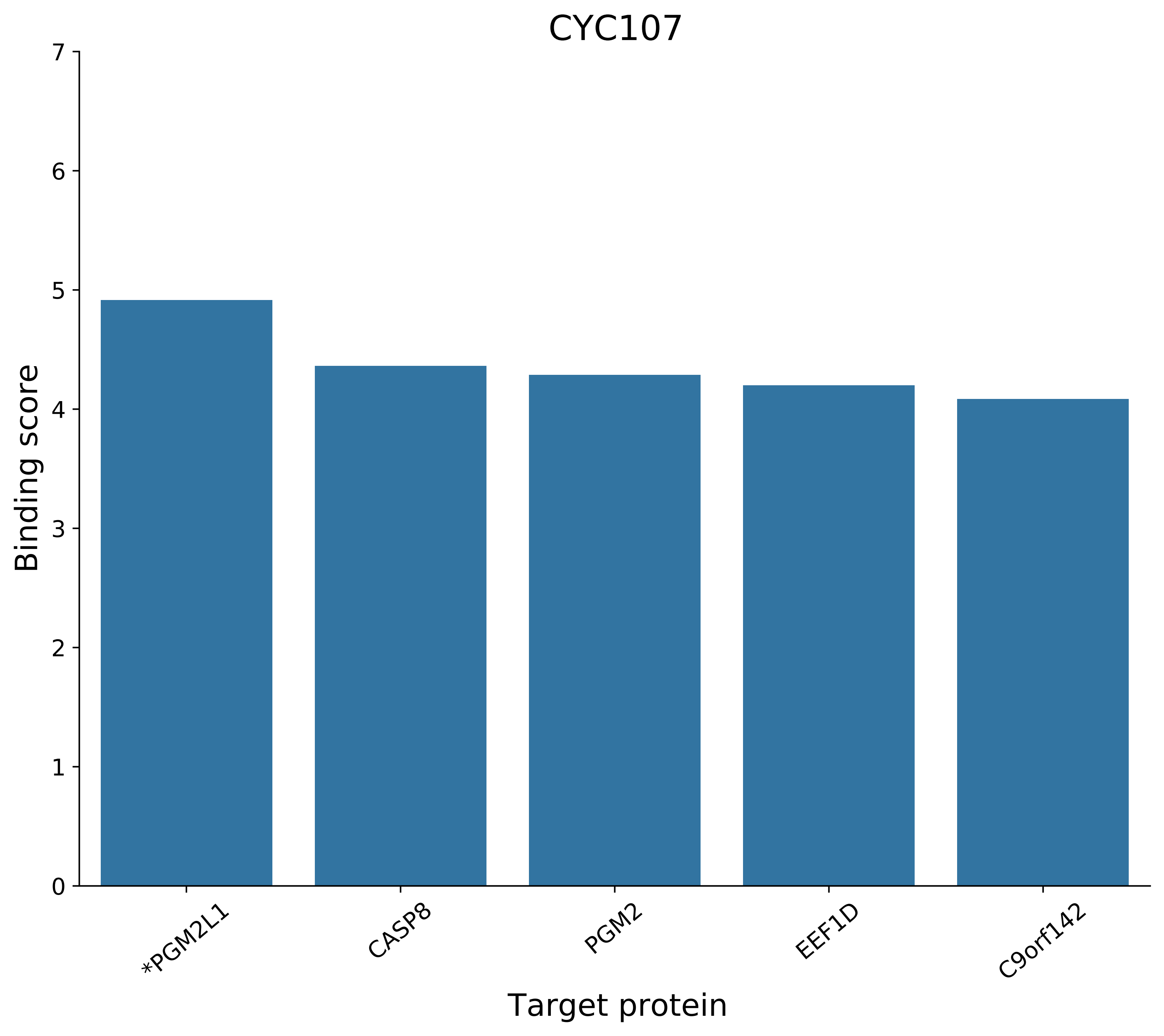
CYC107
Binding score
7
6
5
4
3
2
1
0
*PGM2L1
CASP8
PGM2
EEF1D
C9orf142
Target protein

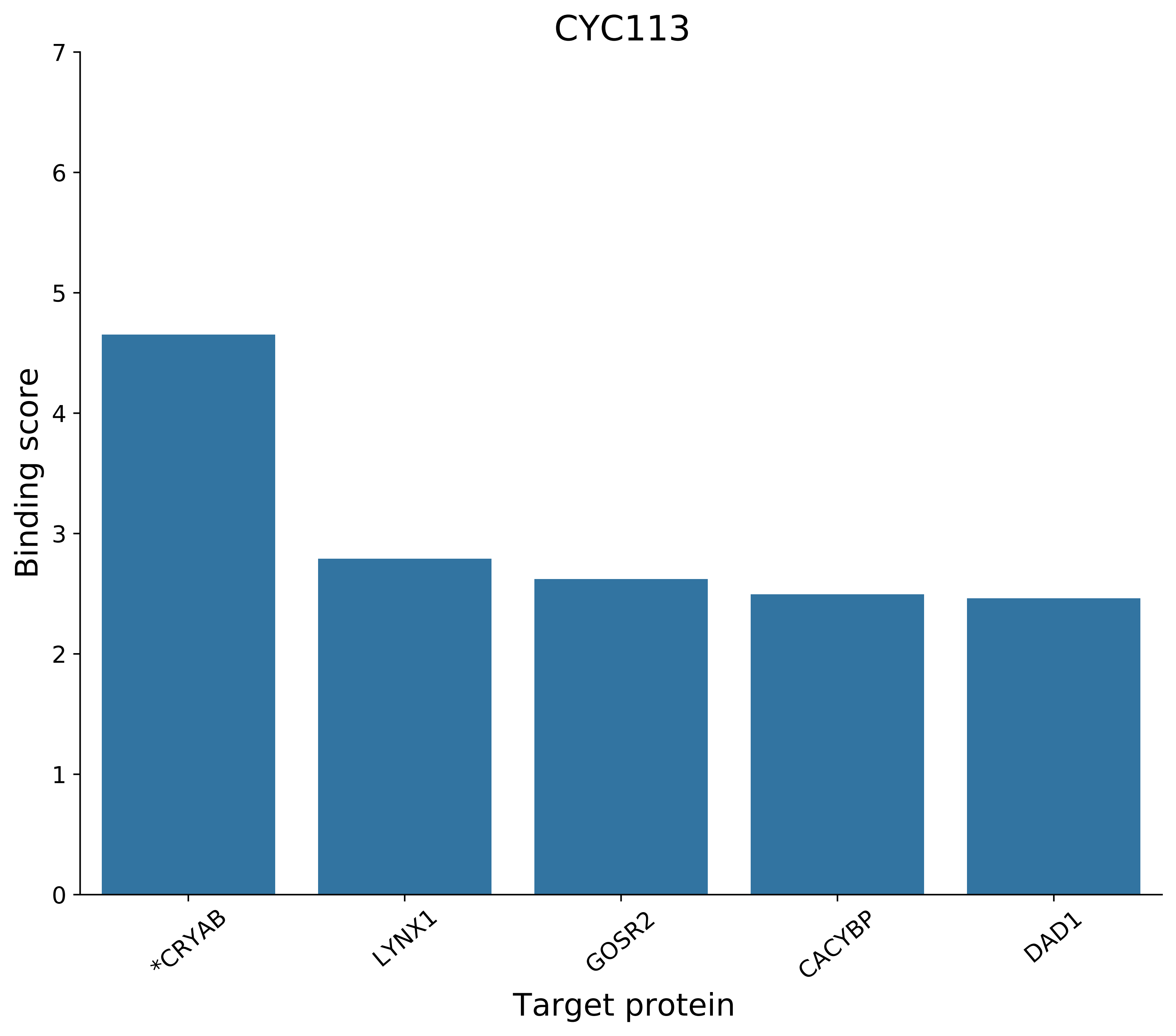
CYC113
Binding score
0
1
2
3
4
5
6
7
*CRYAB
LYNX1
GOSR2
CACYBP
DAD1
Target protein

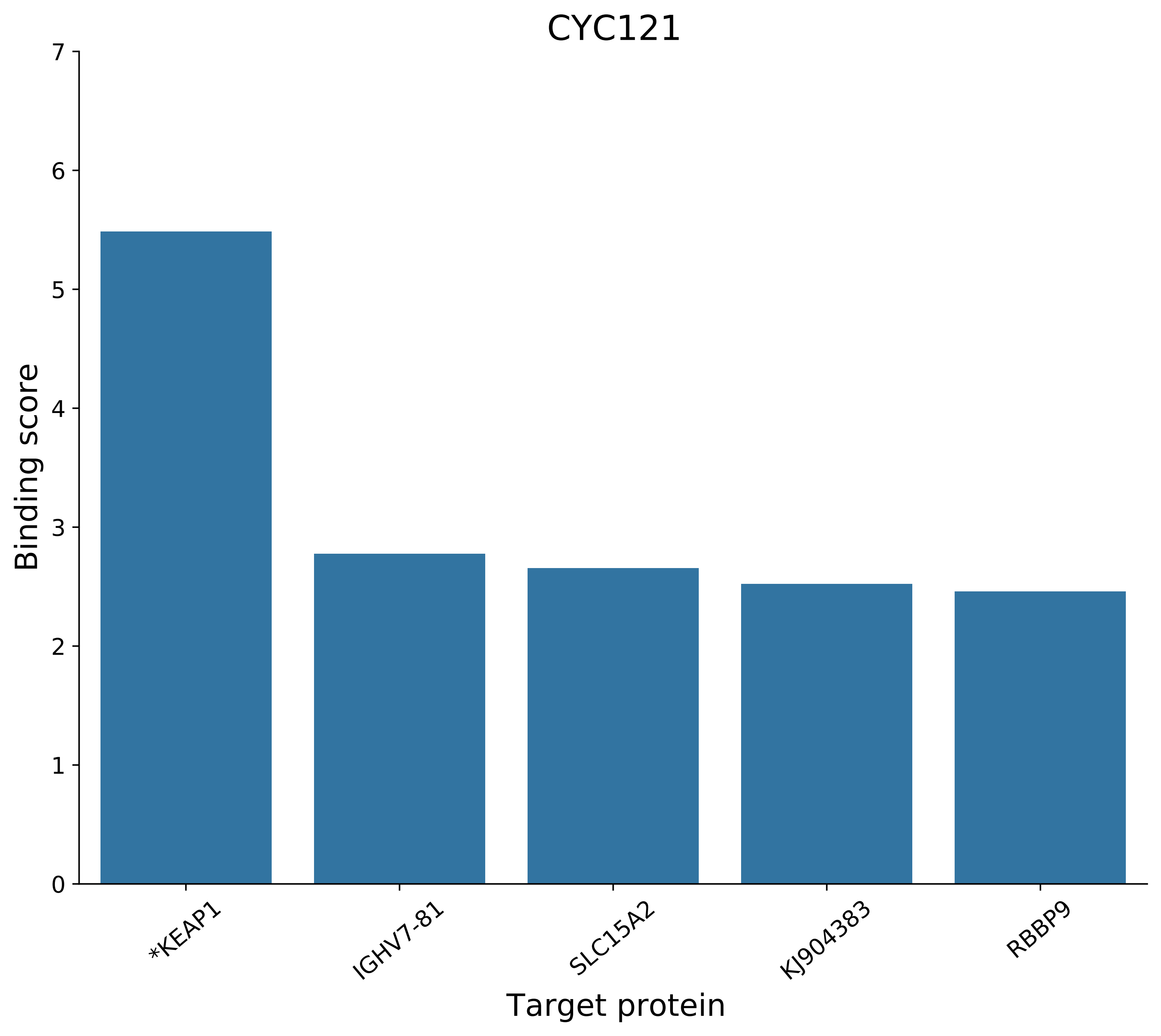
CYC121
Binding score
0
1
2
3
4
5
6
7
*KEAP1
IGHV7-81
SLC15A2
KJ904383
RBBP9
Target protein

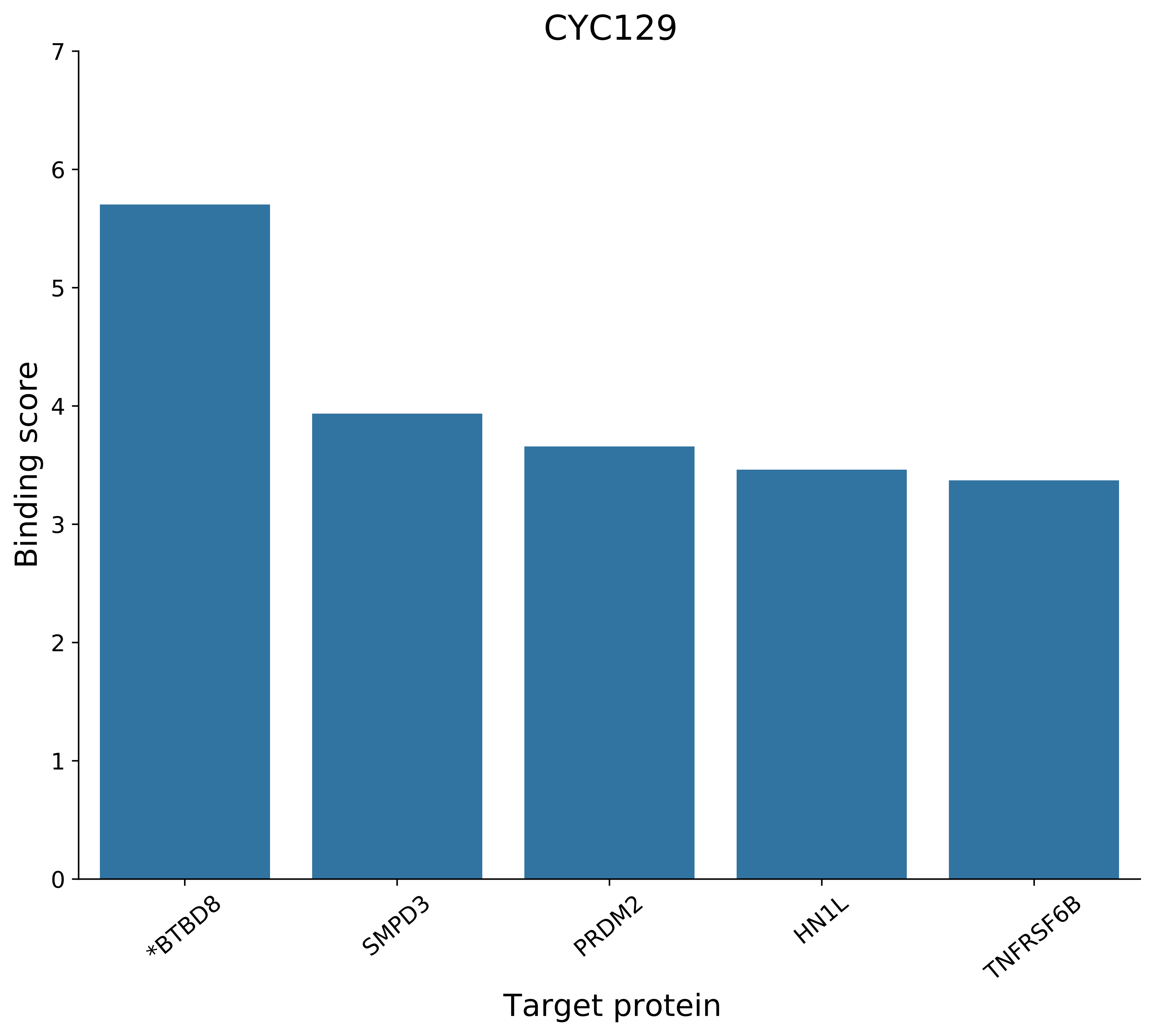
CYC129
Binding score
0
1
2
3
4
5
6
7
*BTBD8
SMPD3
PRDM2
HN1L
TNFRSF6B
Target protein

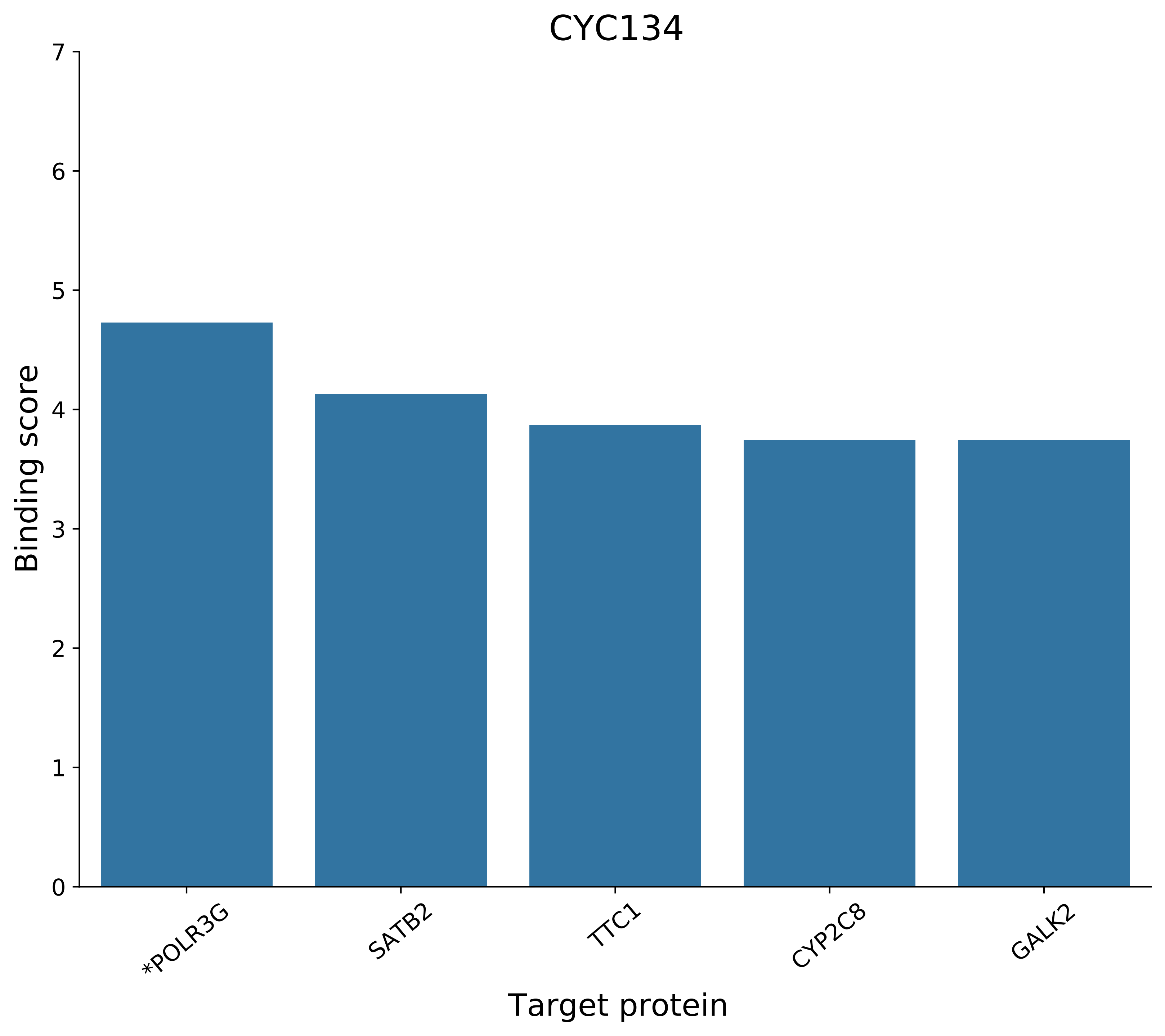

CYC134
Binding score
Target protein
0
1
2
3
4
5
6
7
*POLR3G
SATB2
TTC1
CYP2C8
GALK2

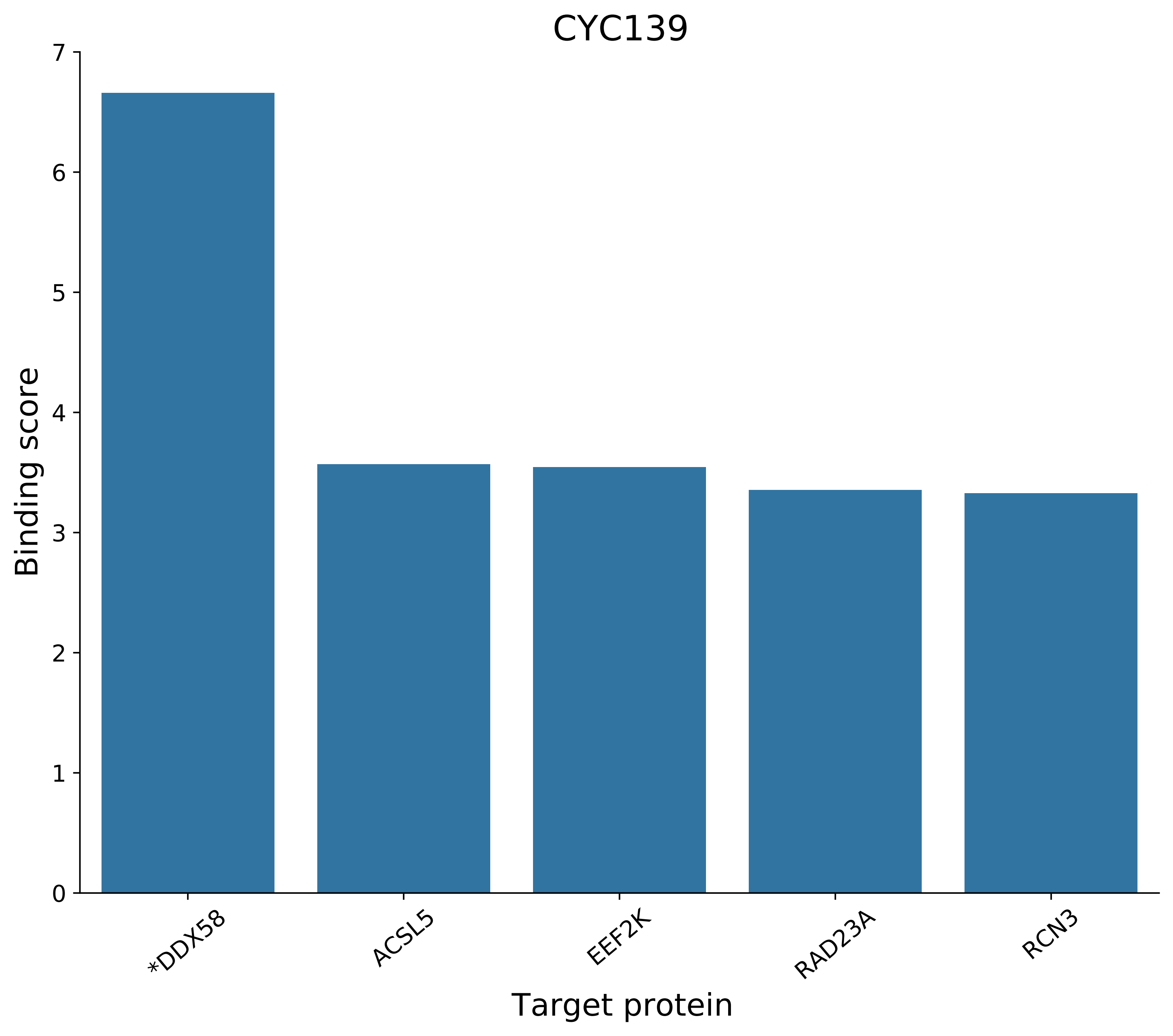
CYC139
Binding score
0
1
2
3
4
5
6
7
*DDX58
ACSL5
EEF2K
RAD23A
RCN3
Target protein

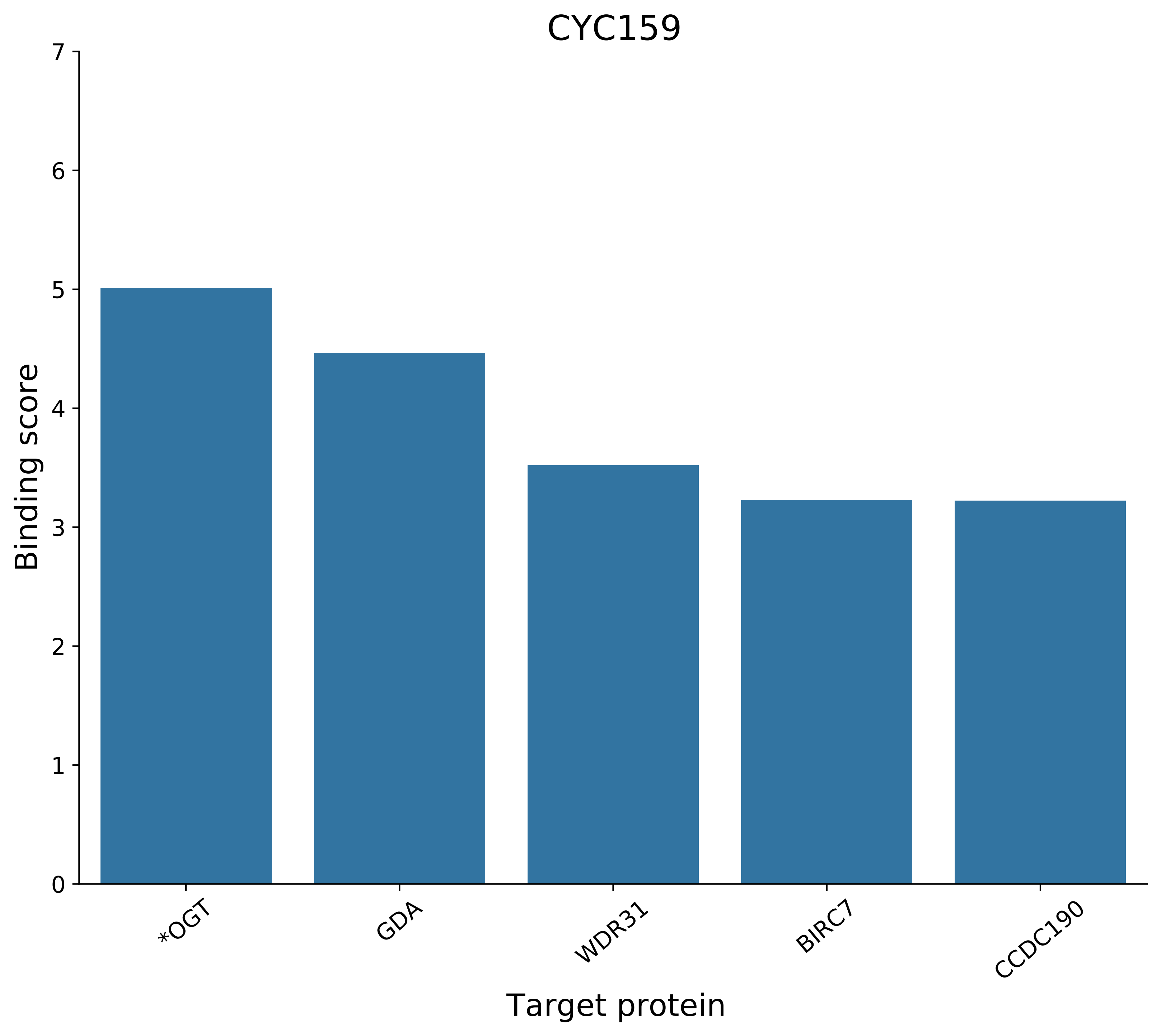

CYC159
Binding score
0
1
2
3
4
5
6
7
*OGT
GDA
WDR31
BIRC7
CCDC190
Target protein

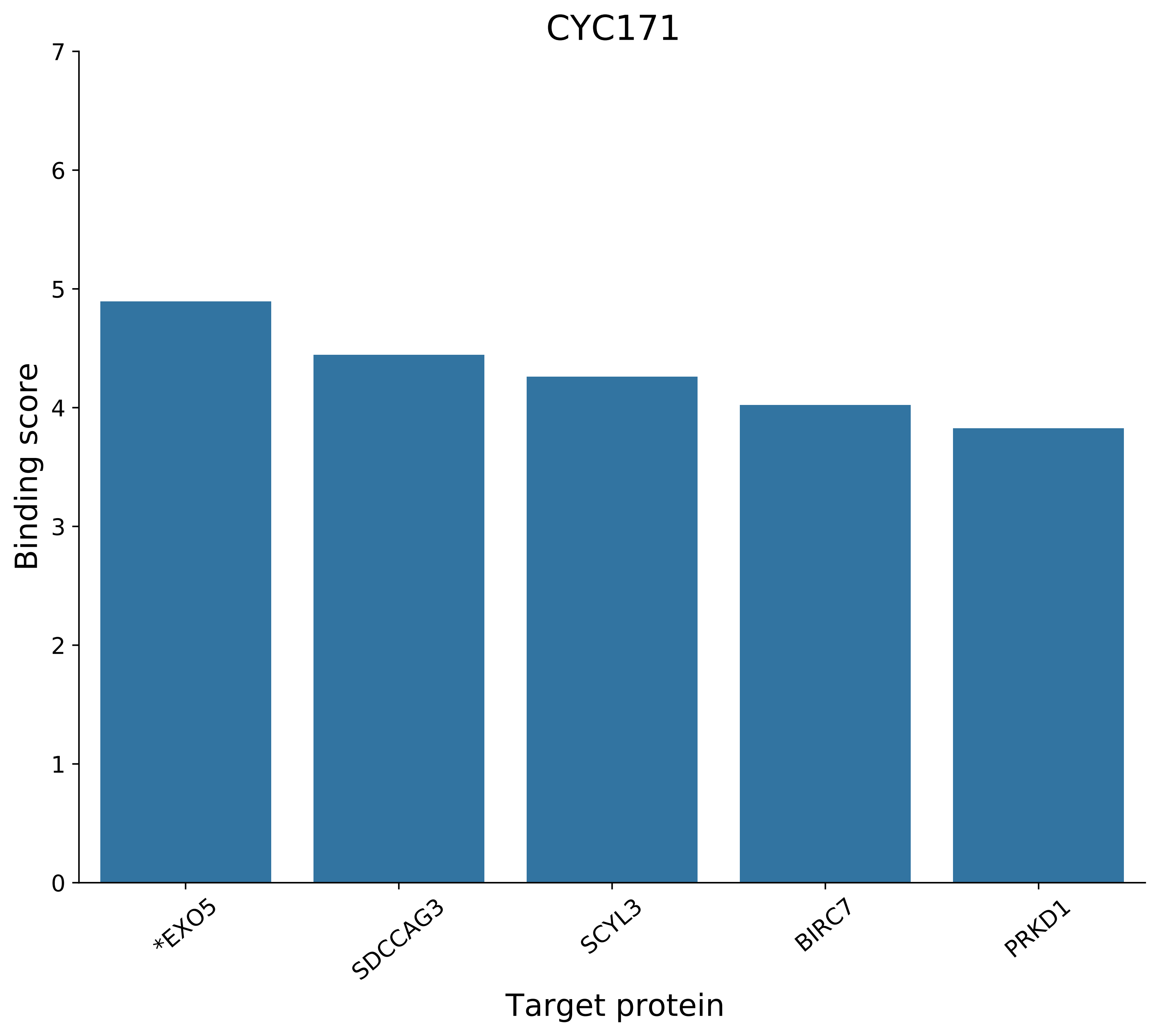

CYC171
Binding score
Target protein
0
1
2
3
4
5
6
7
*EXO5
SDCCAG3
SCYL3
BIRC7
PRKD1

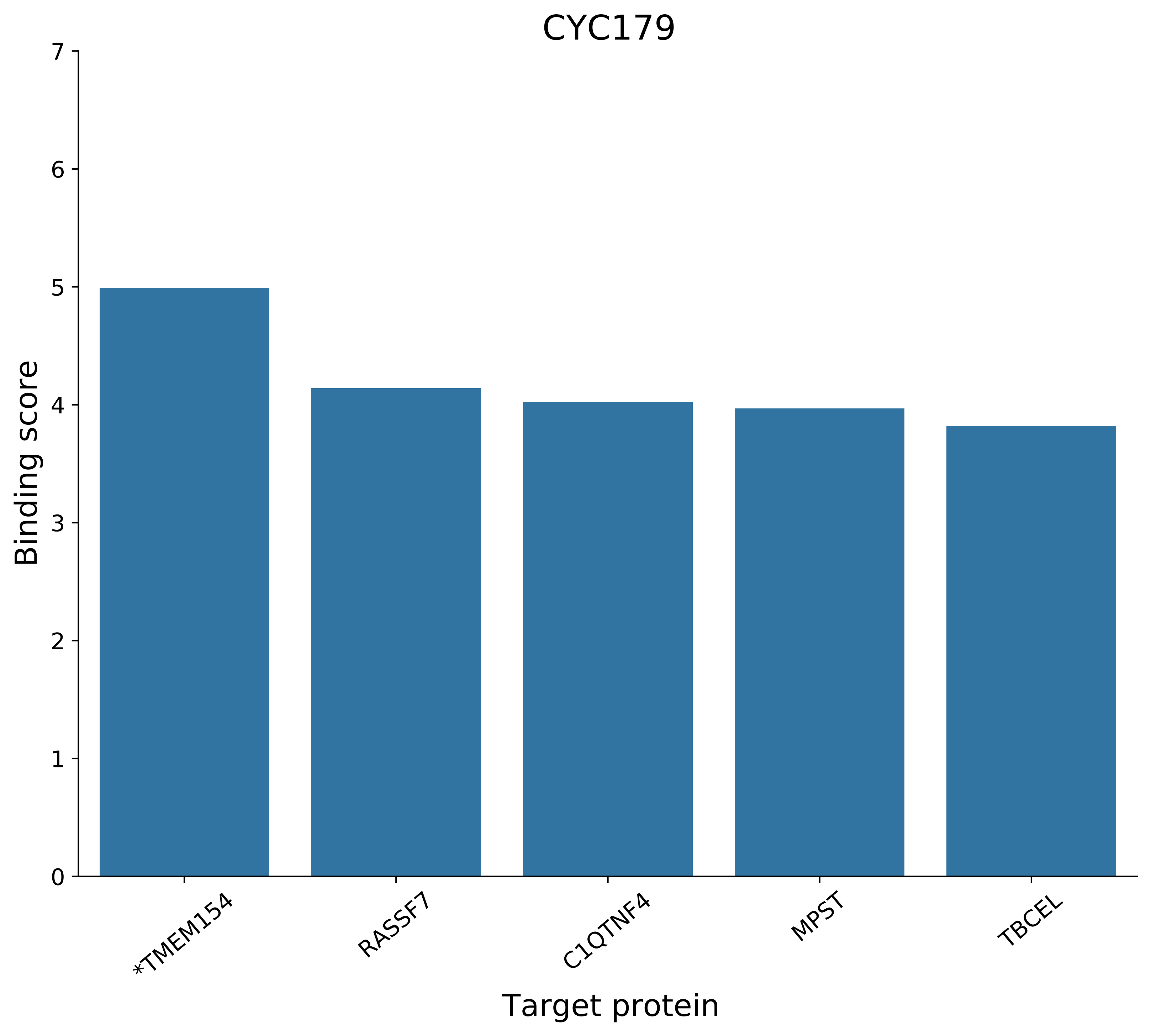
CYC179
Binding score
0
1
2
3
4
5
6
7
*TMEM154
RASSF7
C1QTNF4
MPST
TBCEL
Target protein

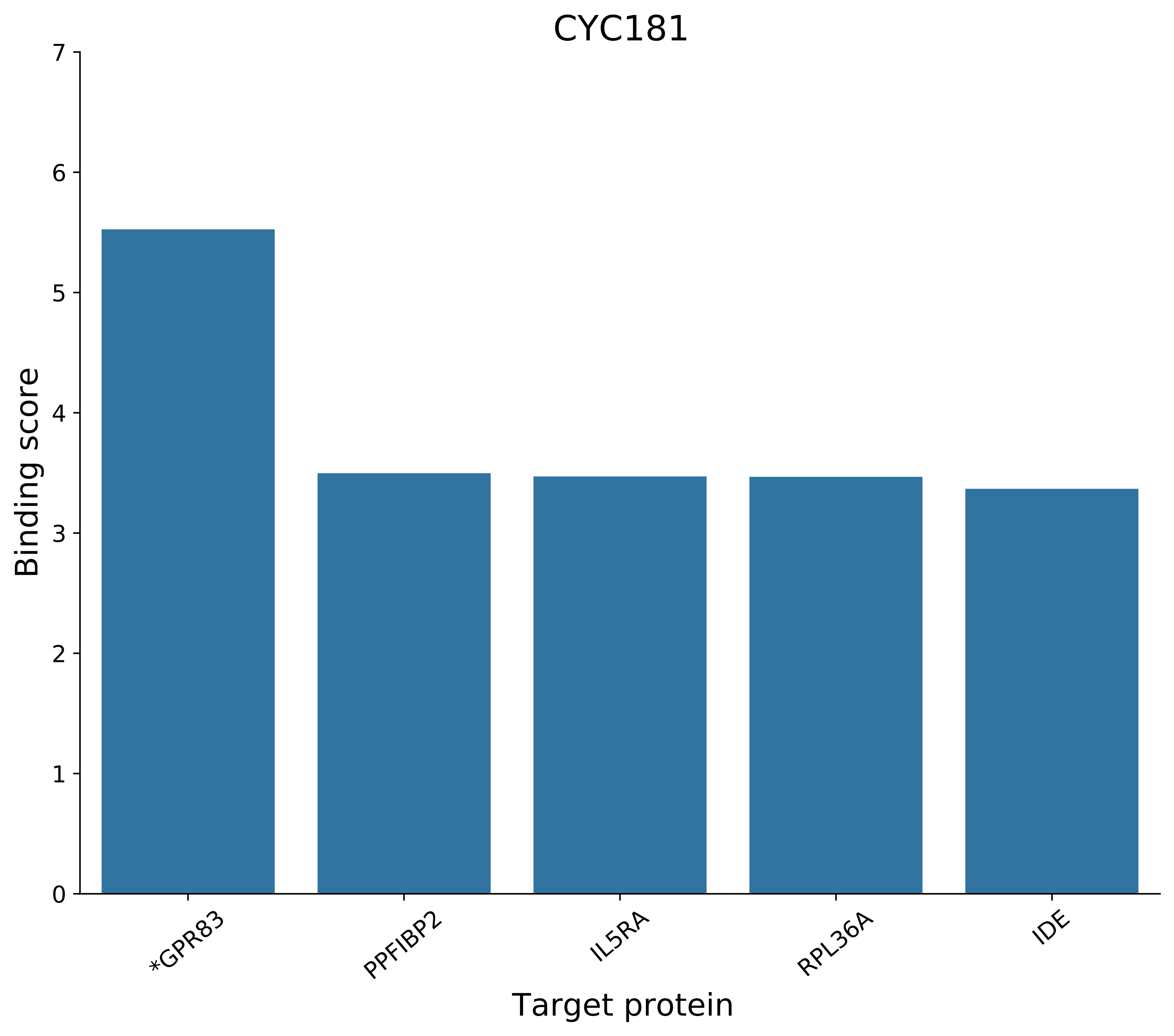
CYC181
Binding score
Target protein
0
1
2
3
4
5
6
7
*GPR83
PPFIBP2
IL5RA
RPL36A
IDE

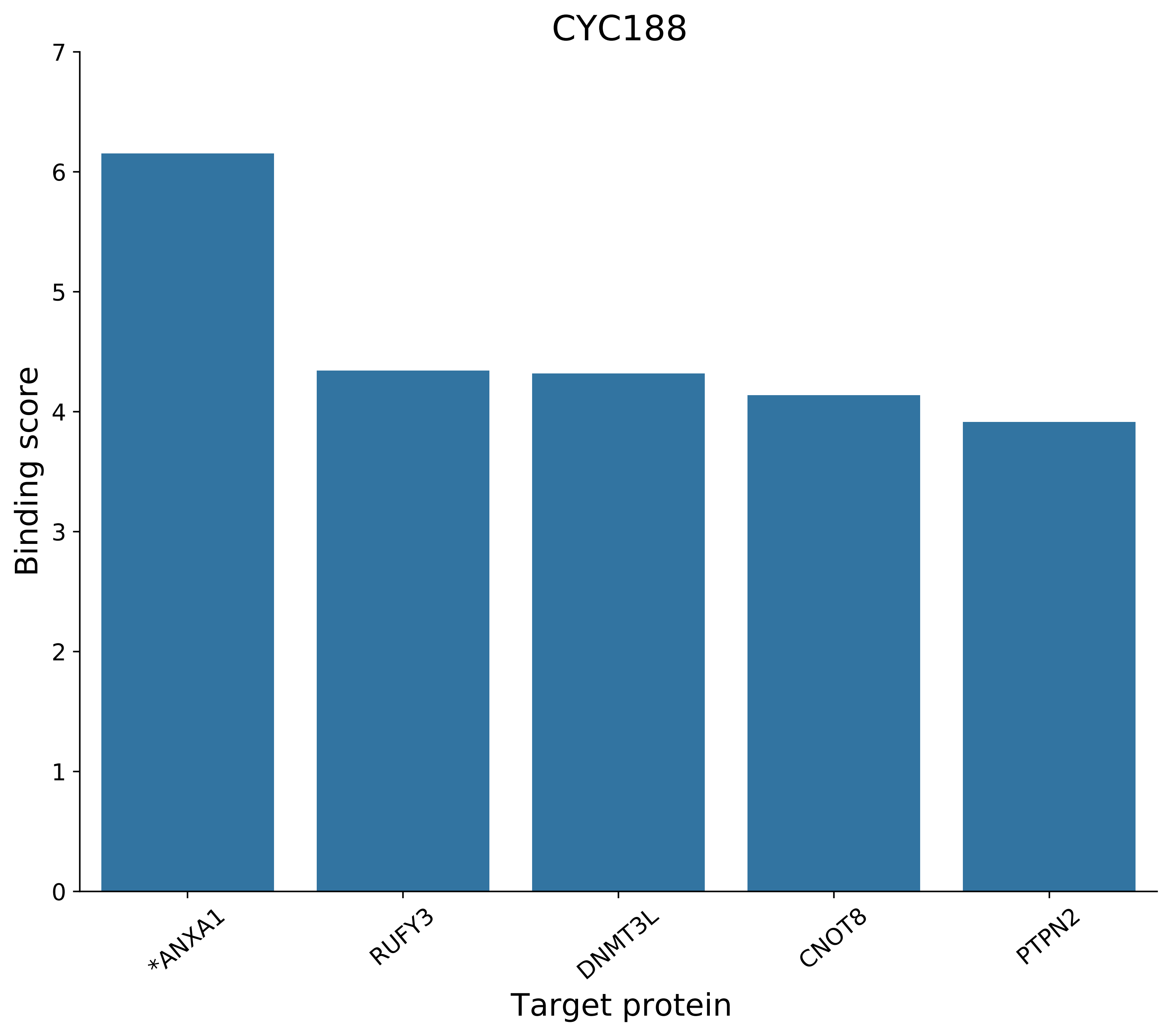

CYC188
Binding score
Target protein
0
1
2
3
4
5
6
7
*ANXA1
RUFY3
DNMT3L
CNOT8
PTPN2

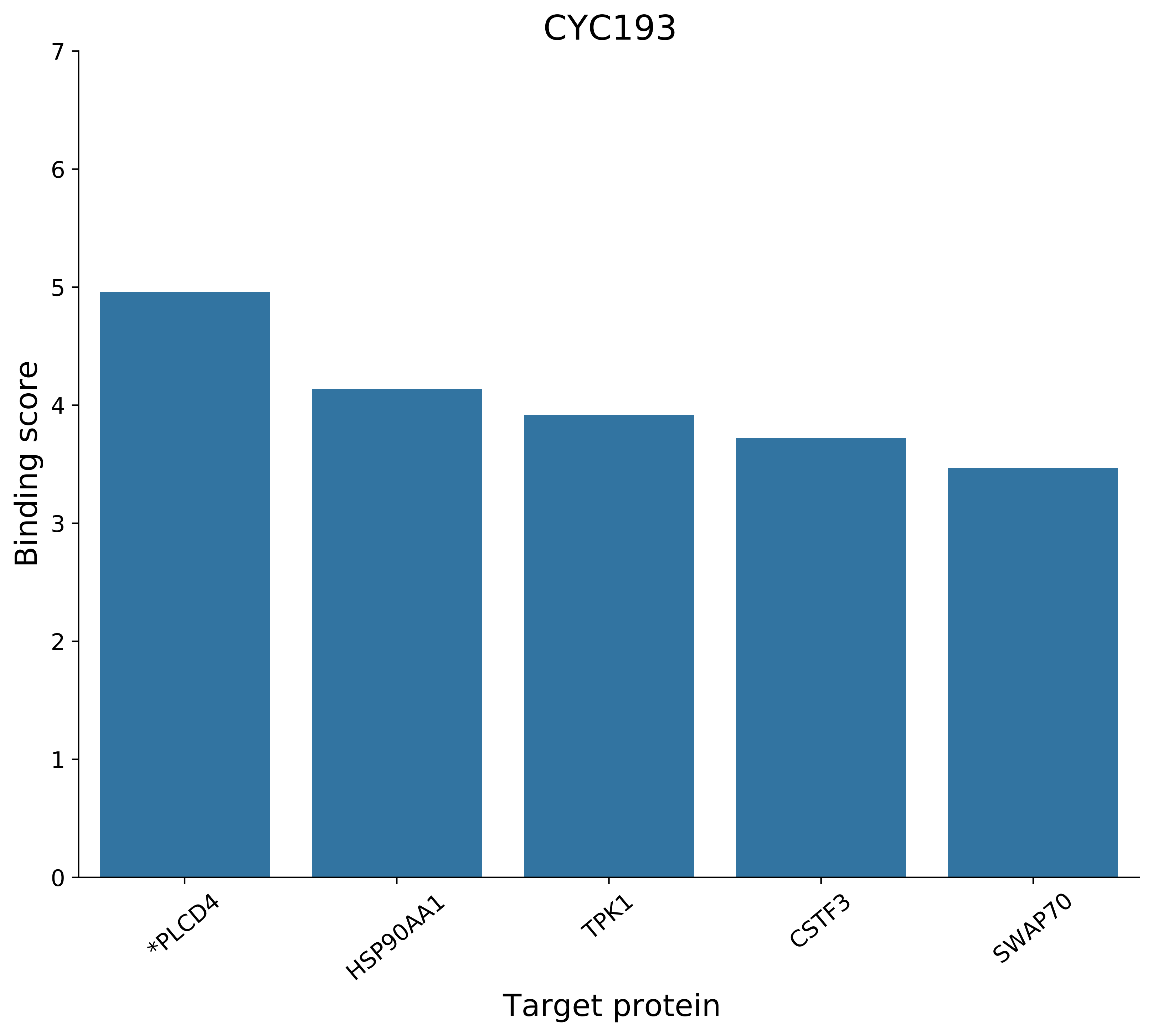
CYC193
Binding score
0
1
2
3
4
5
6
7
*PLCD4
HSP90AA1
TPK1
CSTF3
SWAP70
Target protein

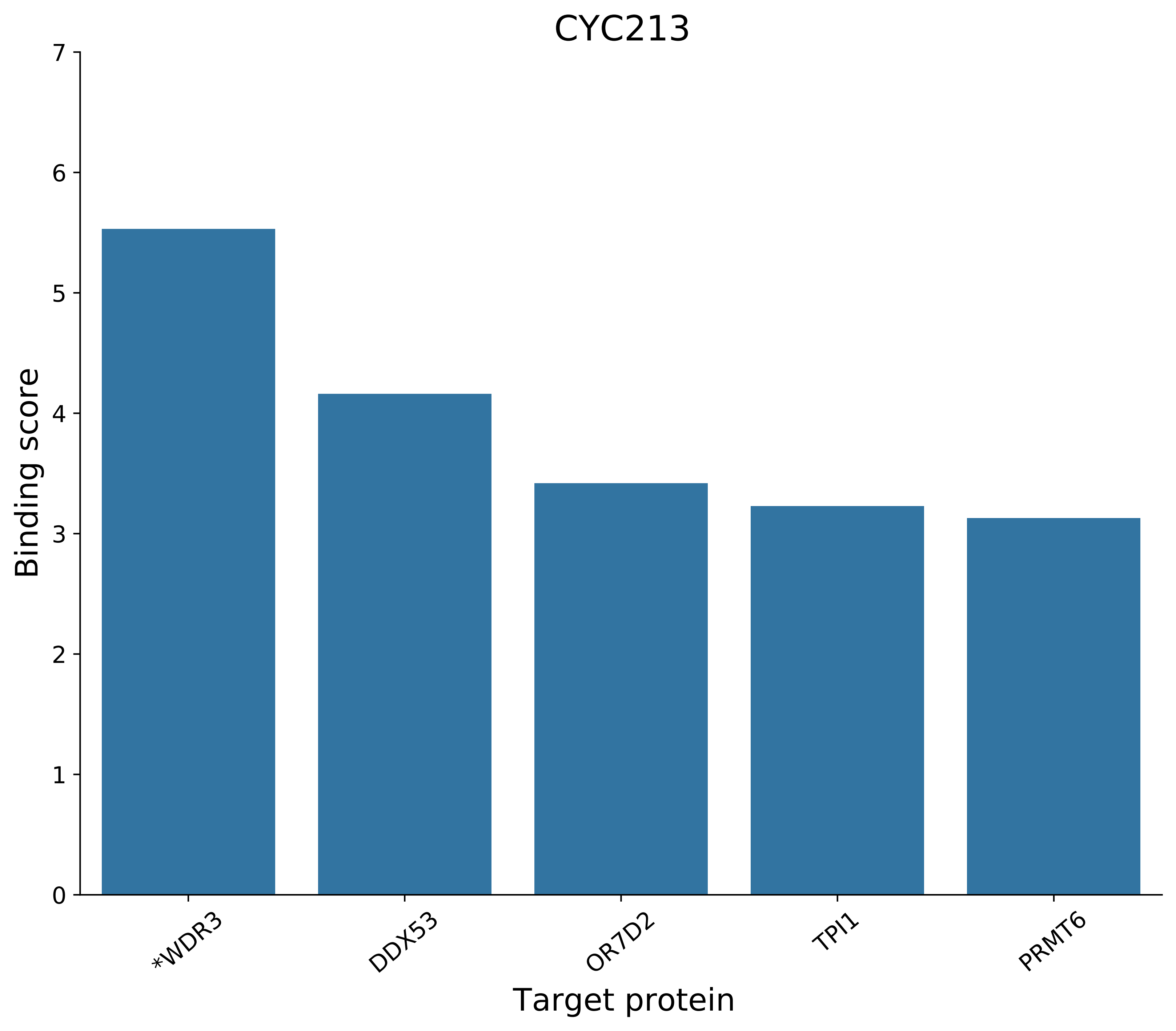

CYC213
Binding score
0
1
2
3
4
5
6
7
*WDR3
DDX53
OR7D2
TPI1
PRMT6
Target protein

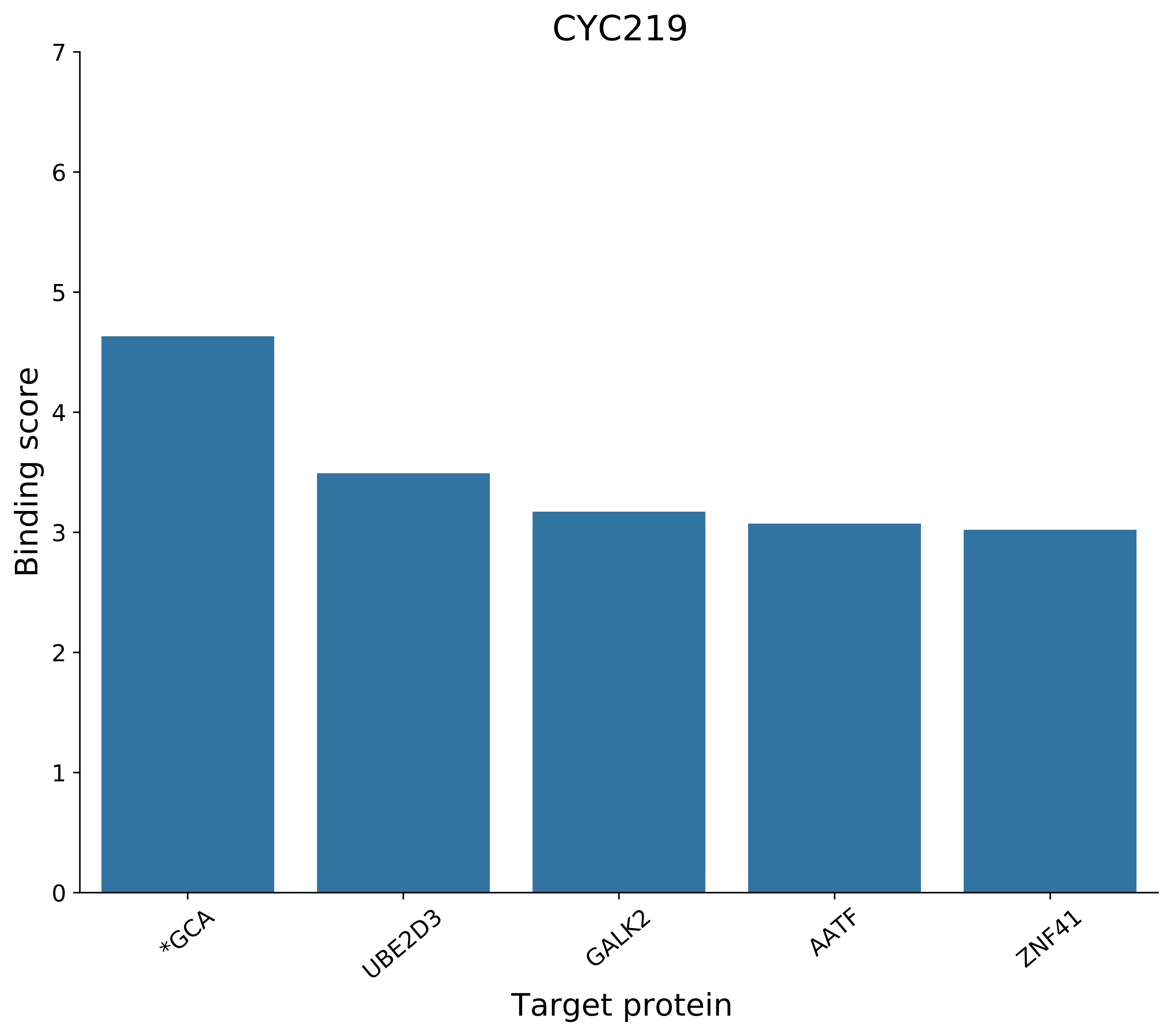
CYC219
Binding score
0
1
2
3
4
5
6
7
*GCA
UBE2D3
GALK2
AATF
ZNF41
Target protein

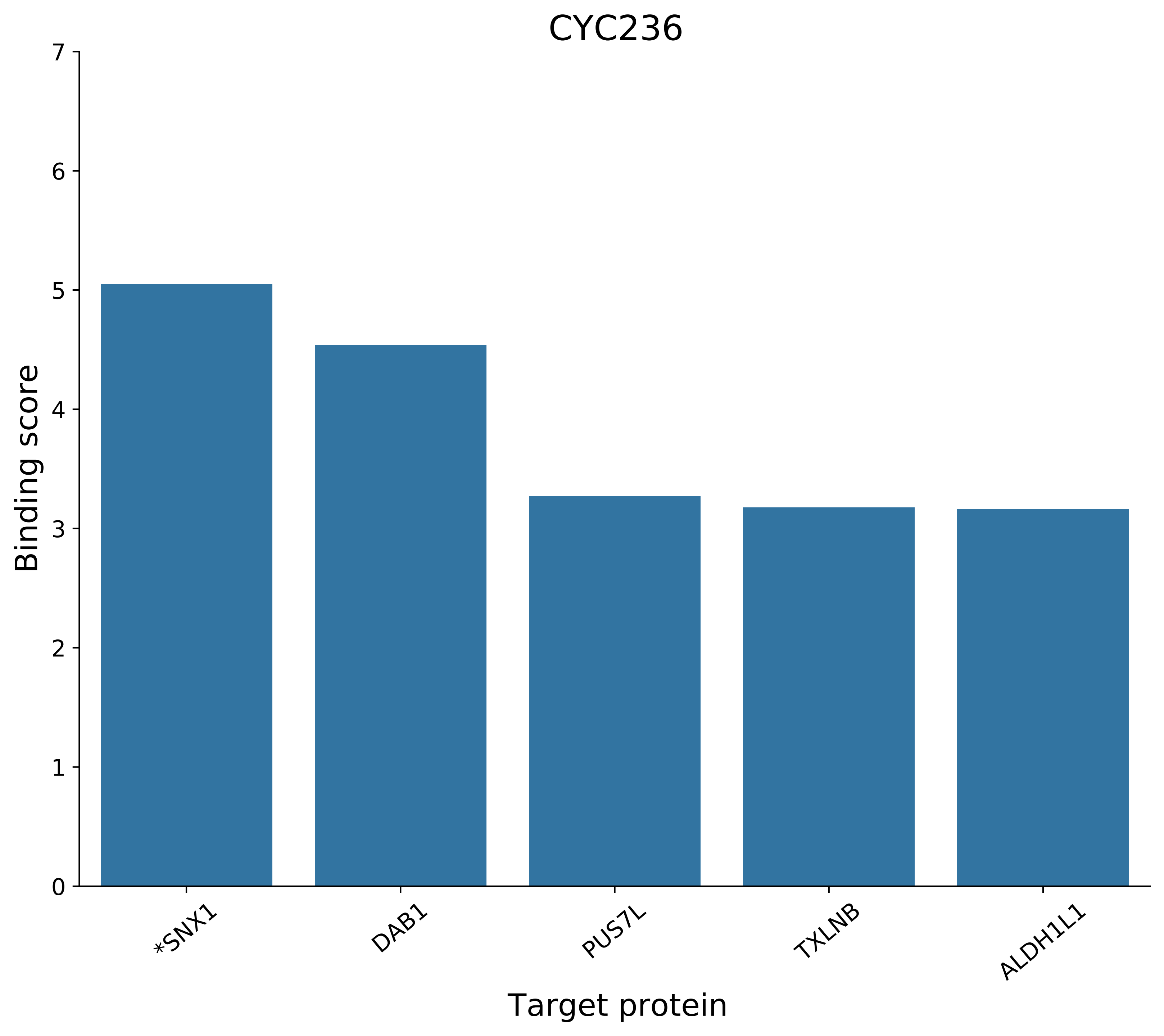
CYC236
Binding score
Target protein
0
1
2
3
4
5
6
7
*SNX1
DAB1
PUS7L
TXLNB
ALDH1L1

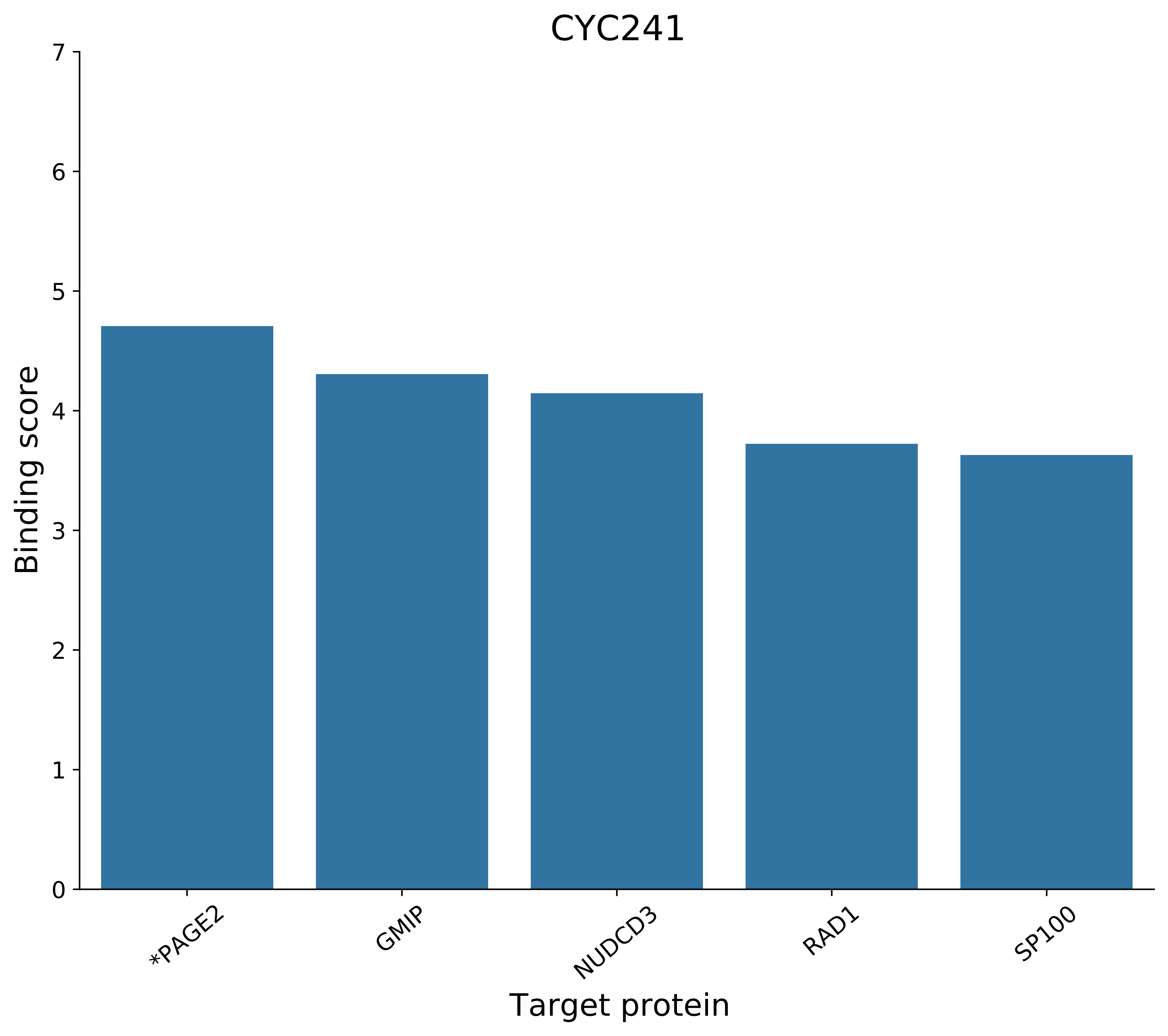

CYC241
Binding score
0
1
2
3
4
5
6
7
*PAGE2
GMIP
NUDCD3
RAD1
SP100
Target protein

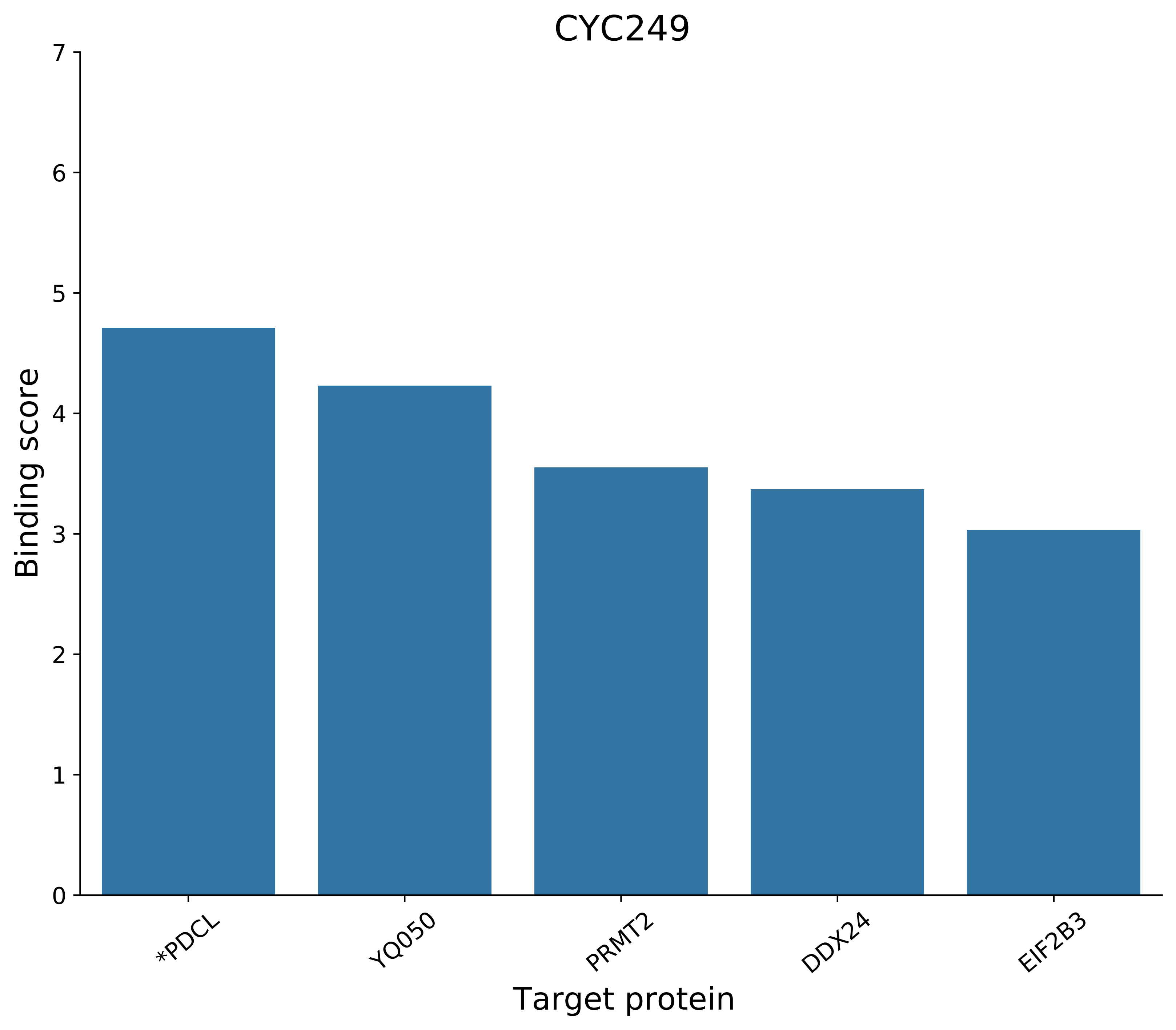
CYC249
Binding score
0
1
2
3
4
5
6
7
*PDCL
YQ050
PRMT2
DDX24
EIF2B3
Target protein

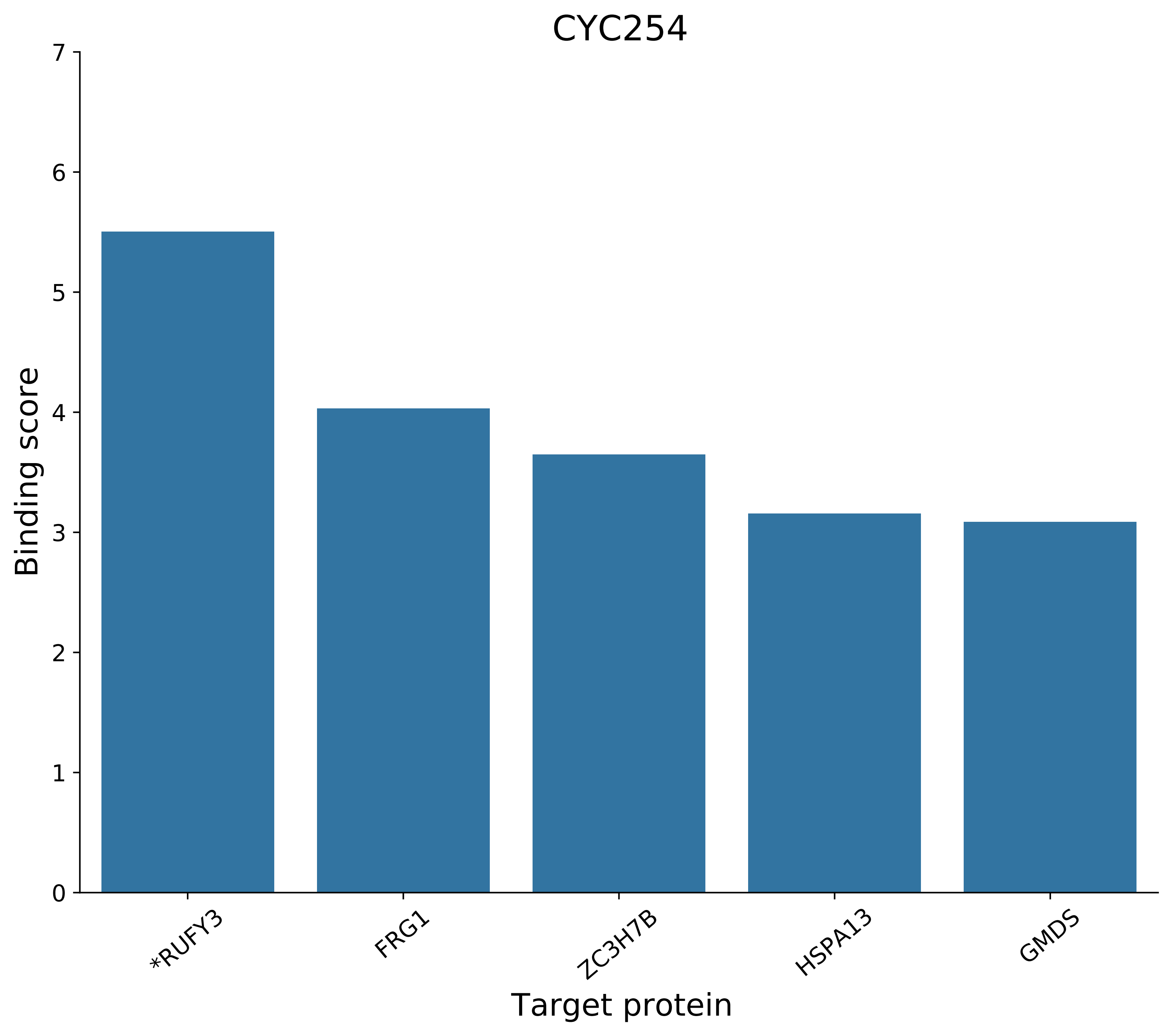
CYC254
Binding score
0
1
2
3
4
5
6
7
*RUFY3
FRG1
ZC3H7B
HSPA13
GMDS
Target protein

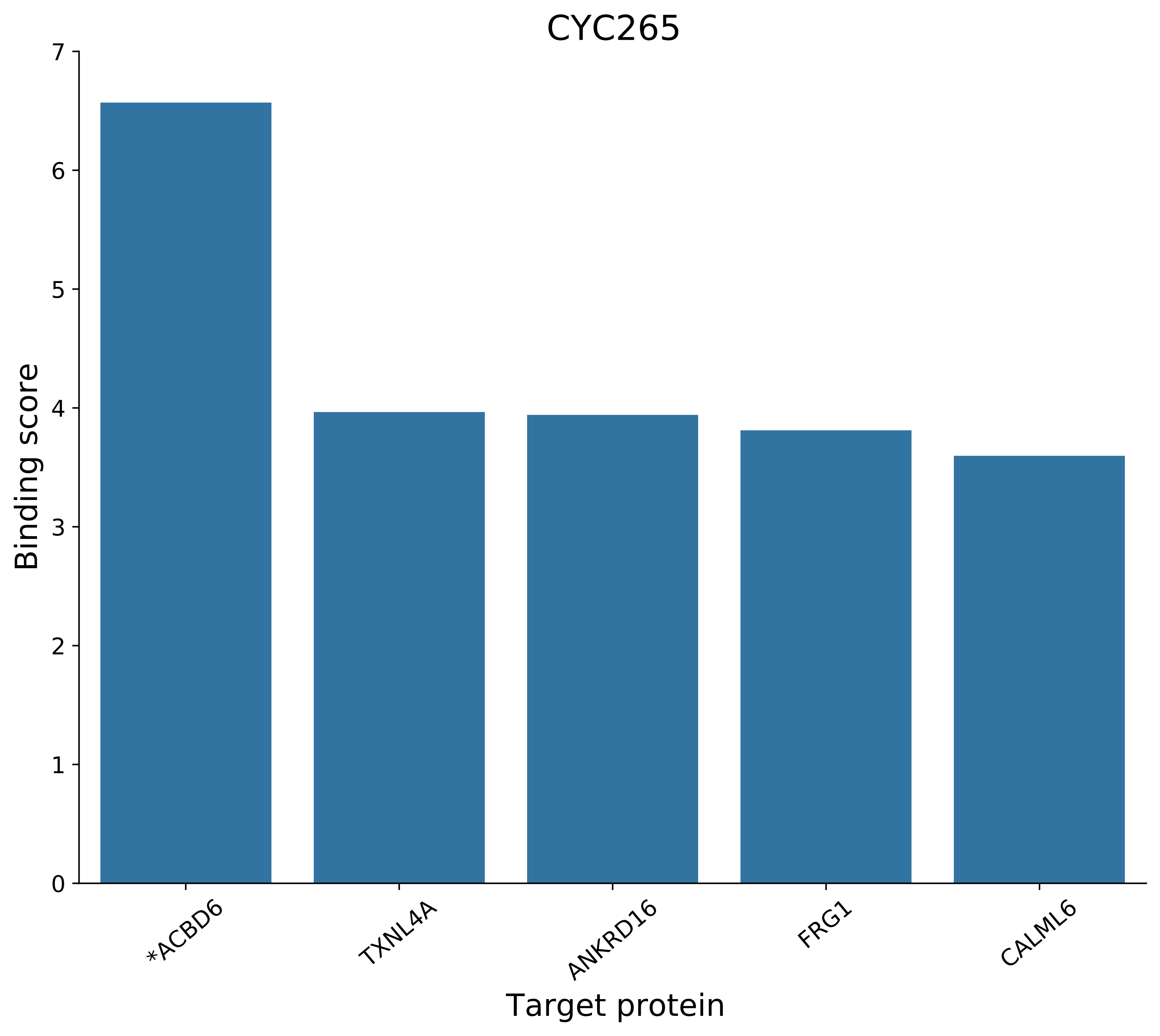
CYC265
Binding score
Target protein
0
1
2
3
4
5
6
7
*ACBD6
TXNL4A
ANKRD16
FRG1
CALML6

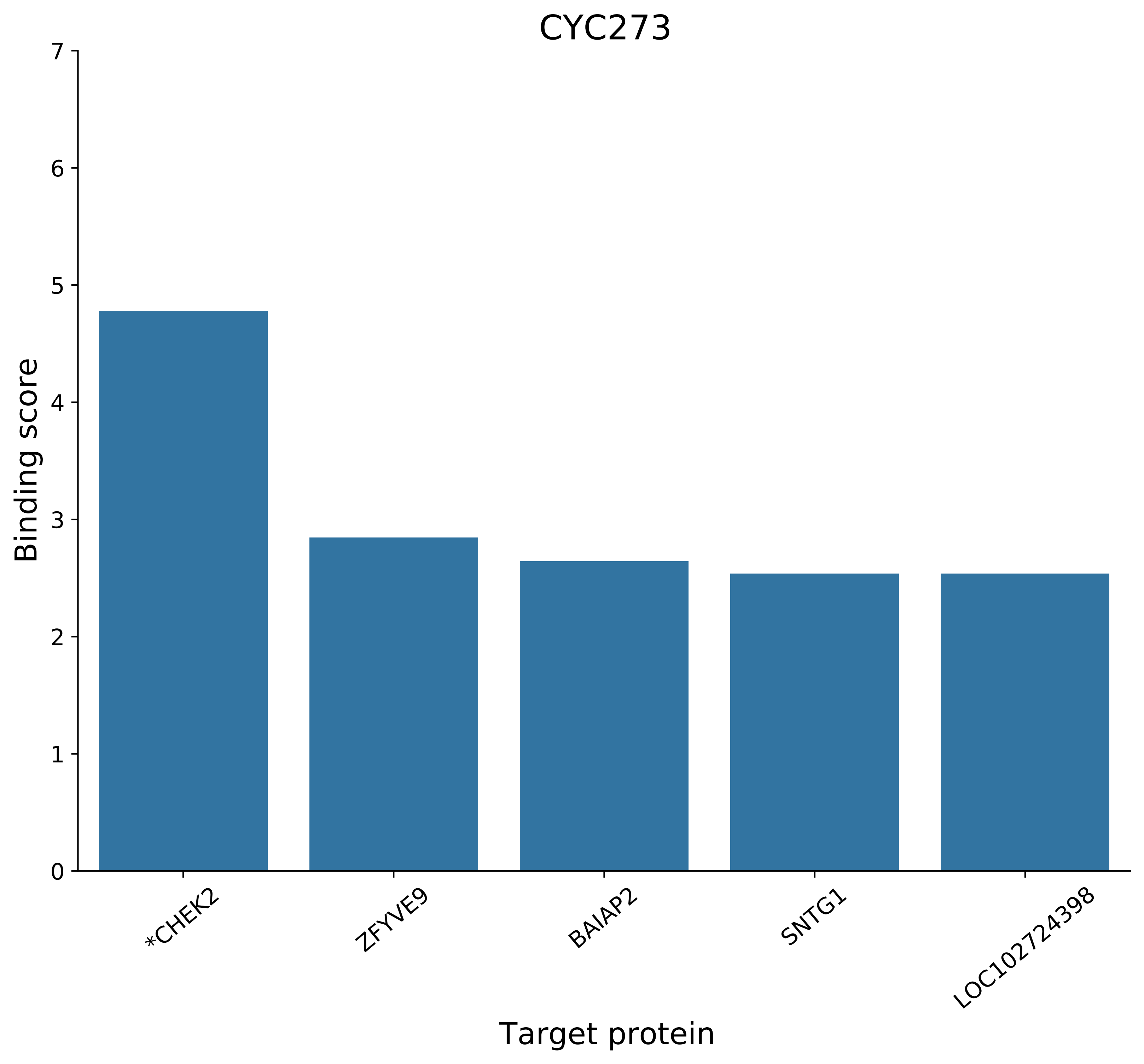

CYC273
Binding score
Target protein
0
1
2
3
4
5
6
7
*CHEK2
ZFYVE9
BAIAP2
SNTG1
LOC102724398

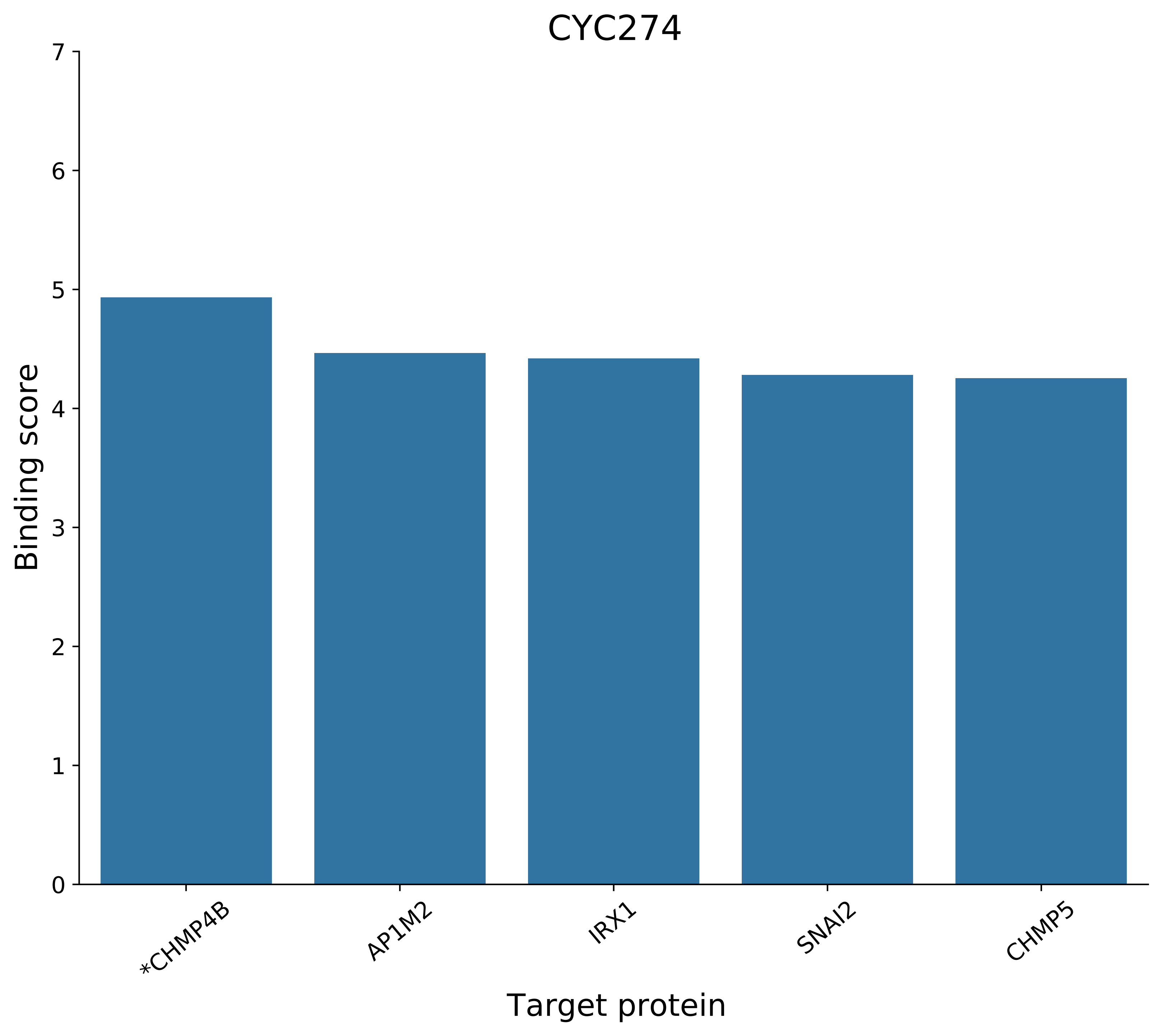
CYC274
Binding score
7
6
5
4
3
2
1
0
*CHMP4B
AP1M2
IRX1
SNAI2
CHMP5
Target protein

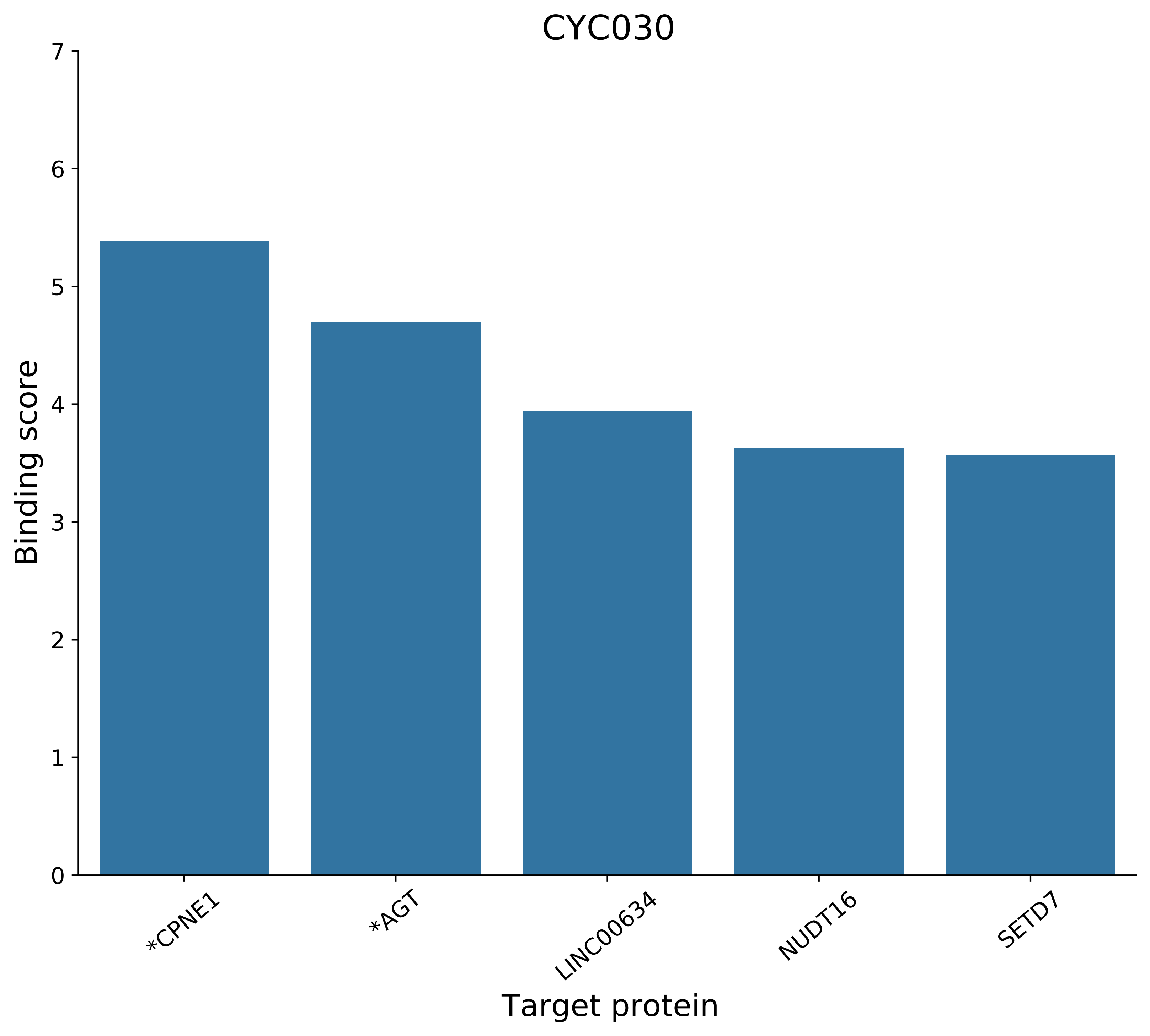

CYC030
Binding score
Target protein
0
1
2
3
4
5
6
7
*CPNE1
*AGT
LINC00634
NUDT16
SETD7

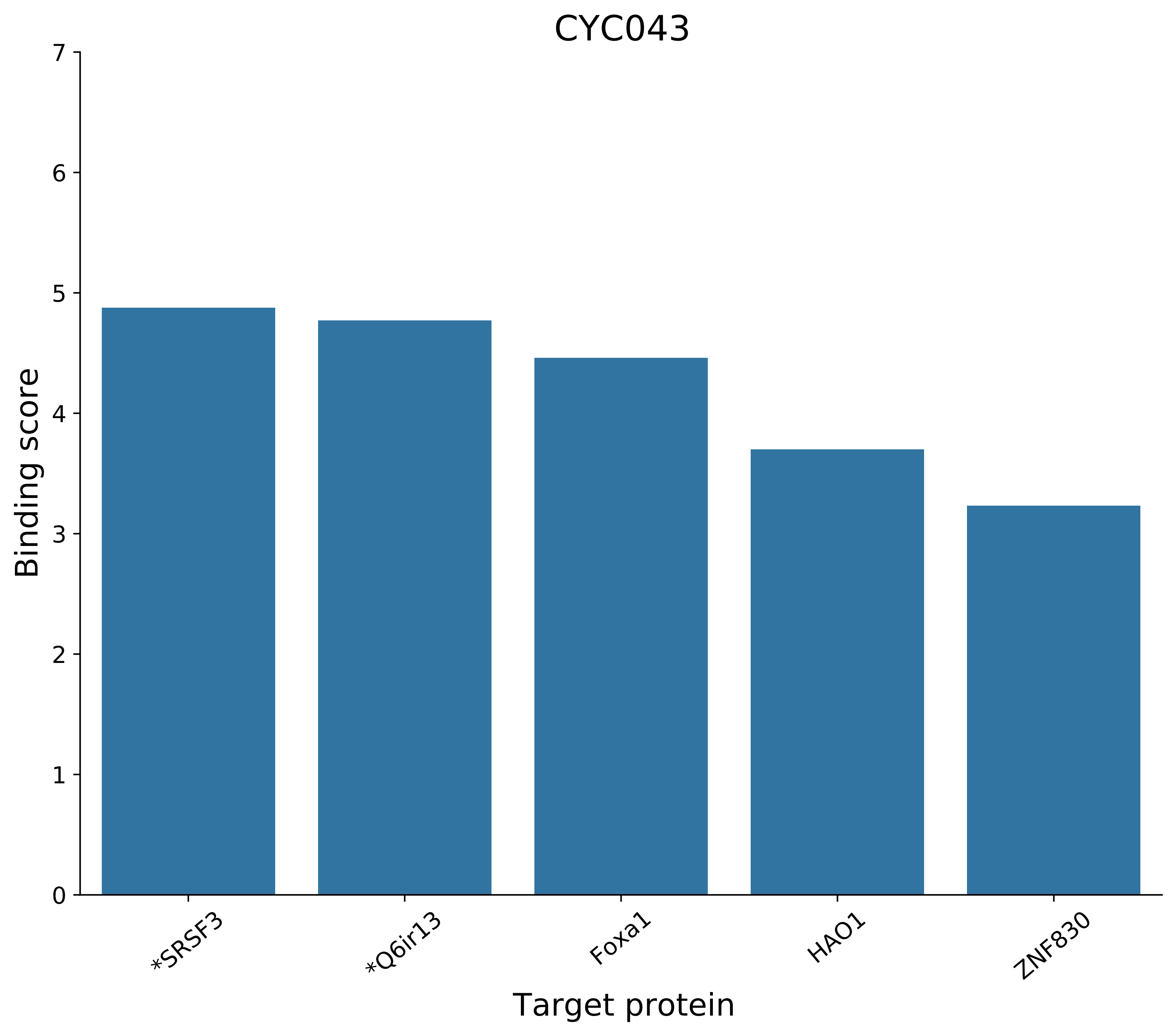
CYC043
Binding score
0
1
2
3
4
5
6
7
*SRSF3
*Q6ir13
Foxa1
HAO1
ZNF830
Target protein

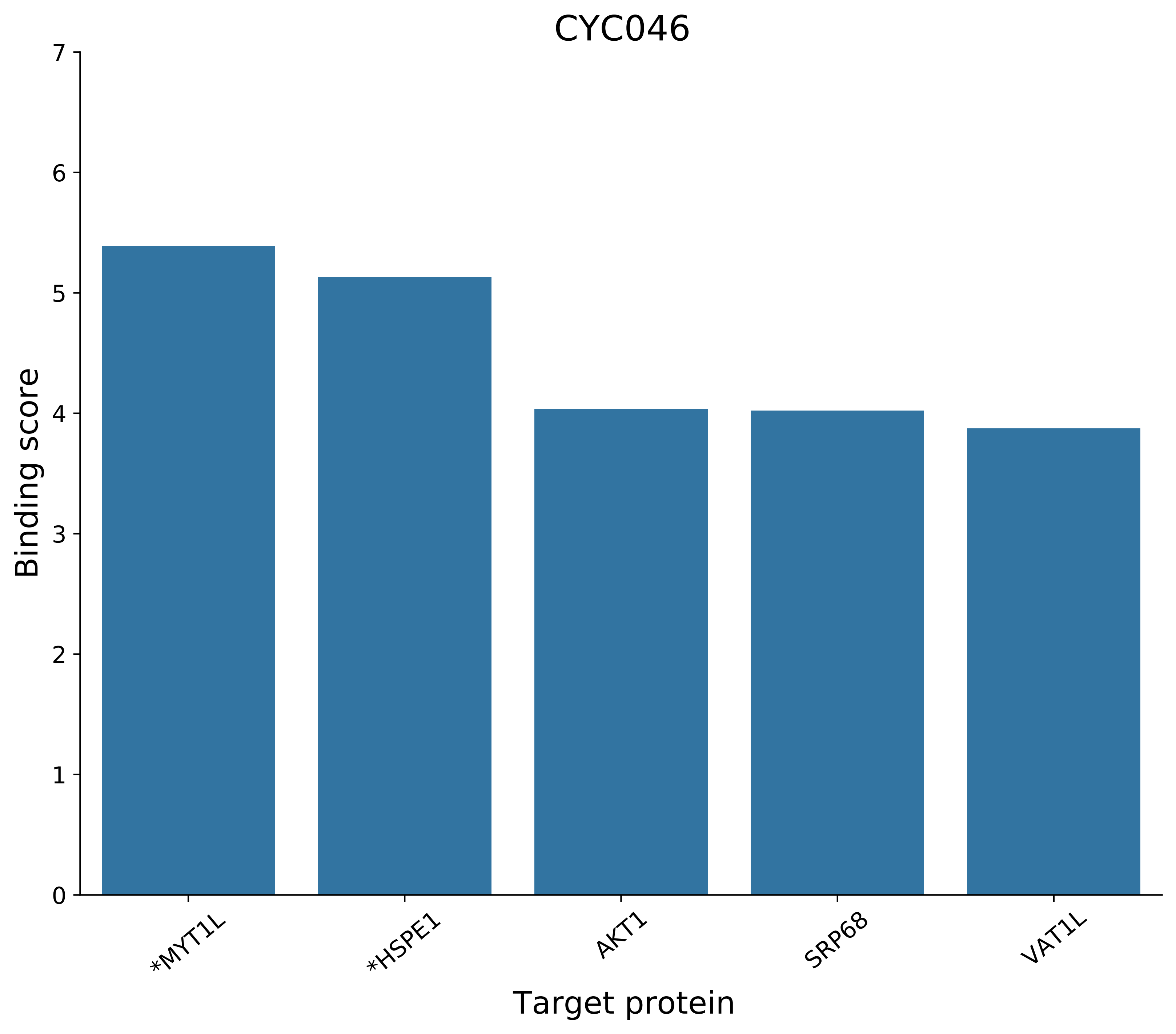
CYC046
Binding score
7
6
5
4
3
2
1
0
*MYT1L
*HSPE1
AKT1
SRP68
VAT1L
Target protein

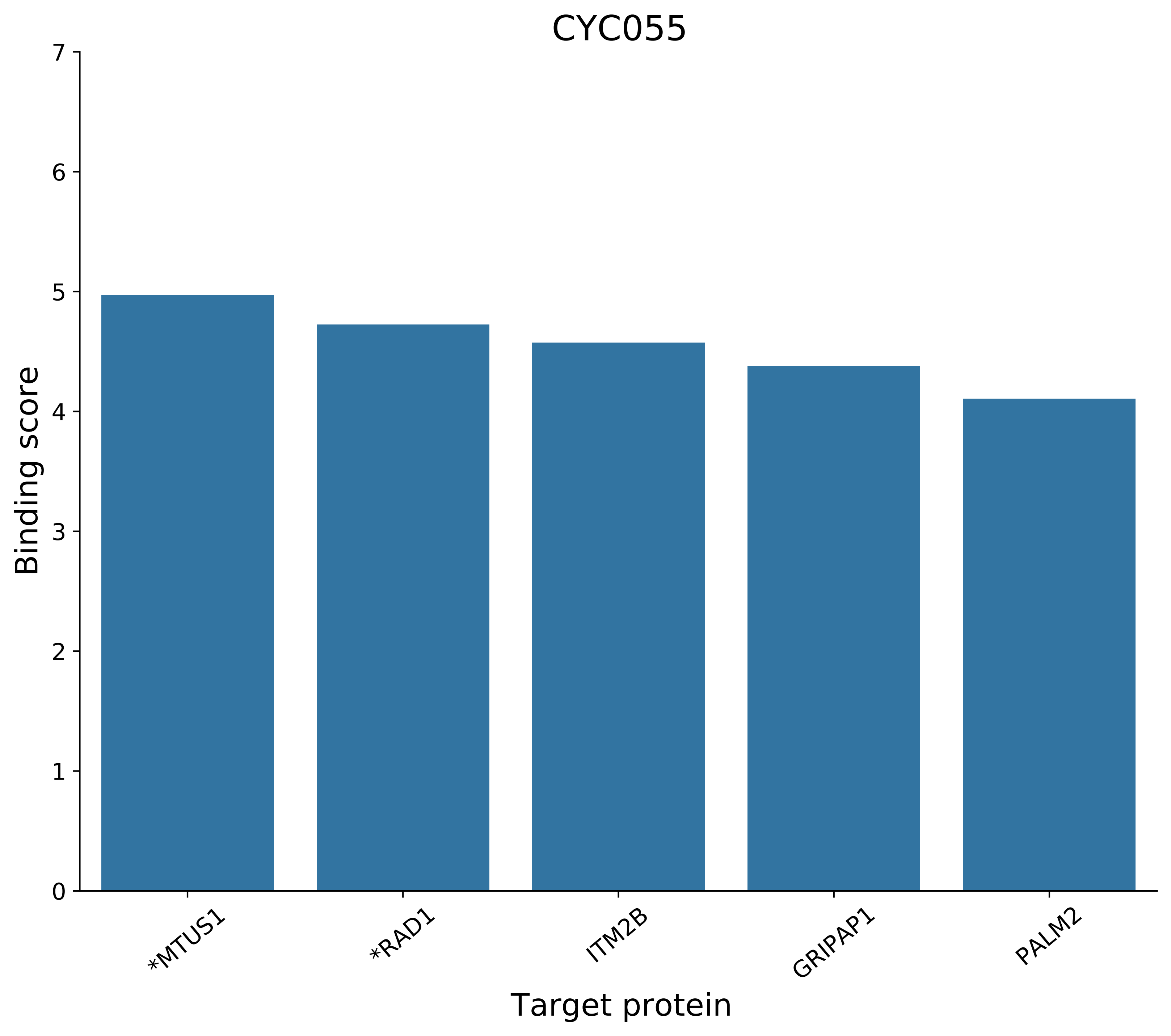
CYC055
Binding score
Target protein
0
1
2
3
4
5
6
7
*MTUS1
*RAD1
ITM2B
GRIPAP1
PALM2

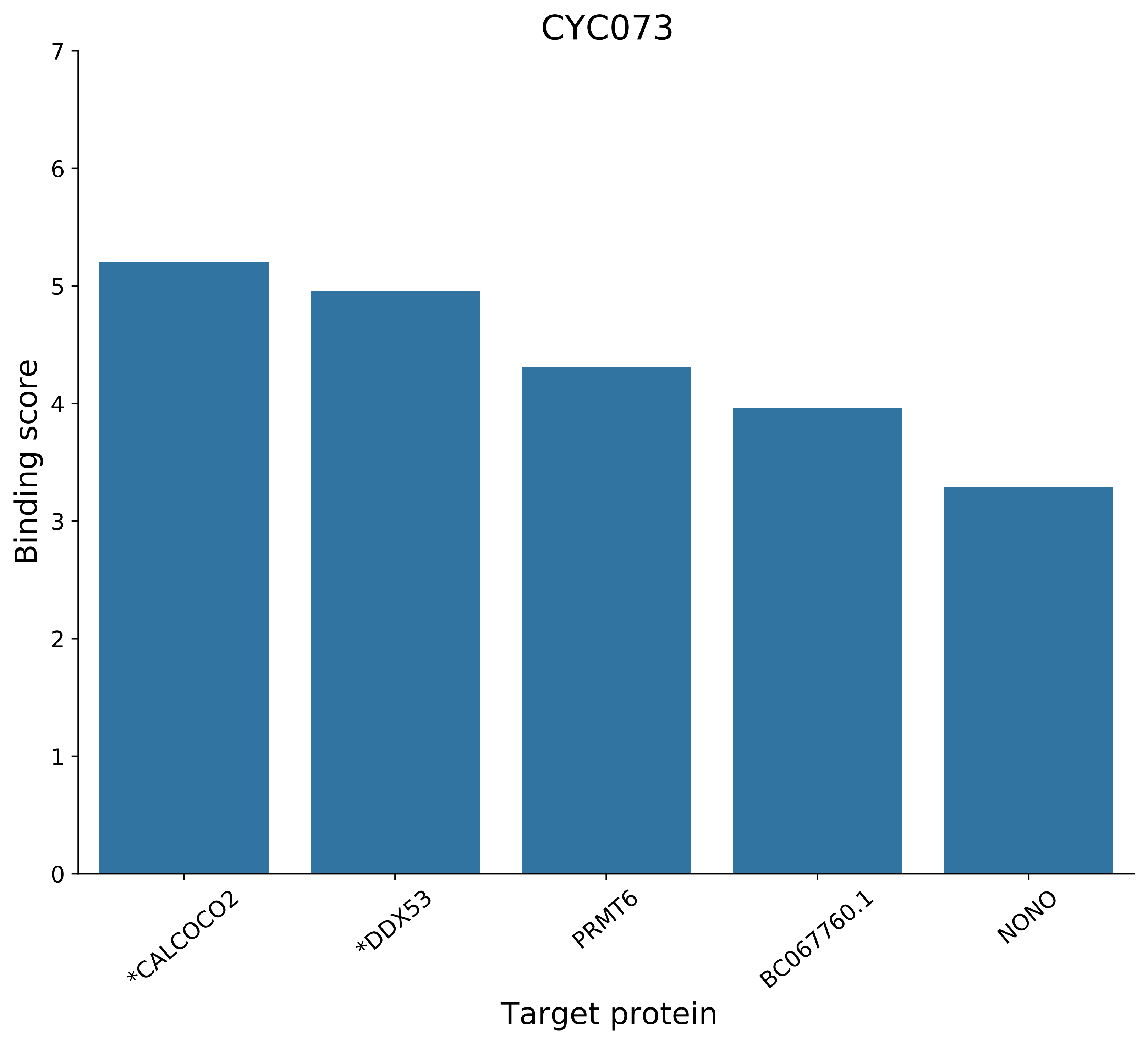
CYC073
Binding score
0
1
2
3
4
5
6
7
*CALCOCO2
*DDX53
PRMT6
BC067760.1
NONO
Target protein

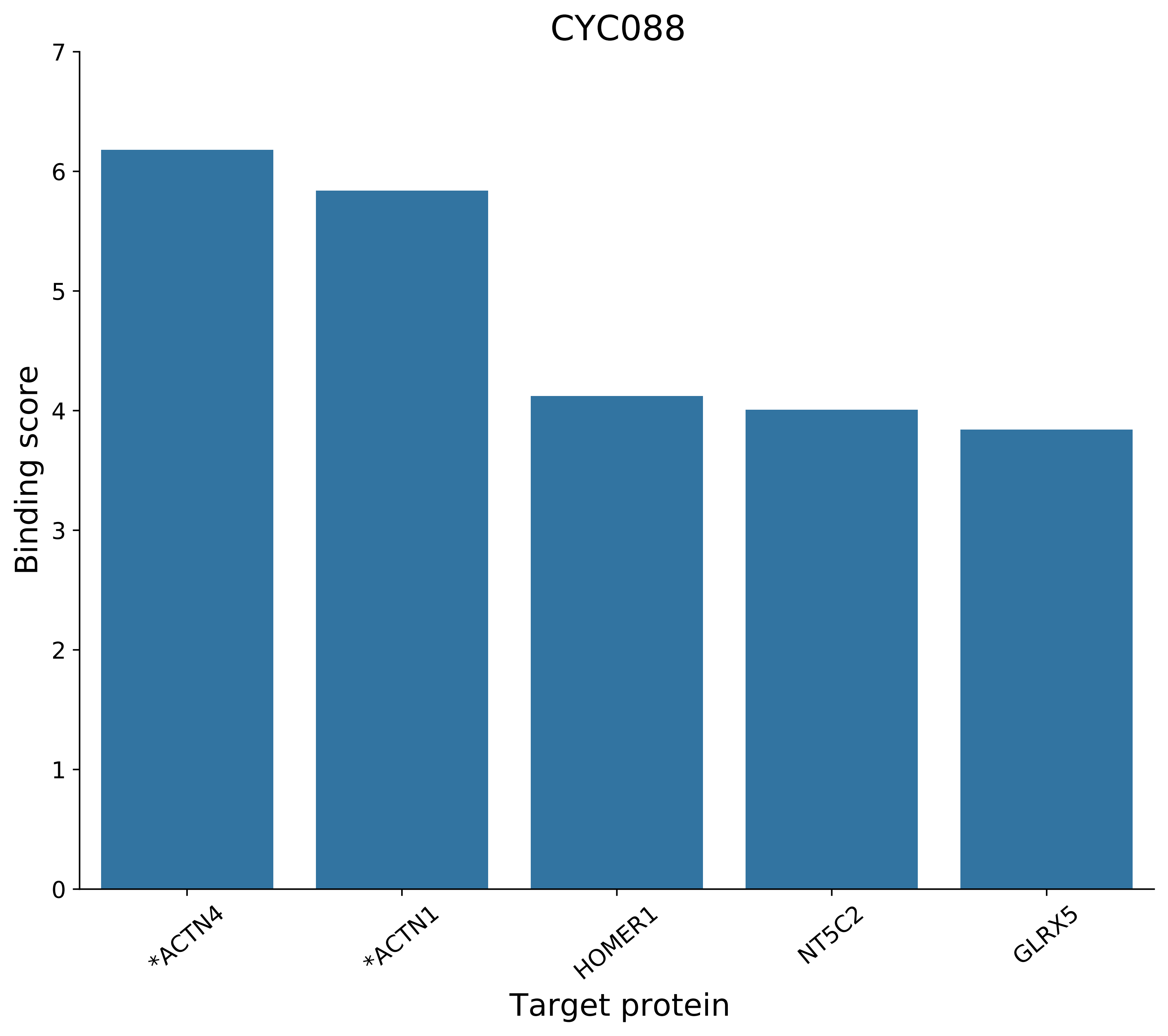

CYC088
Binding score
Target protein
0
1
2
3
4
5
6
7
*ACTN4
*ACTN1
HOMER1
NT5C2
GLRX5

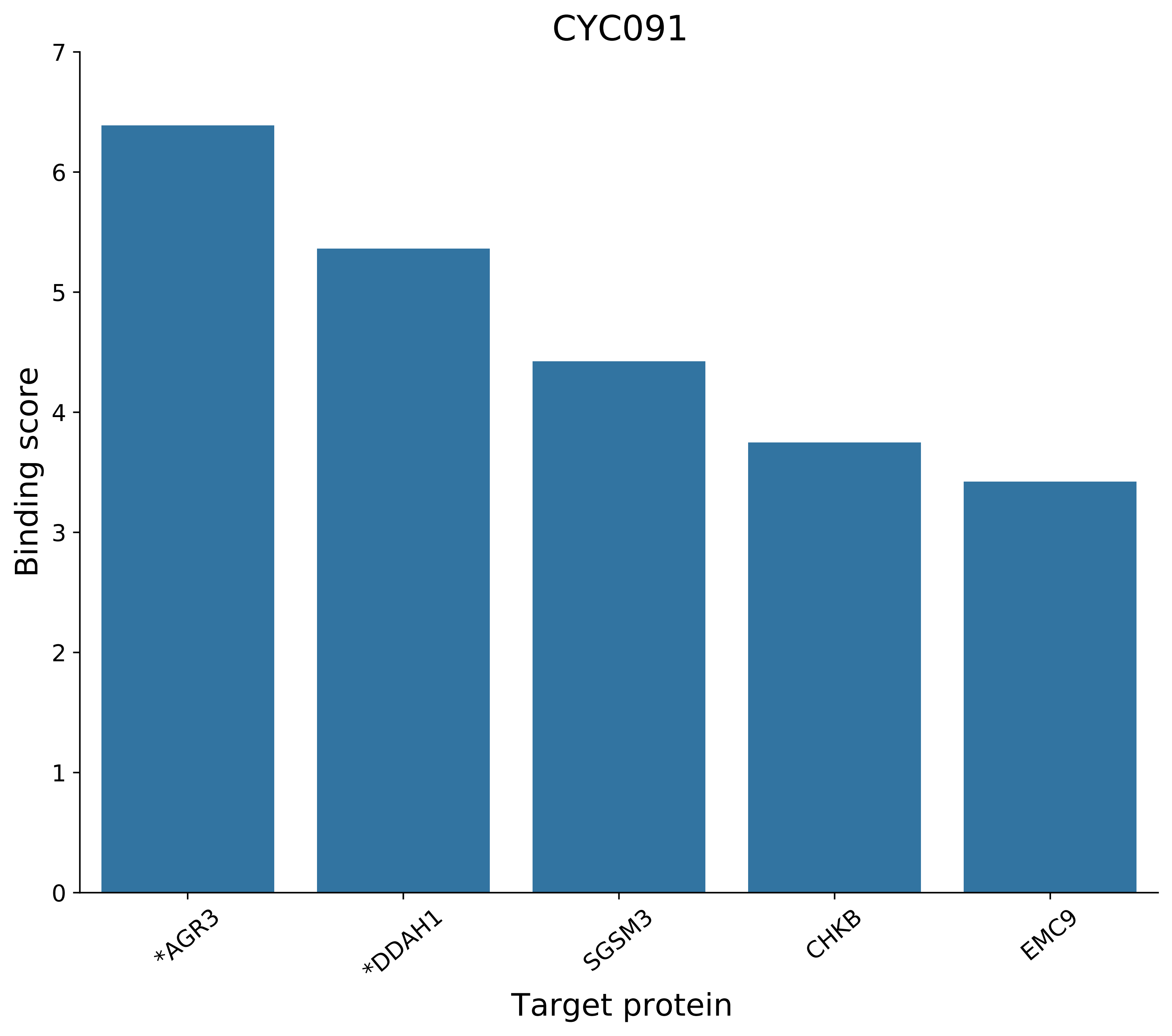
CYC091
Binding score
0
1
2
3
4
5
6
7
*AGR3
*DDAH1
SGSM3
CHKB
EMC9
Target protein

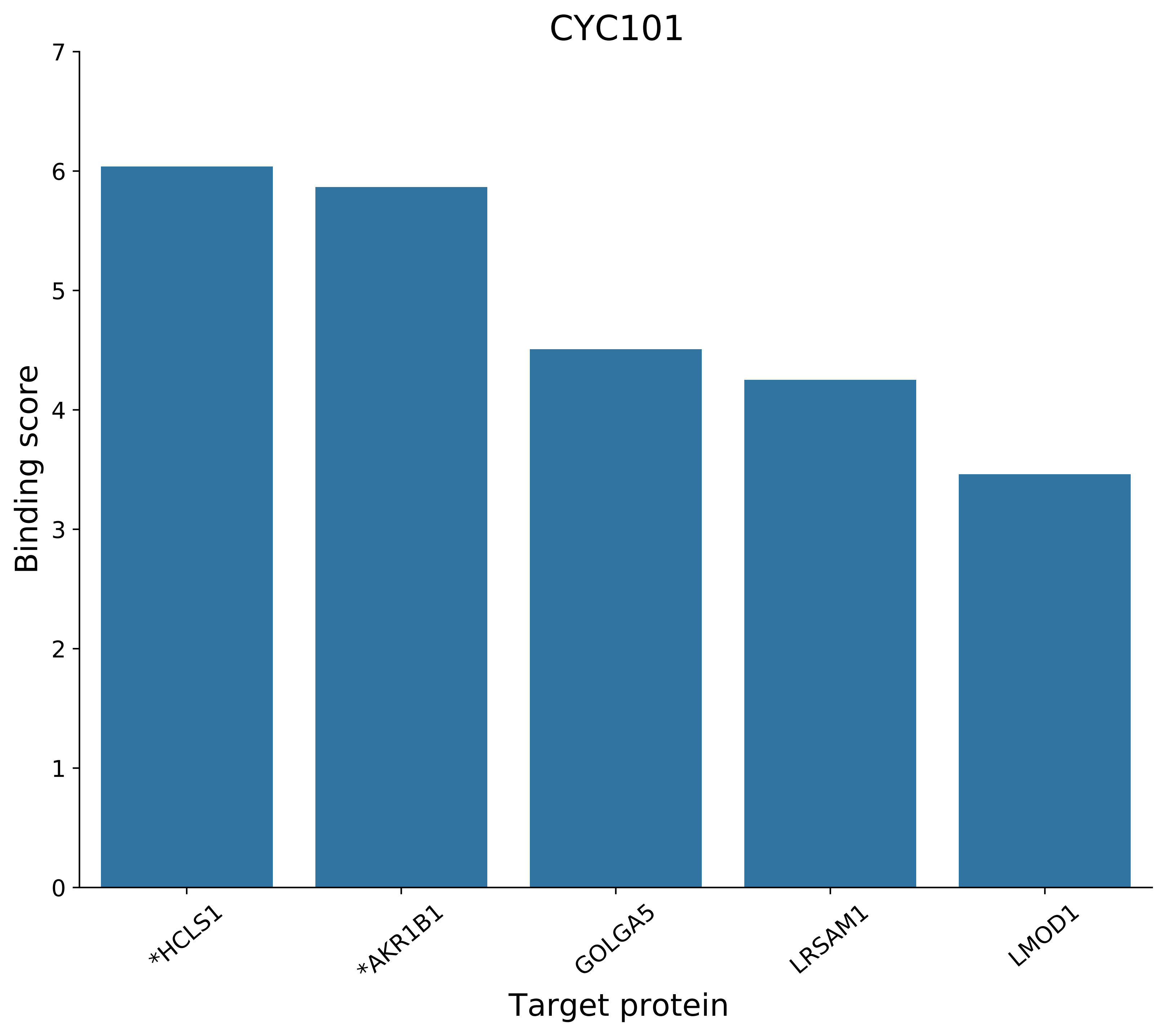

CYC101
Binding score
0
1
2
3
4
5
6
7
*HCLS1
*AKR1B1
GOLGA5
LRSAM1
LMOD1
Target protein

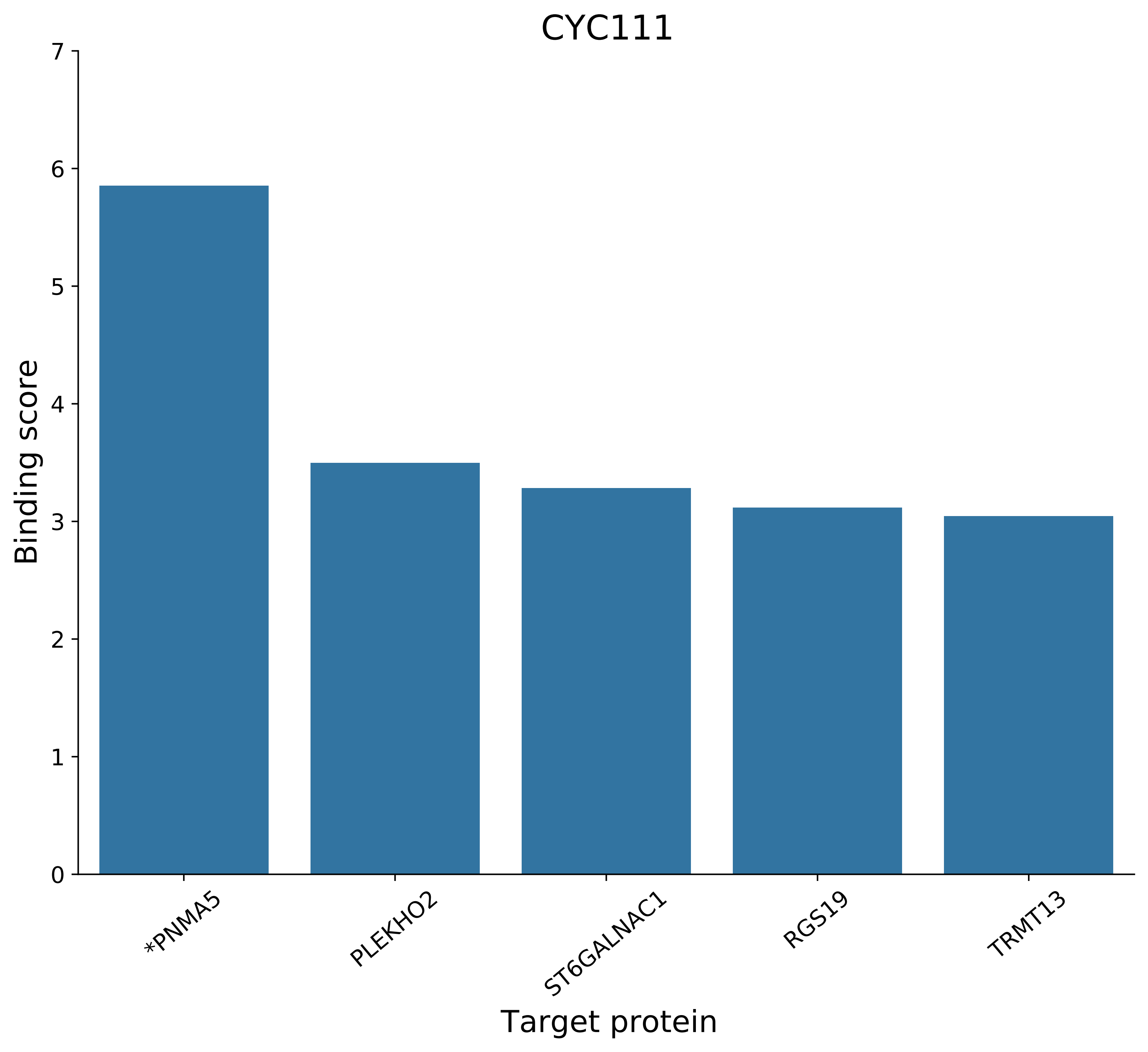

CYC111
Binding score
Target protein
0
1
2
3
4
5
6
7
*PNMA5
PLEKHO2
ST6GALNAC1
RGS19
TRMT13

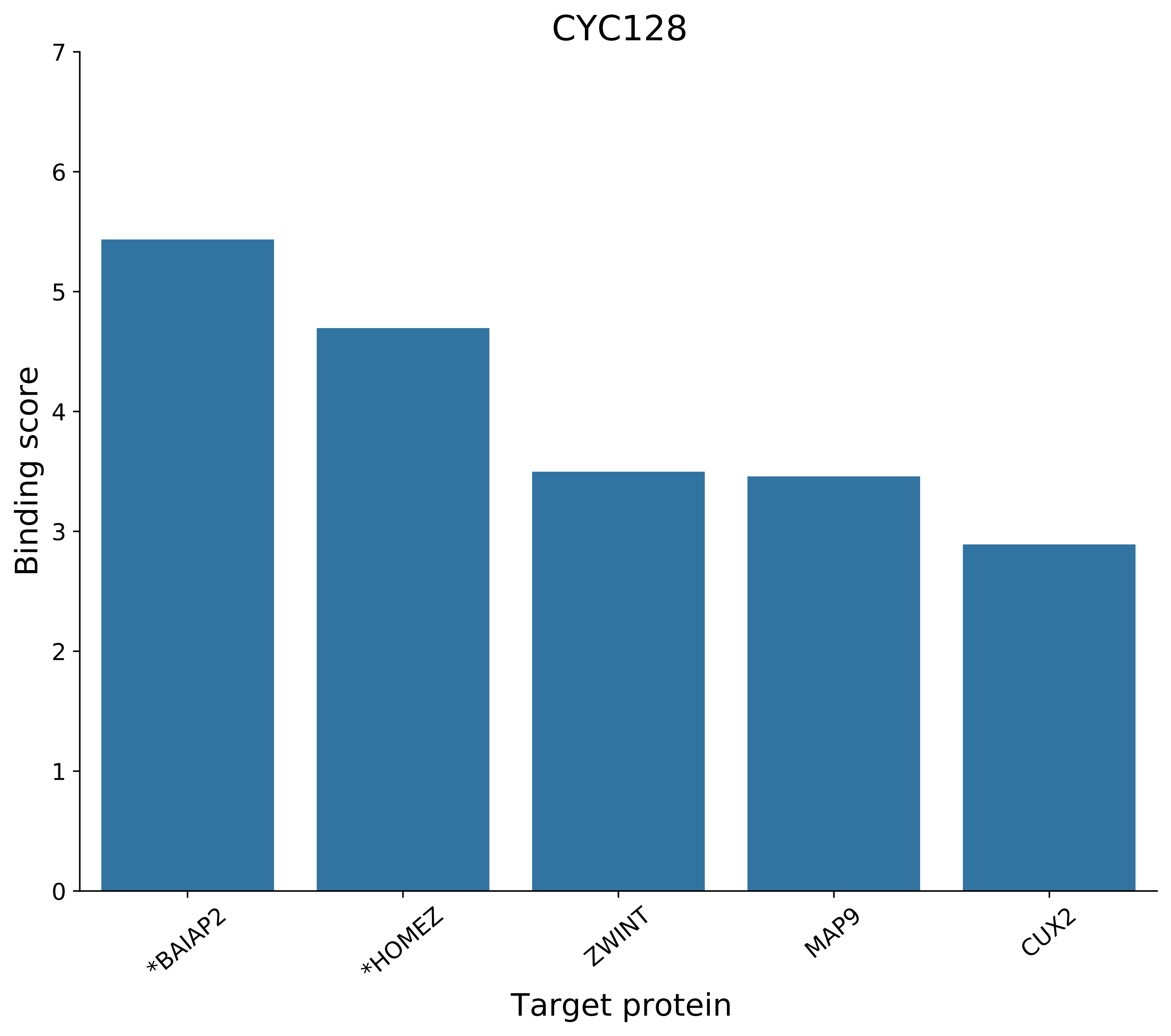
CYC128
Binding score
0
1
2
3
4
5
6
7
*BAIAP2
*HOMEZ
ZWINT
MAP9
CUX2
Target protein

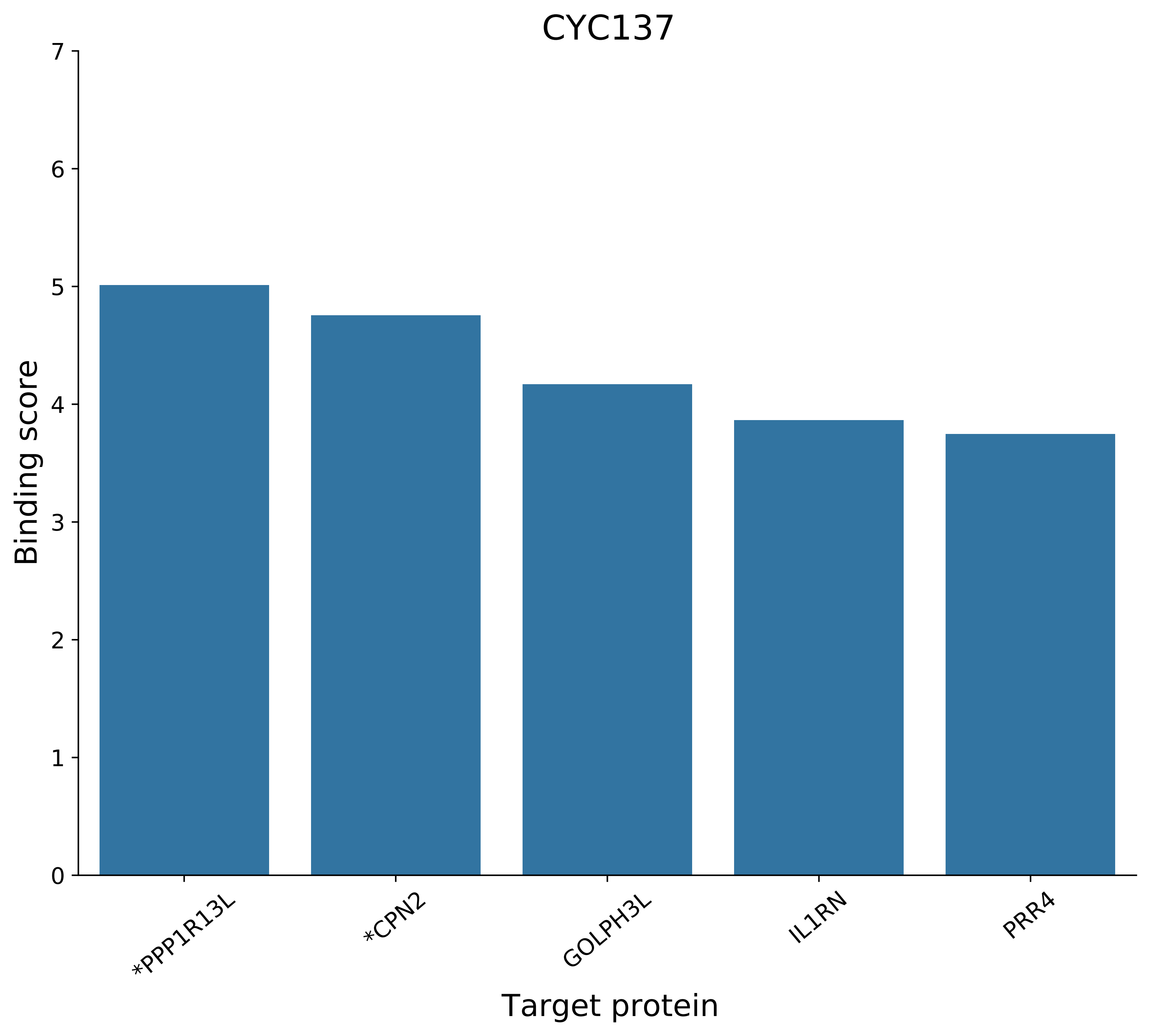
CYC137
Binding score
0
1
2
3
4
5
6
7
*PPP1R13L
*CPN2
GOLPH3L
IL1RN
PRR4
Target protein

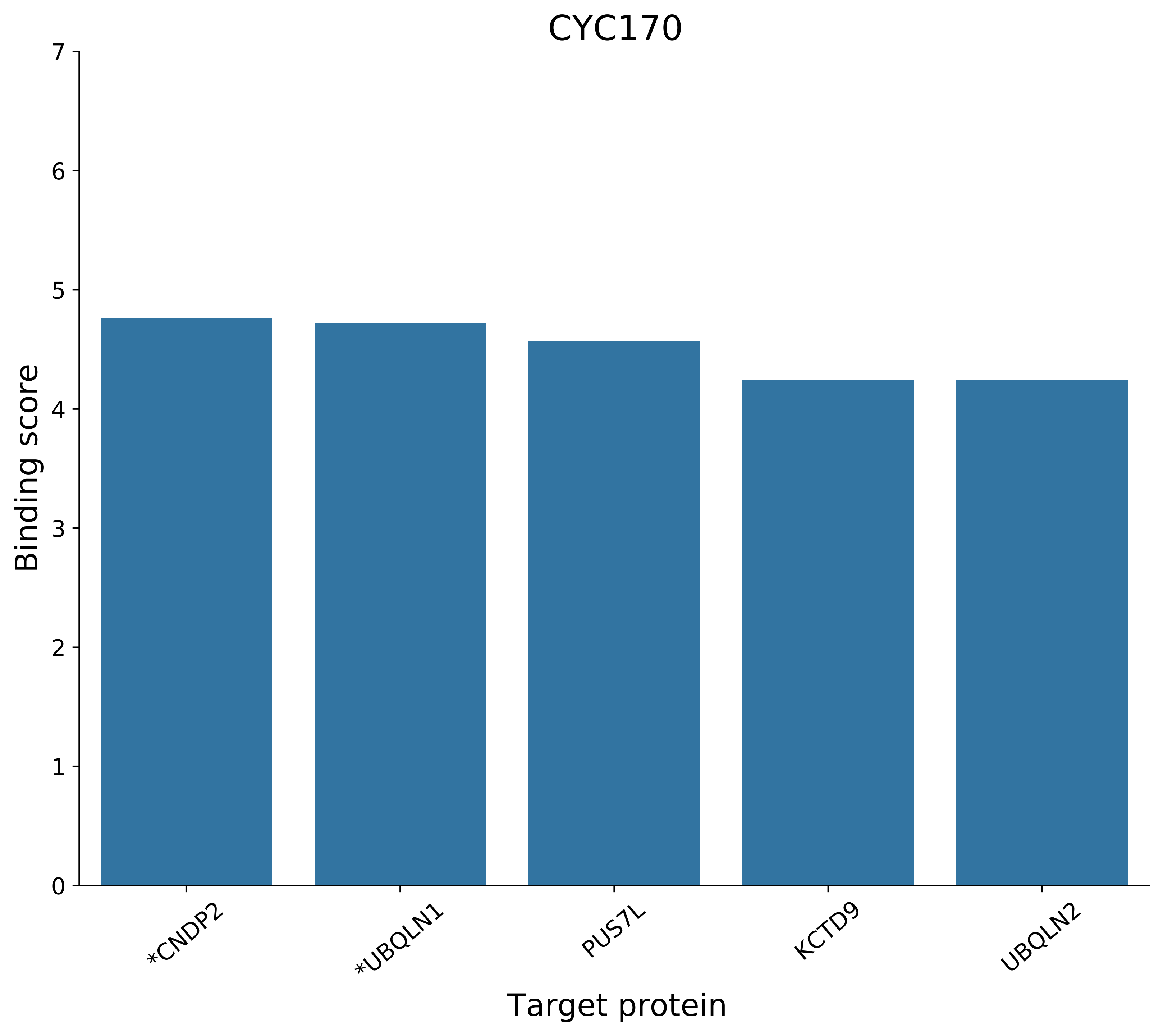
CYC170
Binding score
7
6
5
4
3
2
1
0
*CNDP2
*UBQLN1
PUS7L
KCTD9
UBQLN2
Target protein

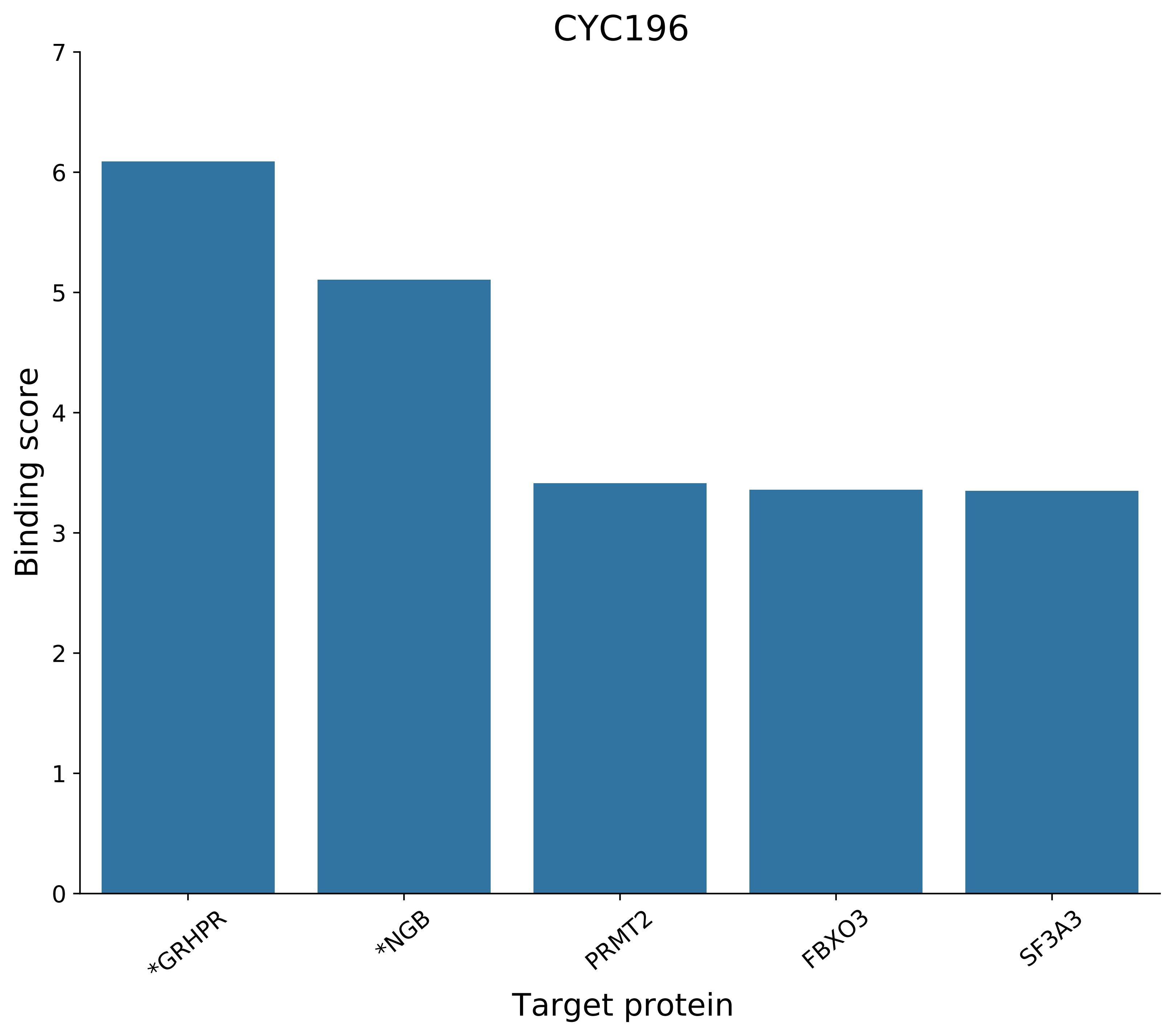
CYC196
Binding score
0
1
2
3
4
5
6
7
*GRHPR
*NGB
PRMT2
FBXO3
SF3A3
Target protein

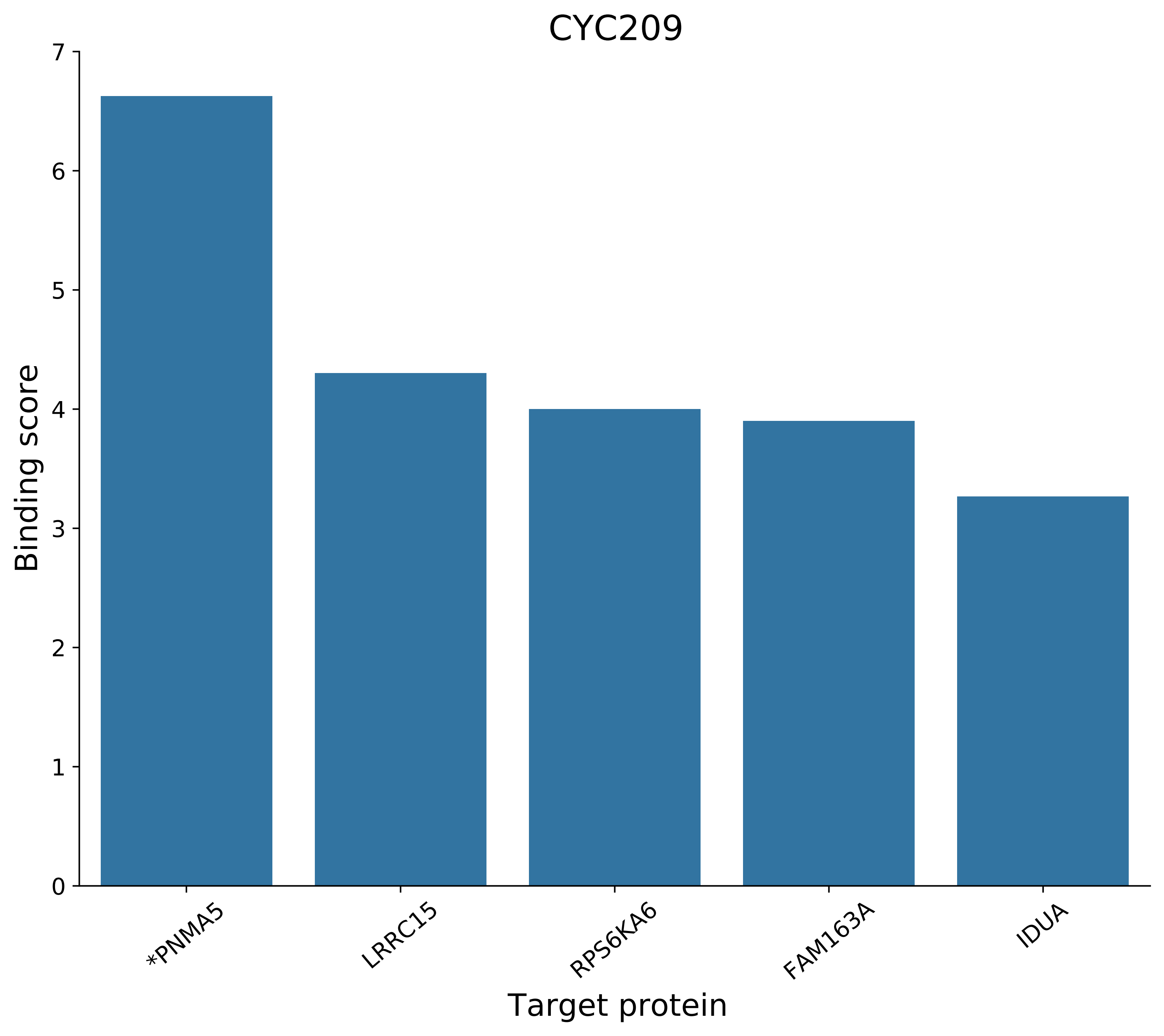
CYC209
Binding score
0
1
2
3
4
5
6
7
*PNMA5
LRRC15
RPS6KA6
FAM163A
IDUA
Target protein

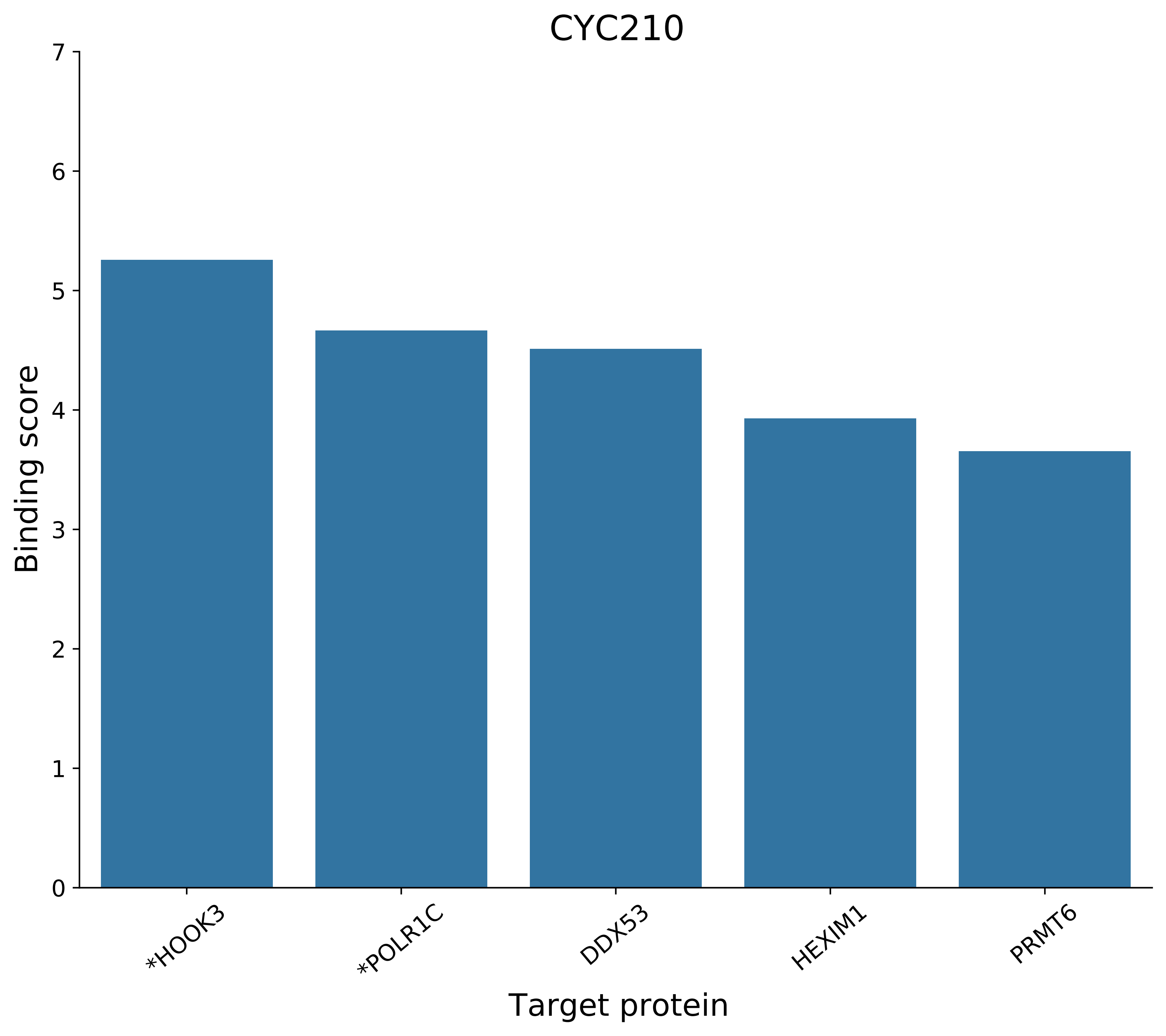
CYC210
Binding score
Target protein
0
1
2
3
4
5
6
7
*HOOK3
*POLR1C
DDX53
HEXIM1
PRMT6

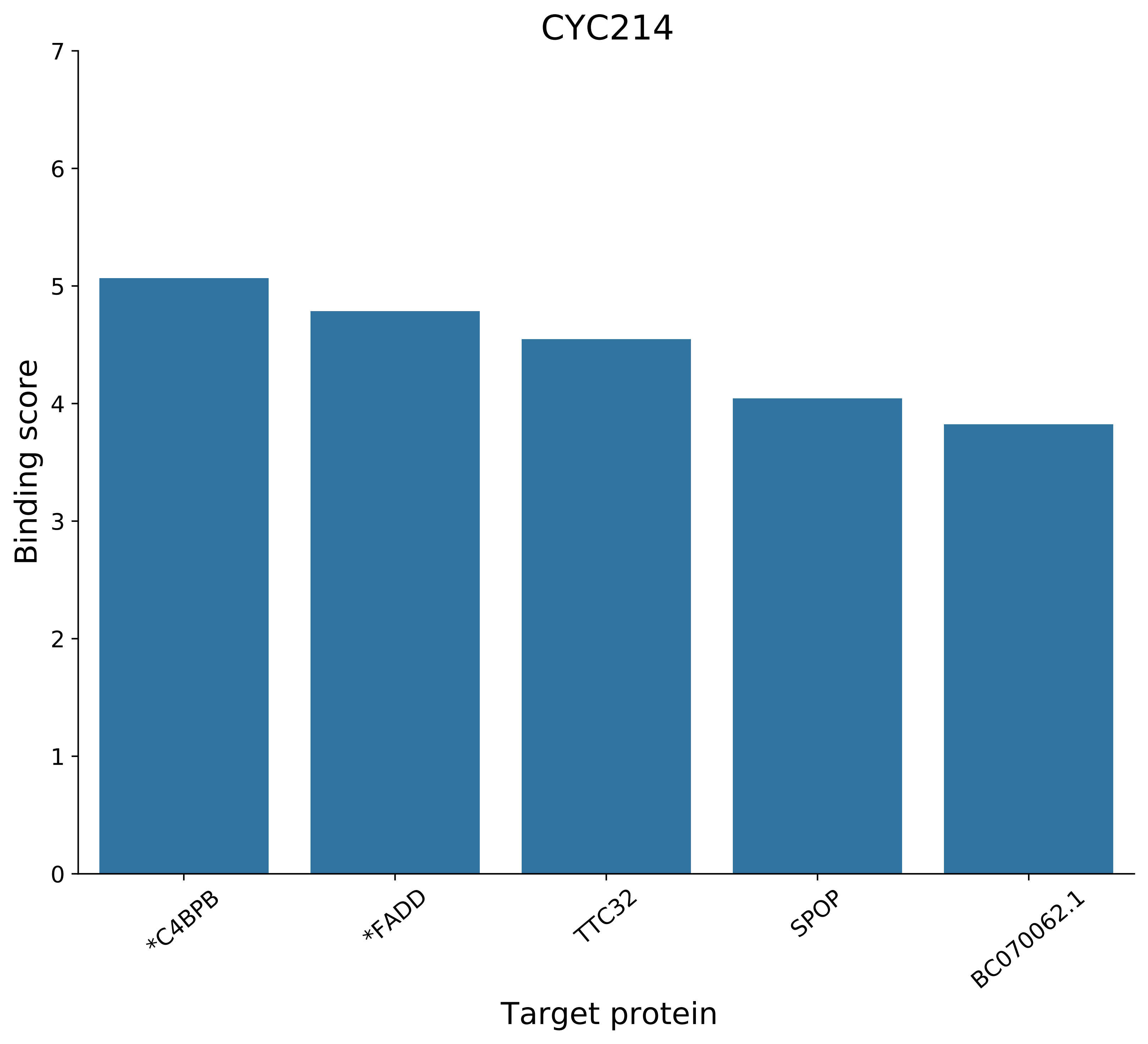
CYC214
Binding score
7
6
5
4
3
2
1
0
*C4BPB
*FADD
TTC32
SPOP
BC070062.1
Target protein

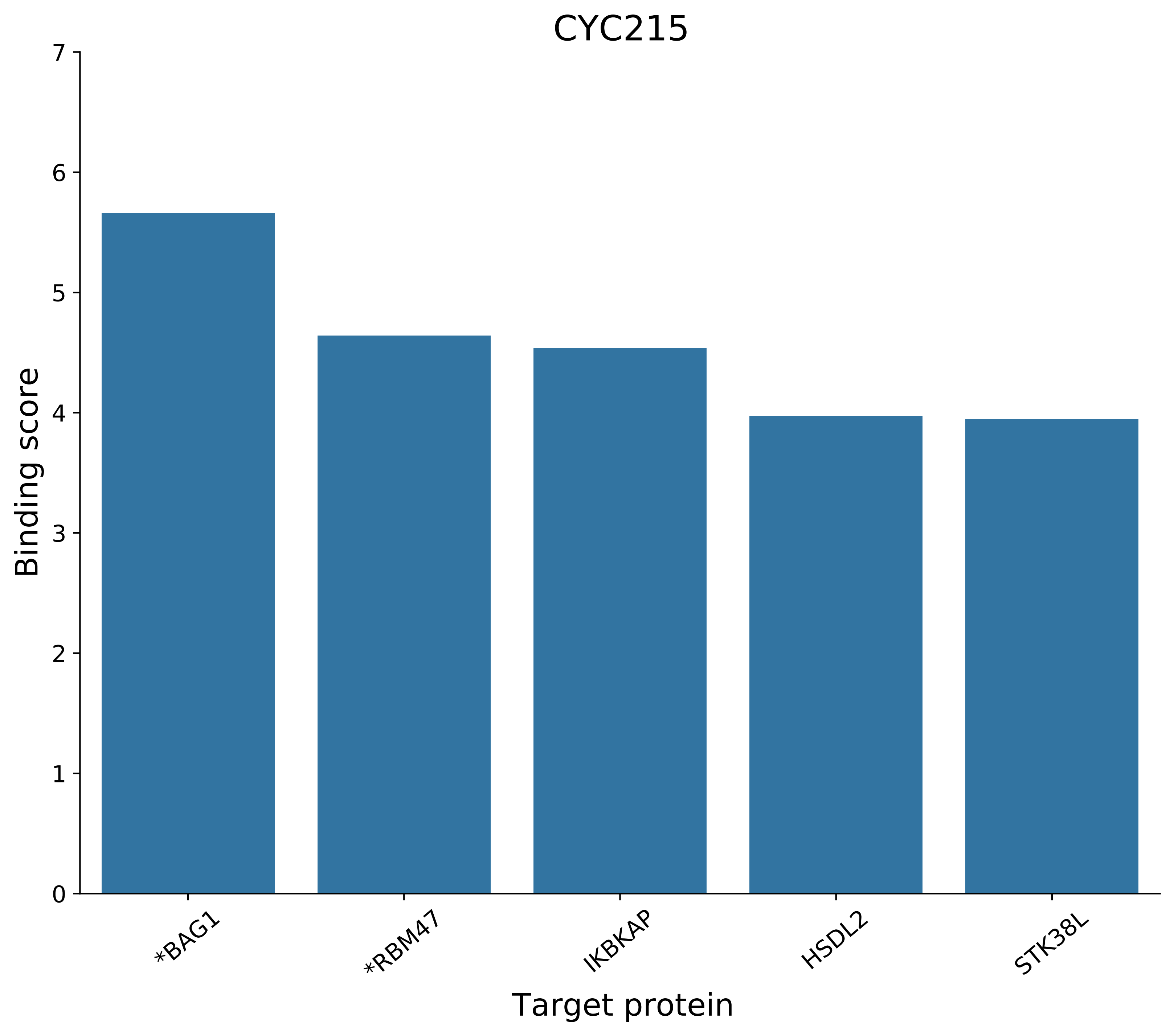
CYC215
Binding score
0
1
2
3
4
5
6
7
*BAG1
*RBM47
IKBKAP
HSDL2
STK38L
Target protein

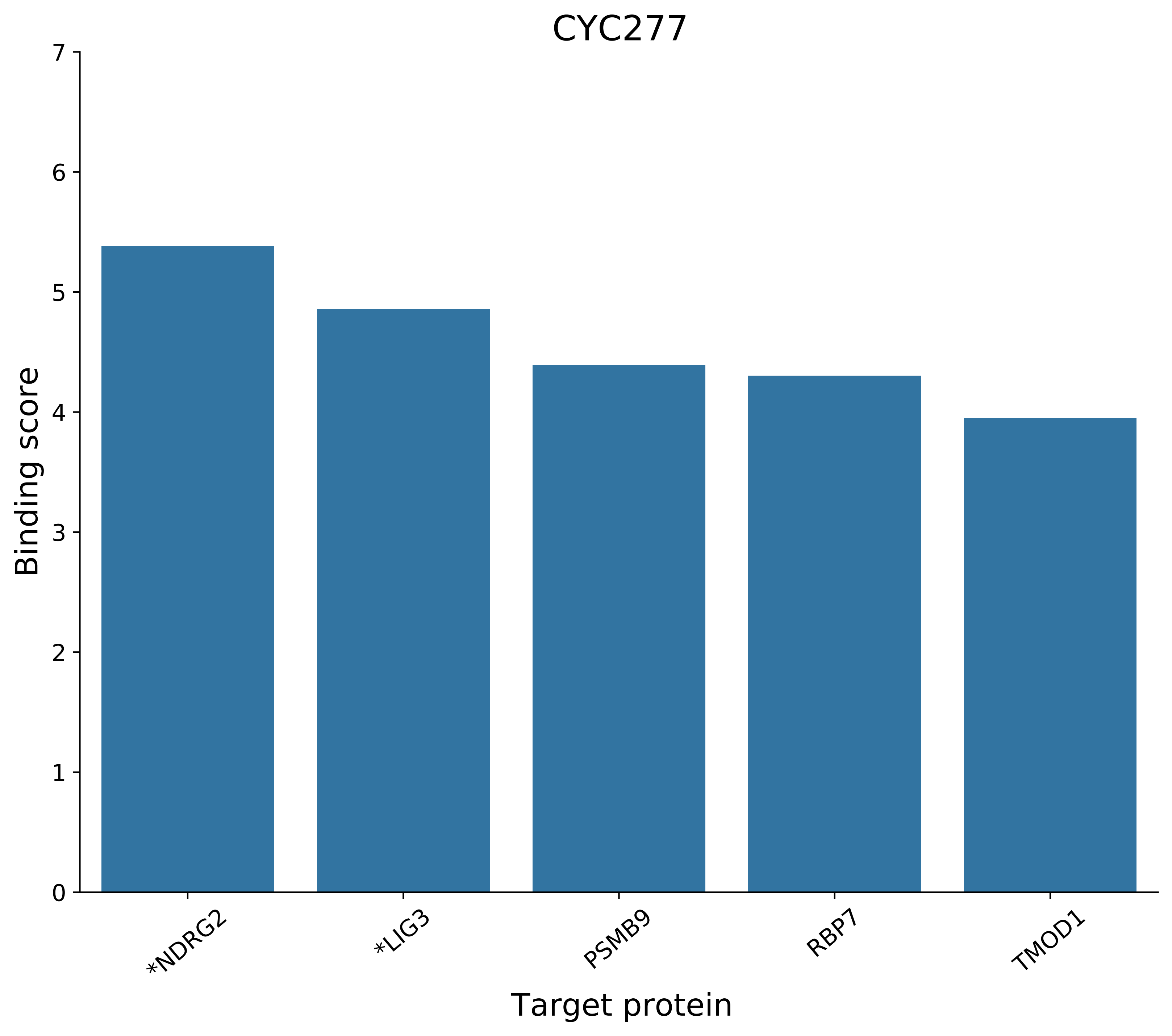

CYC277
Binding score
0
1
2
3
4
5
6
7
*NDRG2
*LIG3
PSMB9
RBP7
TMOD1
Target protein

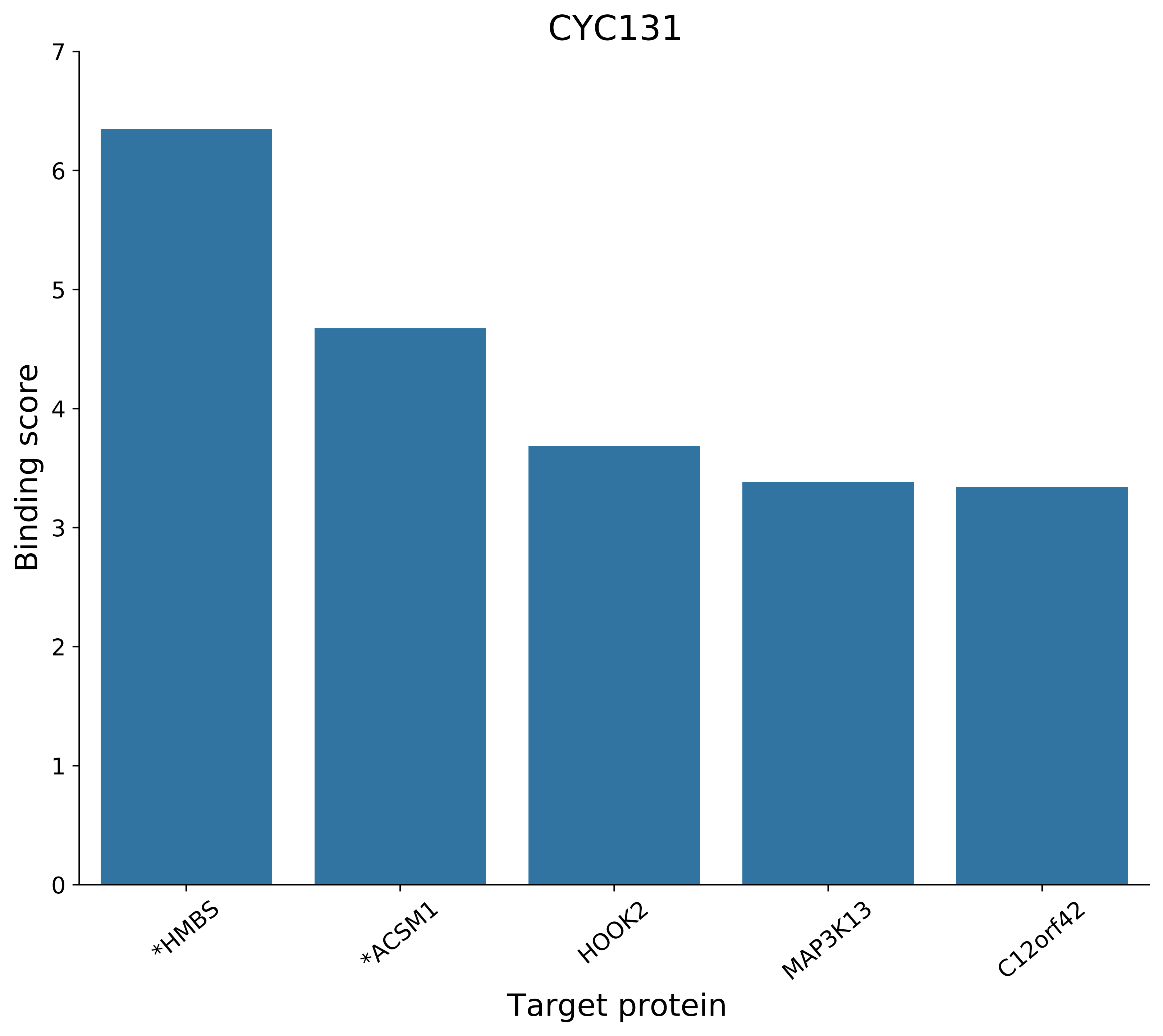

CYC131
Binding score
Target protein
0
1
2
3
4
5
6
7
*HMBS
*ACSM1
HOOK2
MAP3K13
C12orf42

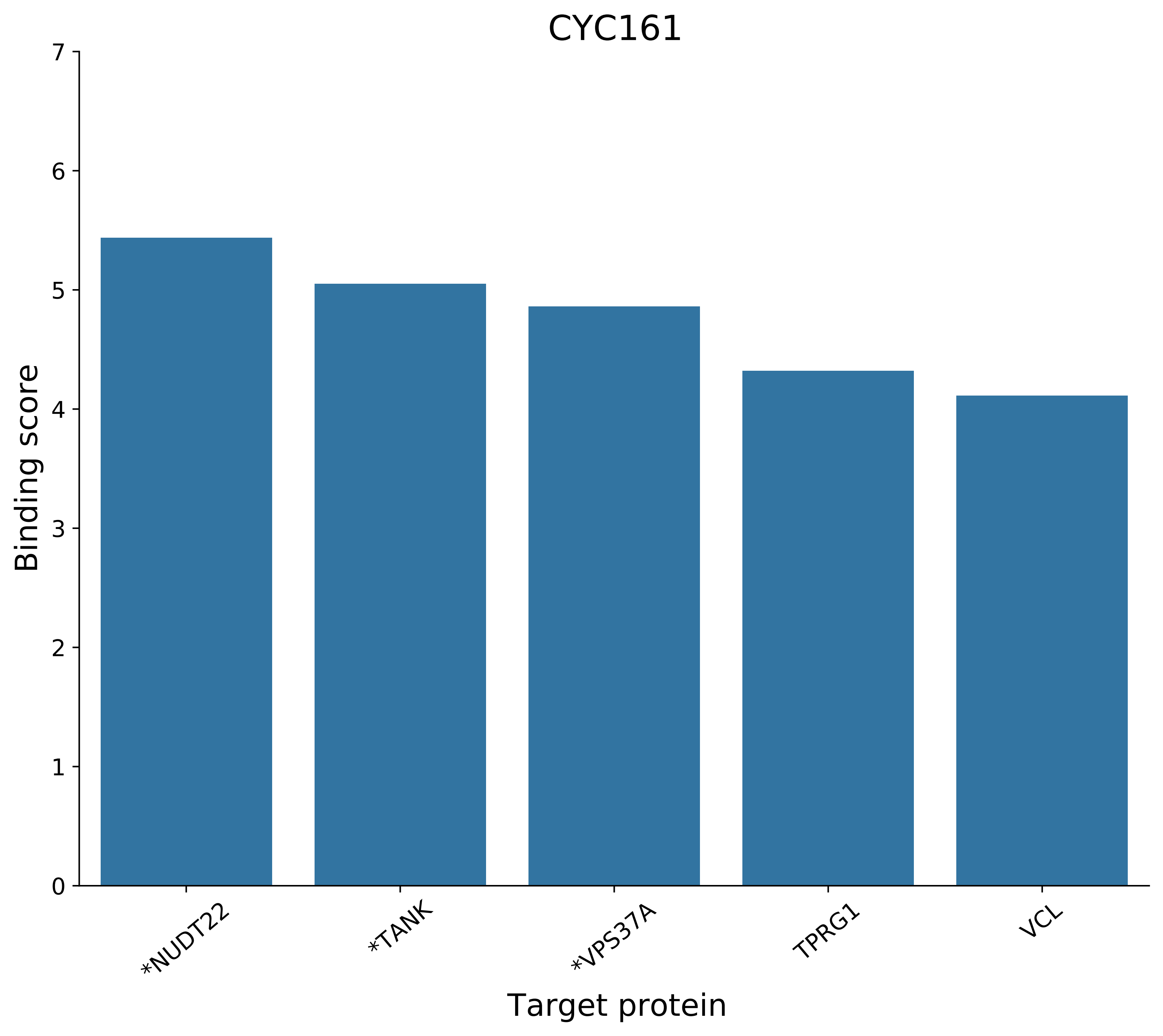
CYC161
Binding score
0
1
2
3
4
5
6
7
*NUDT22
*TANK
*VPS37A
TPRG1
VCL
Target protein

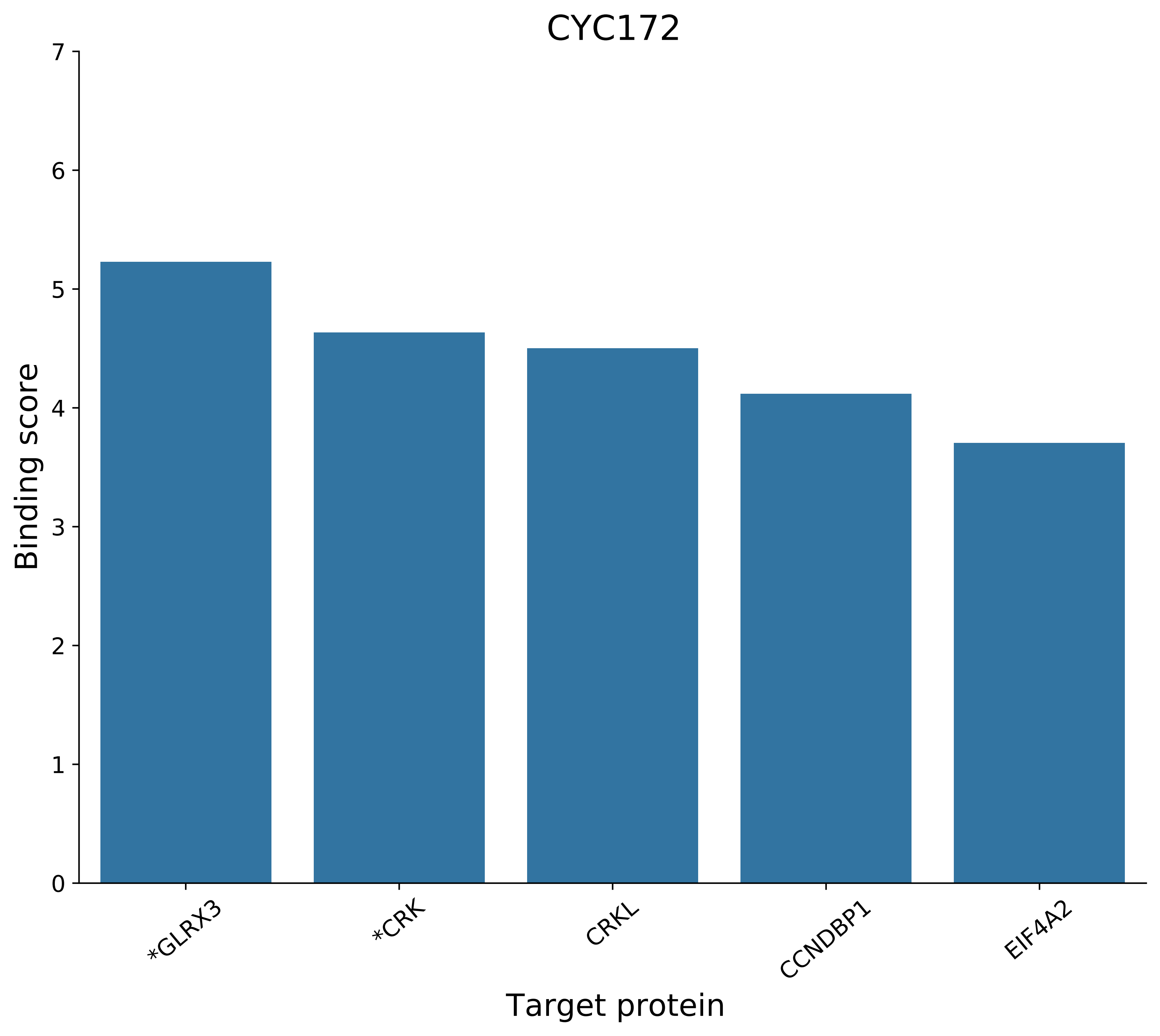
CYC172
Binding score
0
1
2
3
4
5
6
7
*GLRX3
*CRK
CRKL
CCNDBP1
EIF4A2
Target protein

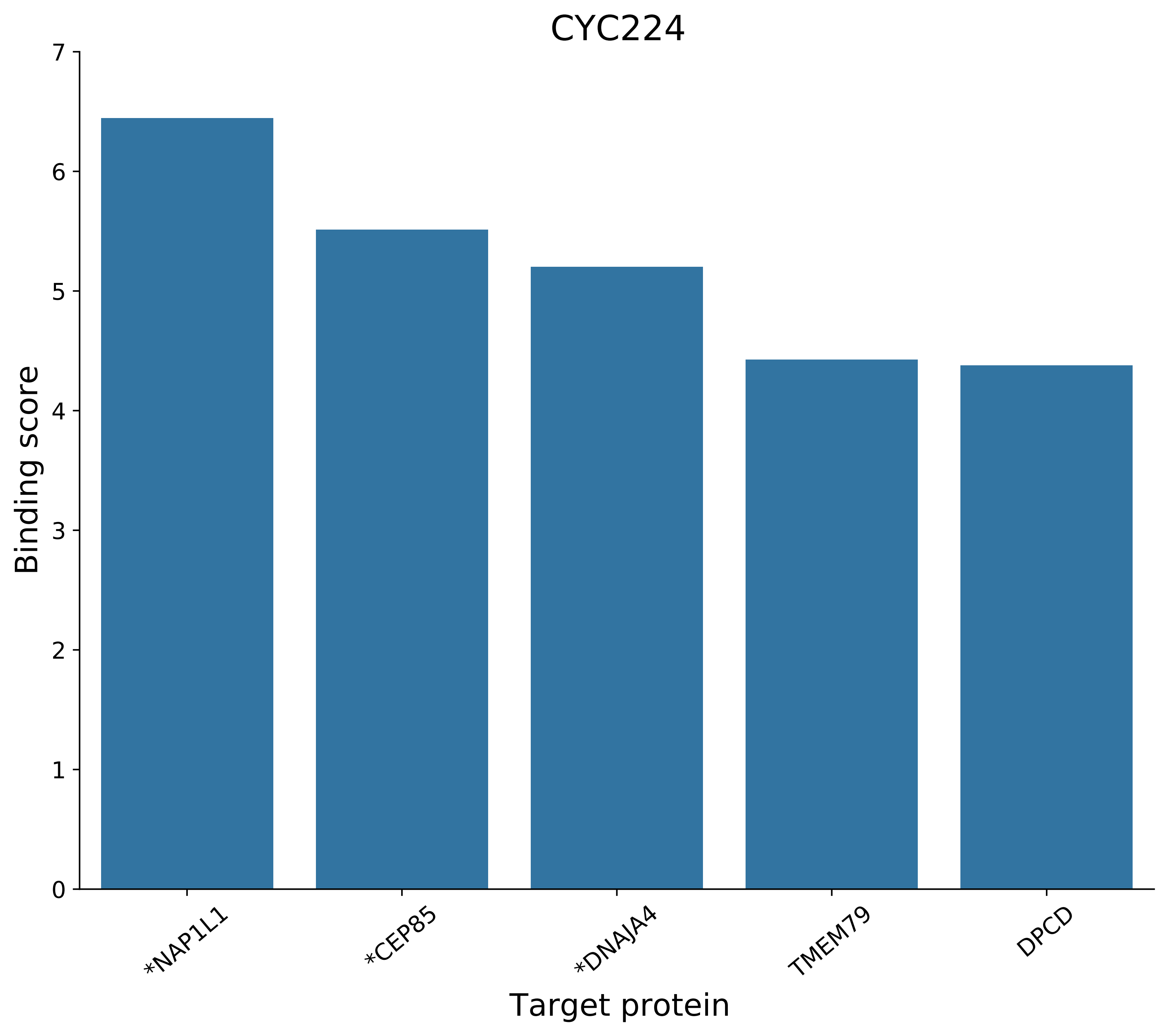
CYC224
Binding score
7
6
5
4
3
2
1
0
*NAP1L1
*CEP85
*DNAJA4
TMEM79
DPCD
Target protein

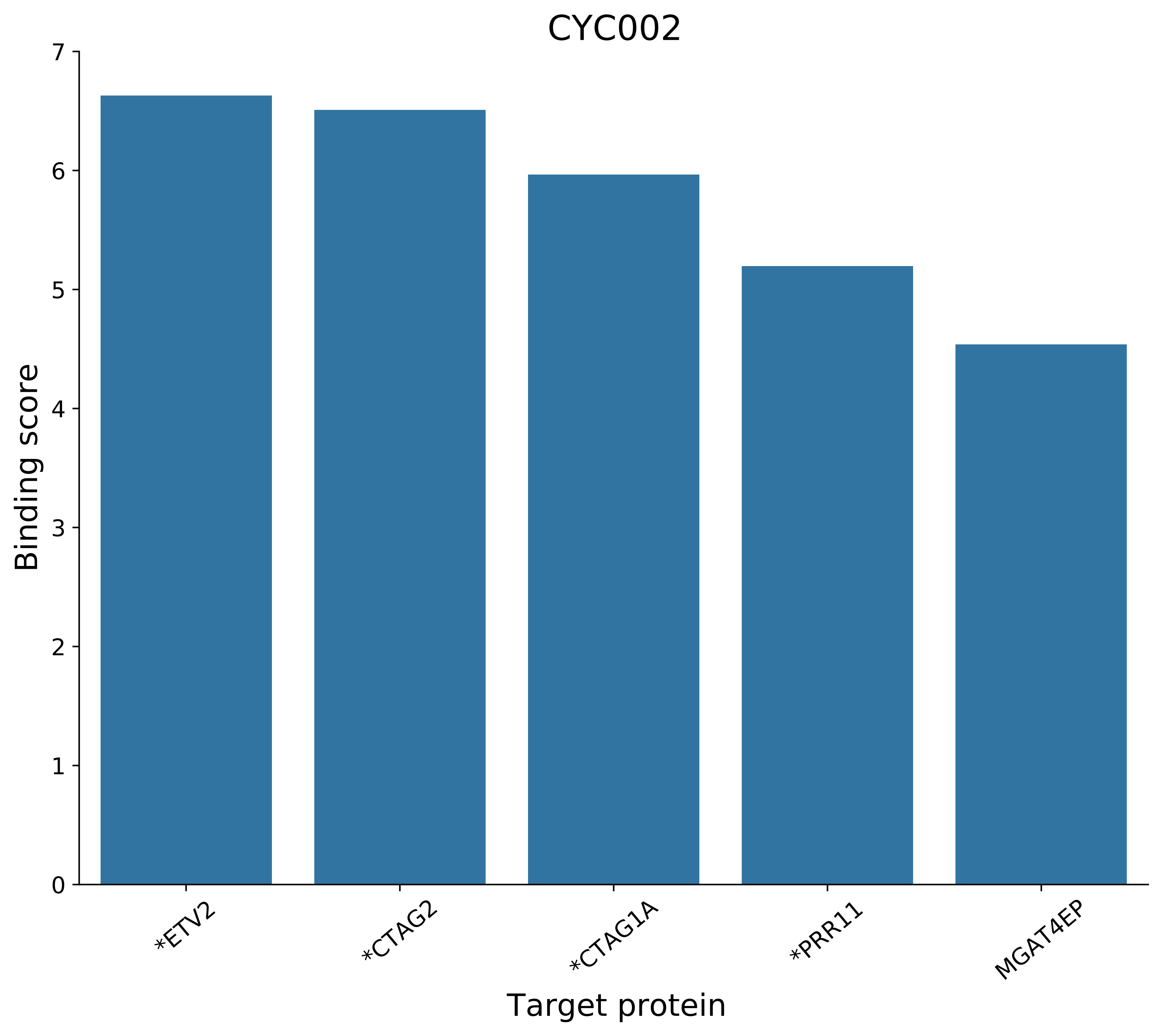
CYC002
Binding score
Target protein
0
1
2
3
4
5
6
7
*ETV2
*CTAG2
*CTAG1A
*PRR11
MGAT4EP

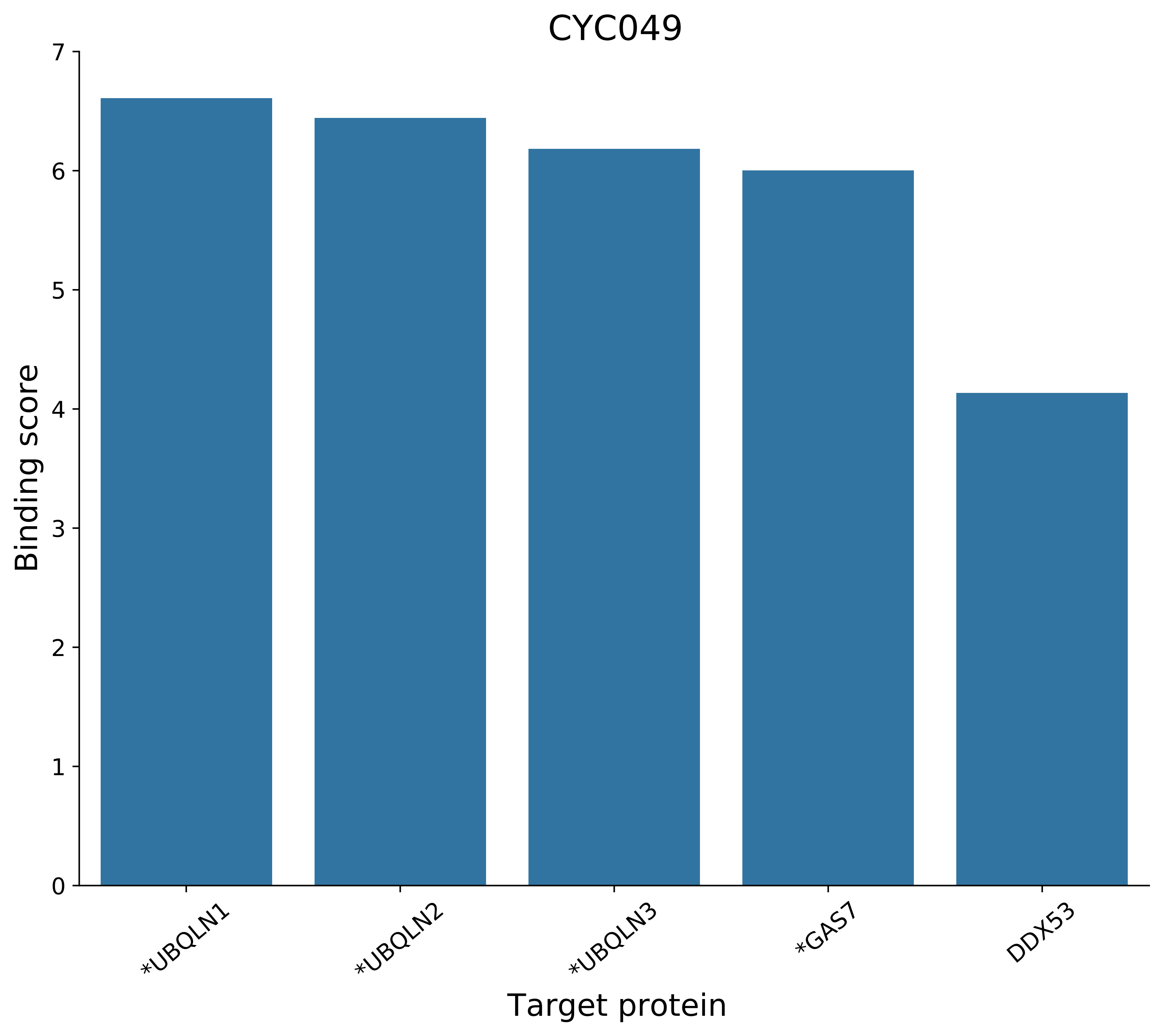
CYC049
Binding score
0
1
2
3
4
5
6
7
*UBQLN1
*UBQLN2
*UBQLN3
*GAS7
DDX53
Target protein

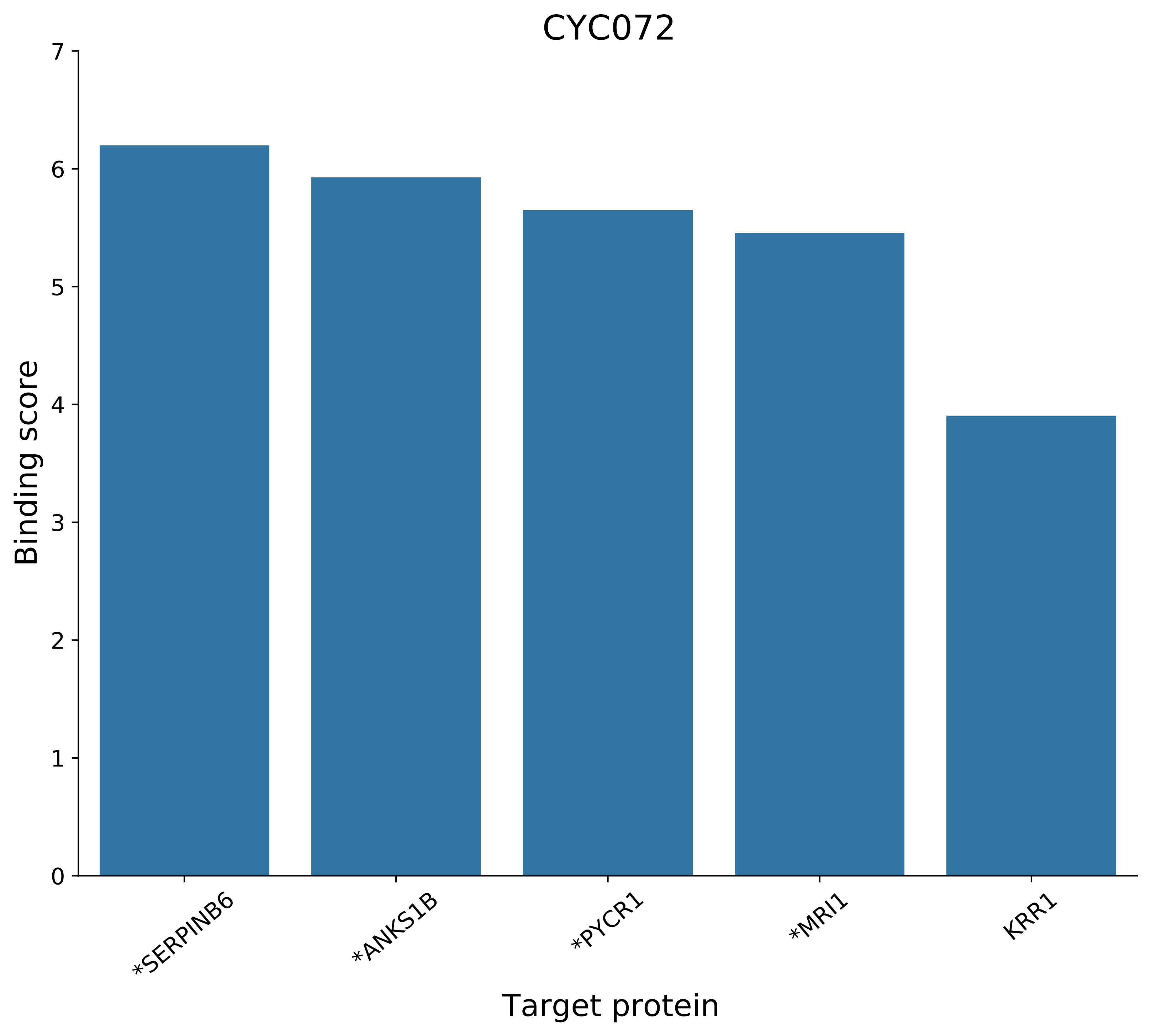
CYC072
Binding score
Target protein
0
1
2
3
4
5
6
7
*SERPINB6
*ANKS1B
*PYCR1
*MRI1
KRR1

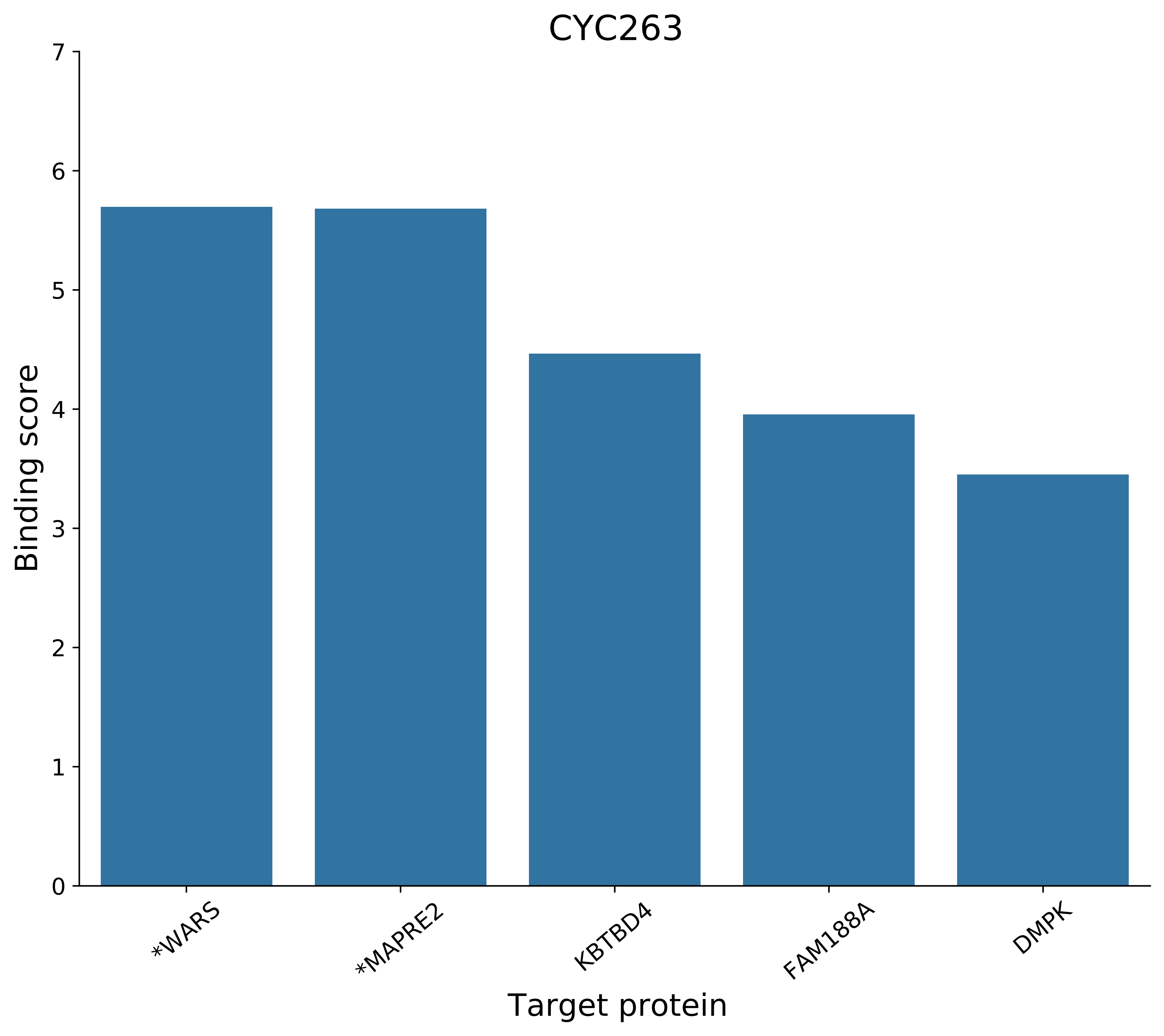

CYC263
Binding score
7
6
5
4
3
2
1
0
*WARS
*MAPRE2
KBTBD4
FAM188A
DMPK
Target protein

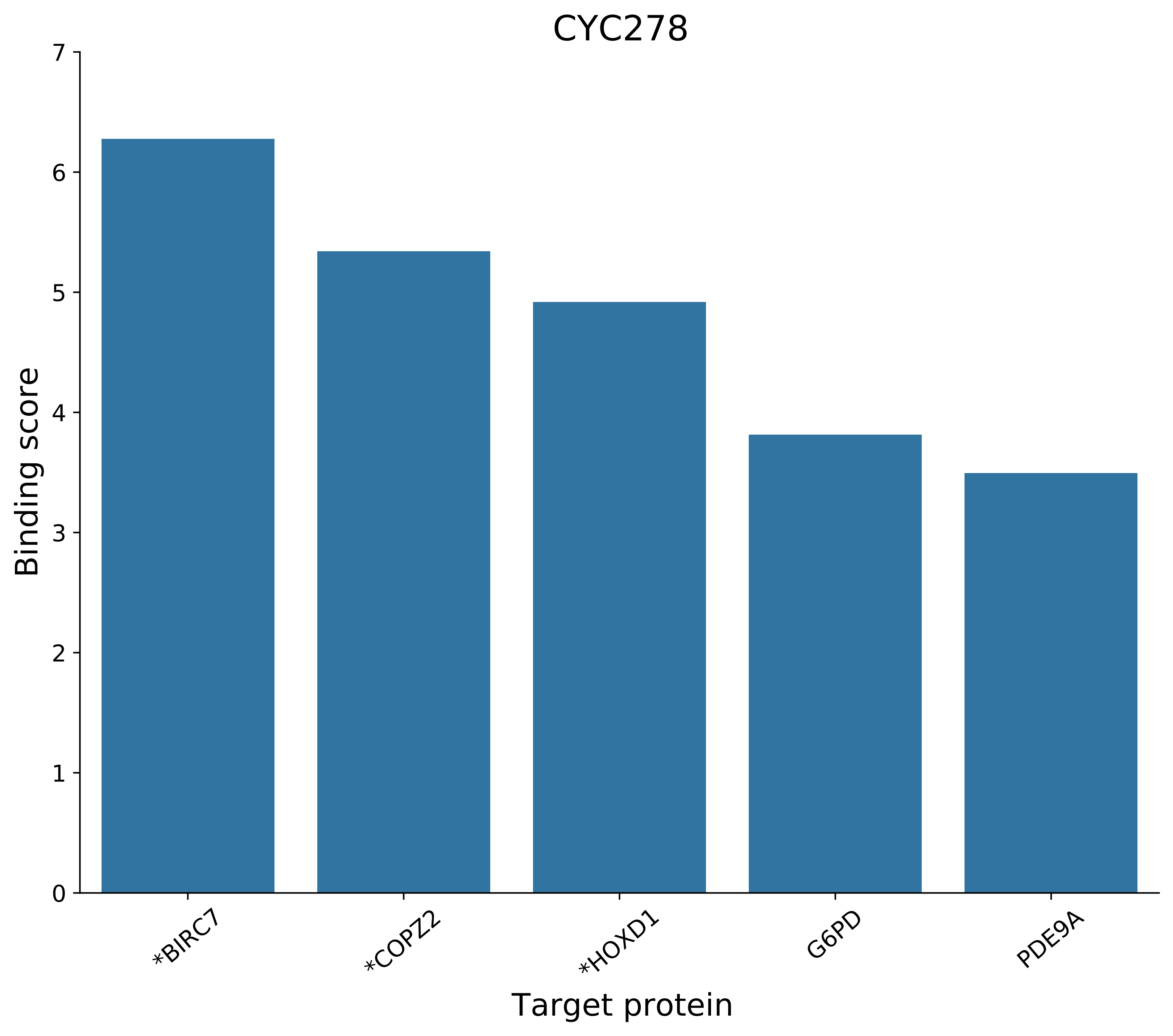
CYC278
Binding score
0
1
2
3
4
5
6
7
*BIRC7
*COPZ2
*HOXD1
G6PD
PDE9A
Target protein

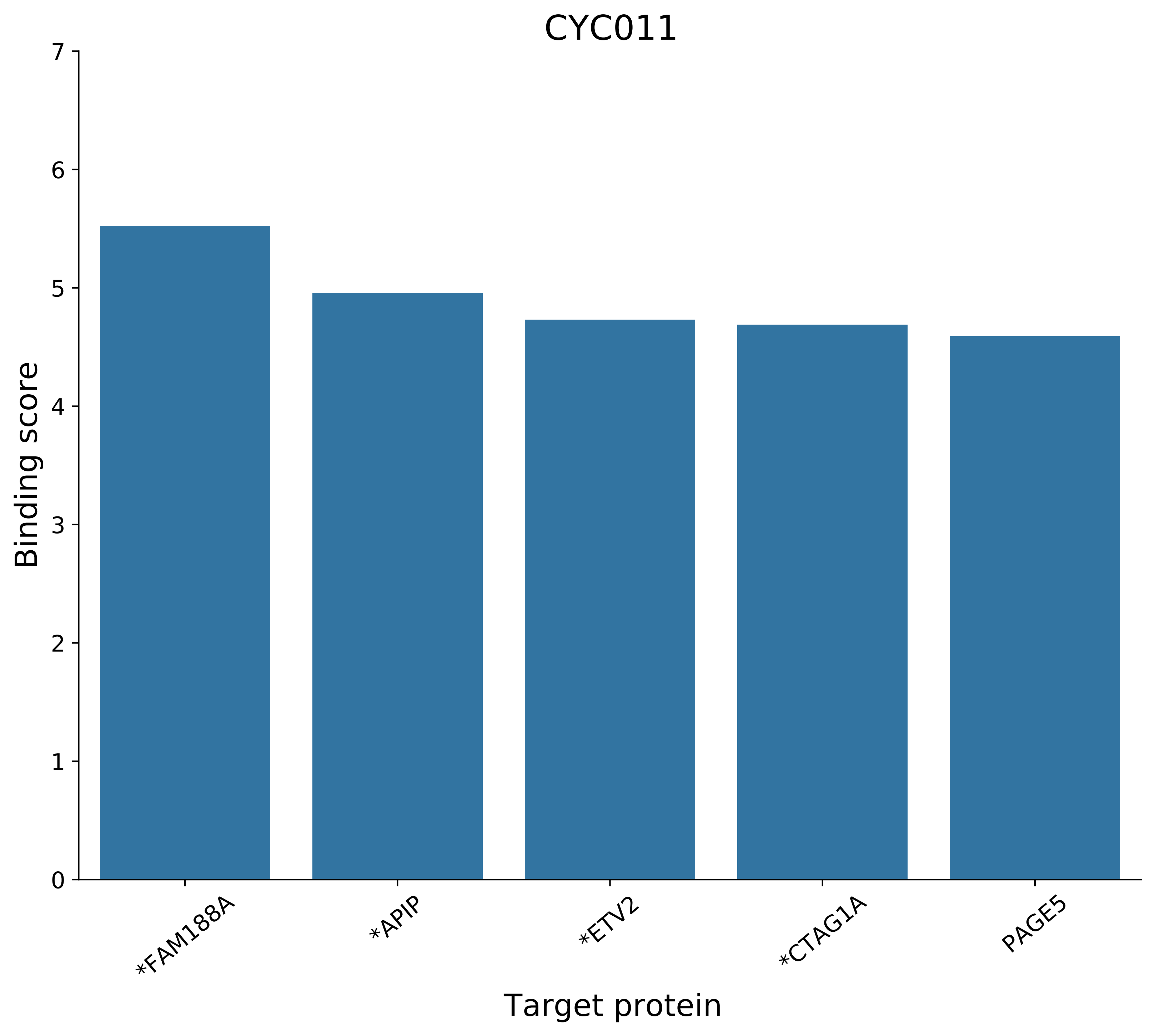

CYC011
Binding score
0
1
2
3
4
5
6
7
*FAM188A
*APIP
*ETV2
*CTAG1A
PAGE5
Target protein

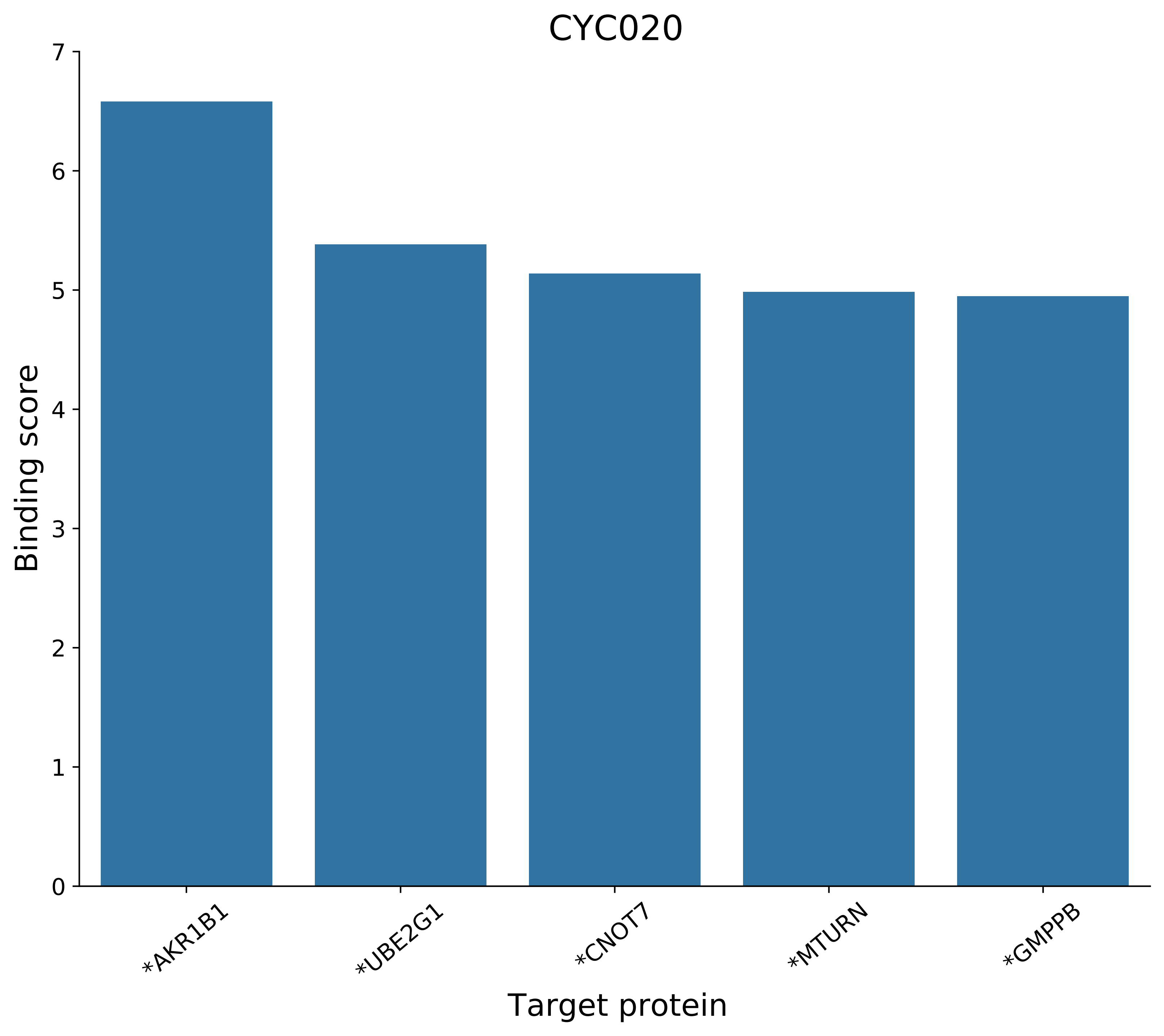

CYC020
Binding score
Target protein
*AKR1B1
*UBE2G1
*CNOT7
*MTURN
*GMPPB
0
1
2
3
4
5
6
7

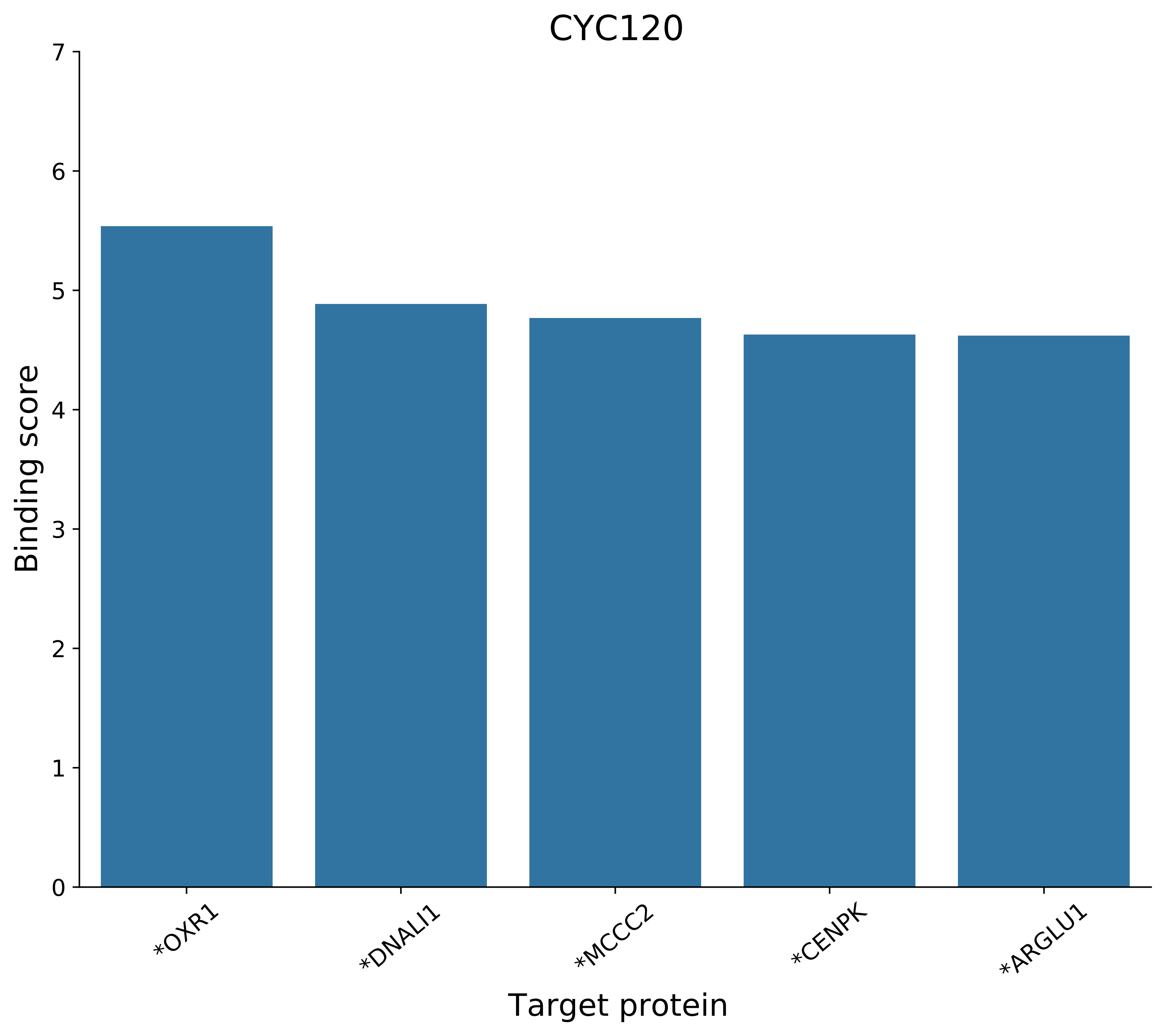

CYC120
Binding score
Target protein
0
1
2
3
4
5
6
7
*OXR1
*DNALI1
*MCCC2
*CENPK
*ARGLU1

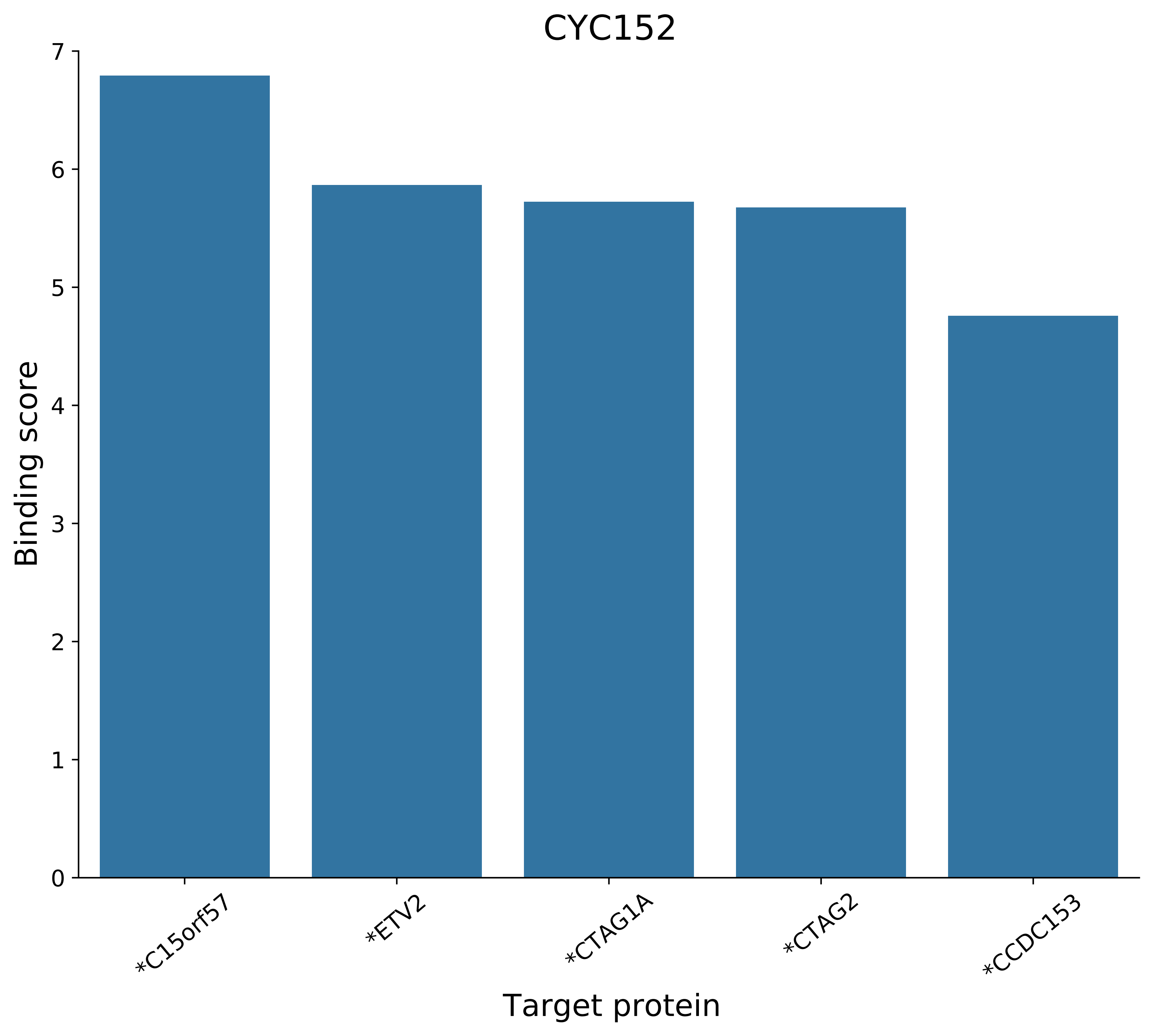
CYC152
Binding score
Target protein
0
1
2
3
4
5
6
7
*C15orf57
*ETV2
*CTAG1A
*CTAG2
*CCDC153

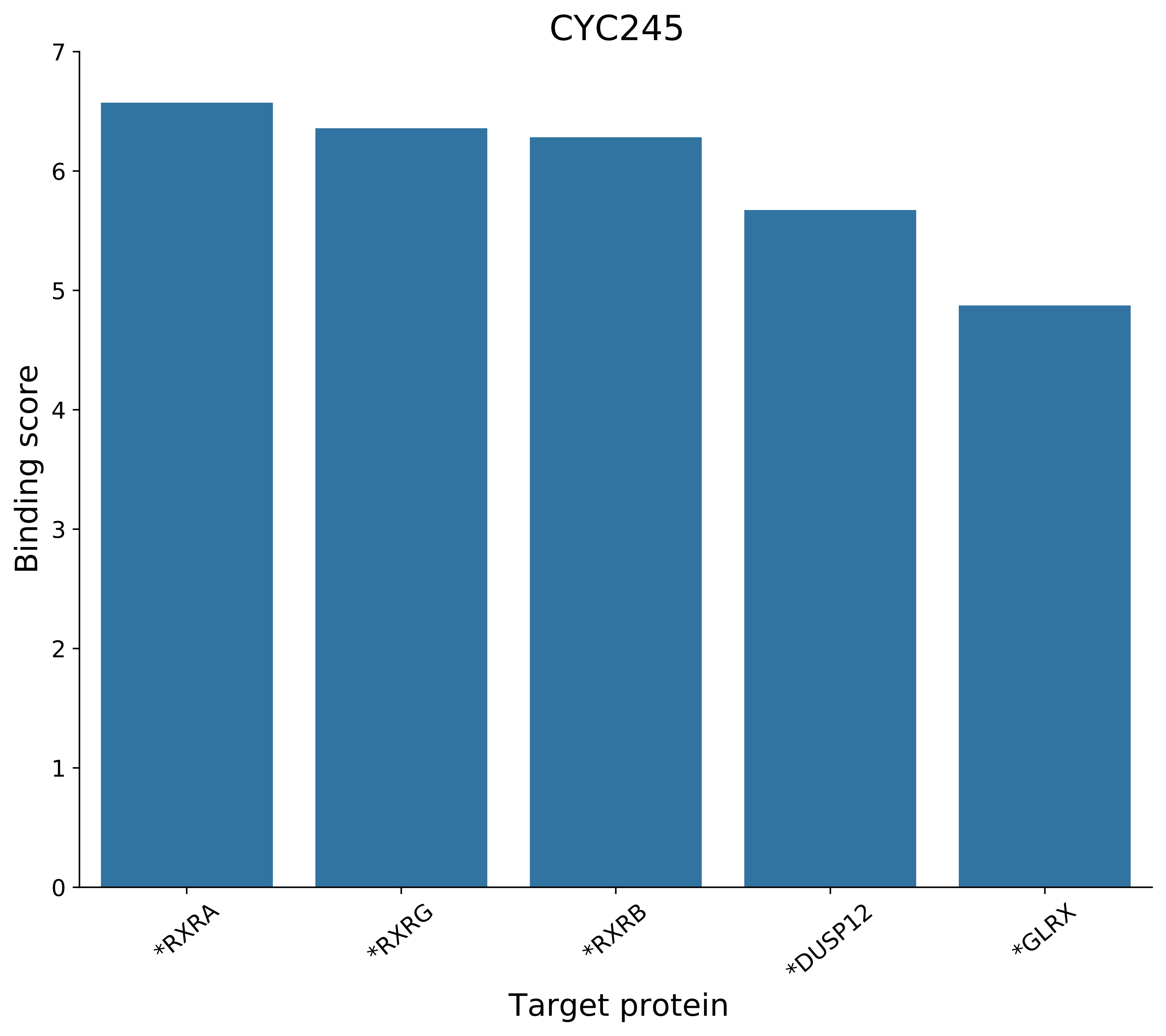
CYC245
Binding score
Target protein
*RXRA
*RXRG
*RXRB
*DUSP12
*GLRX
0
1
2
3
4
5
6
7

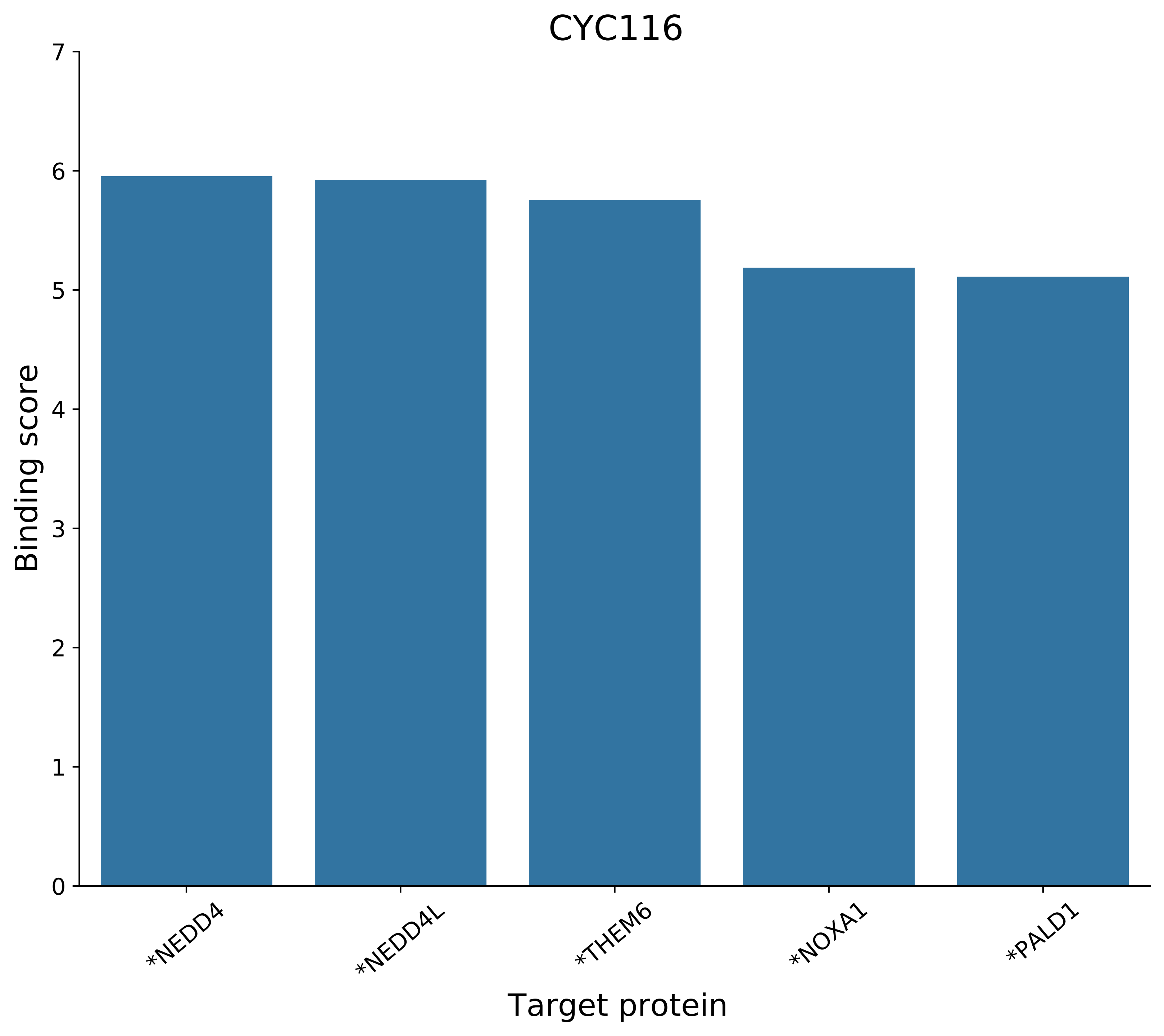
CYC116
Binding score
Target protein
0
1
2
3
4
5
6
7
*NEDD4
*NEDD4L
*THEM6
*NOXA1
*PALD1

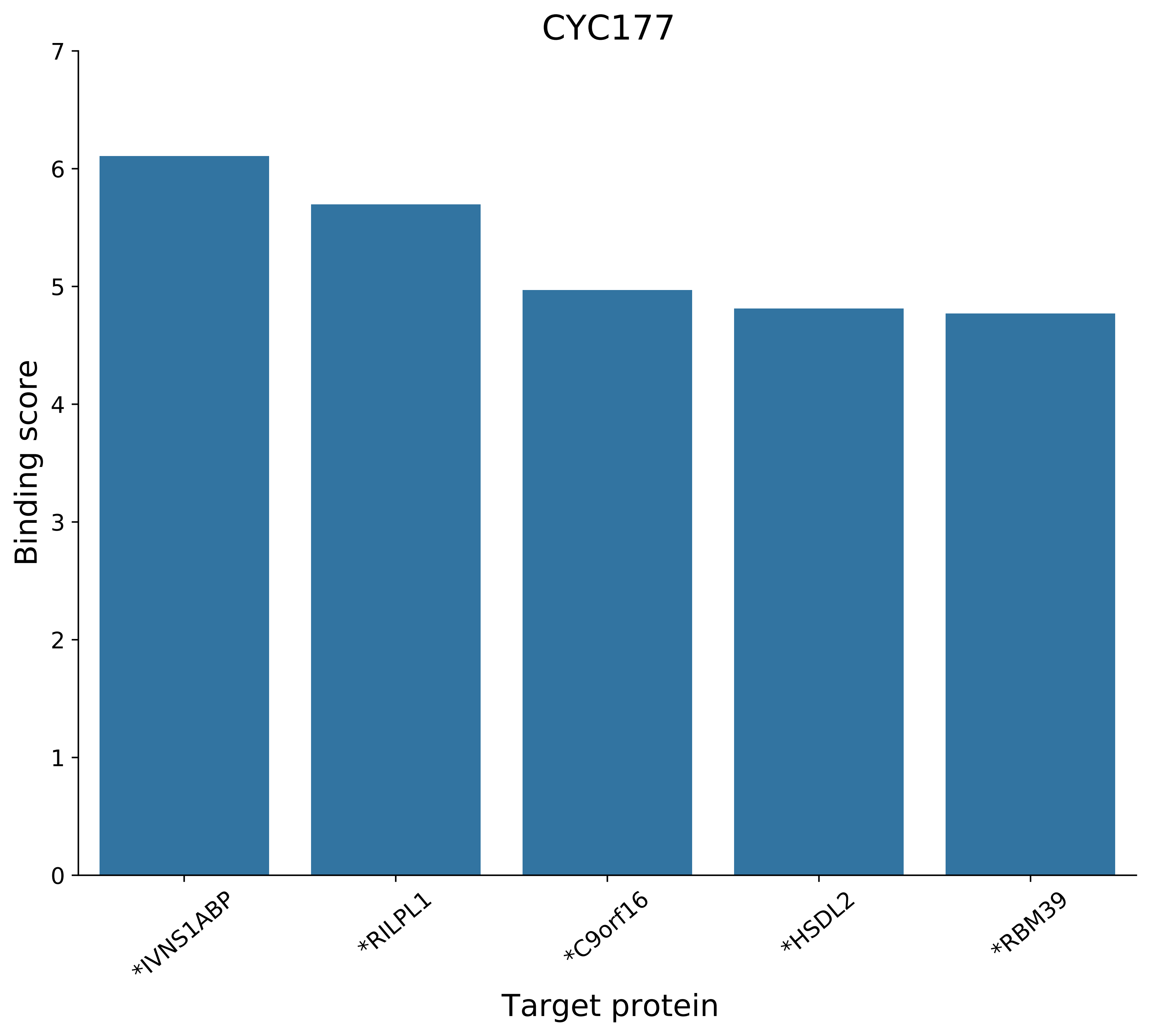
CYC177
Binding score
0
1
2
3
4
5
6
7
*IVNS1ABP
*RILPL1
*C9orf16
*HSDL2
*RBM39
Target protein

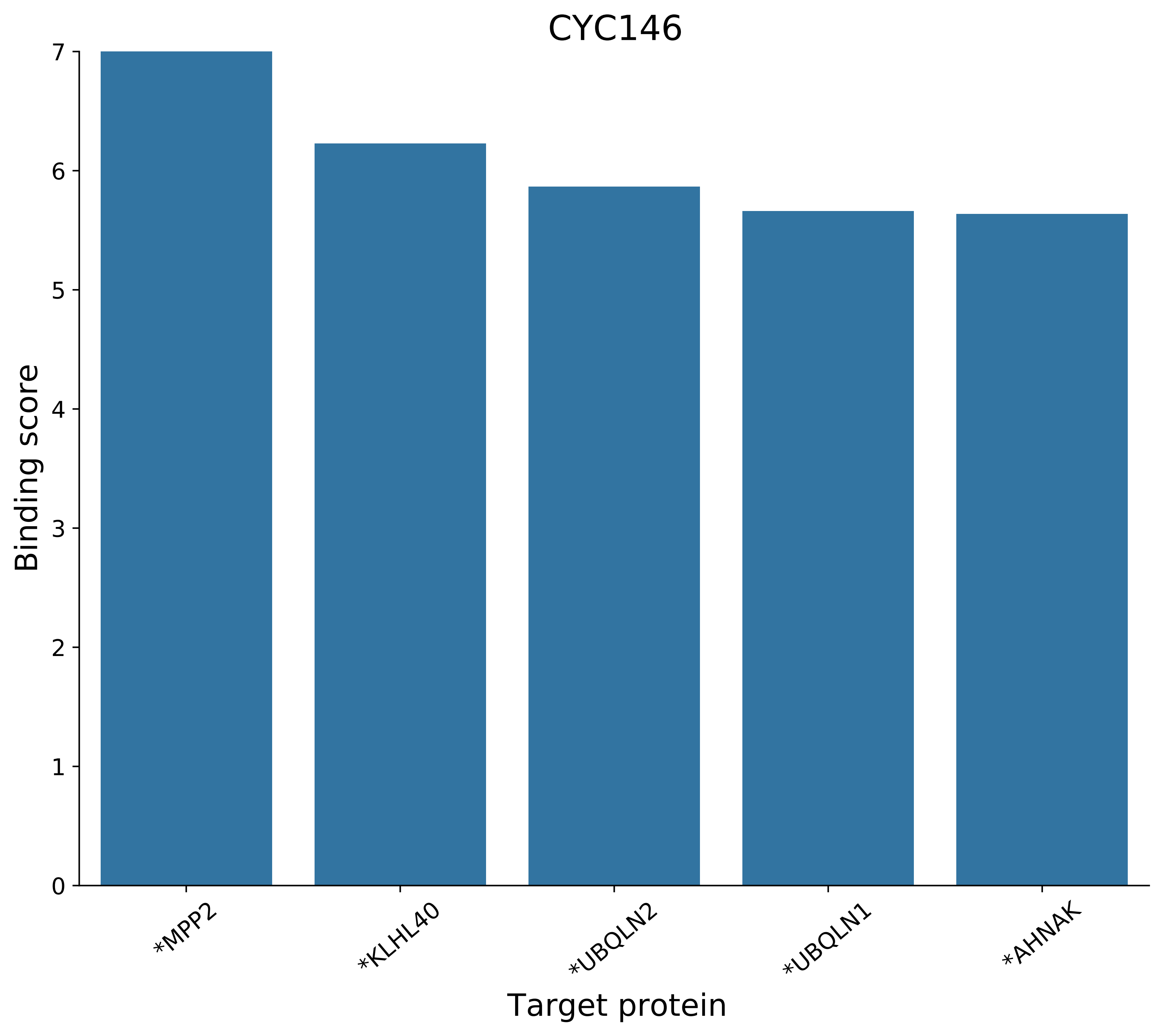
CYC146
Binding score
Target protein
0
1
2
3
4
5
6
7
*MPP2
*KLHL40
*UBQLN2
*UBQLN1
*AHNAK

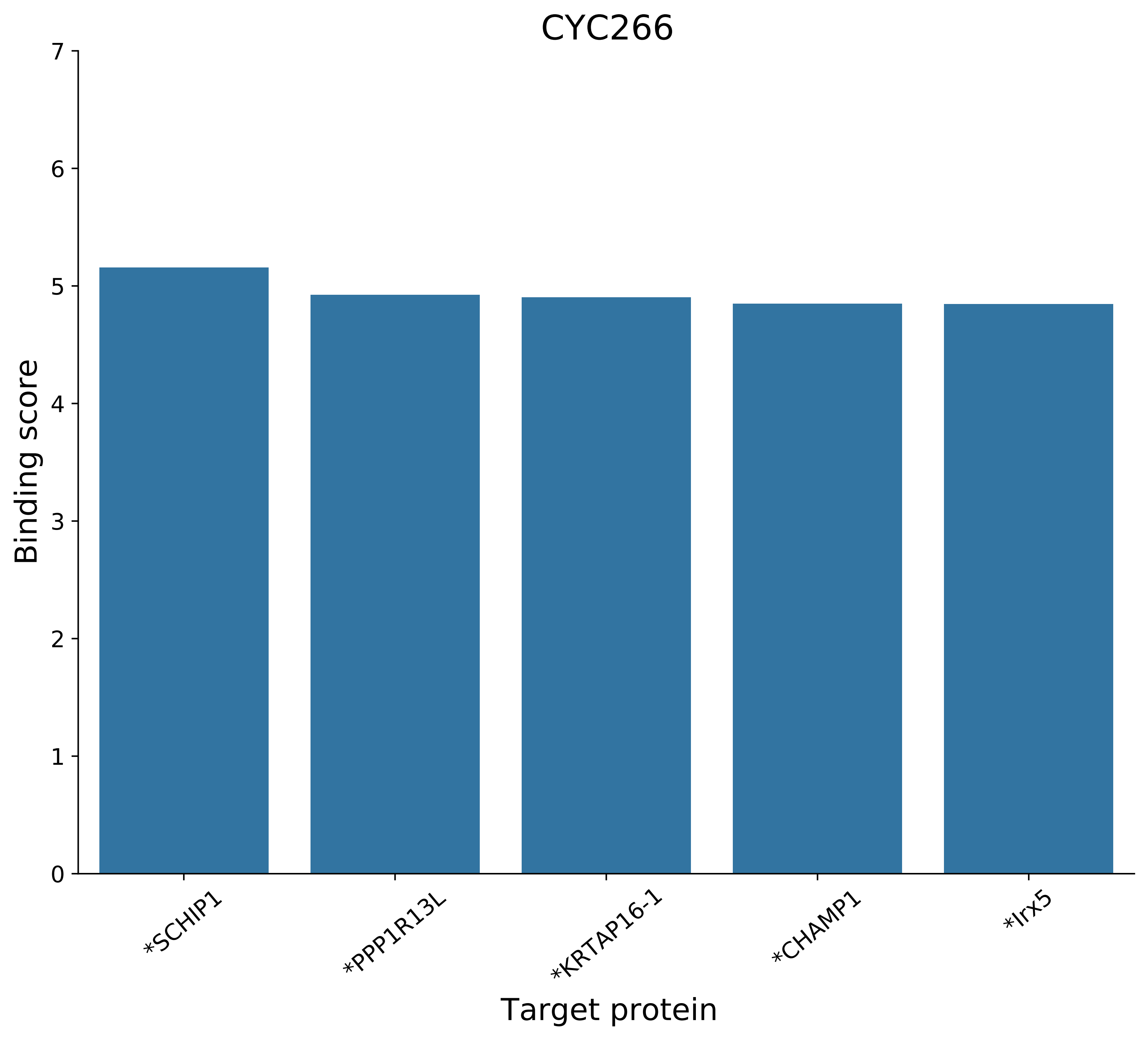
CYC266
Binding score
Target protein
0
1
2
3
4
5
6
7
*SCHIP1
*PPP1R13L
*KRTAP16-1
*CHAMP1
*Irx5

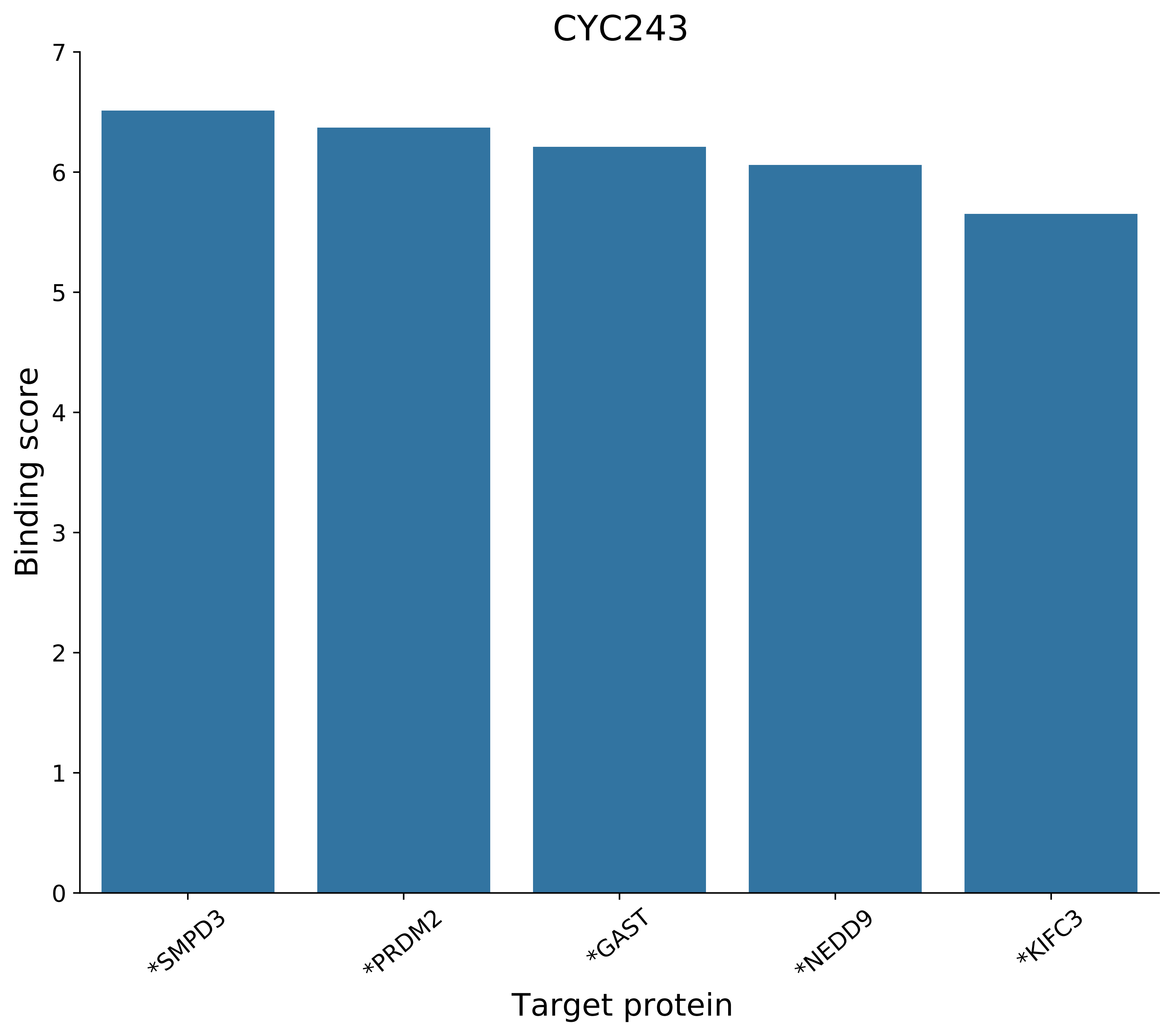
CYC243
Binding score
0
1
2
3
4
5
6
7
*SMPD3
*PRDM2
*GAST
*NEDD9
*KIFC3
Target protein

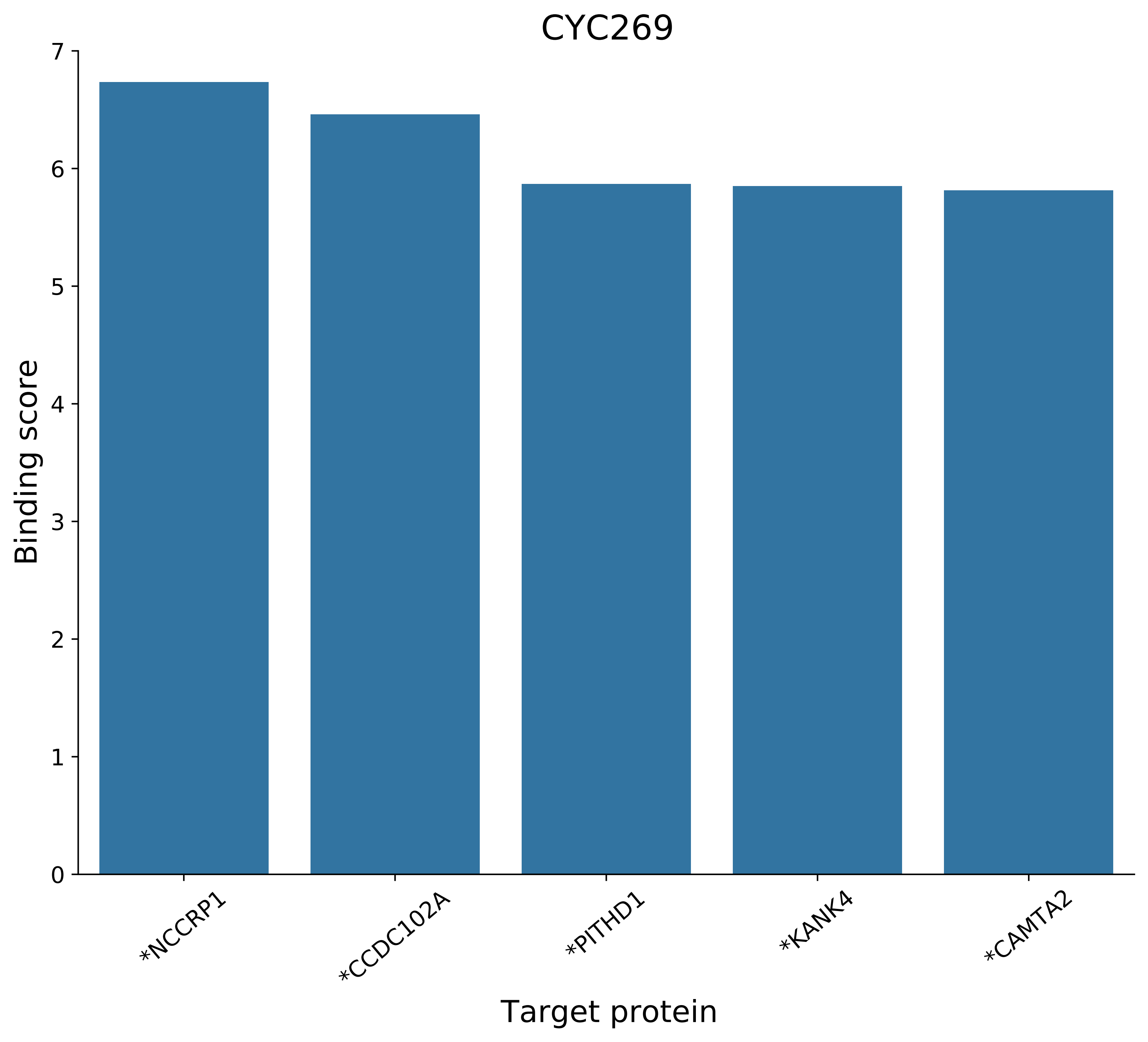
CYC269
Binding score
0
1
2
3
4
5
6
7
*NCCRP1
*CCDC102A
*PITHD1
*KANK4
*CAMTA2
Target protein

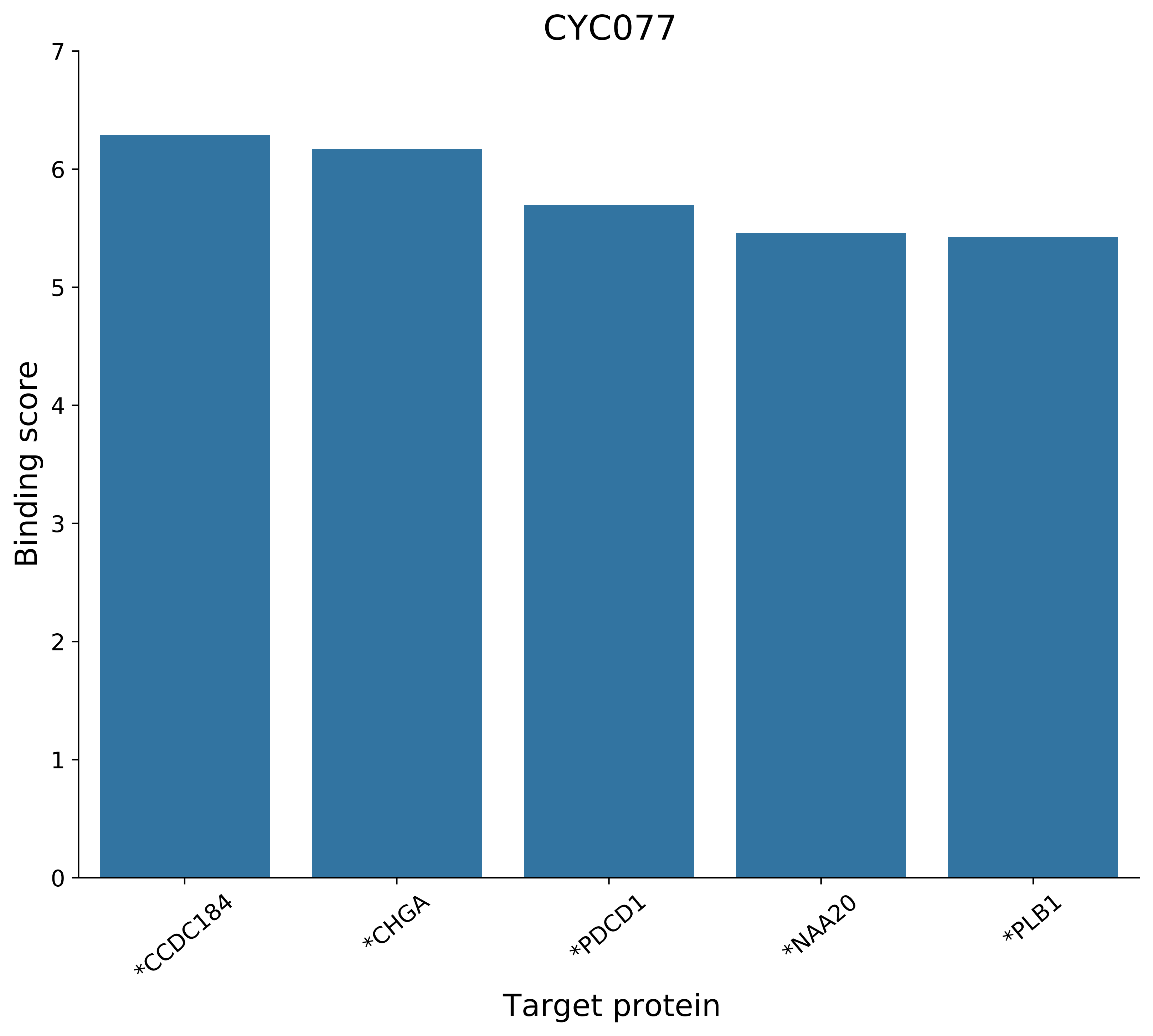
CYC077
Binding score
Target protein
0
1
2
3
4
5
6
7
*CCDC184
*CHGA
*PDCD1
*NAA20
*PLB1

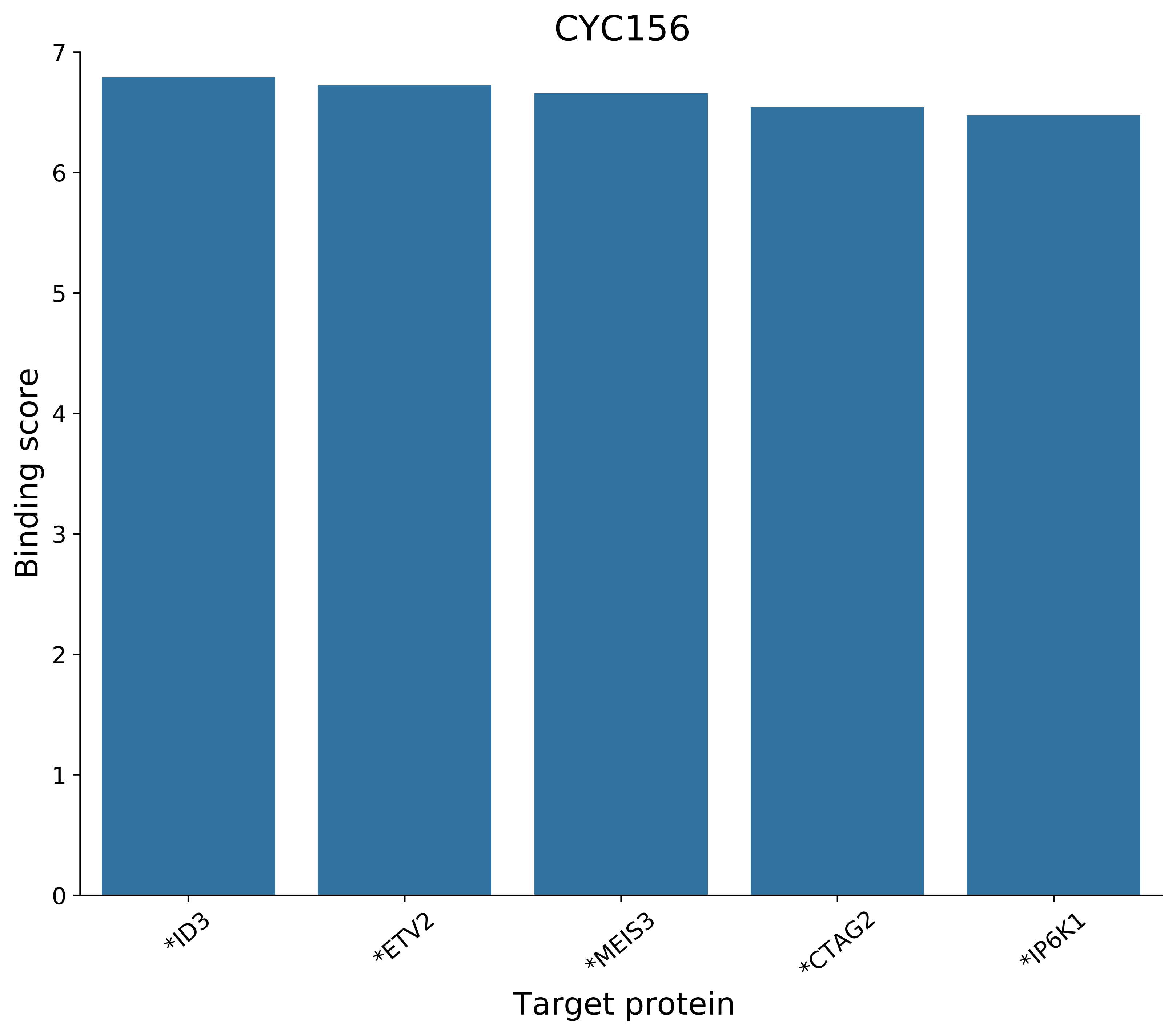

CYC156
Binding score
0
1
2
3
4
5
6
7
*ID3
*ETV2
*MEIS3
*CTAG2
*IP6K1
Target protein

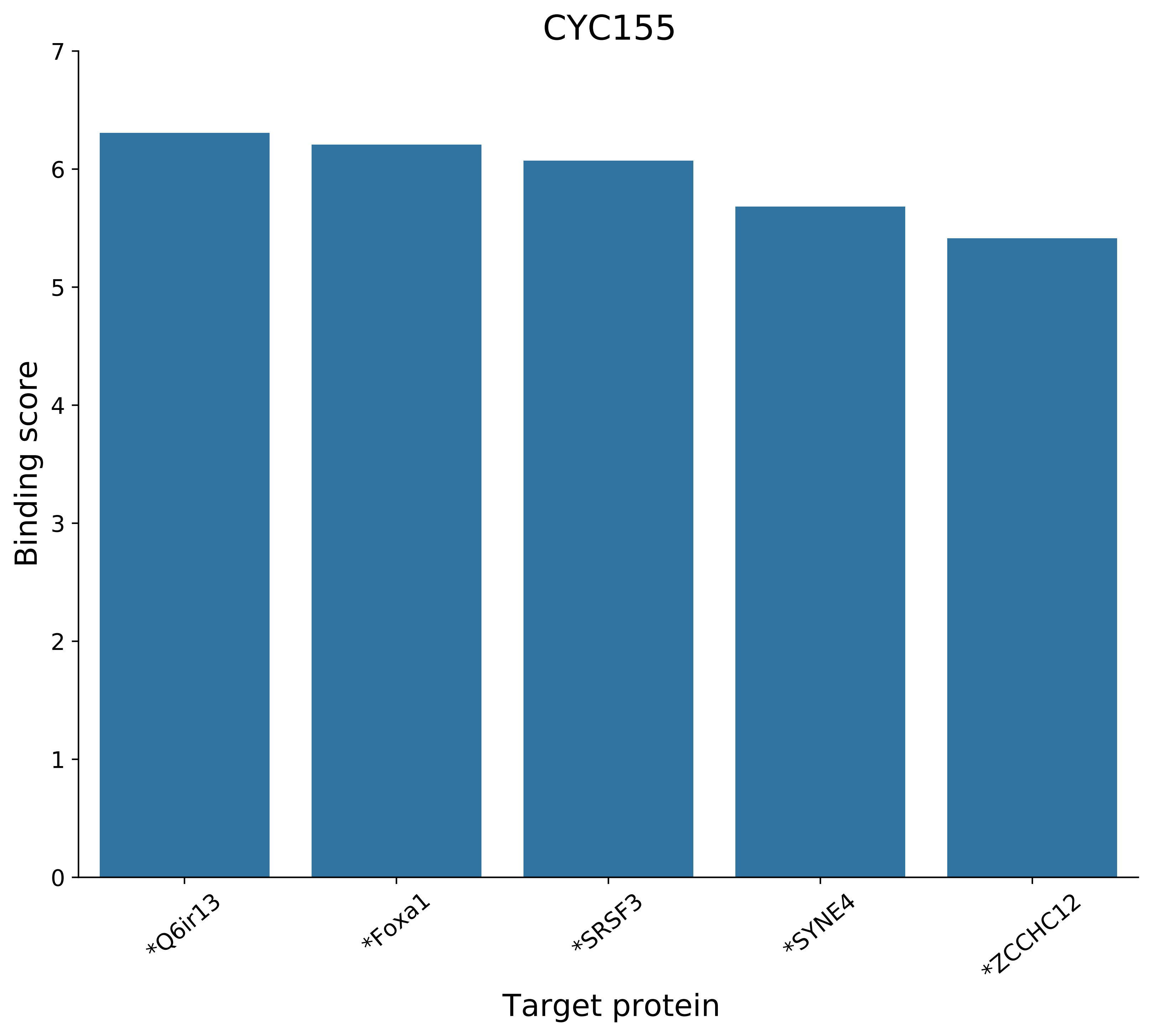

CYC155
Binding score
Target protein
0
1
2
3
4
5
6
7
*Q6ir13
*Foxa1
*SRSF3
*SYNE4
*ZCCHC12

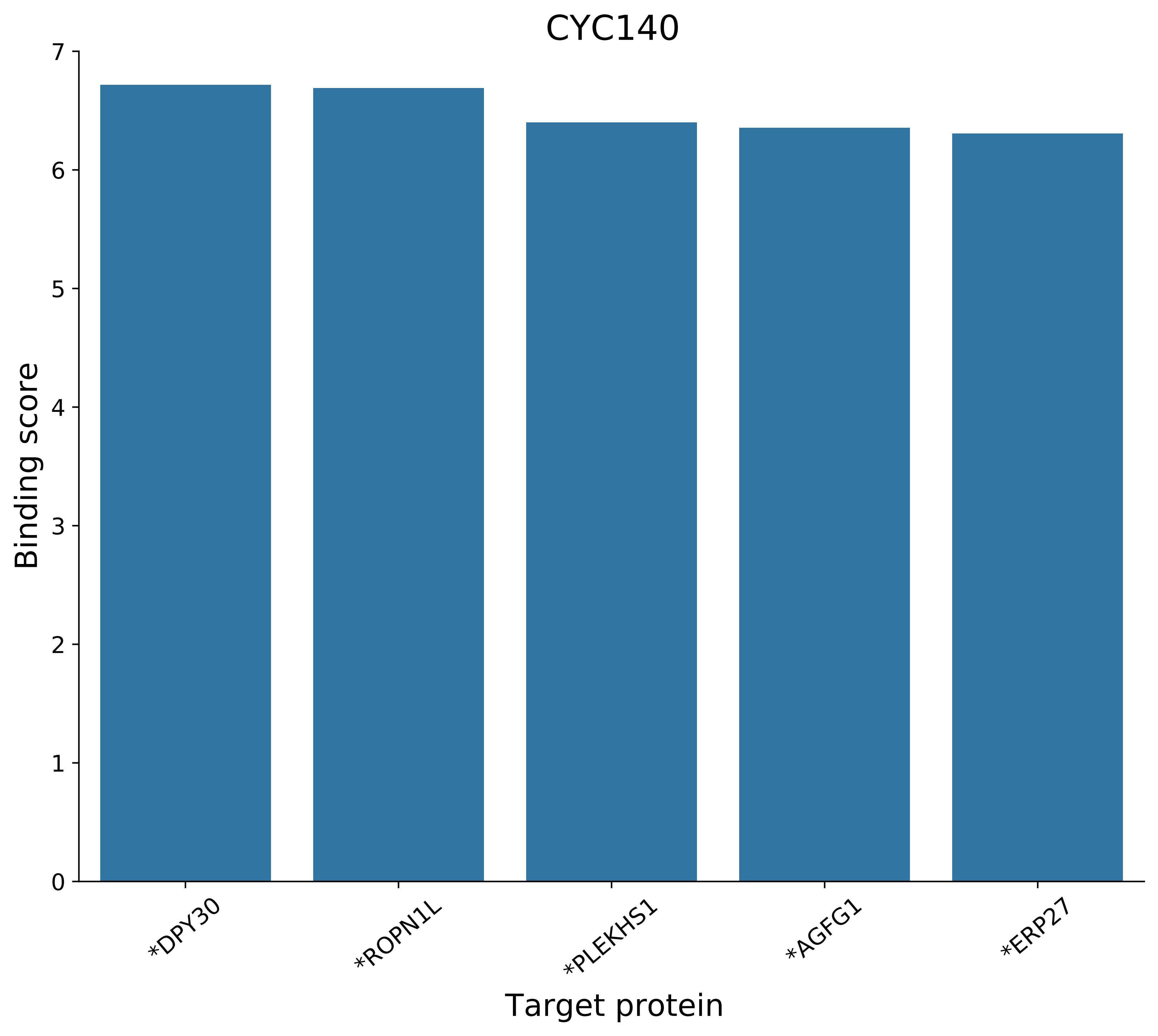

CYC140
Binding score
Target protein
0
1
2
3
4
5
6
7
*DPY30
*ROPN1L
*PLEKHS1
*AGFG1
*ERP27

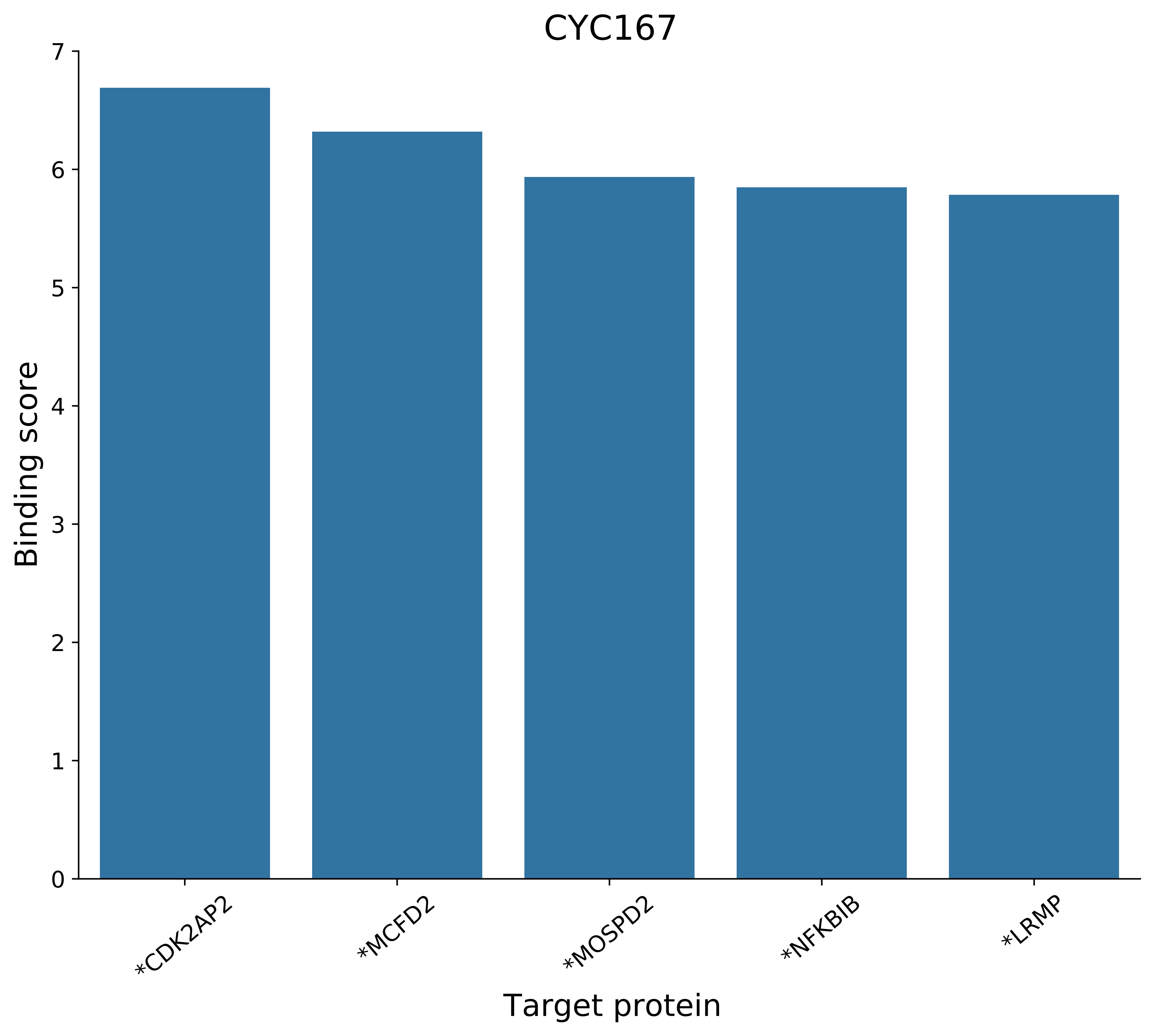
CYC167
Binding score
Target protein
0
1
2
3
4
5
6
7
*CDK2AP2
*MCFD2
*MOSPD2
*NFKBIB
*LRMP

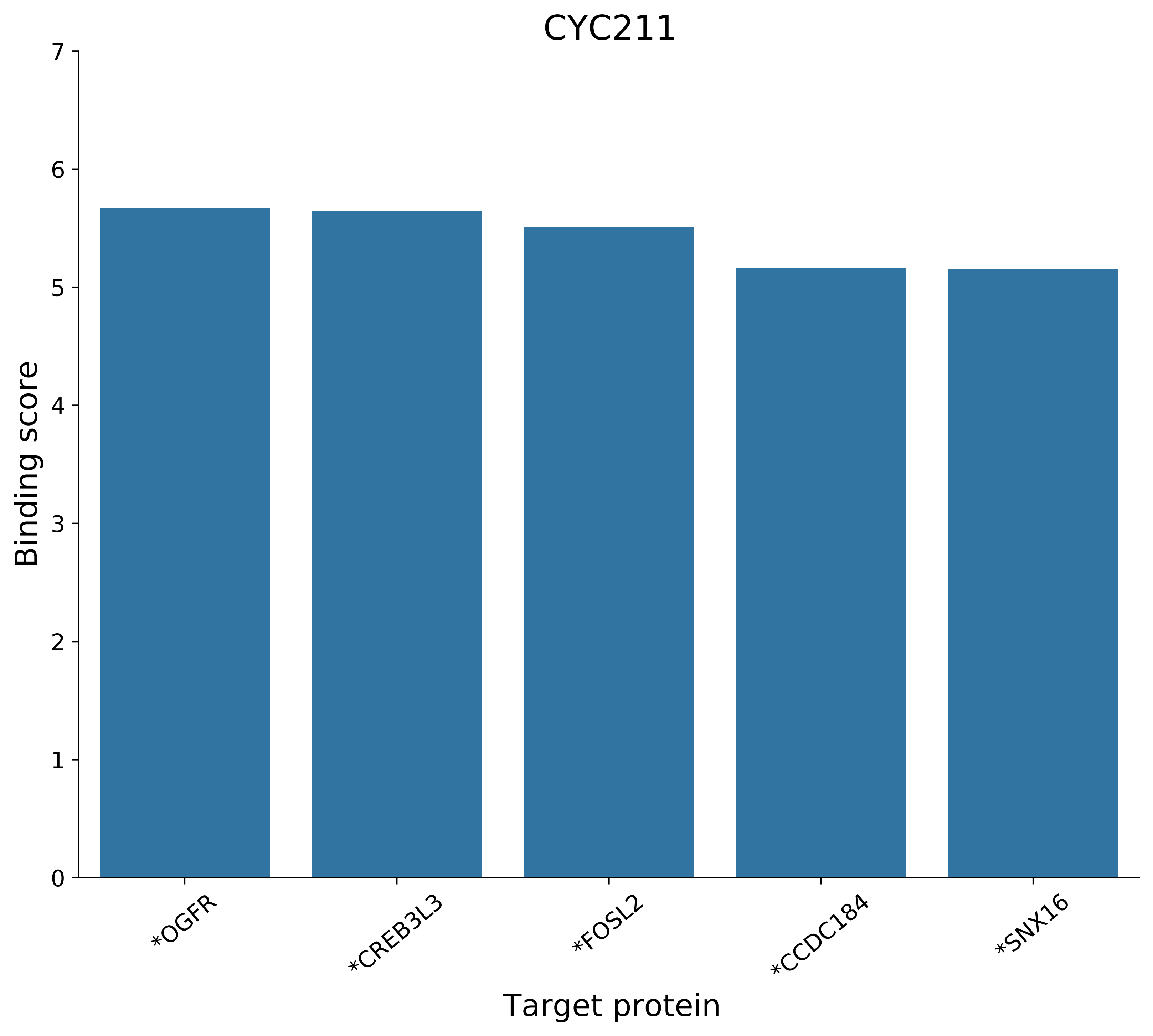
CYC211
Binding score
0
1
2
3
4
5
6
7
*OGFR
*CREB3L3
*FOSL2
*CCDC184
*SNX16
Target protein

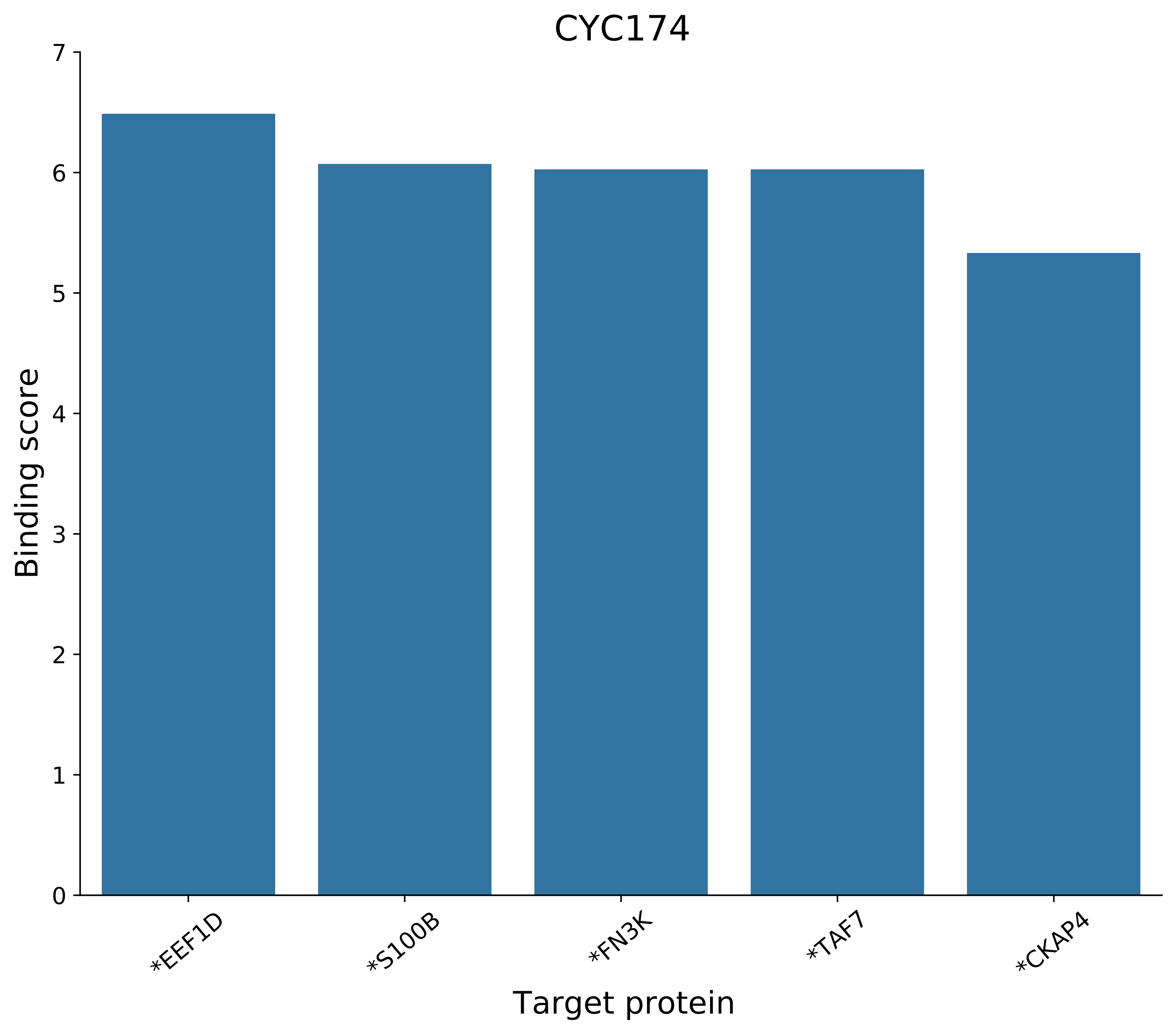

CYC174
Binding score
0
1
2
3
4
5
6
7
*EEF1D
*S100B
*FN3K
*TAF7
*CKAP4
Target protein

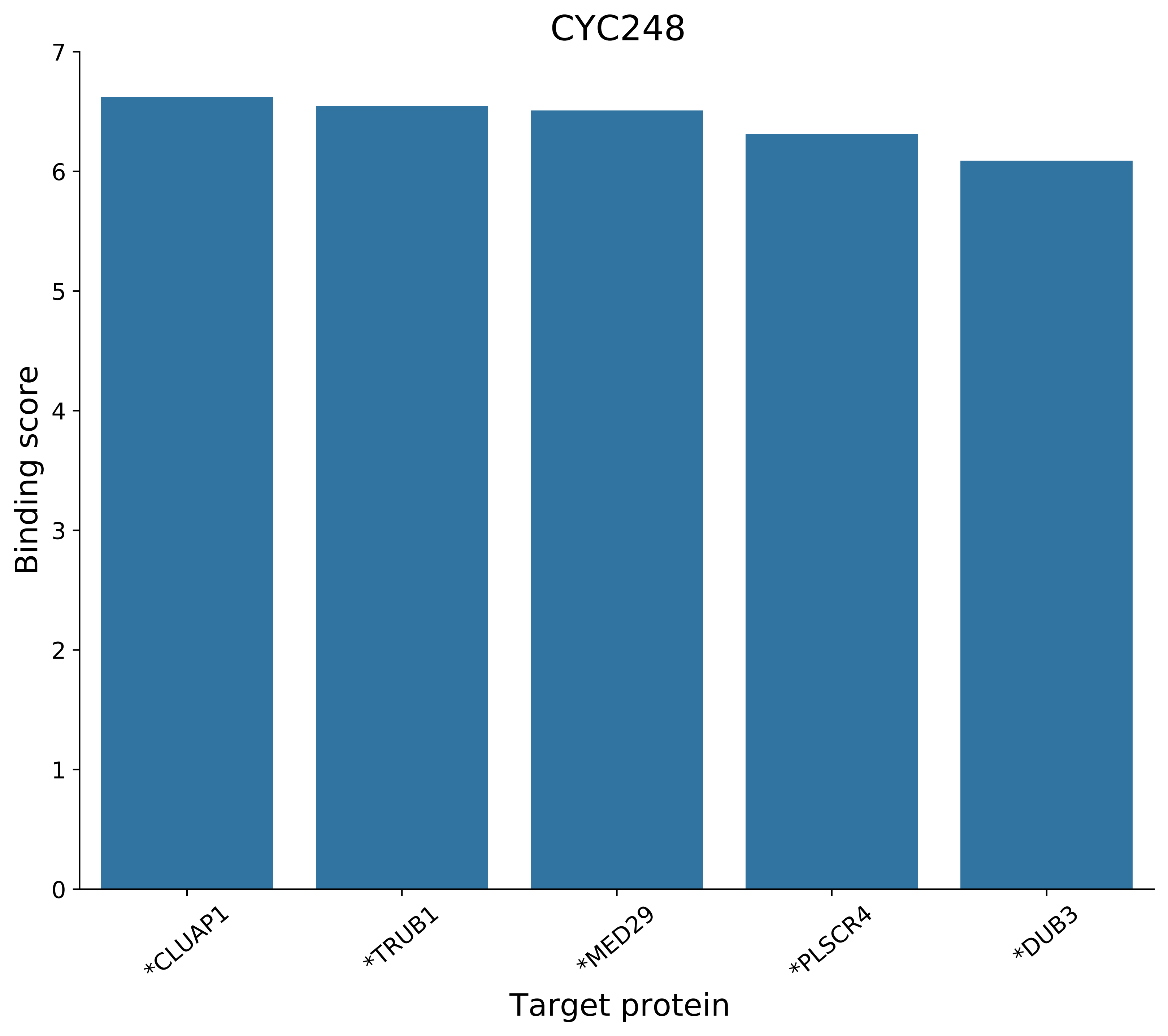
CYC248
Binding score
0
1
2
3
4
5
6
7
*CLUAP1
*TRUB1
*MED29
*PLSCR4
*DUB3
Target protein

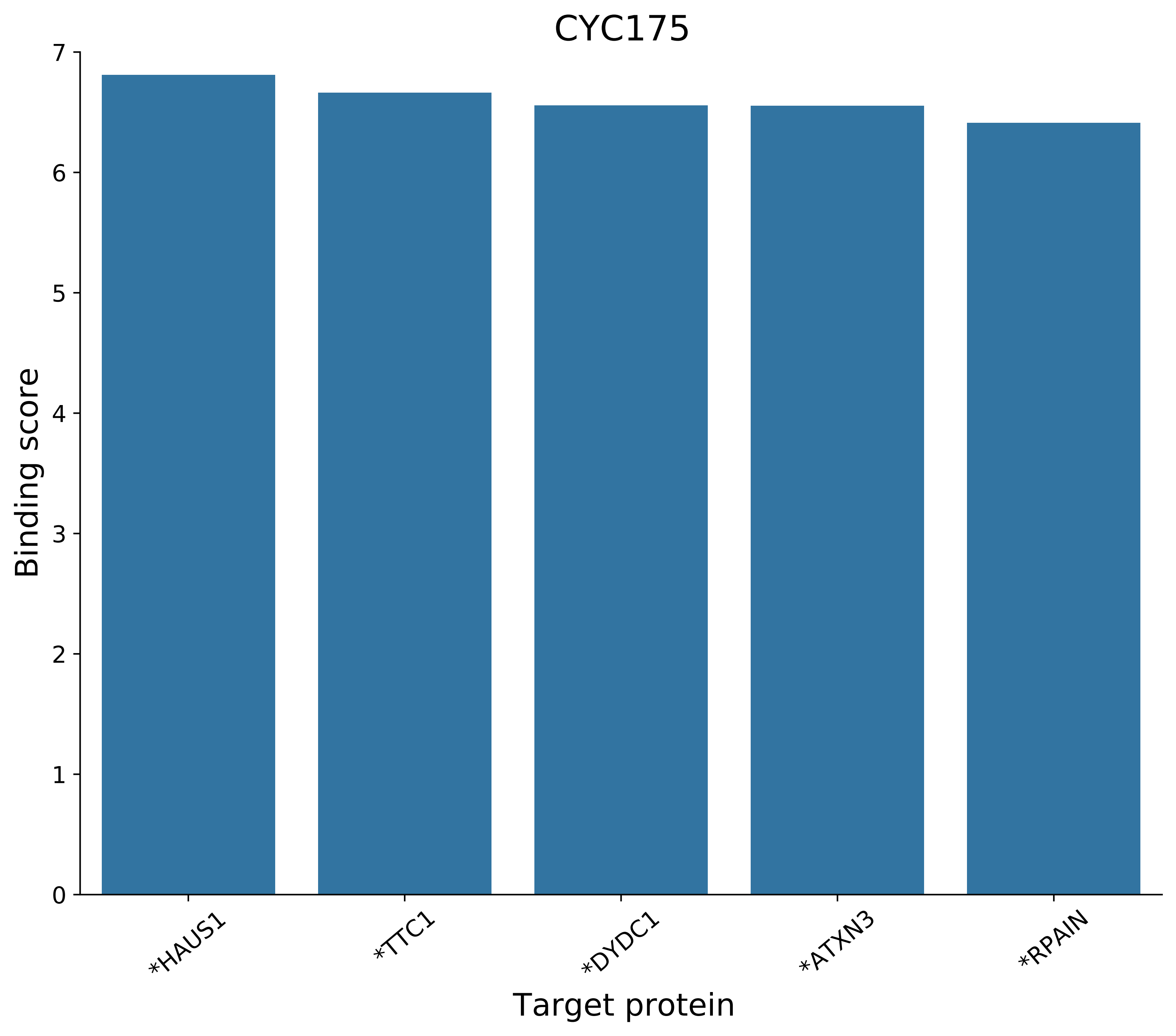
CYC175
Binding score
Target protein
0
1
2
3
4
5
6
7
*HAUS1
*TTC1
*DYDC1
*ATXN3
*RPAIN

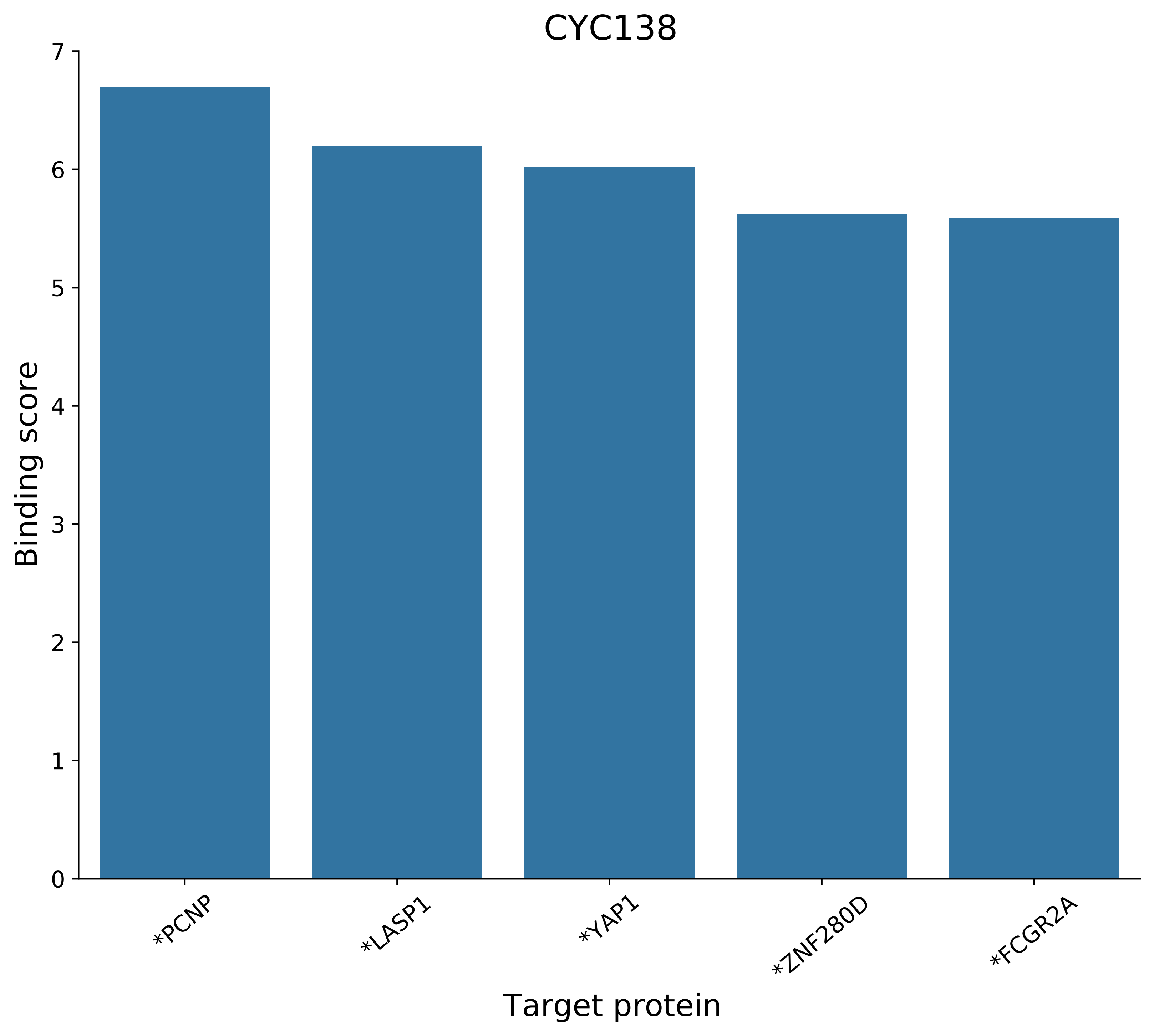

CYC138
Binding score
Target protein
7
6
5
4
3
2
1
0
*PCNP
*LASP1
*YAP1
*ZNF280D
*FCGR2A

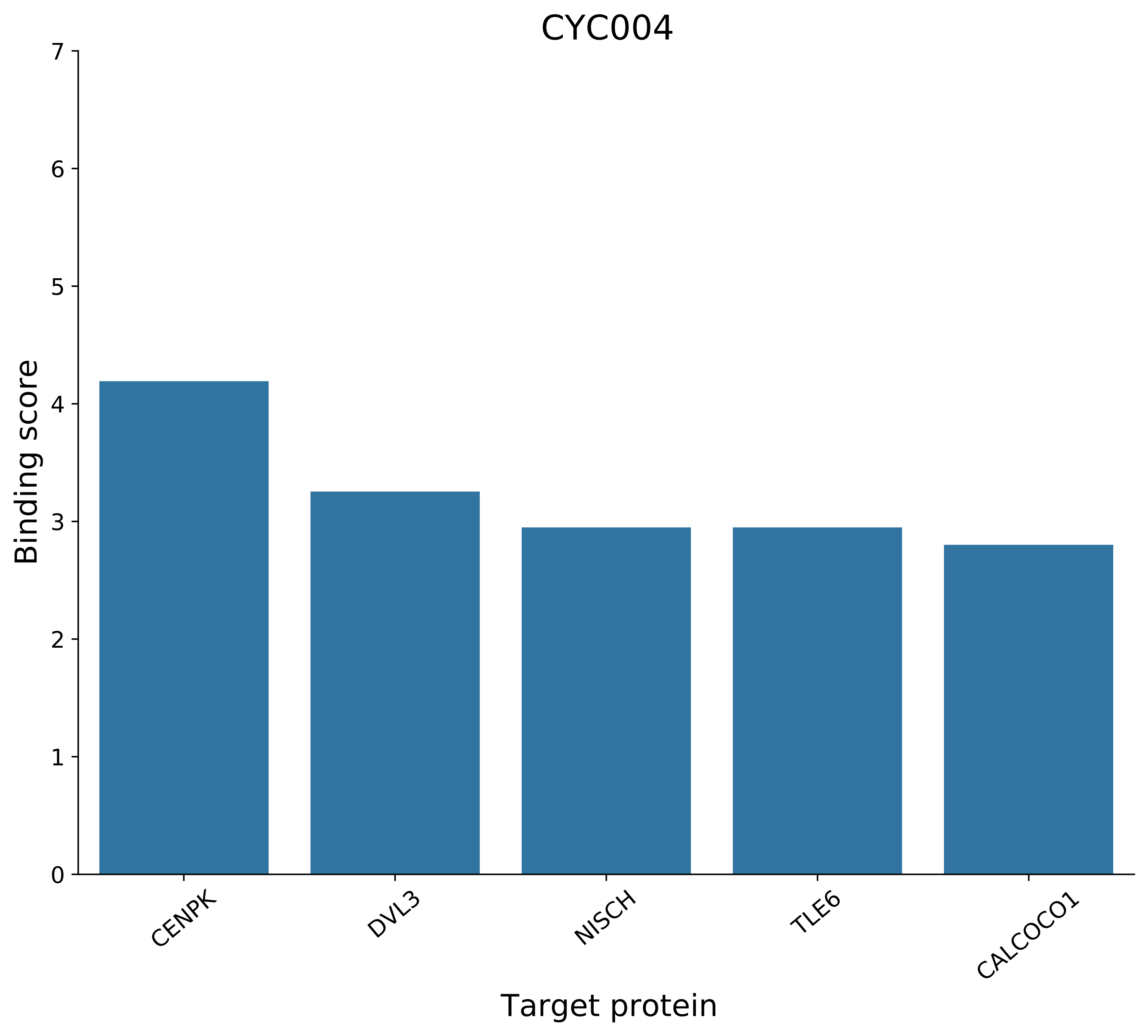

CYC004
Binding score
Target protein
0
1
2
3
4
5
6
7
CENPK
DVL3
NISCH
TLE6
CALCOCO1

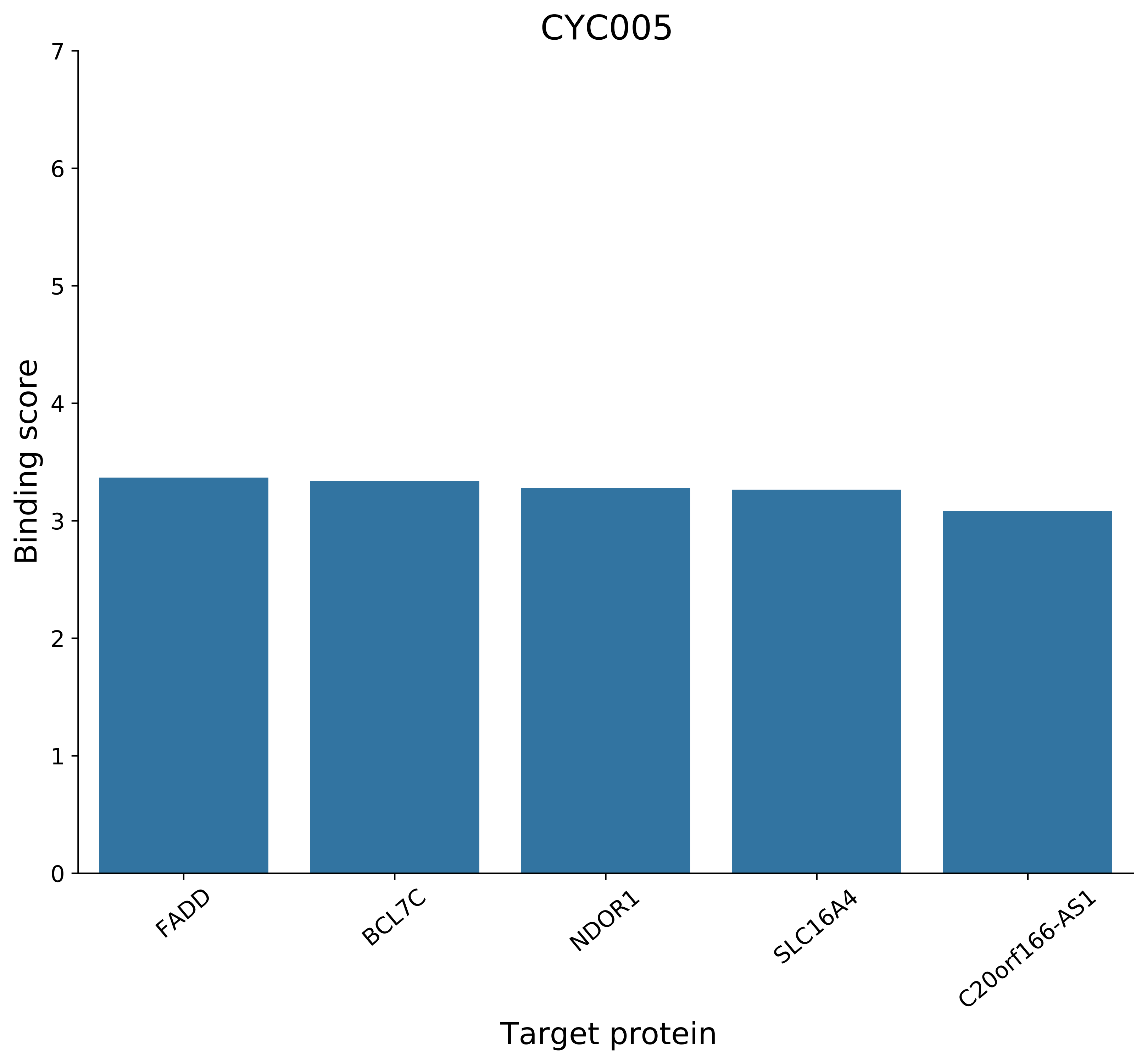
CYC005
Binding score
Target protein
0
1
2
3
4
5
6
7
FADD
BCL7C
NDOR1
SLC16A4
C20orf166-AS1

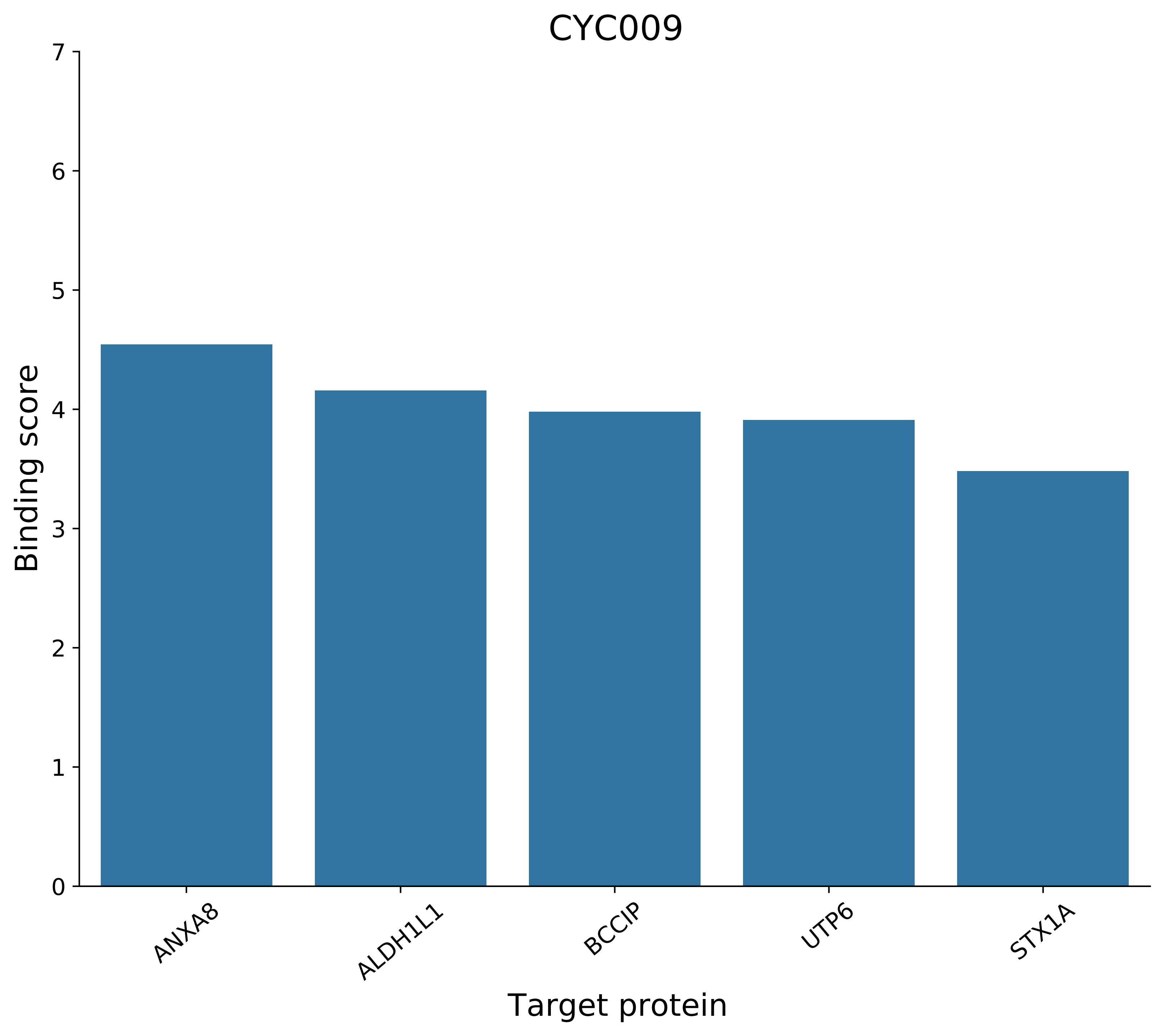
CYC009
Binding score
Target protein
0
1
2
3
4
5
6
7
ANXA8
ALDH1L1
BCCIP
UTP6
STX1A

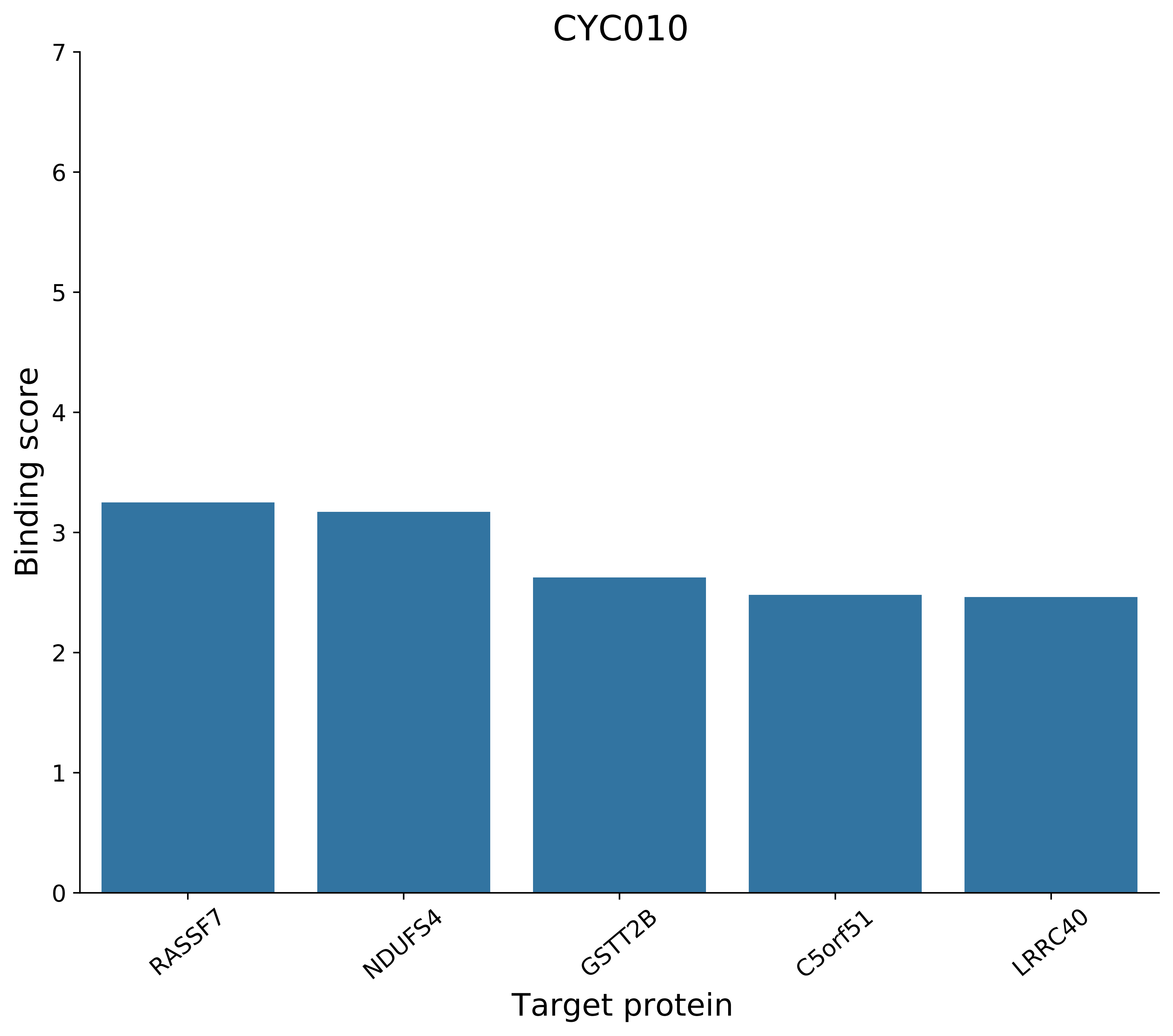

CYC010
Binding score
0
1
2
3
4
5
6
7
RASSF7
NDUFS4
GSTT2B
C5orf51
LRRC40
Target protein

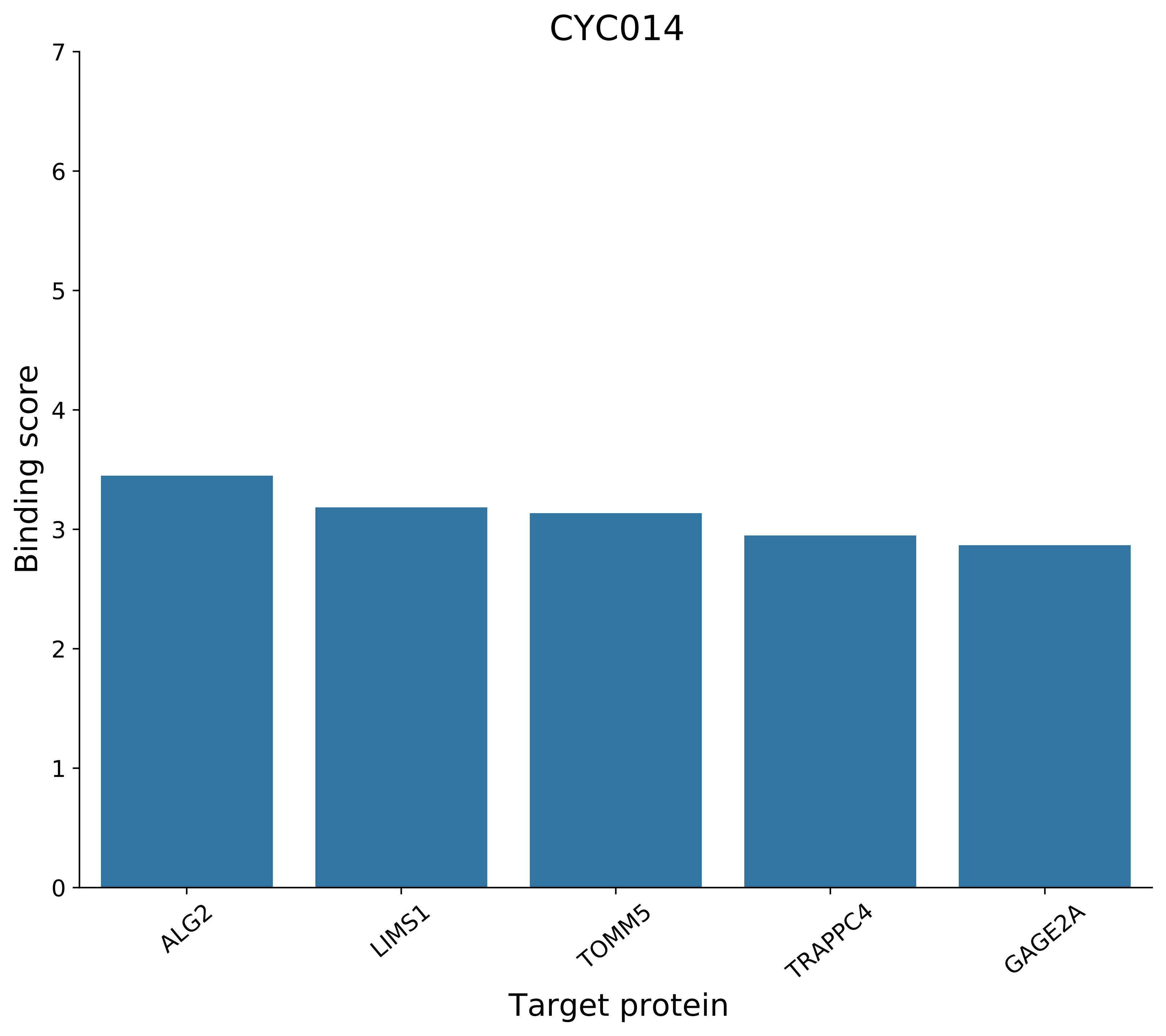
CYC014
Binding score
0
1
2
3
4
5
6
7
ALG2
LIMS1
TOMM5
TRAPPC4
GAGE2A
Target protein

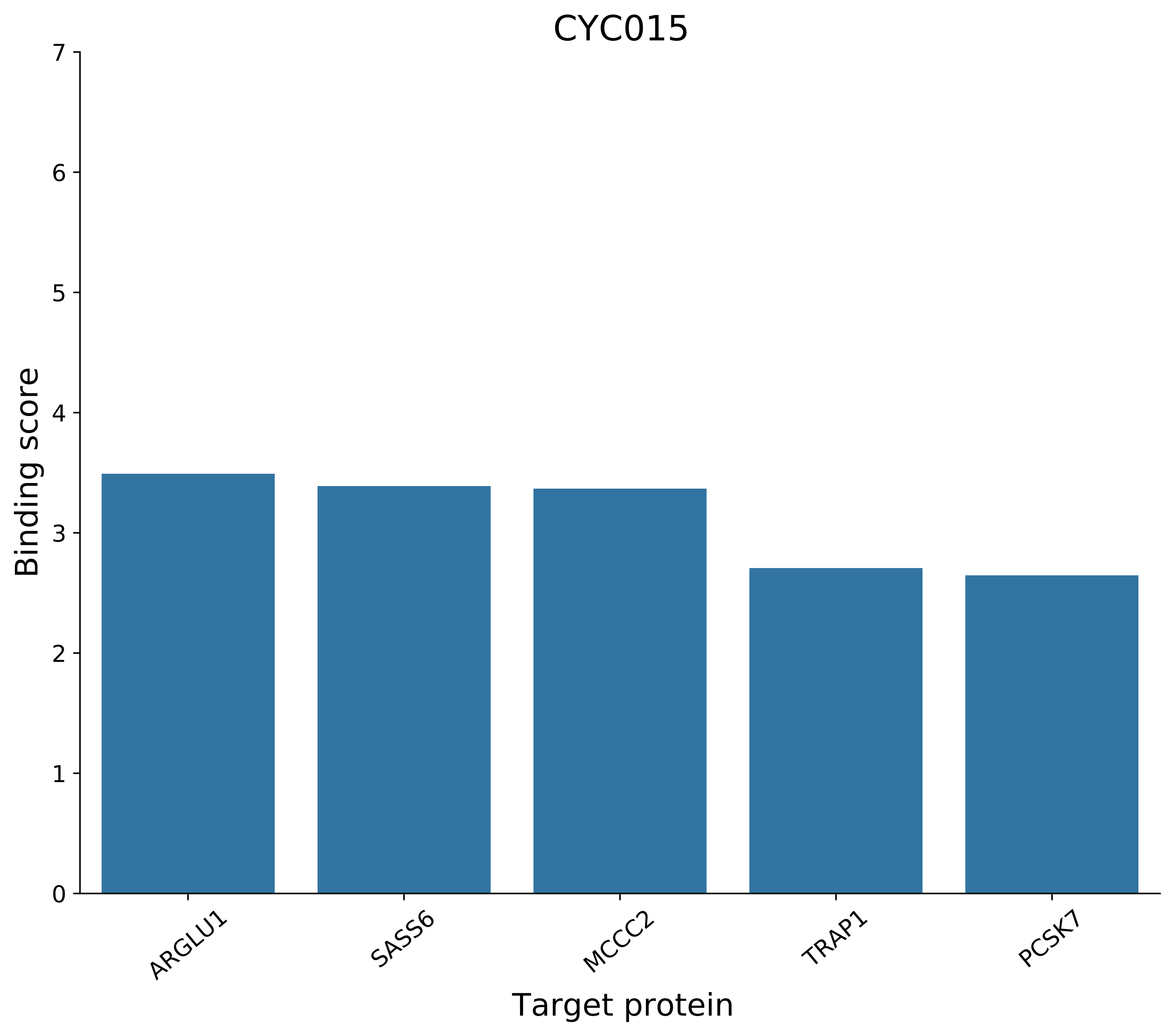

CYC015
Binding score
Target protein
0
1
2
3
4
5
6
7
ARGLU1
SASS6
MCCC2
TRAP1
PCSK7

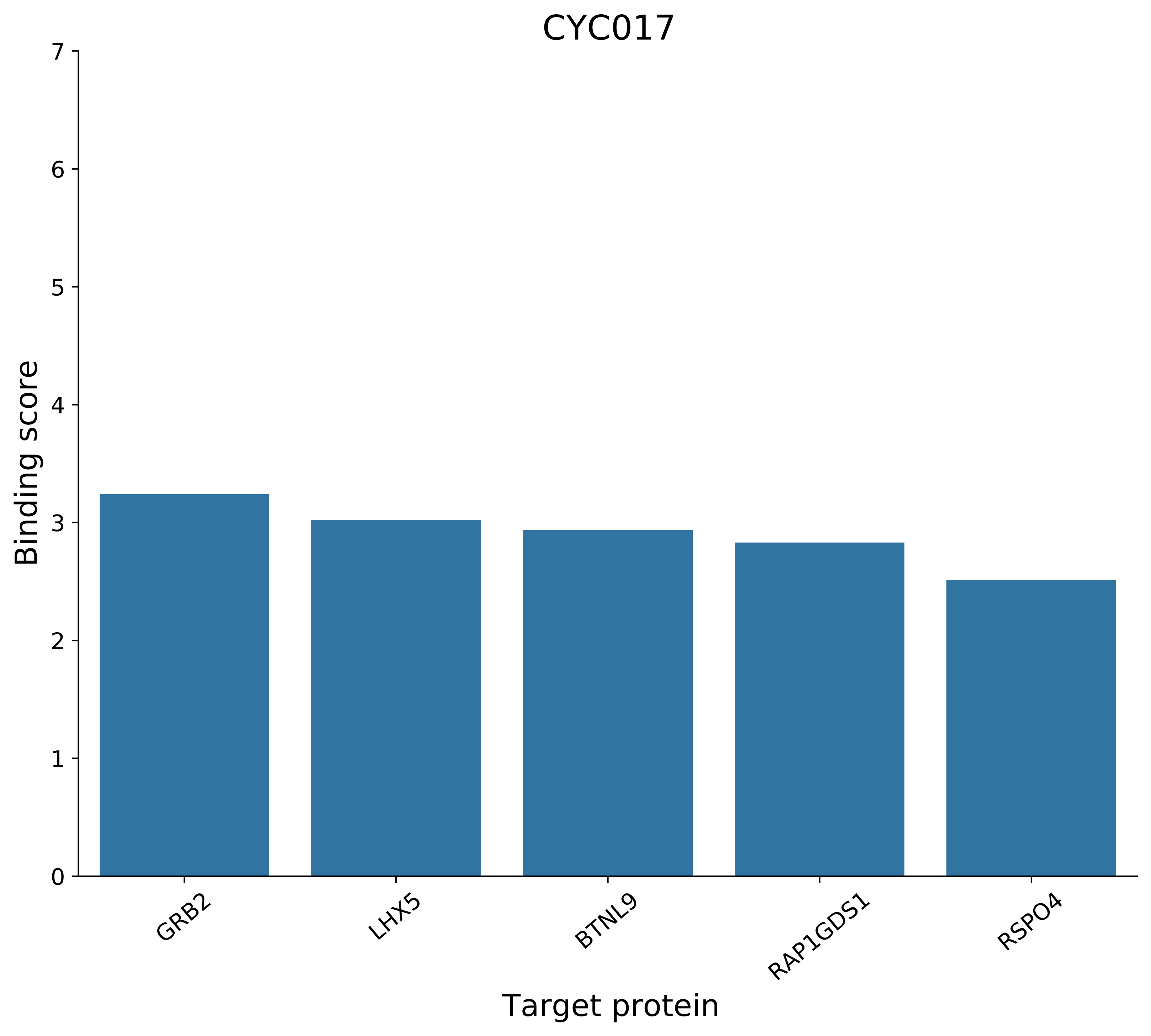
CYC017
Binding score
Target protein
0
1
2
3
4
5
6
7
GRB2
LHX5
BTNL9
RAP1GDS1
RSPO4

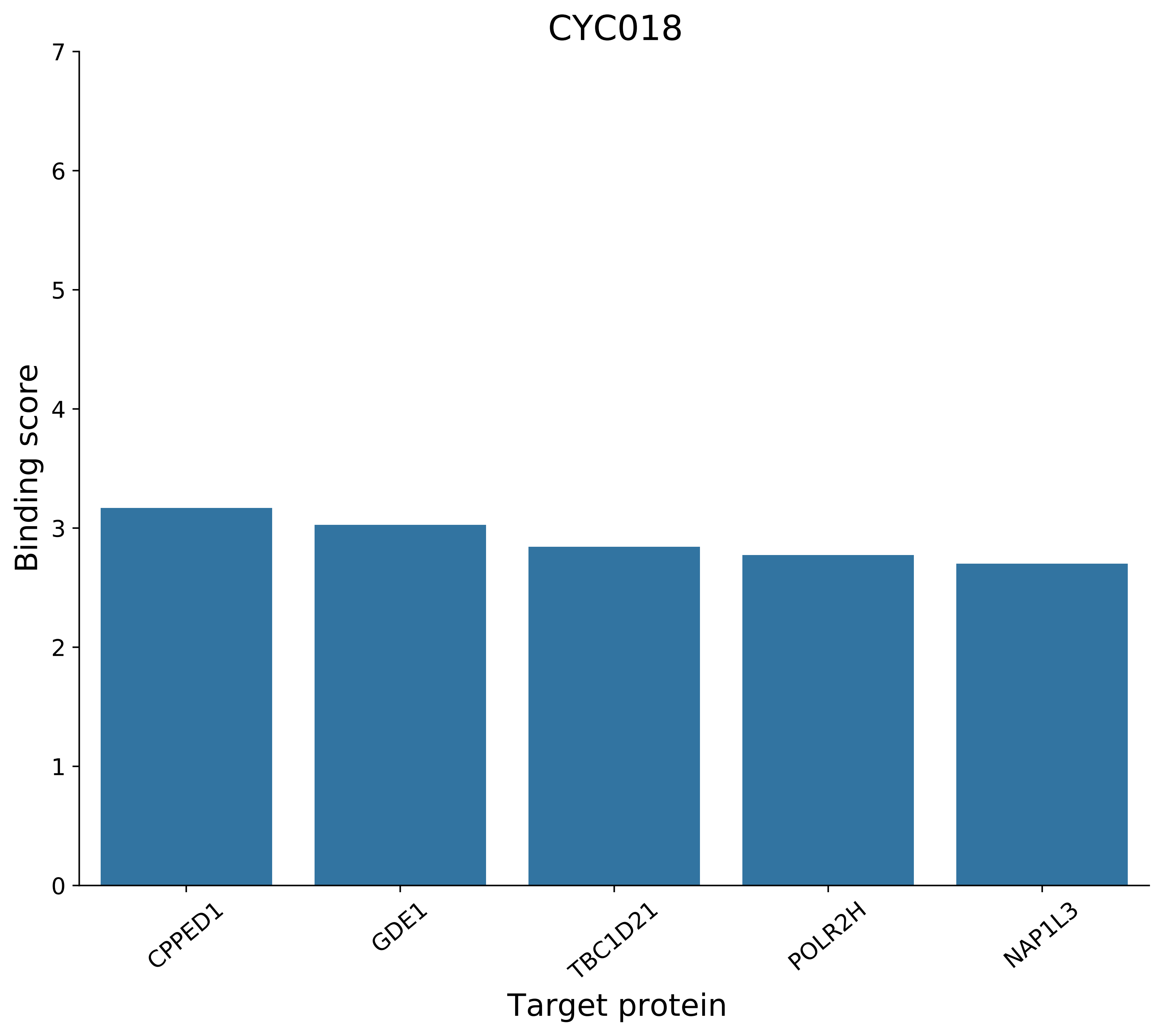
CYC018
Binding score
Target protein
0
1
2
3
4
5
6
7
CPPED1
GDE1
TBC1D21
POLR2H
NAP1L3

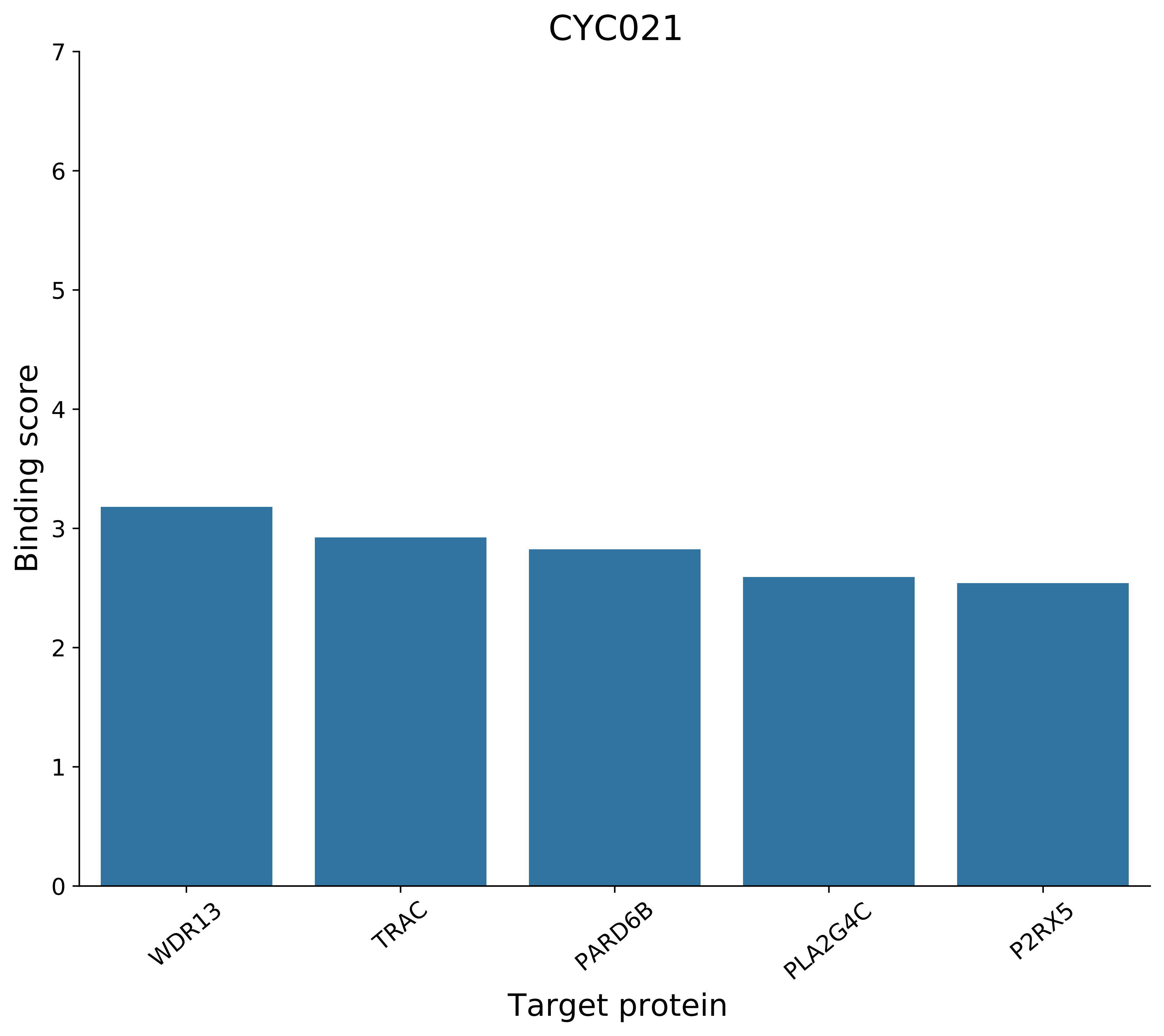

CYC021
Binding score
Target protein
0
1
2
3
4
5
6
7
WDR13
TRAC
PARD6B
PLA2G4C
P2RX5

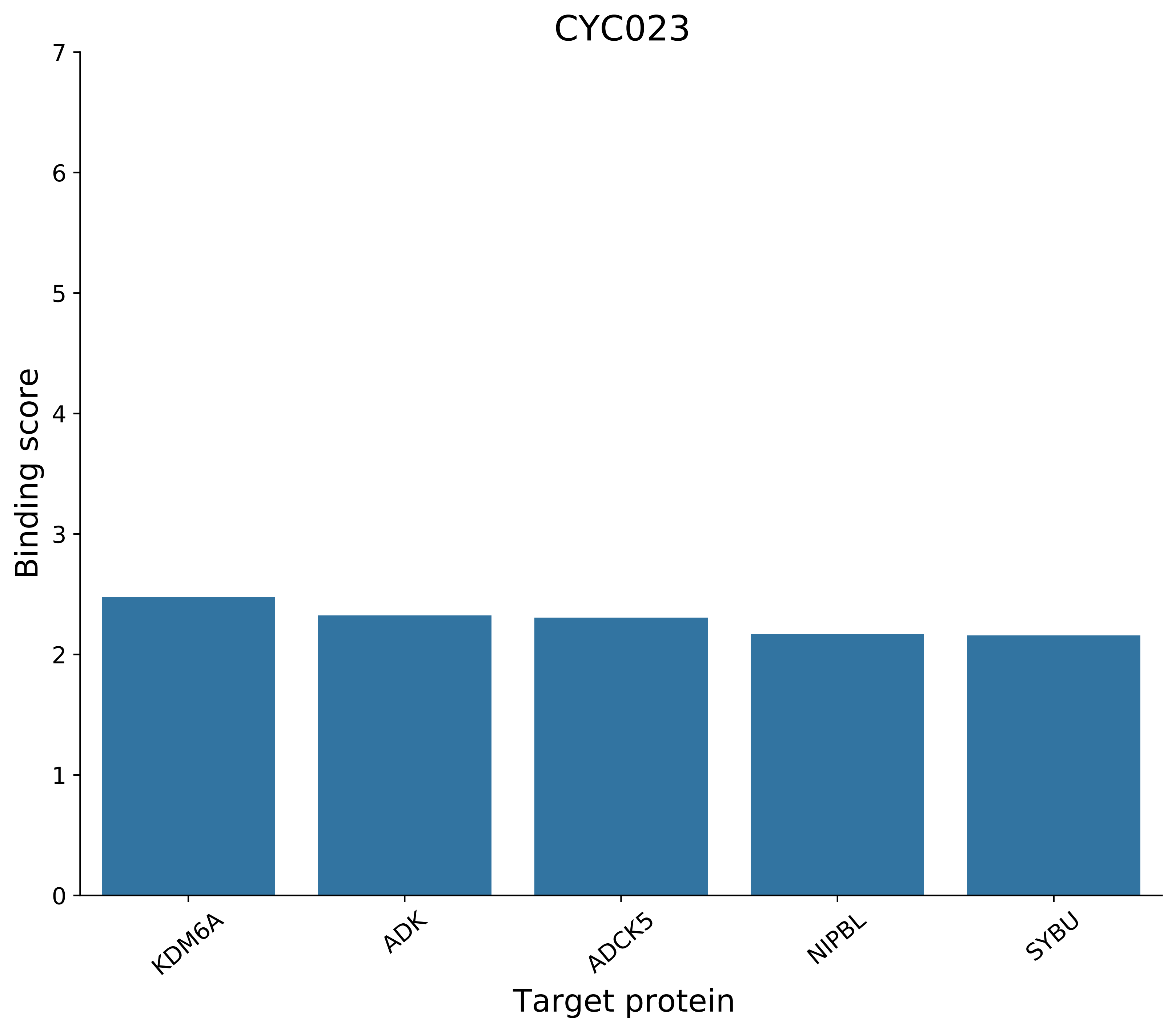
CYC023
Binding score
7
6
5
4
3
2
1
0
KDM6A
ADK
ADCK5
NIPBL
SYBU
Target protein

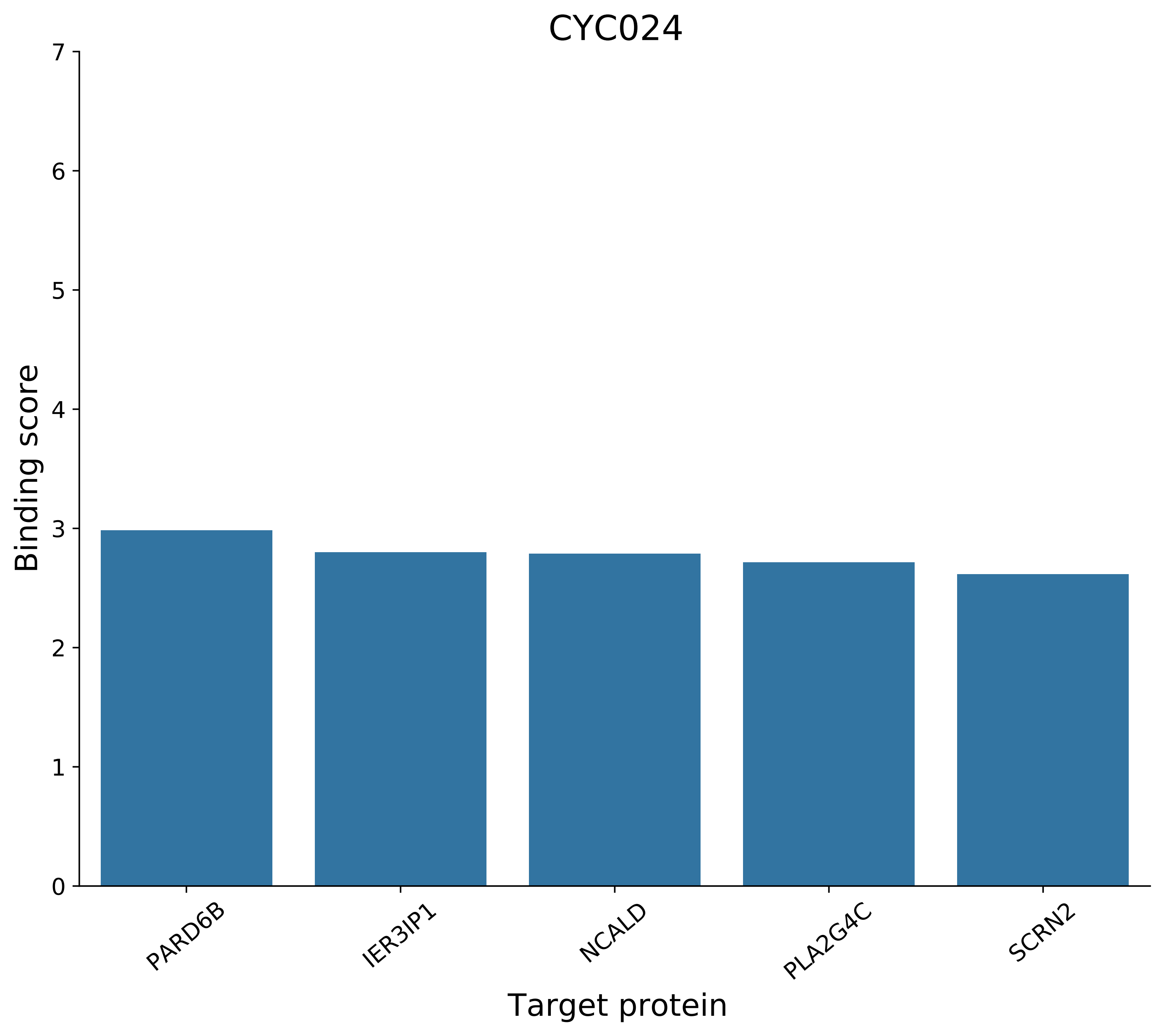
CYC024
Binding score
0
1
2
3
4
5
6
7
PARD6B
IER3IP1
NCALD
PLA2G4C
SCRN2
Target protein

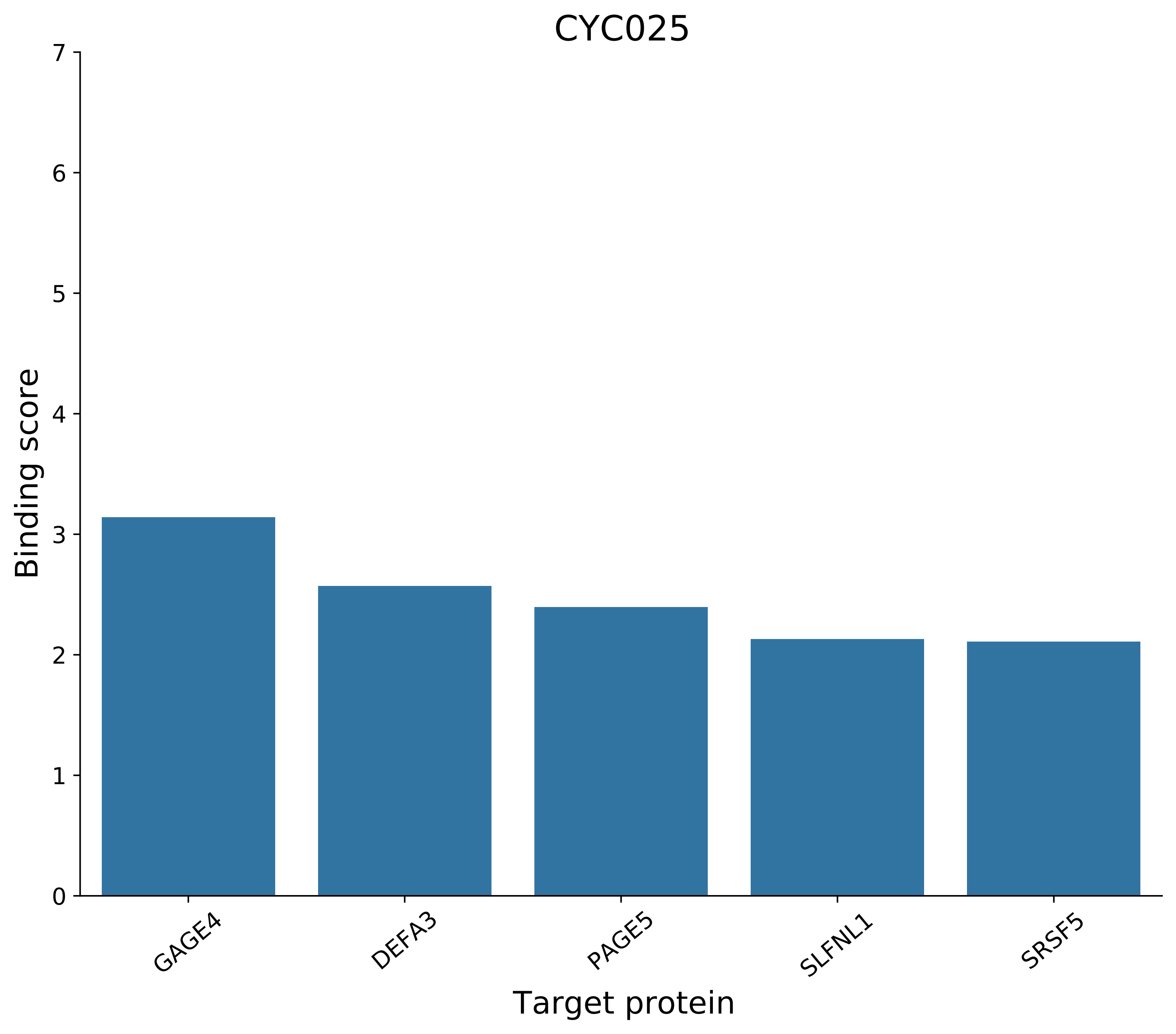

CYC025
Binding score
Target protein
0
1
2
3
4
5
6
7
GAGE4
DEFA3
PAGE5
SLFNL1
SRSF5

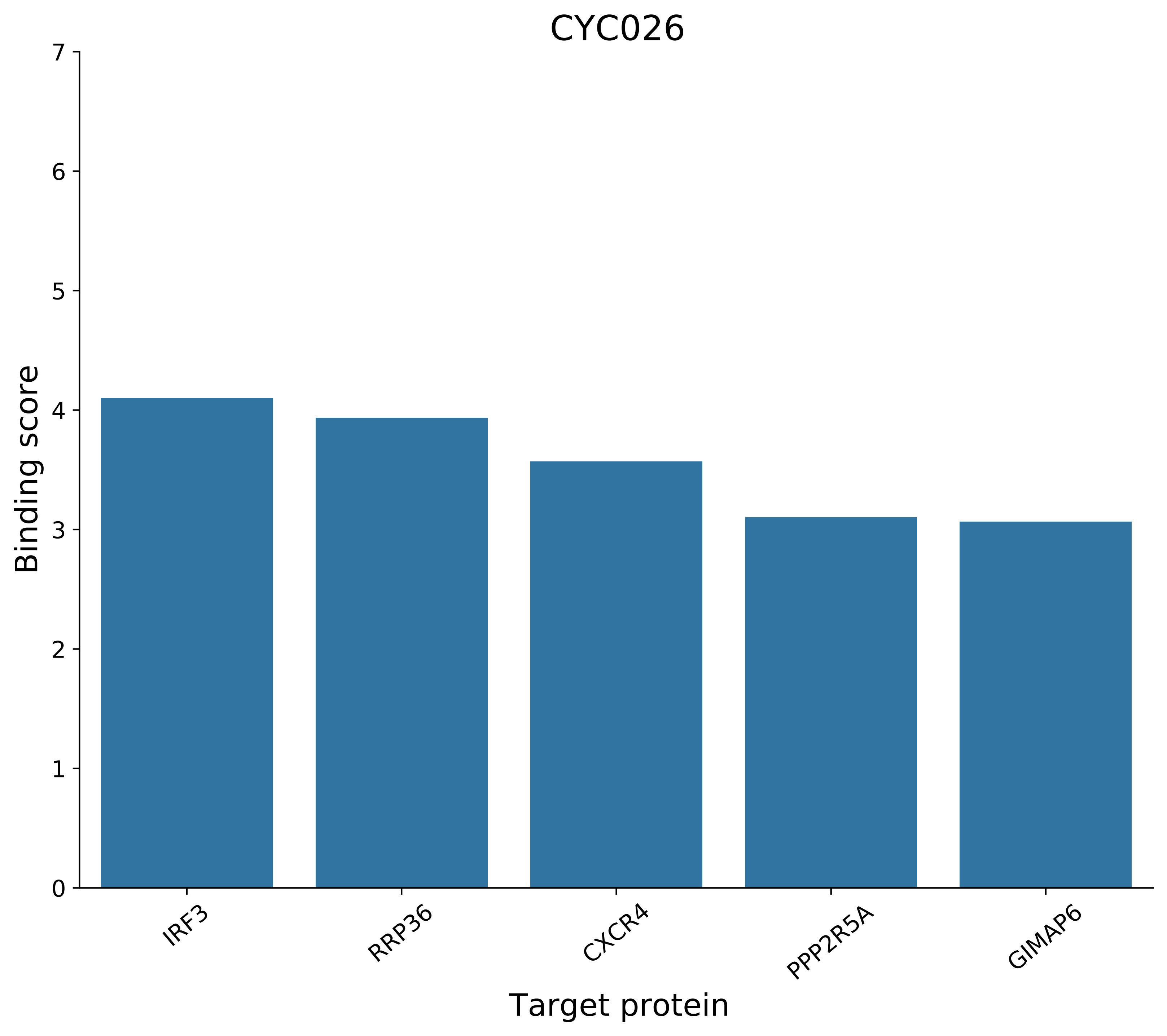
CYC026
Binding score
0
1
2
3
4
5
6
7
IRF3
RRP36
CXCR4
PPP2R5A
GIMAP6
Target protein

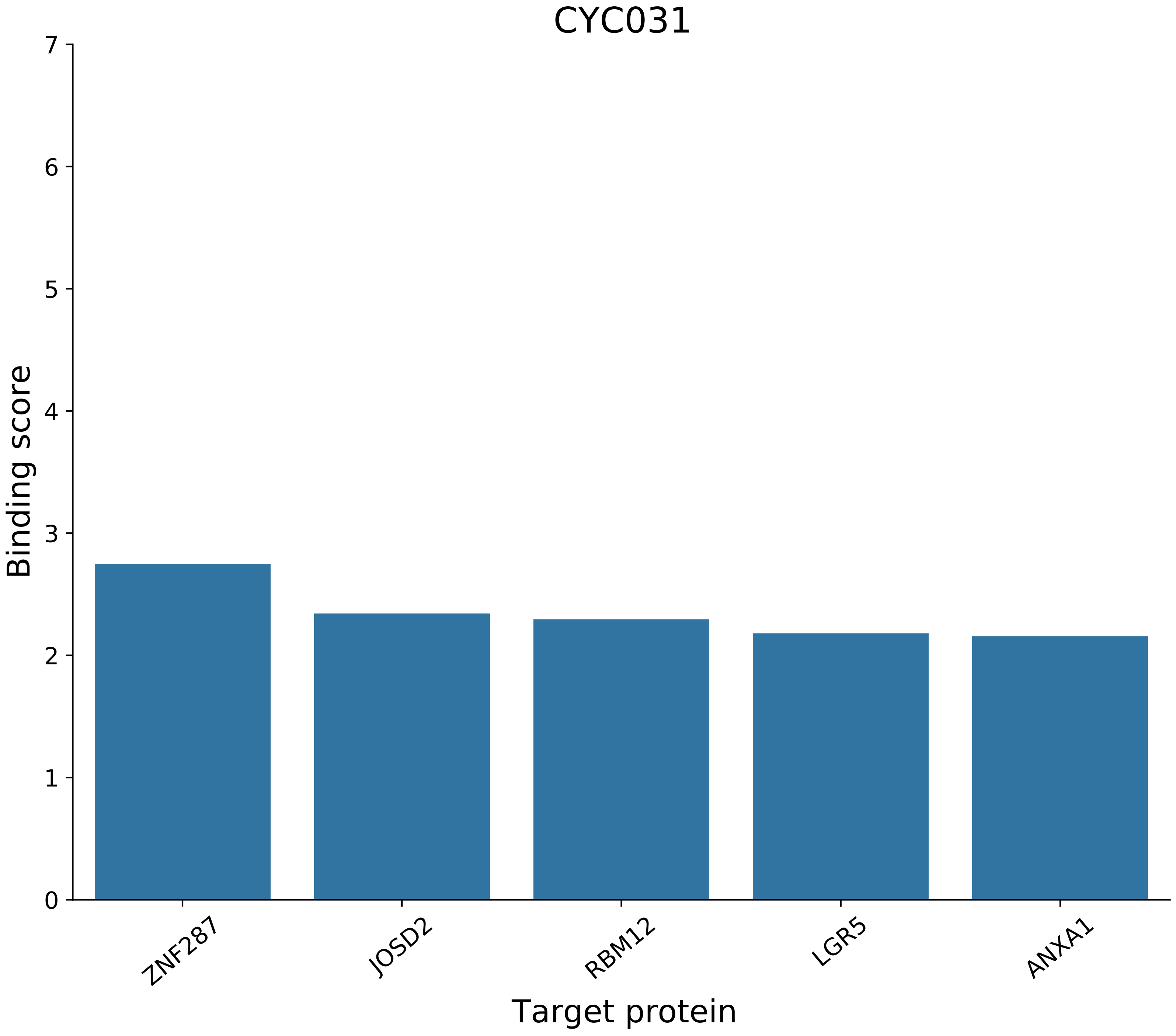
CYC031
Binding score
7
6
5
4
3
2
1
0
ZNF287
JOSD2
RBM12
LGR5
ANXA1
Target protein

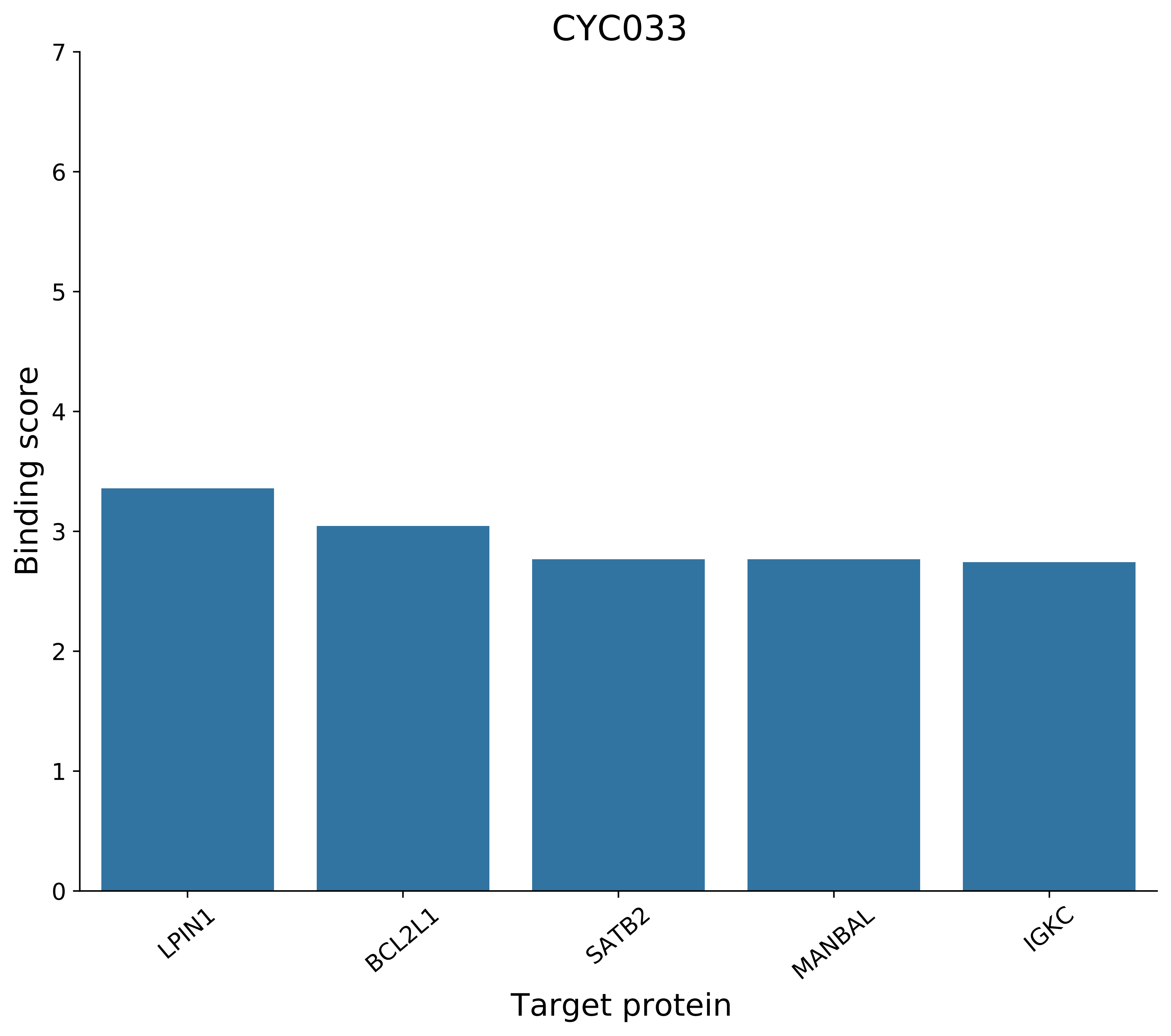
CYC033
Binding score
0
1
2
3
4
5
6
7
LPIN1
BCL2L1
SATB2
MANBAL
IGKC
Target protein

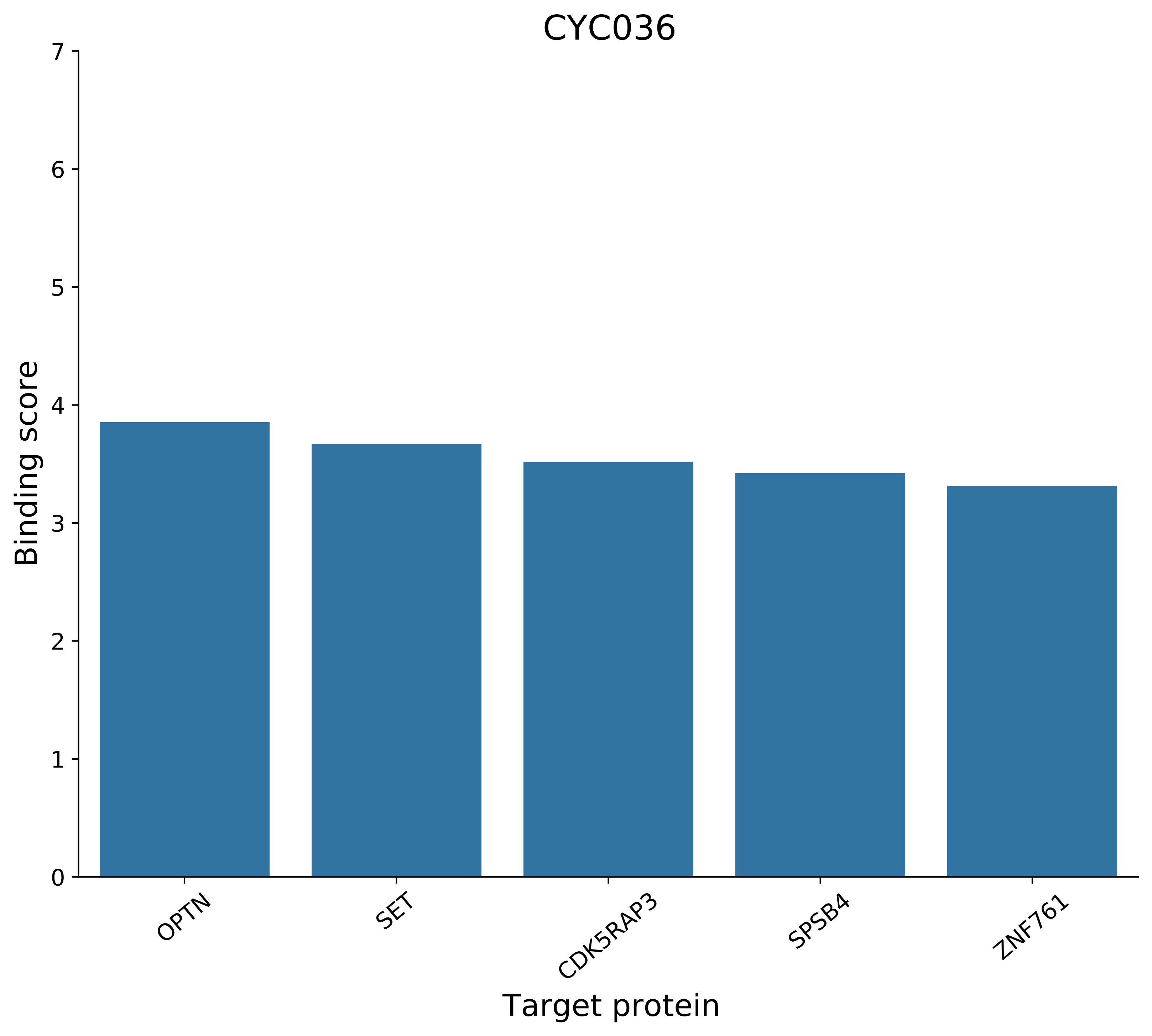

CYC036
Binding score
0
1
2
3
4
5
6
7
OPTN
SET
CDK5RAP3
SPSB4
ZNF761
Target protein

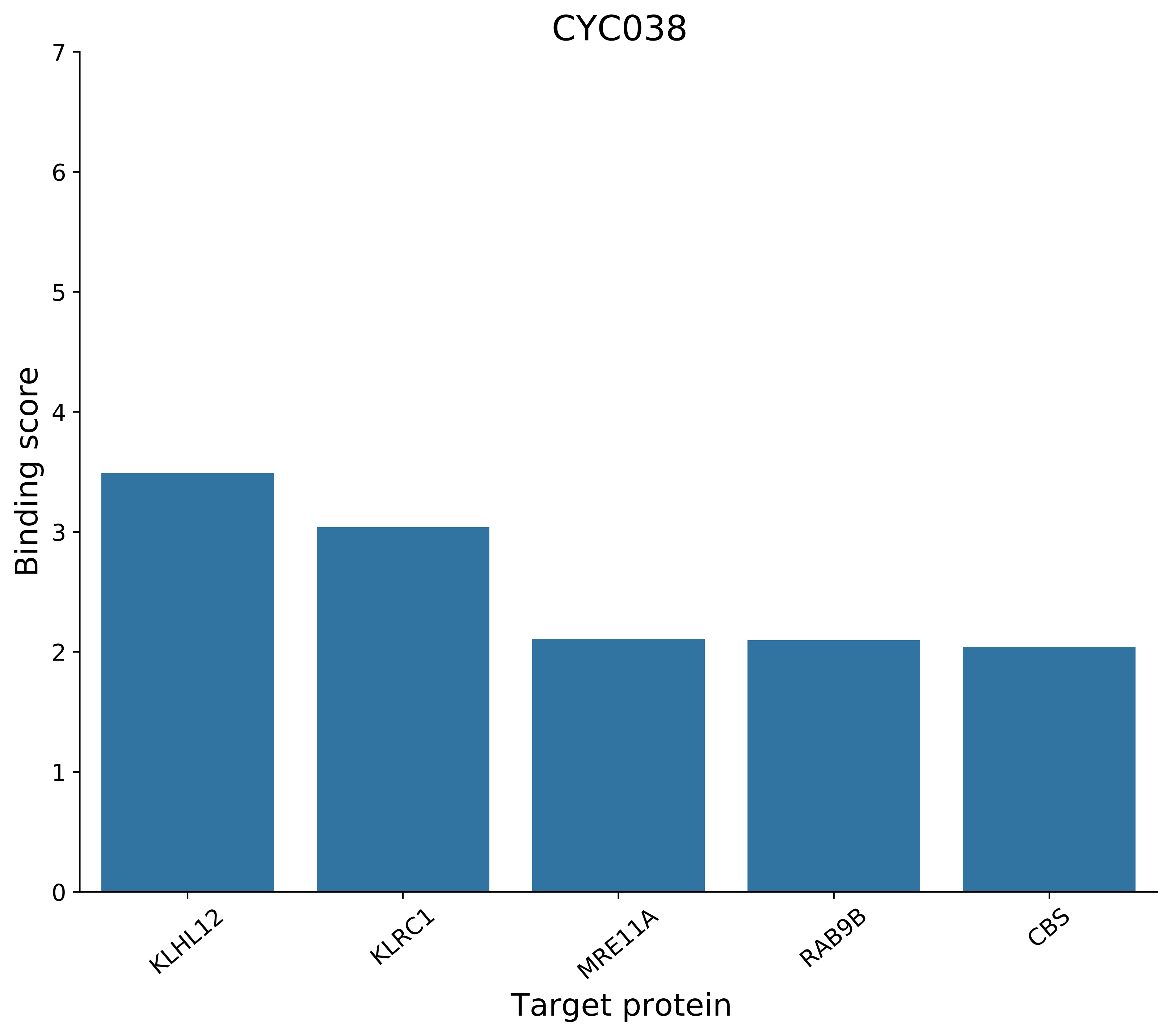
CYC038
Binding score
Target protein
0
1
2
3
4
5
6
7
KLHL12
KLRC1
MRE11A
RAB9B
CBS

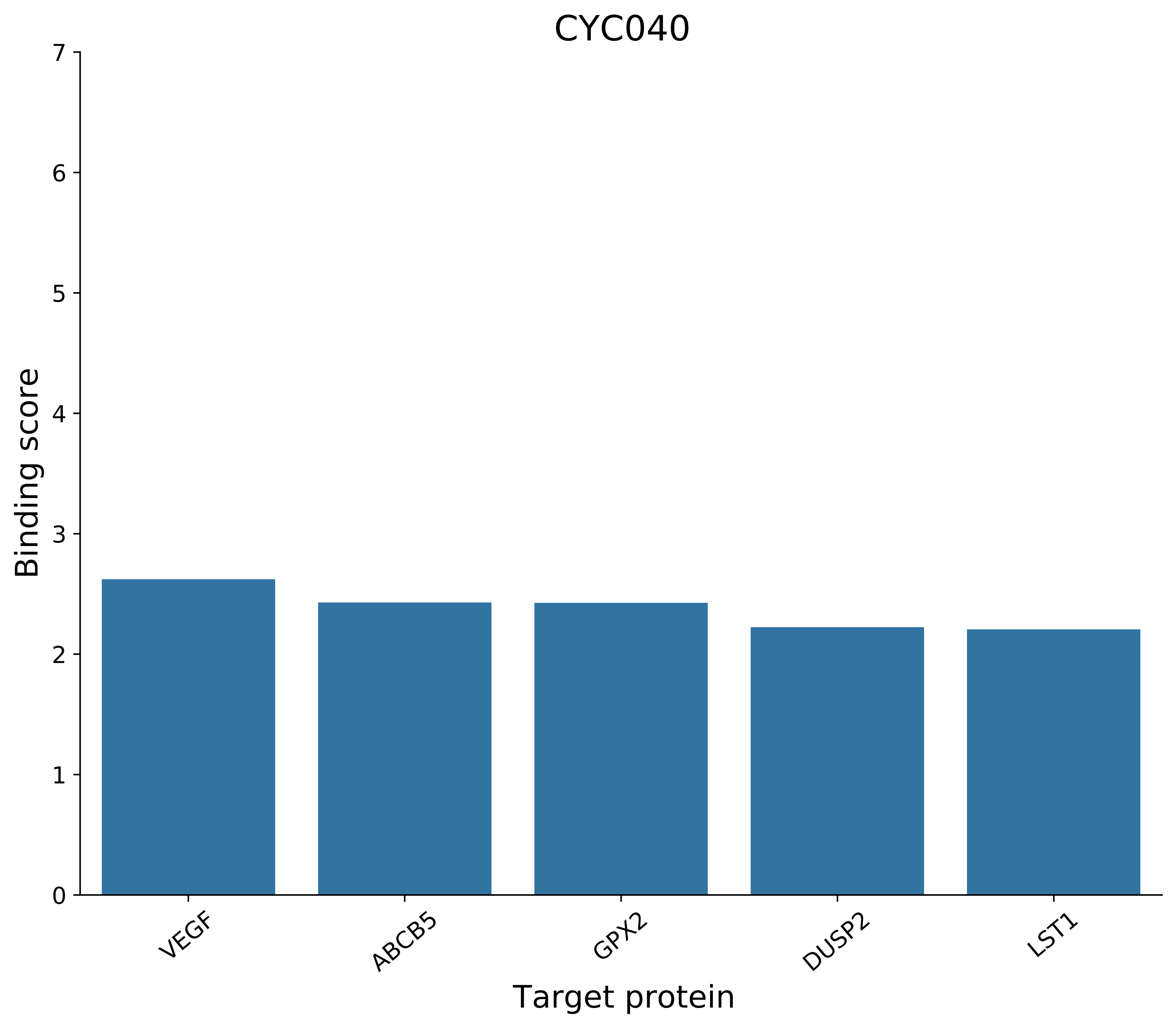
CYC040
Binding score
0
1
2
3
4
5
6
7
VEGF
ABCB5
GPX2
DUSP2
LST1
Target protein

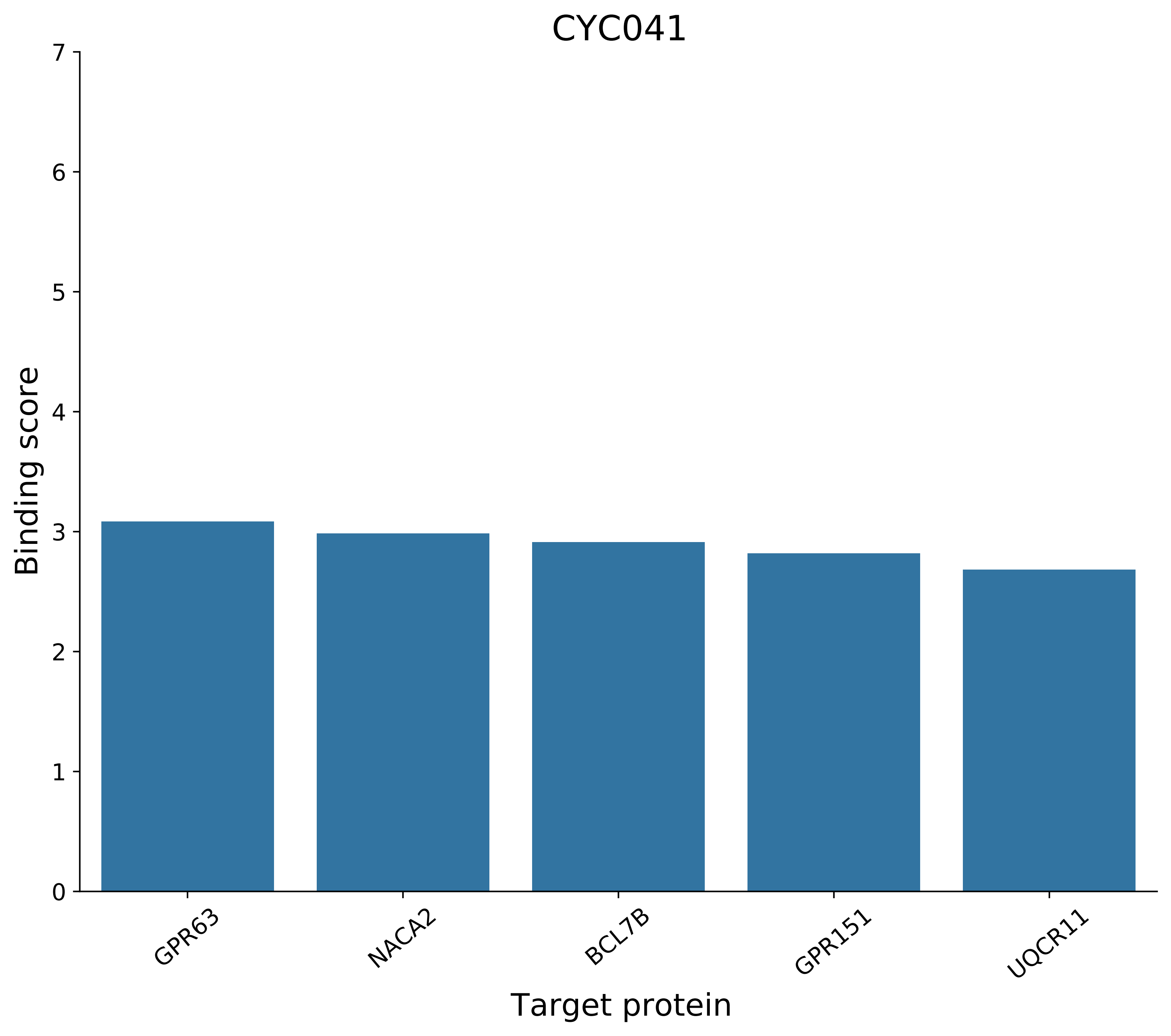
CYC041
Binding score
0
1
2
3
4
5
6
7
GPR63
NACA2
BCL7B
GPR151
UQCR11
Target protein

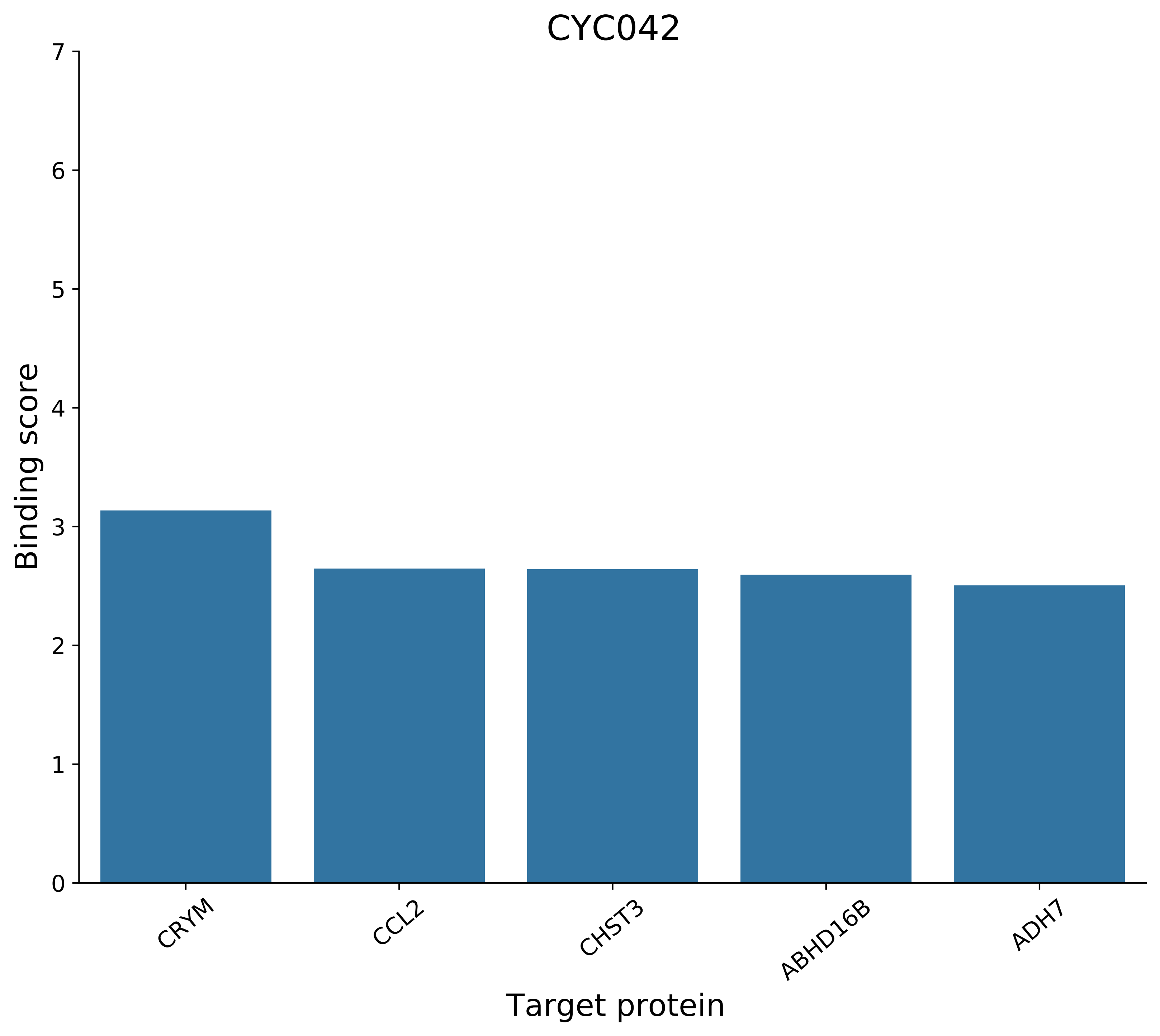
CYC042
Binding score
Target protein
0
1
2
3
4
5
6
7
CRYM
CCL2
CHST3
ABHD16B
ADH7

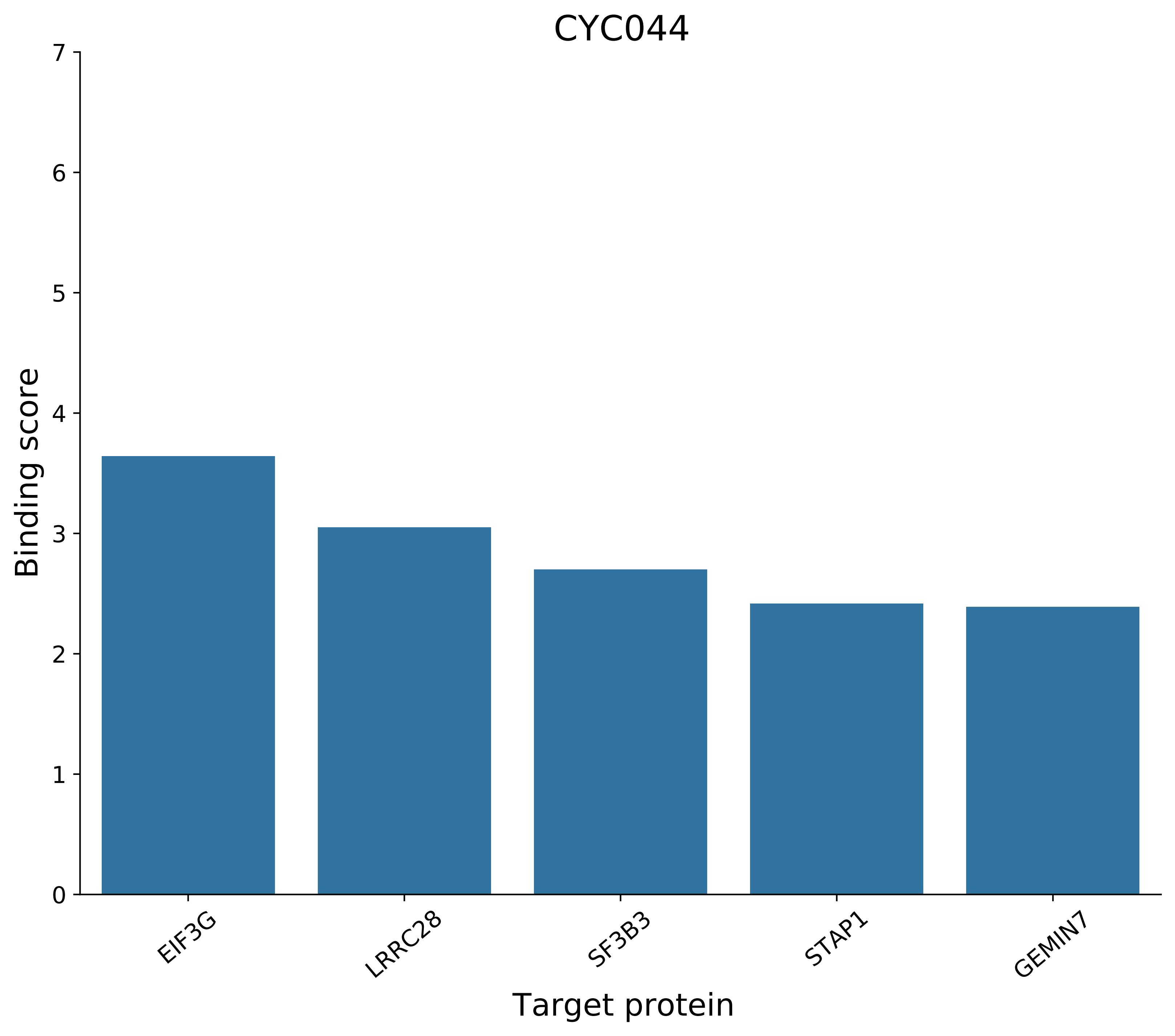
CYC044
Binding score
0
1
2
3
4
5
6
7
EIF3G
LRRC28
SF3B3
STAP1
GEMIN7
Target protein

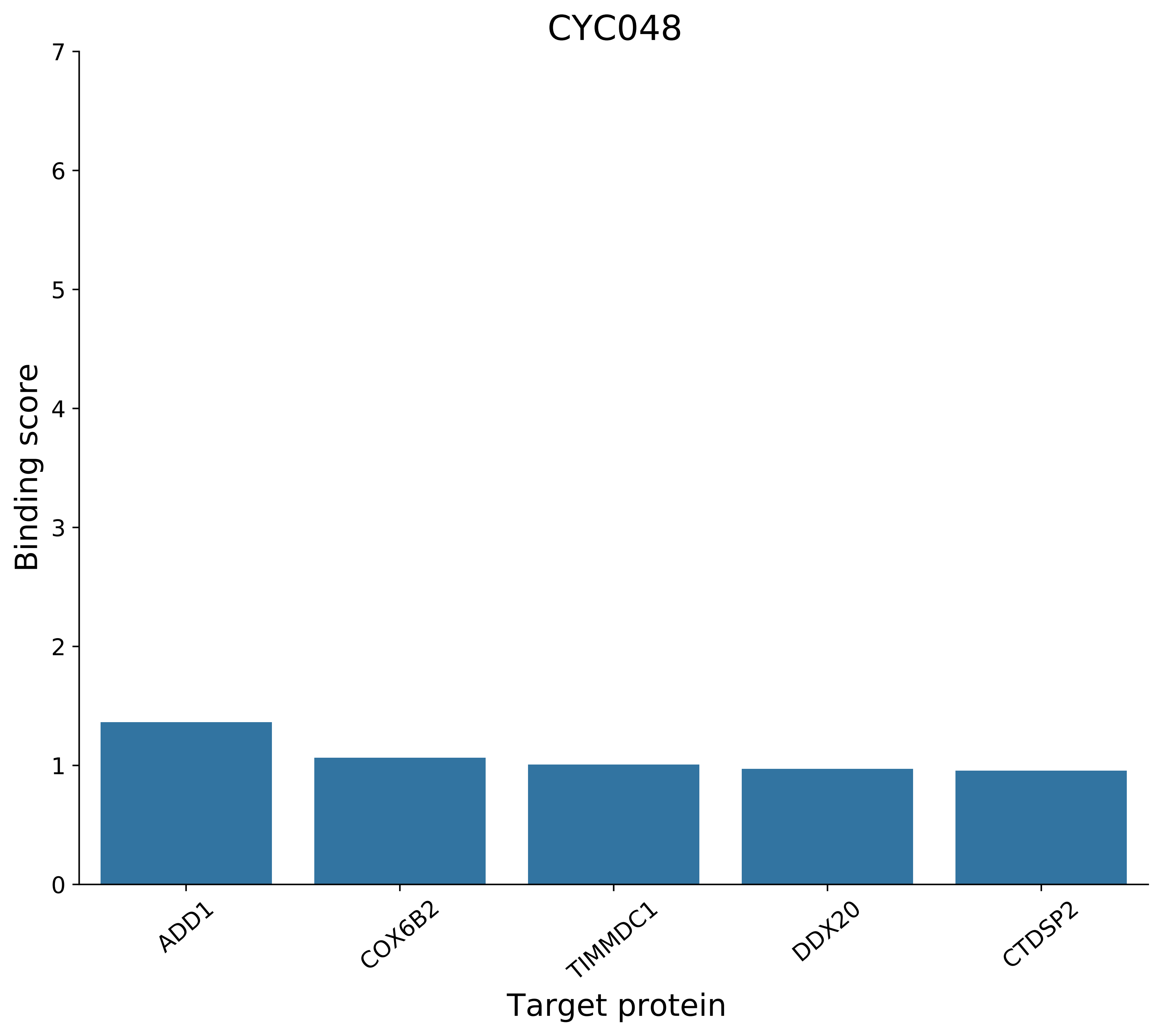
CYC048
Binding score
0
1
2
3
4
5
6
7
ADD1
COX6B2
TIMMDC1
DDX20
CTDSP2
Target protein

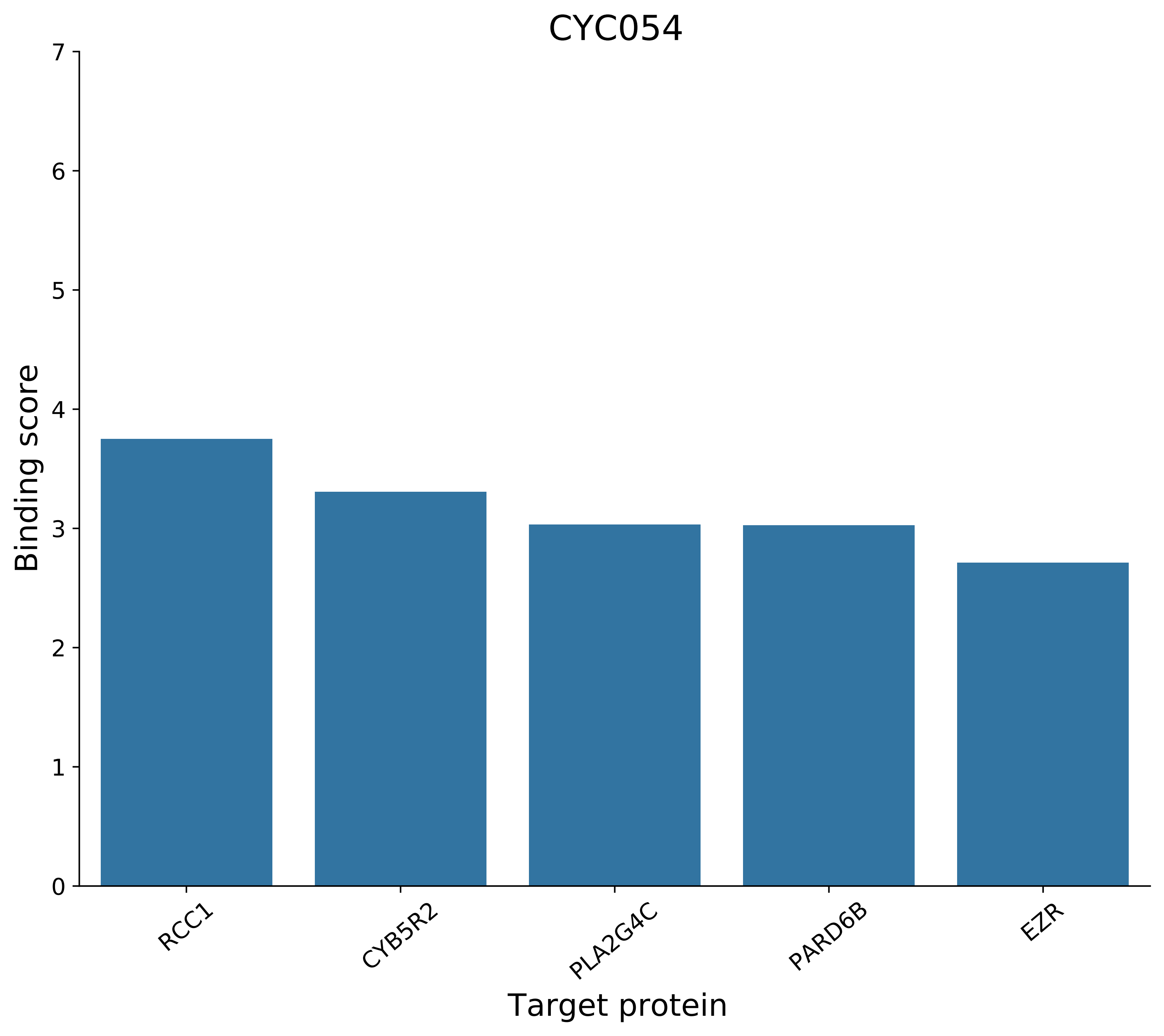
CYC054
Binding score
0
1
2
3
4
5
6
7
RCC1
CYB5R2
PLA2G4C
PARD6B
EZR
Target protein

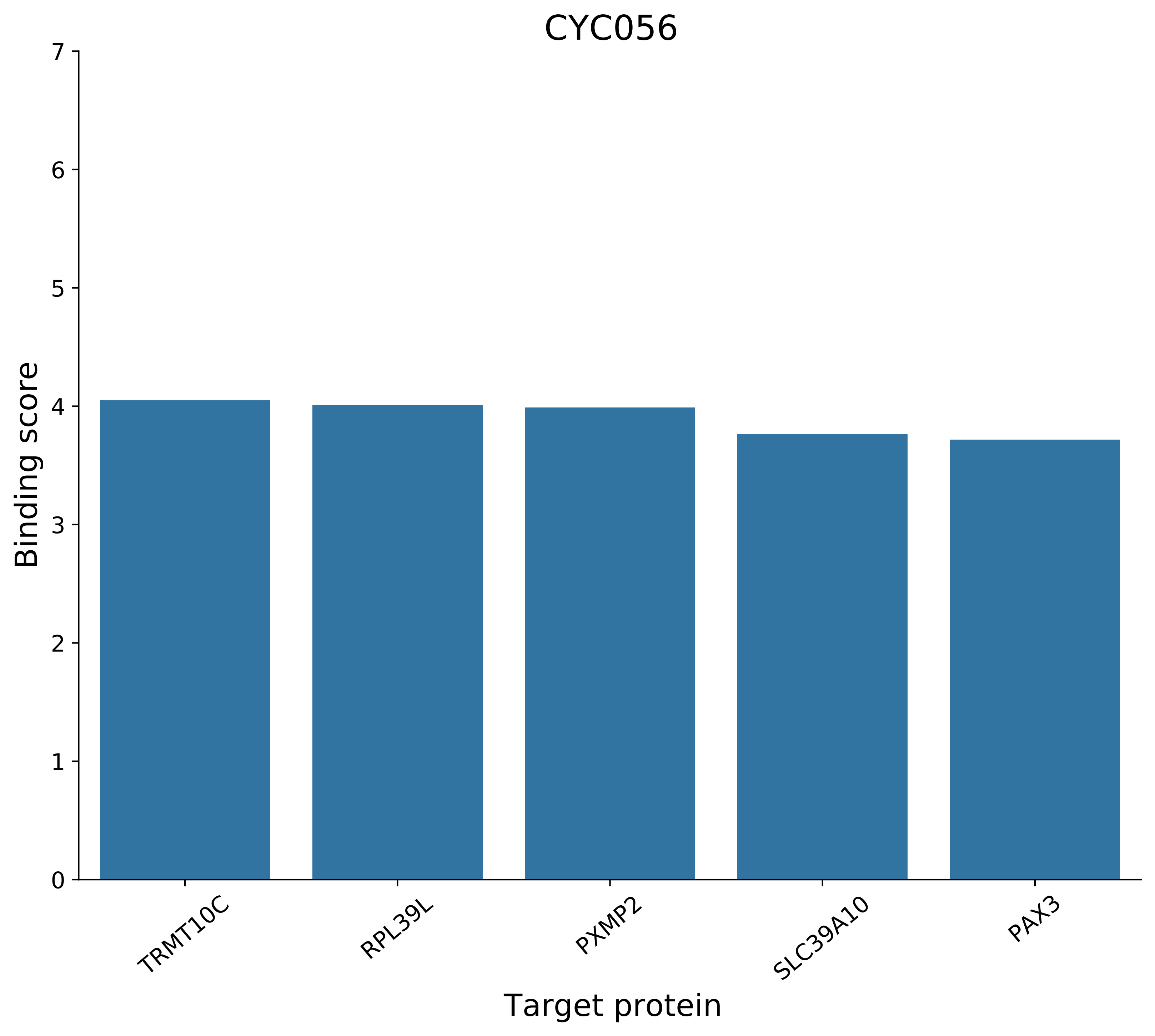

CYC056
Binding score
Target protein
0
1
2
3
4
5
6
7
TRMT10C
RPL39L
PXMP2
SLC39A10
PAX3

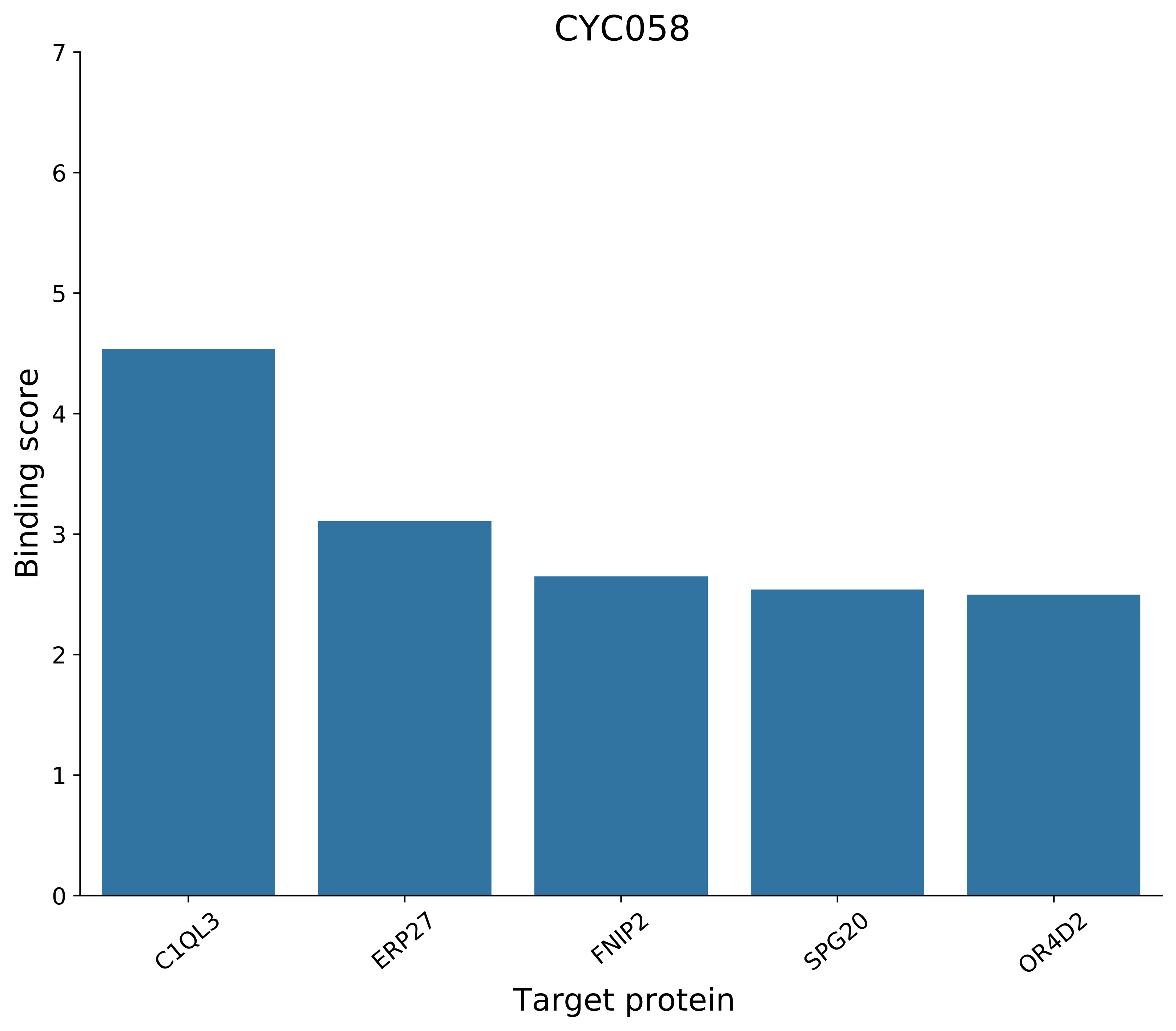

CYC058
Binding score
Target protein
0
1
2
3
4
5
6
7
C1QL3
ERP27
FNIP2
SPG20
OR4D2

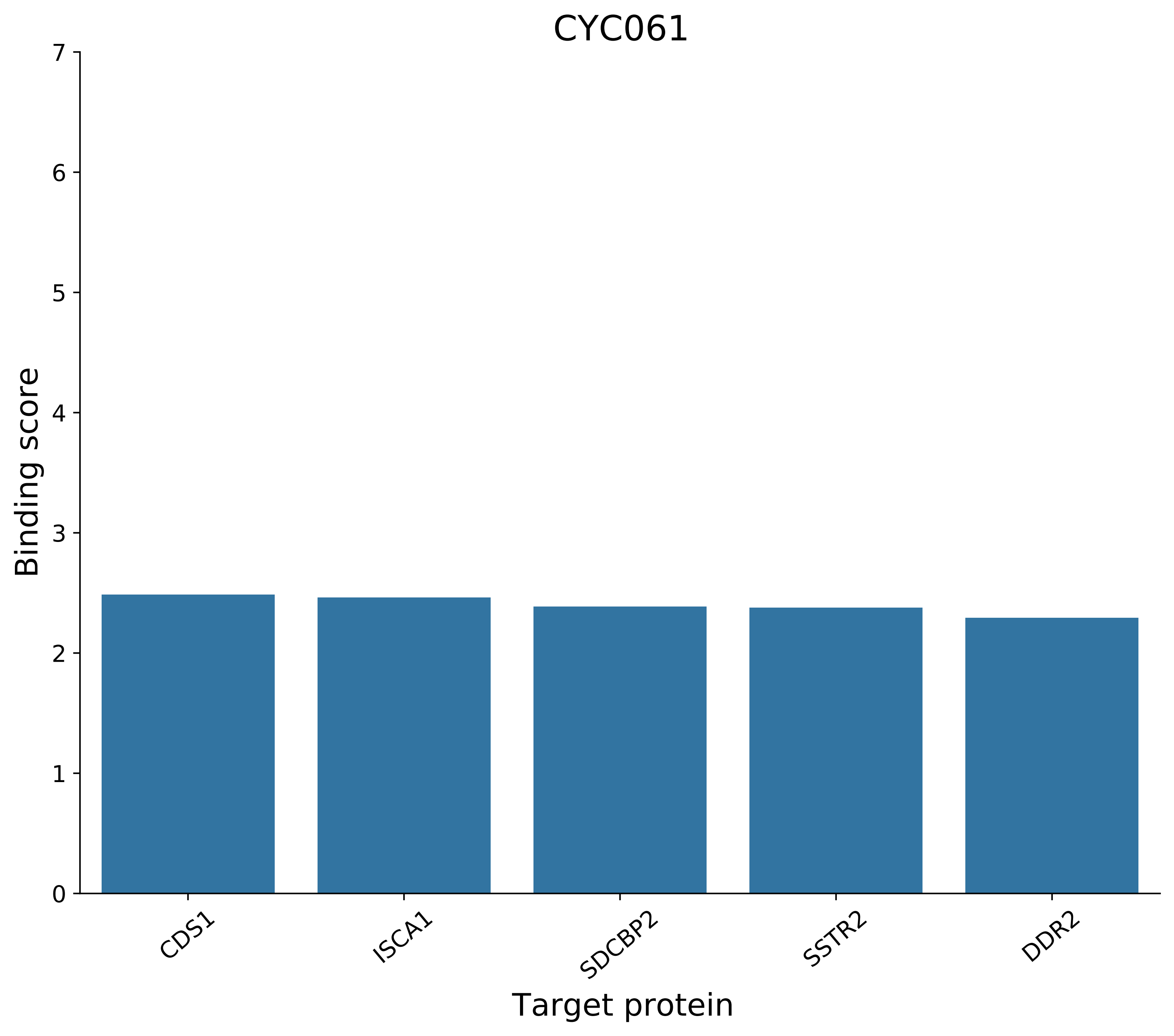
CYC061
Binding score
0
1
2
3
4
5
6
7
CDS1
ISCA1
SDCBP2
SSTR2
DDR2
Target protein

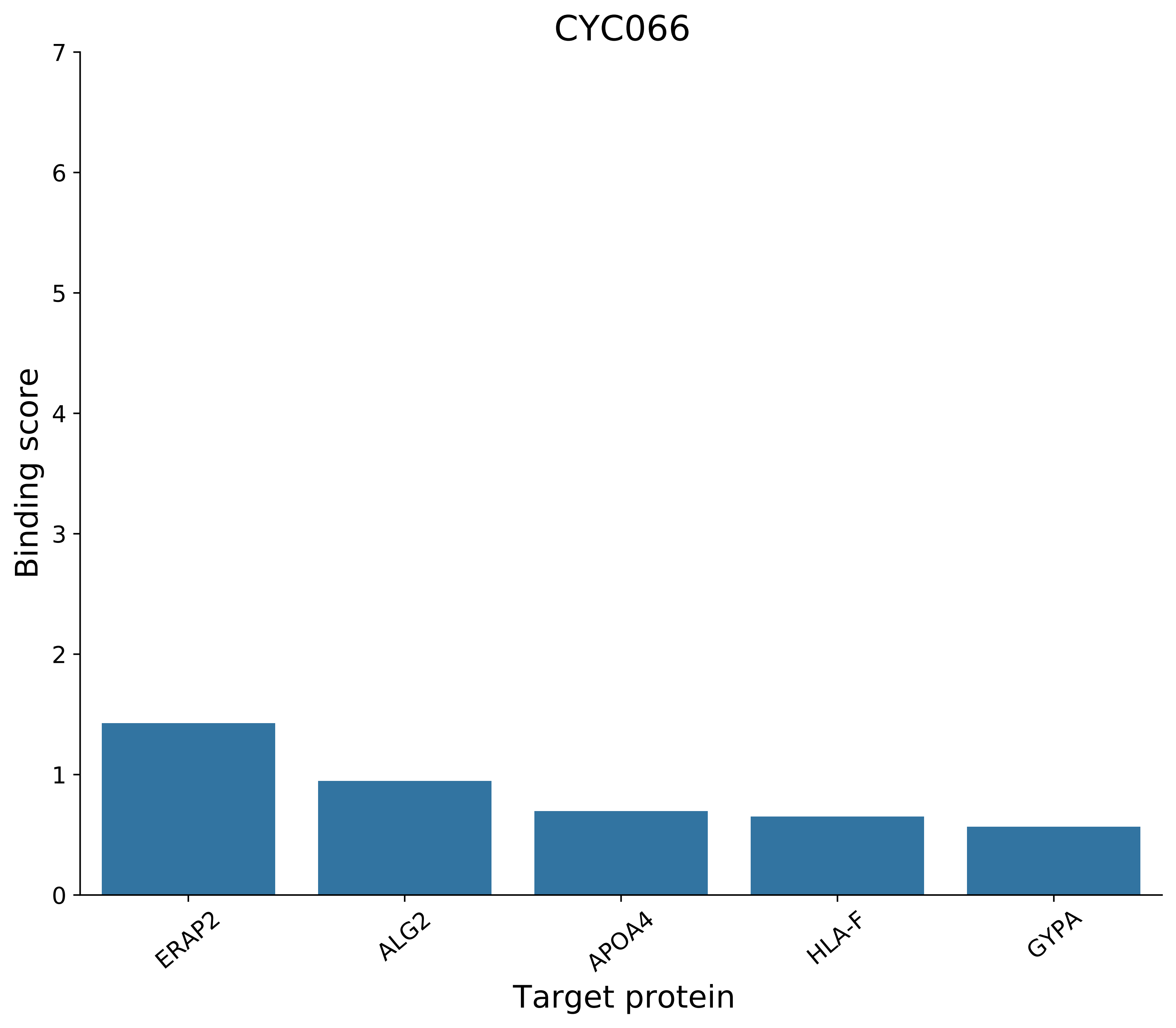
CYC066
Binding score
0
1
2
3
4
5
6
7
ERAP2
ALG2
APOA4
HLA-F
GYPA
Target protein

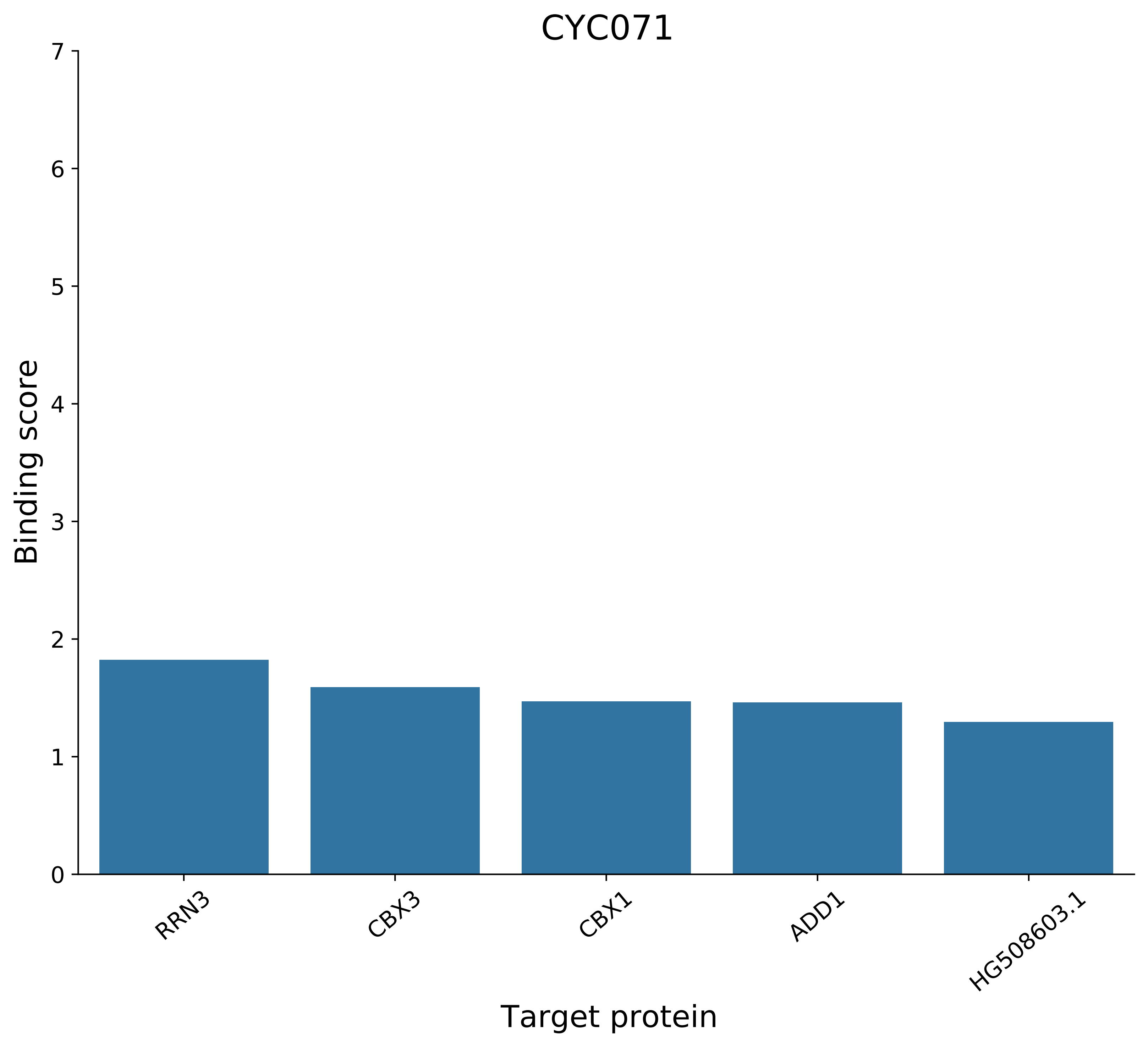
CYC071
Binding score
0
1
2
3
4
5
6
7
RRN3
CBX3
CBX1
ADD1
HG508603.1
Target protein

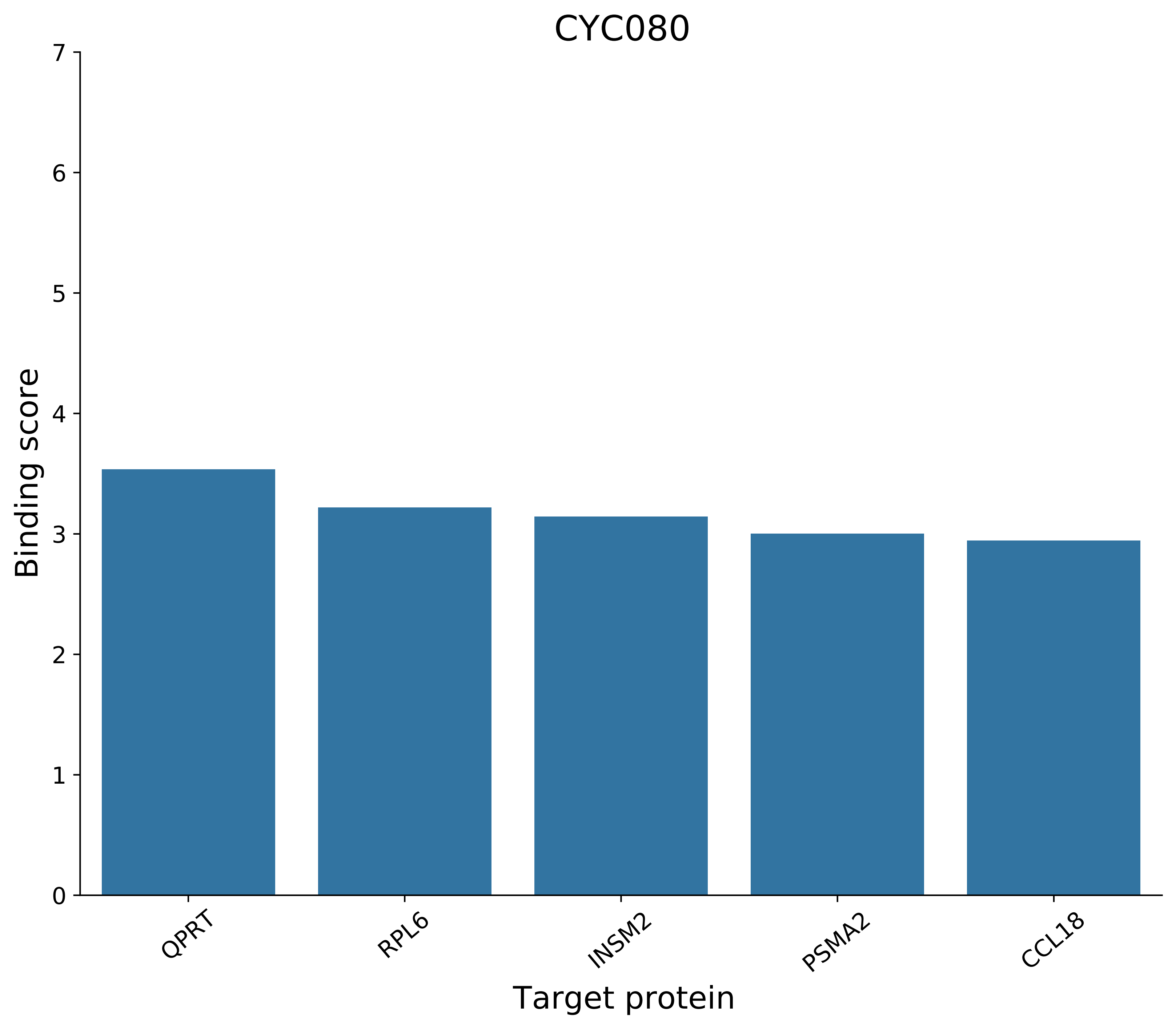
CYC080
Binding score
Target protein
0
1
2
3
4
5
6
7
QPRT
RPL6
INSM2
PSMA2
CCL18

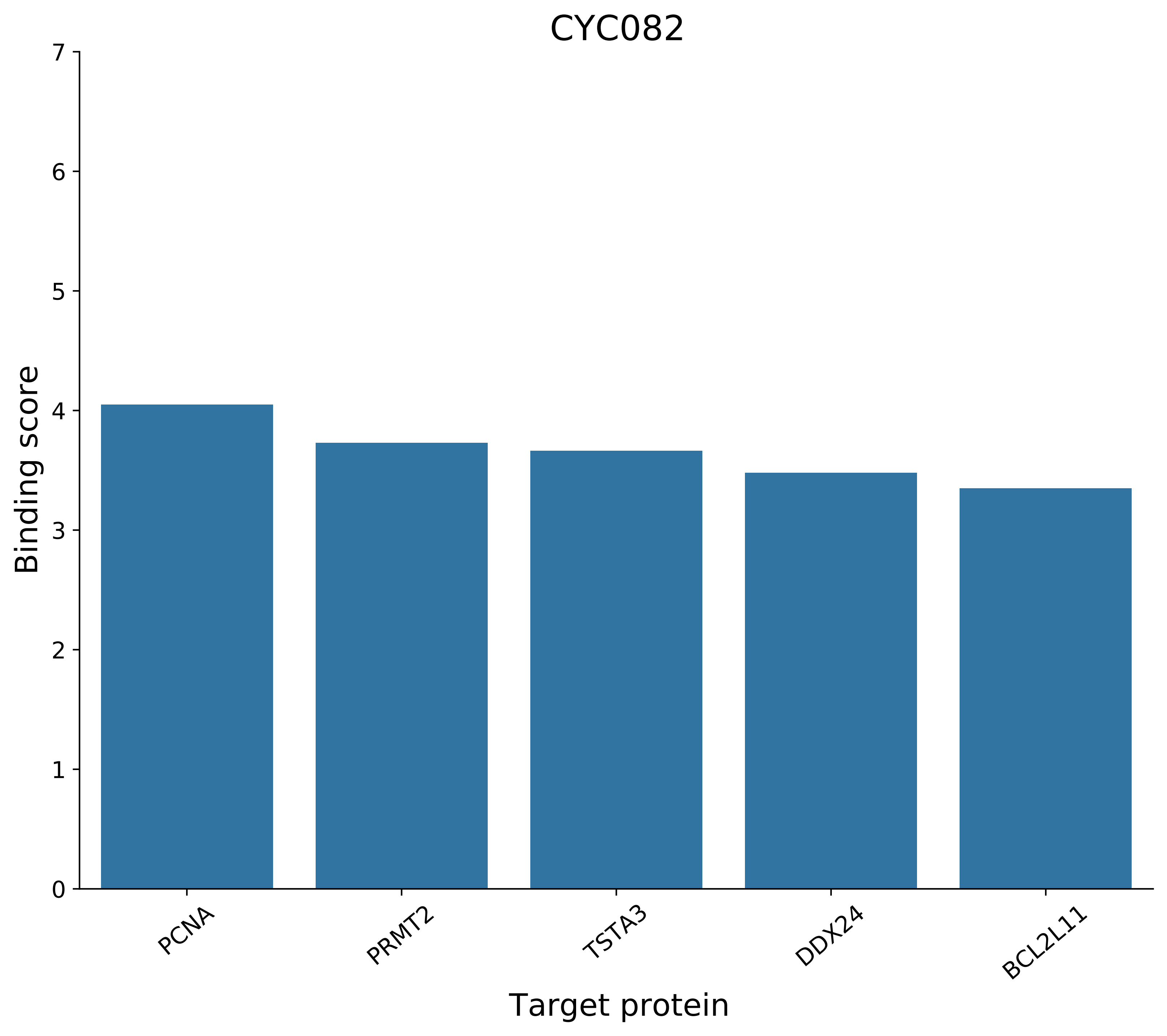

CYC082
Binding score
0
1
2
3
4
5
6
7
PCNA
PRMT2
TSTA3
DDX24
BCL2L11
Target protein

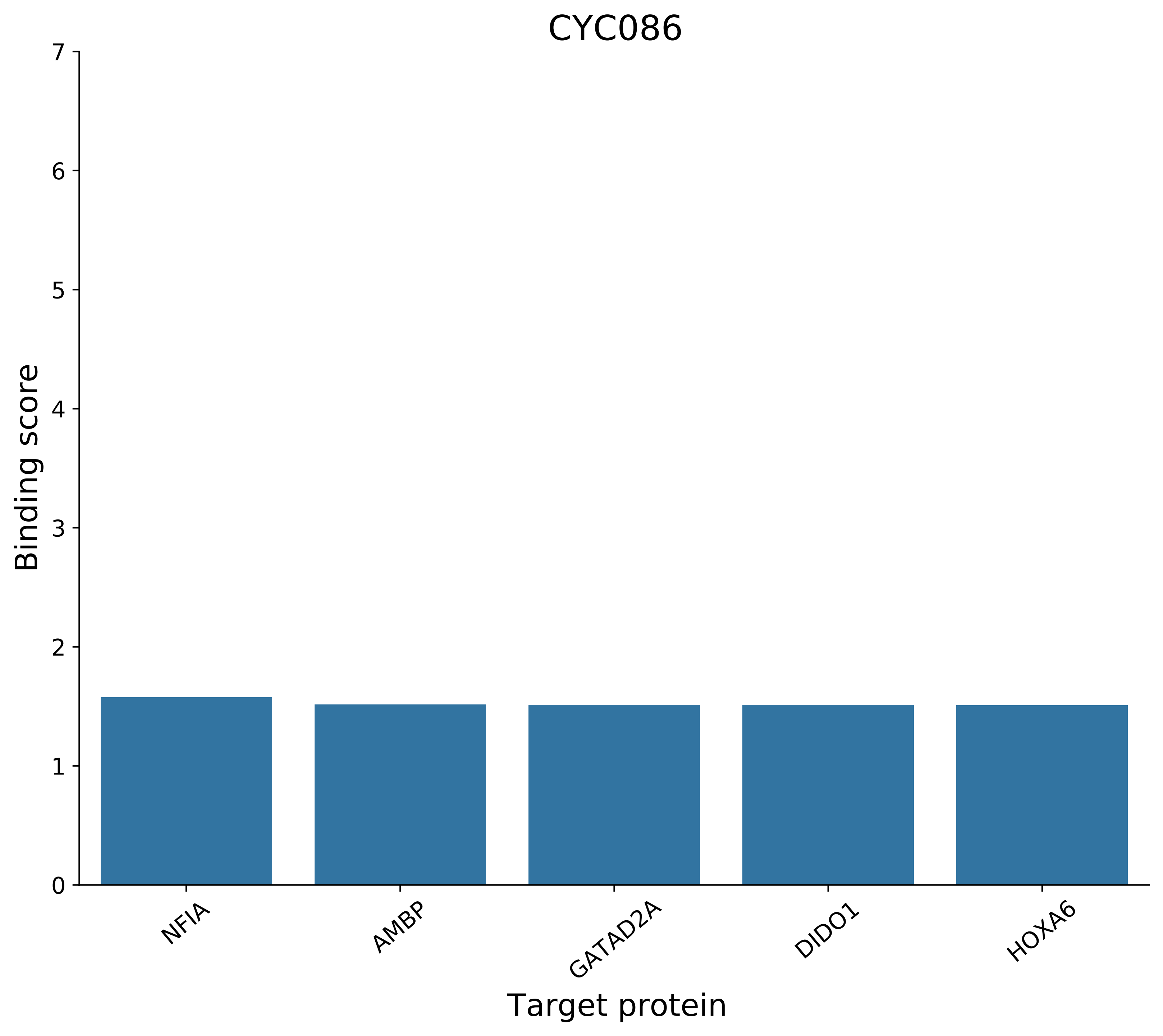
CYC086
Binding score
0
1
2
3
4
5
6
7
NFIA
AMBP
GATAD2A
DIDO1
HOXA6
Target protein

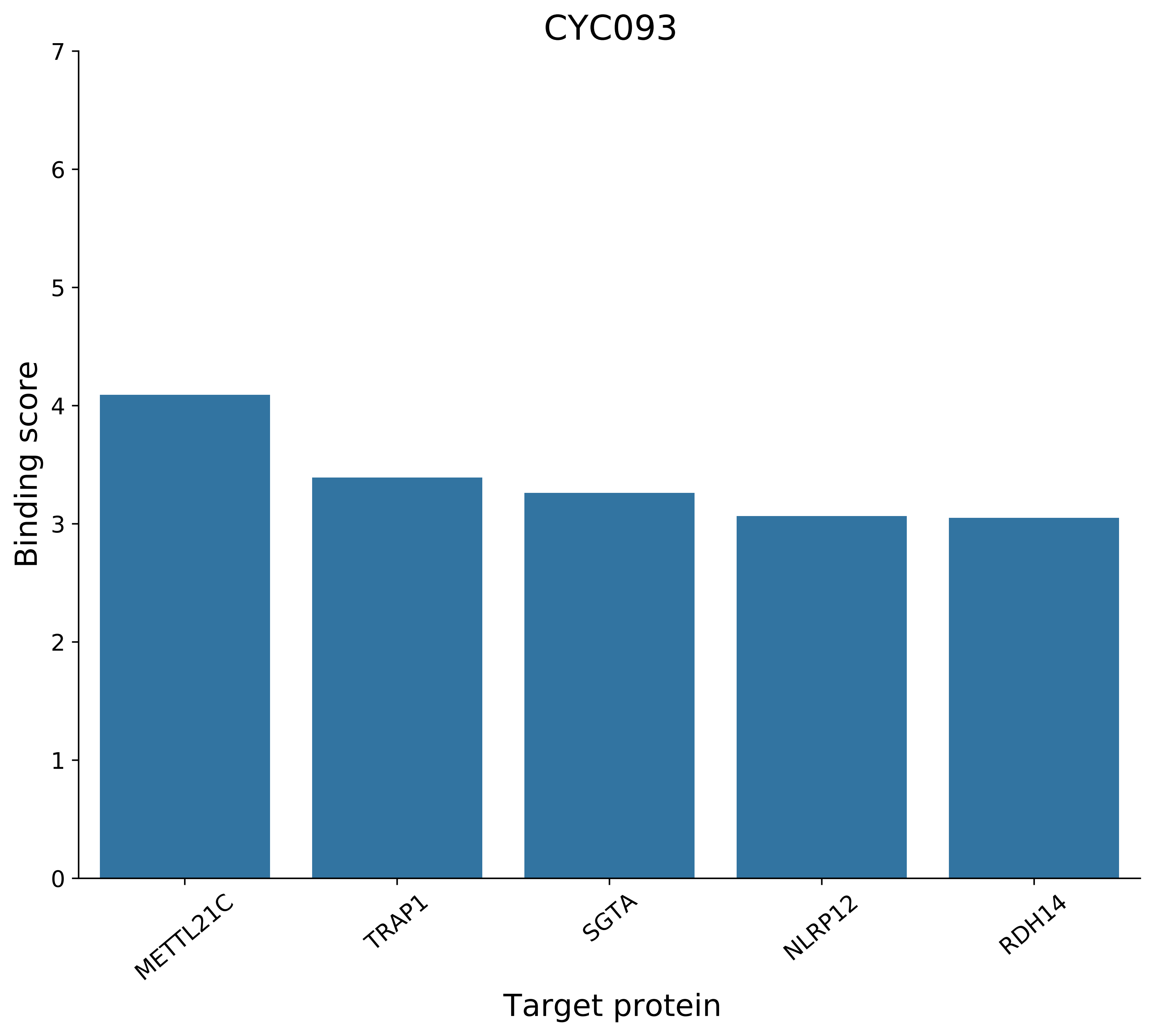
CYC093
Binding score
Target protein
0
1
2
3
4
5
6
7
METTL21C
TRAP1
SGTA
NLRP12
RDH14

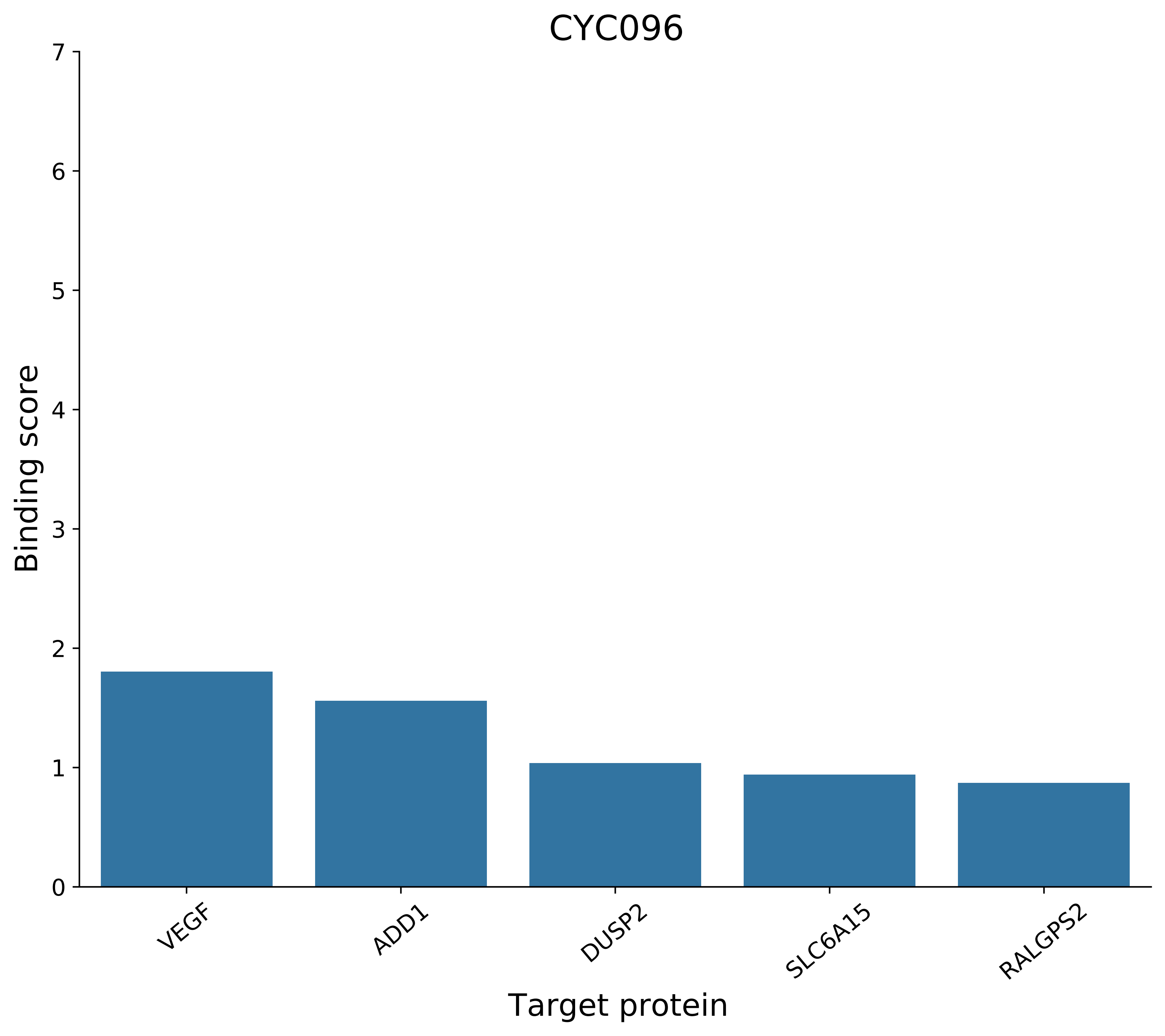
CYC096
Binding score
0
1
2
3
4
5
6
7
VEGF
ADD1
DUSP2
SLC6A15
RALGPS2
Target protein

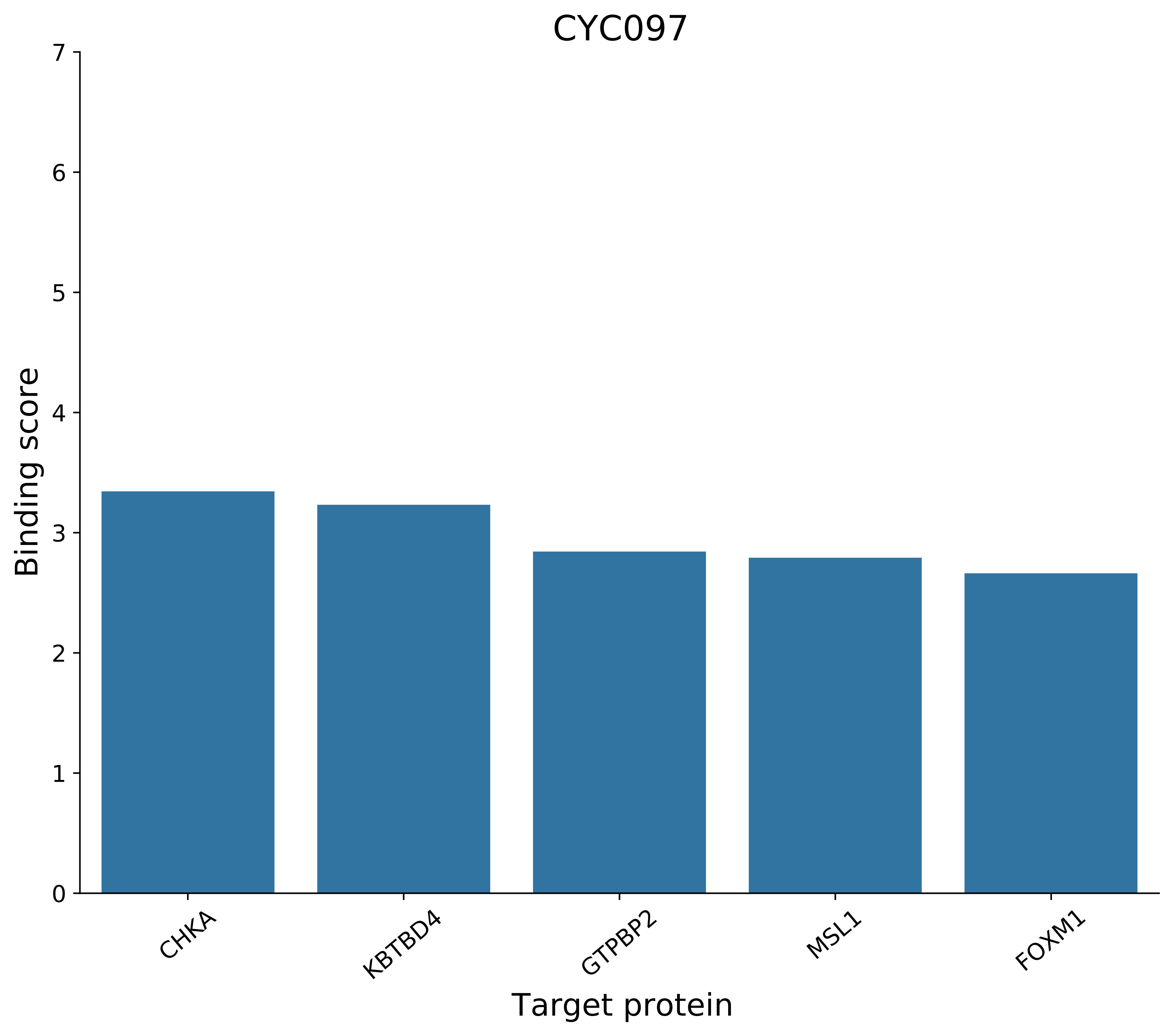

CYC097
Binding score
0
1
2
3
4
5
6
7
CHKA
KBTBD4
GTPBP2
MSL1
FOXM1
Target protein

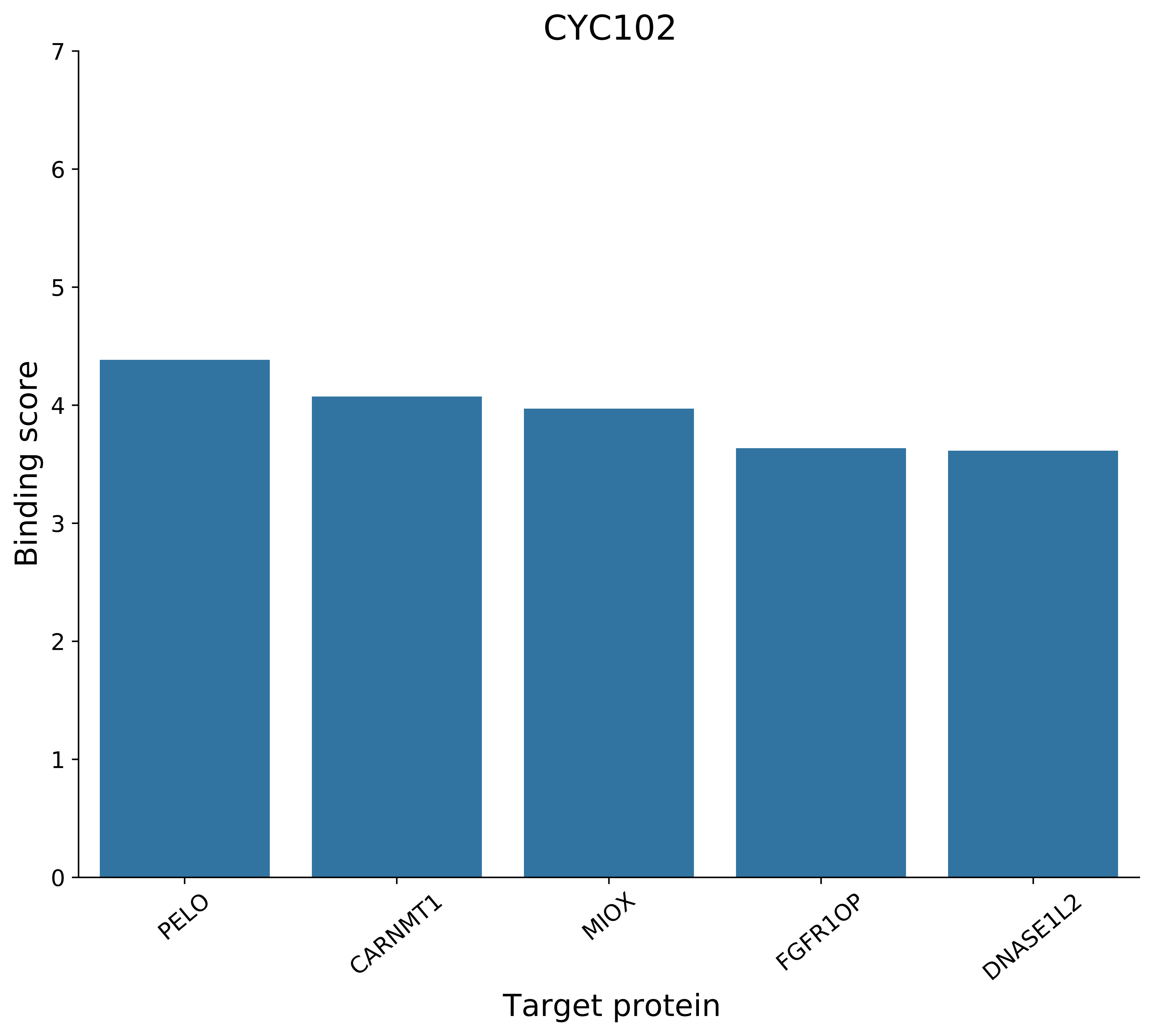

CYC102
Binding score
Target protein
0
1
2
3
4
5
6
7
PELO
CARNMT1
MIOX
FGFR1OP
DNASE1L2

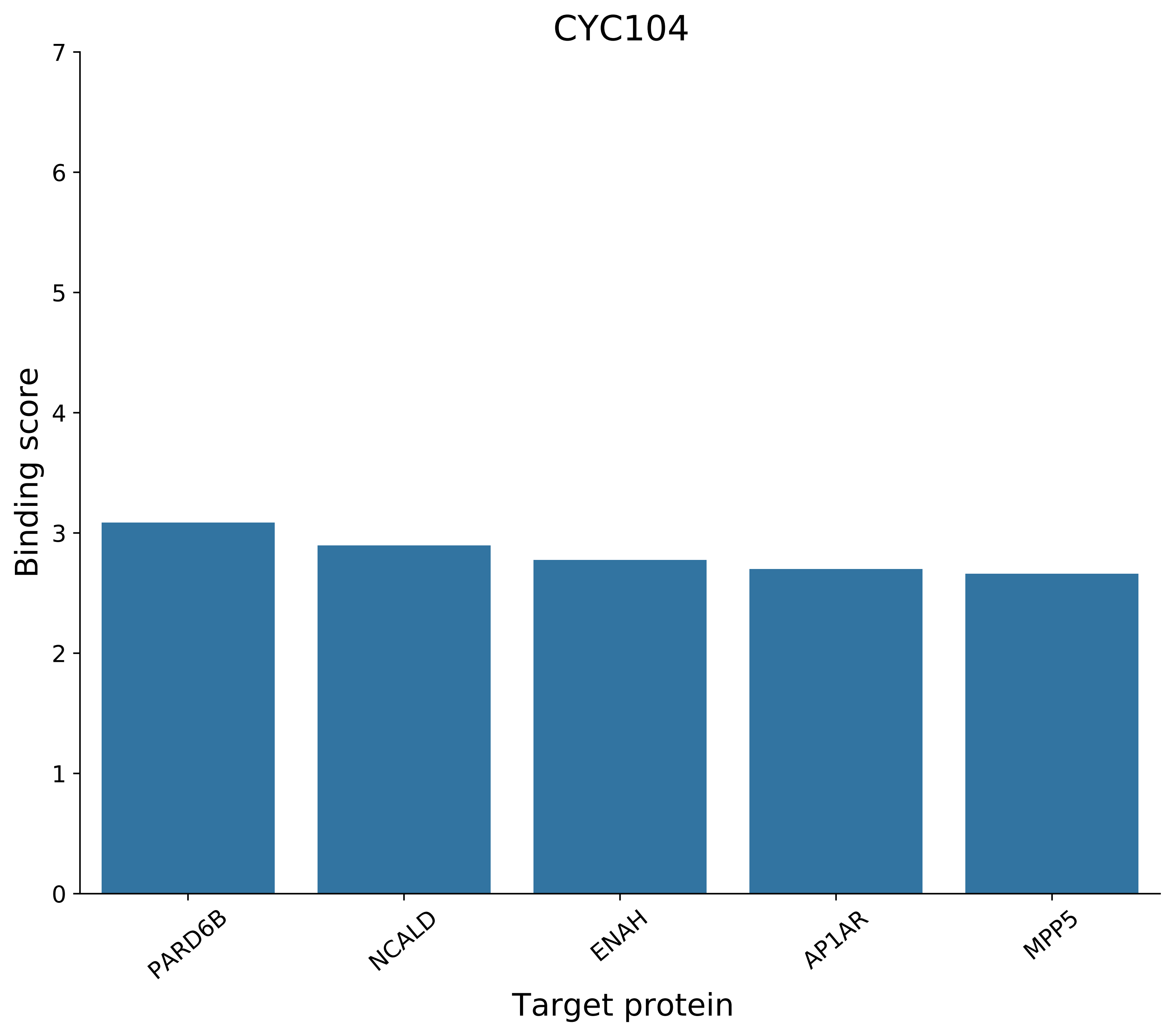
CYC104
Binding score
0
1
2
3
4
5
6
7
PARD6B
NCALD
ENAH
AP1AR
MPP5
Target protein

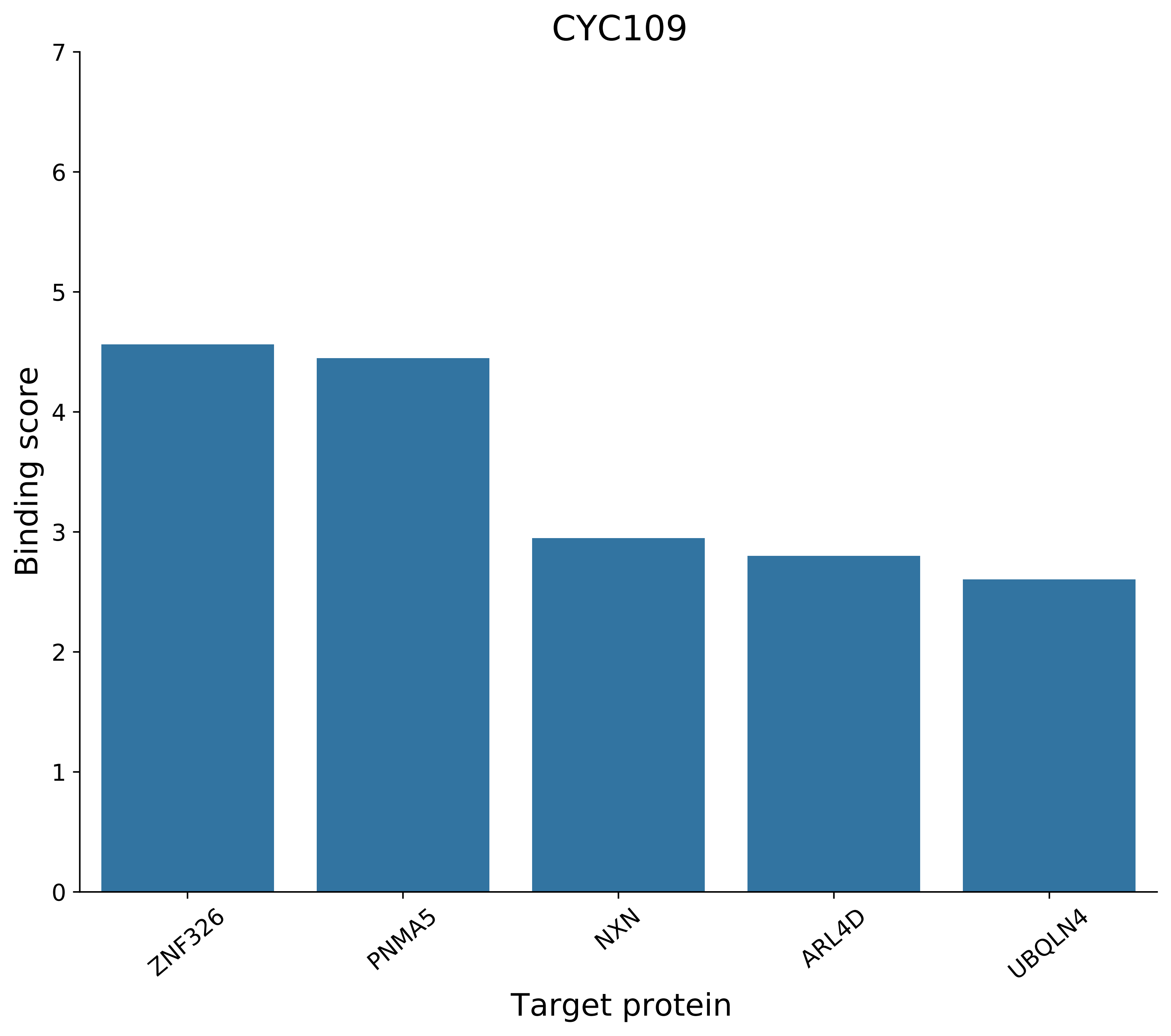
CYC109
Binding score
0
1
2
3
4
5
6
7
ZNF326
PNMA5
NXN
ARL4D
UBQLN4
Target protein

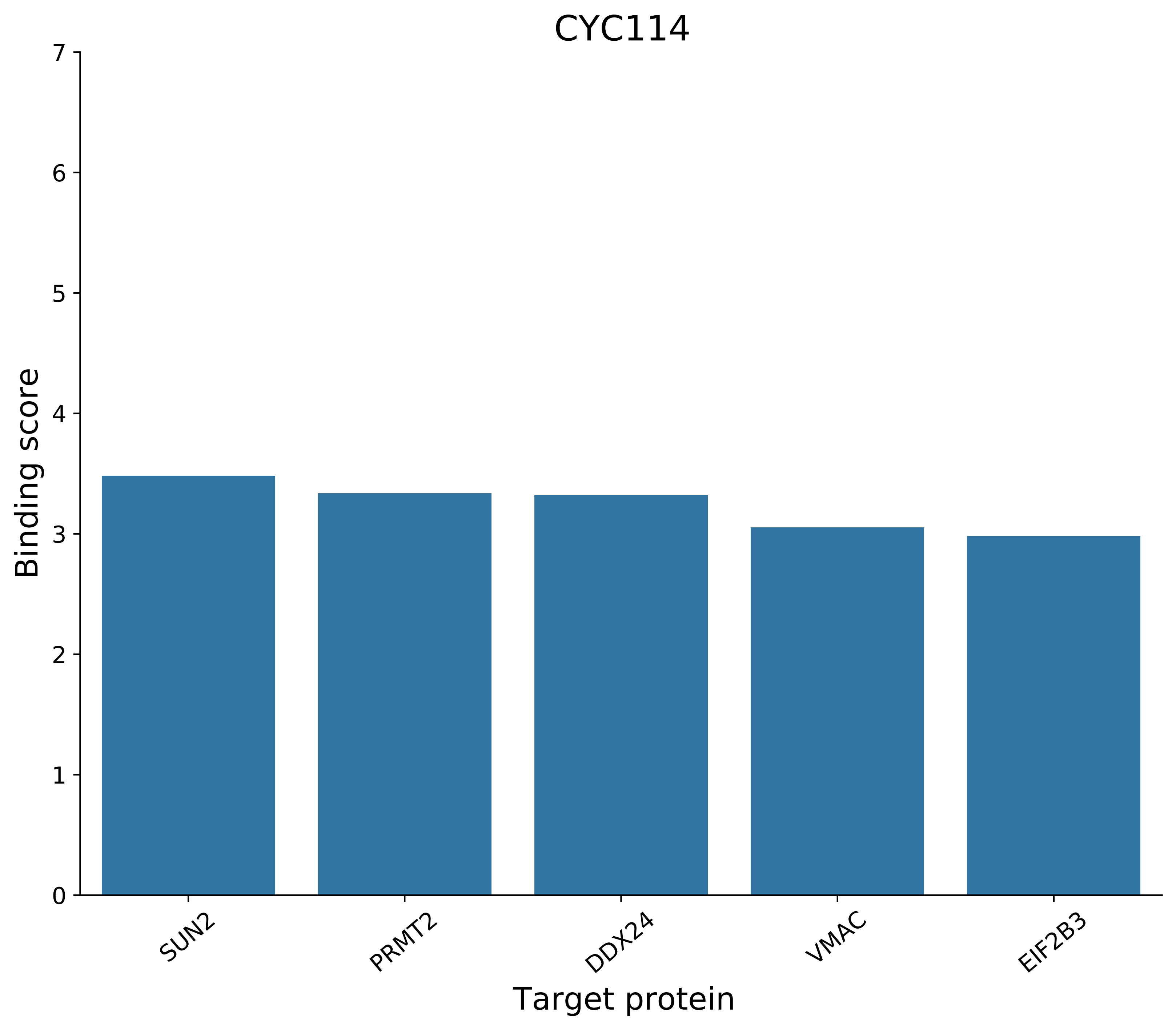
CYC114
Binding score
7
6
5
4
3
2
1
0
SUN2
PRMT2
DDX24
VMAC
EIF2B3
Target protein

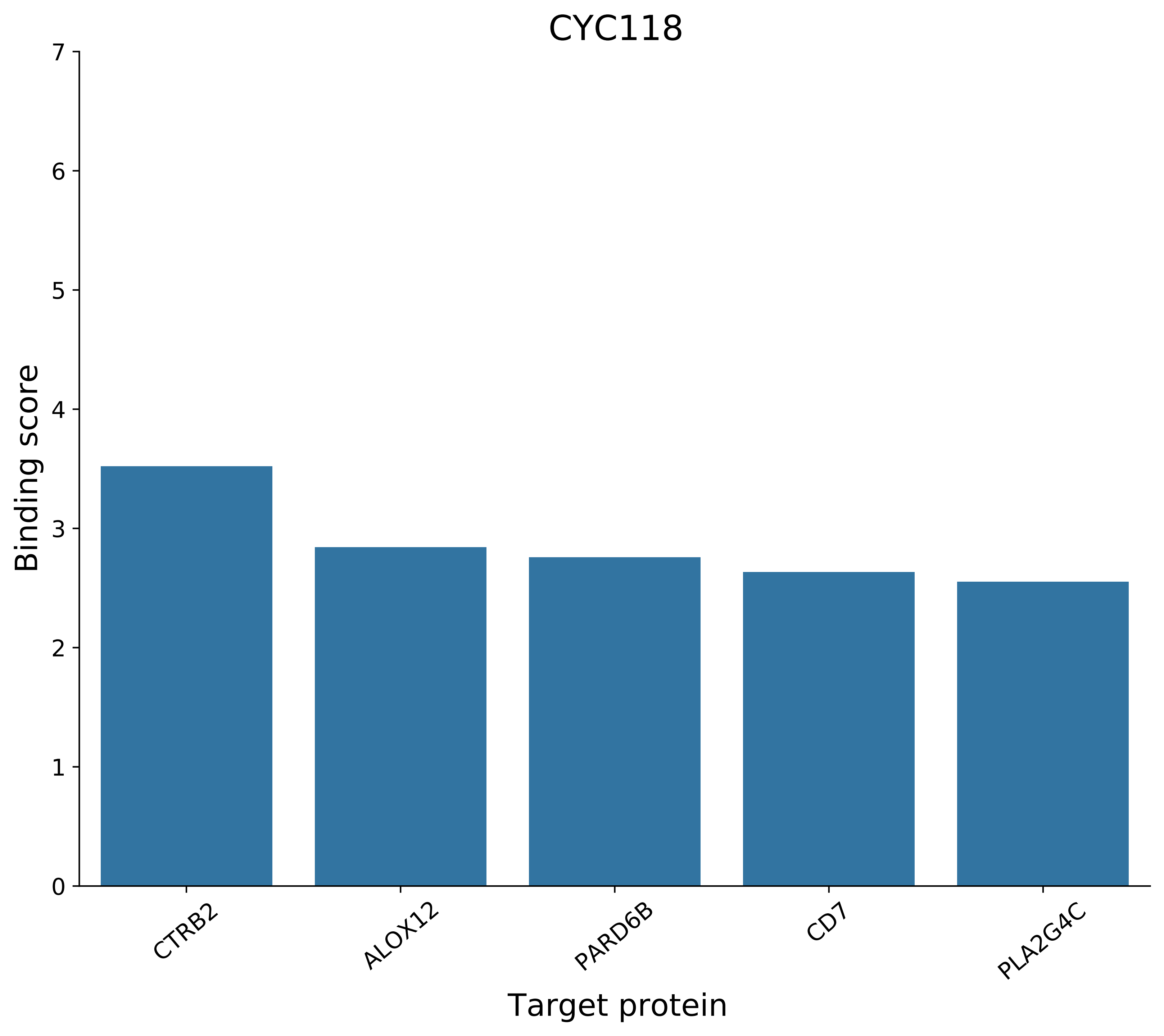
CYC118
Binding score
0
1
2
3
4
5
6
7
CTRB2
ALOX12
PARD6B
CD7
PLA2G4C
Target protein

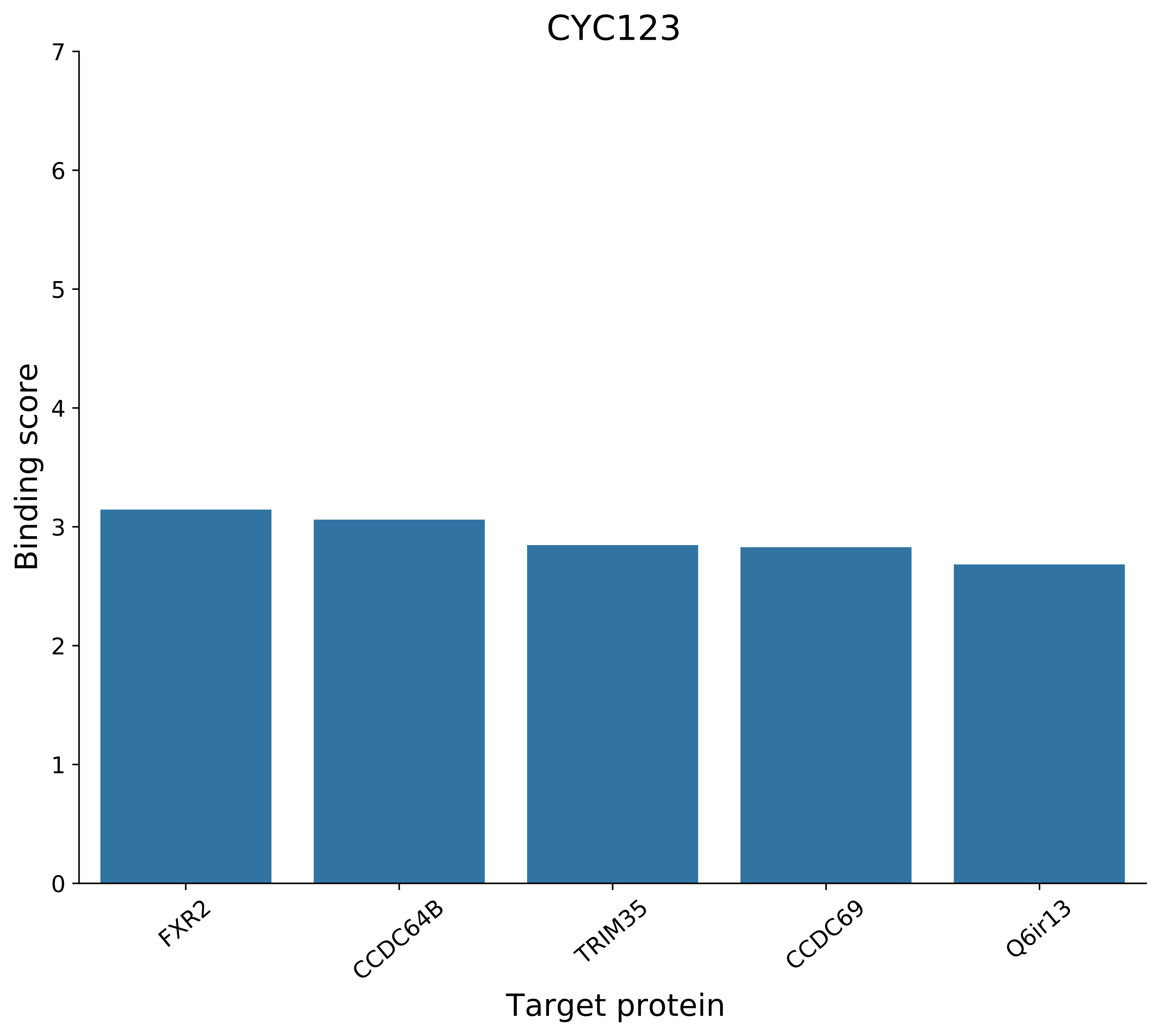
CYC123
Binding score
Target protein
0
1
2
3
4
5
6
7
FXR2
CCDC64B
TRIM35
CCDC69
Q6ir13

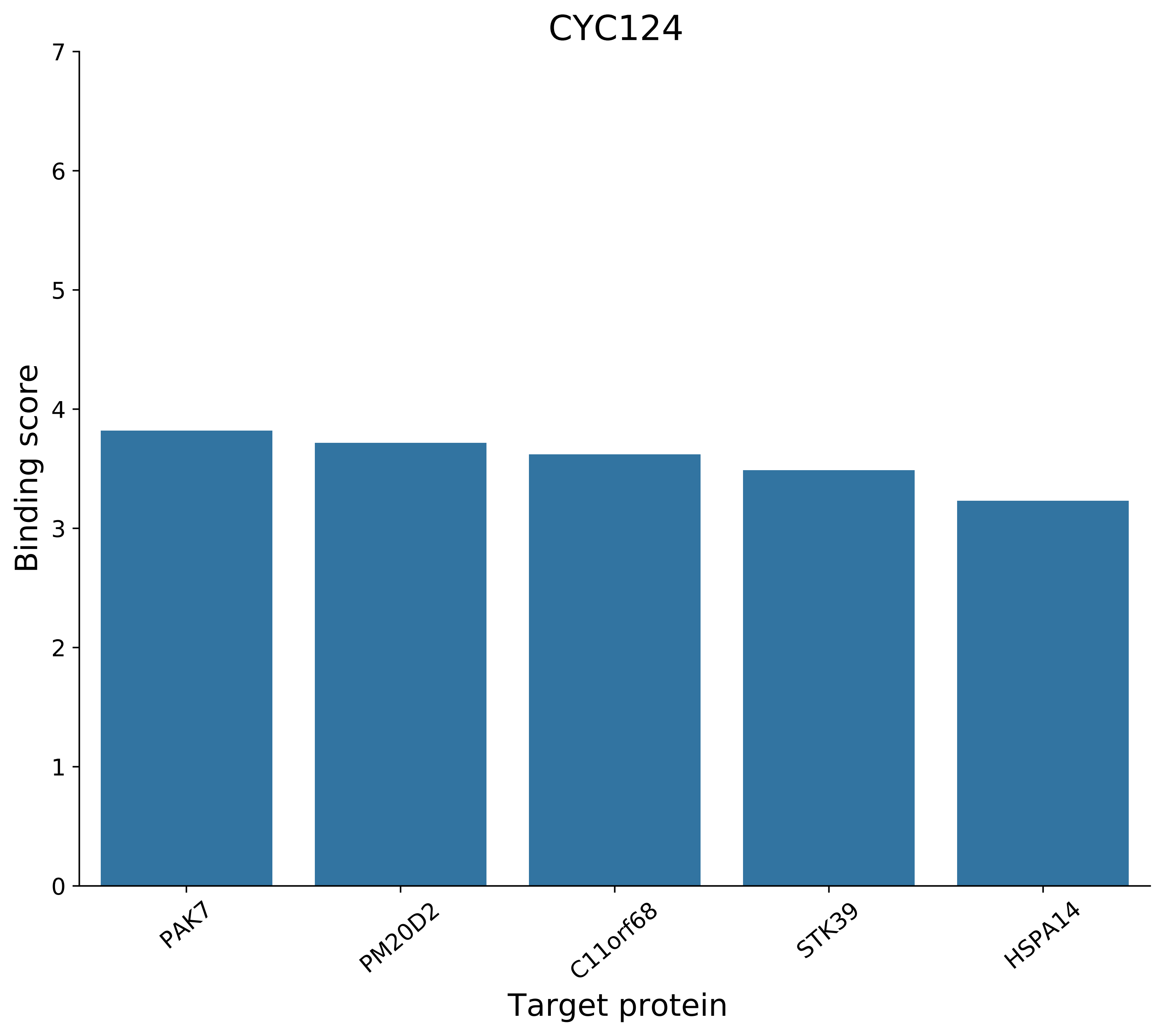
CYC124
Binding score
0
1
2
3
4
5
6
7
PAK7
PM20D2
C11orf68
STK39
HSPA14
Target protein

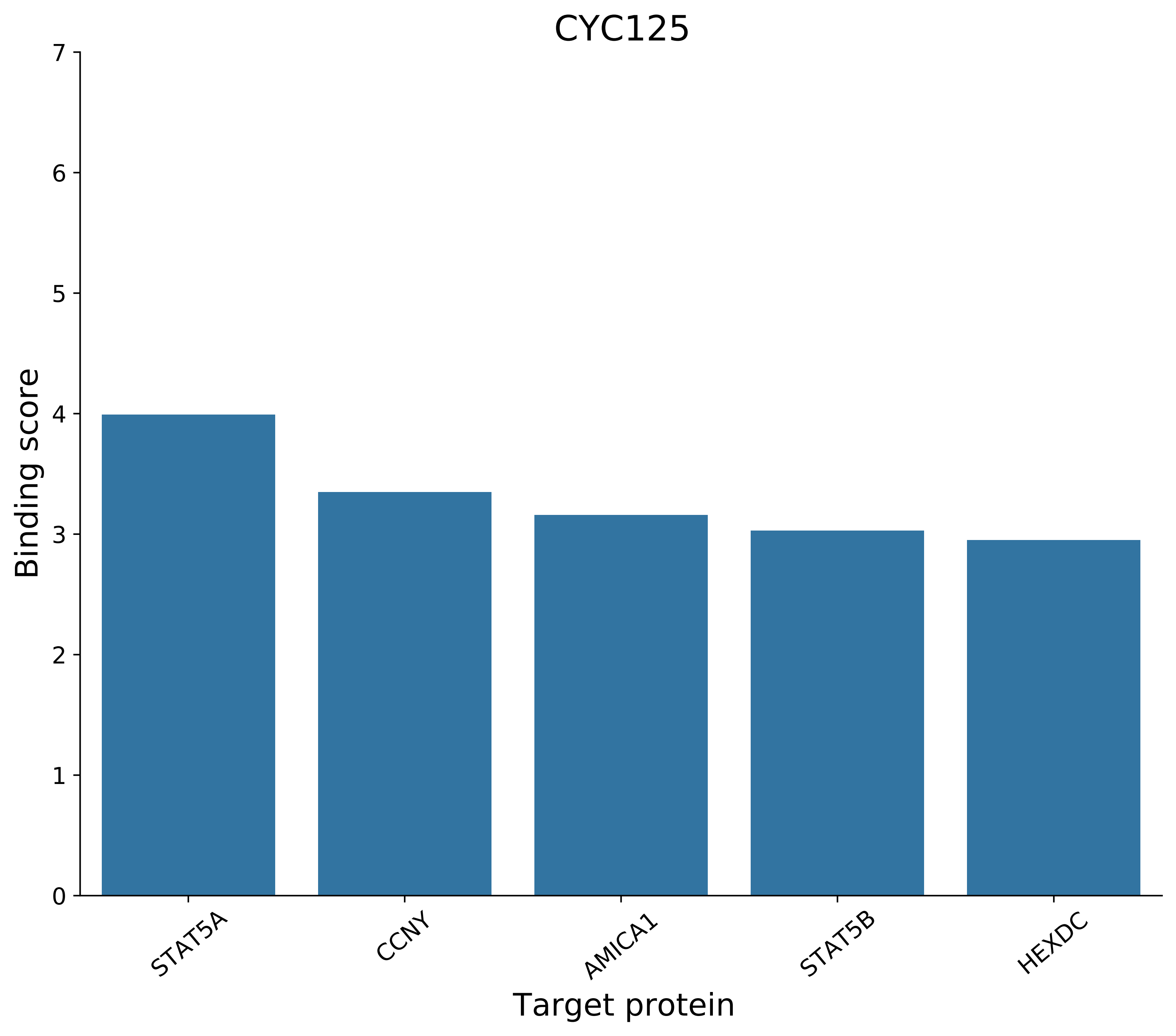

CYC125
Binding score
Target protein
0
1
2
3
4
5
6
7
STAT5A
CCNY
AMICA1
STAT5B
HEXDC

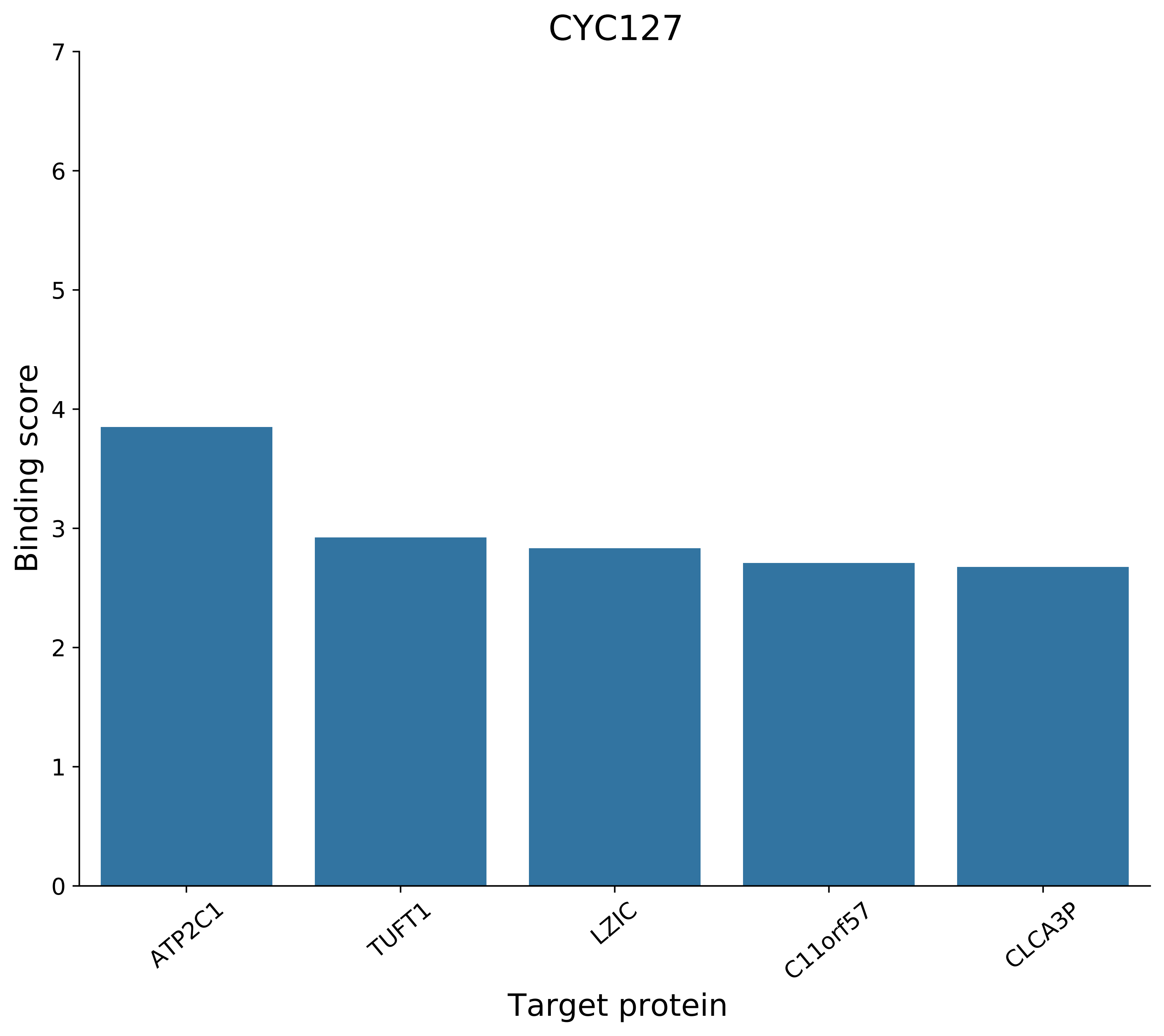

CYC127
Binding score
0
1
2
3
4
5
6
7
ATP2C1
TUFT1
LZIC
C11orf57
CLCA3P
Target protein

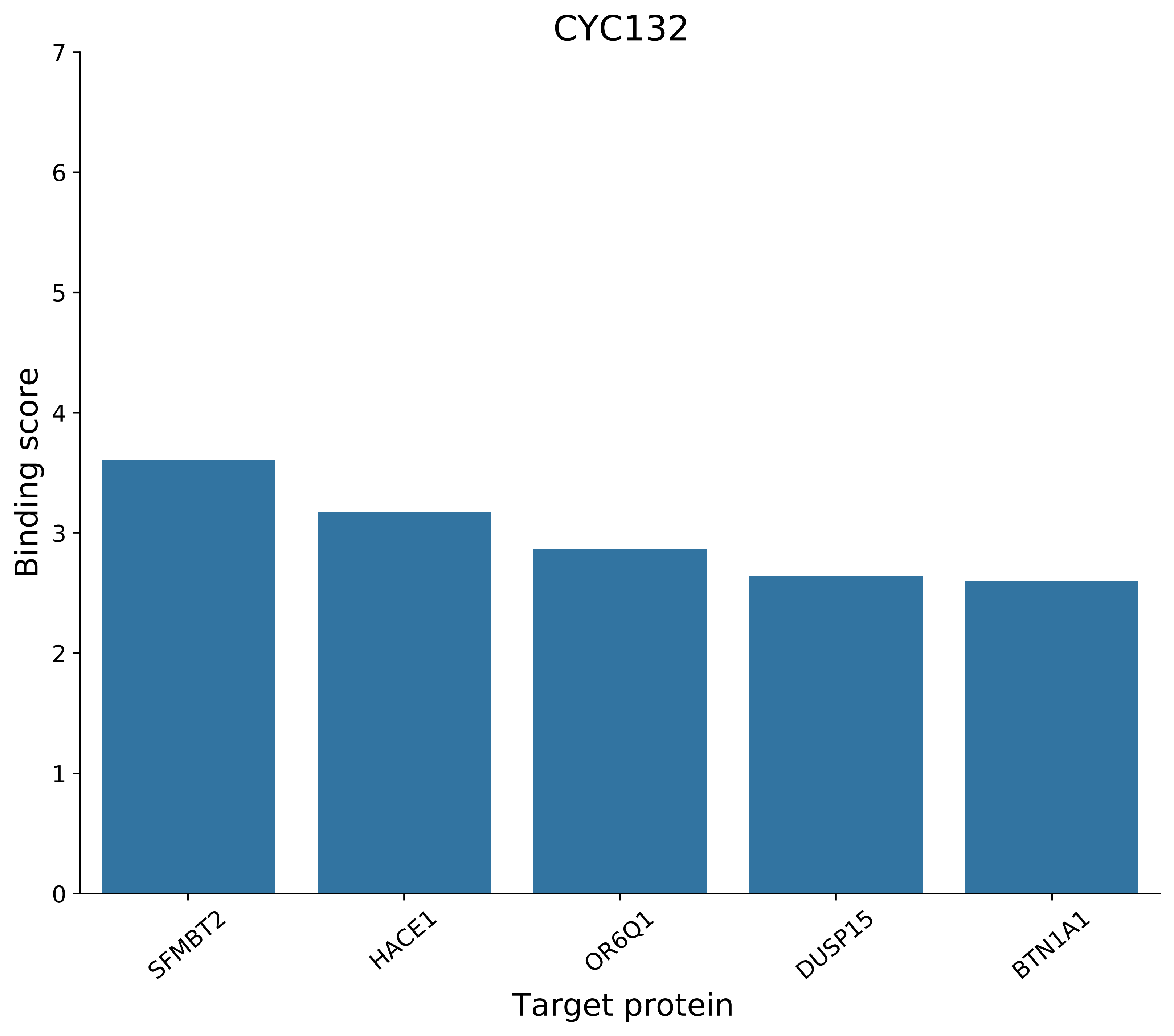
CYC132
Binding score
0
1
2
3
4
5
6
7
SFMBT2
HACE1
OR6Q1
DUSP15
BTN1A1
Target protein

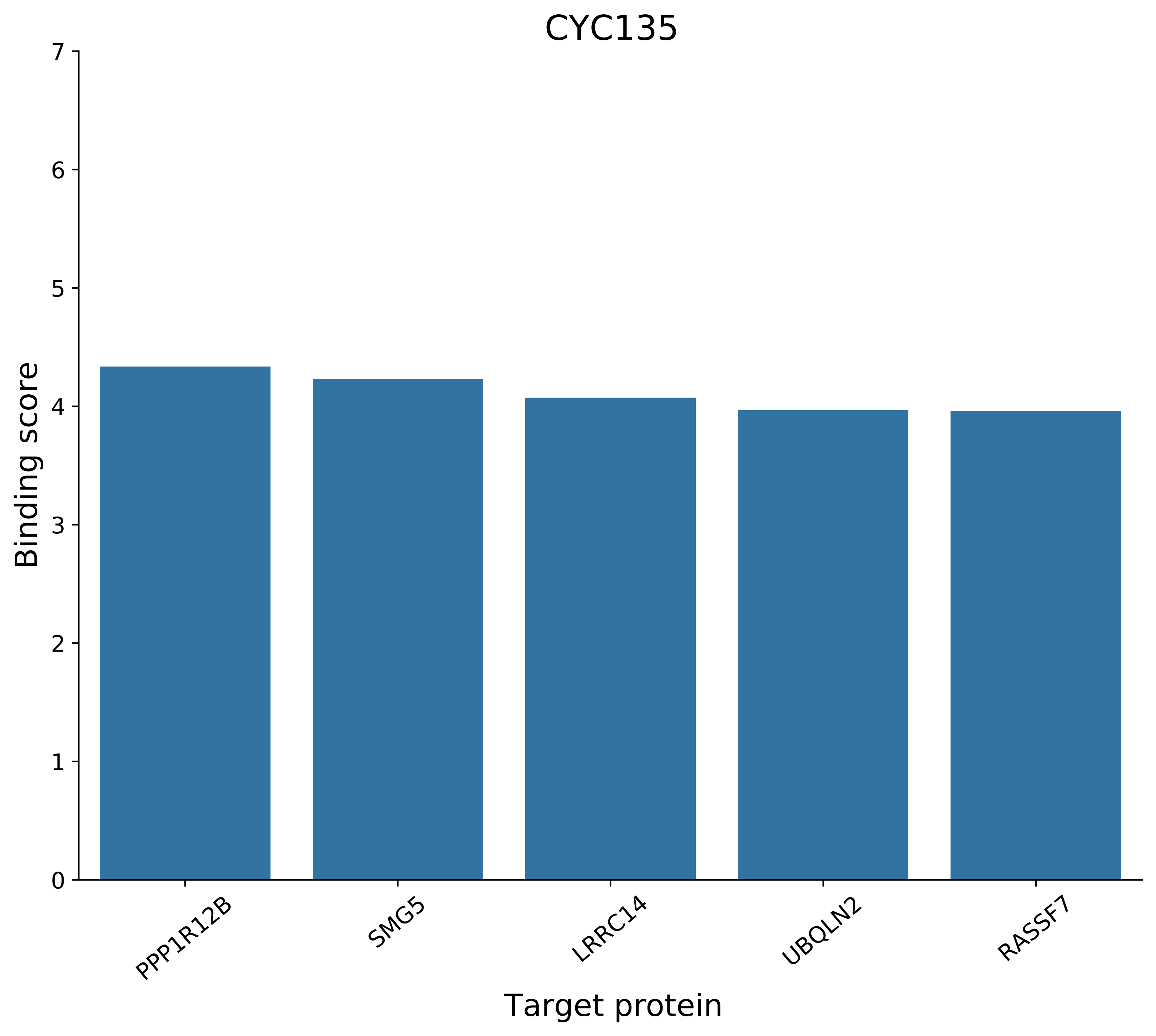
CYC135
Binding score
Target protein
0
1
2
3
4
5
6
7
PPP1R12B
SMG5
LRRC14
UBQLN2
RASSF7

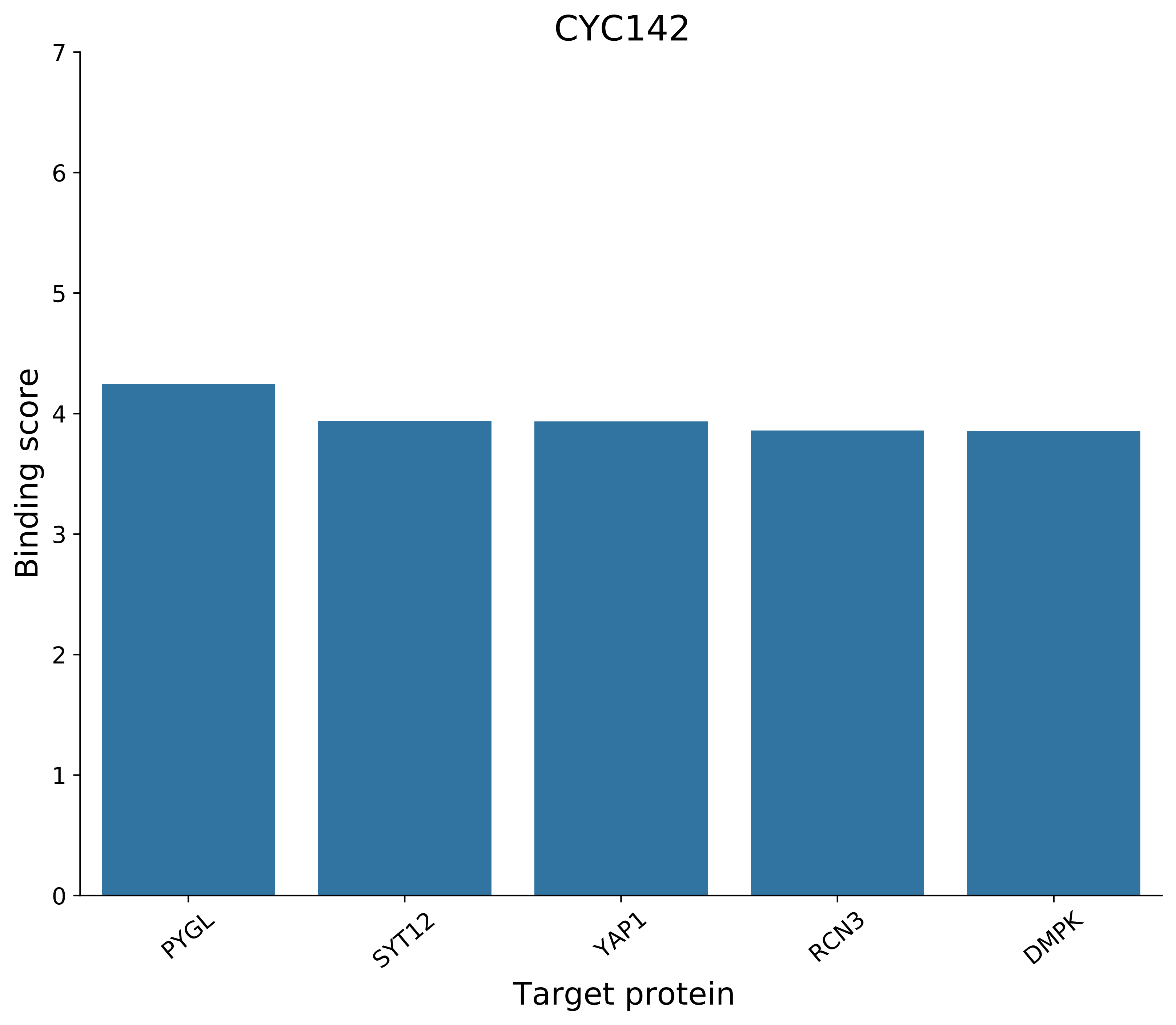
CYC142
Binding score
0
1
2
3
4
5
6
7
PYGL
SYT12
YAP1
RCN3
DMPK
Target protein

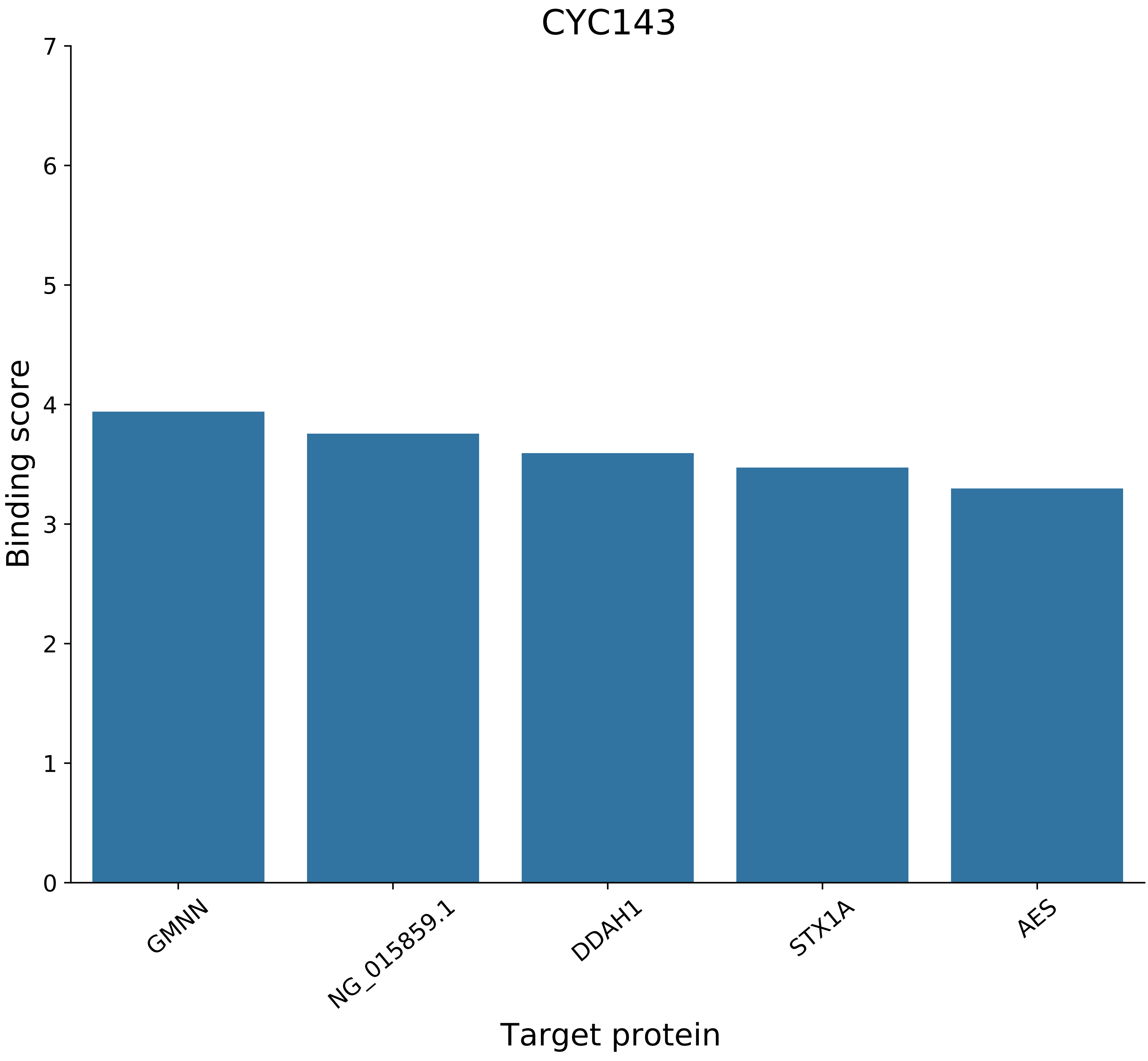

CYC143
Binding score
Target protein
0
1
2
3
4
5
6
7
GMNN
NG_015859.1
DDAH1
STX1A
AES

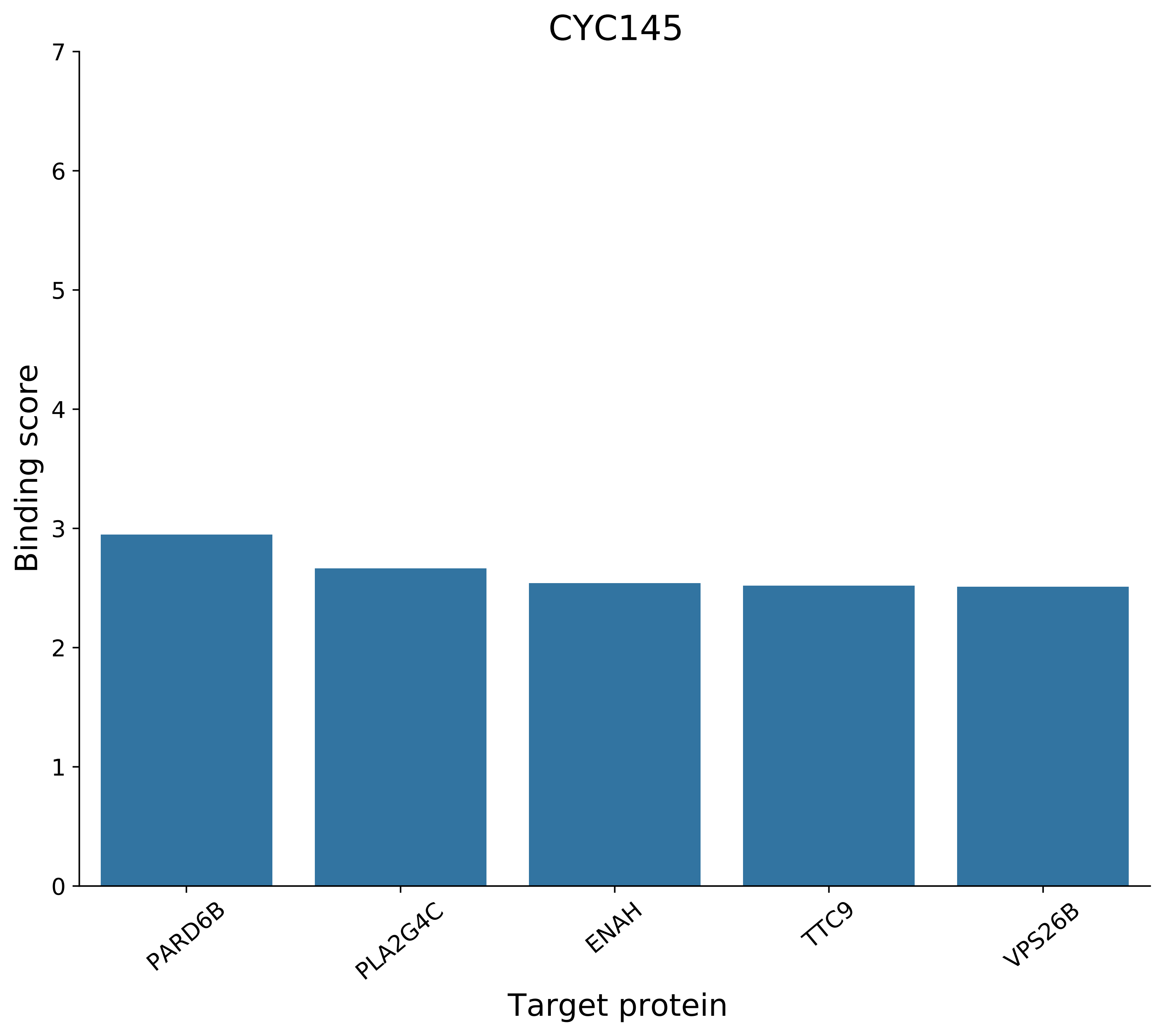
CYC145
Binding score
0
1
2
3
4
5
6
7
PARD6B
PLA2G4C
ENAH
TTC9
VPS26B
Target protein

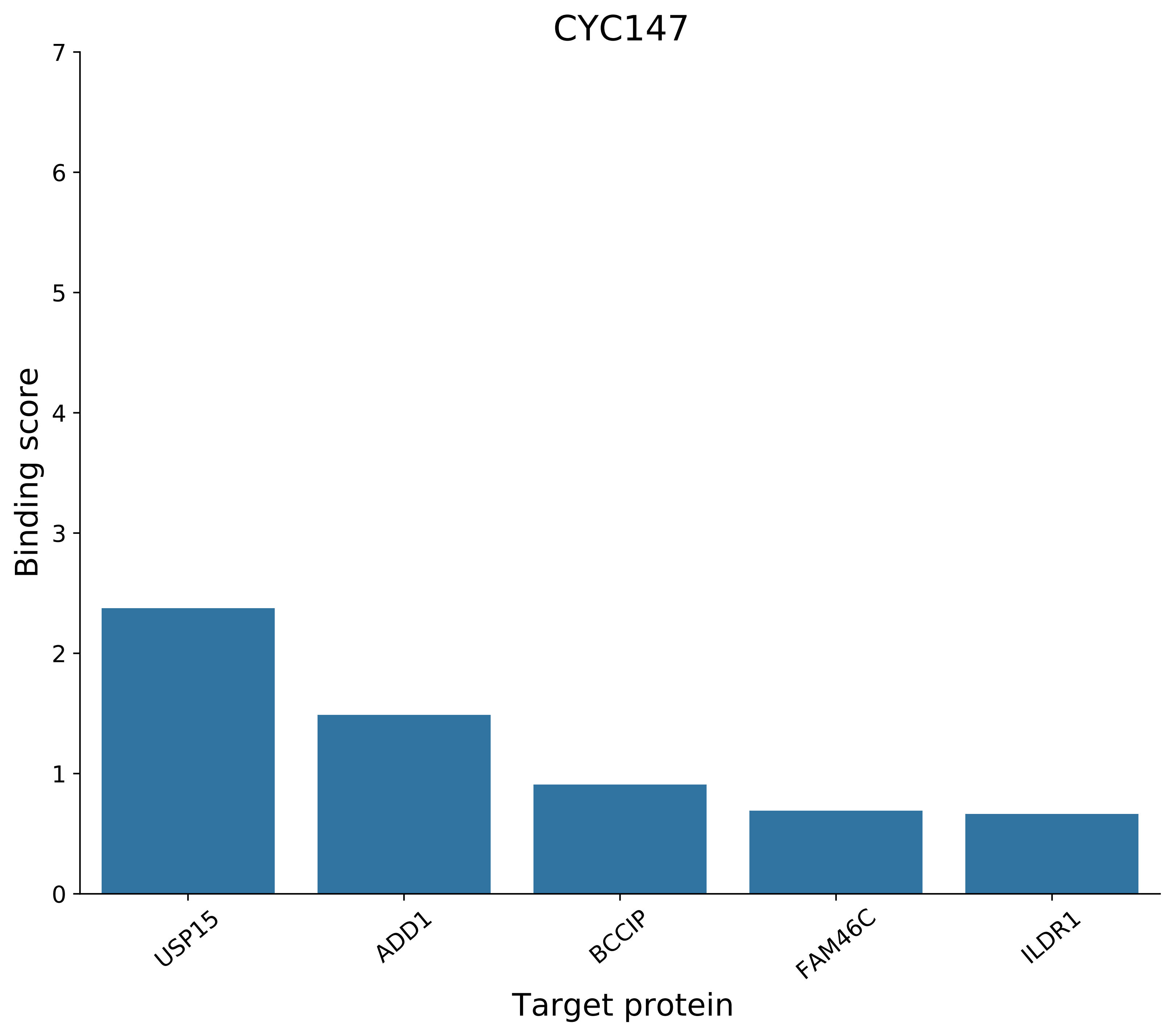
CYC147
Binding score
0
1
2
3
4
5
6
7
USP15
ADD1
BCCIP
FAM46C
ILDR1
Target protein

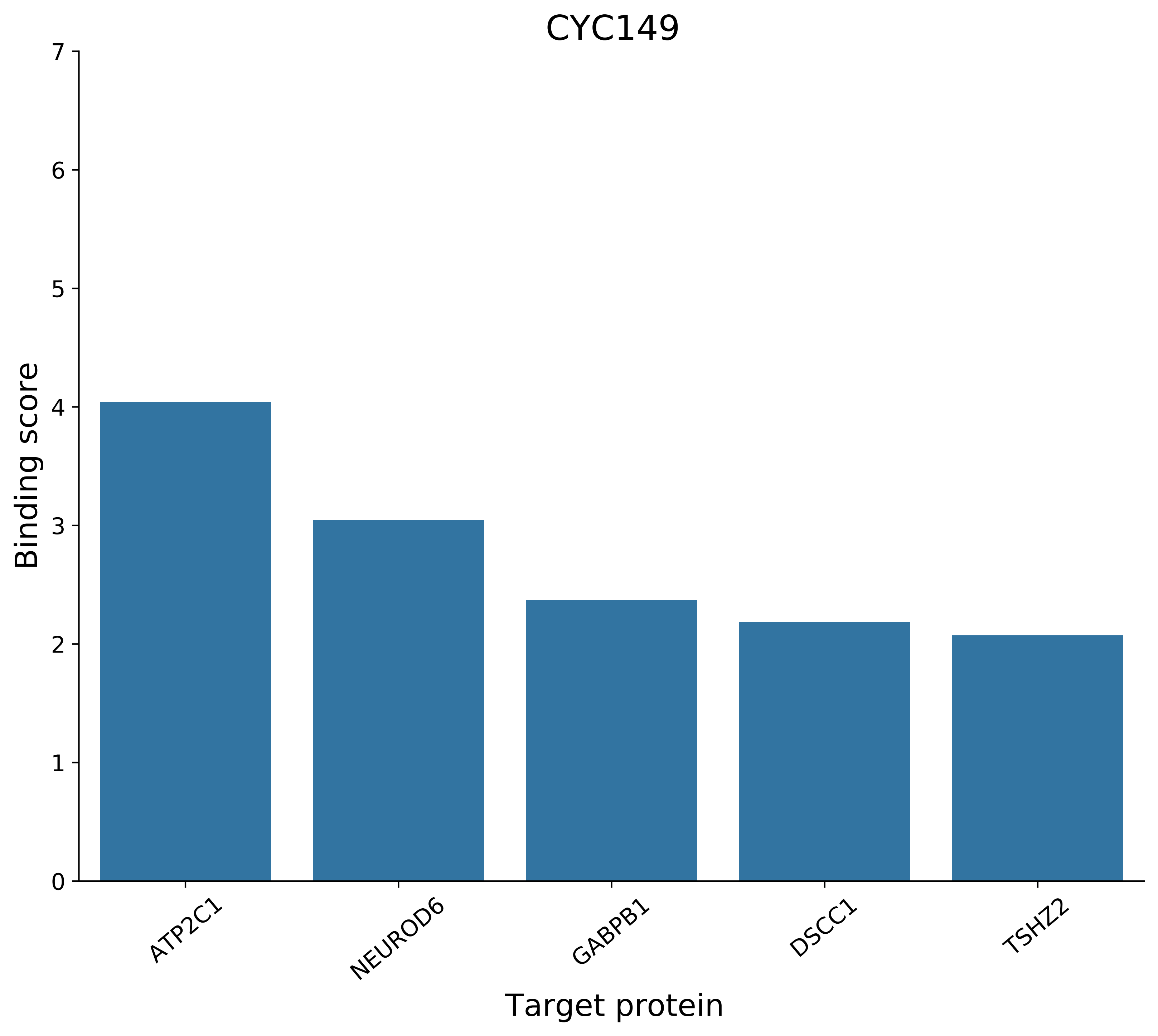

CYC149
Binding score
Target protein
0
1
2
3
4
5
6
7
ATP2C1
NEUROD6
GABPB1
DSCC1
TSHZ2

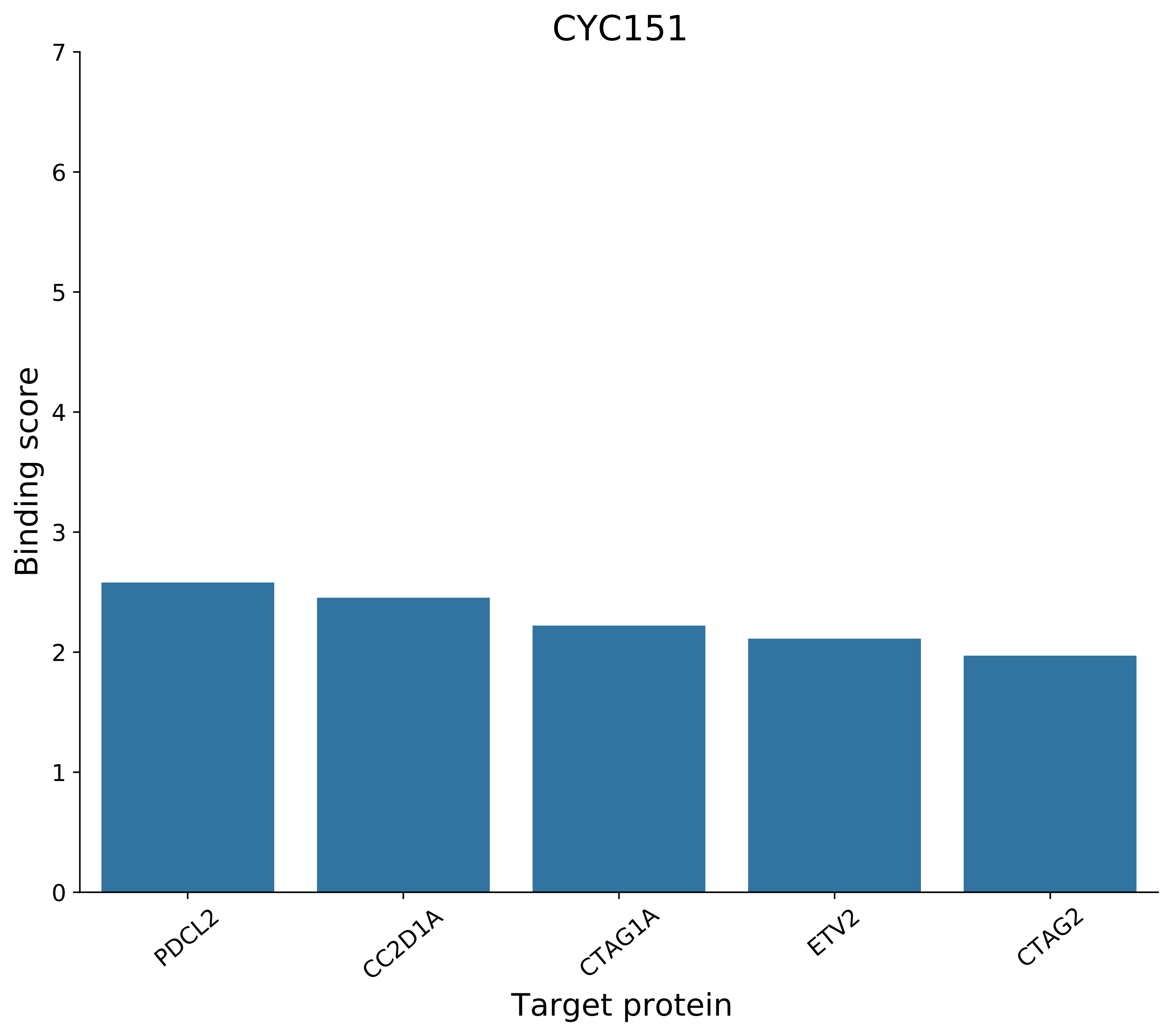

CYC151
Binding score
7
6
5
4
3
2
1
0
PDCL2
CC2D1A
CTAG1A
ETV2
CTAG2
Target protein

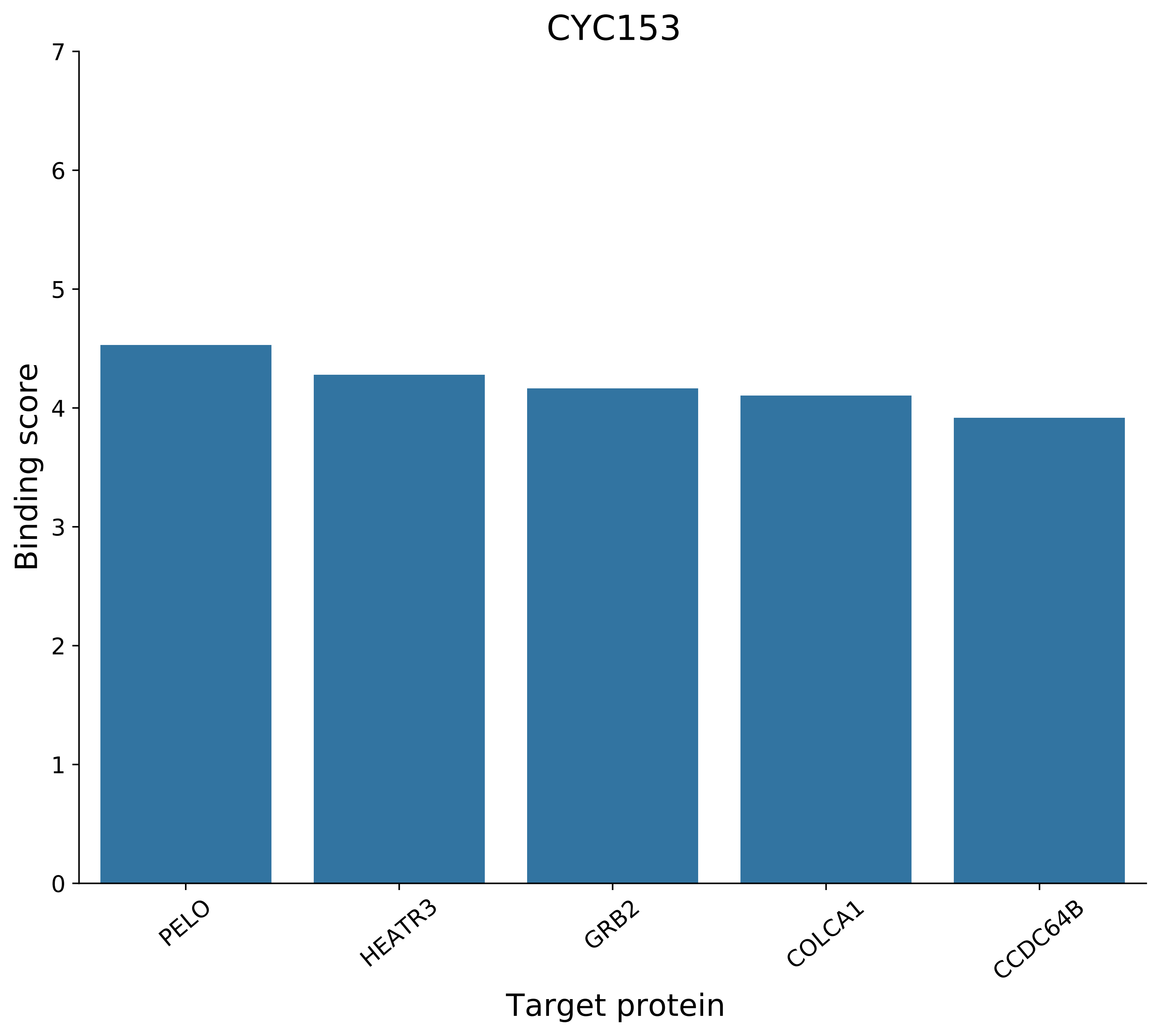
CYC153
Binding score
Target protein
0
1
2
3
4
5
6
7
PELO
HEATR3
GRB2
COLCA1
CCDC64B

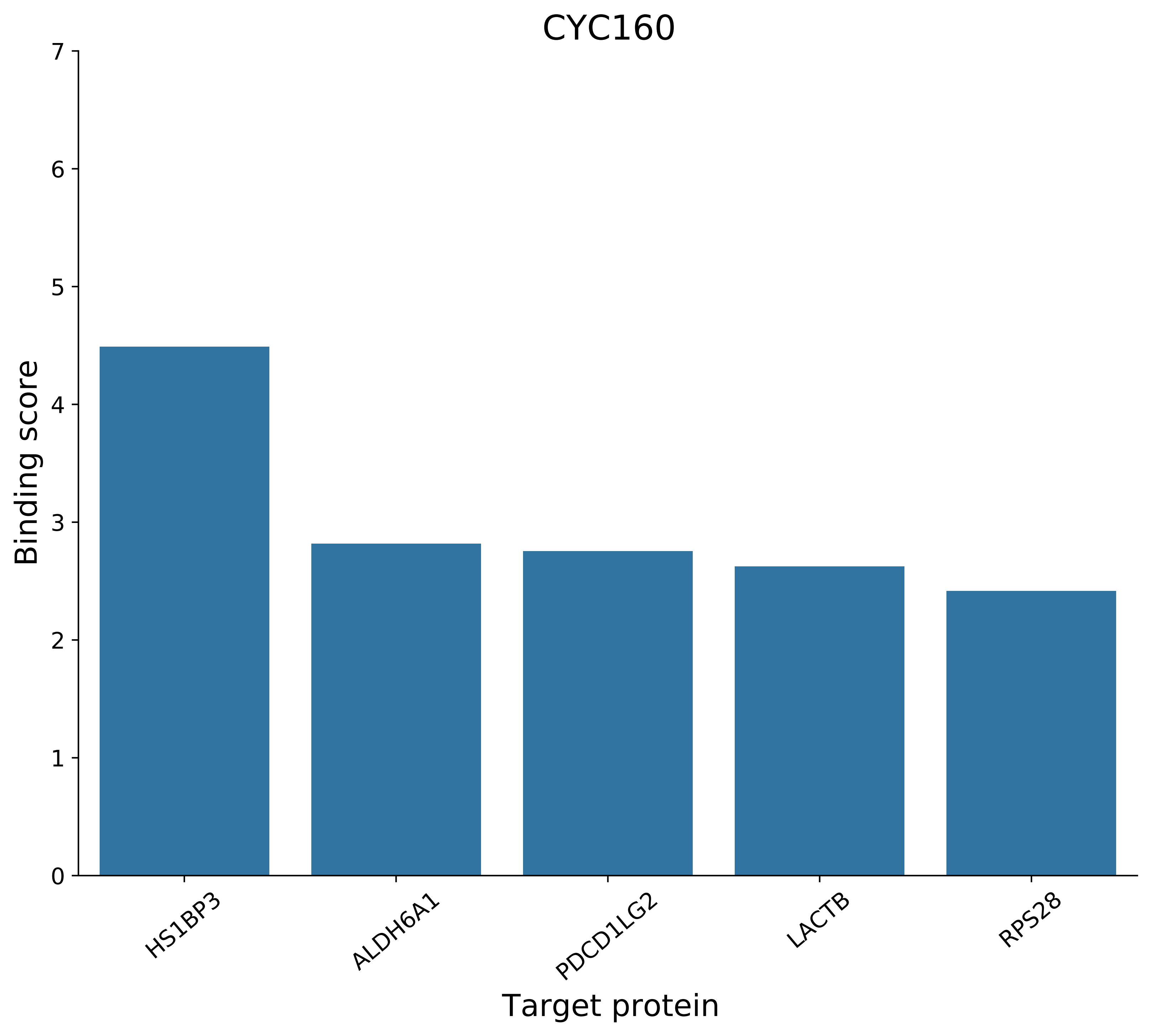

CYC160
Binding score
0
1
2
3
4
5
6
7
HS1BP3
ALDH6A1
PDCD1LG2
LACTB
RPS28
Target protein

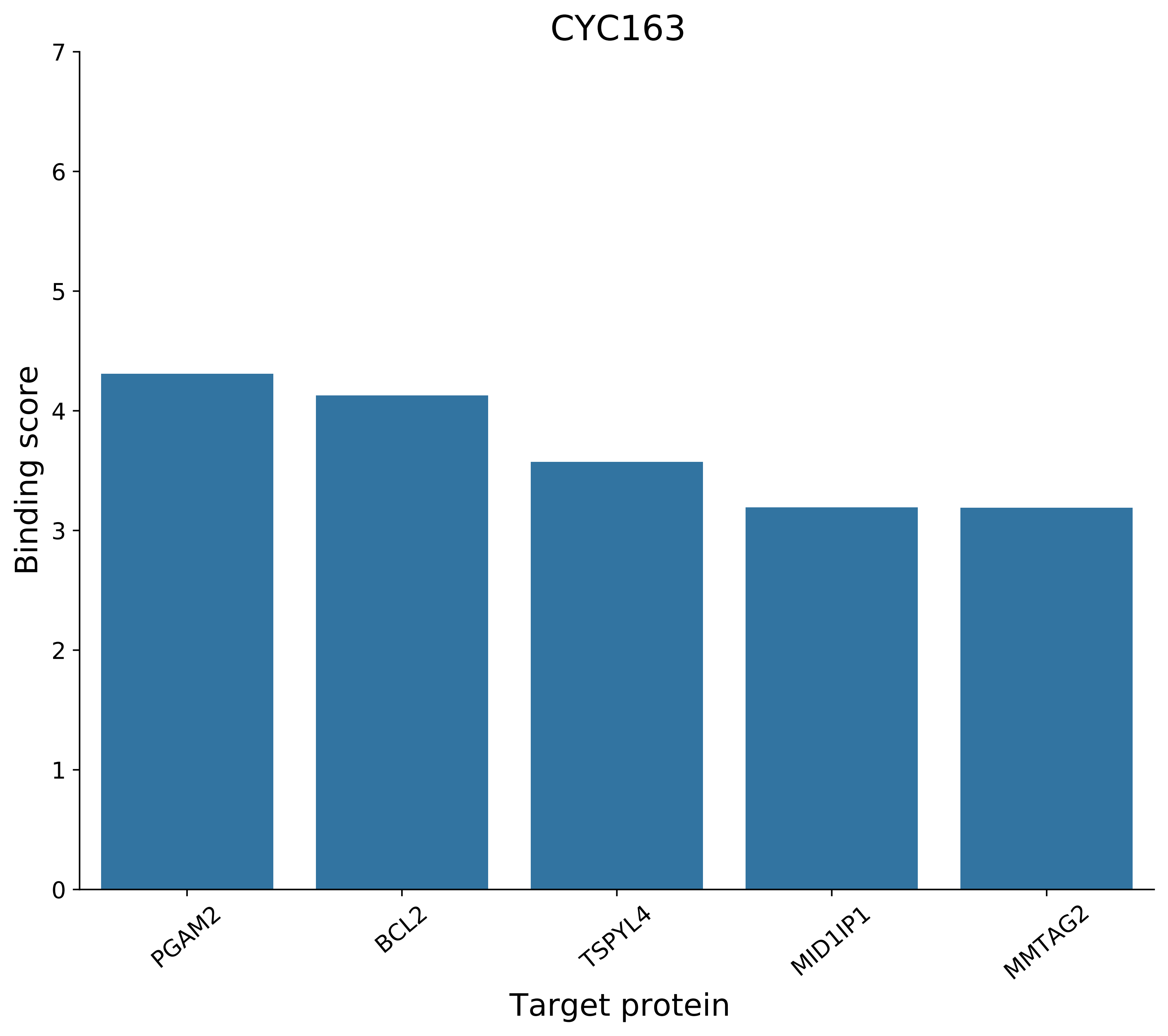
CYC163
Binding score
0
1
2
3
4
5
6
7
PGAM2
BCL2
TSPYL4
MID1IP1
MMTAG2
Target protein

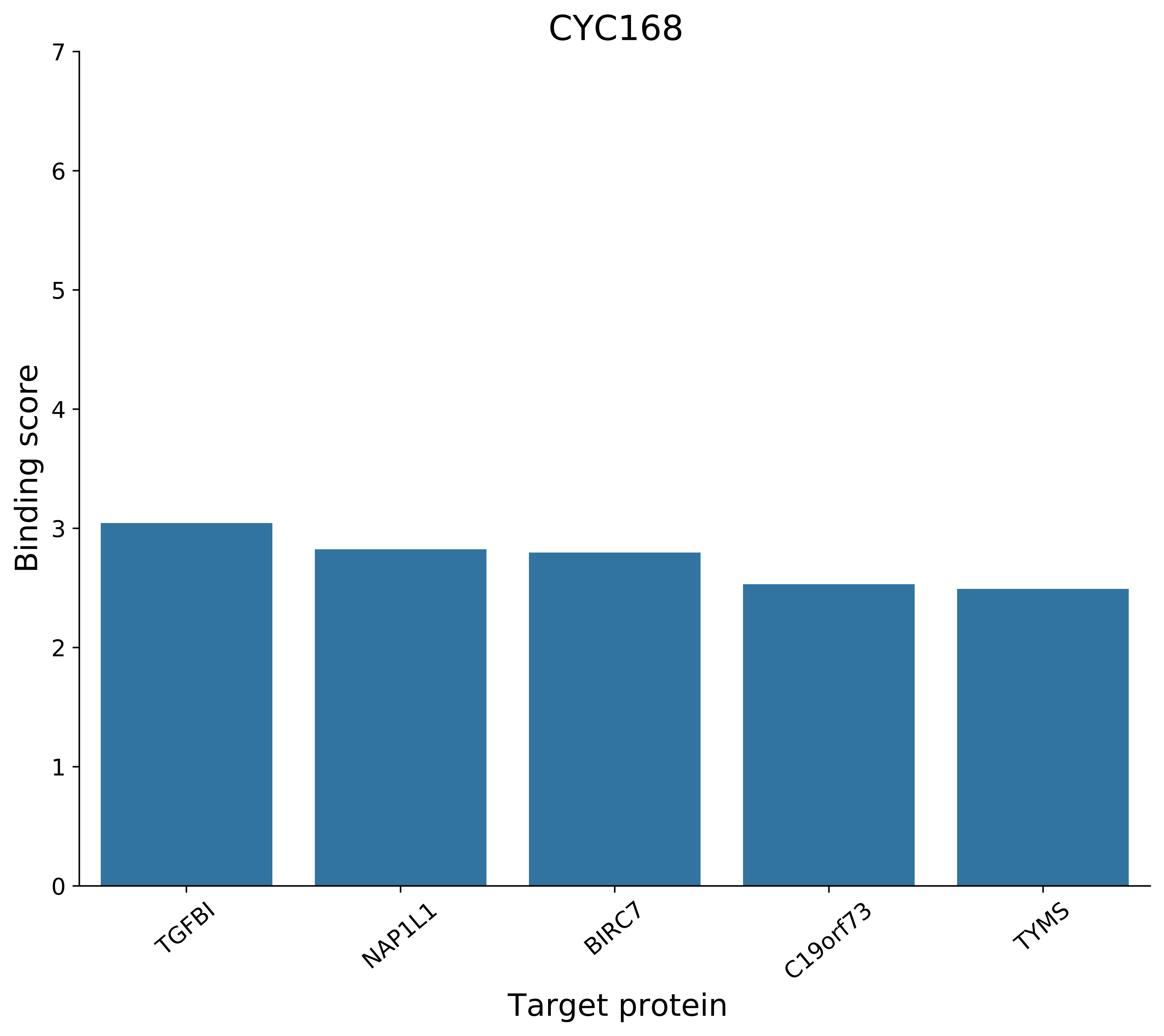
CYC168
Binding score
Target protein
0
1
2
3
4
5
6
7
TGFBI
NAP1L1
BIRC7
C19orf73
TYMS

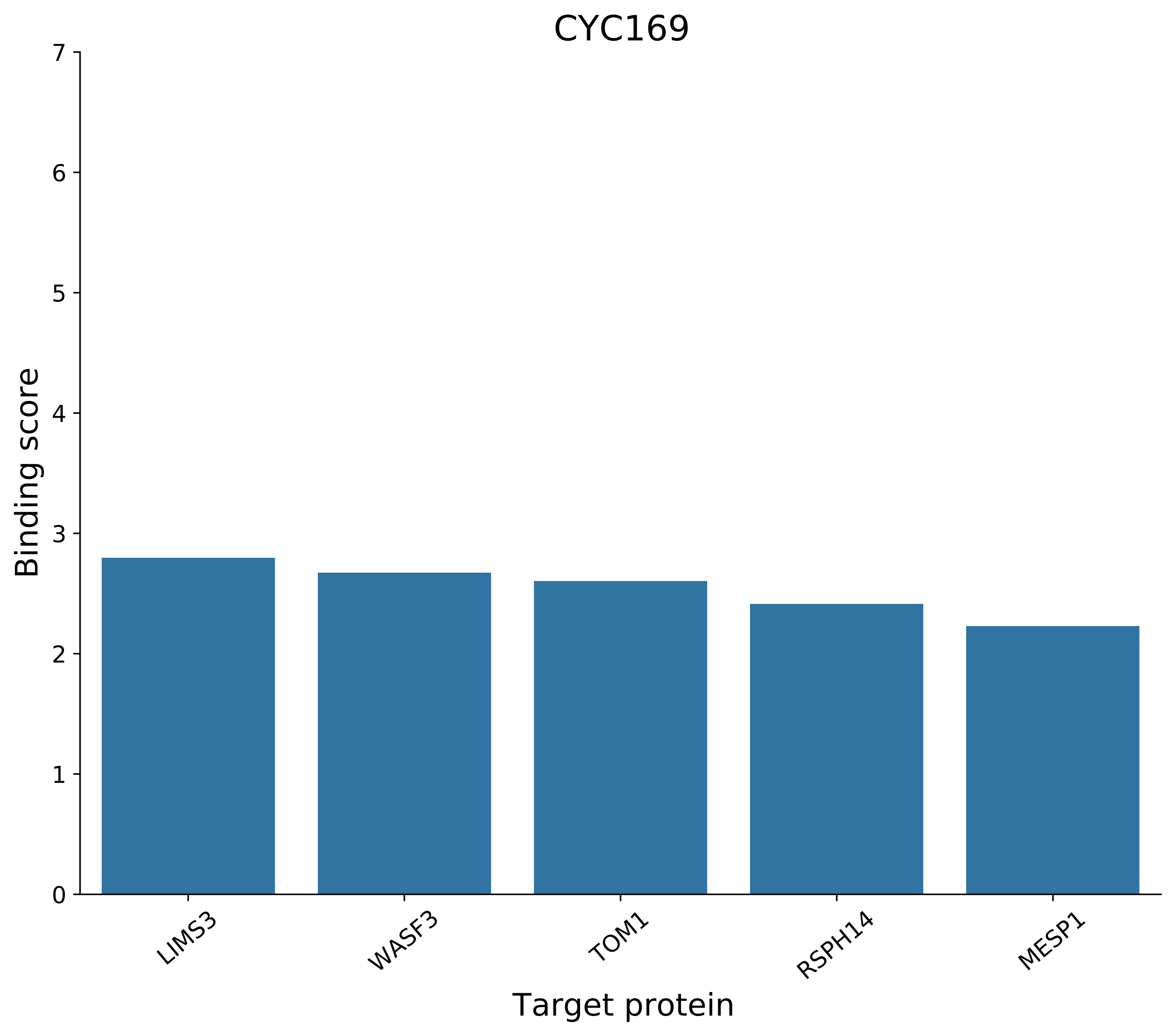
CYC169
Binding score
0
1
2
3
4
5
6
7
LIMS3
WASF3
TOM1
RSPH14
MESP1
Target protein

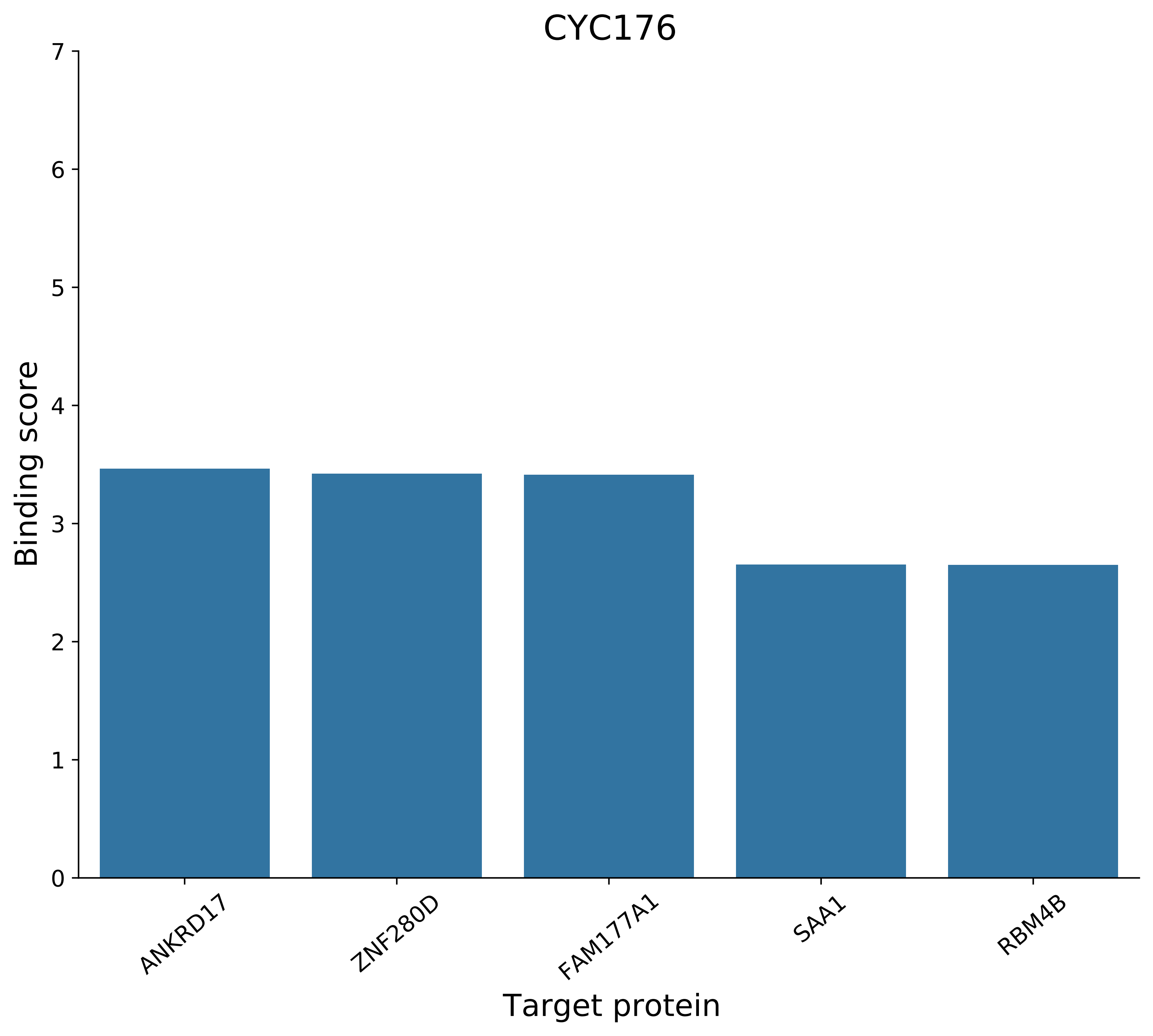

CYC176
Binding score
0
1
2
3
4
5
6
7
ANKRD17
ZNF280D
FAM177A1
SAA1
RBM4B
Target protein

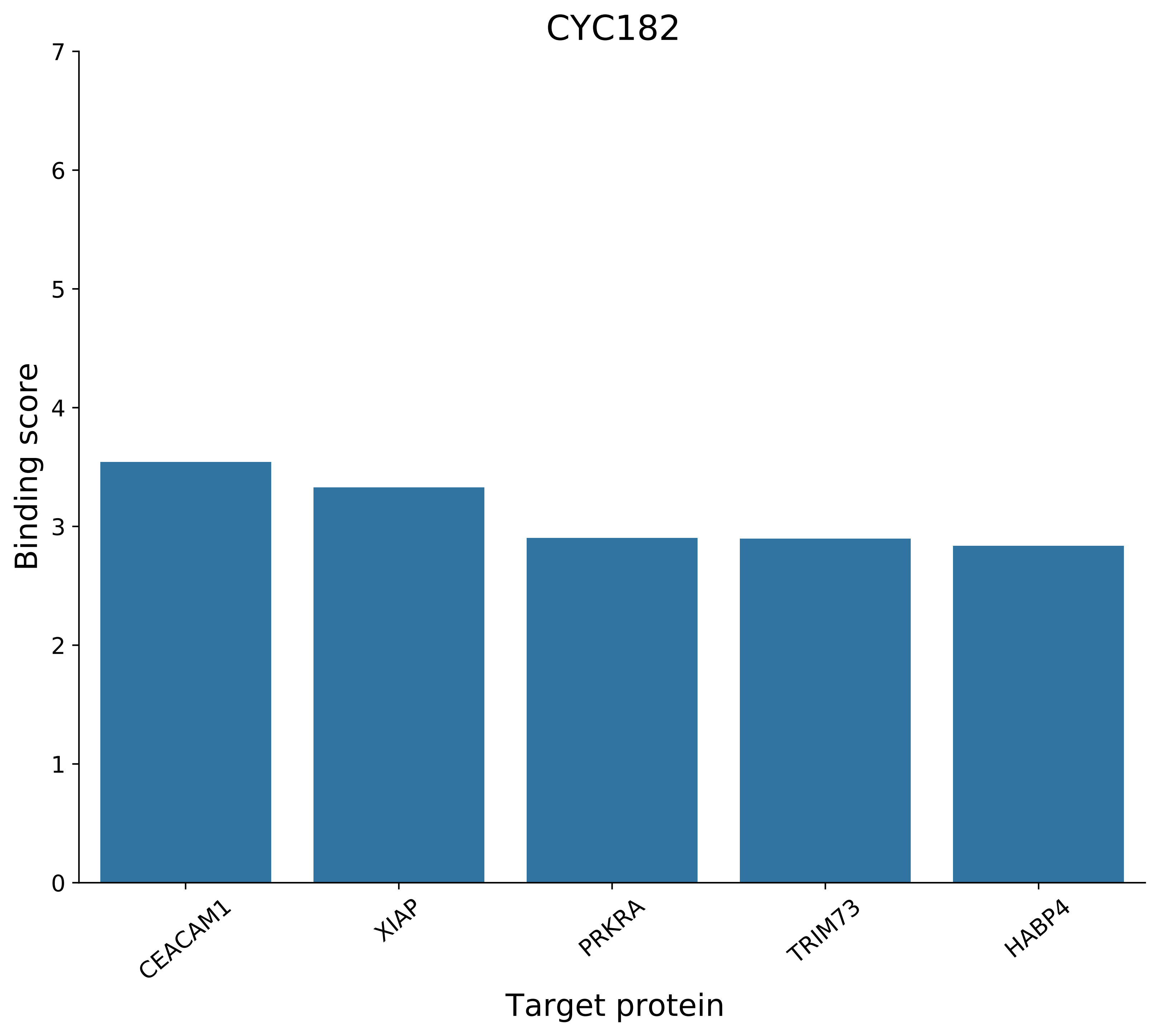

CYC182
Binding score
Target protein
0
1
2
3
4
5
6
7
CEACAM1
XIAP
PRKRA
TRIM73
HABP4

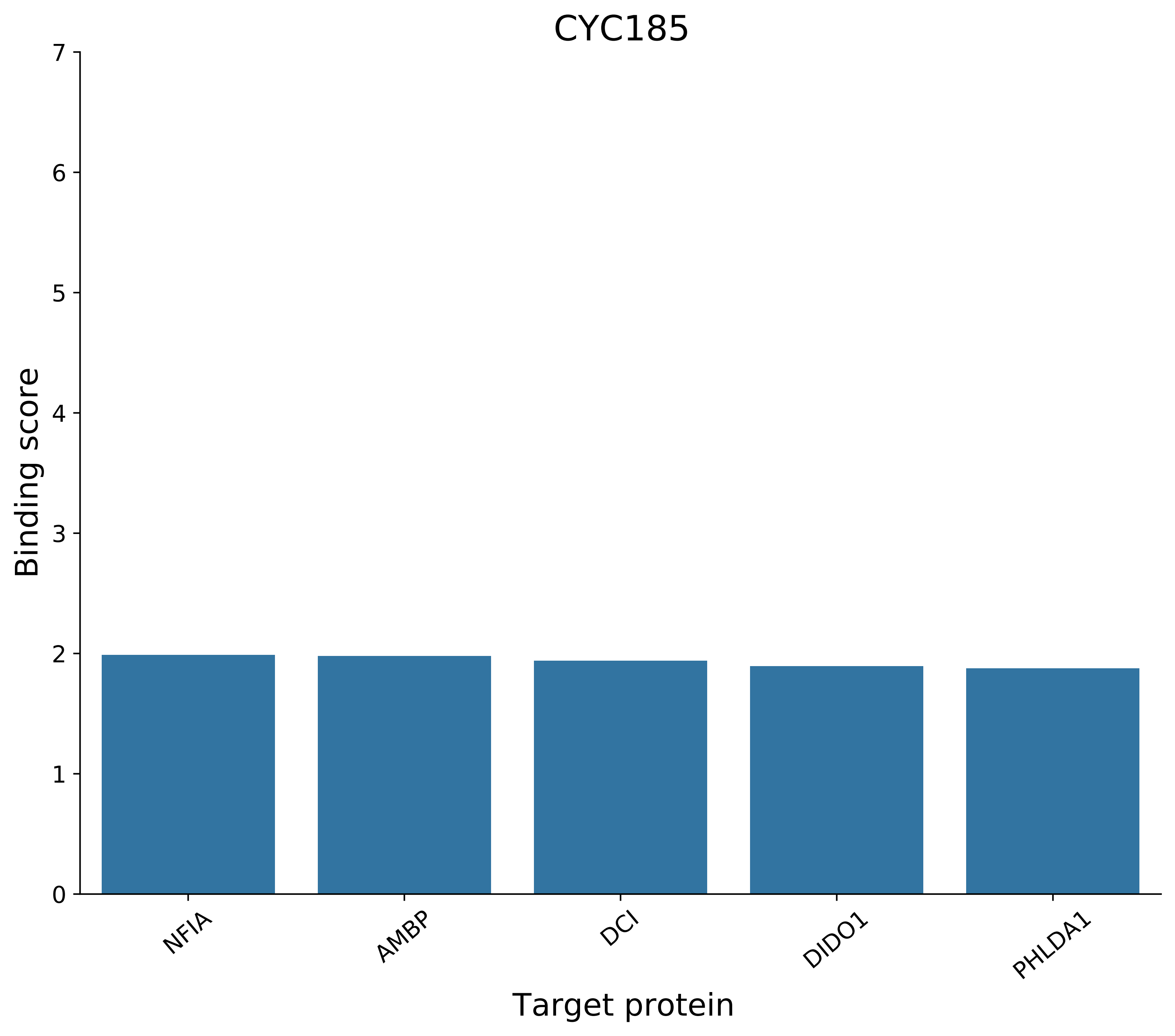
CYC185
Binding score
Target protein
0
1
2
3
4
5
6
7
NFIA
AMBP
DCI
DIDO1
PHLDA1

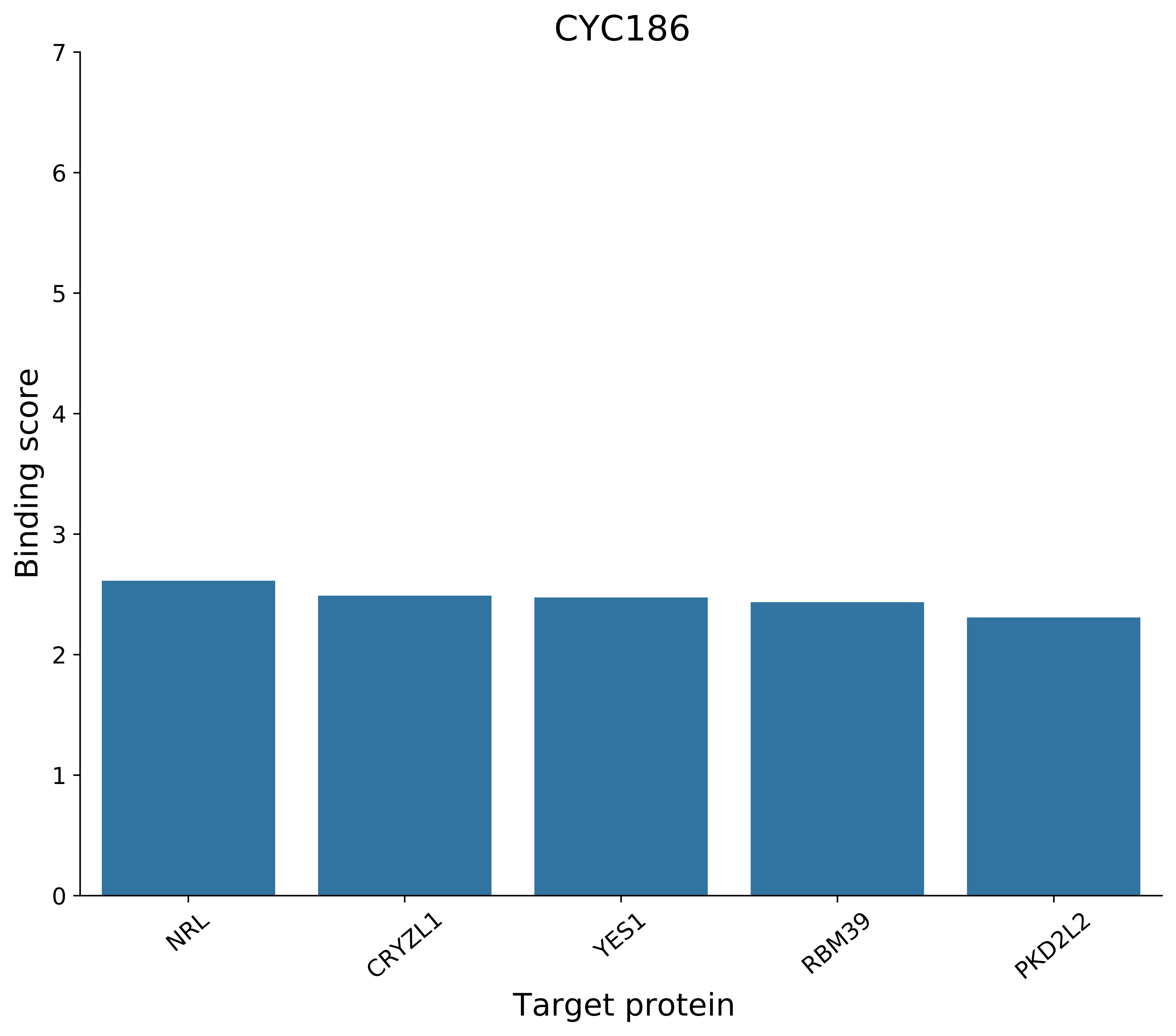
CYC186
Binding score
Target protein
0
1
2
3
4
5
6
7
NRL
CRYZL1
YES1
RBM39
PKD2L2

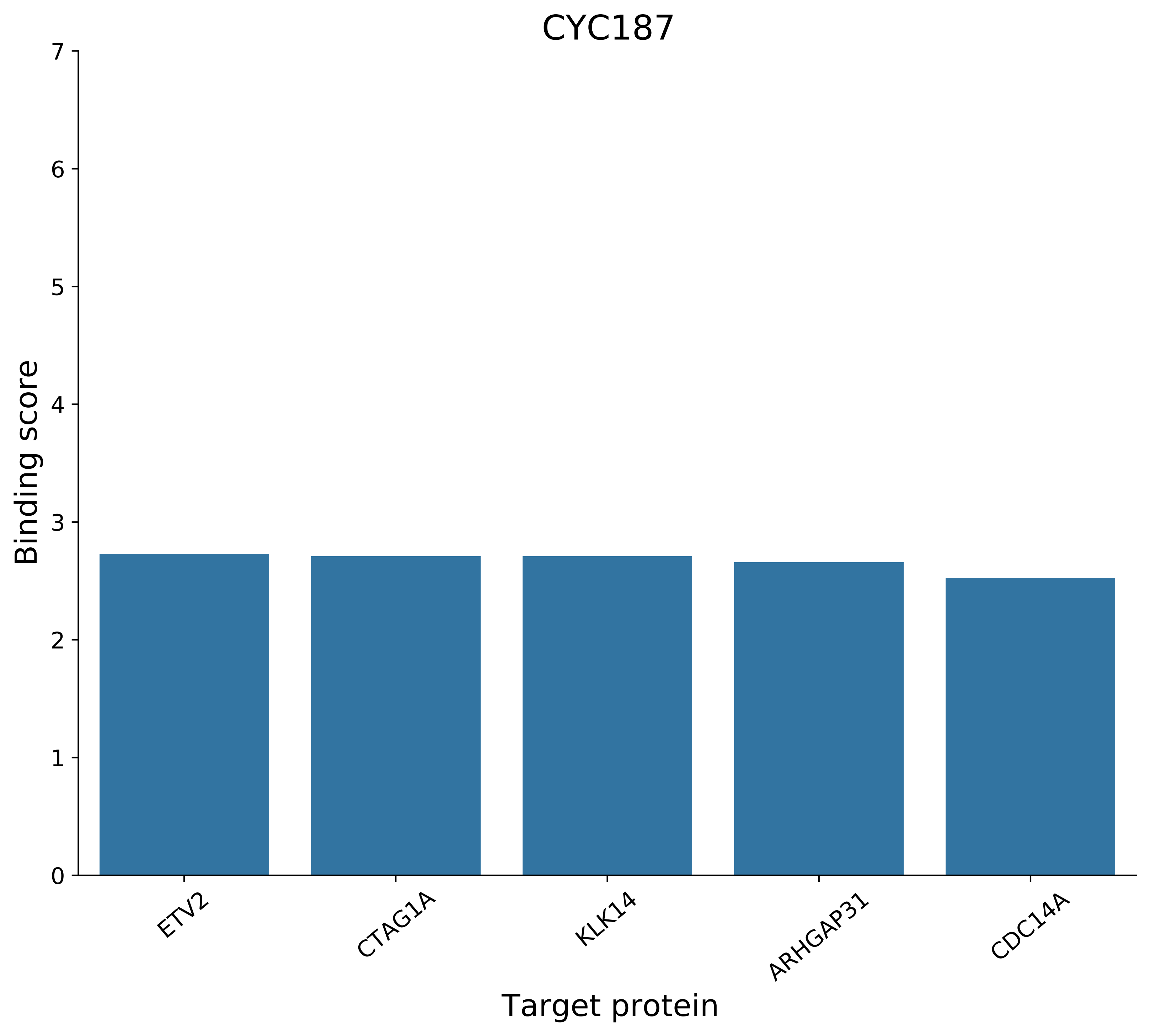
CYC187
Binding score
0
1
2
3
4
5
6
7
ETV2
CTAG1A
KLK14
ARHGAP31
CDC14A
Target protein

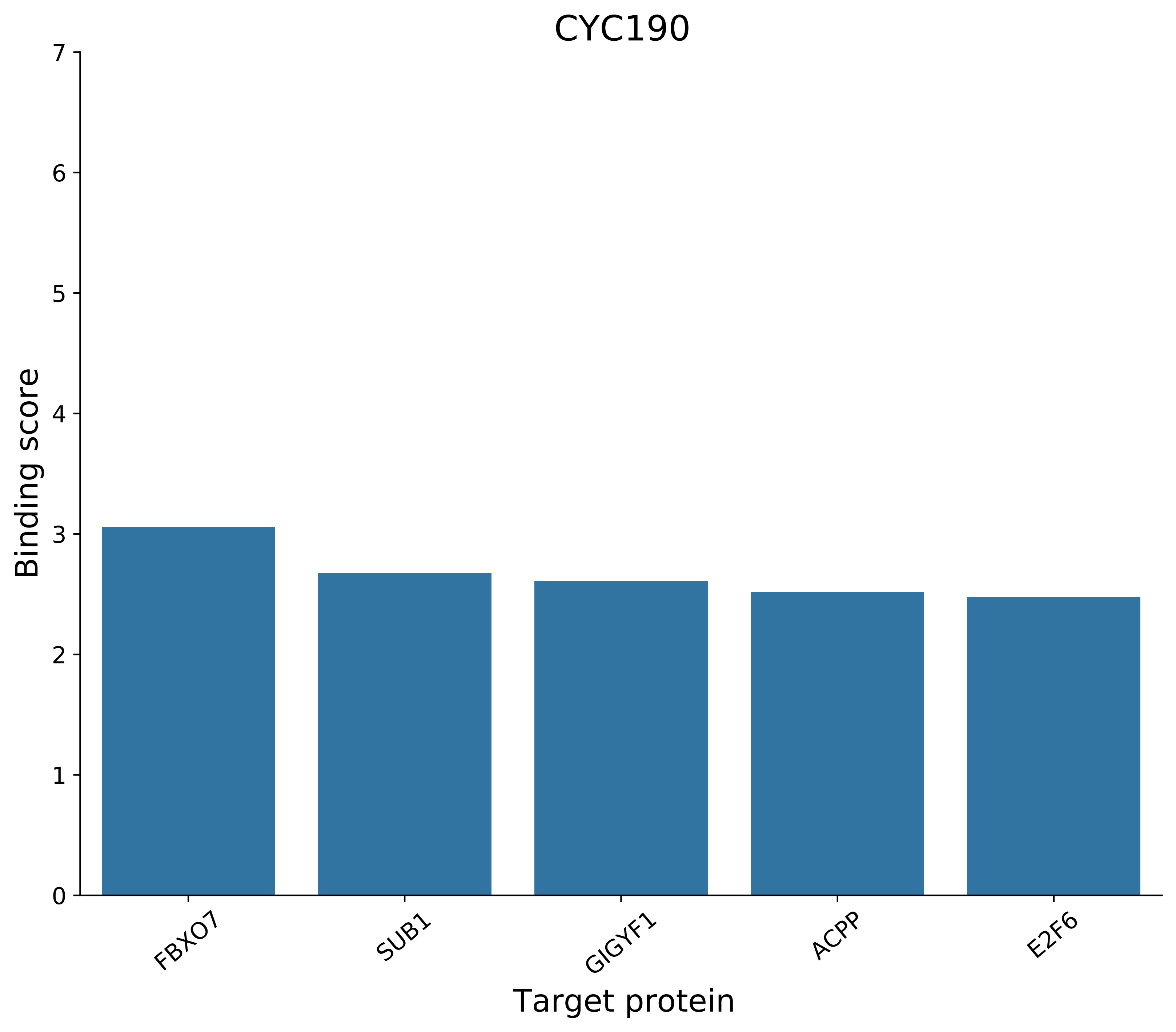
CYC190
Binding score
0
1
2
3
4
5
6
7
FBXO7
SUB1
GIGYF1
ACPP
E2F6
Target protein

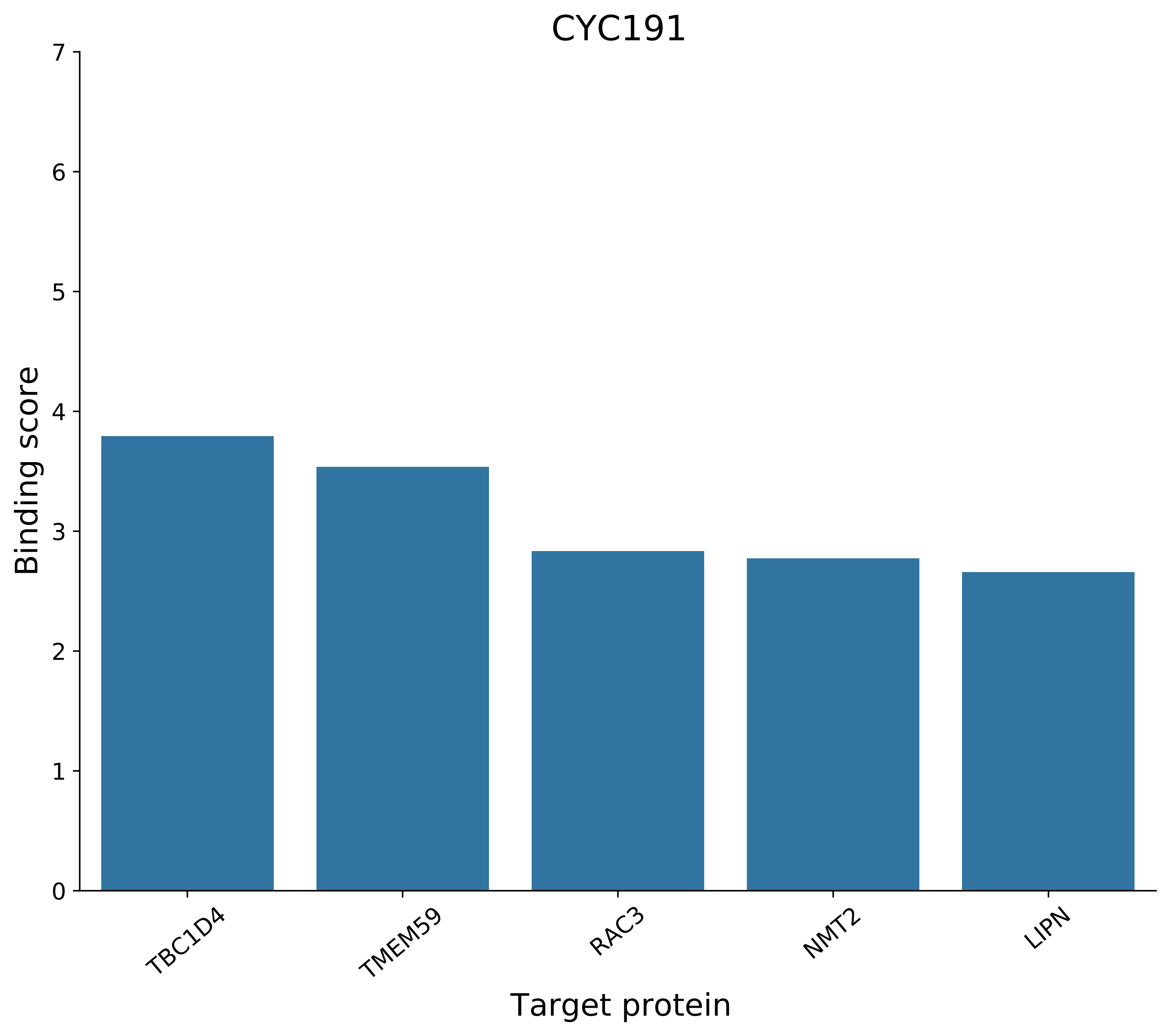
CYC191
Binding score
0
1
2
3
4
5
6
7
TBC1D4
TMEM59
RAC3
NMT2
LIPN
Target protein

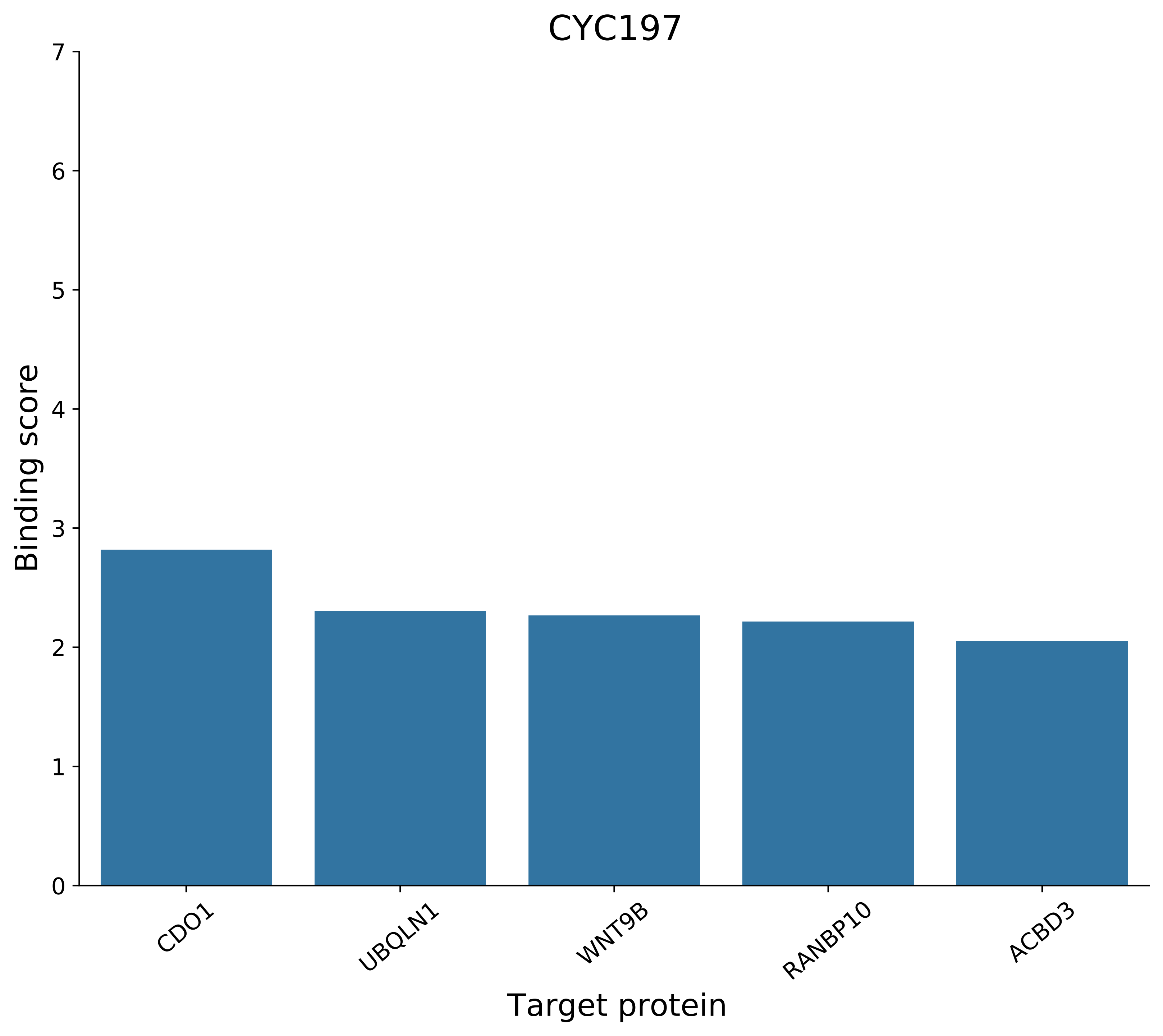
CYC197
Binding score
0
1
2
3
4
5
6
7
CDO1
UBQLN1
WNT9B
RANBP10
ACBD3
Target protein

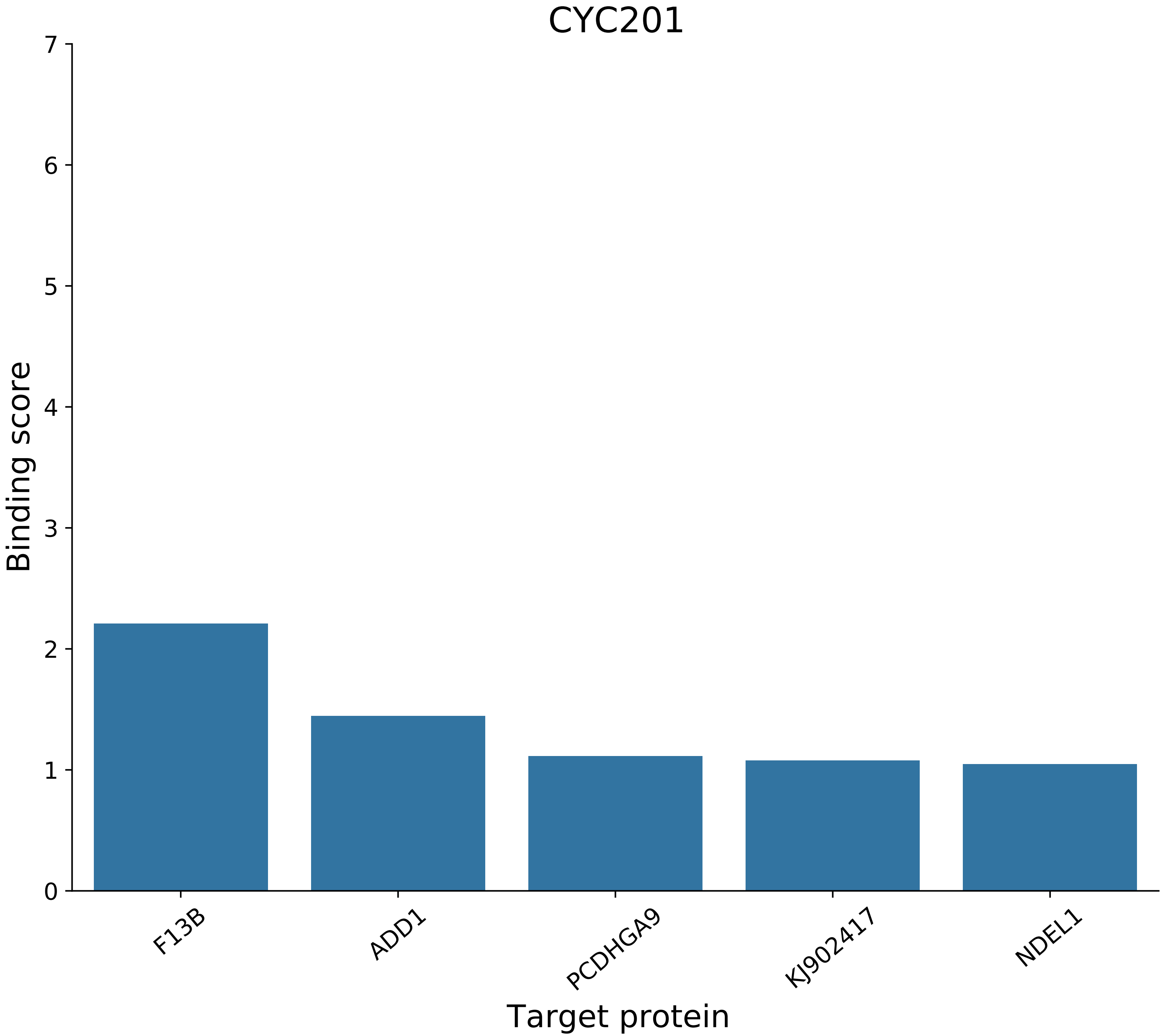
CYC201
Binding score
Target protein
0
1
2
3
4
5
6
7
F13B
ADD1
PCDHGA9
KJ902417
NDEL1

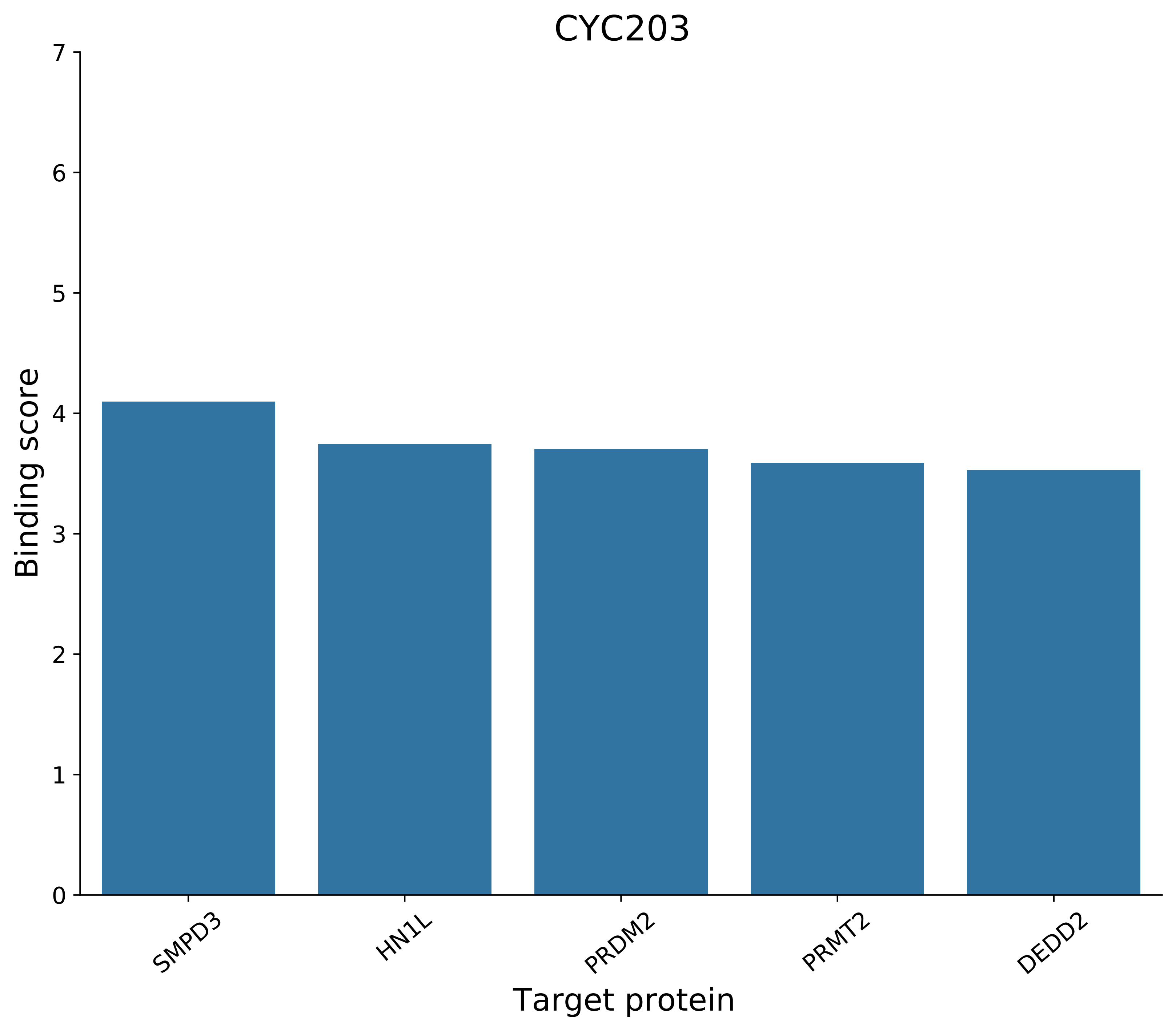
CYC203
Binding score
0
1
2
3
4
5
6
7
SMPD3
HN1L
PRDM2
PRMT2
DEDD2
Target protein

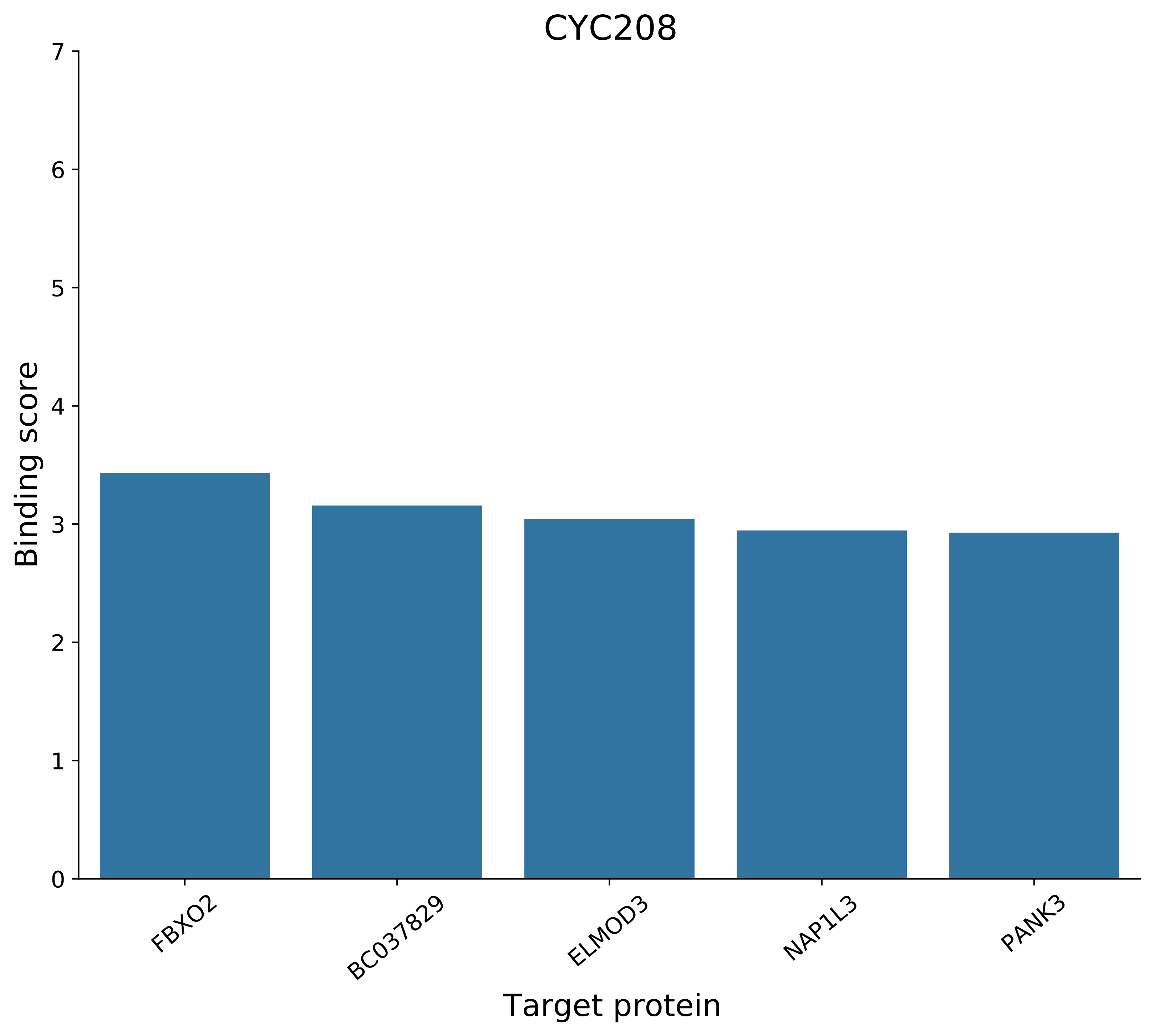

CYC208
Binding score
0
1
2
3
4
5
6
7
FBXO2
BC037829
ELMOD3
NAP1L3
PANK3
Target protein

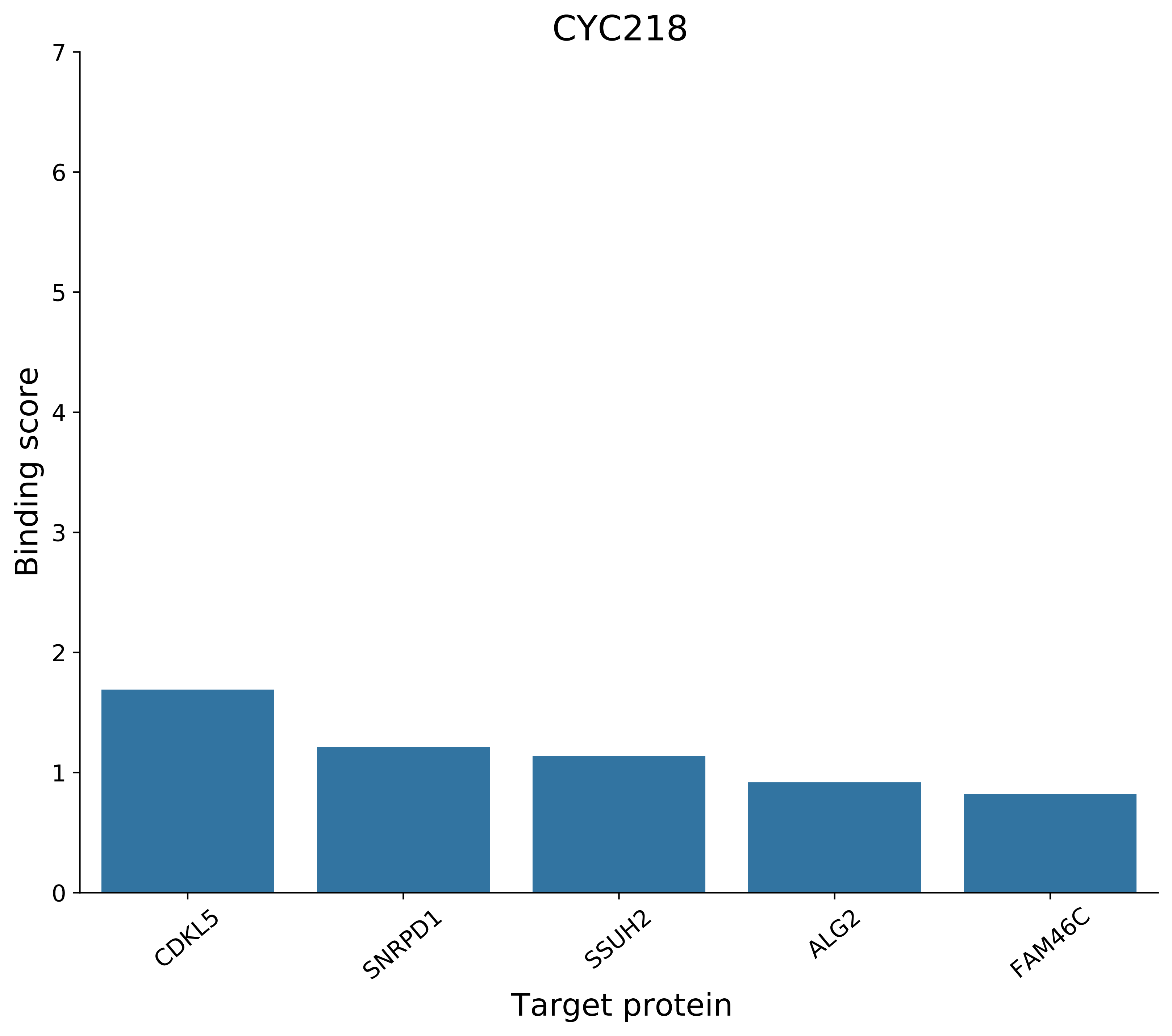

CYC218
Binding score
0
1
2
3
4
5
6
7
CDKL5
SNRPD1
SSUH2
ALG2
FAM46C
Target protein

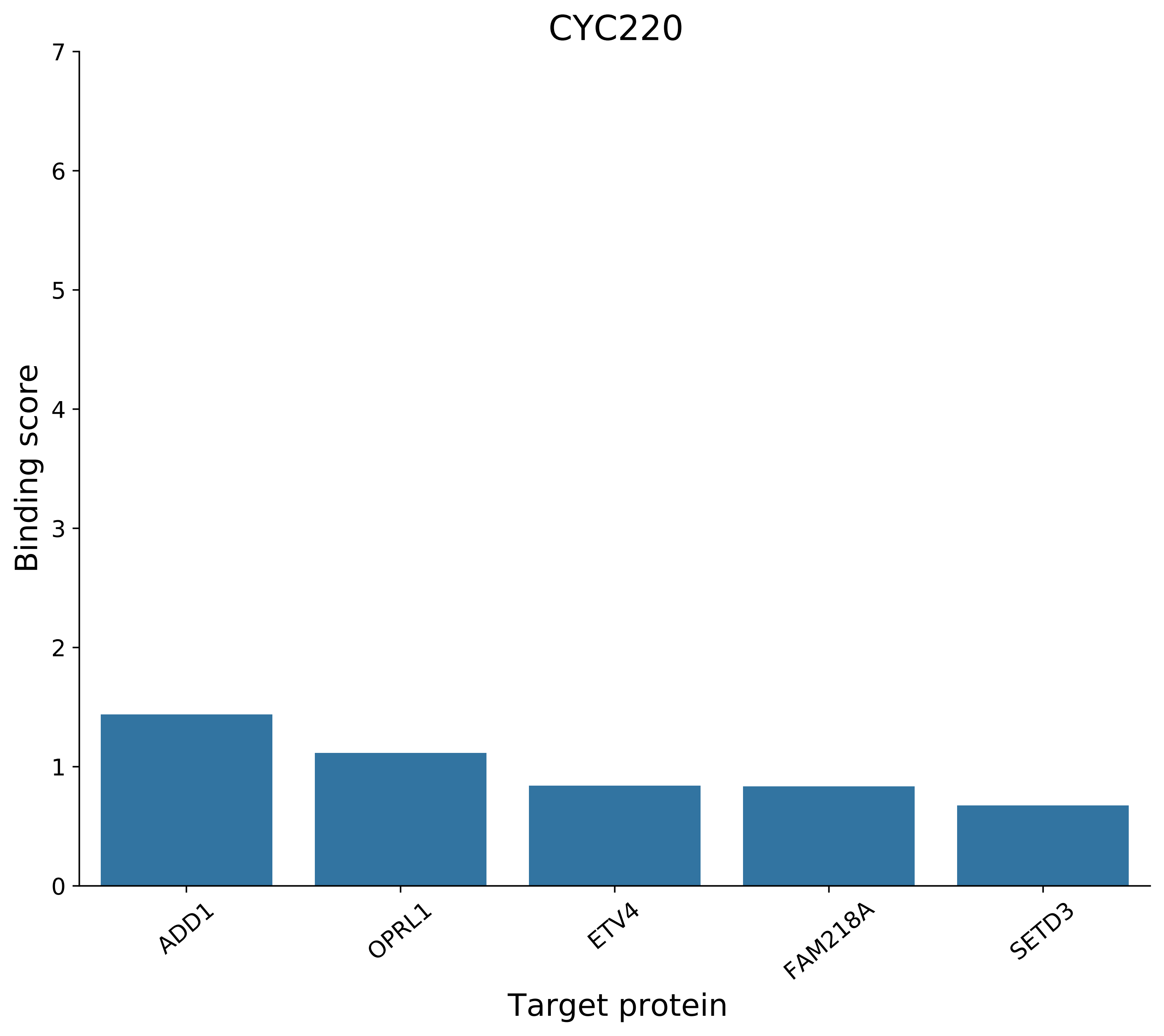
CYC220
Binding score
0
1
2
3
4
5
6
7
ADD1
OPRL1
ETV4
FAM218A
SETD3
Target protein

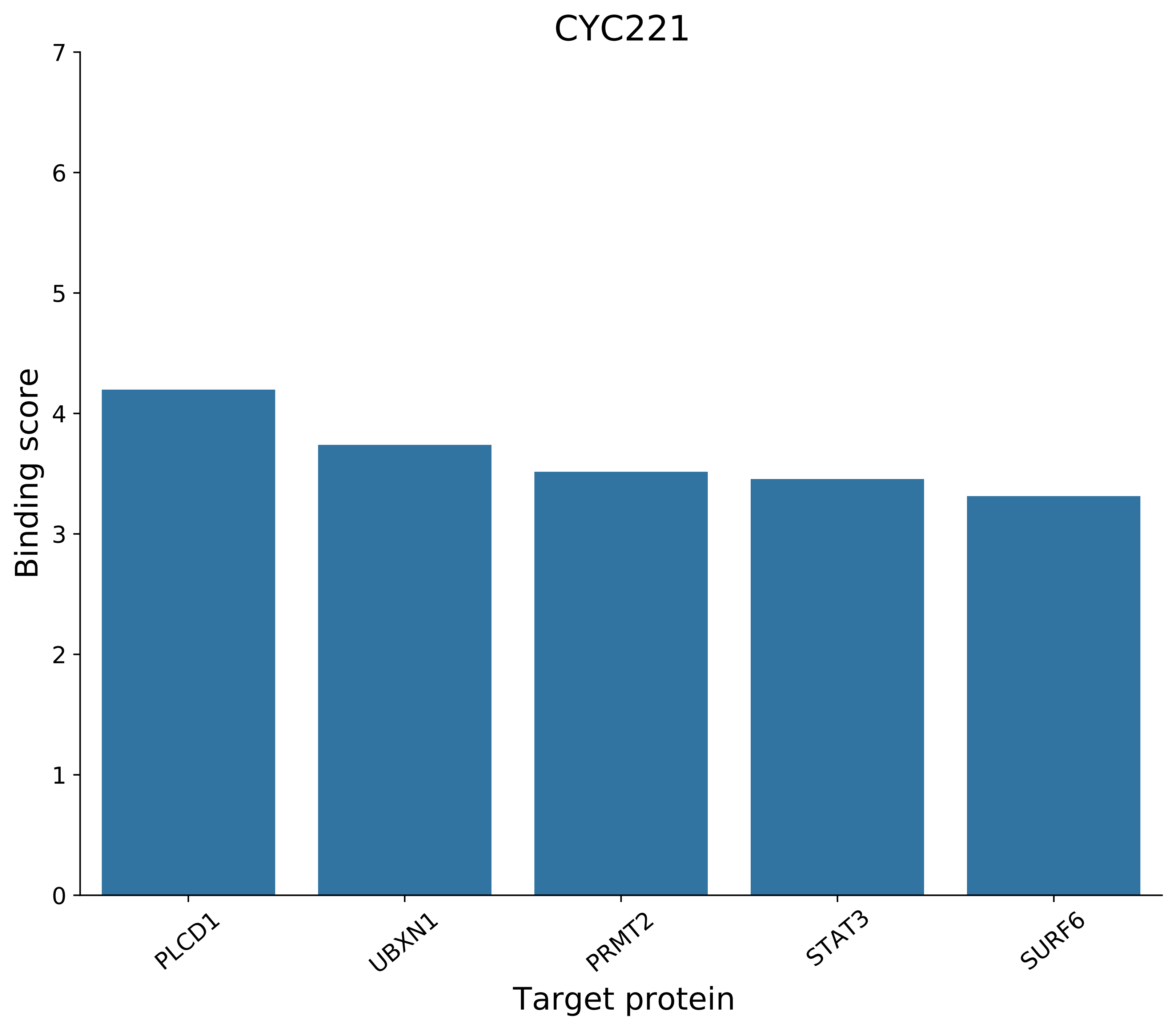
CYC221
7
6
5
4
3
2
1
0
Binding score
PLCD1
UBXN1
PRMT2
STAT3
SURF6
Target protein

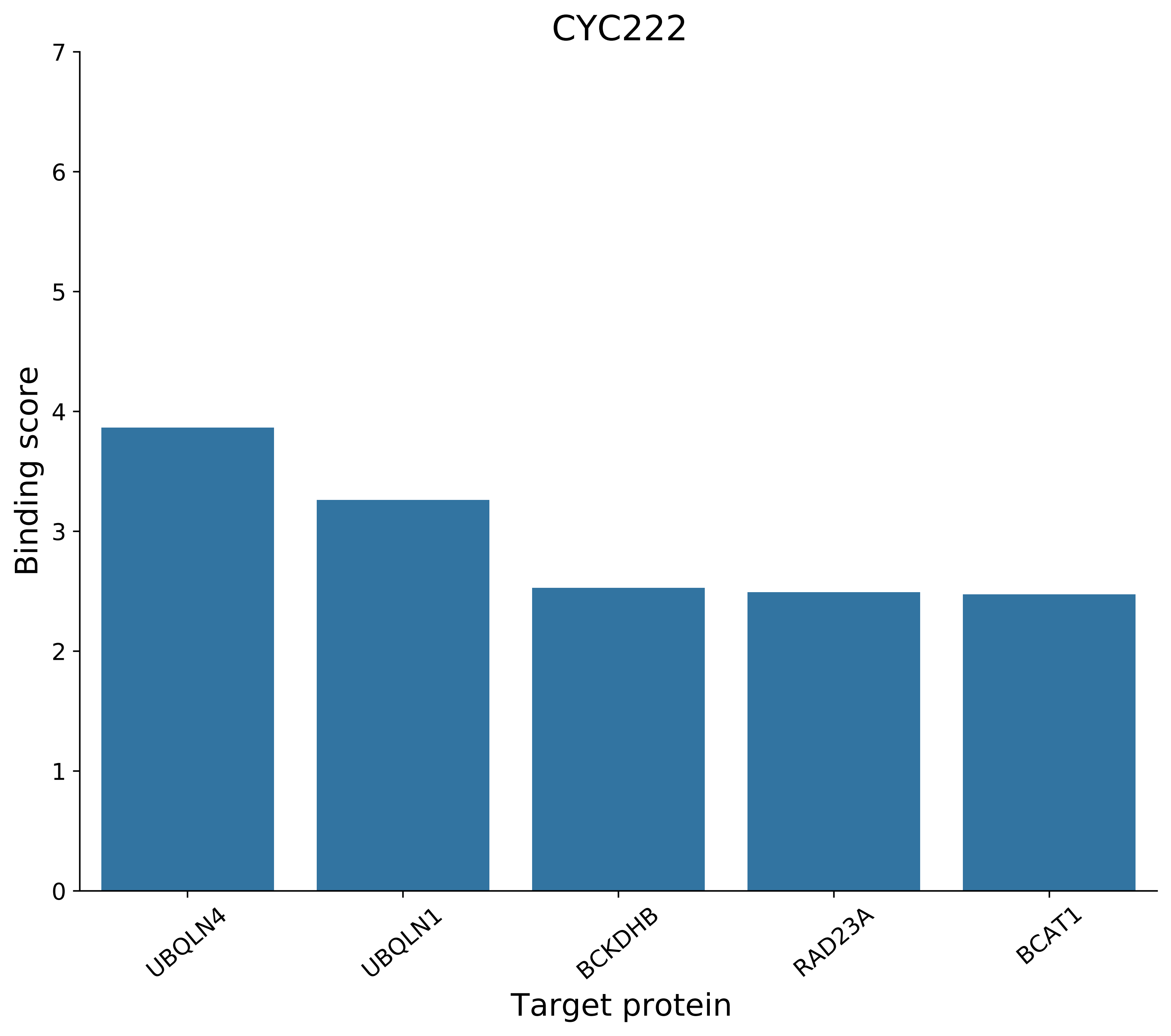
CYC222
Binding score
0
1
2
3
4
5
6
7
UBQLN4
UBQLN1
BCKDHB
RAD23A
BCAT1
Target protein

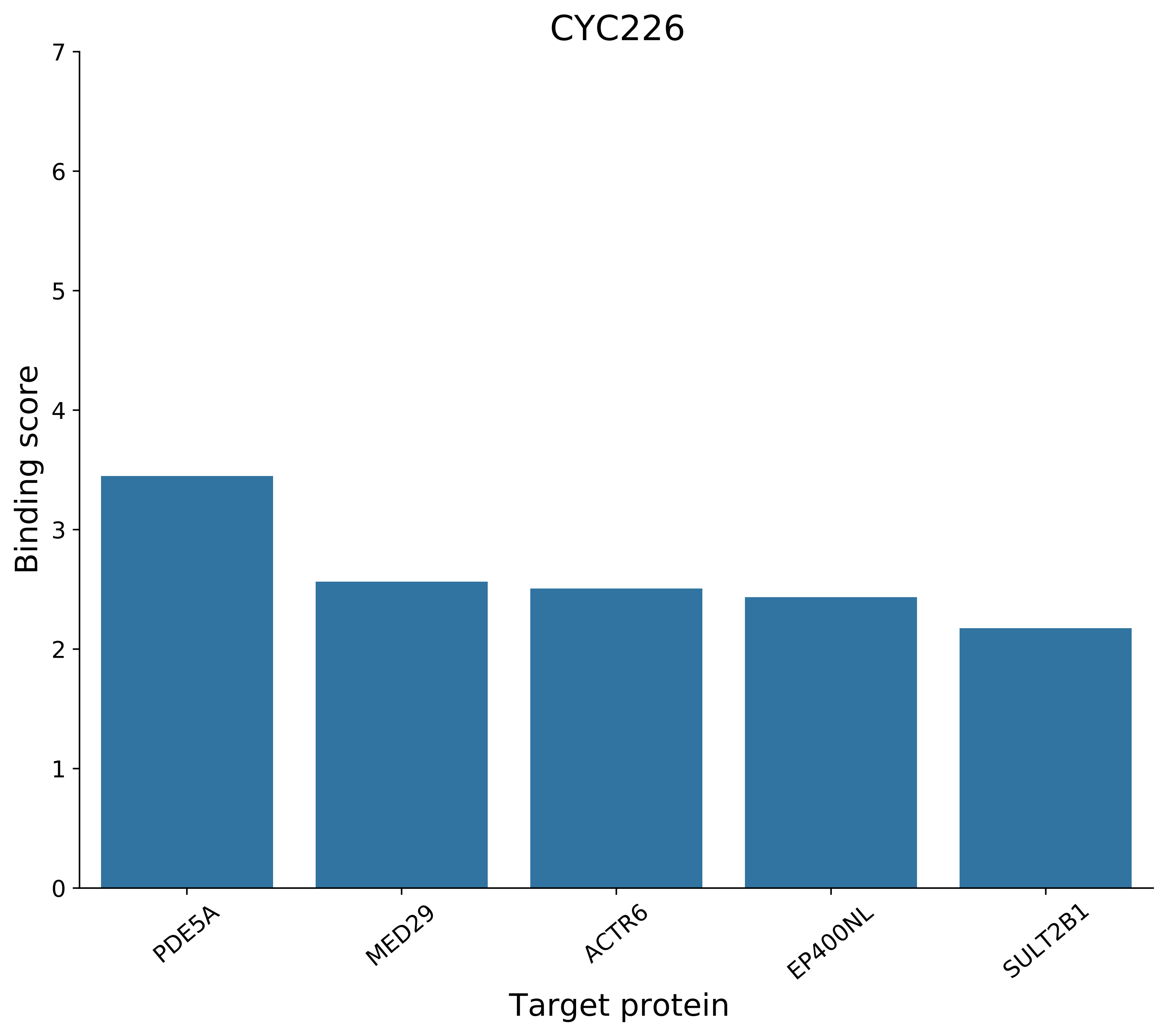
CYC226
Binding score
0
1
2
3
4
5
6
7
PDE5A
MED29
ACTR6
EP400NL
SULT2B1
Target protein

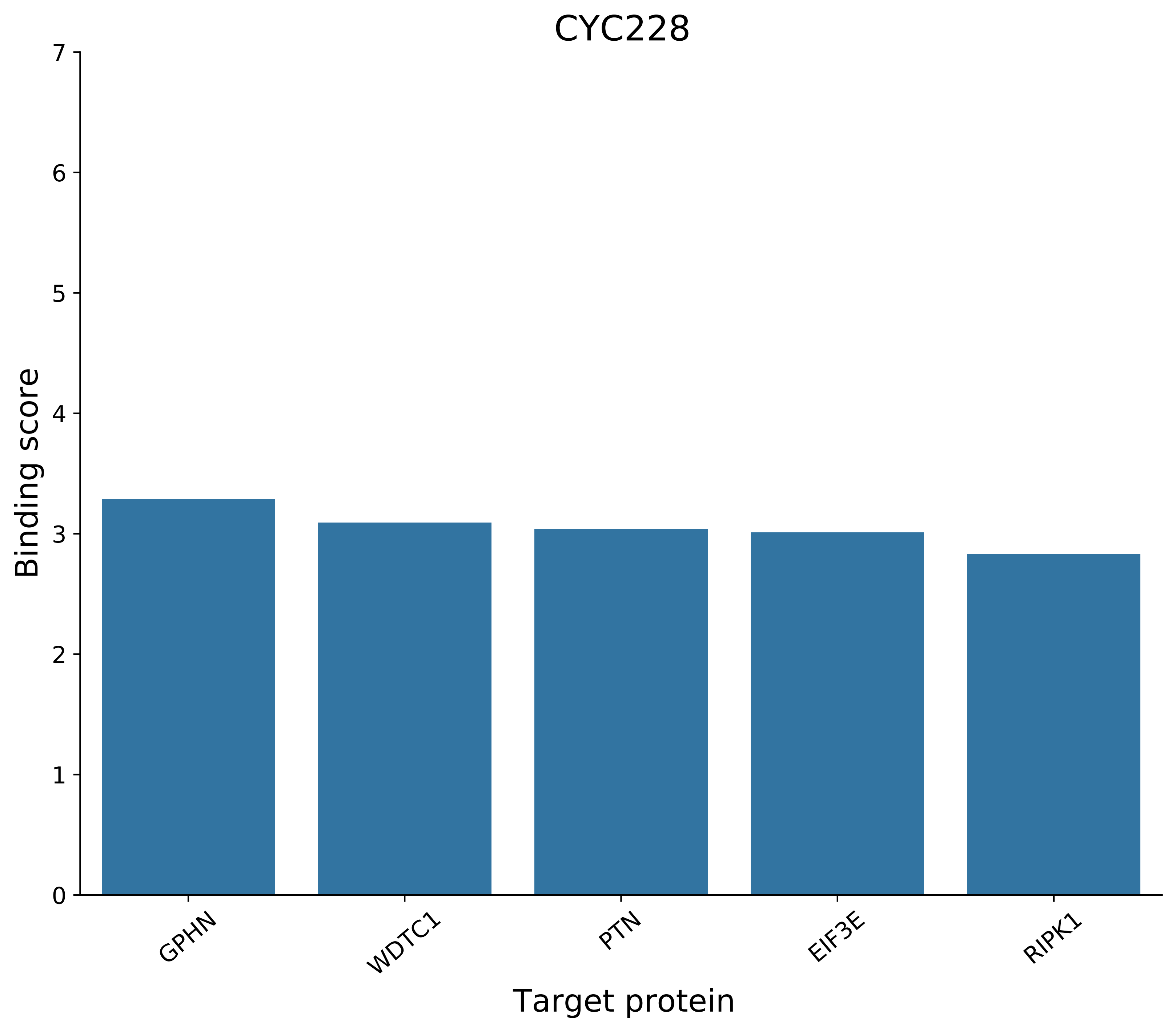

CYC228
Binding score
0
1
2
3
4
5
6
7
GPHN
WDTC1
PTN
EIF3E
RIPK1
Target protein

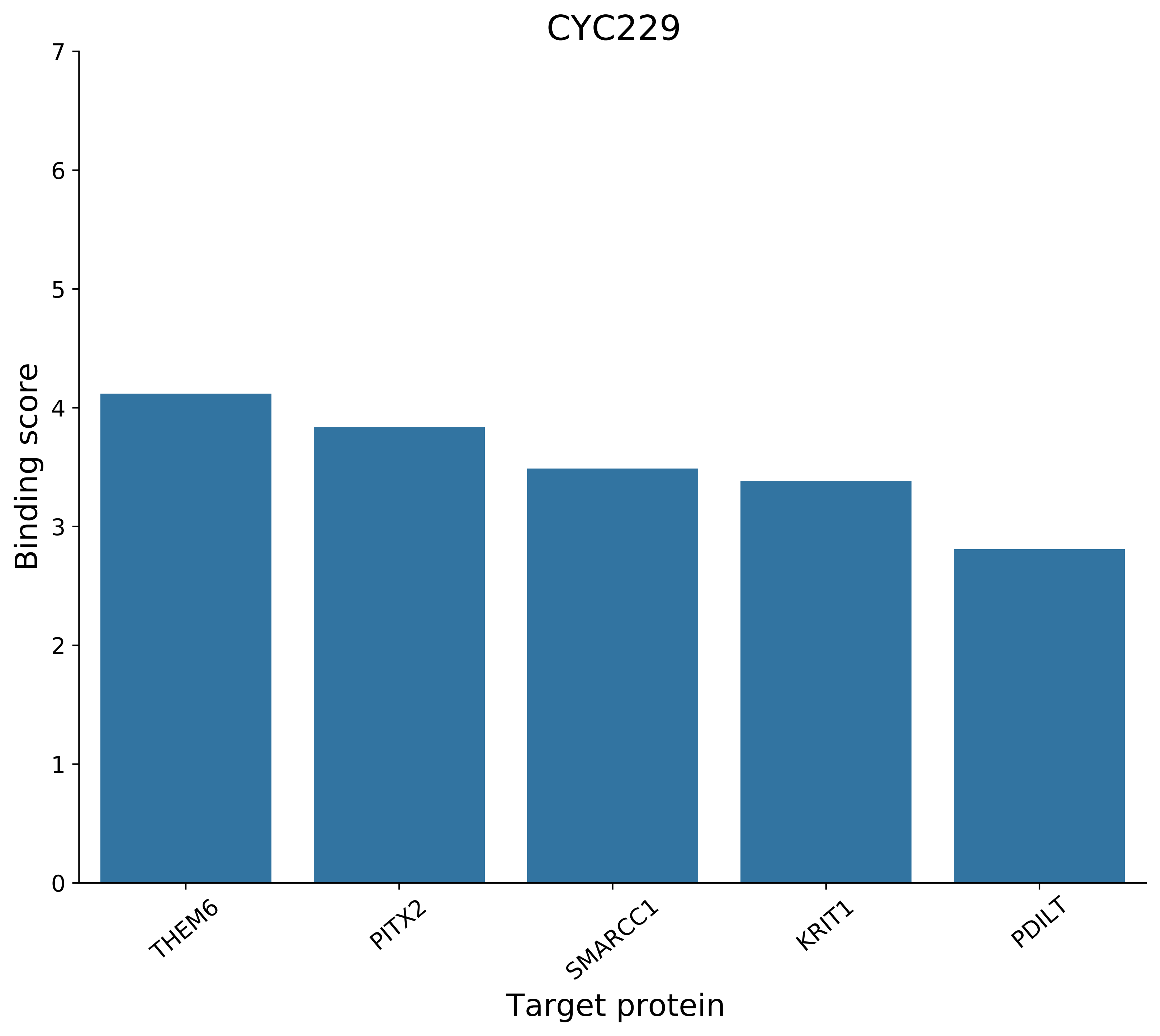
CYC229
Binding score
0
1
2
3
4
5
6
7
THEM6
PITX2
SMARCC1
KRIT1
PDILT
Target protein

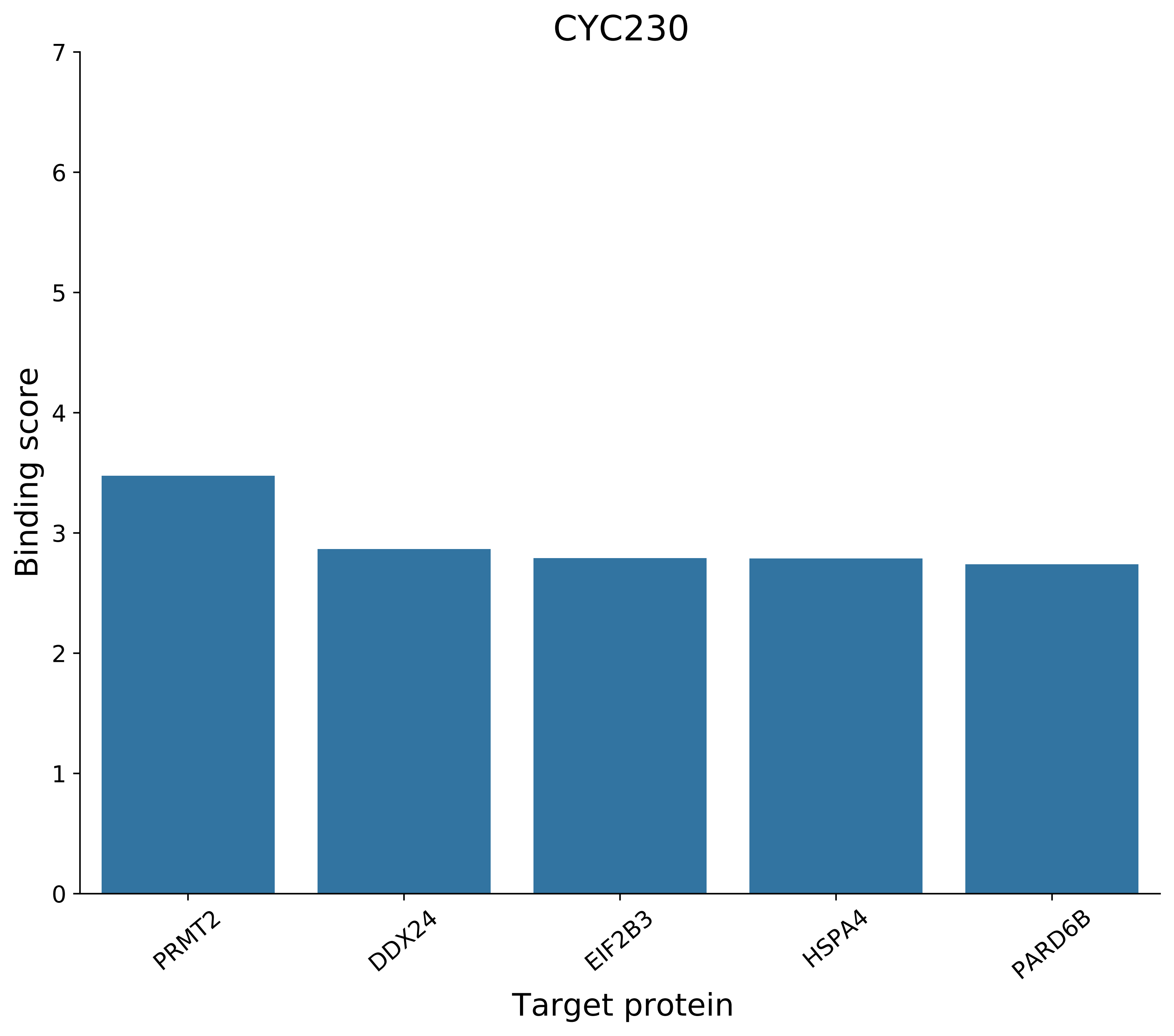

CYC230
Binding score
7
6
5
4
3
2
1
0
PRMT2
DDX24
EIF2B3
HSPA4
PARD6B
Target protein

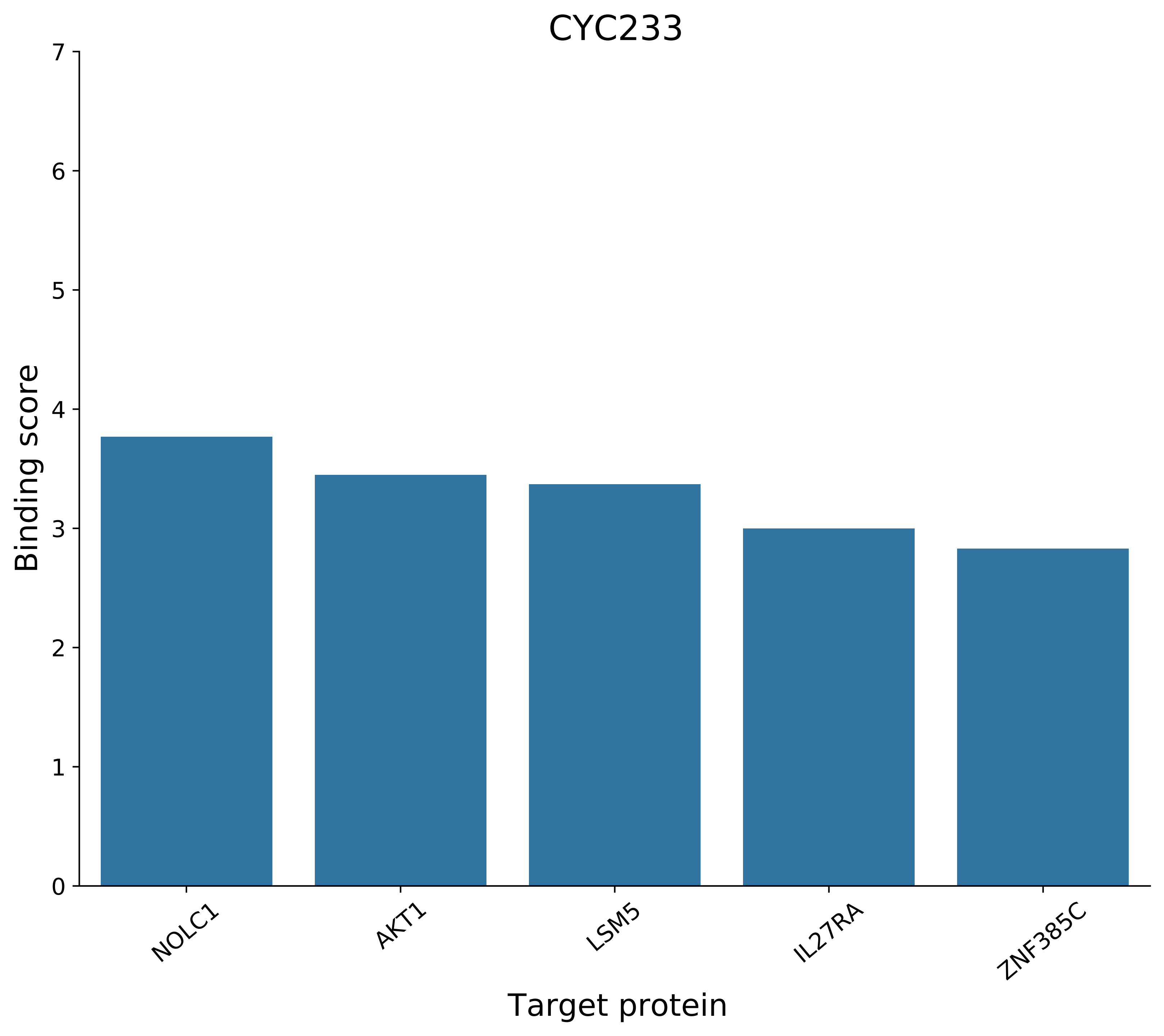
CYC233
Binding score
0
1
2
3
4
5
6
7
NOLC1
AKT1
LSM5
IL27RA
ZNF385C
Target protein

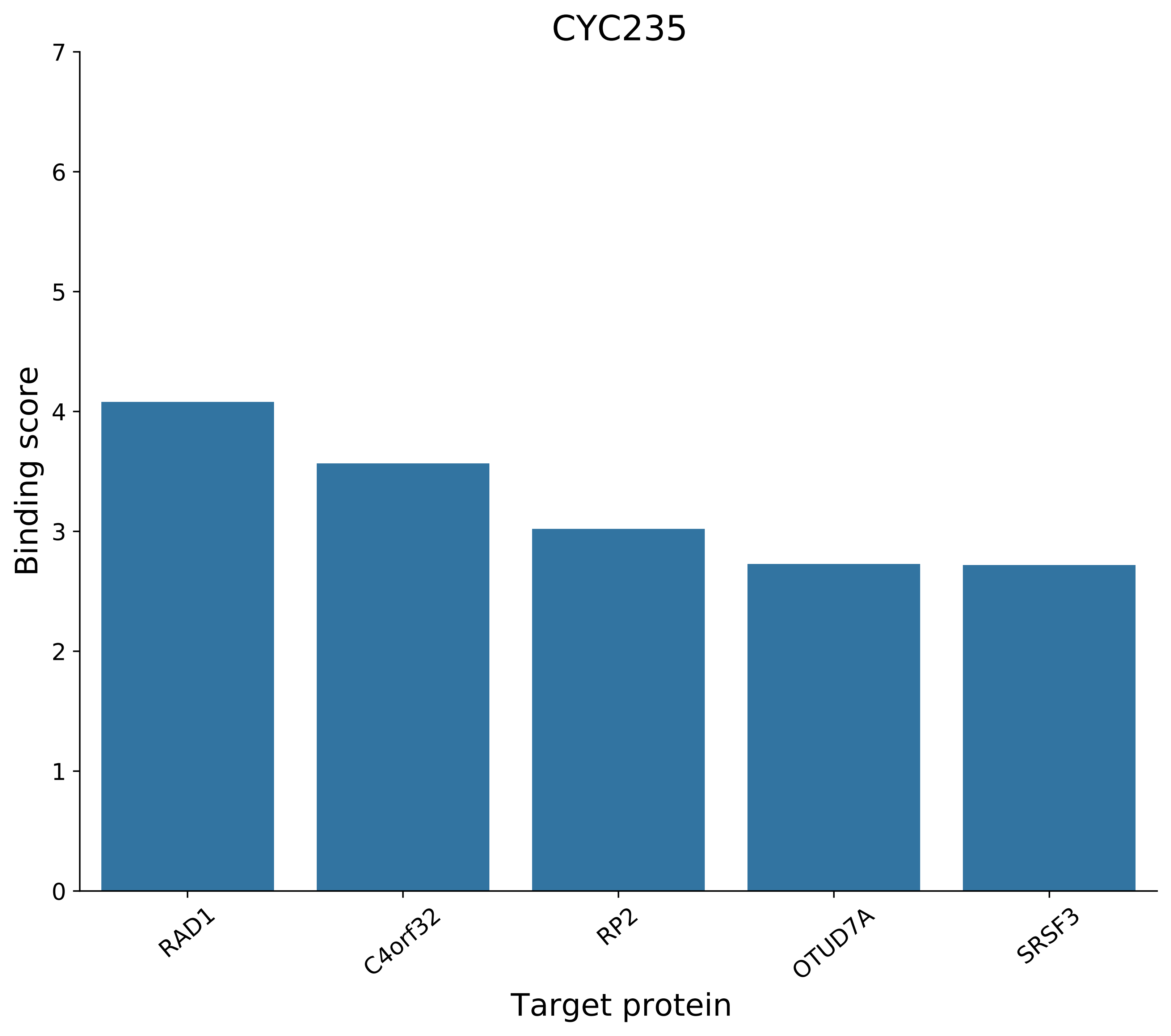
CYC235
Binding score
0
1
2
3
4
5
6
7
RAD1
C4orf32
RP2
OTUD7A
SRSF3
Target protein

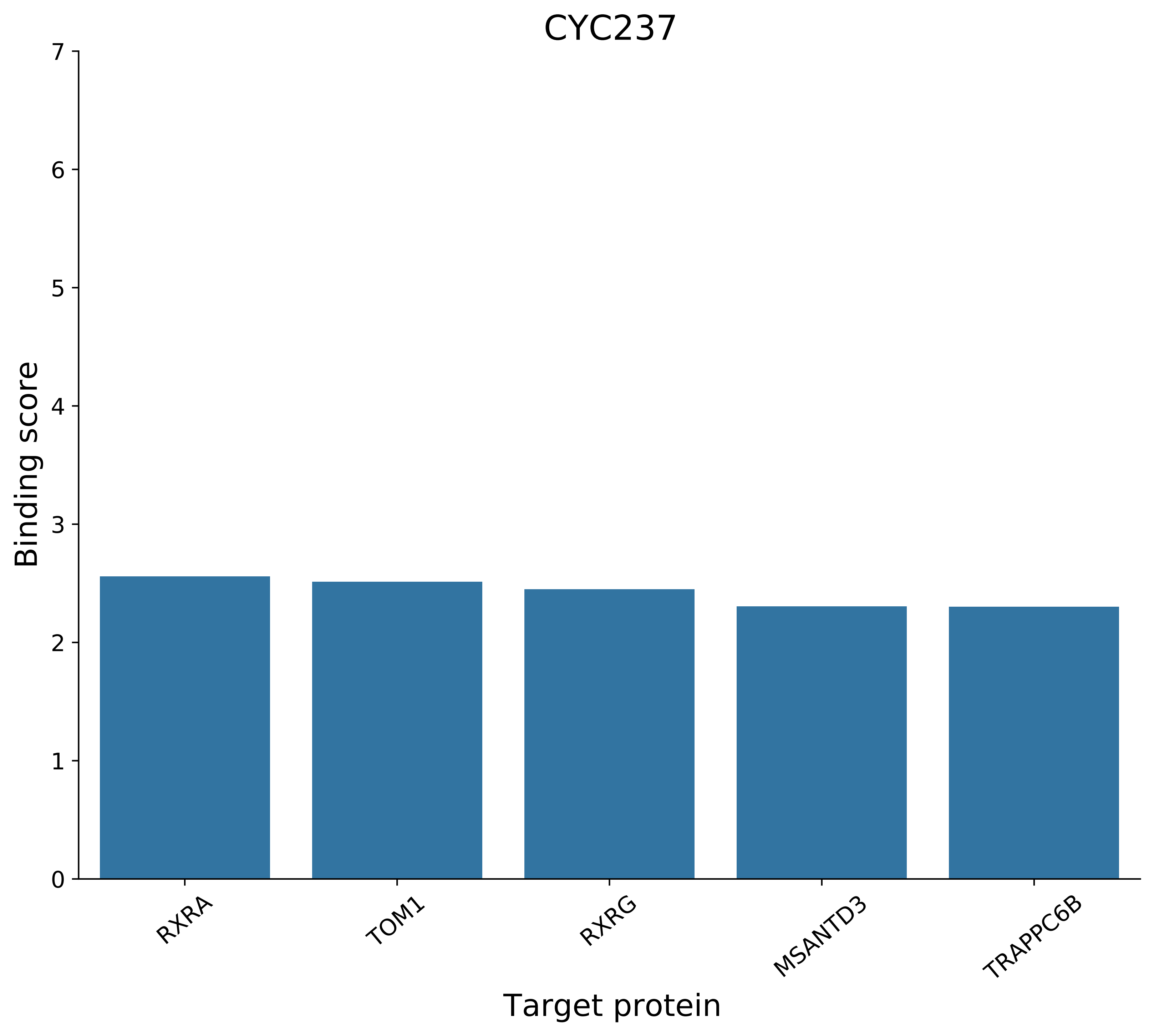
CYC237
Binding score
0
1
2
3
4
5
6
7
RXRA
TOM1
RXRG
MSANTD3
TRAPPC6B
Target protein

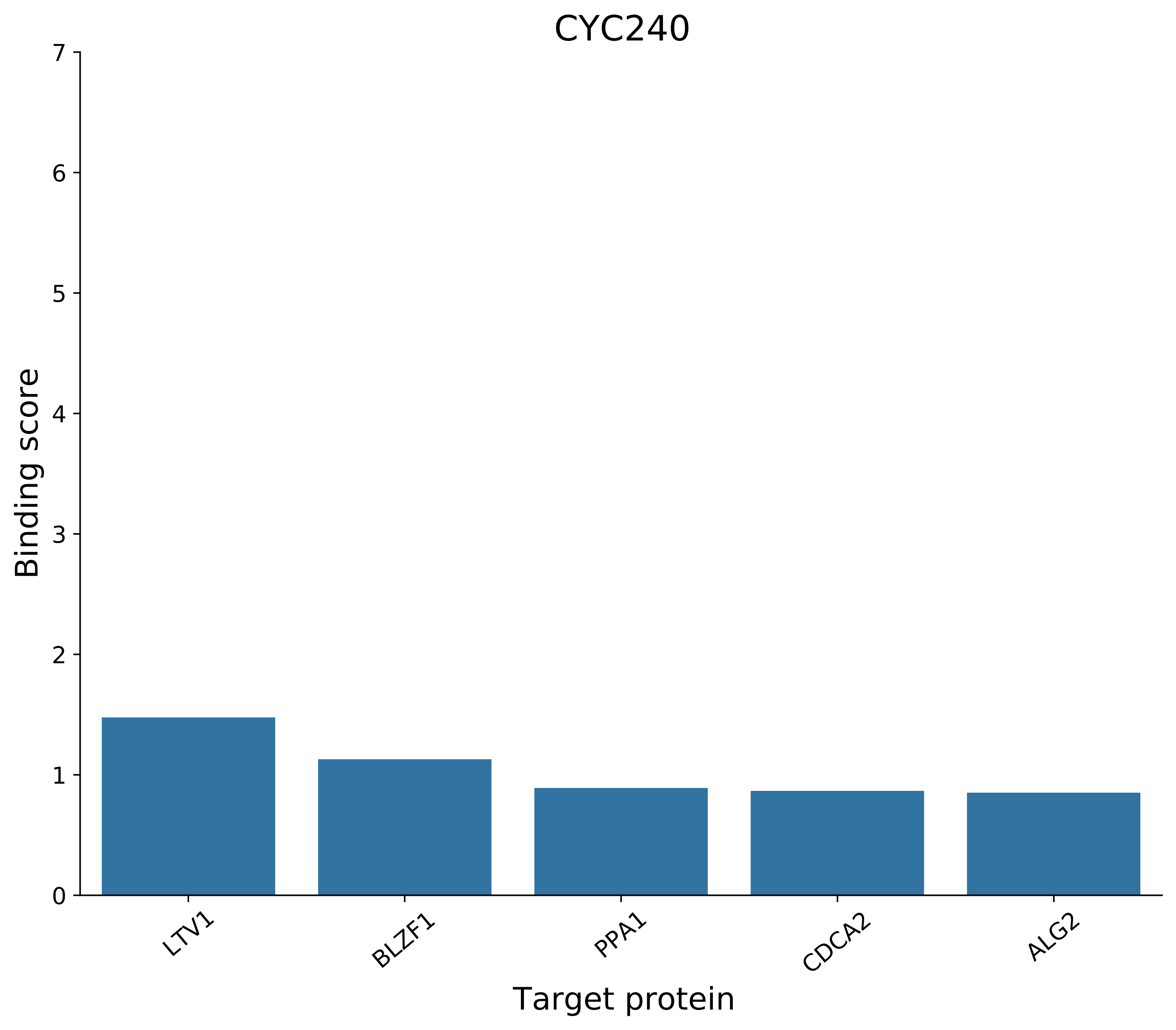

CYC240
Binding score
7
6
5
4
3
2
1
0
LTV1
BLZF1
PPA1
CDCA2
ALG2
Target protein

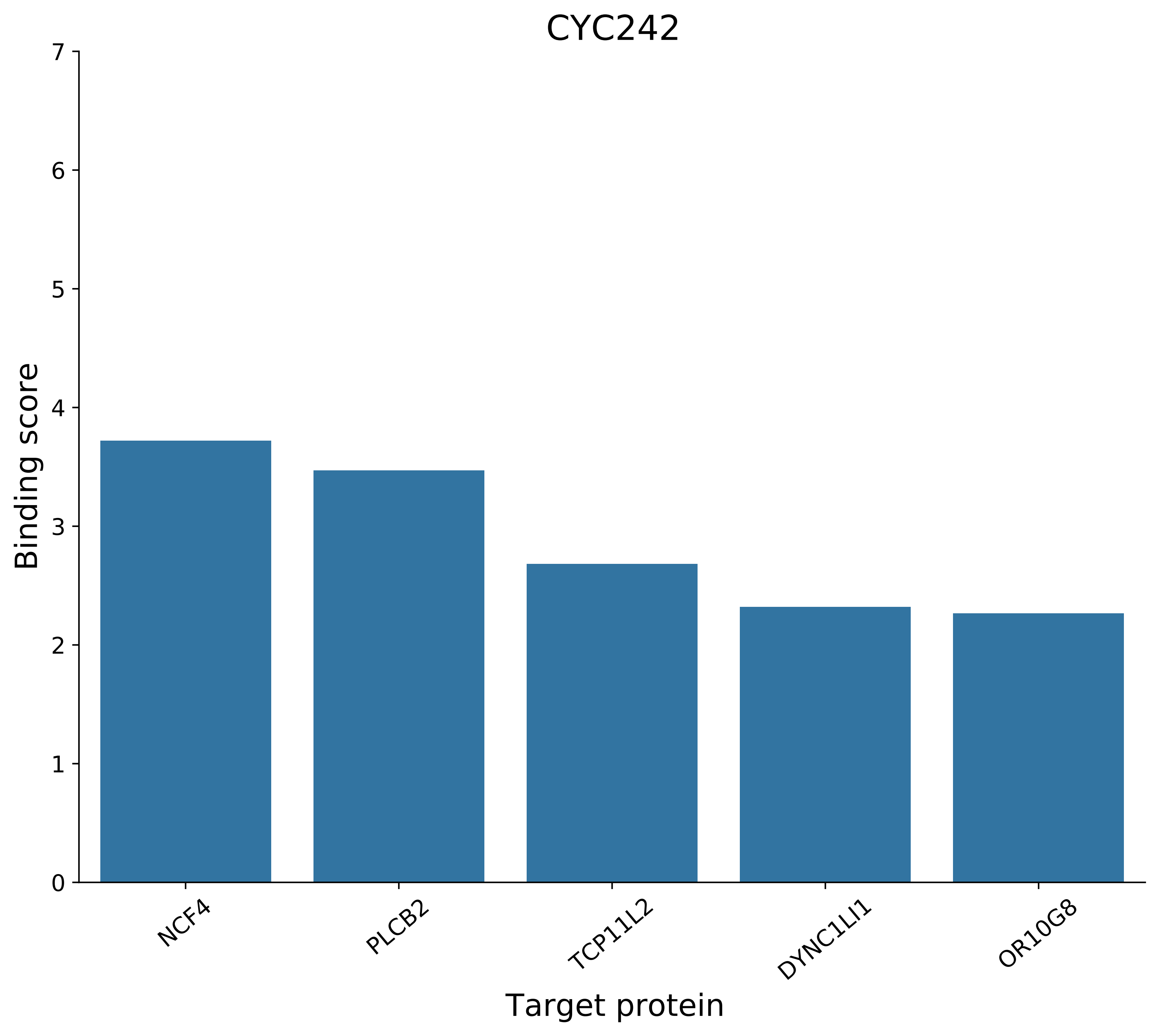

CYC242
Binding score
7
6
5
4
3
2
1
0
NCF4
PLCB2
TCP11L2
DYNC1LI1
OR10G8
Target protein

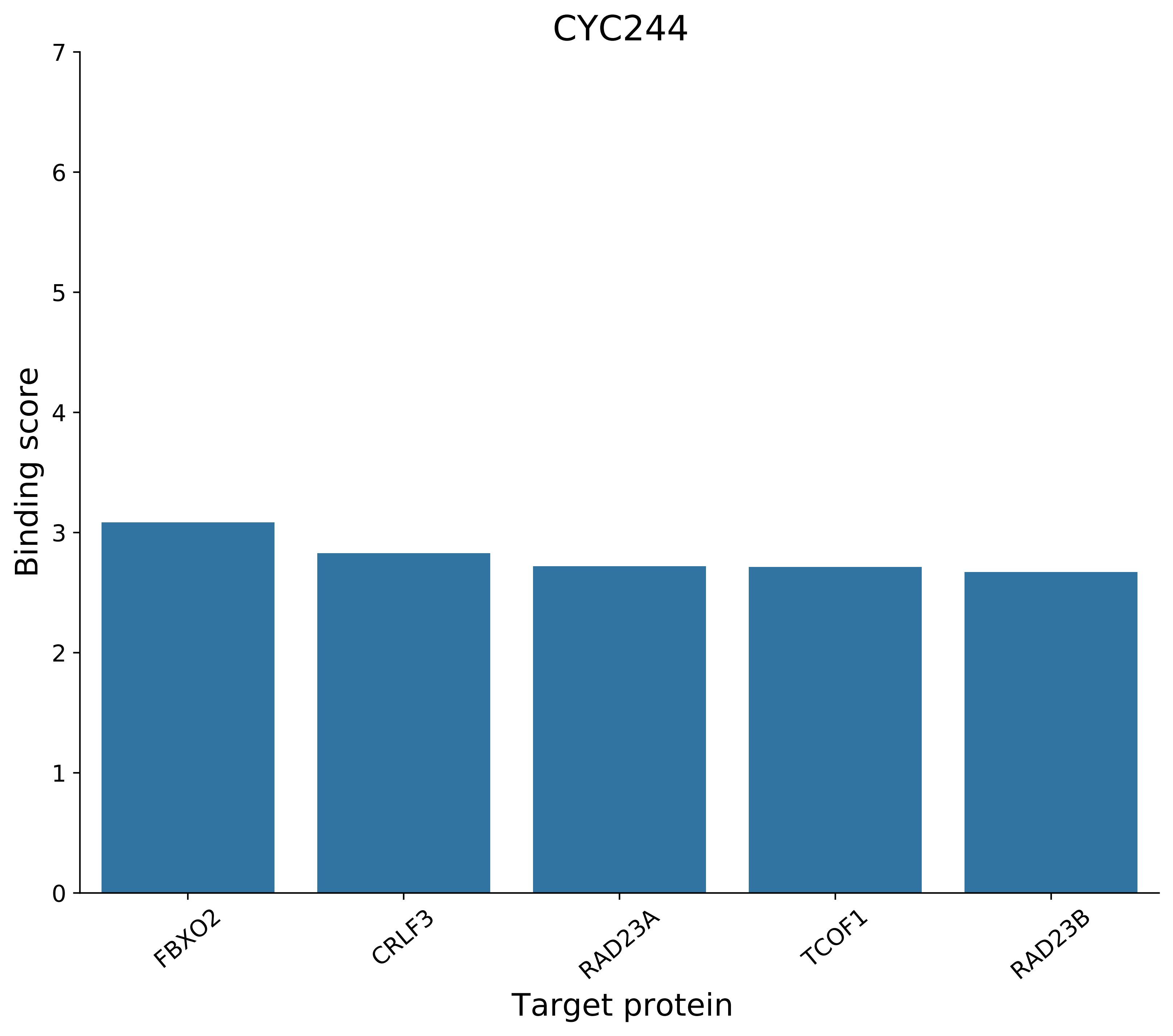
CYC244
Binding score
0
1
2
3
4
5
6
7
FBXO2
CRLF3
RAD23A
TCOF1
RAD23B
Target protein

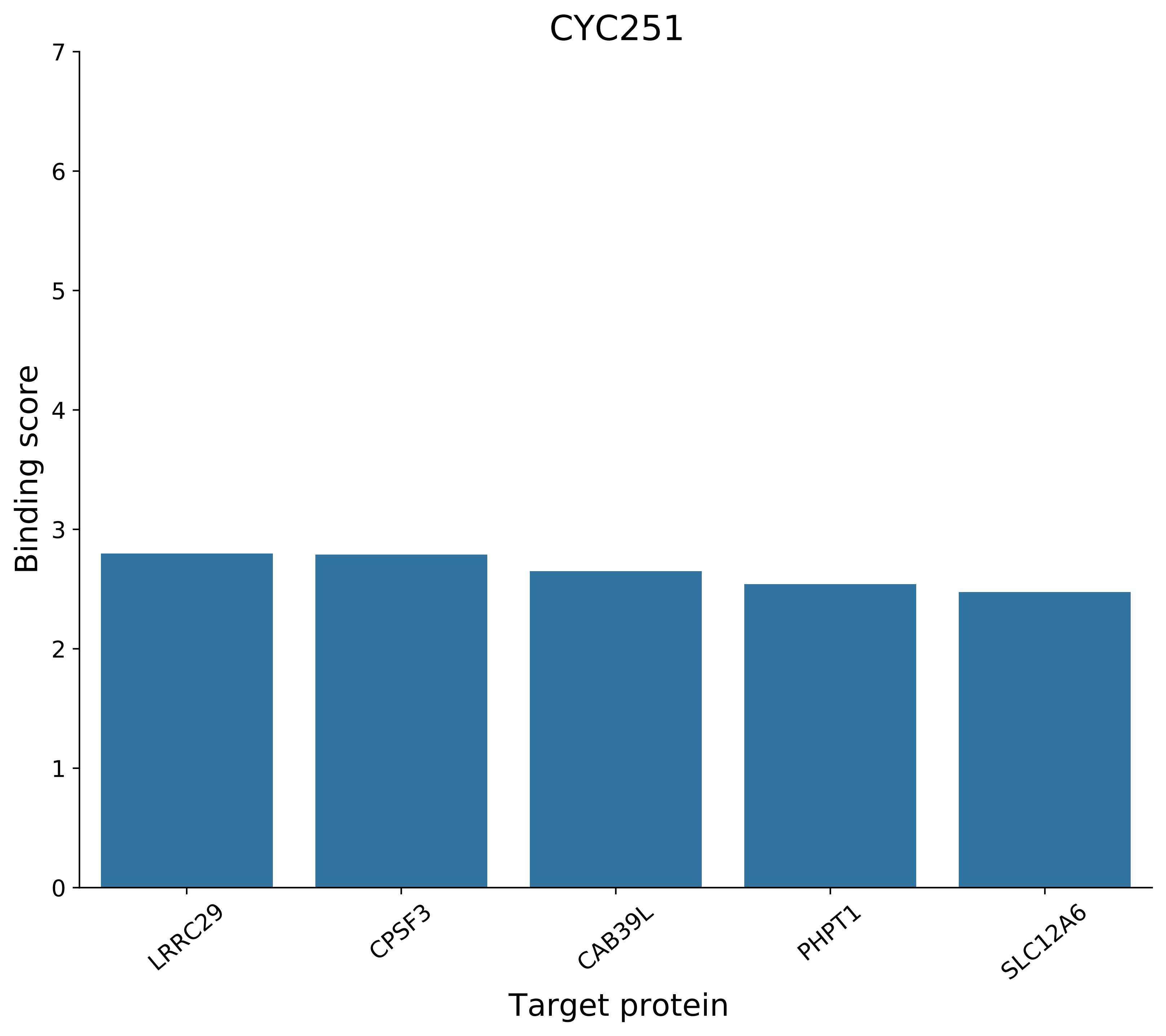
CYC251
Binding score
7
6
5
4
3
2
1
0
LRRC29
CPSF3
CAB39L
PHPT1
SLC12A6
Target protein

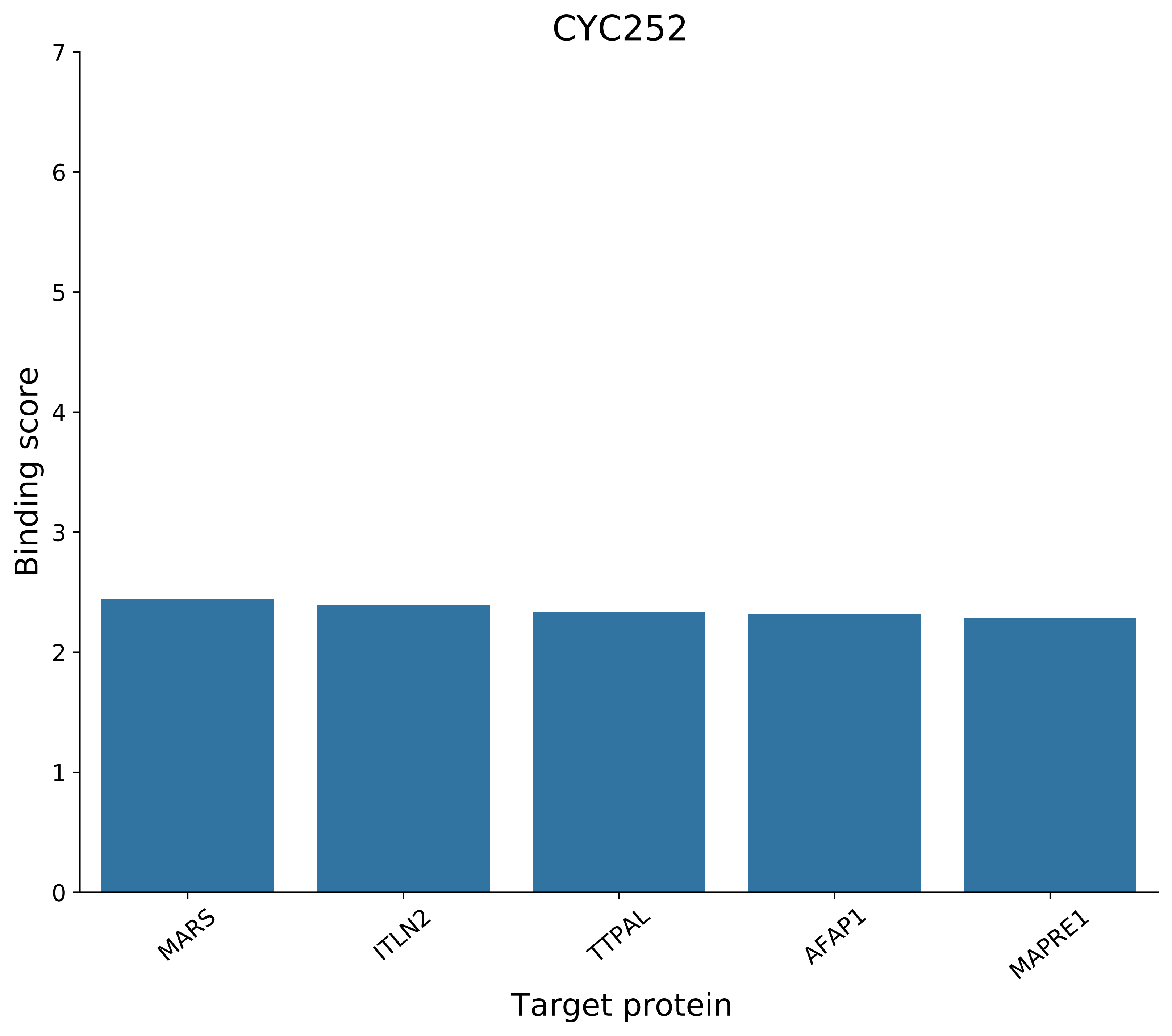
CYC252
Binding score
Target protein
0
1
2
3
4
5
6
7
MARS
ITLN2
TTPAL
AFAP1
MAPRE1

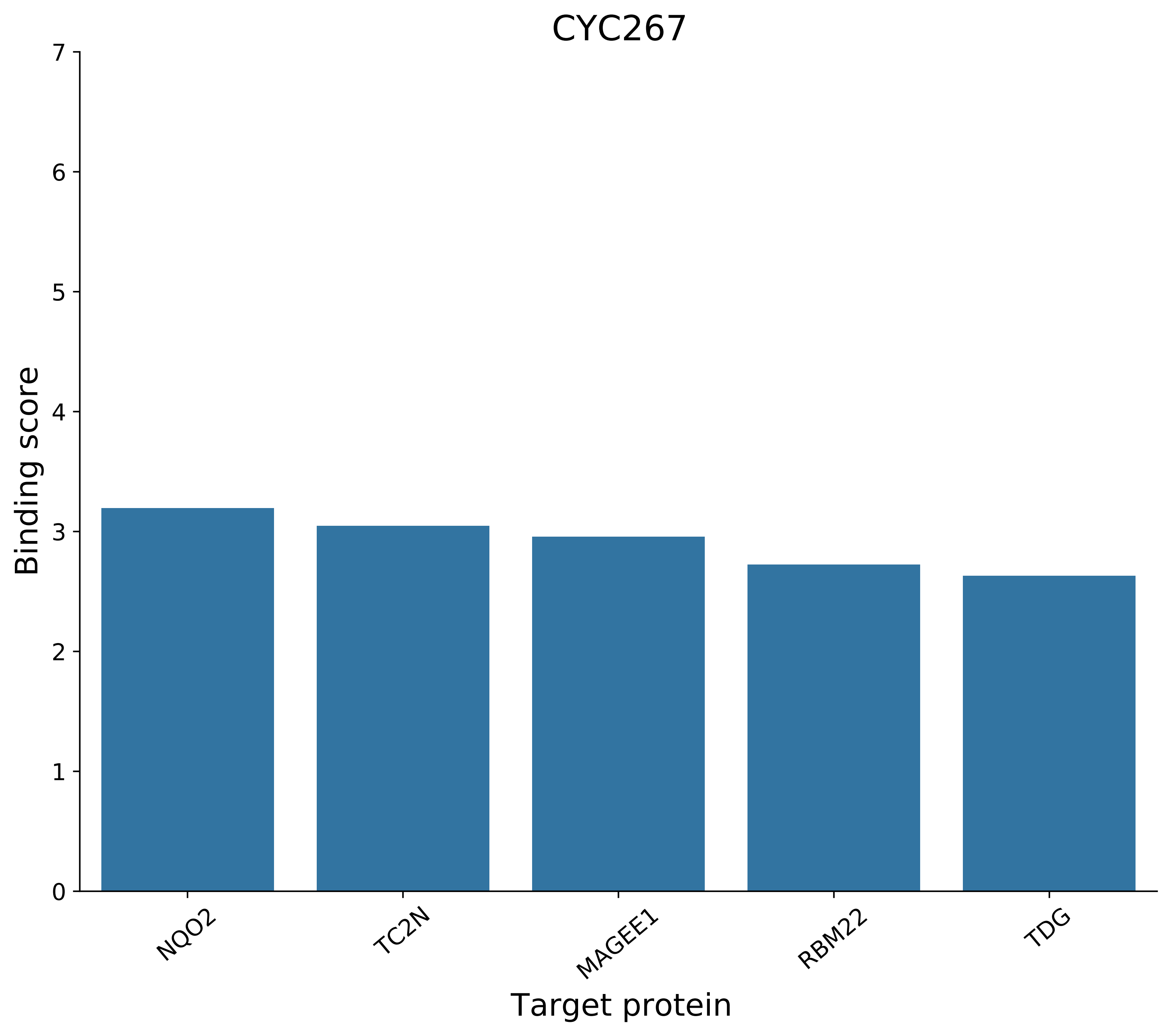
CYC267
Binding score
0
1
2
3
4
5
6
7
NQO2
TC2N
MAGEE1
RBM22
TDG
Target protein

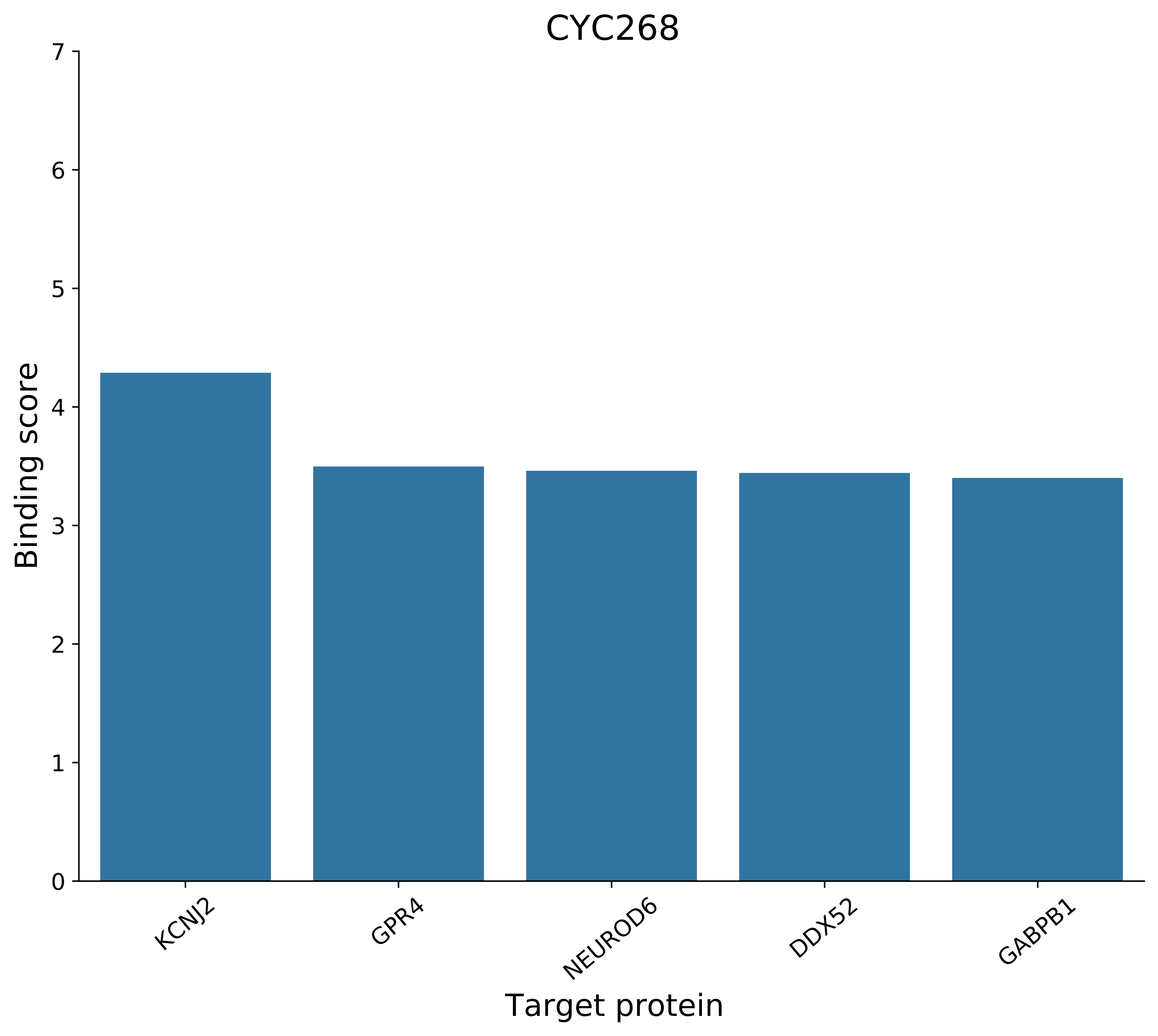
CYC268
Binding score
0
1
2
3
4
5
6
7
KCNJ2
GPR4
NEUROD6
DDX52
GABPB1
Target protein

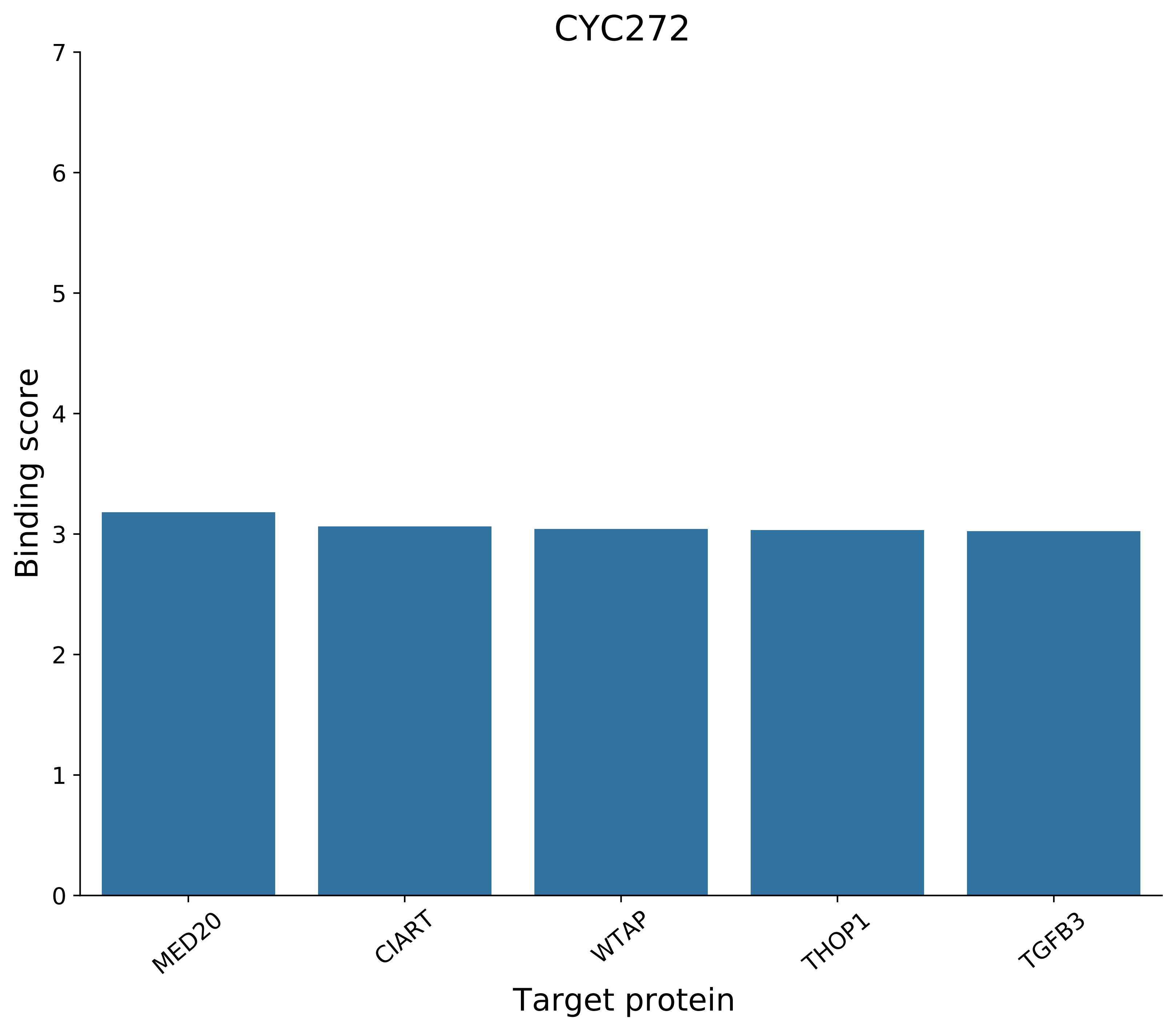
CYC272
Binding score
Target protein
0
1
2
3
4
5
6
7
MED20
CIART
WTAP
THOP1
TGFB3

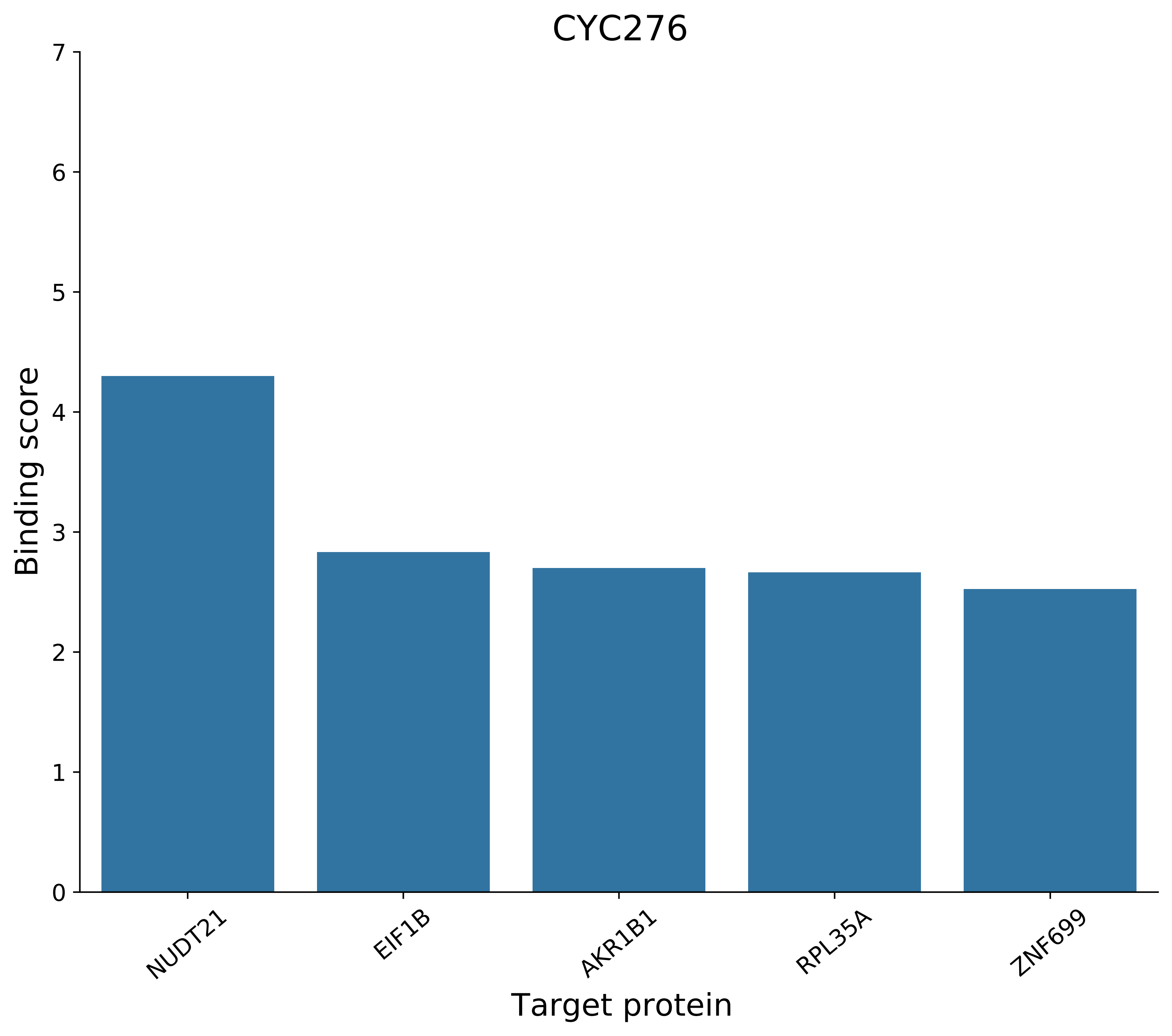

CYC276
Binding score
Target protein
0
1
2
3
4
5
6
7
NUDT21
EIF1B
AKR1B1
RPL35A
ZNF699

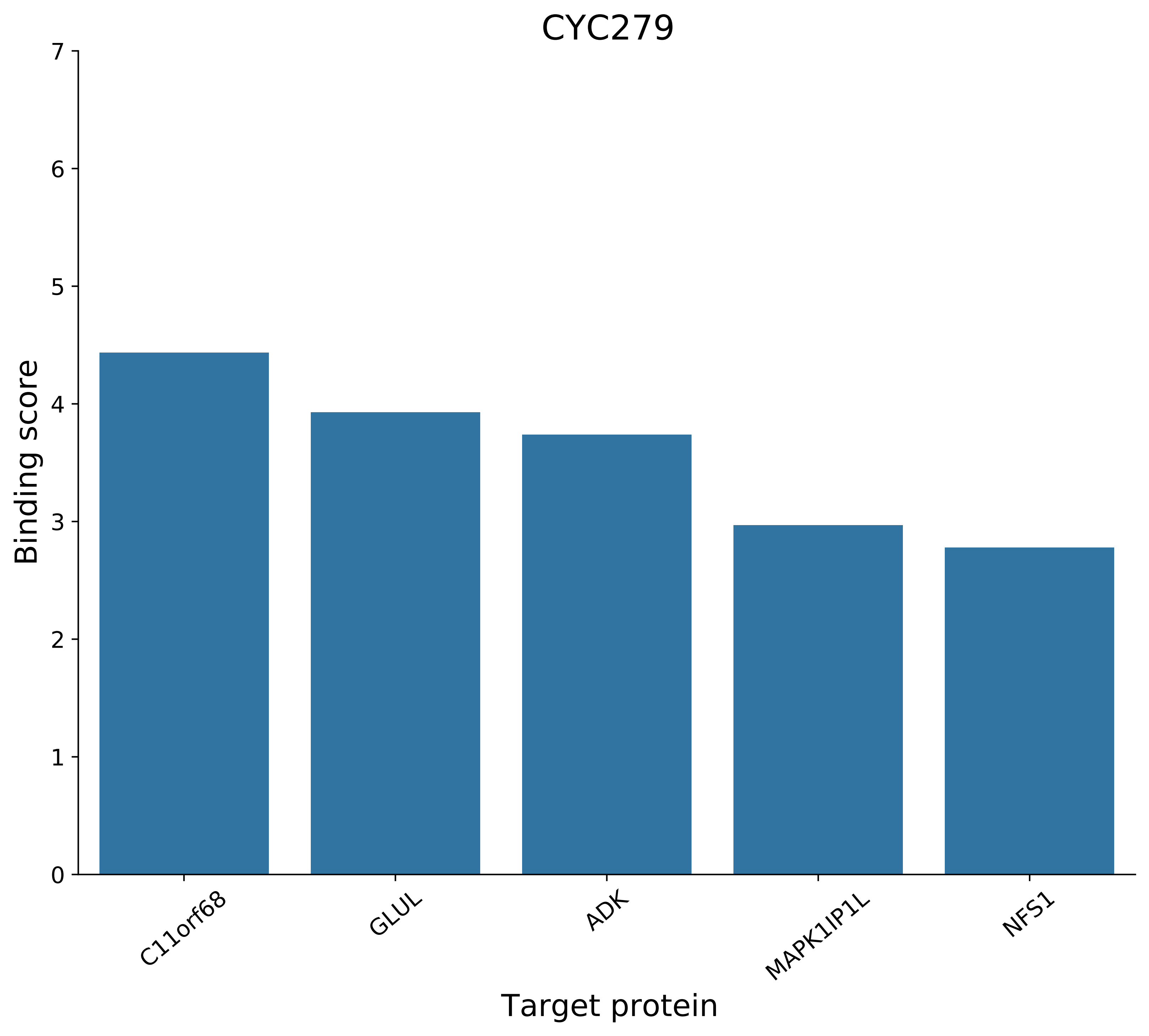

CYC279
Binding score
Target protein
C11orf68
GLUL
ADK
MAPK1IP1L
NFS1
0
1
2
3
4
5
6
7

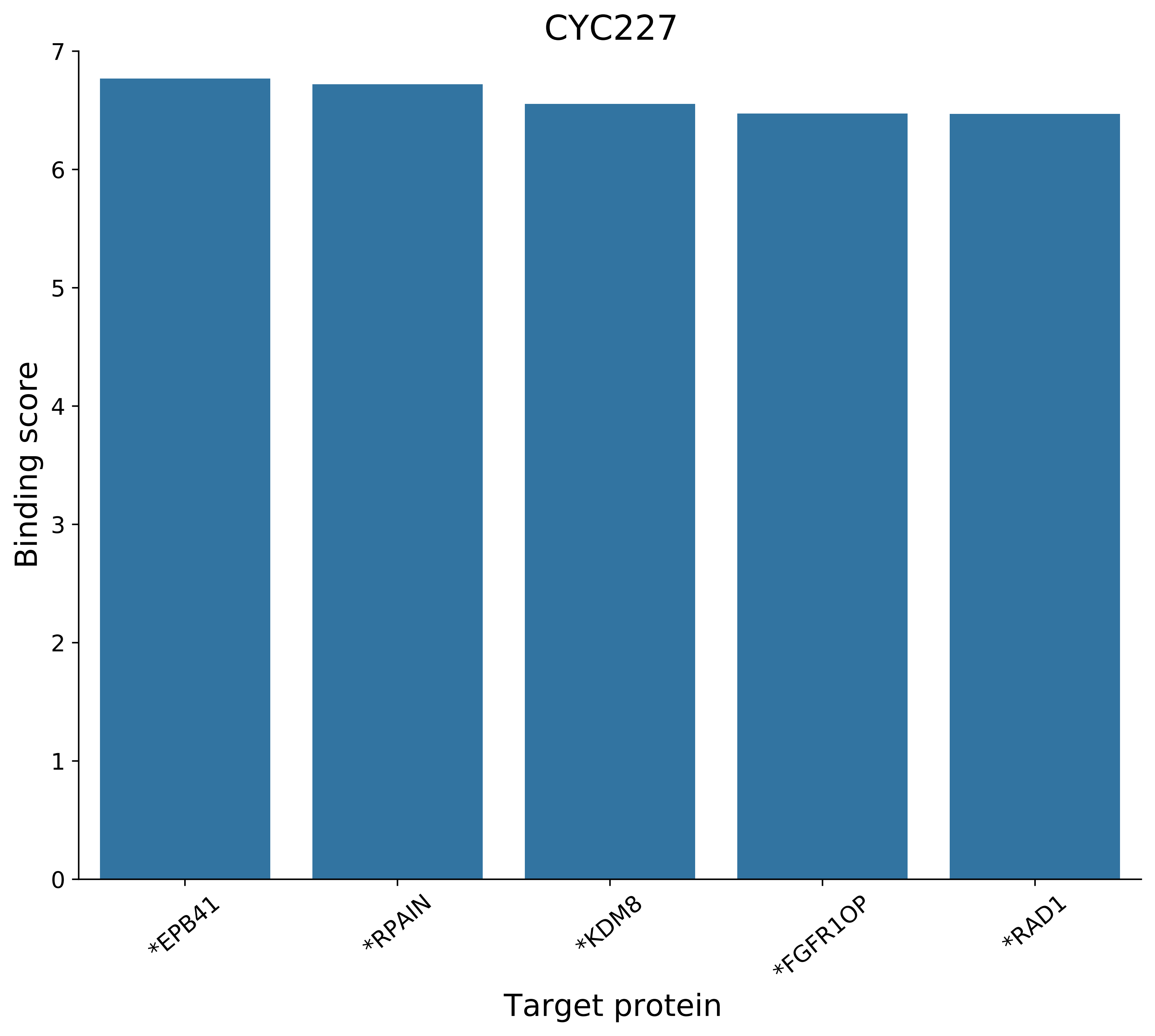
CYC227
Binding score
0
1
2
3
4
5
6
7
*EPB41
*RPAIN
*KDM8
*FGFR1OP
*RAD1
Target protein

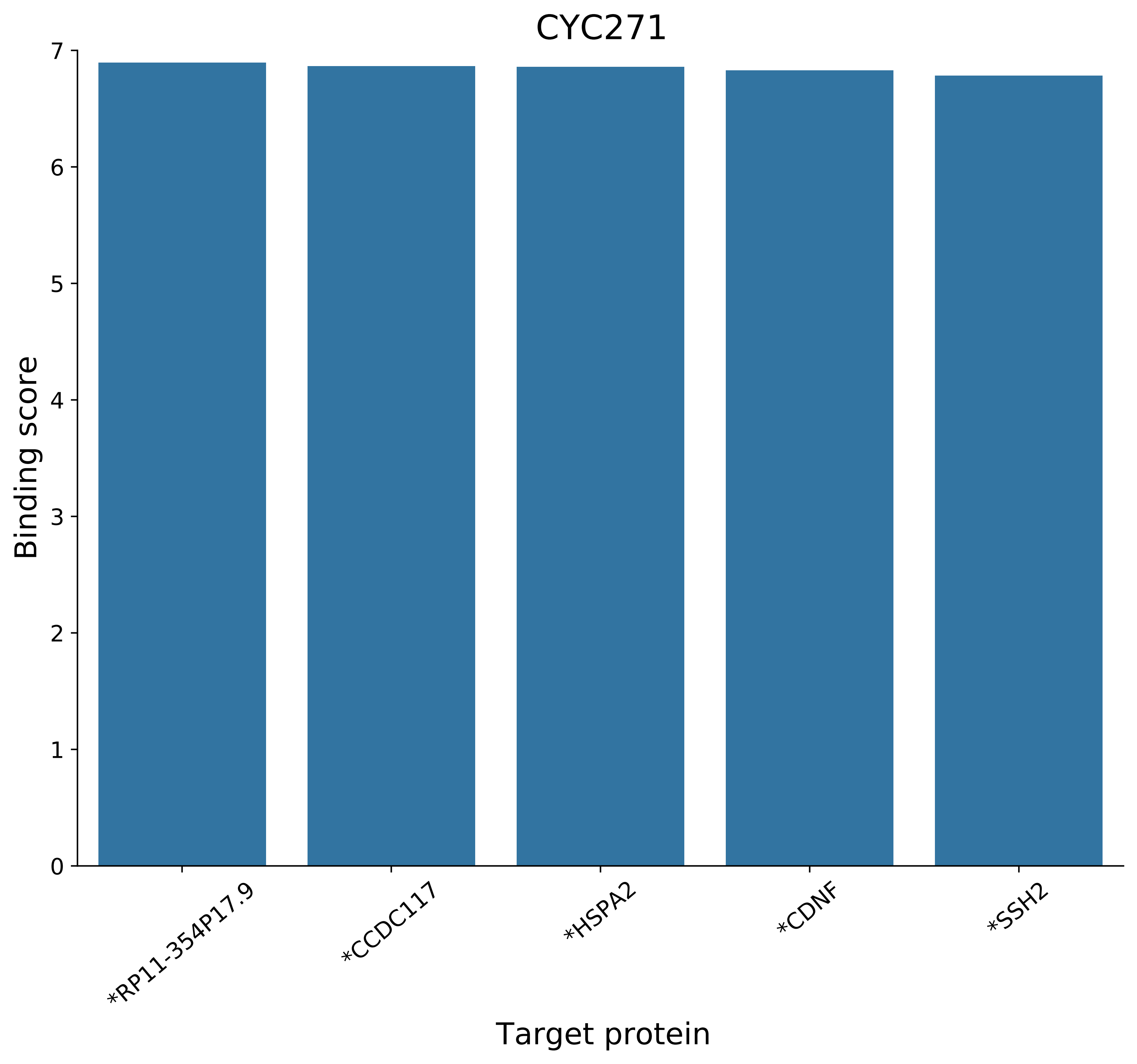
CYC271
Binding score
7
6
5
4
3
2
1
0
*RP11-354P17.9
*CCDC117
*HSPA2
*CDNF
*SSH2
Target protein
