## Supplementary Figures for "The landscape of high-affinity human antibodies against intratumoral antigens": Supplementary Figure 5.pdf

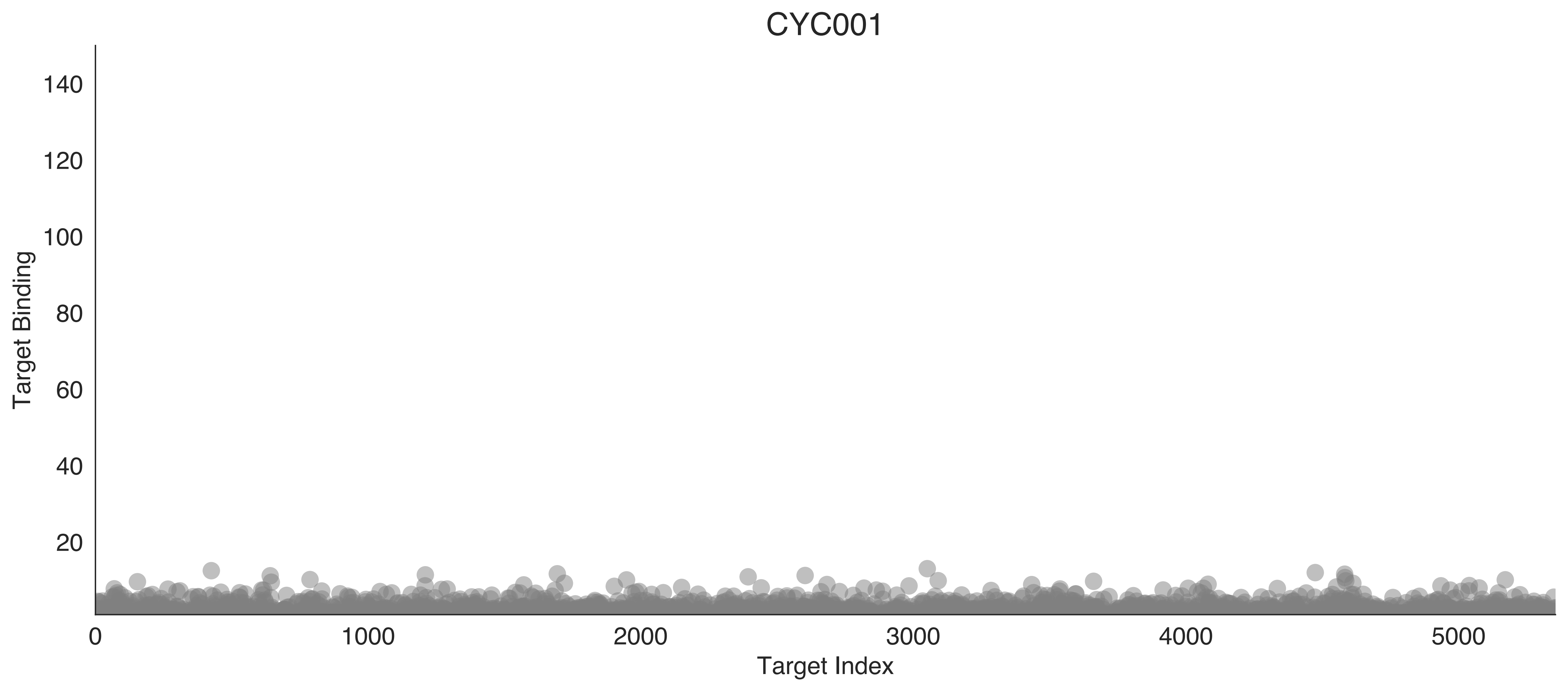
CYC001
Target Binding
140
120
100
80
60
40
20
0
1000
2000
3000
4000
5000
Target Index

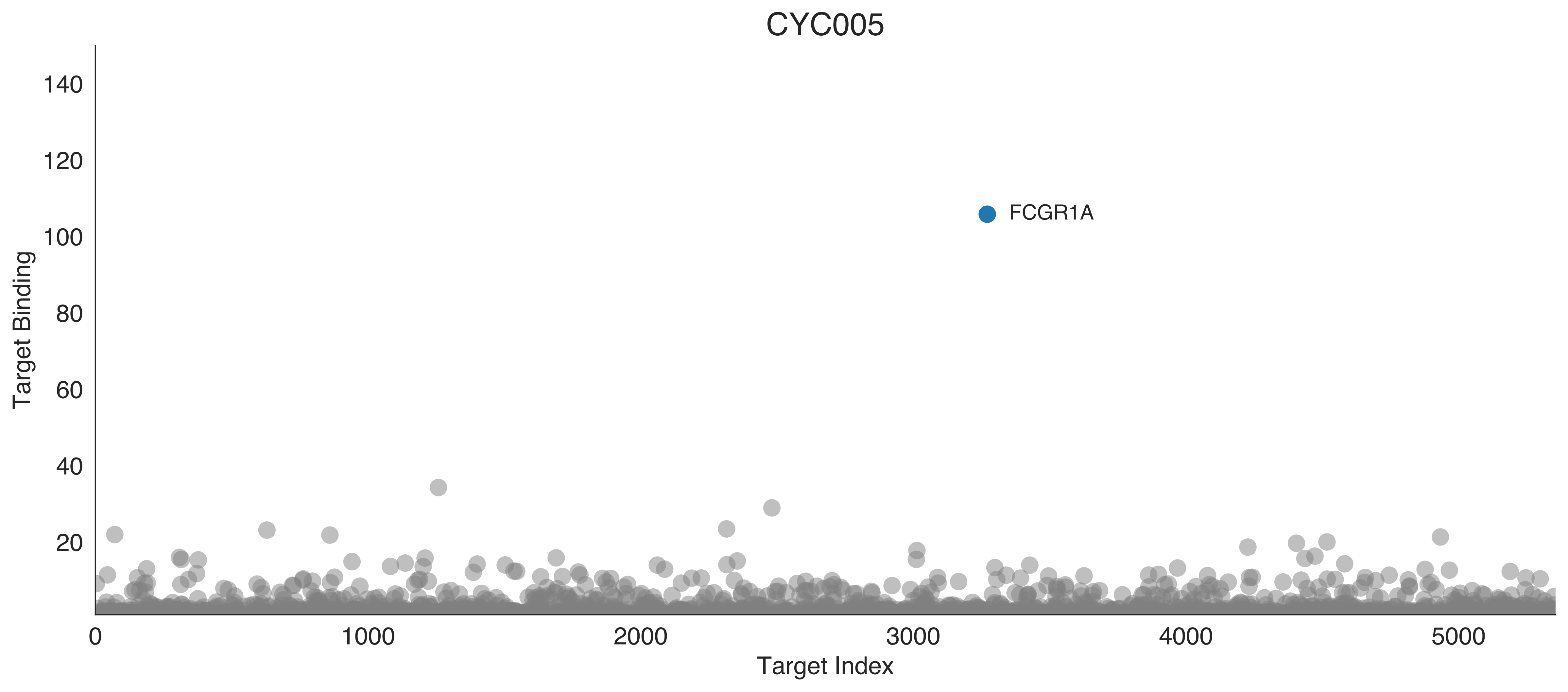
CYC005
FCGR1A
Target Binding
Target Index
0
20
40
60
80
100
120
140
1000
2000
3000
4000
5000

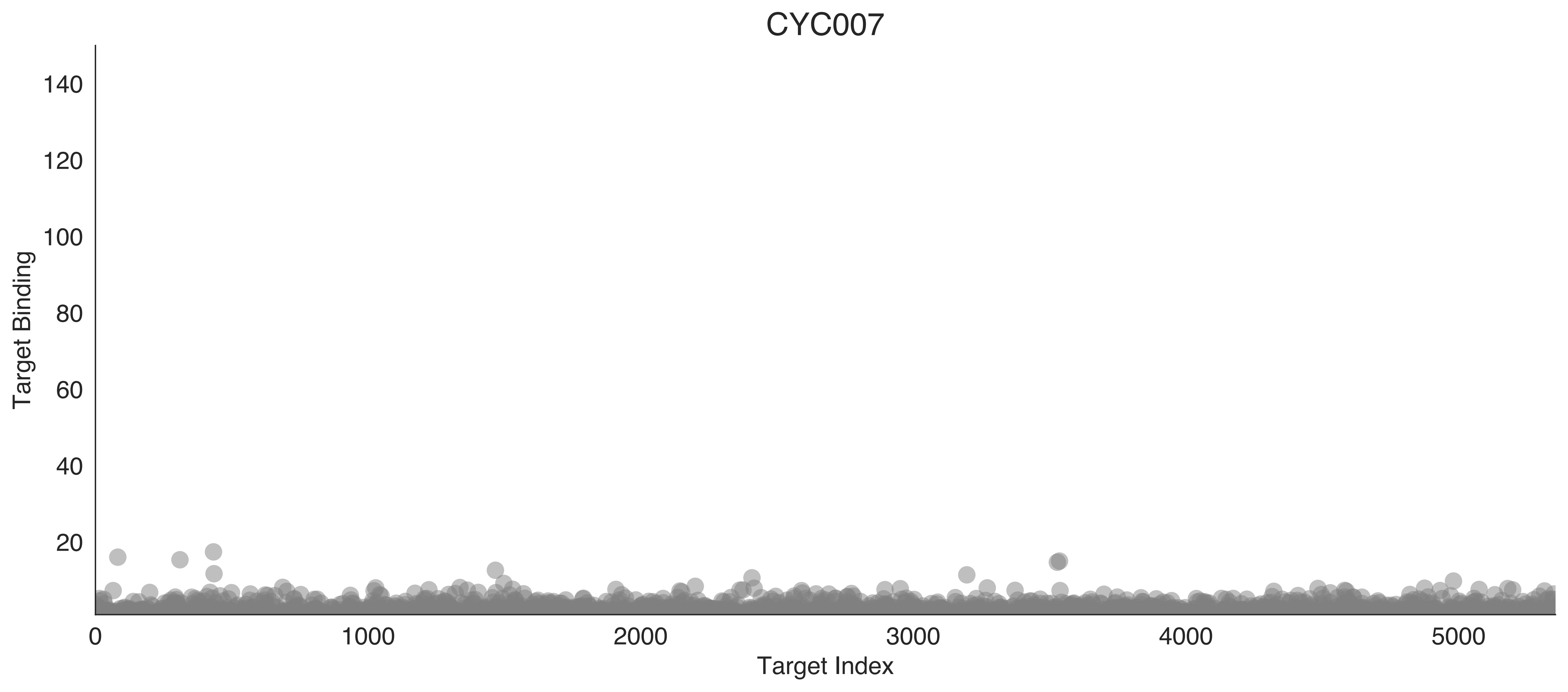
CYC007
Target Binding
140
120
100
80
60
40
20
0
1000
2000
3000
4000
5000
Target Index

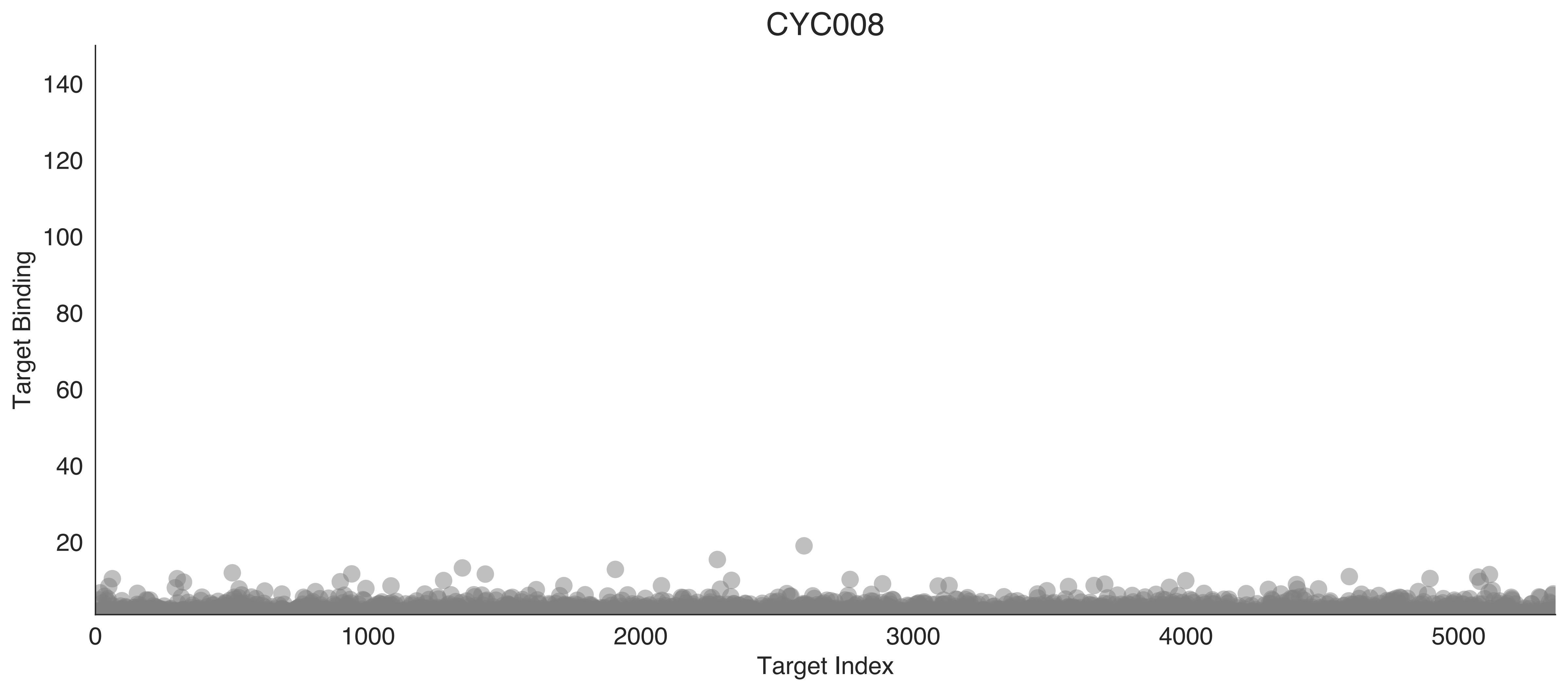
CYC008
Target Binding
140
120
100
80
60
40
20
0
1000
2000
3000
4000
5000
Target Index

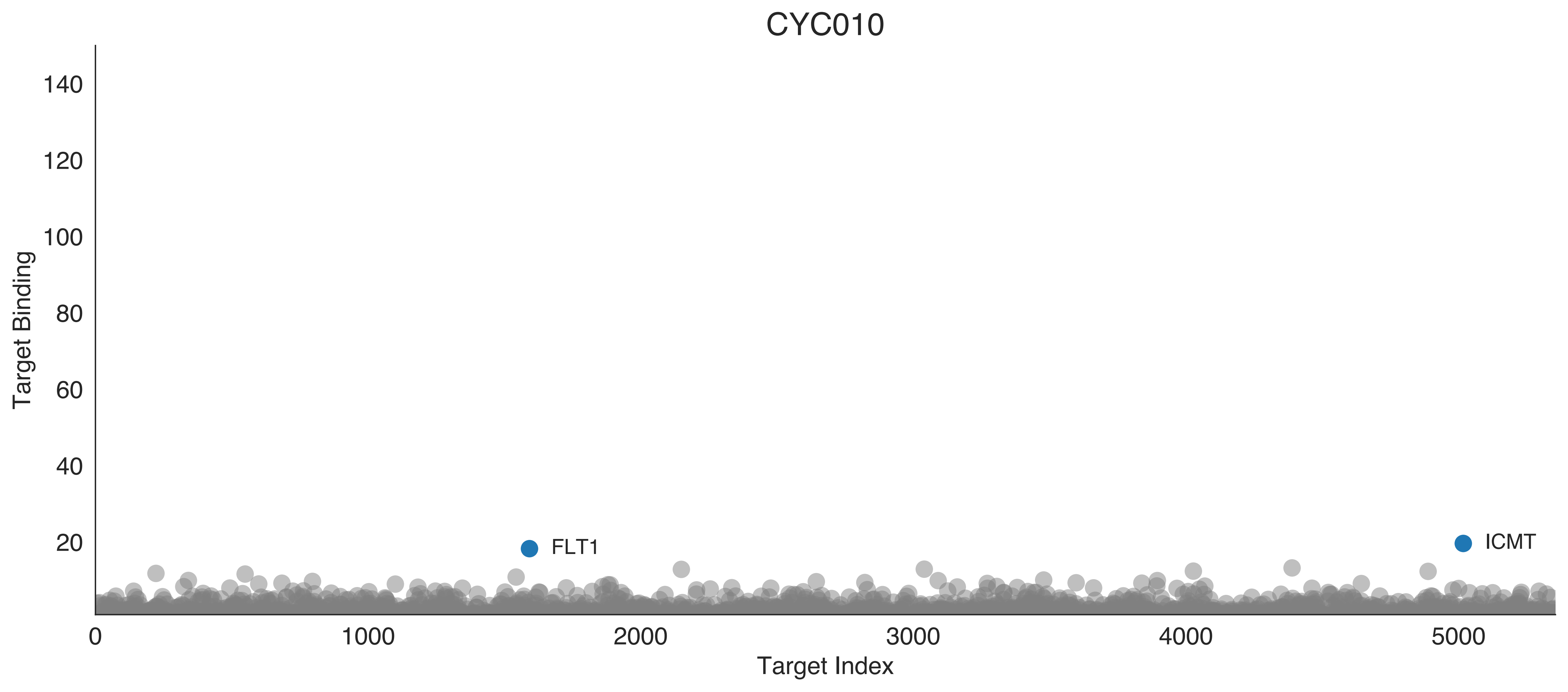

CYC010
Target Binding
Target Index
FLT1
ICMT
140
120
100
80
60
40
20
0
1000
2000
3000
4000
5000

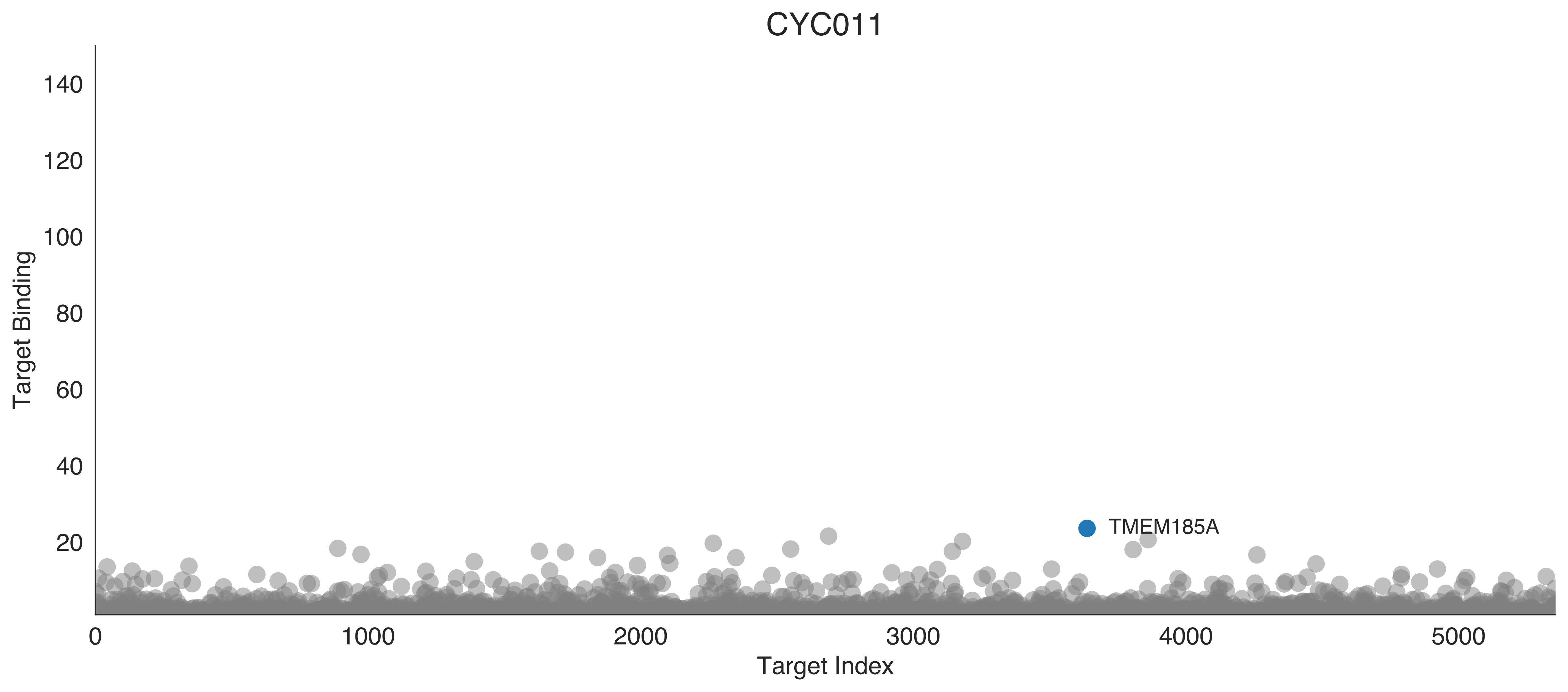
CYC011
Target Binding
Target Index
TMEM185A
0
20
40
60
80
100
120
140
1000
2000
3000
4000
5000

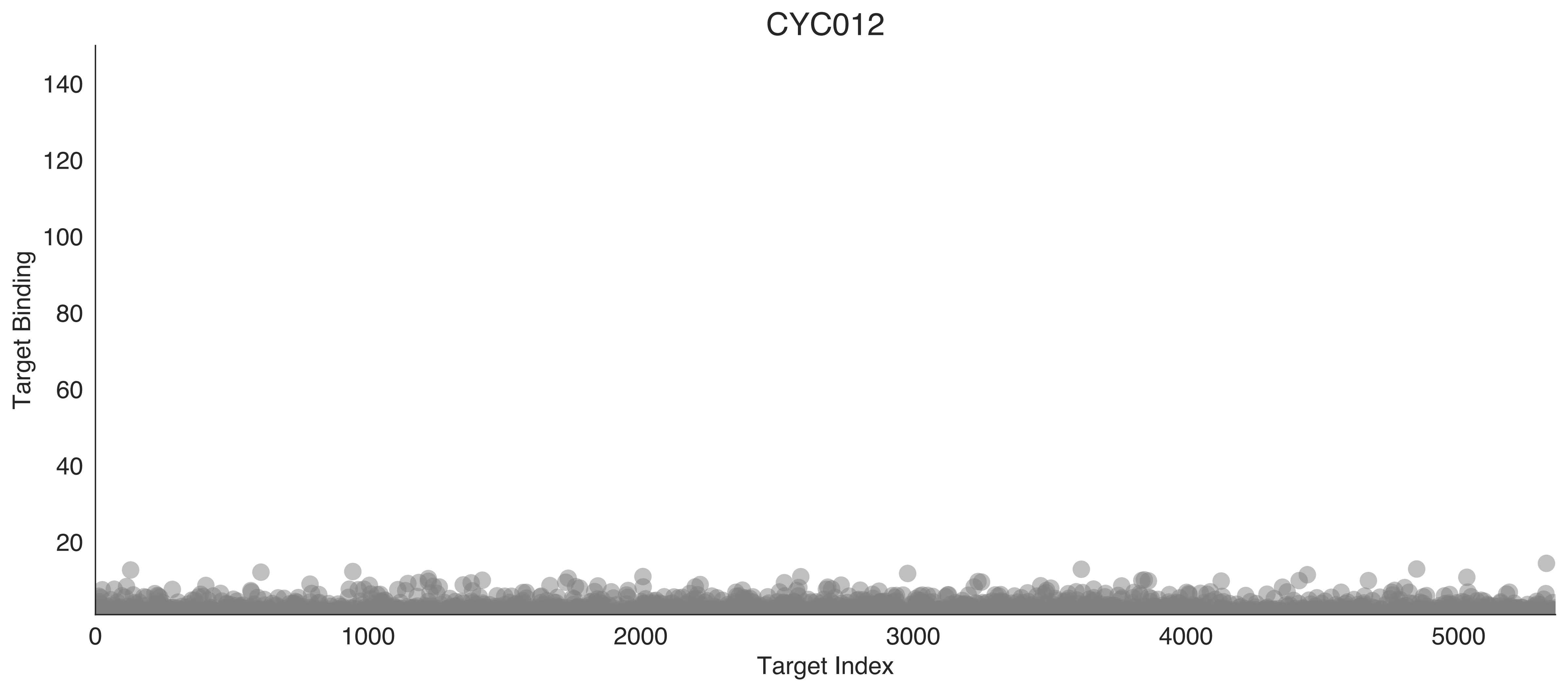
CYC012
Target Binding
Target Index
0
20
40
60
80
100
120
140
1000
2000
3000
4000
5000

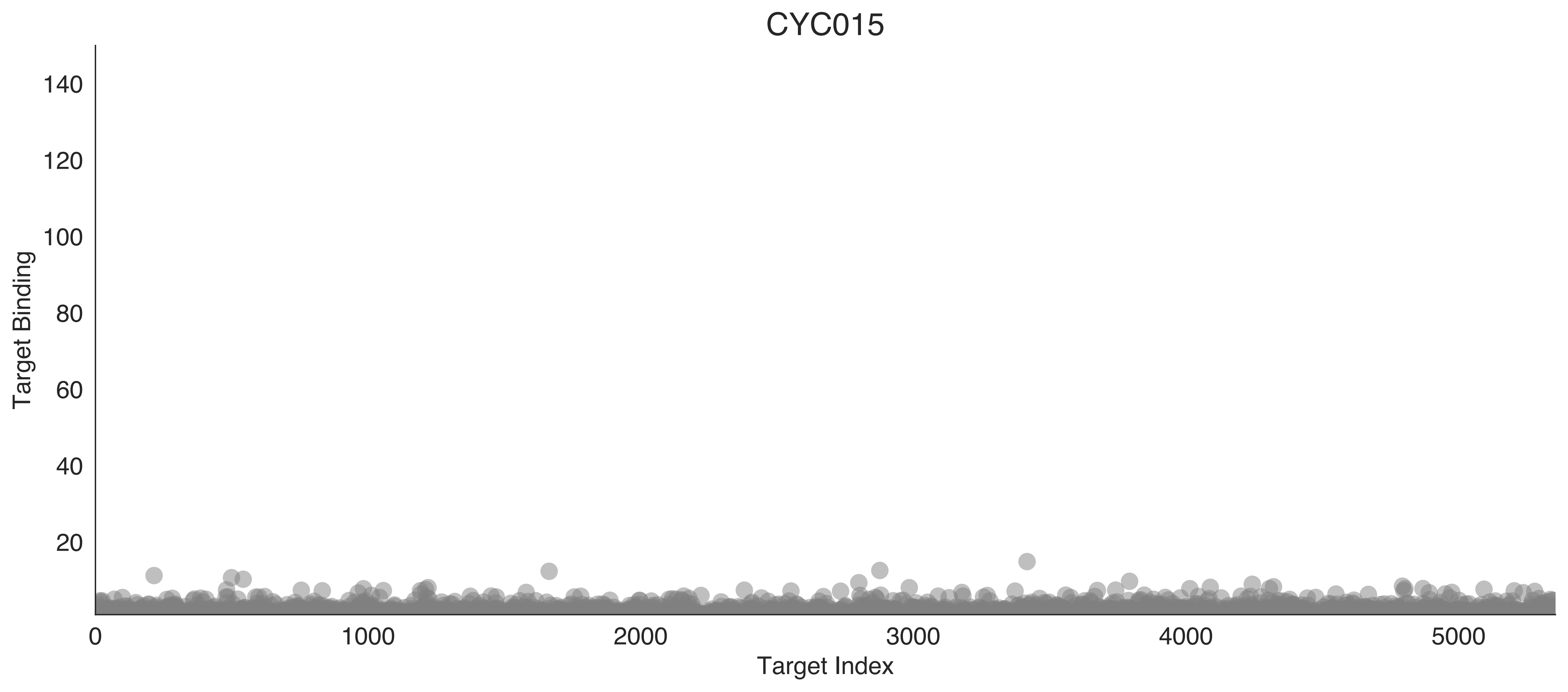

CYC015
Target Binding
Target Index
140
120
100
80
60
40
20
0
1000
2000
3000
4000
5000

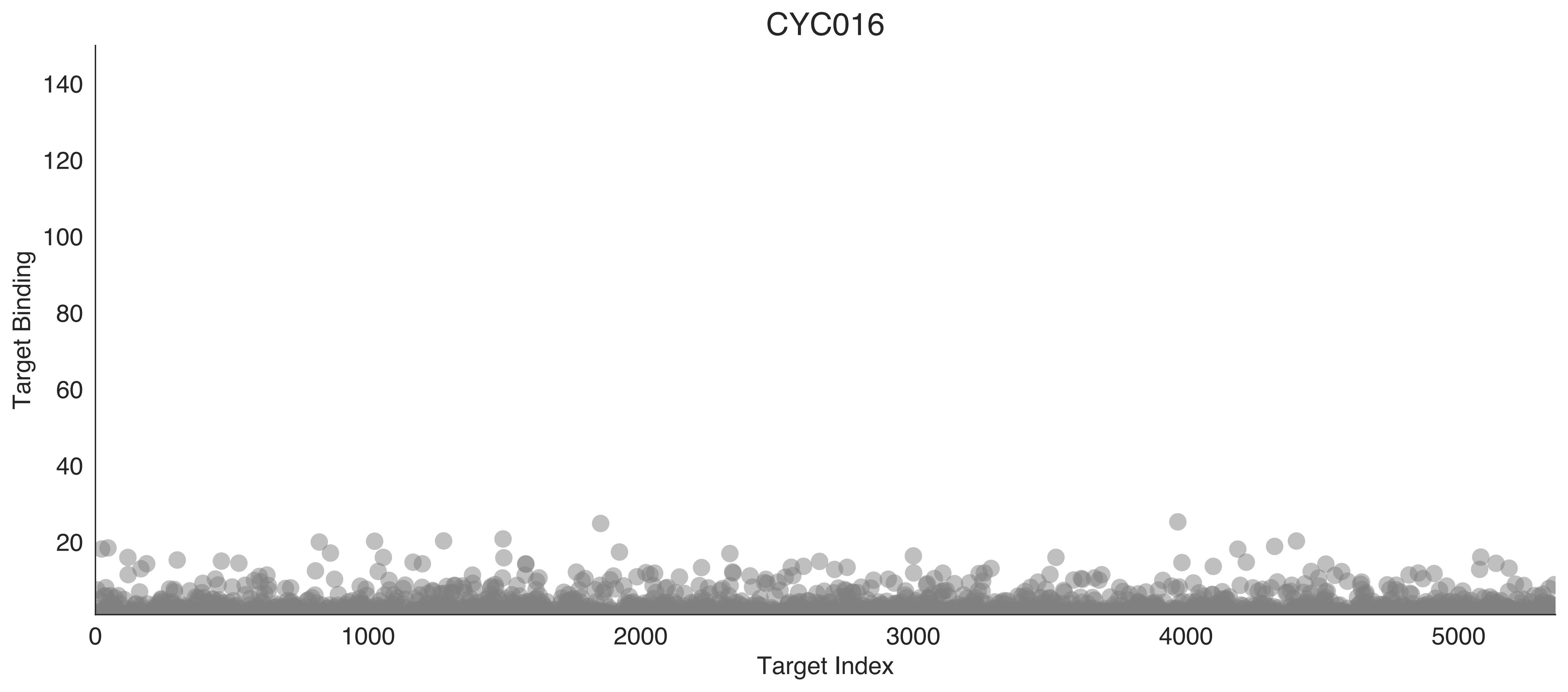

CYC016
Target Binding
140
120
100
80
60
40
20
0
1000
2000
3000
4000
5000
Target Index

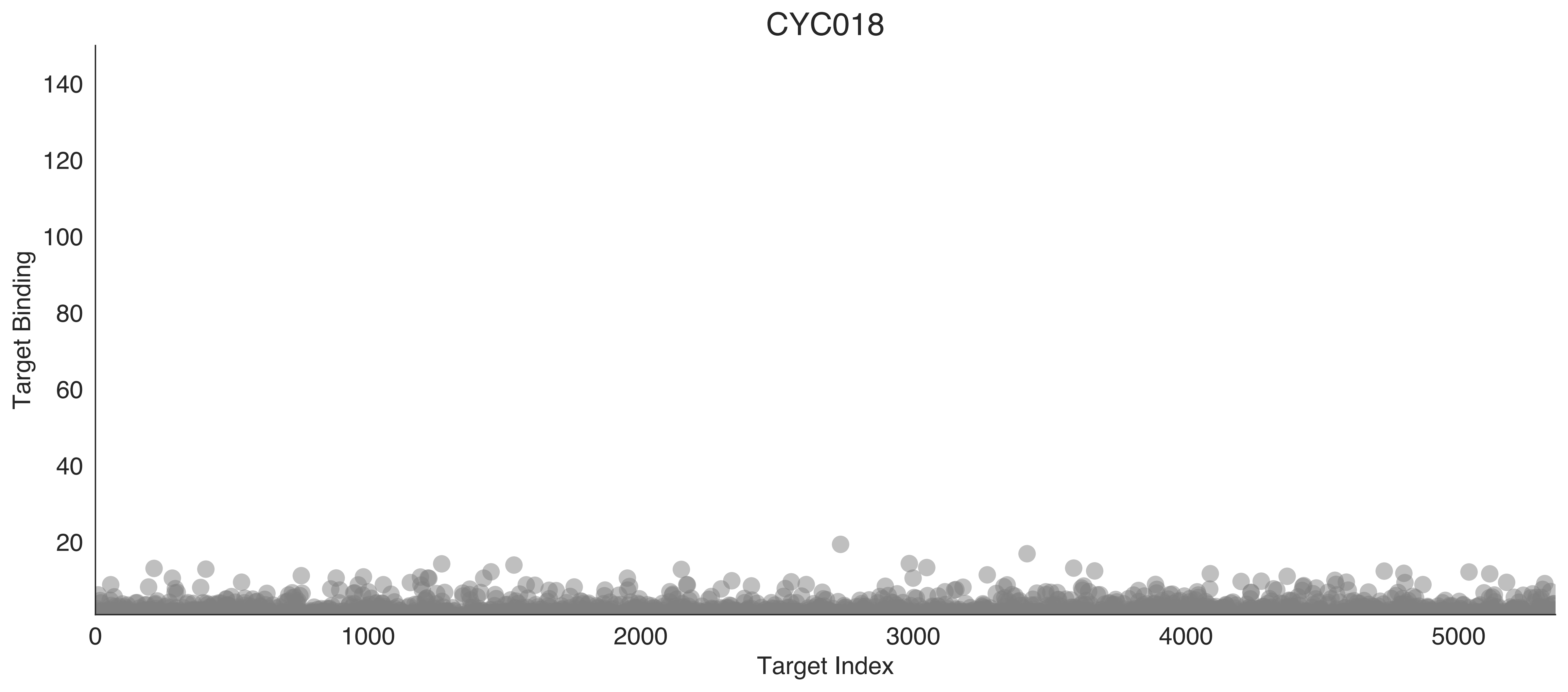

CYC018
Target Binding
140
120
100
80
60
40
20
0
1000
2000
3000
4000
5000
Target Index

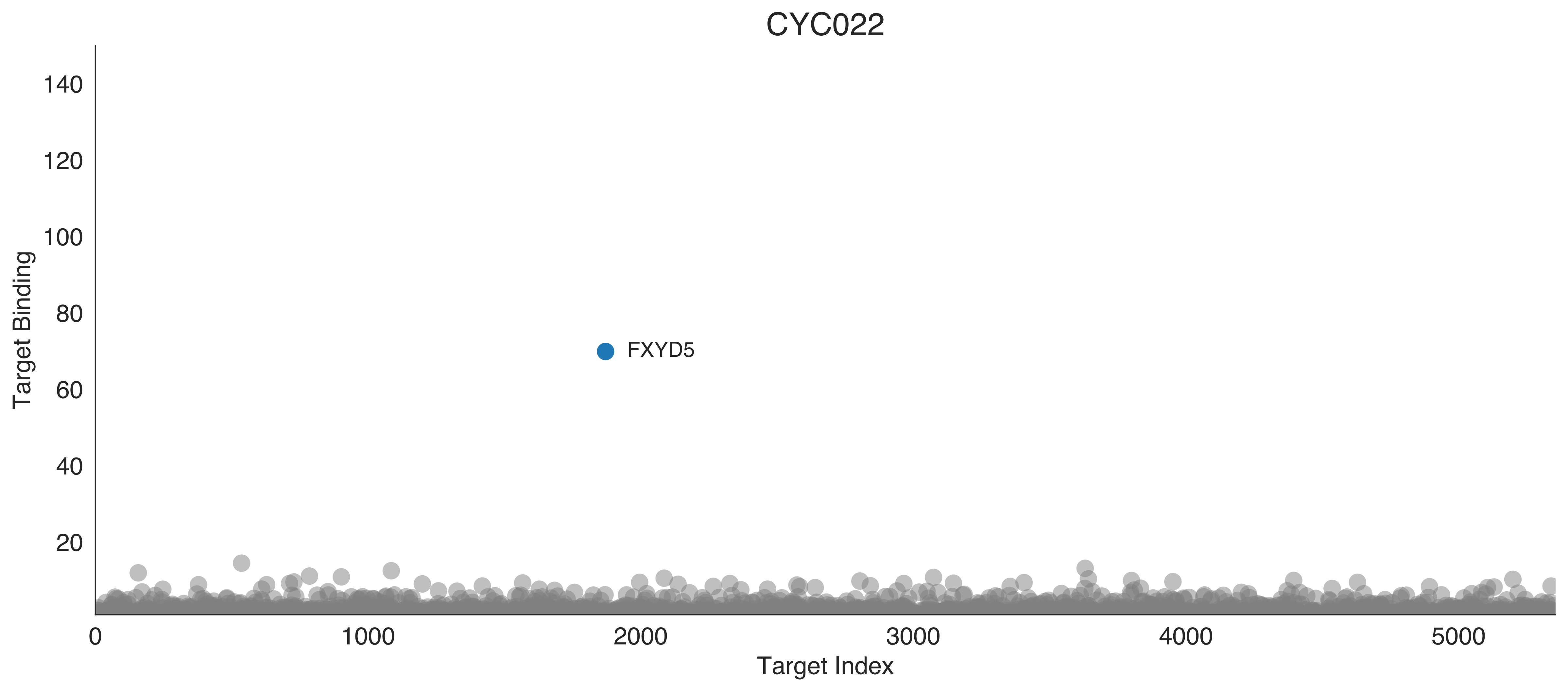

CYC022
Target Binding
Target Index
140
120
100
80
60
40
20
0
1000
2000
3000
4000
5000
FXYD5

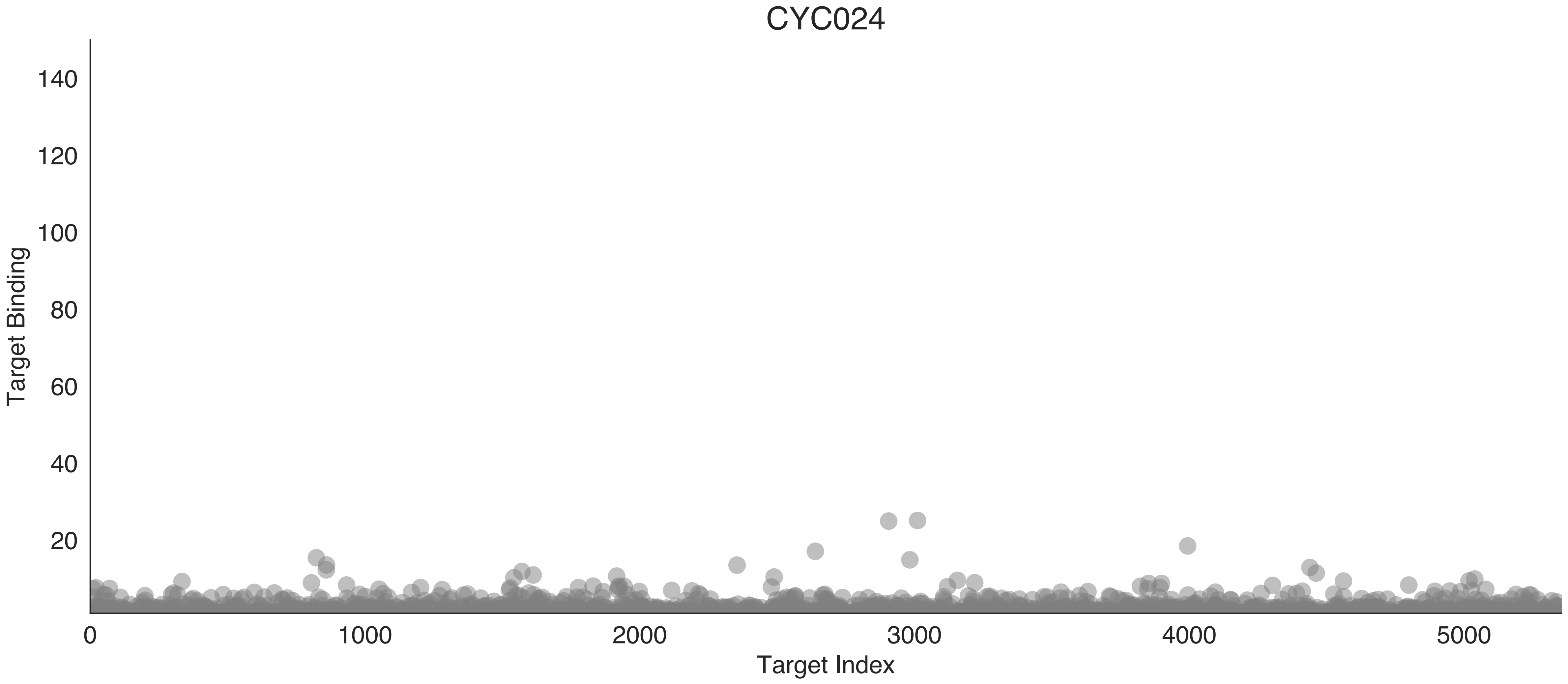
CYC024
Target Binding
140
120
100
80
60
40
20
0
1000
2000
3000
4000
5000
Target Index

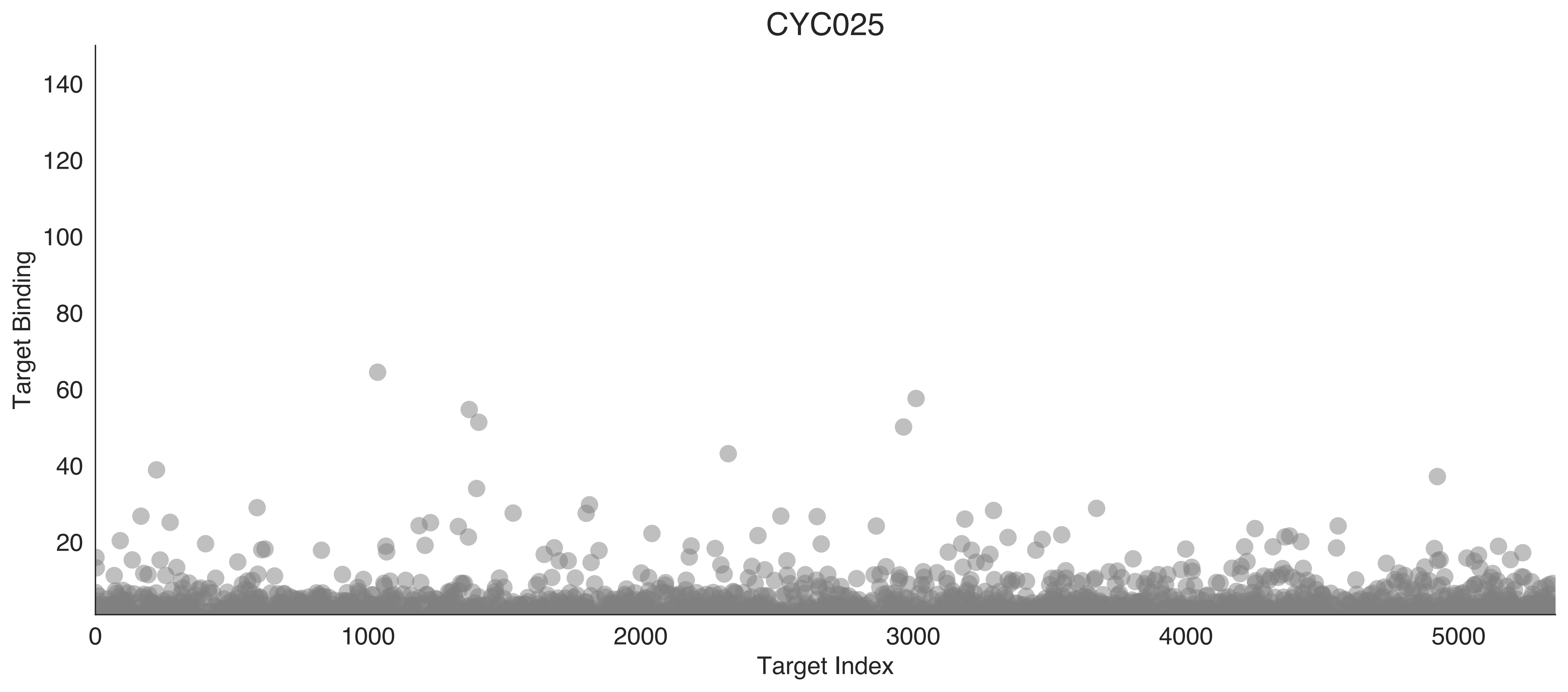
CYC025
Target Binding
Target Index
0
20
40
60
80
100
120
140
1000
2000
3000
4000
5000

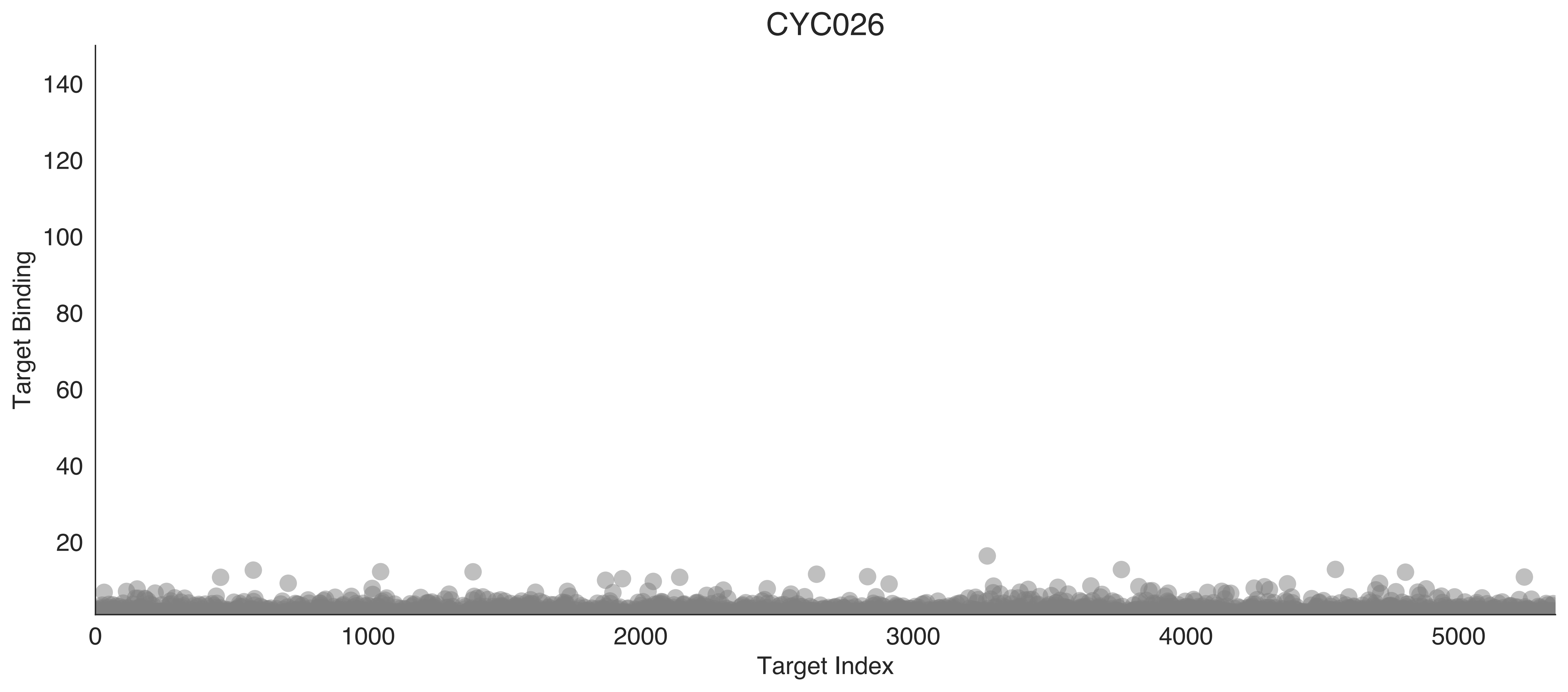

CYC026
Target Binding
Target Index
140
120
100
80
60
40
20
0
1000
2000
3000
4000
5000

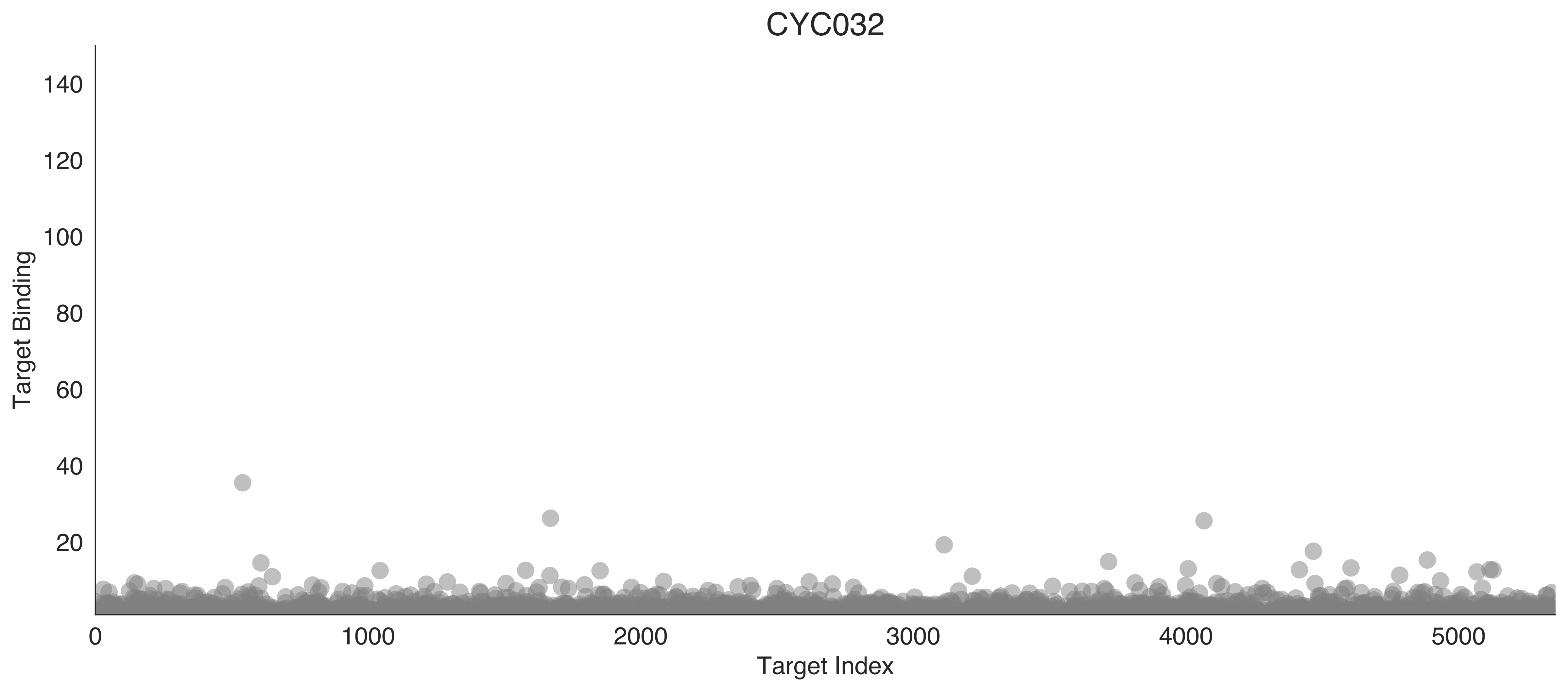

CYC032
Target Binding
140
120
100
80
60
40
20
0
1000
2000
3000
4000
5000
Target Index

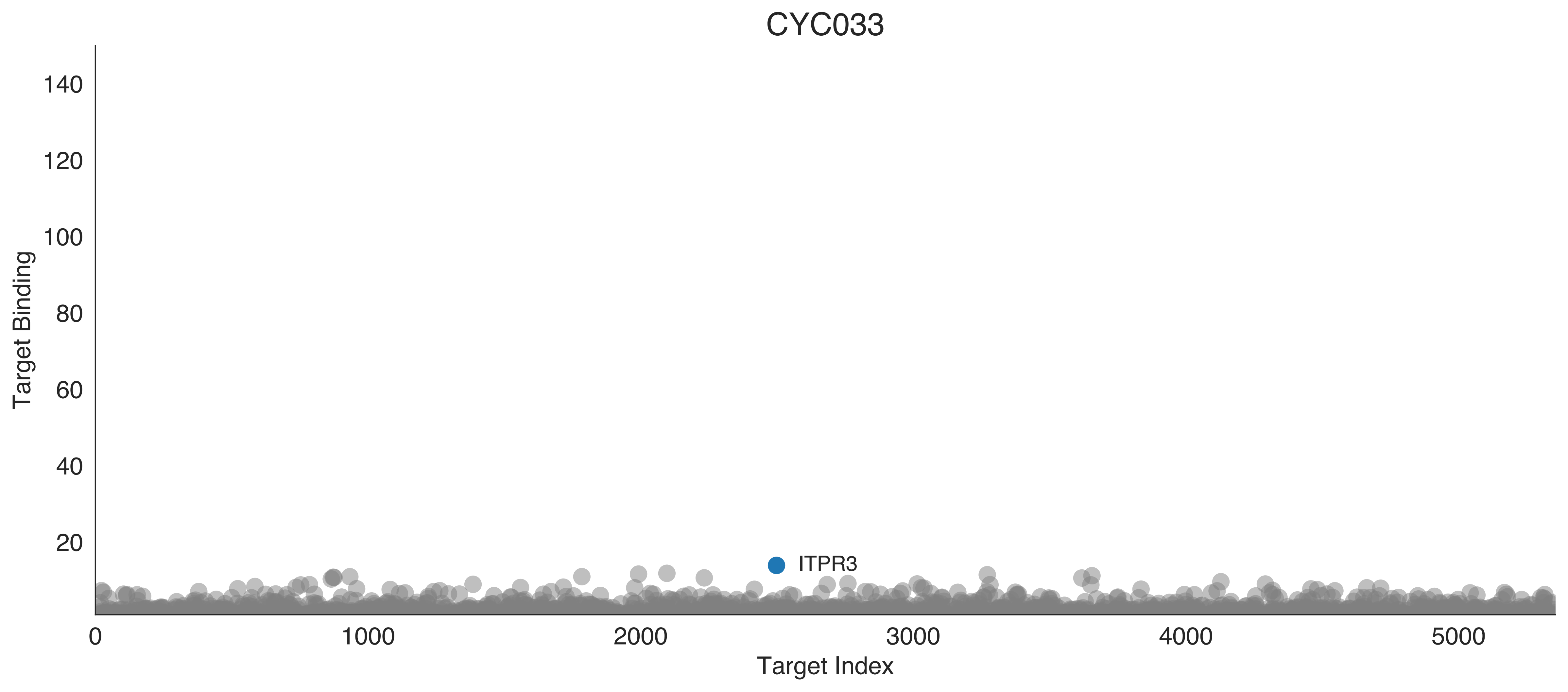
CYC033
Target Binding
140
120
100
80
60
40
20
ITPR3
0
1000
2000
3000
4000
5000
Target Index

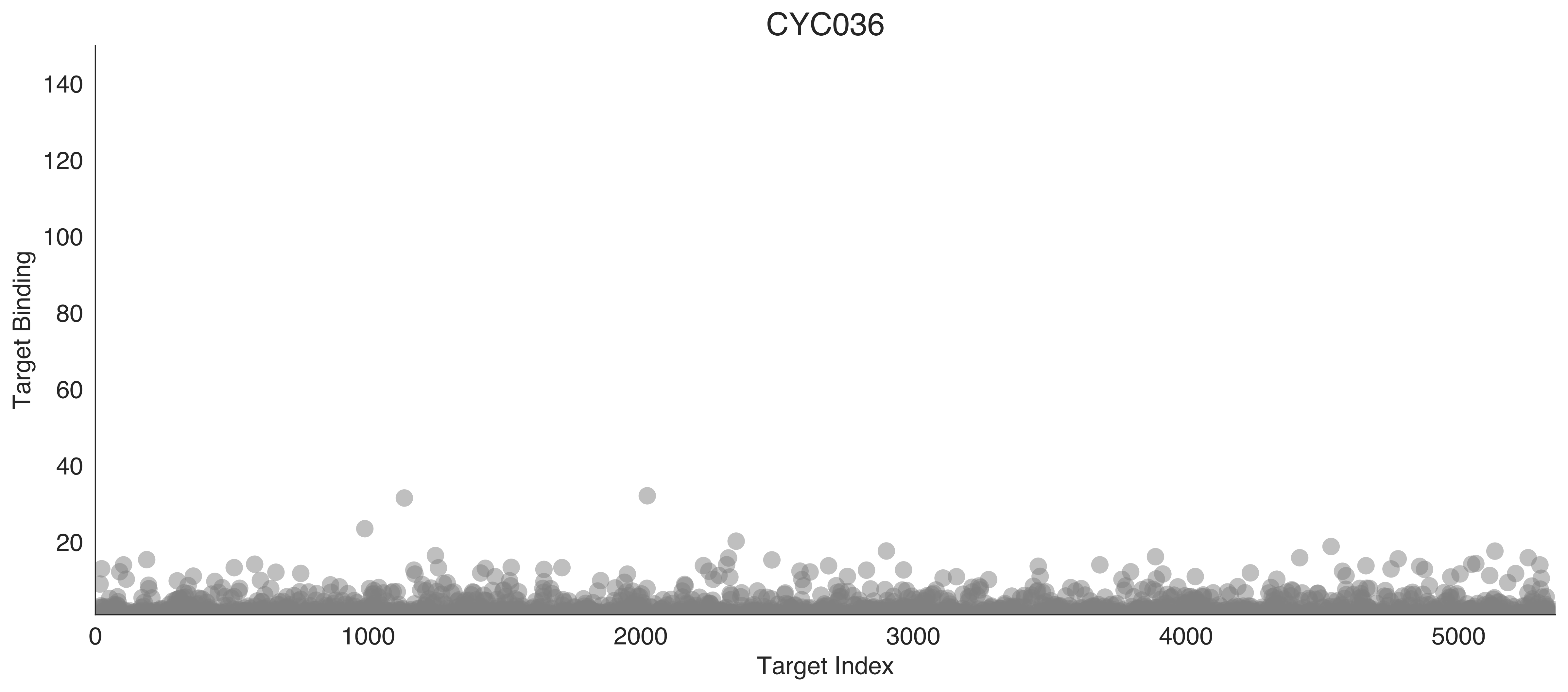
CYC036
Target Binding
140
120
100
80
60
40
20
0
1000
2000
3000
4000
5000
Target Index

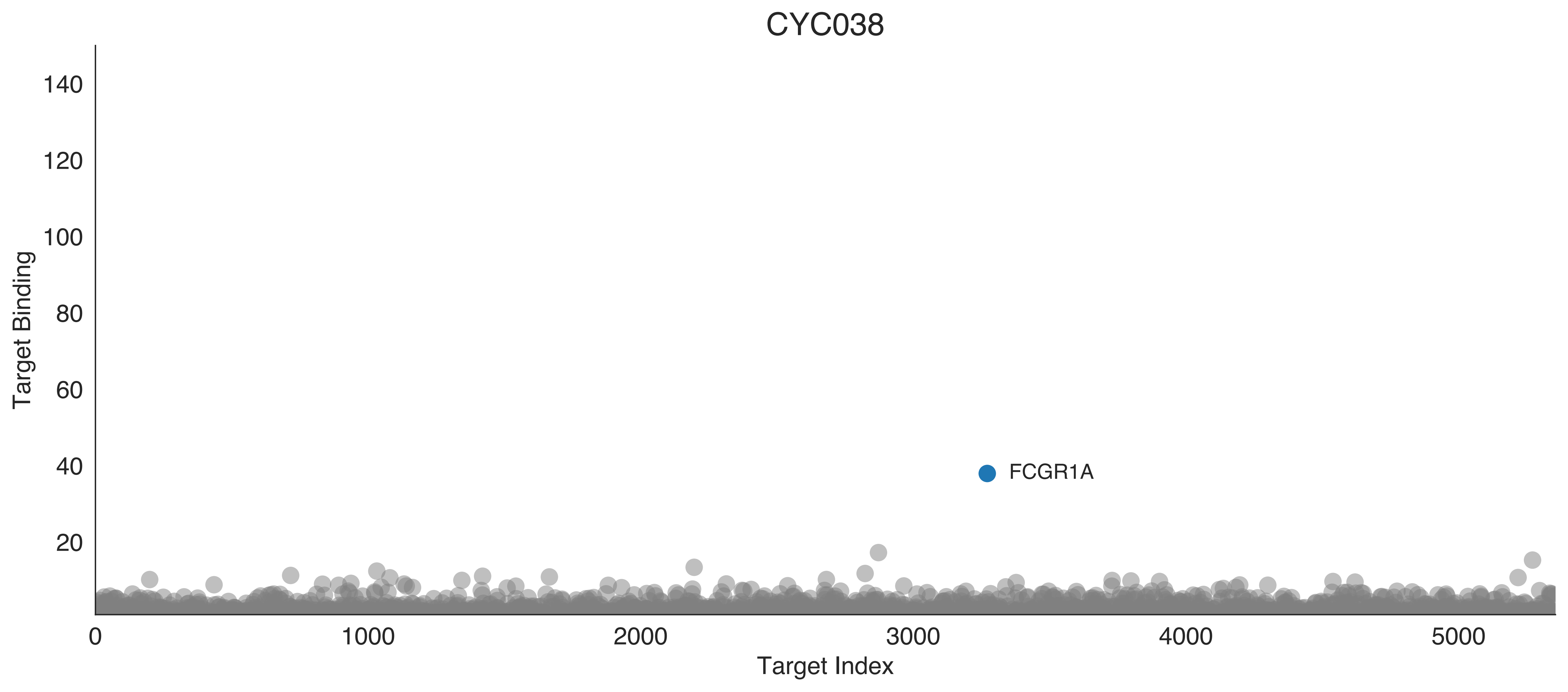

CYC038
Target Binding
140
120
100
80
60
40
20
FCGR1A
0
1000
2000
3000
4000
5000
Target Index

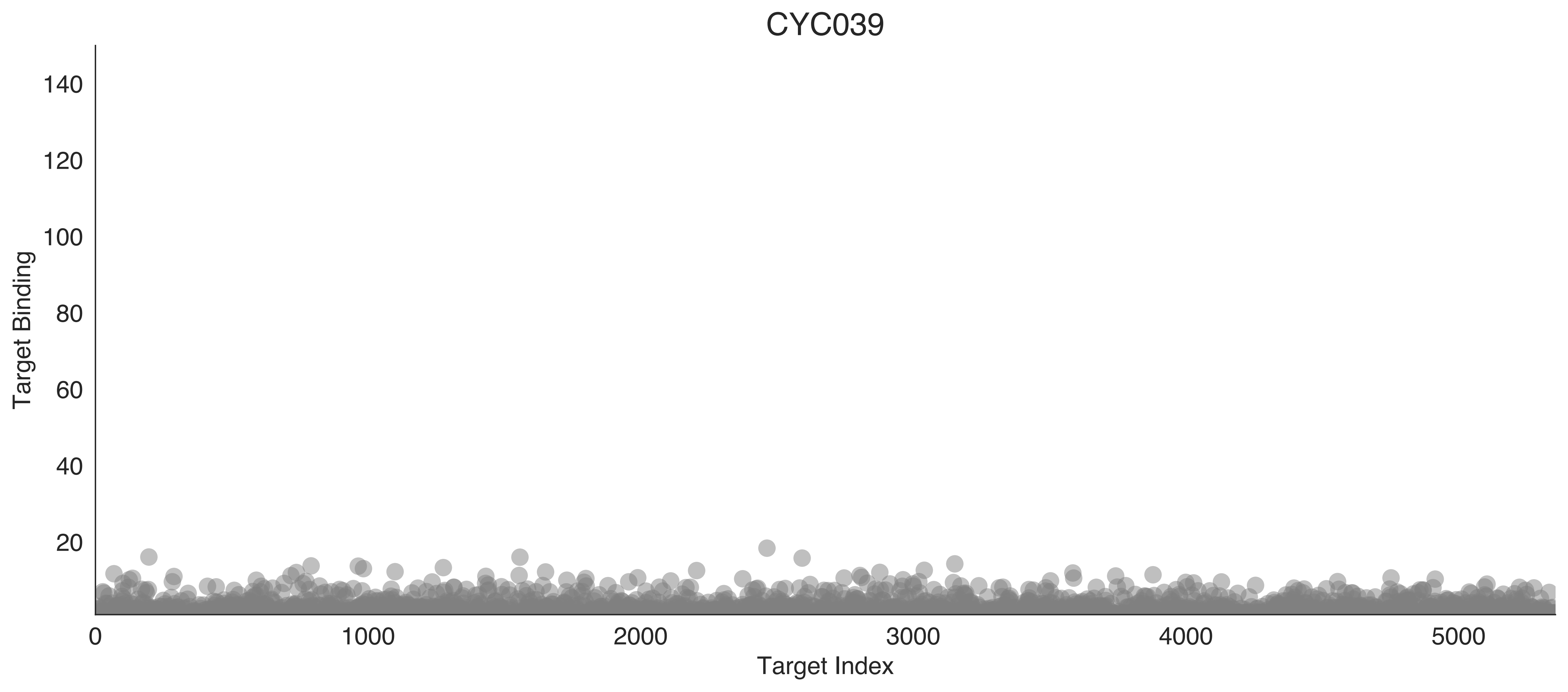

CYC039
Target Binding
Target Index
0
20
40
60
80
100
120
140
1000
2000
3000
4000
5000

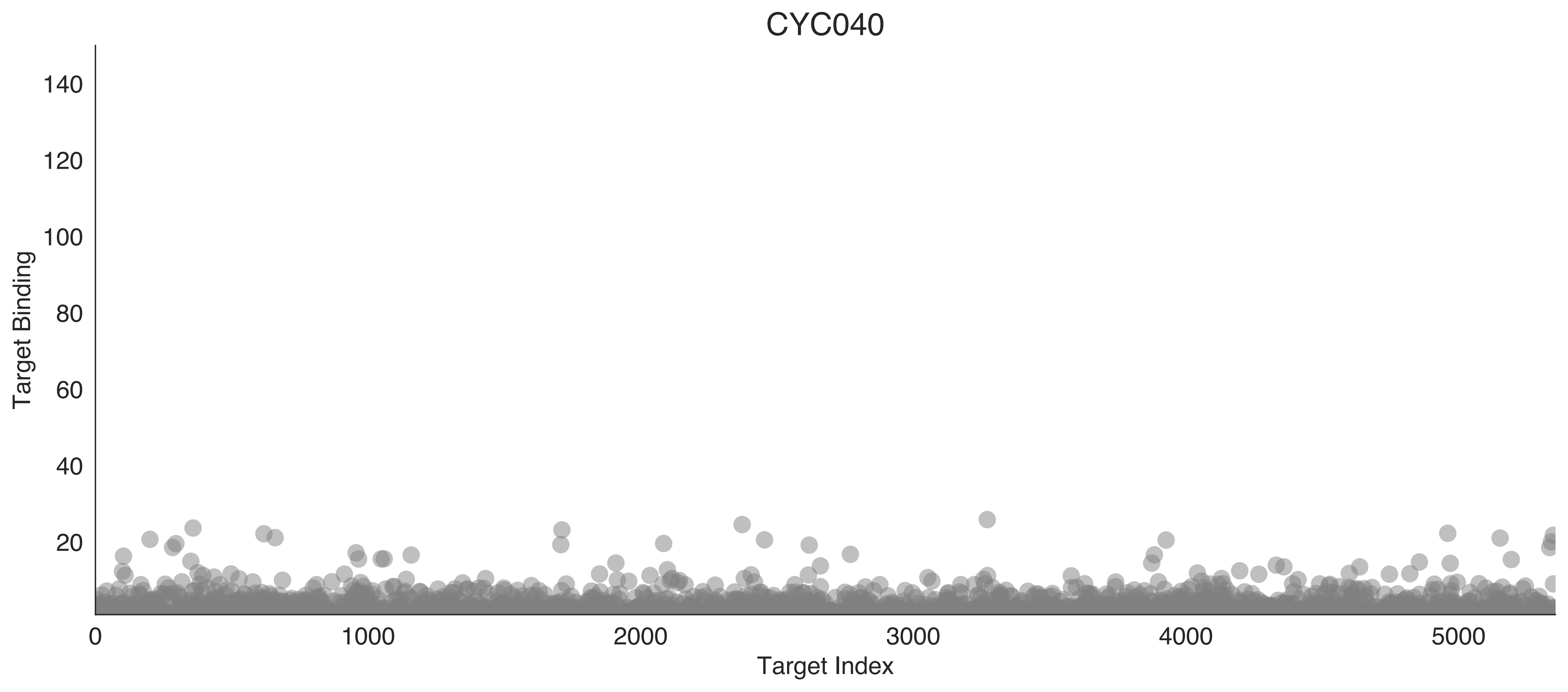

CYC040
Target Binding
Target Index
140
120
100
80
60
40
20
0
1000
2000
3000
4000
5000

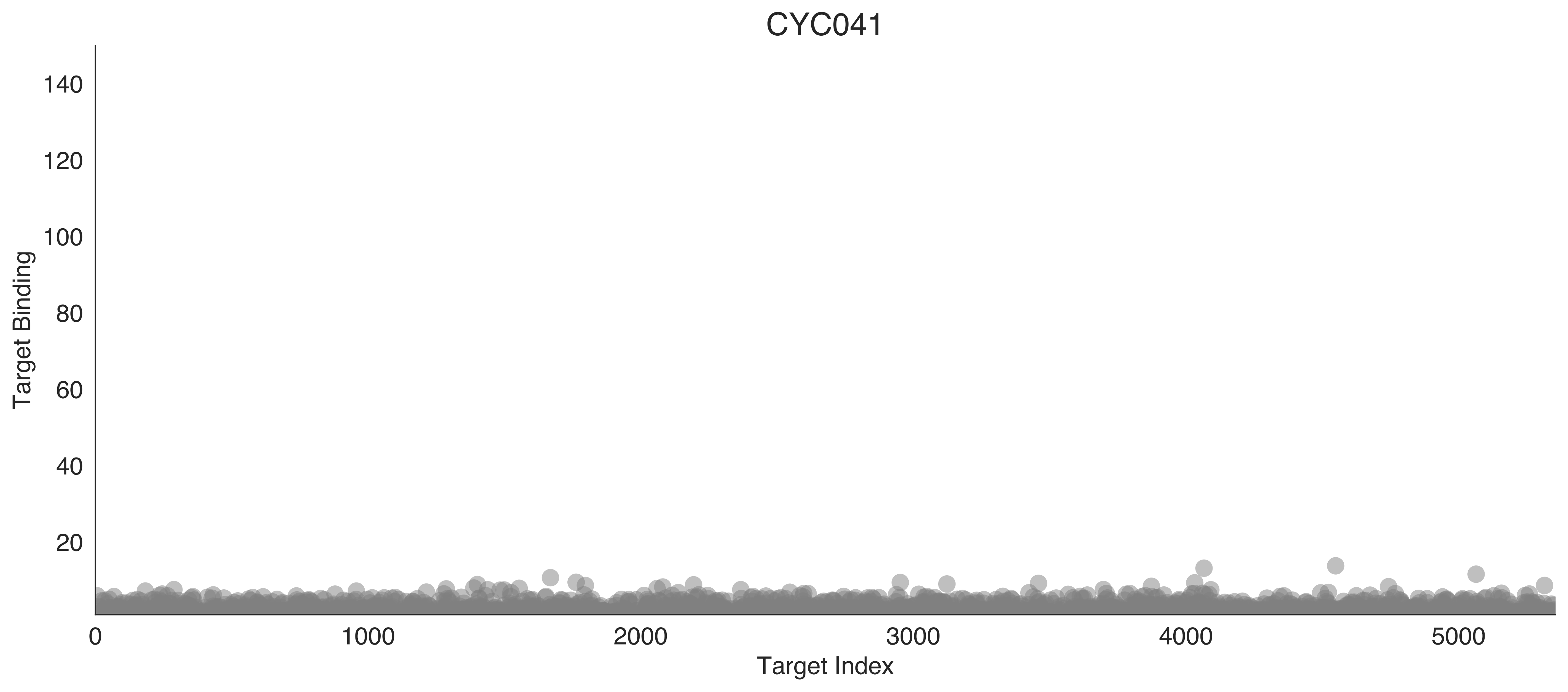

CYC041
Target Binding
140
120
100
80
60
40
20
0
1000
2000
3000
4000
5000
Target Index

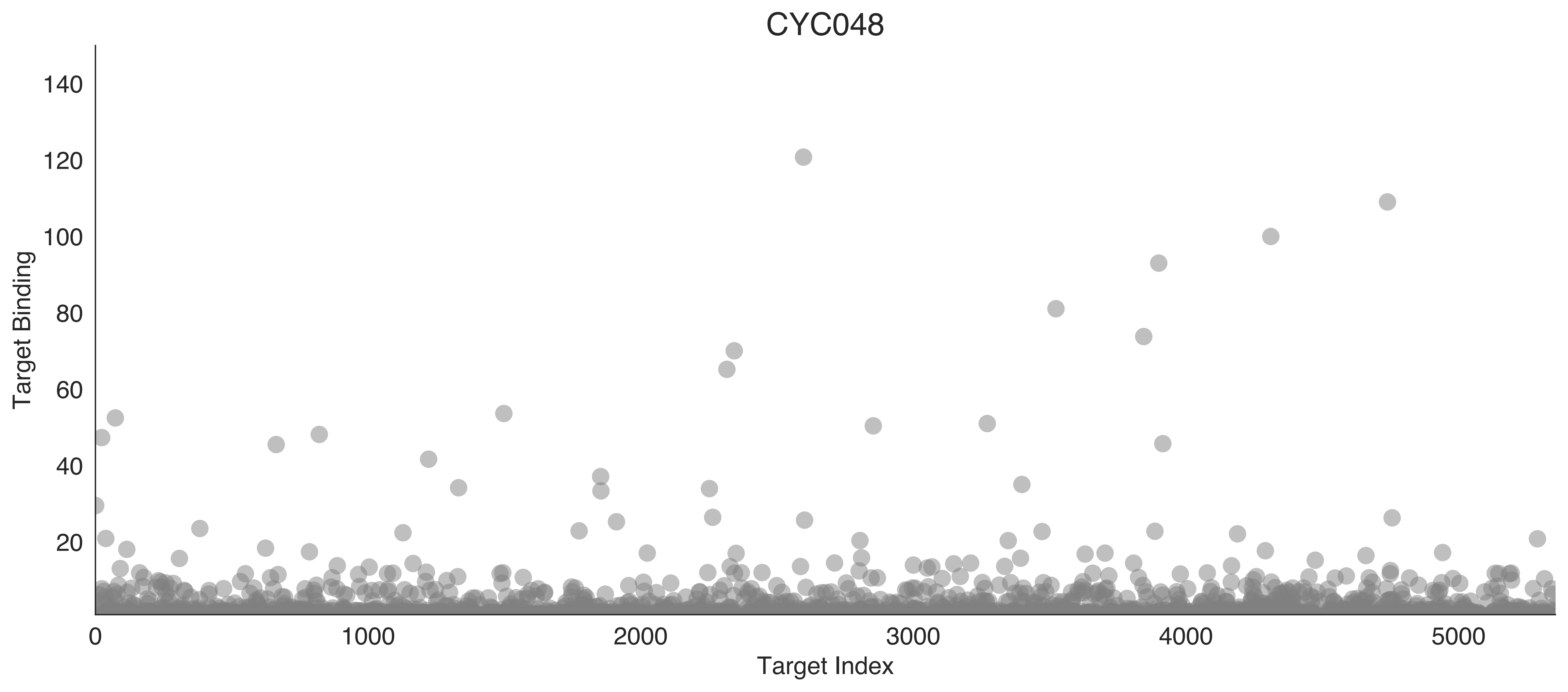
CYC048
Target Binding
Target Index
0
20
40
60
80
100
120
140
1000
2000
3000
4000
5000

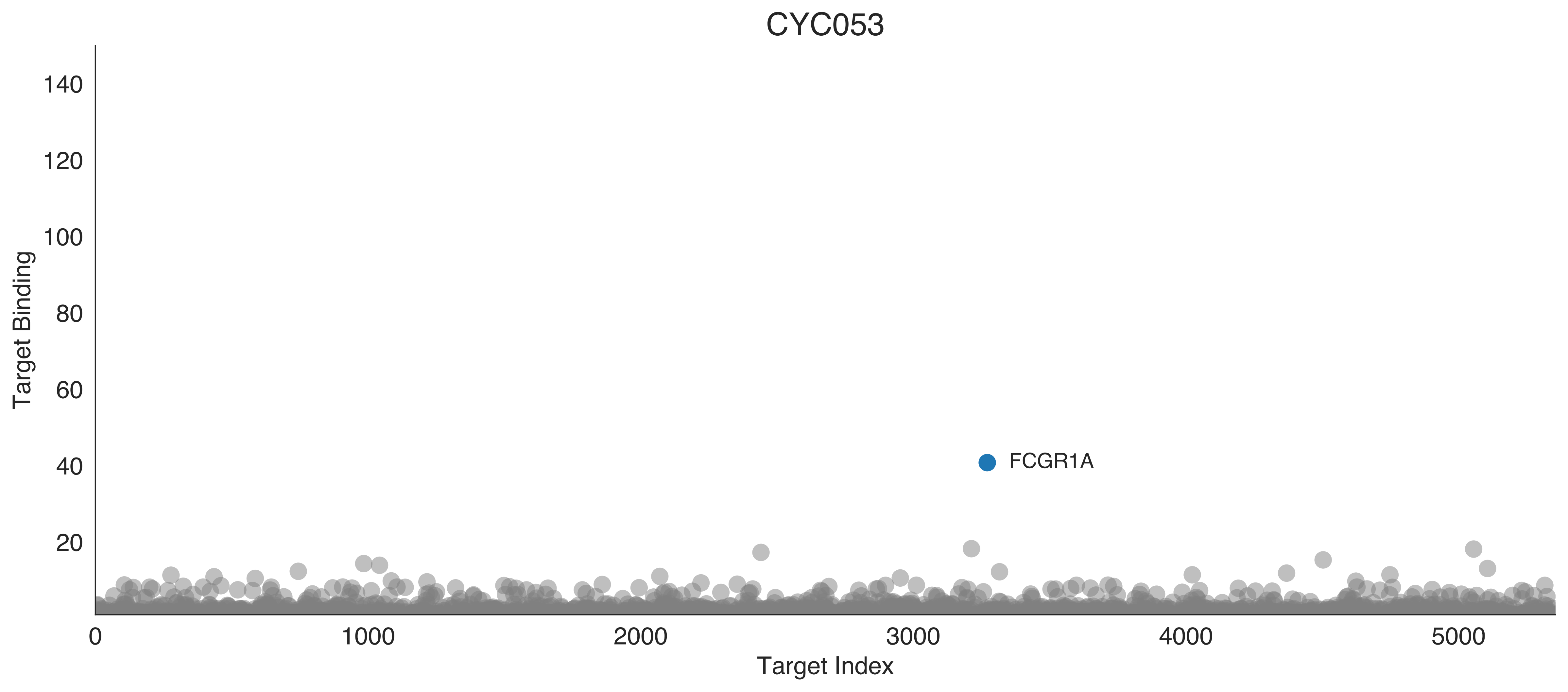

CYC053
Target Binding
Target Index
FCGR1A
140
120
100
80
60
40
20
0
1000
2000
3000
4000
5000

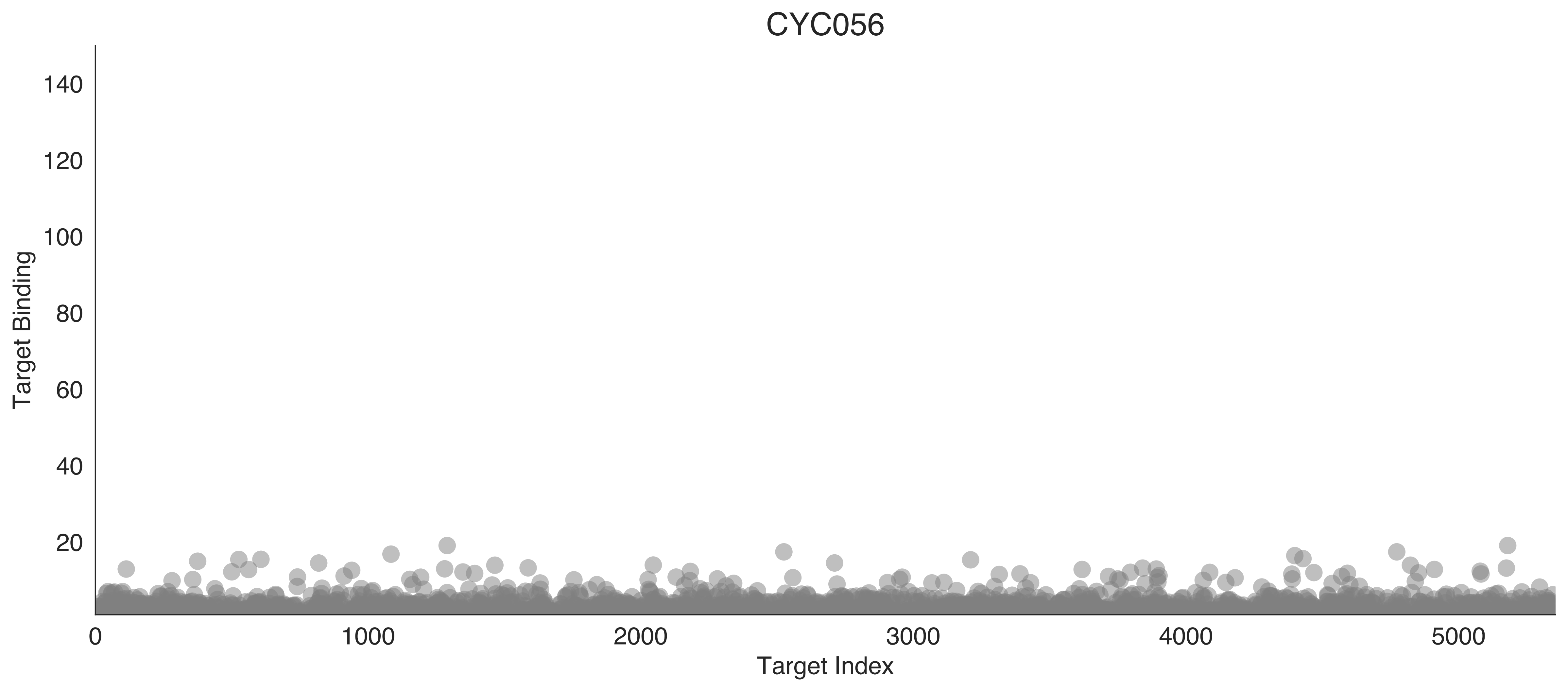
CYC056
Target Binding
Target Index
140
120
100
80
60
40
20
0
1000
2000
3000
4000
5000

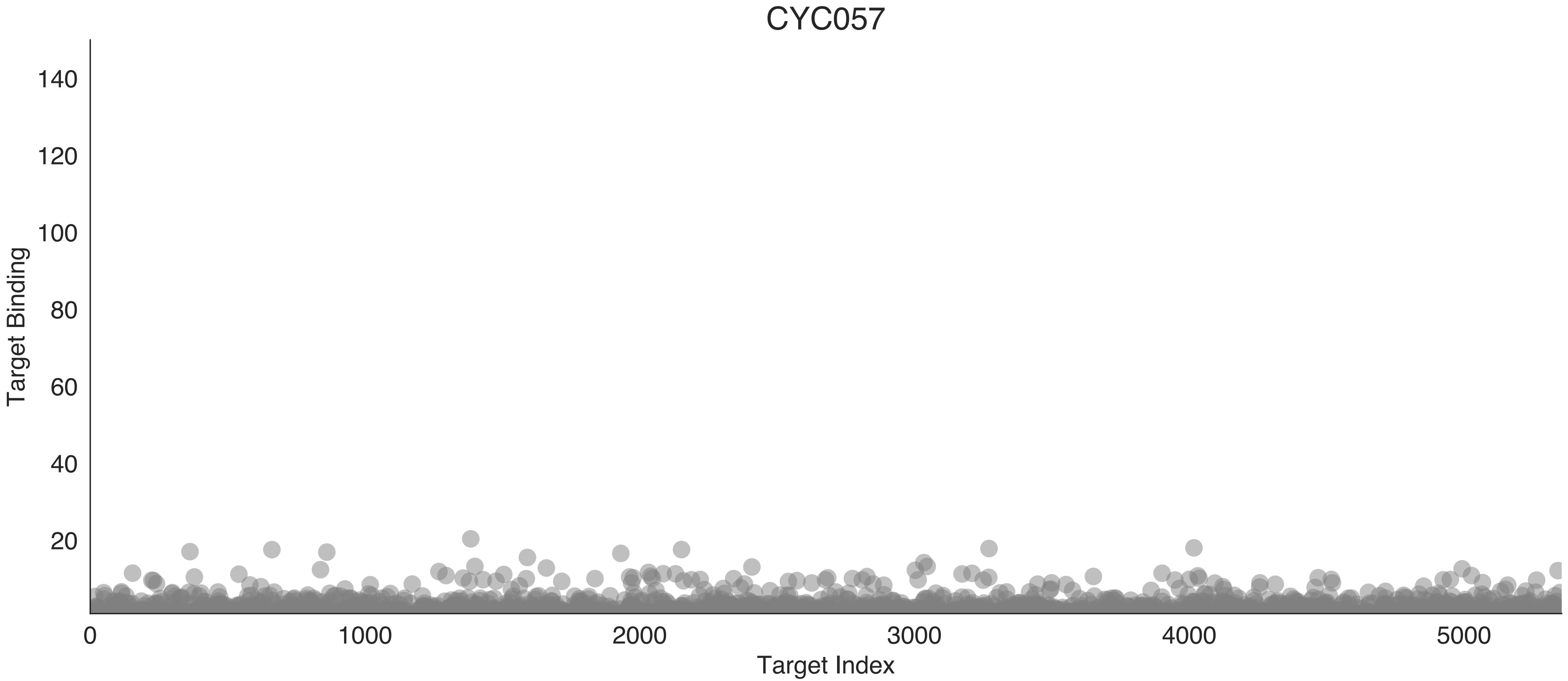

CYC057
Target Binding
Target Index
0
20
40
60
80
100
120
140
0
1000
2000
3000
4000
5000

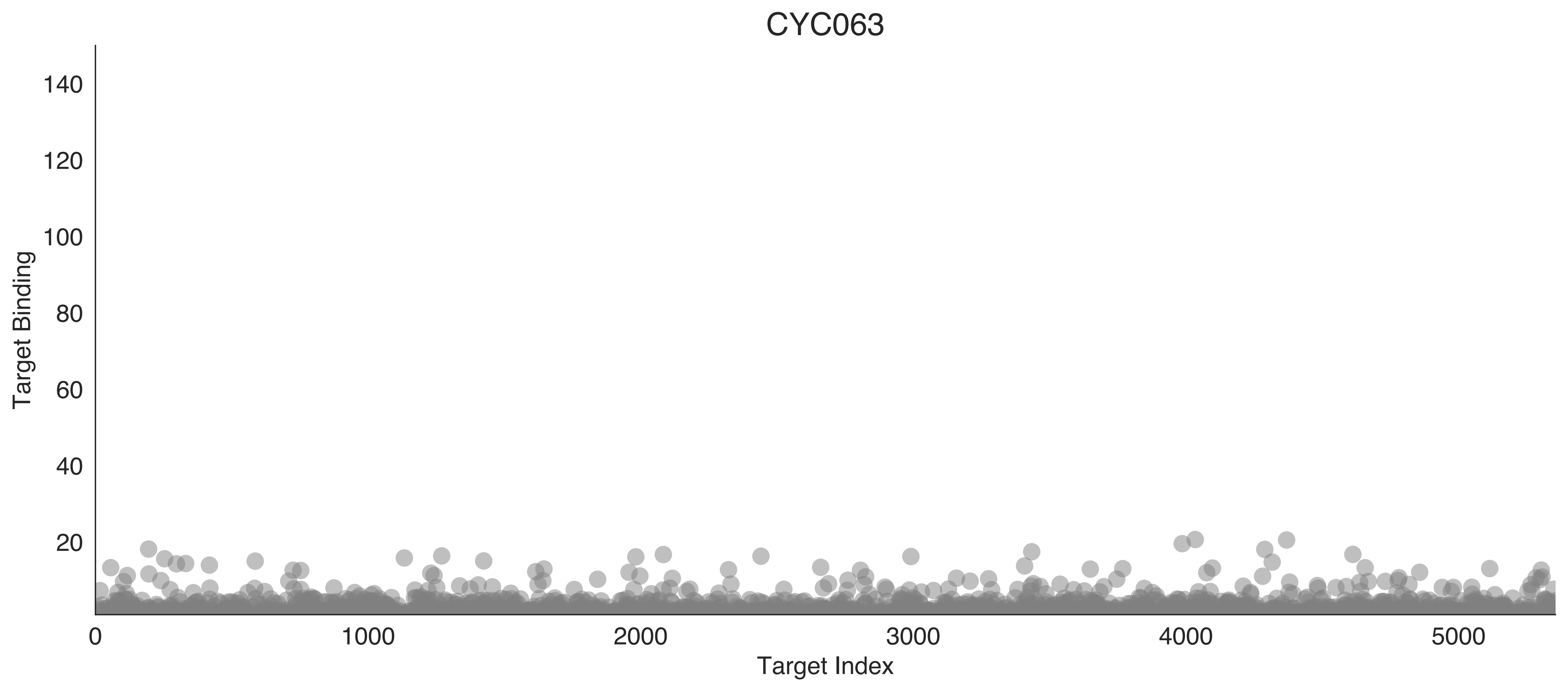
CYC063
Target Binding
Target Index
140
120
100
80
60
40
20
0
1000
2000
3000
4000
5000

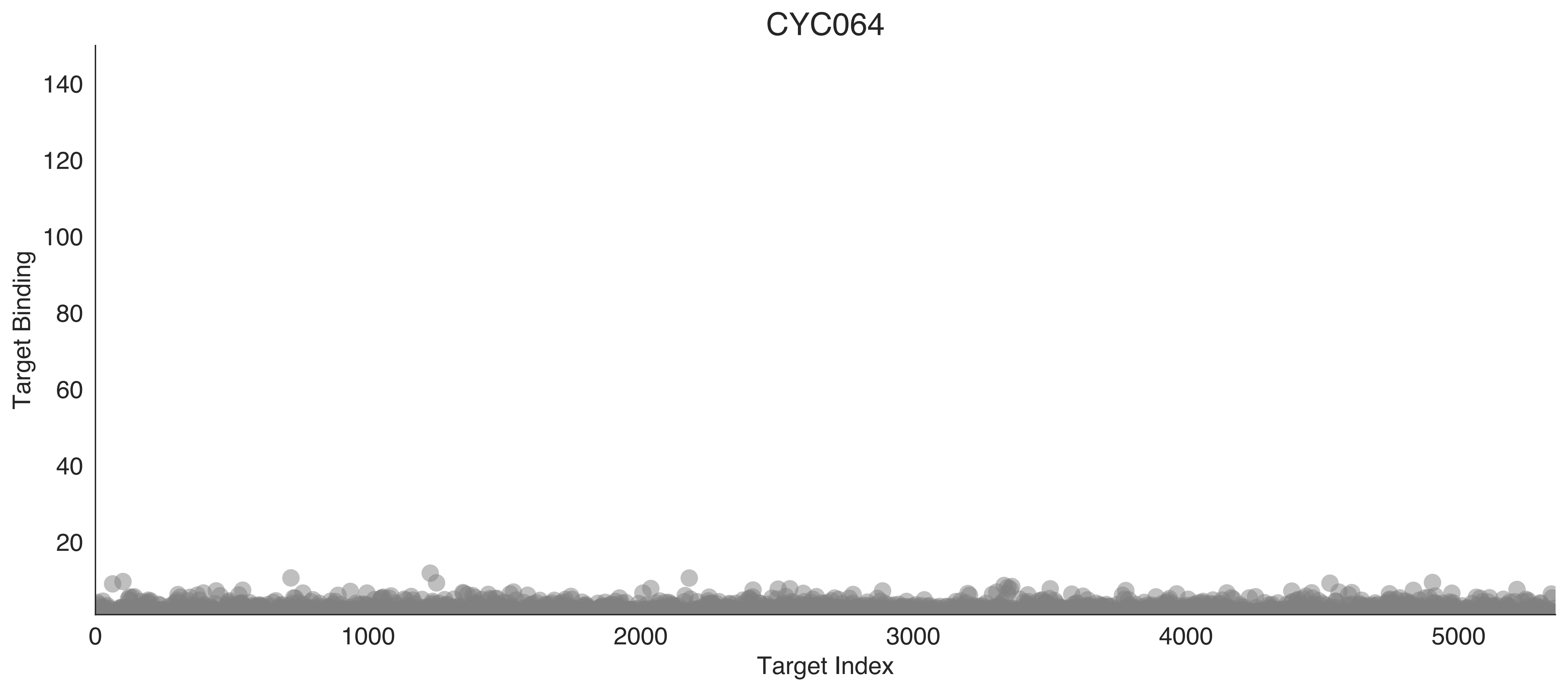
CYC064
Target Binding
140
120
100
80
60
40
20
0
1000
2000
3000
4000
5000
Target Index

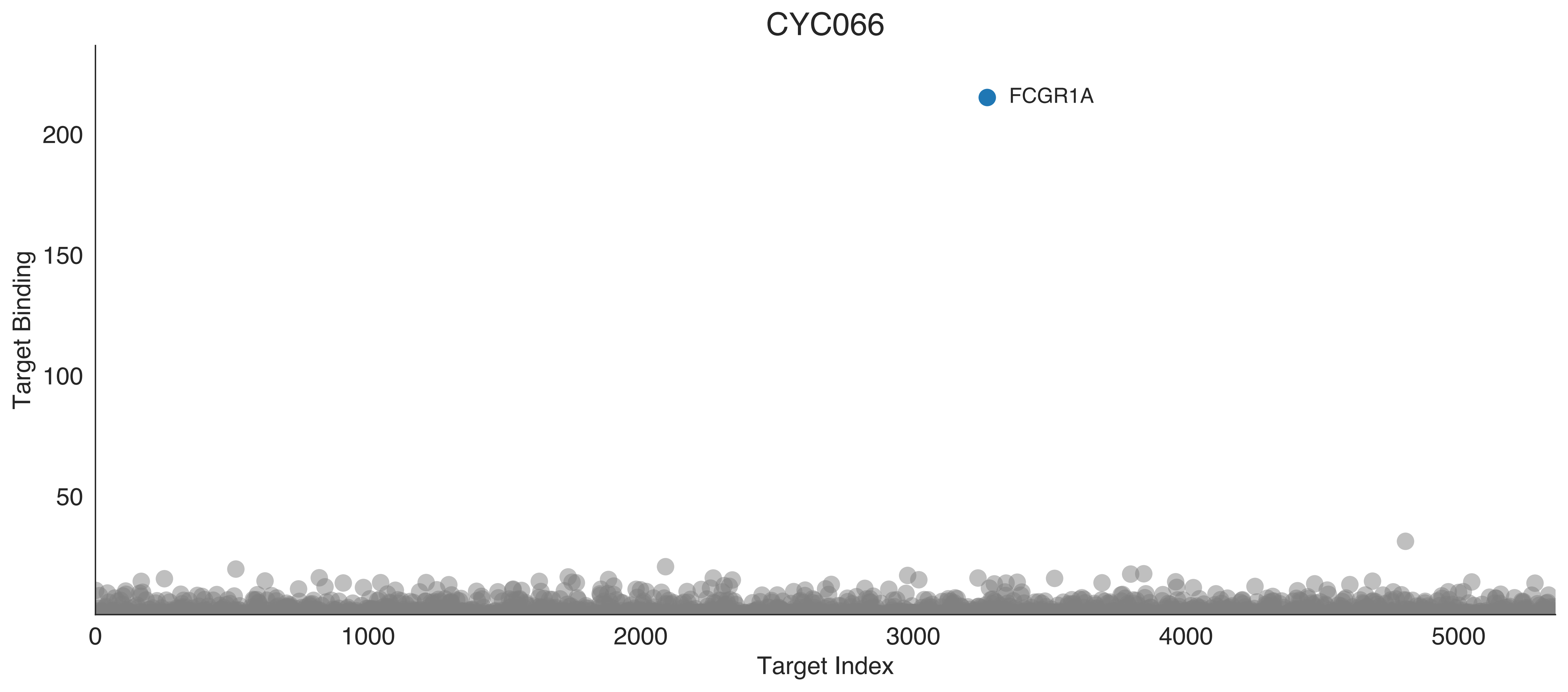

CYC066
FCGR1A
Target Binding
Target Index
0
50
100
150
200
0
1000
2000
3000
4000
5000

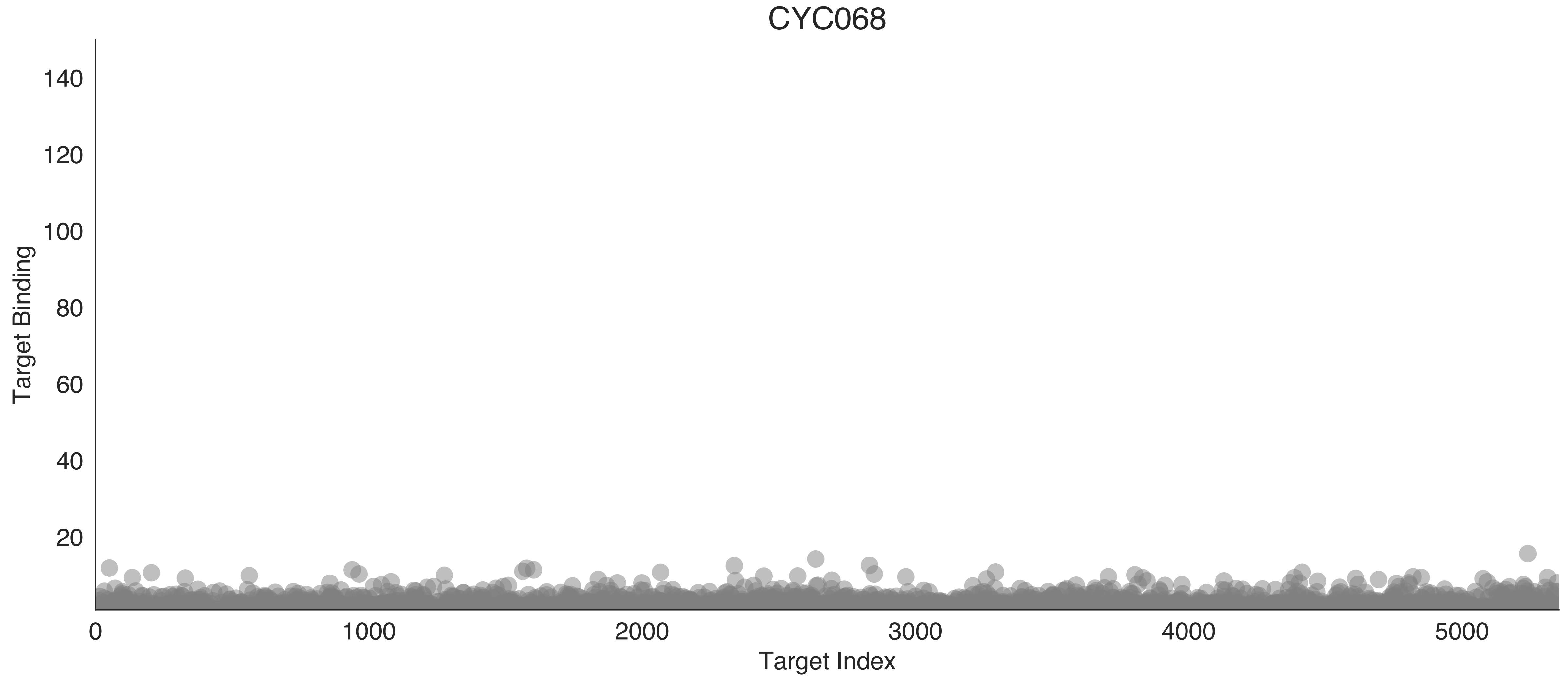

CYC068
Target Binding
140
120
100
80
60
40
20
0
1000
2000
3000
4000
5000
Target Index

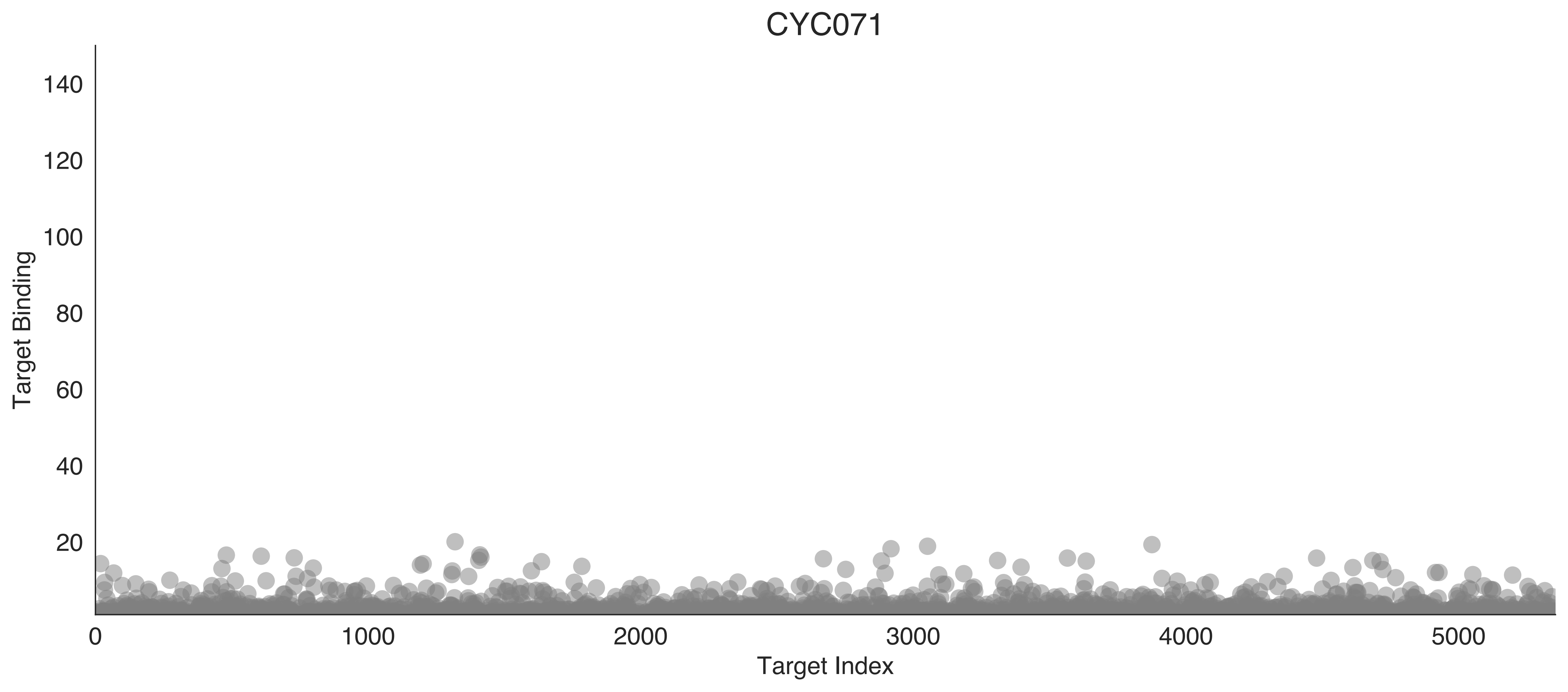
CYC071
Target Binding
140
120
100
80
60
40
20
0
1000
2000
3000
4000
5000
Target Index

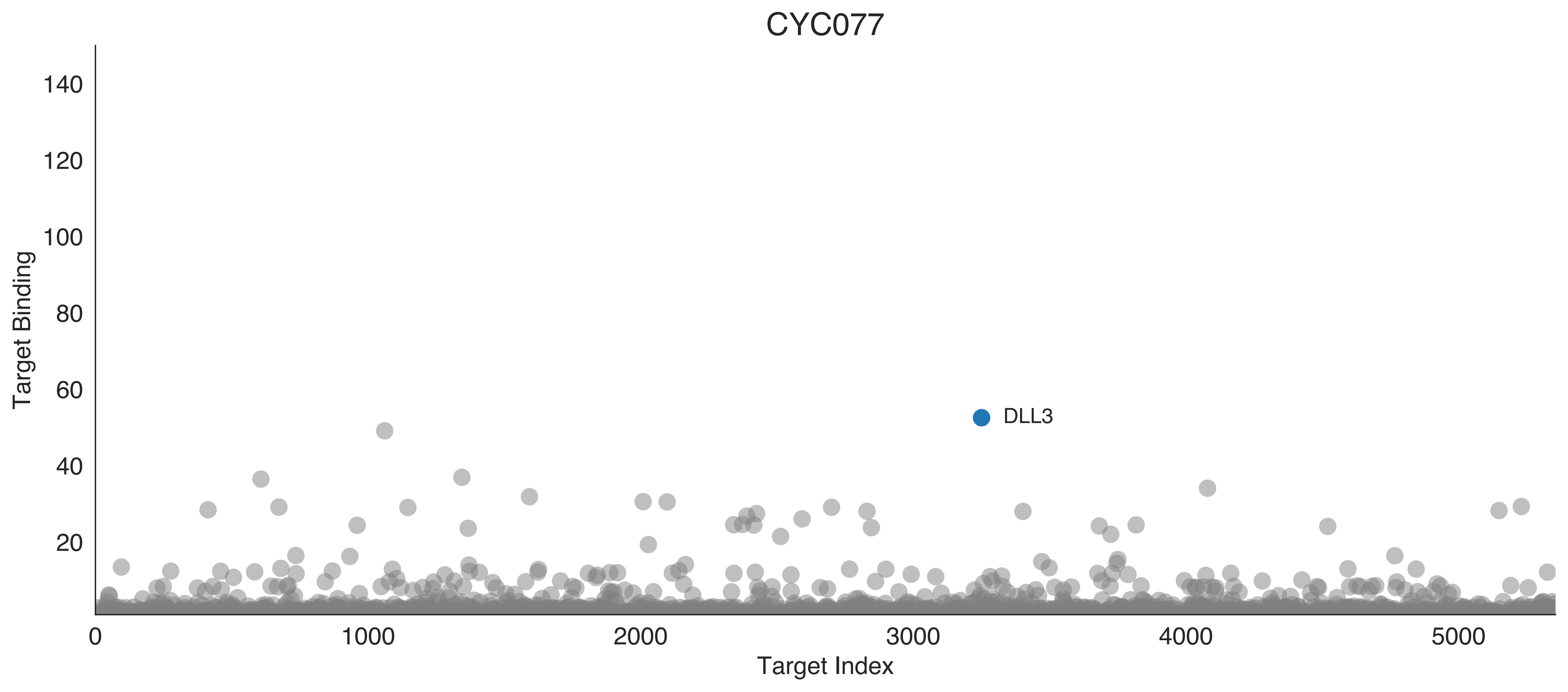

CYC077
Target Binding
Target Index
DLL3
140
120
100
80
60
40
20
0
1000
2000
3000
4000
5000

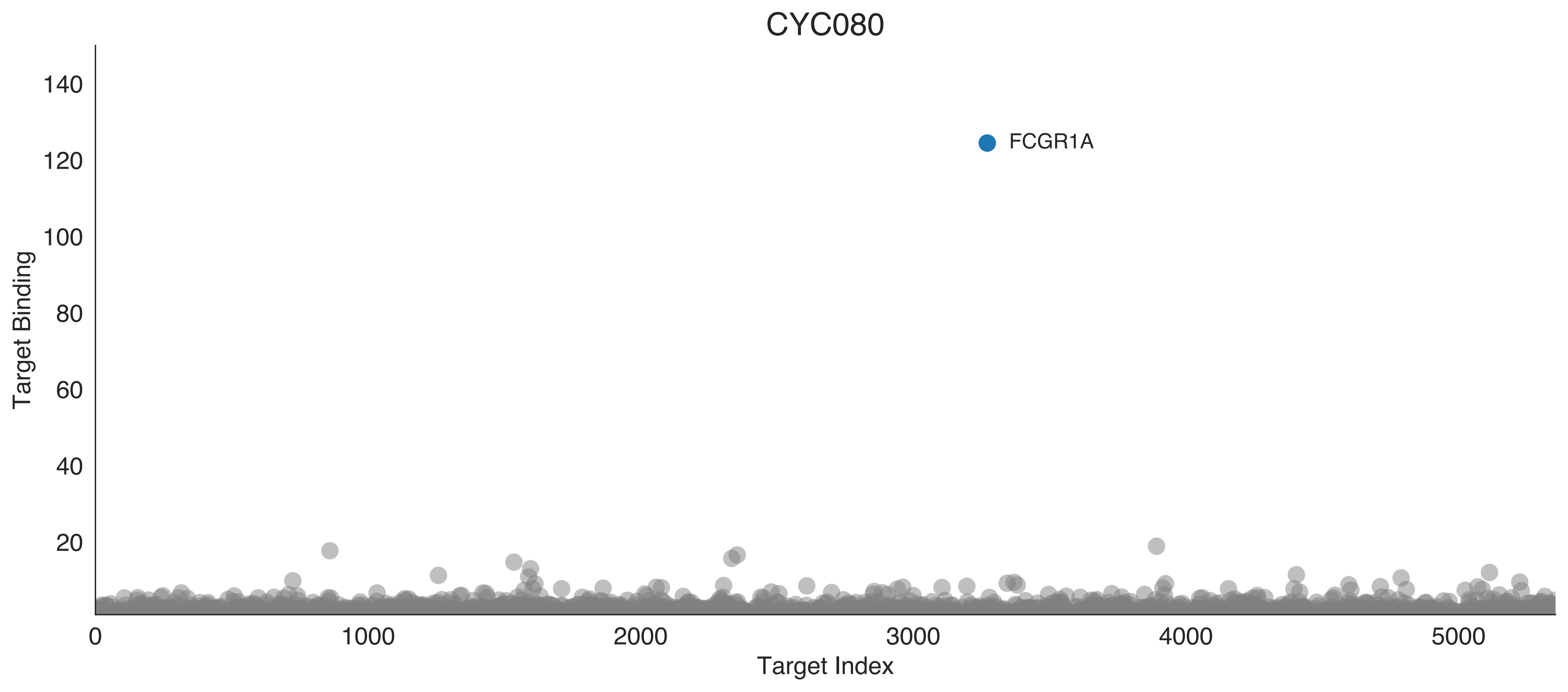

CYC080
FCGR1A
Target Binding
Target Index
0
20
40
60
80
100
120
140
1000
2000
3000
4000
5000

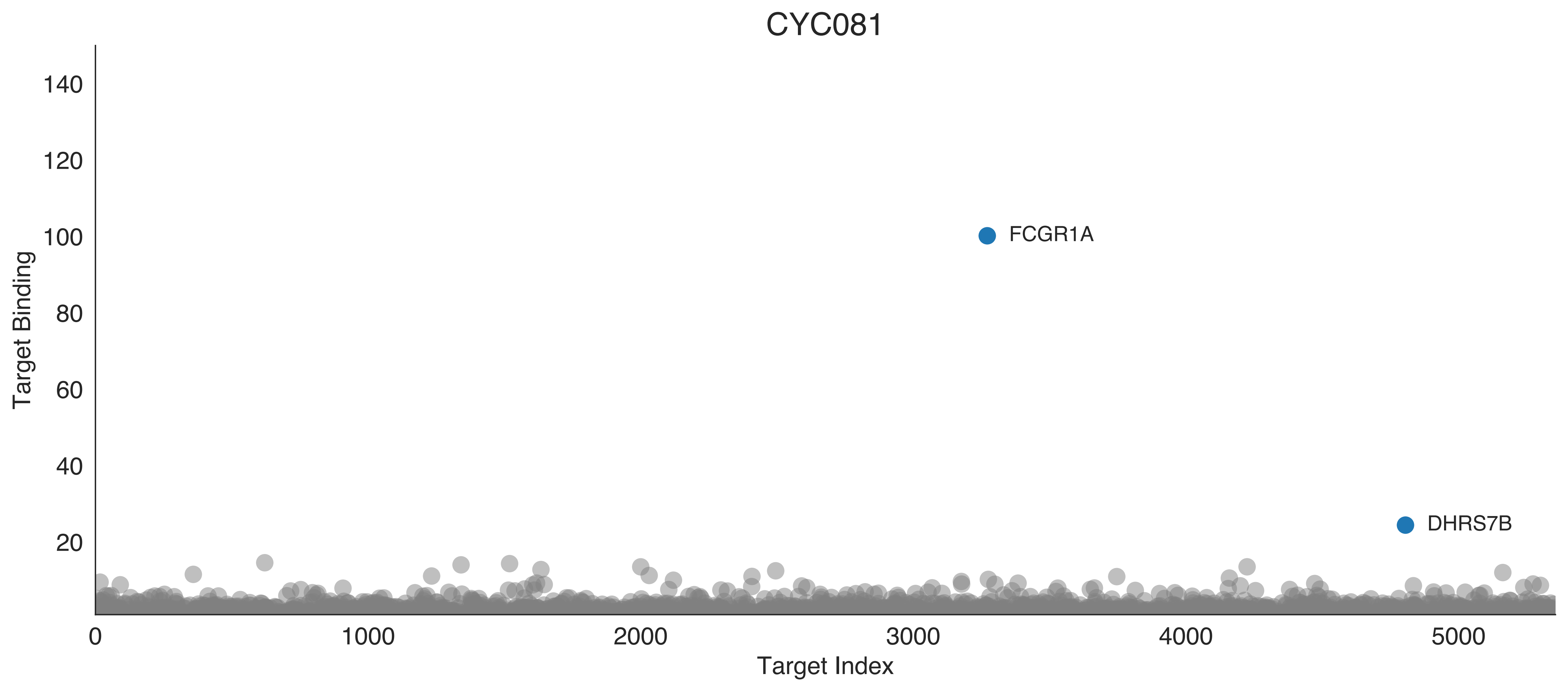

CYC081
Target Binding
Target Index
140
120
100
80
60
40
20
0
1000
2000
3000
4000
5000
FCGR1A
DHRS7B

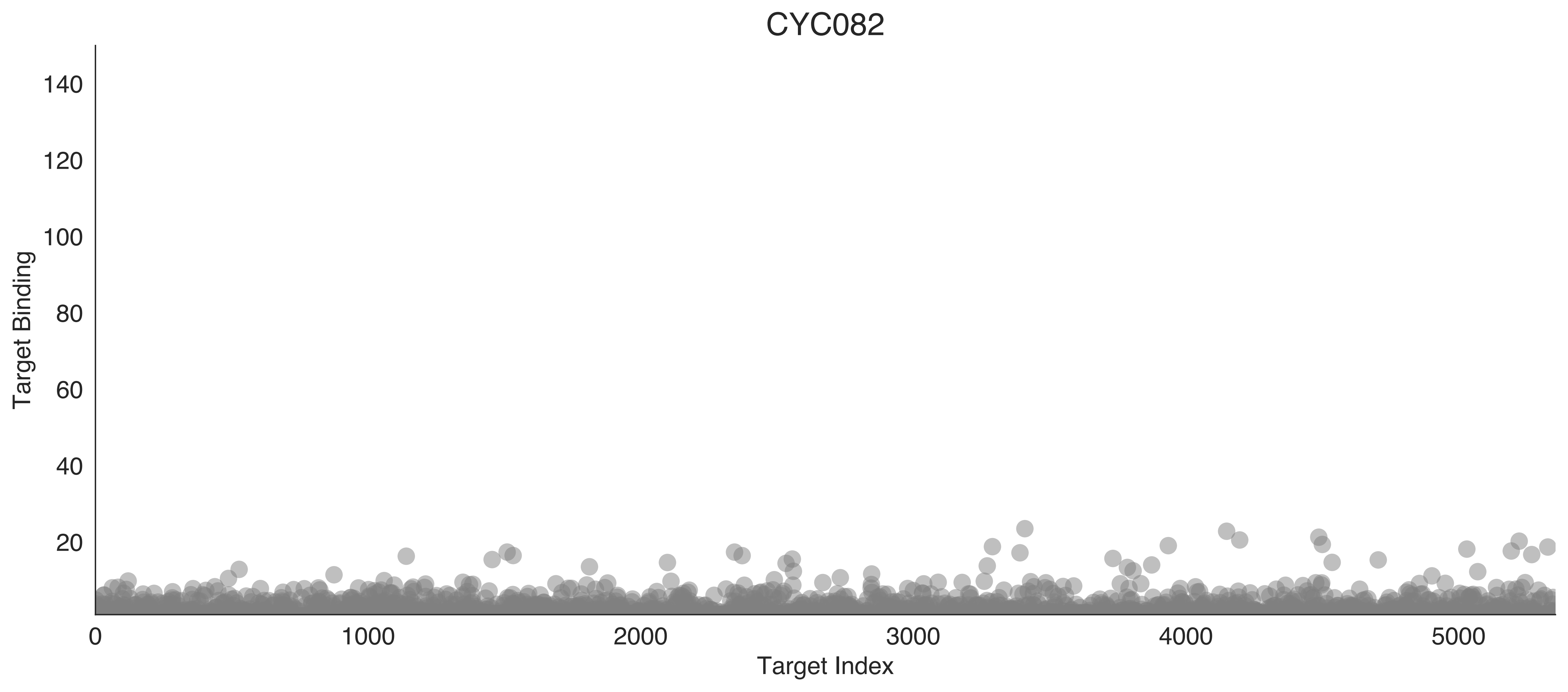

CYC082
Target Binding
Target Index
0
20
40
60
80
100
120
140
0
1000
2000
3000
4000
5000

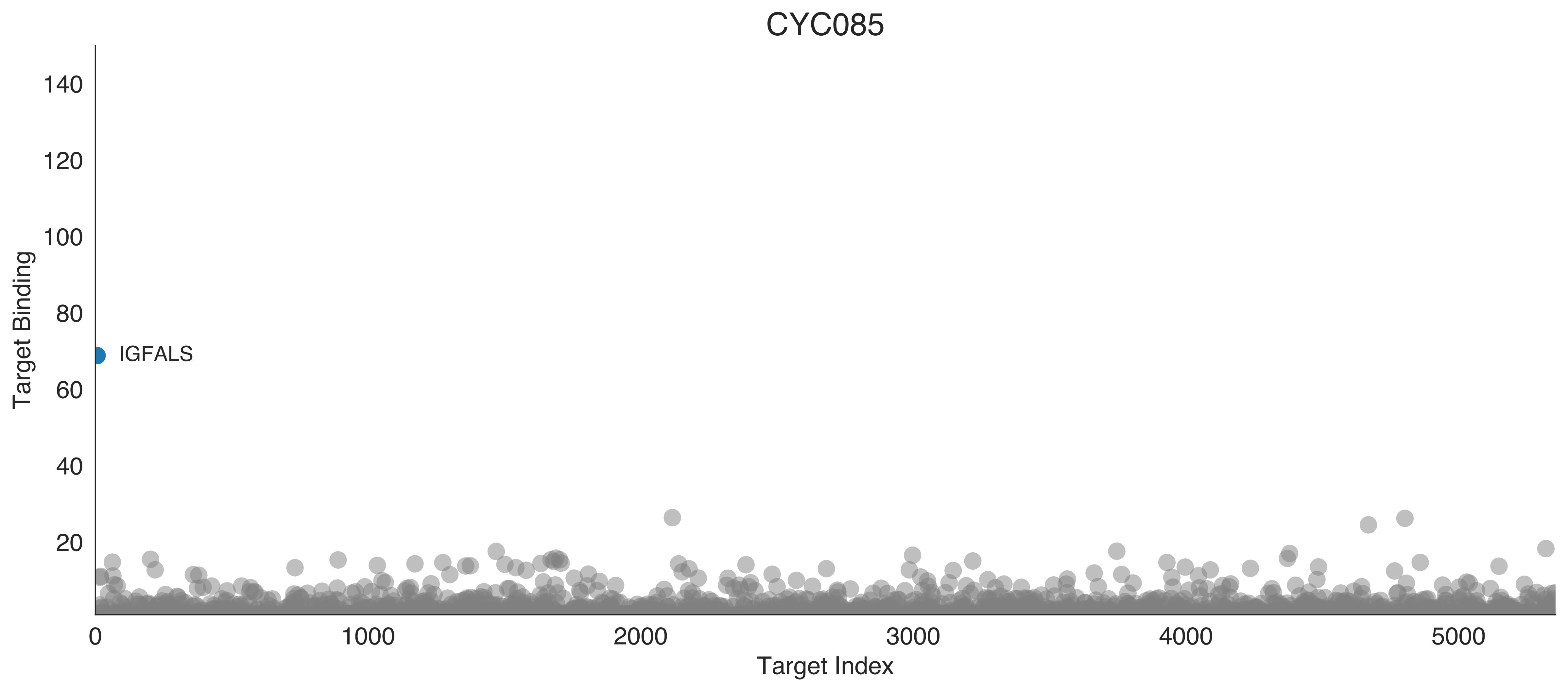

CYC085
Target Binding
Target Index
IGFALS
0
20
40
60
80
100
120
140
1000
2000
3000
4000
5000

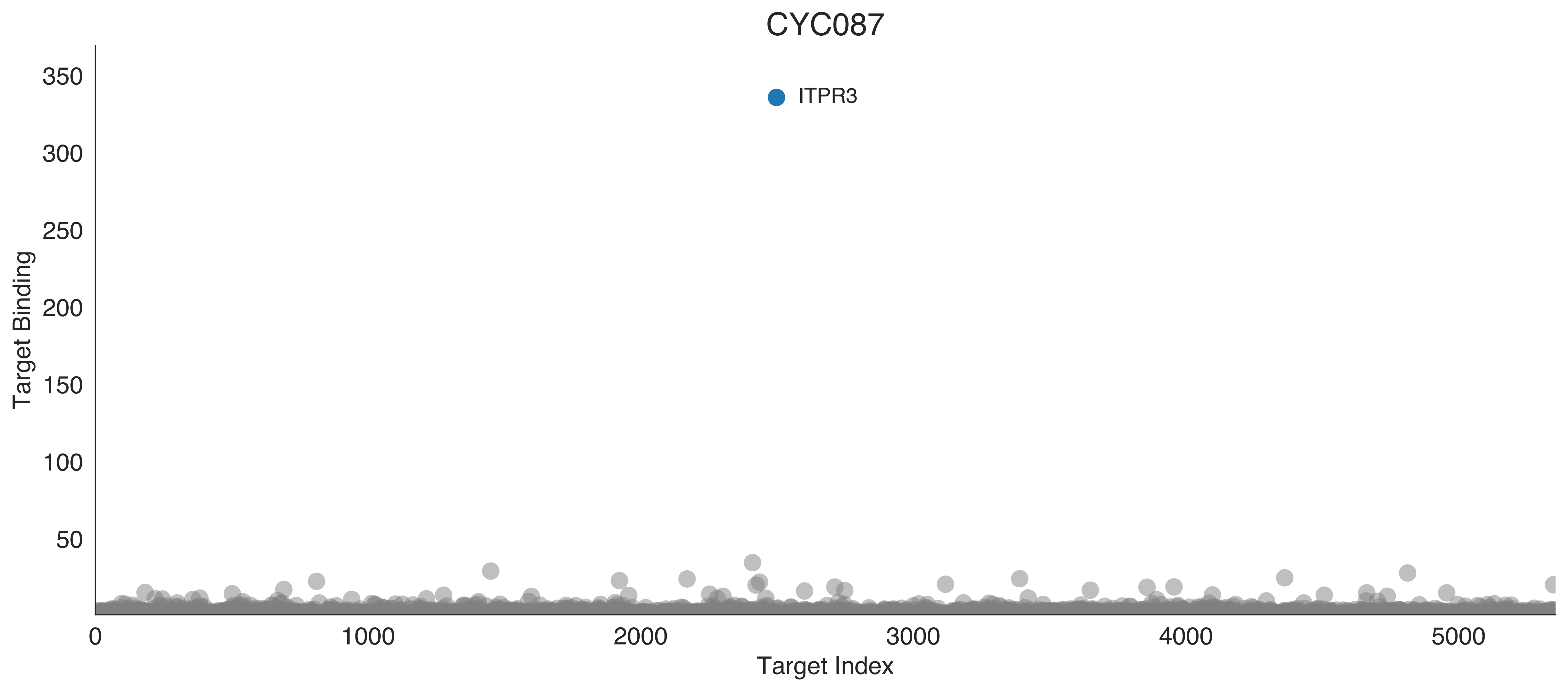

CYC087
ITPR3
Target Binding
350
300
250
200
150
100
50
0
1000
2000
3000
4000
5000
Target Index

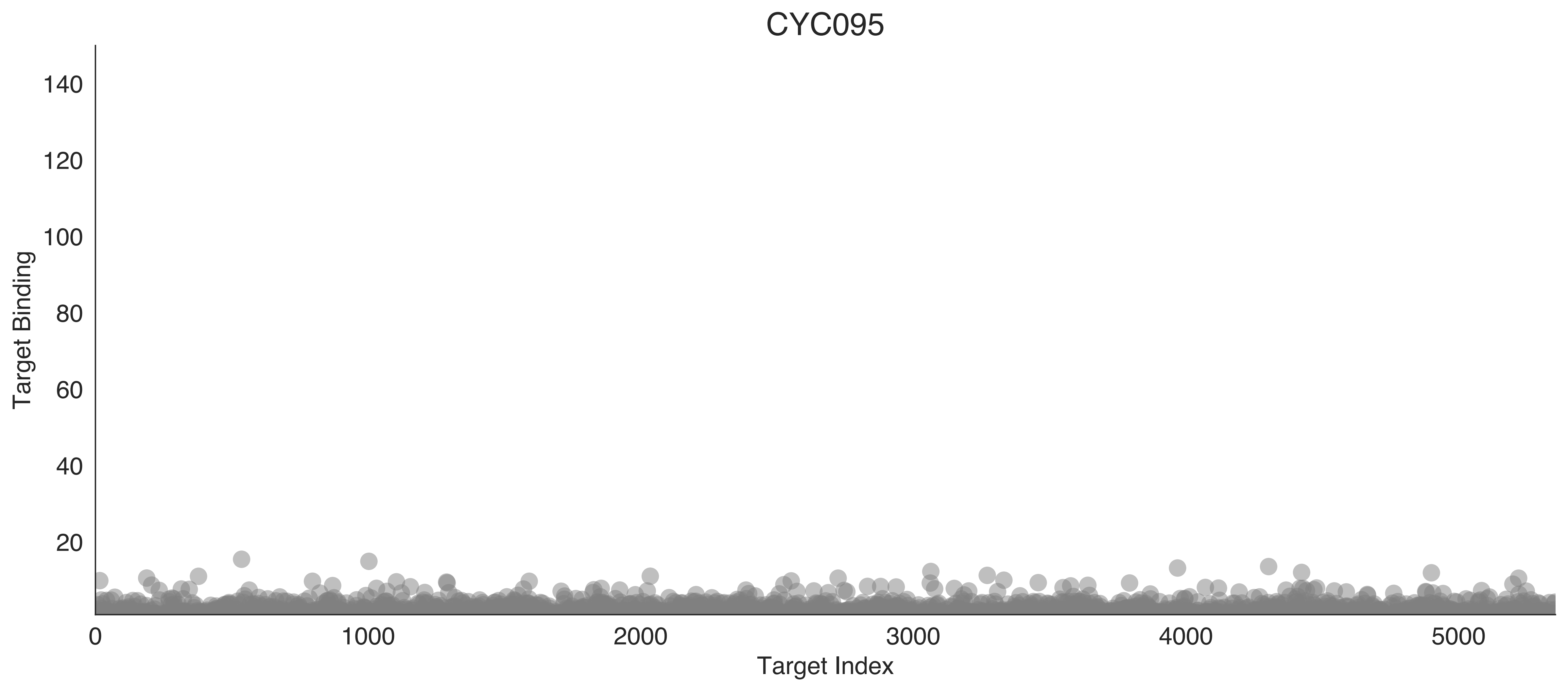
CYC095
Target Binding
140
120
100
80
60
40
20
0
1000
2000
3000
4000
5000
Target Index

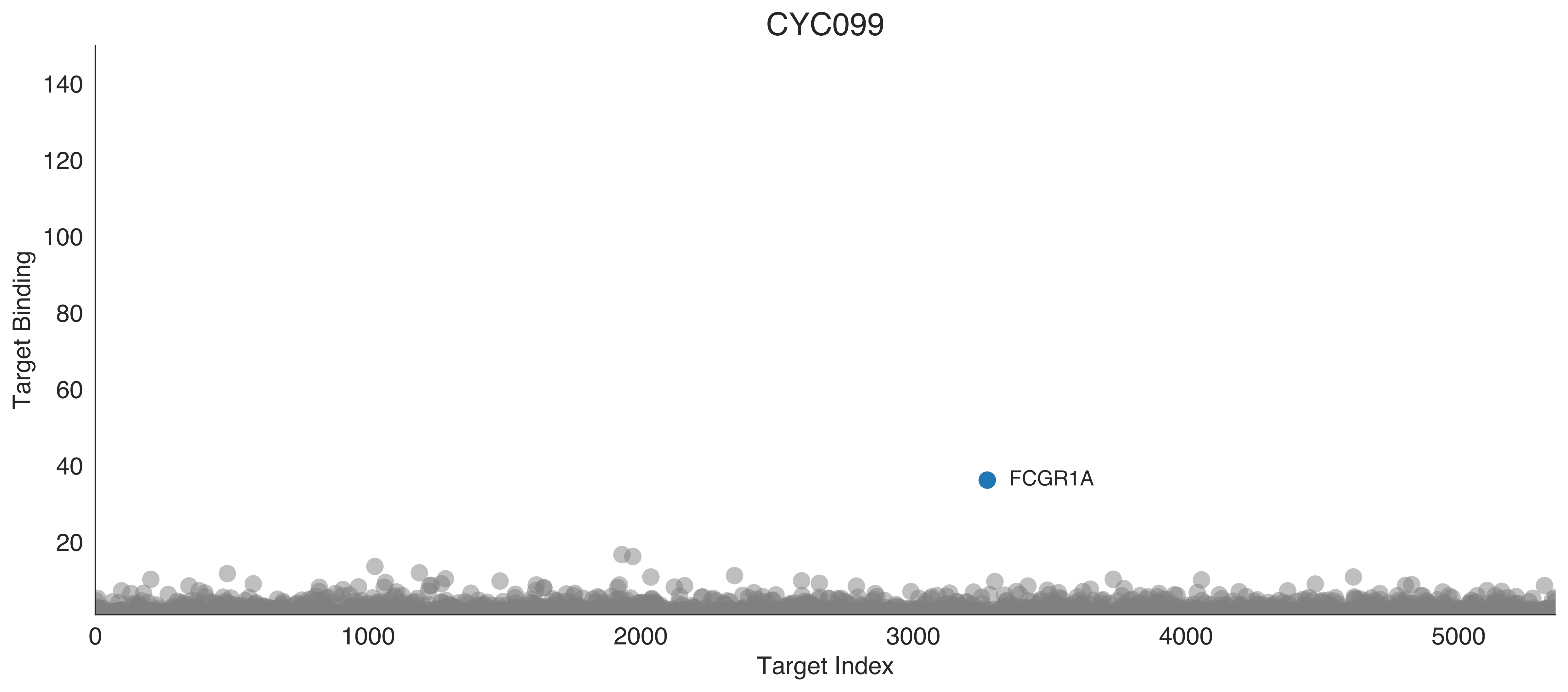

CYC099
Target Binding
Target Index
140
120
100
80
60
40
20
0
1000
2000
3000
4000
5000
FCGR1A

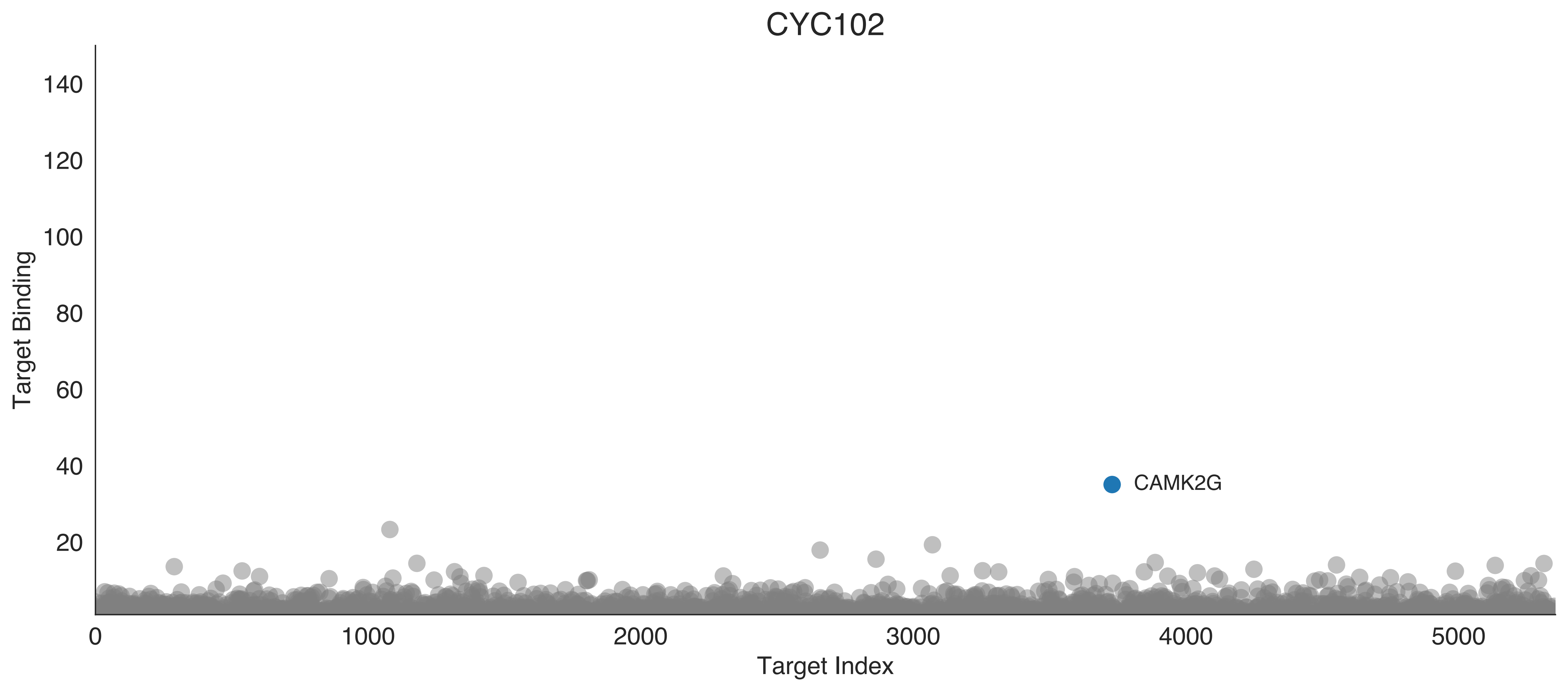
CYC102
Target Binding
140
120
100
80
60
40
20
CAMK2G
0
1000
2000
3000
4000
5000
Target Index

CYC103

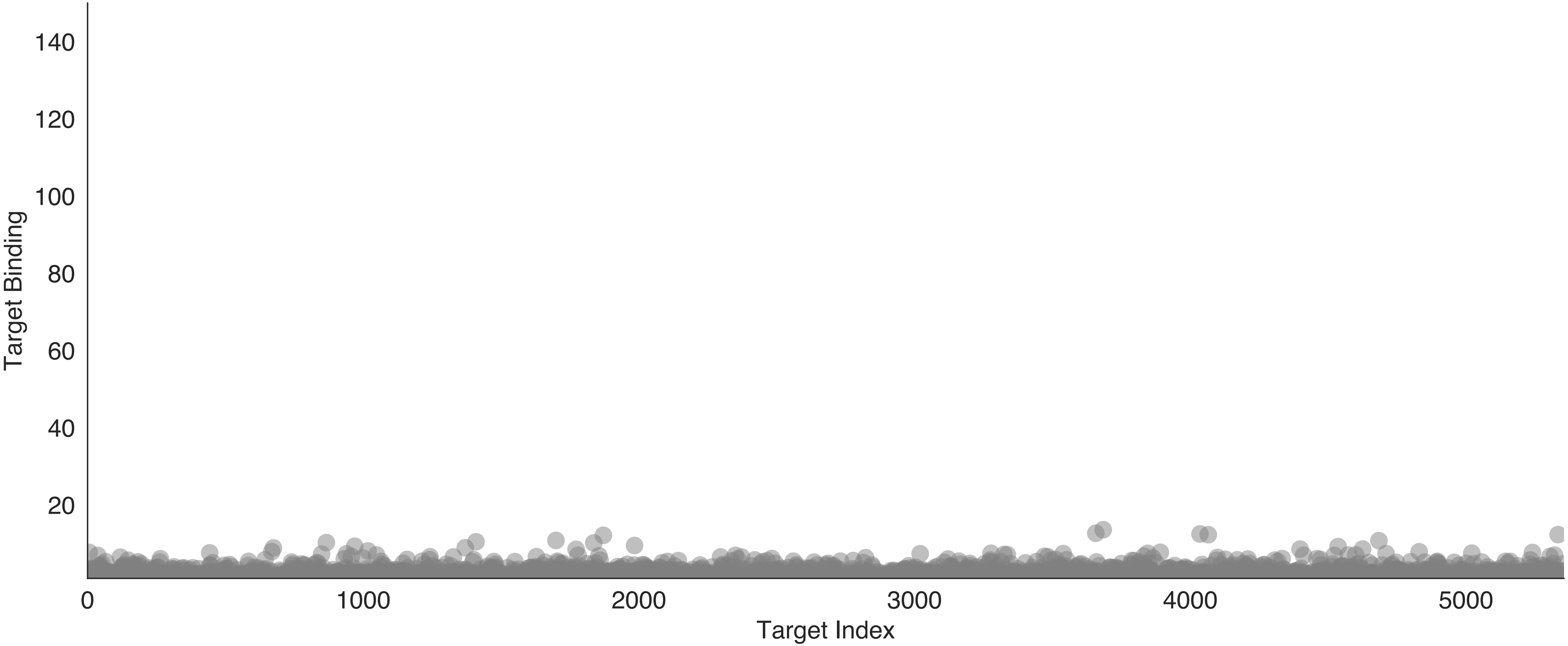

Target Binding
140
120
100
80
60
40
20
0
1000
2000
3000
4000
5000
Target Index

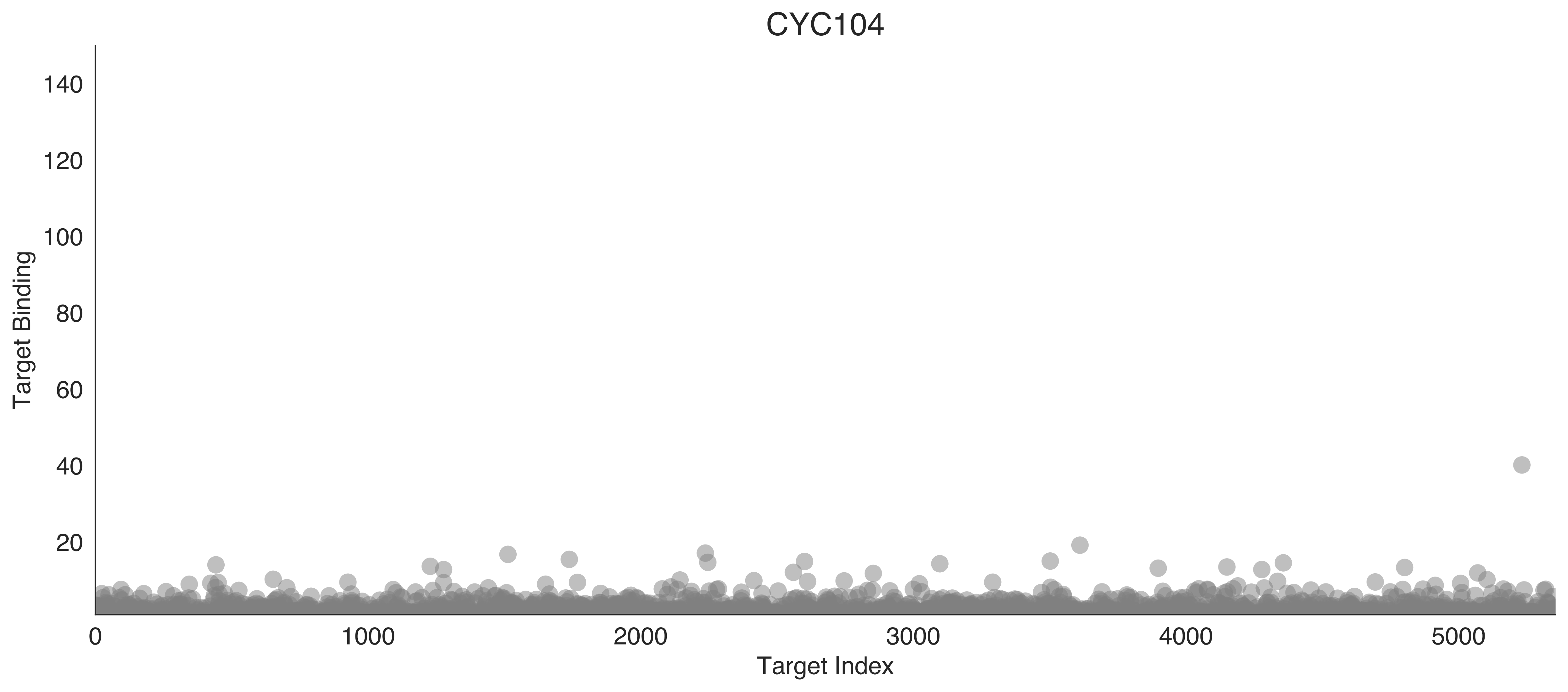
CYC104
Target Binding
Target Index
0
20
40
60
80
100
120
140
1000
2000
3000
4000
5000

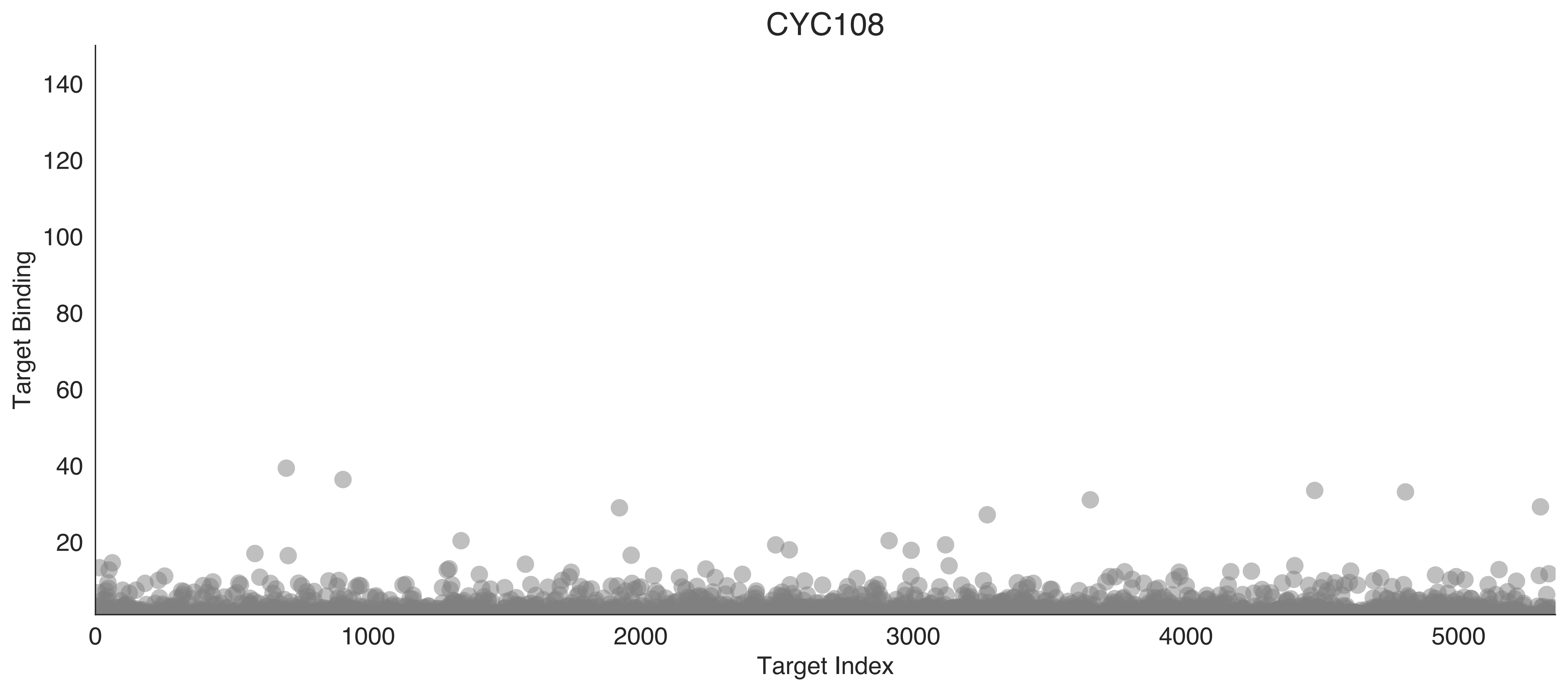
CYC108
Target Binding
140
120
100
80
60
40
20
0
1000
2000
3000
4000
5000
Target Index

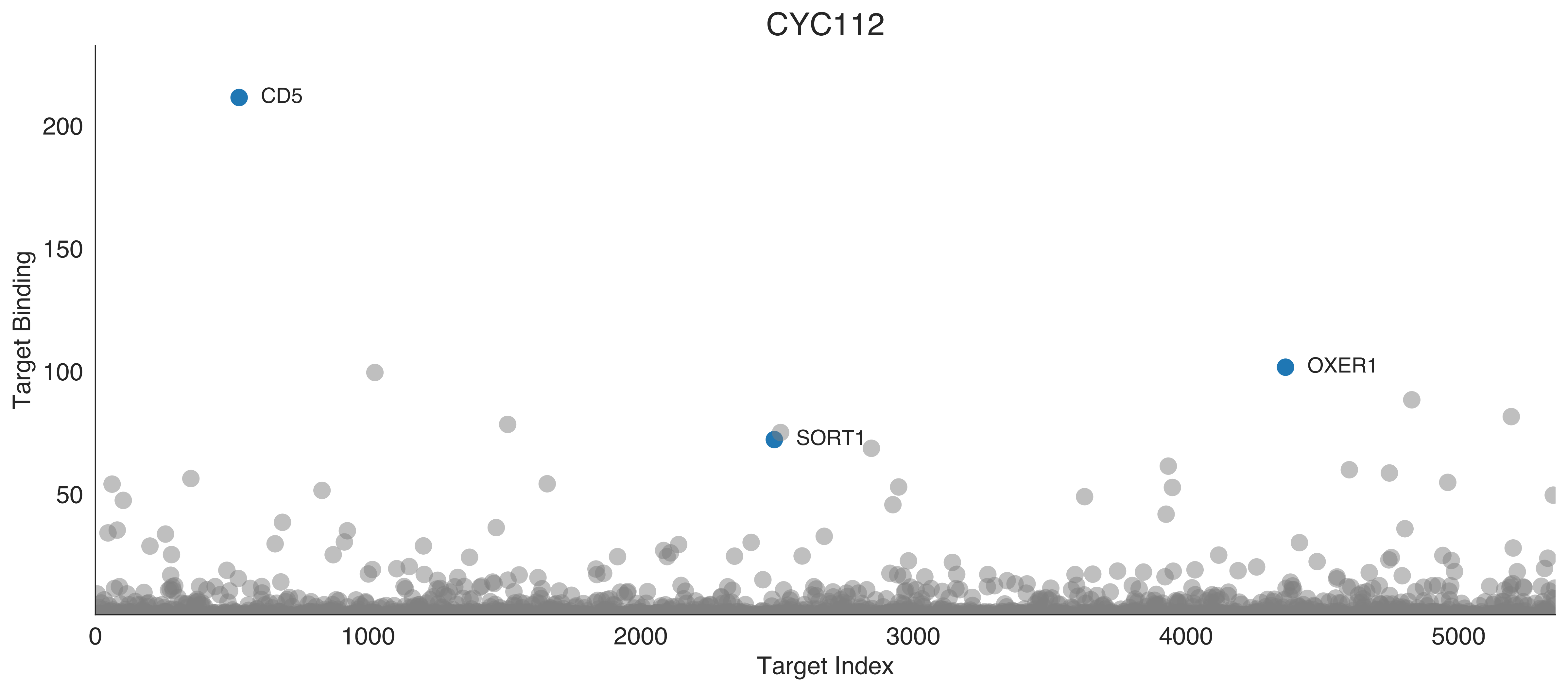

CYC112
CD5
SORT1
OXER1
Target Binding
Target Index
200
150
100
50
0
1000
2000
3000
4000
5000

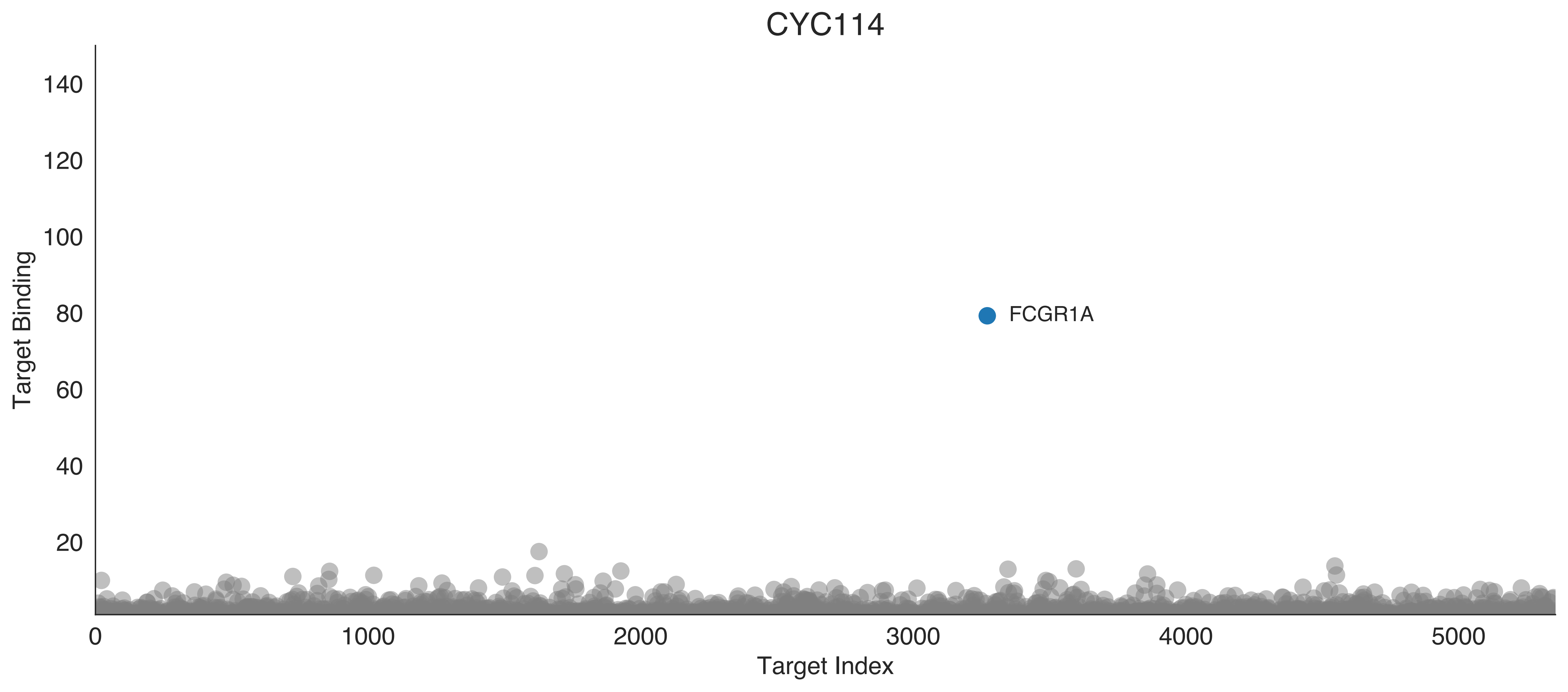
CYC114
Target Binding
Target Index
FCGR1A
140
120
100
80
60
40
20
0
1000
2000
3000
4000
5000

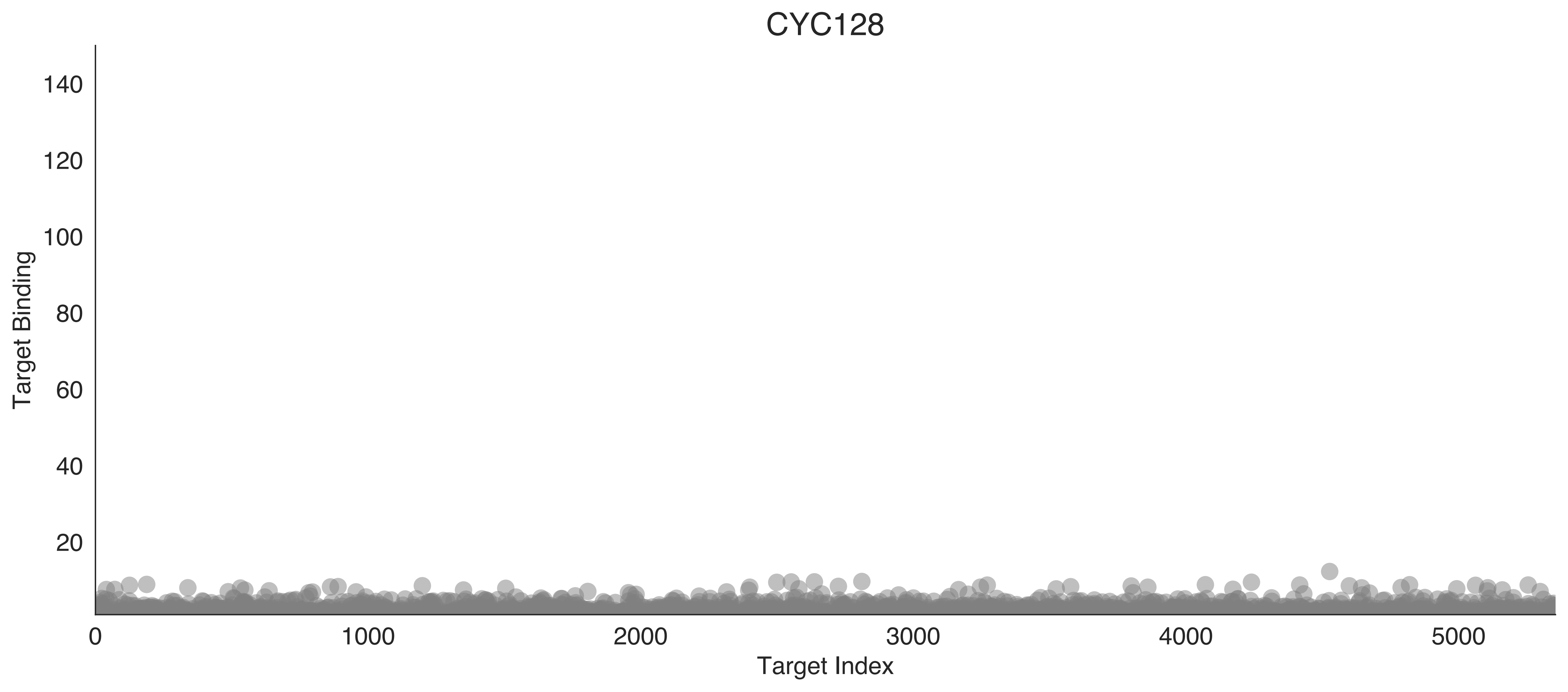
CYC128
Target Binding
Target Index
140
120
100
80
60
40
20
0
1000
2000
3000
4000
5000

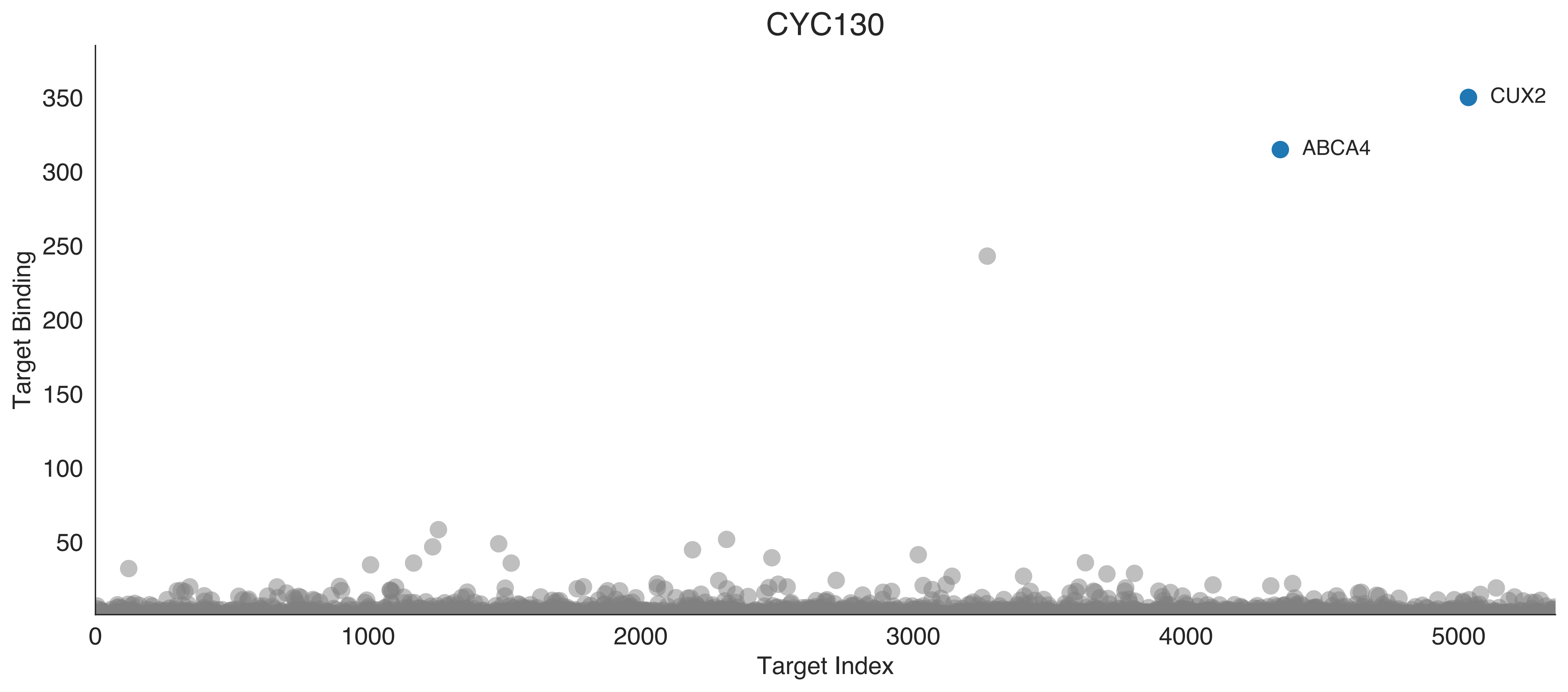

CYC130
CUX2
ABCA4
Target Binding
Target Index
0
50
100
150
200
250
300
350
1000
2000
3000
4000
5000

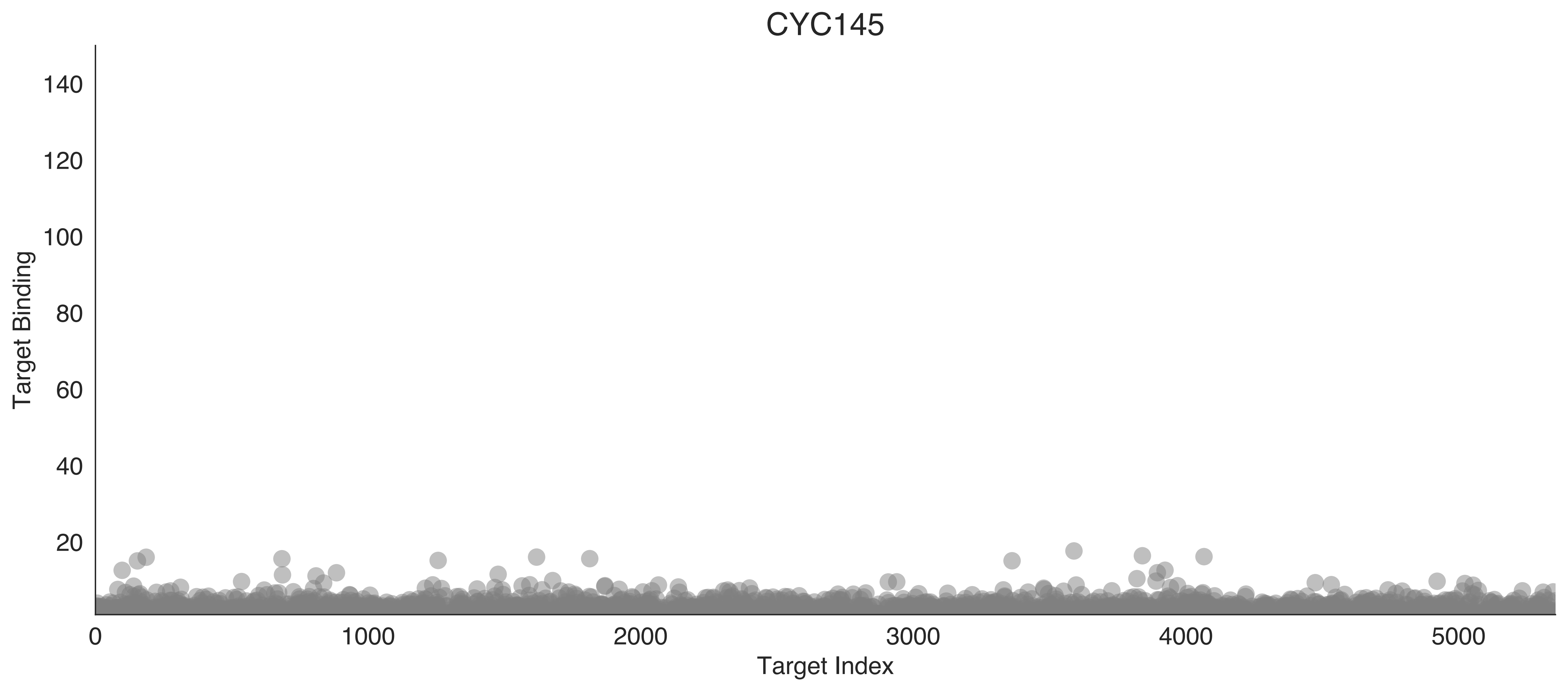

CYC145
Target Binding
Target Index
0
20
40
60
80
100
120
140
1000
2000
3000
4000
5000

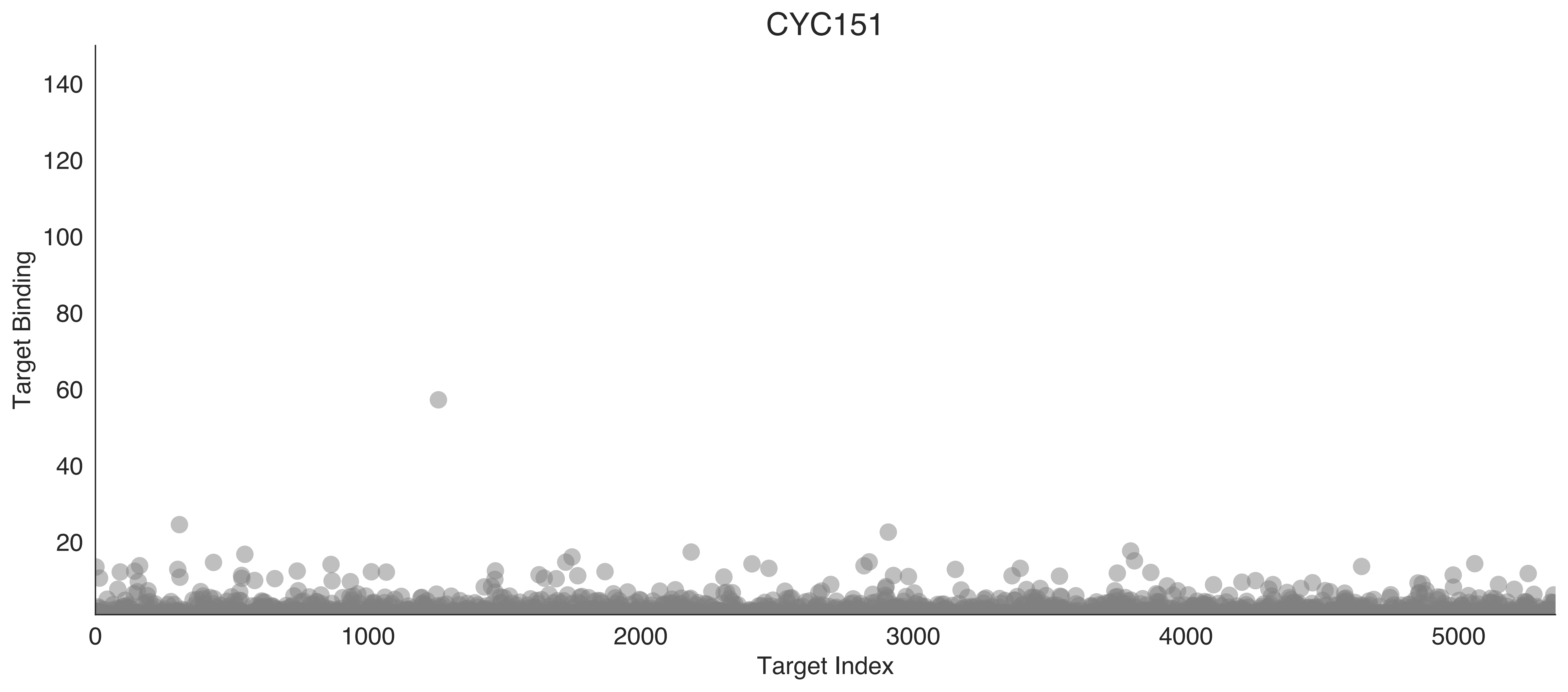

CYC151
Target Binding
140
120
100
80
60
40
20
0
1000
2000
3000
4000
5000
Target Index

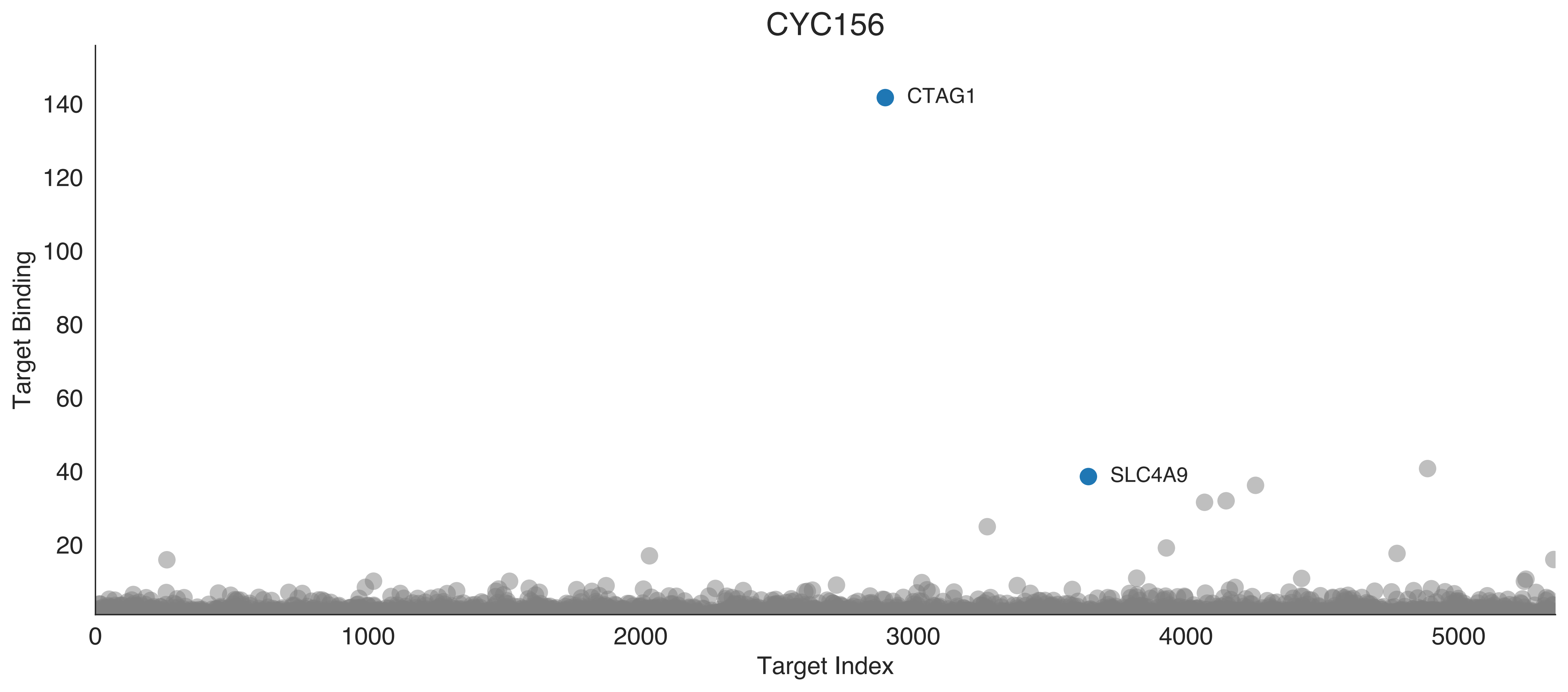
CYC156
Target Binding
Target Index
0
20
40
60
80
100
120
140
1000
2000
3000
4000
5000
CTAG1
SLC4A9

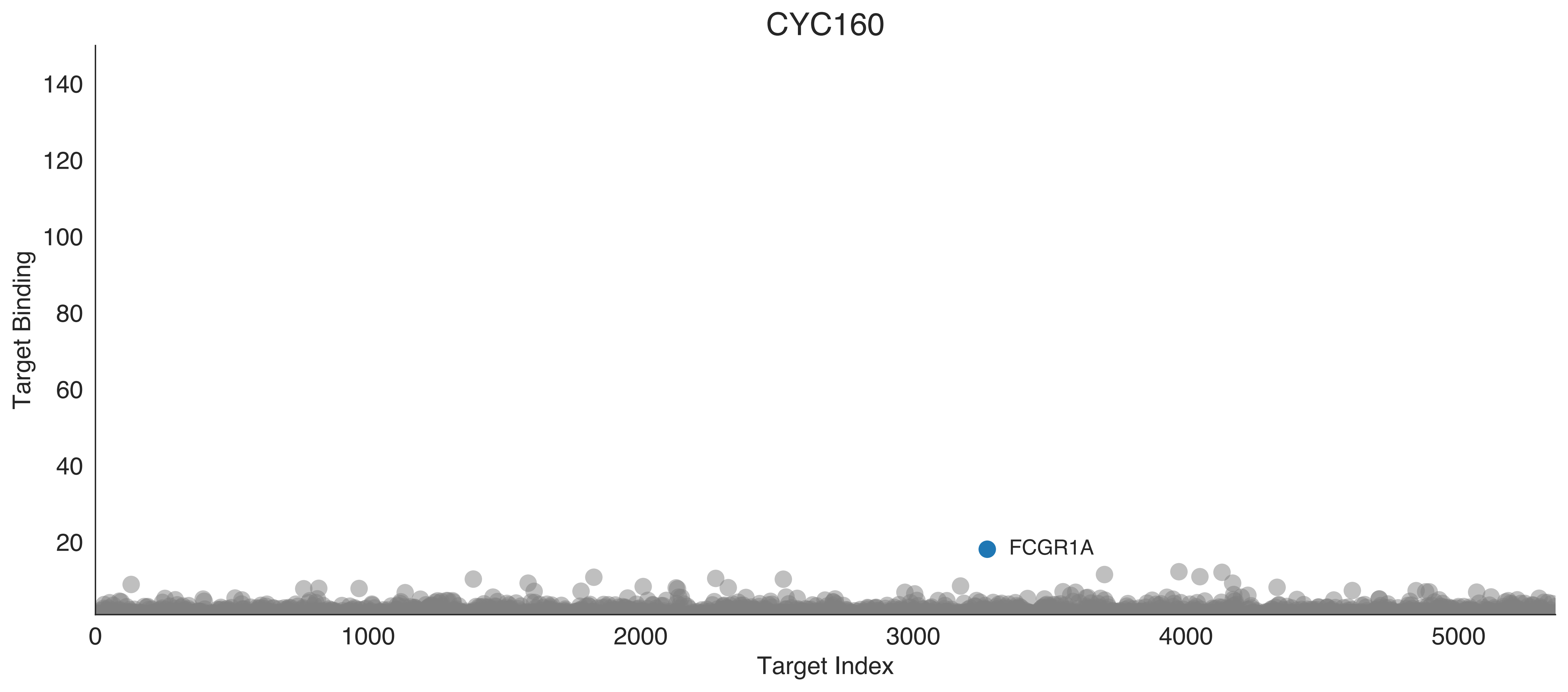
CYC160
Target Binding
140
120
100
80
60
40
20
FCGR1A
0
1000
2000
3000
4000
5000
Target Index

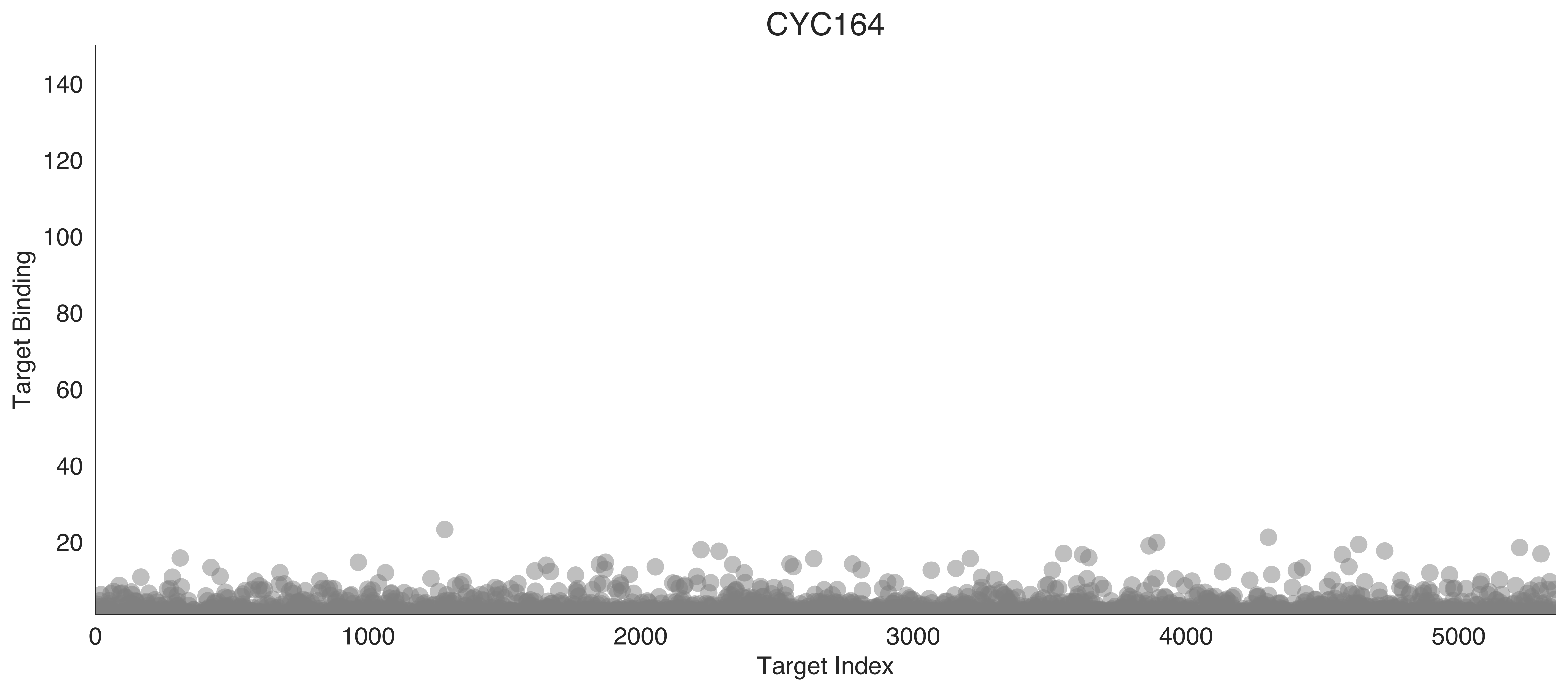
CYC164
Target Binding
Target Index
0
20
40
60
80
100
120
140
0
1000
2000
3000
4000
5000

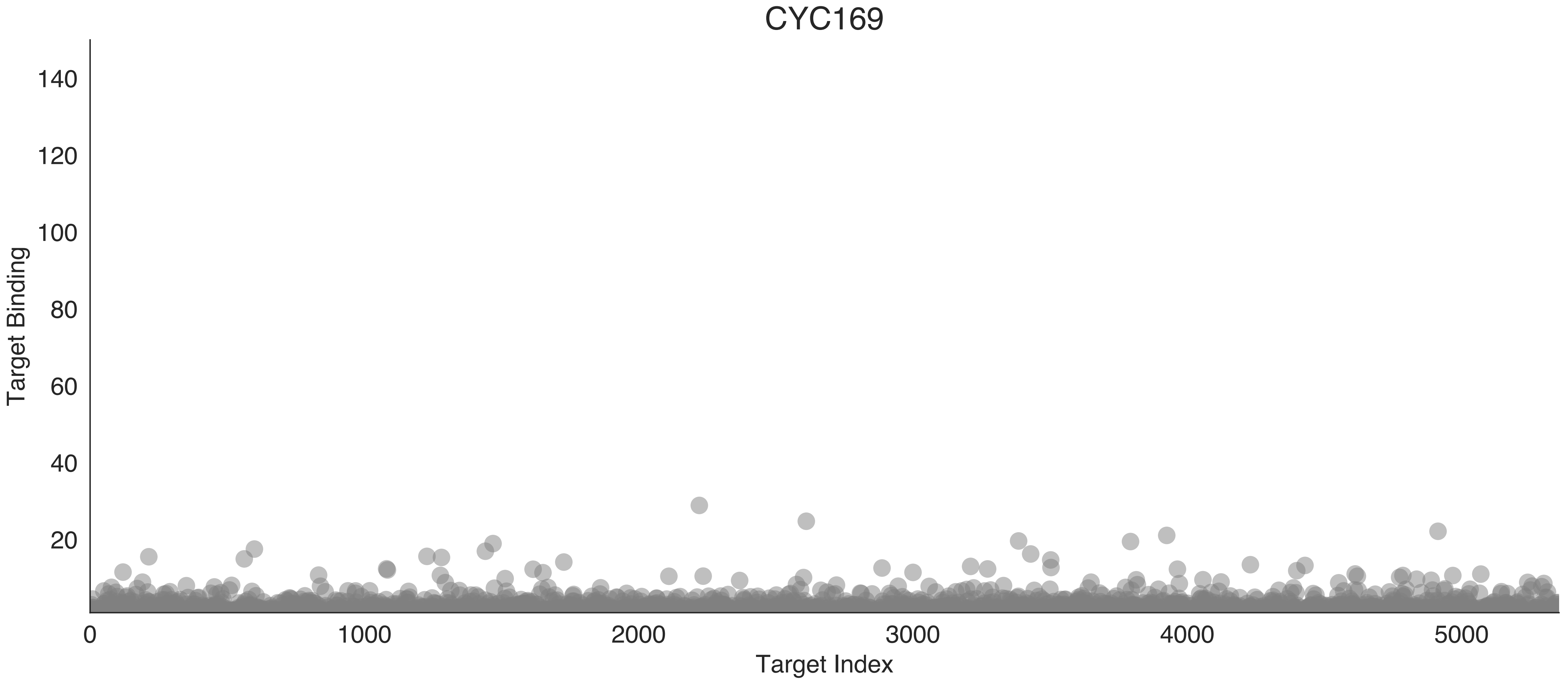

CYC169
Target Binding
140
120
100
80
60
40
20
0
1000
2000
3000
4000
5000
Target Index

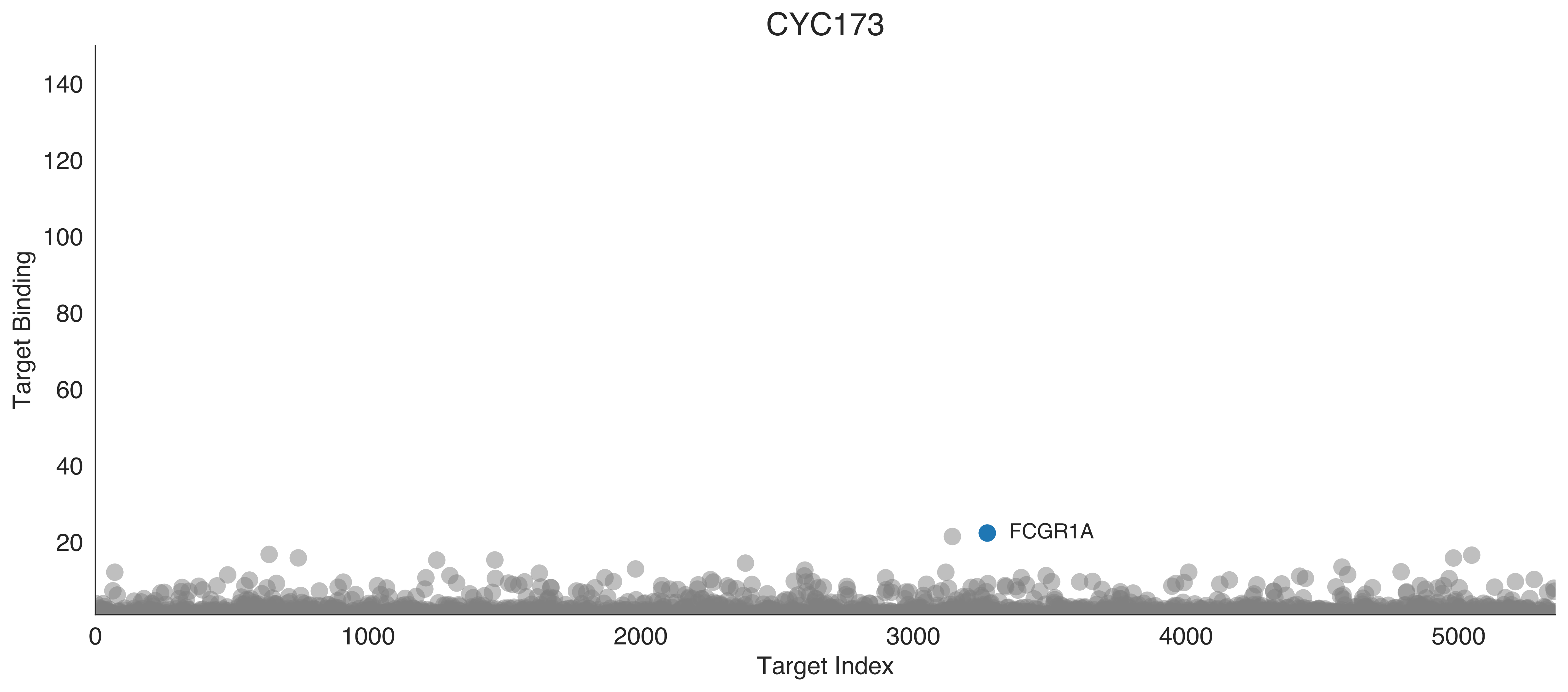
CYC173
Target Binding
Target Index
FCGR1A
0
20
40
60
80
100
120
140
1000
2000
3000
4000
5000

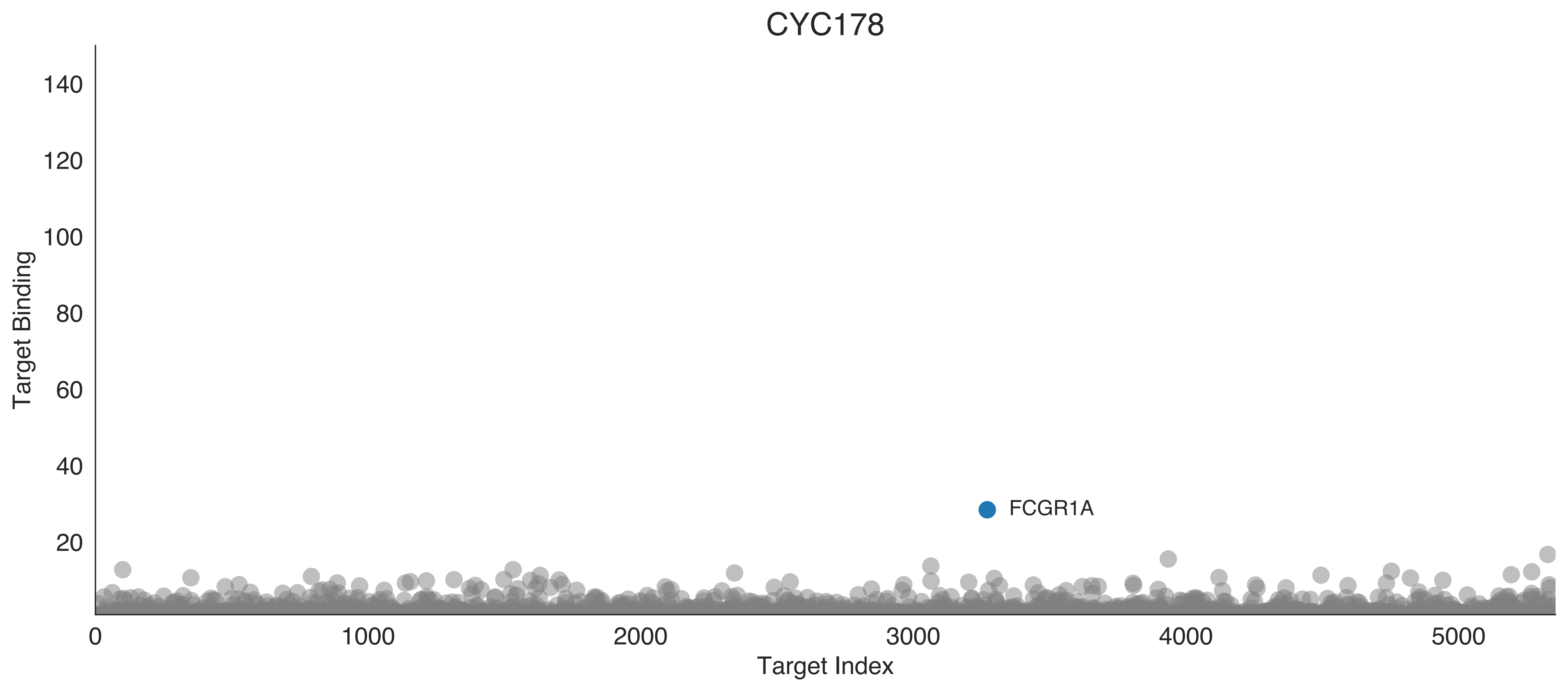

CYC178
Target Binding
140
120
100
80
60
40
20
FCGR1A
0
1000
2000
3000
4000
5000
Target Index

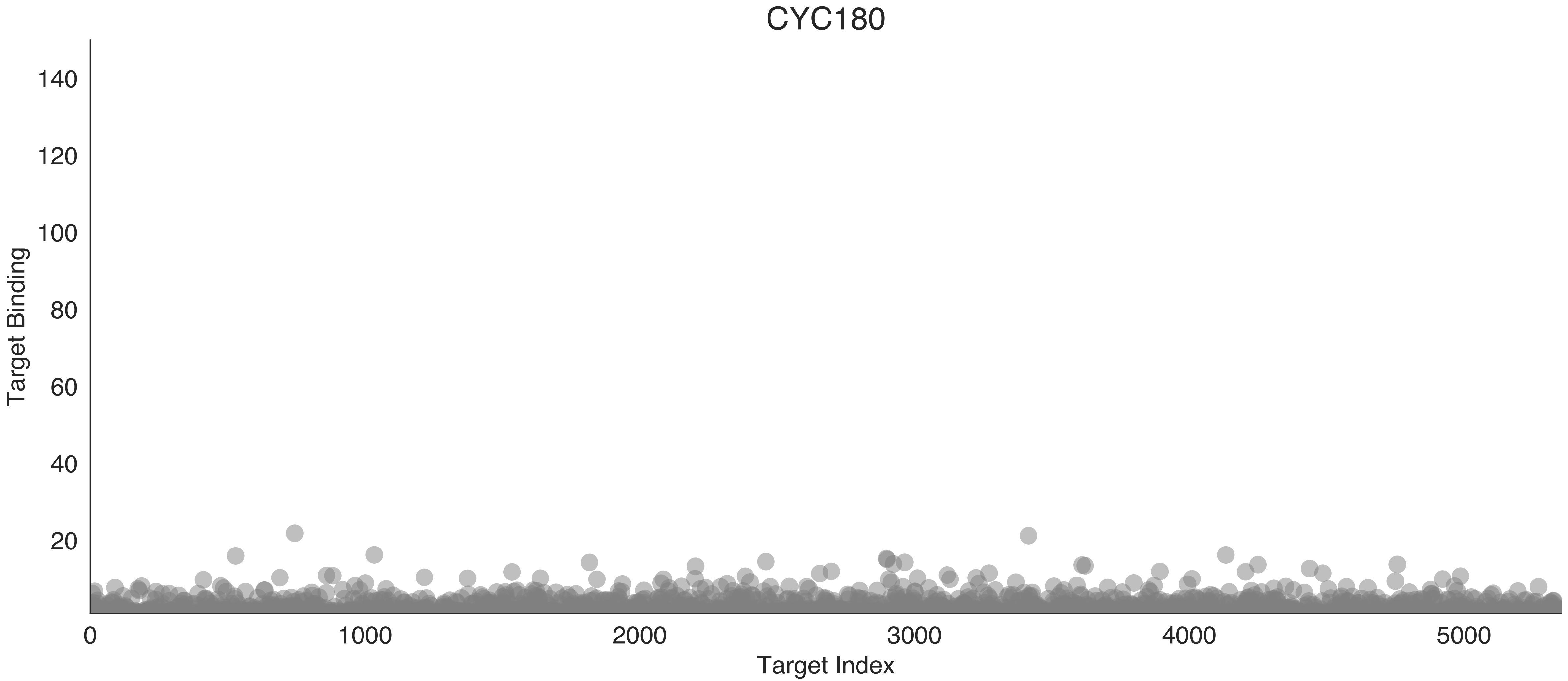

CYC180
Target Binding
140
120
100
80
60
40
20
0
1000
2000
3000
4000
5000
Target Index

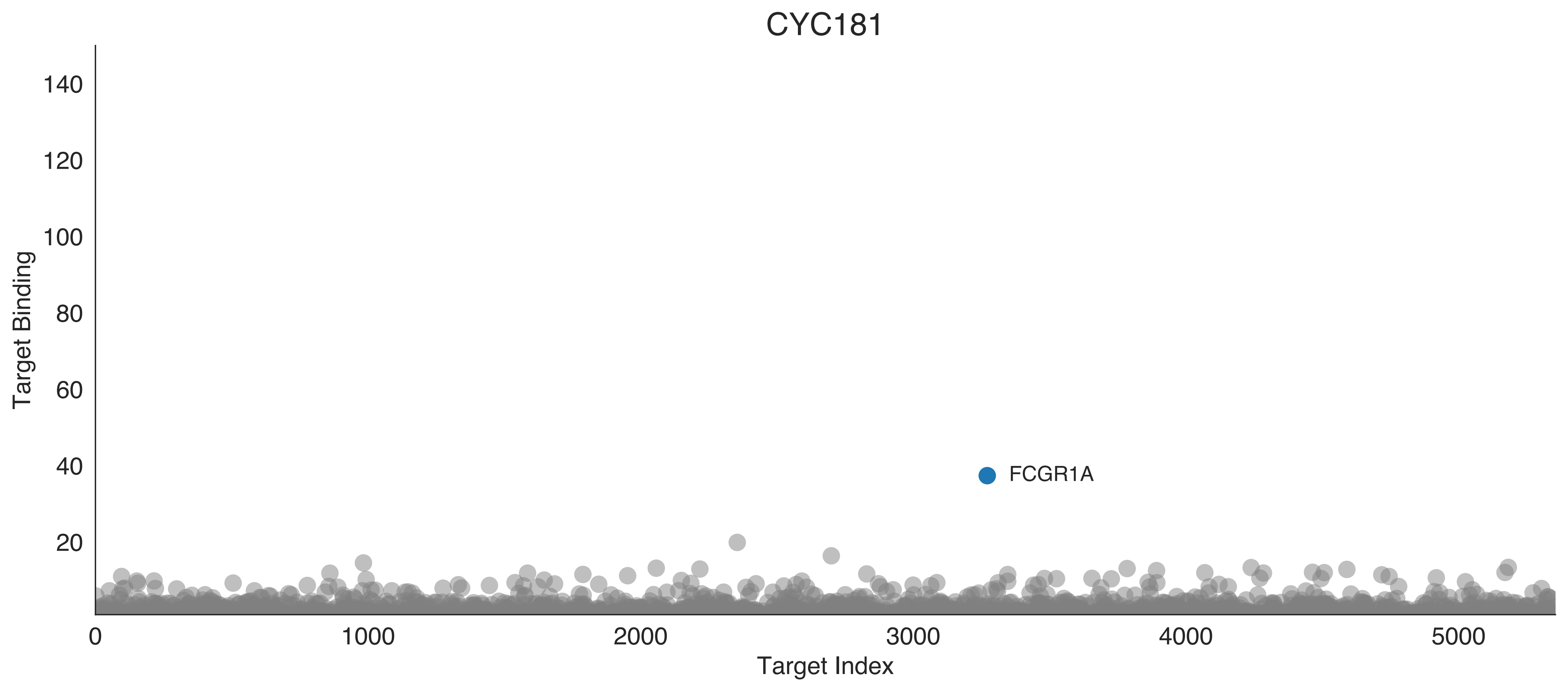

CYC181
140
120
100
80
60
40
20
Target Binding
FCGR1A
0
1000
2000
3000
4000
5000
Target Index

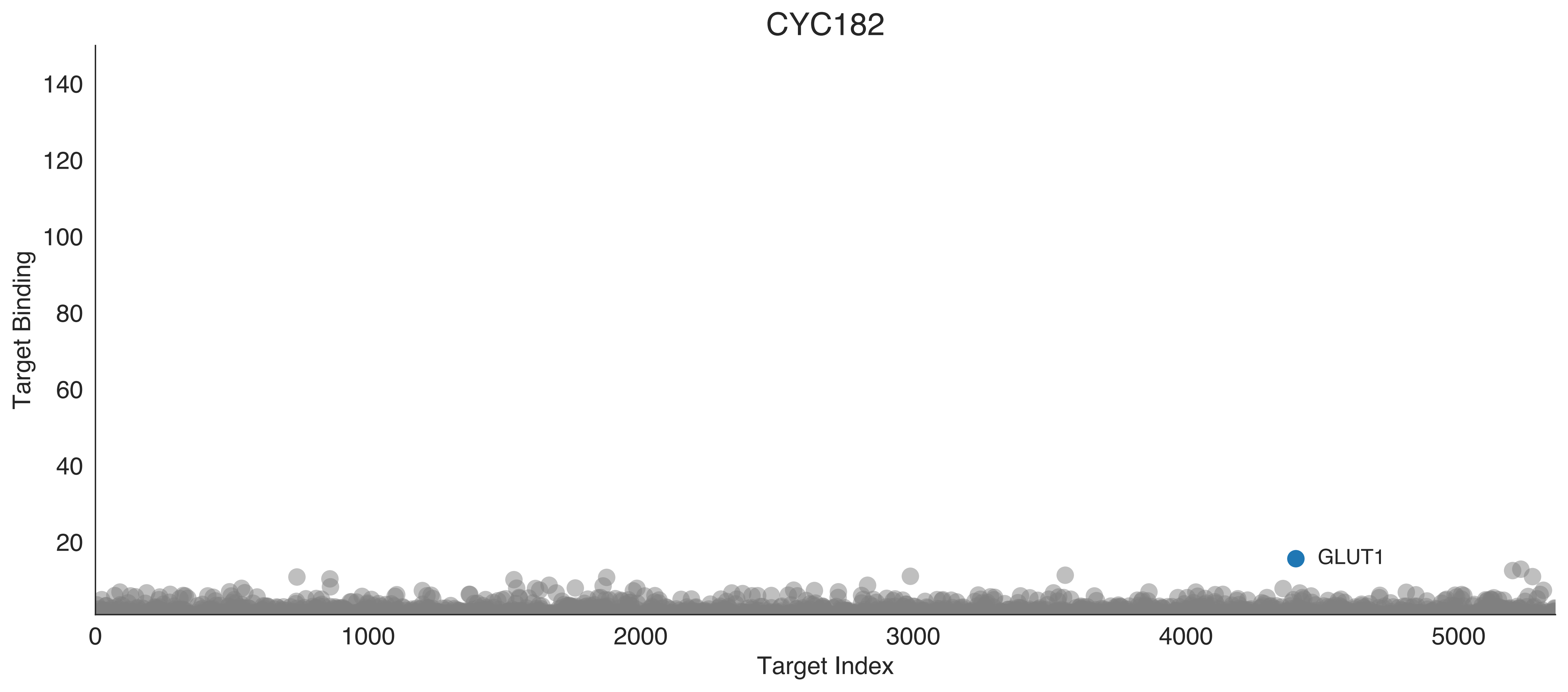

CYC182
Target Binding
Target Index
GLUT1
140
120
100
80
60
40
20
0
1000
2000
3000
4000
5000

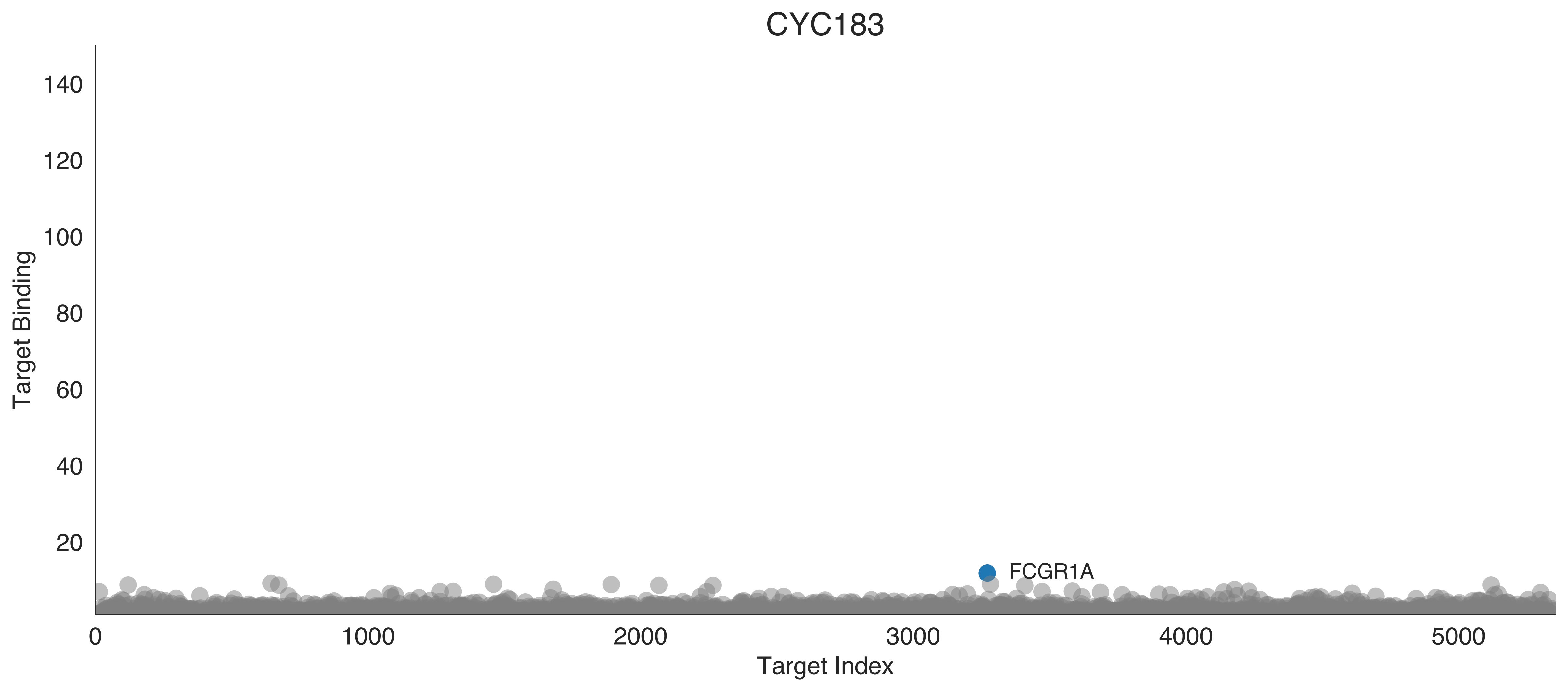

CYC183
Target Binding
Target Index
140
120
100
80
60
40
20
0
1000
2000
3000
4000
5000
FCGR1A

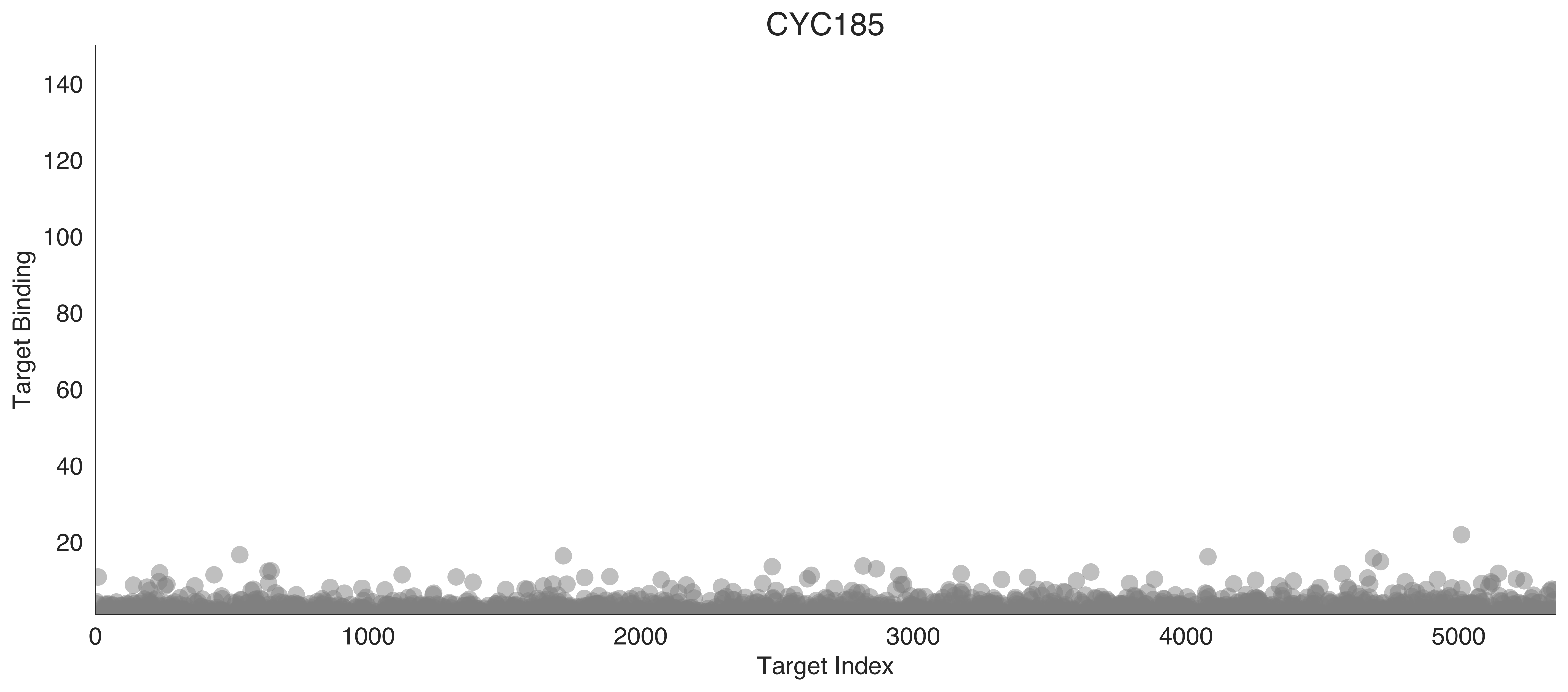

CYC185
Target Binding
140
120
100
80
60
40
20
0
1000
2000
3000
4000
5000
Target Index

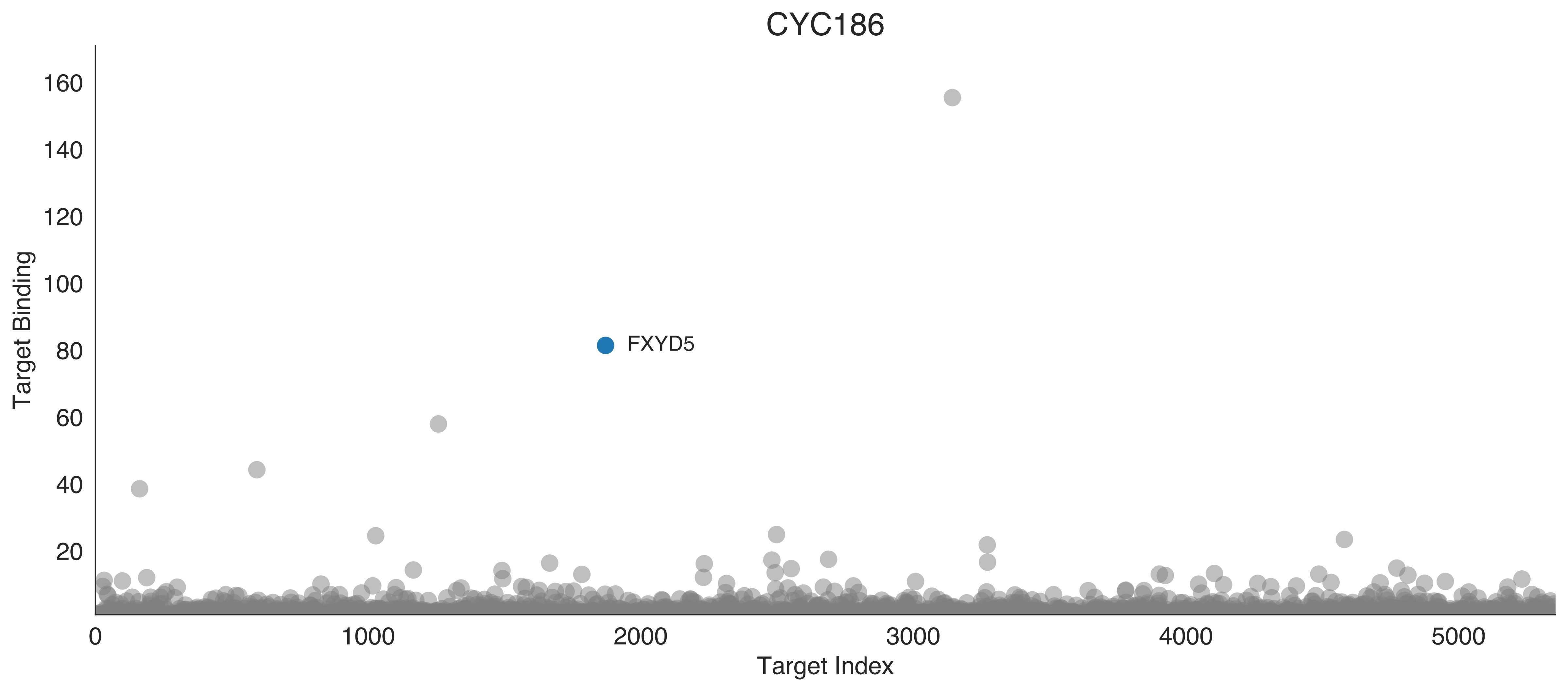
CYC186
Target Binding
160
140
120
100
80
60
40
20
FXYD5
0
1000
2000
3000
4000
5000
Target Index

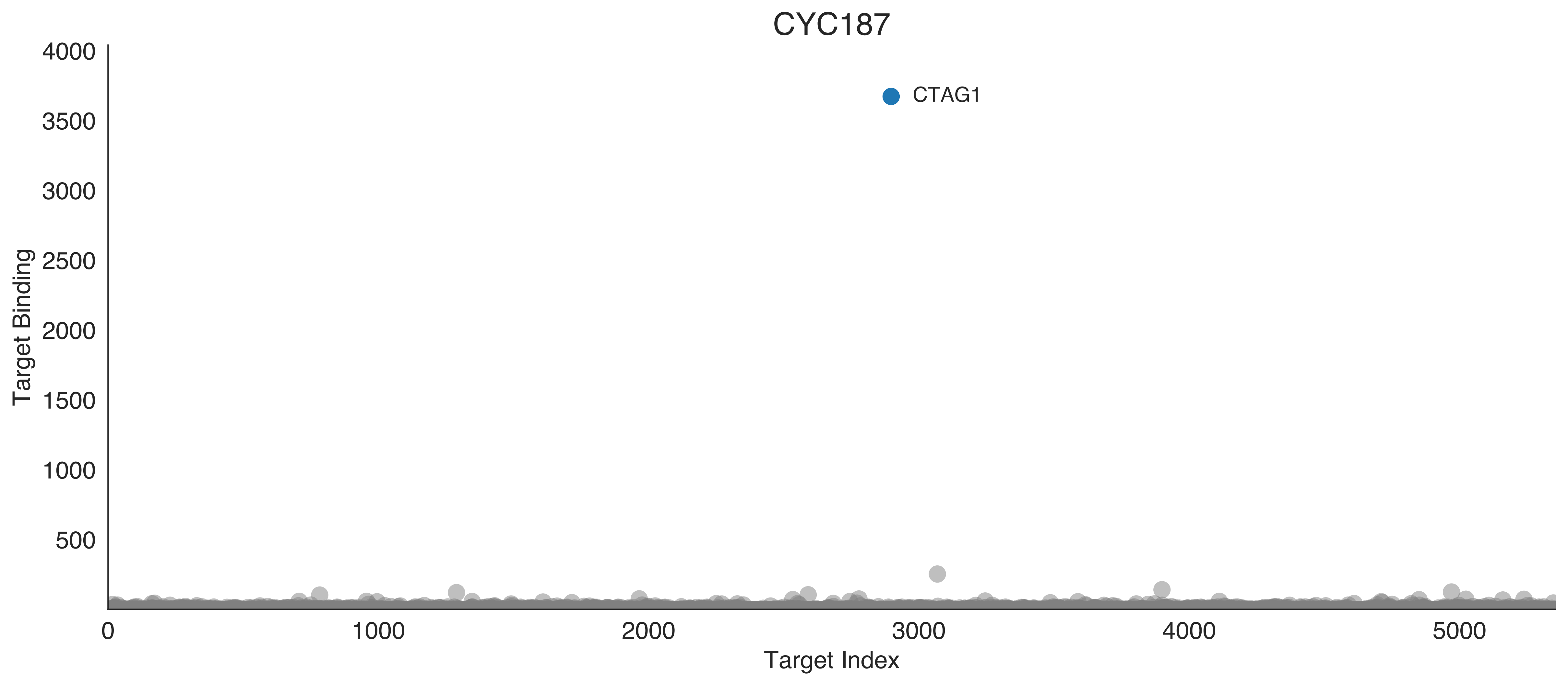

CYC187
CTAG1
Target Binding
4000
3500
3000
2500
2000
1500
1000
500
0
1000
2000
3000
4000
5000
Target Index

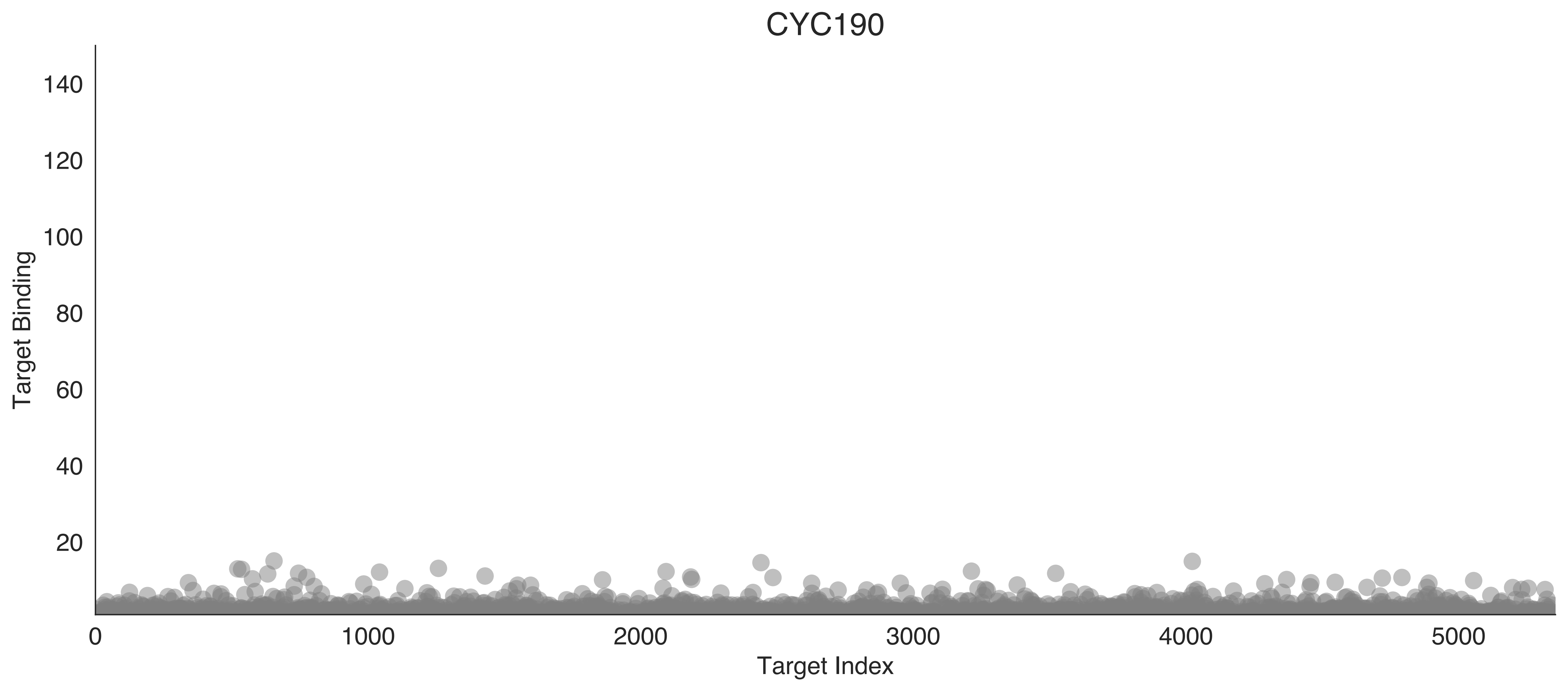

CYC190
Target Binding
Target Index
140
120
100
80
60
40
20
0
1000
2000
3000
4000
5000

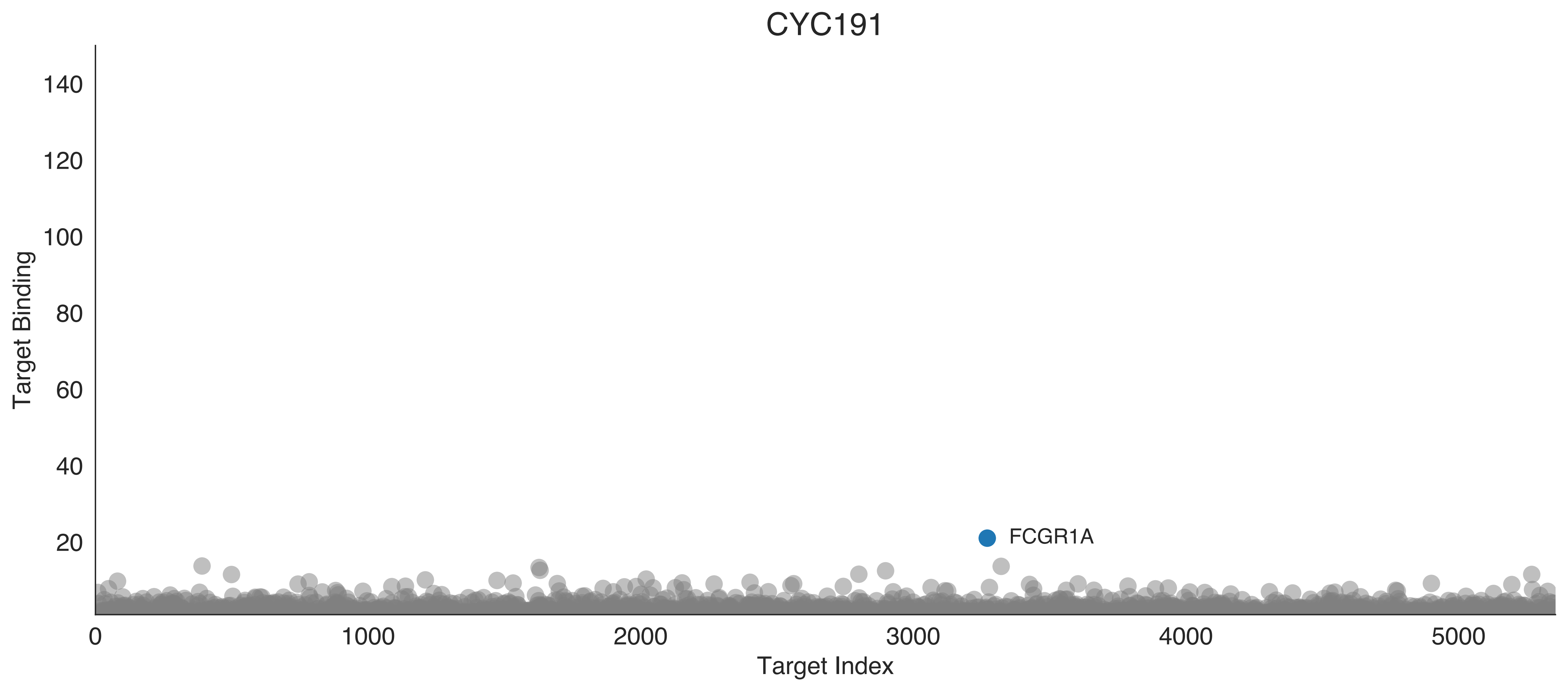

CYC191
Target Binding
140
120
100
80
60
40
20
0
1000
2000
3000
4000
5000
Target Index
FCGR1A

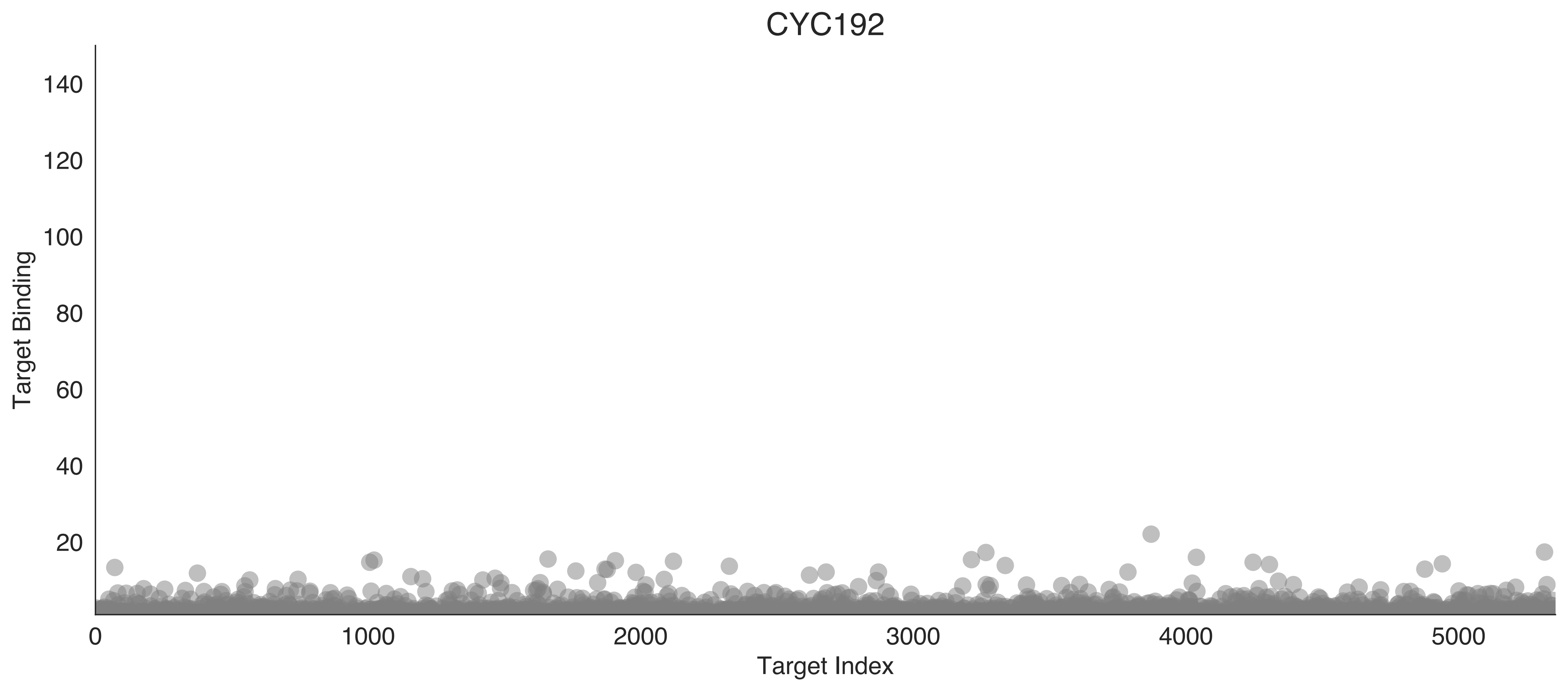
CYC192
Target Binding
140
120
100
80
60
40
20
0
1000
2000
3000
4000
5000
Target Index

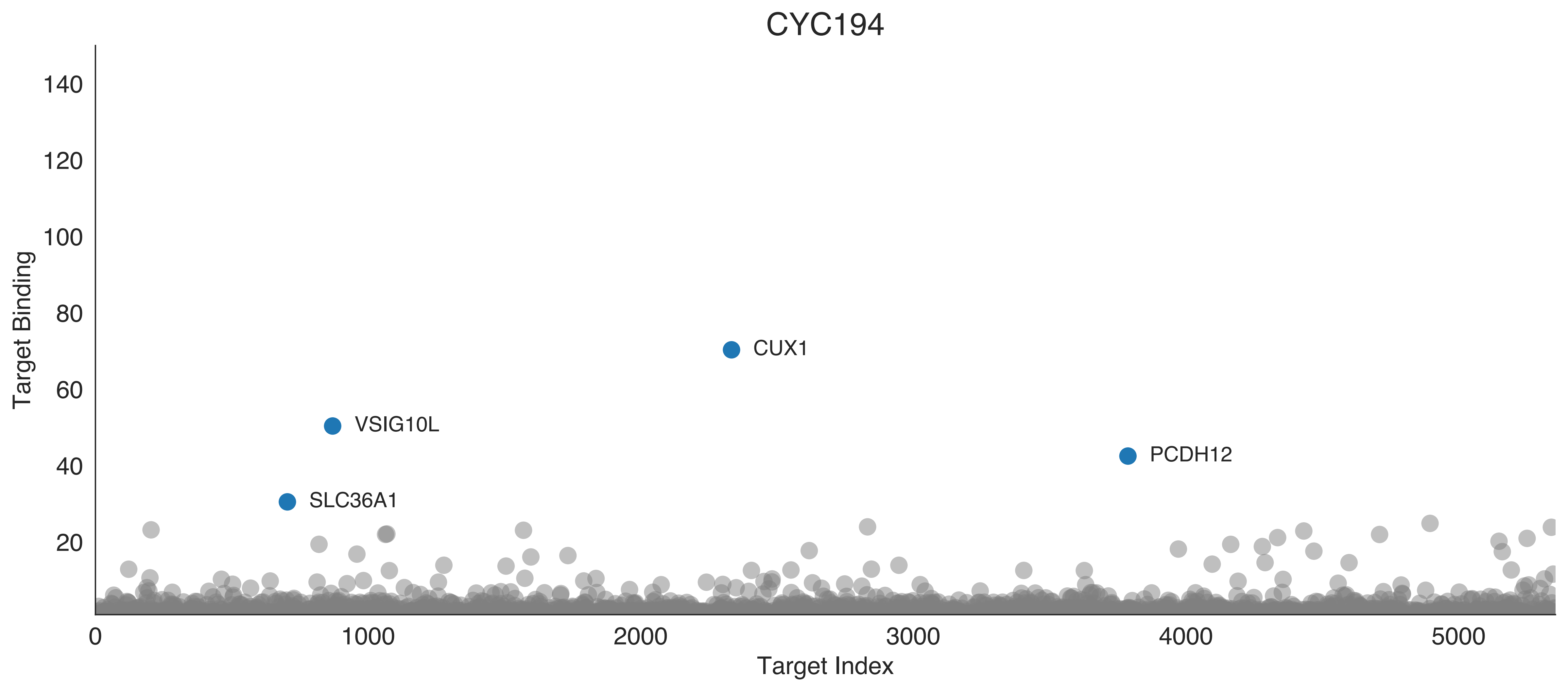
CYC194
Target Binding
140
120
100
80
60
40
20
CUX1
VSIG10L
PCDH12
SLC36A1
0
1000
2000
3000
4000
5000
Target Index

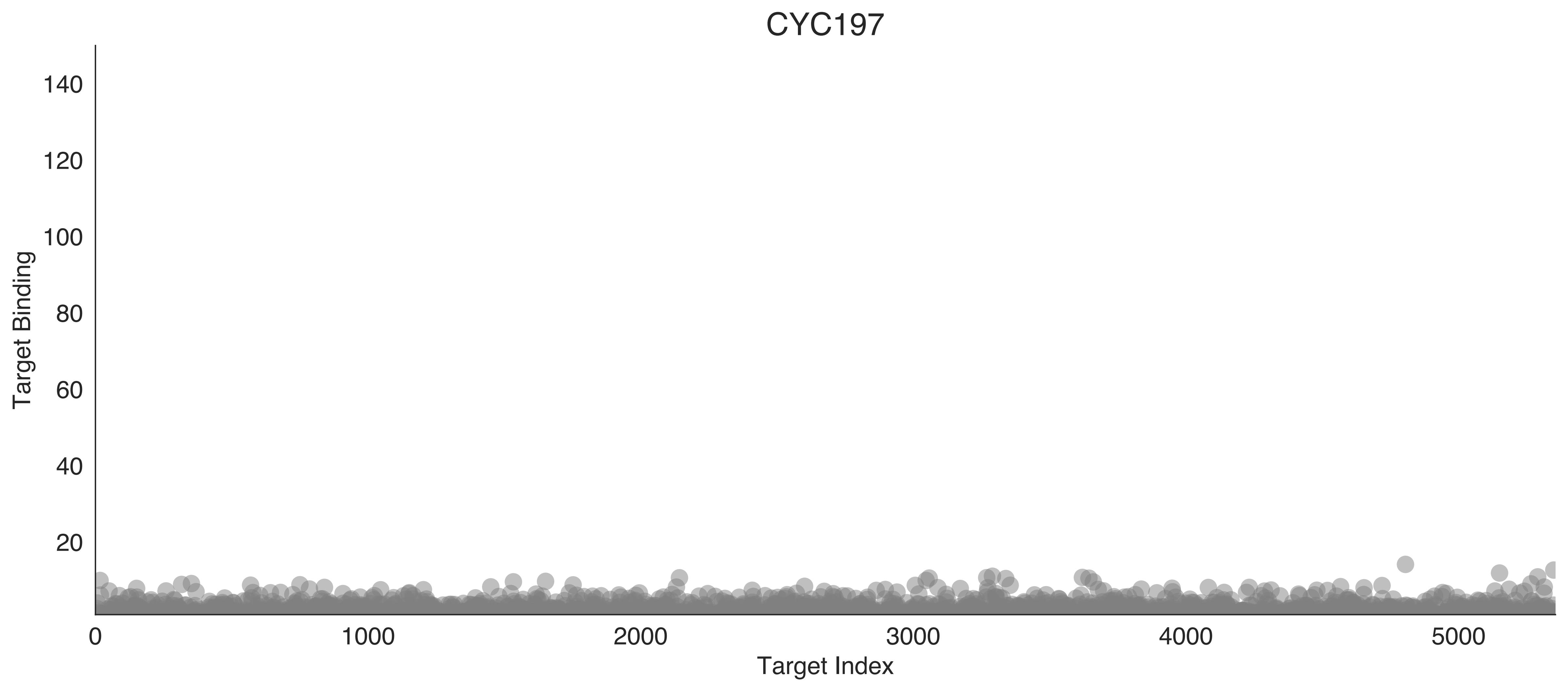

CYC197
Target Binding
Target Index
140
120
100
80
60
40
20
0
1000
2000
3000
4000
5000

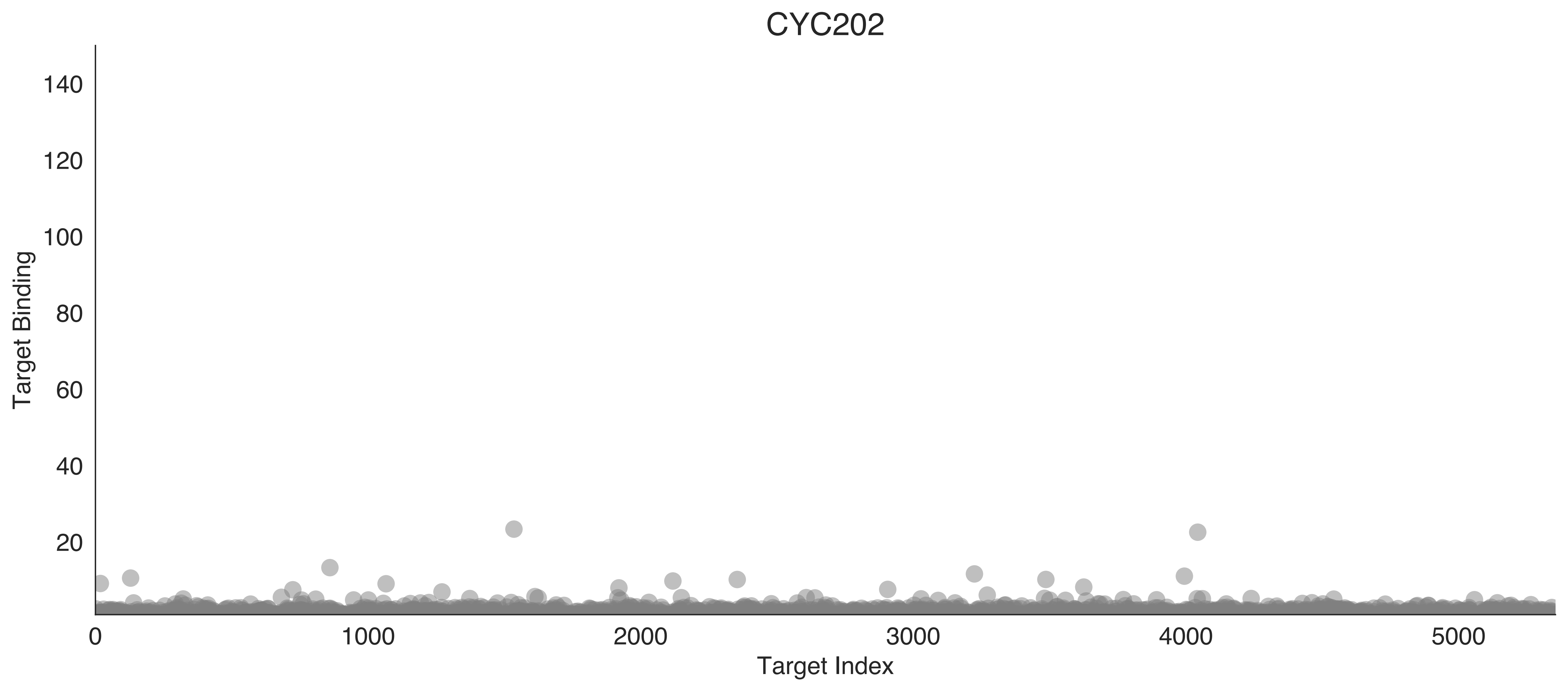

CYC202
Target Binding
Target Index
0
20
40
60
80
100
120
140
1000
2000
3000
4000
5000

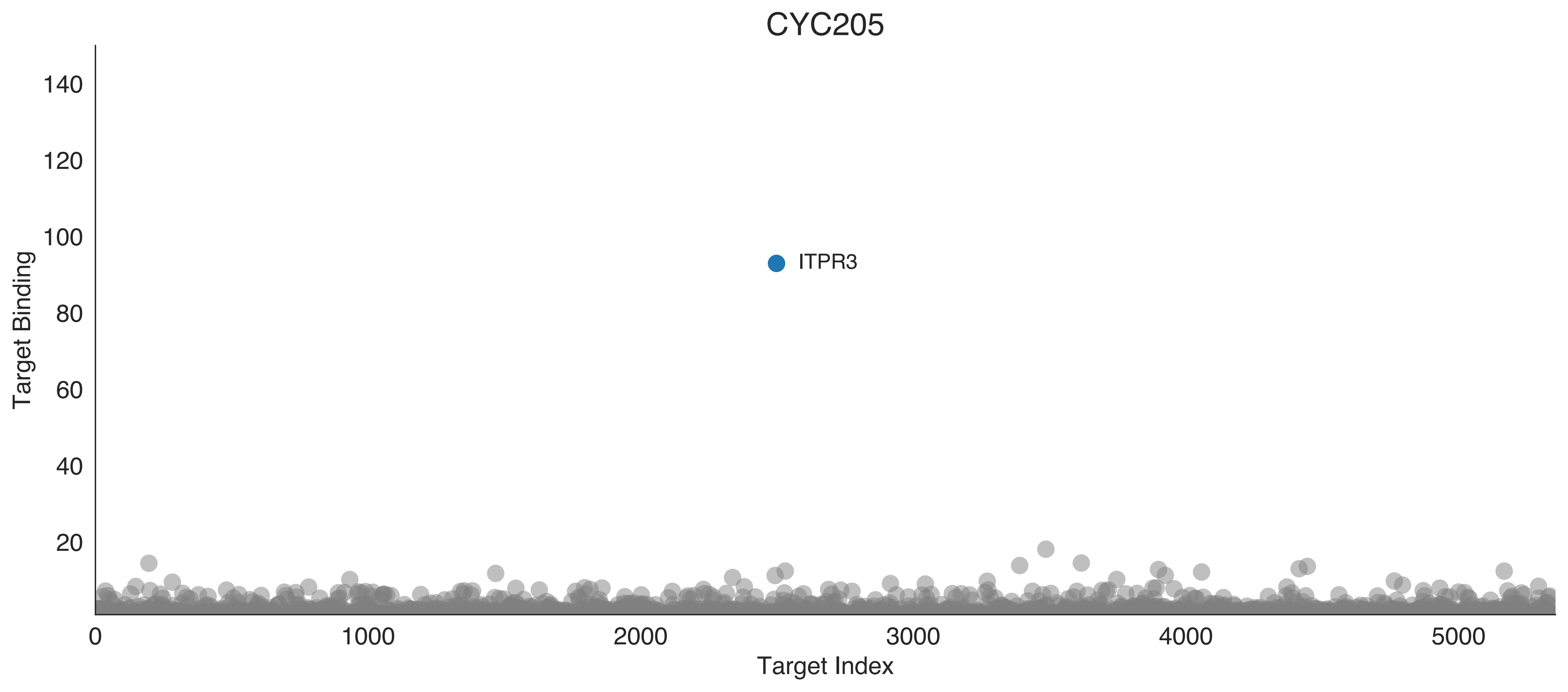
CYC205
Target Binding
140
120
100
80
60
40
20
ITPR3
0
1000
2000
3000
4000
5000
Target Index

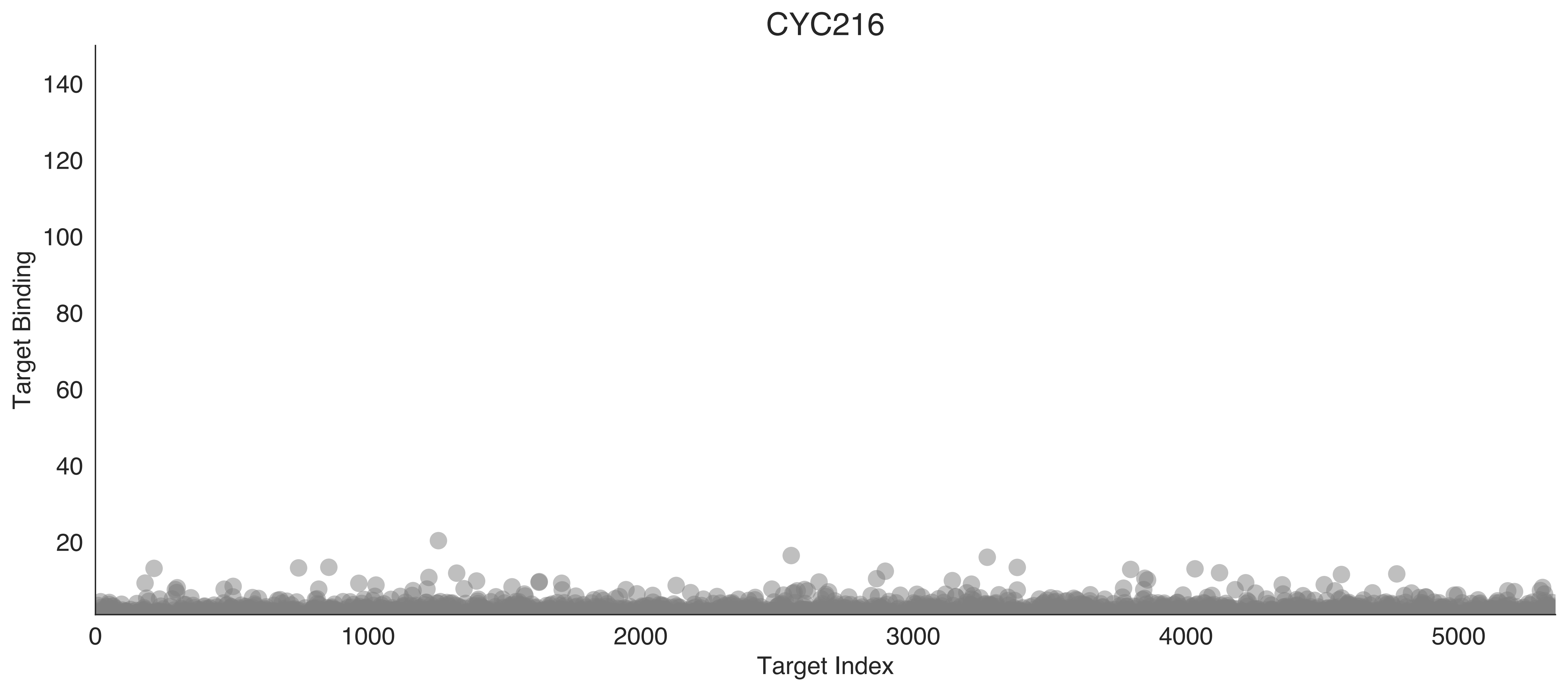

CYC216
Target Binding
Target Index
0
20
40
60
80
100
120
140
0
1000
2000
3000
4000
5000

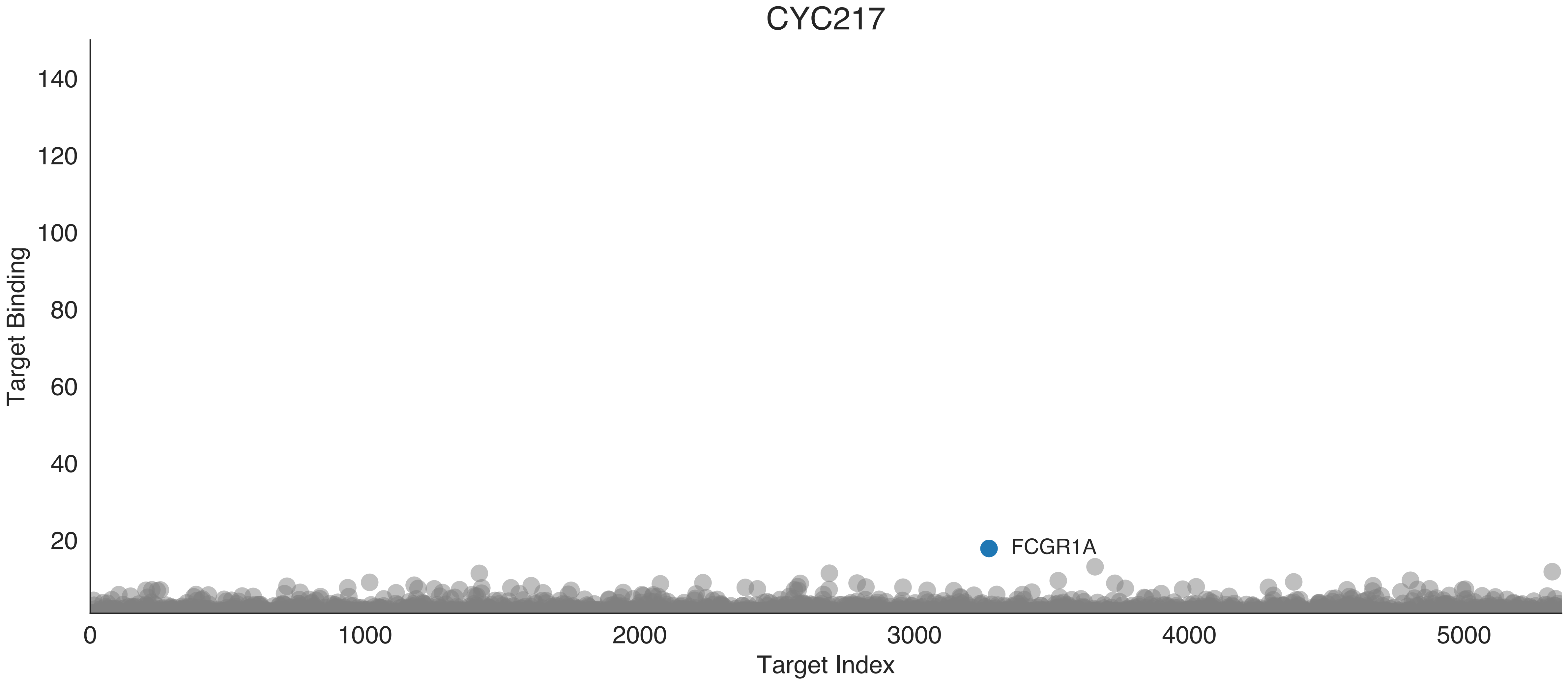

CYC217
Target Binding
140
120
100
80
60
40
20
FCGR1A
0
1000
2000
3000
4000
5000
Target Index

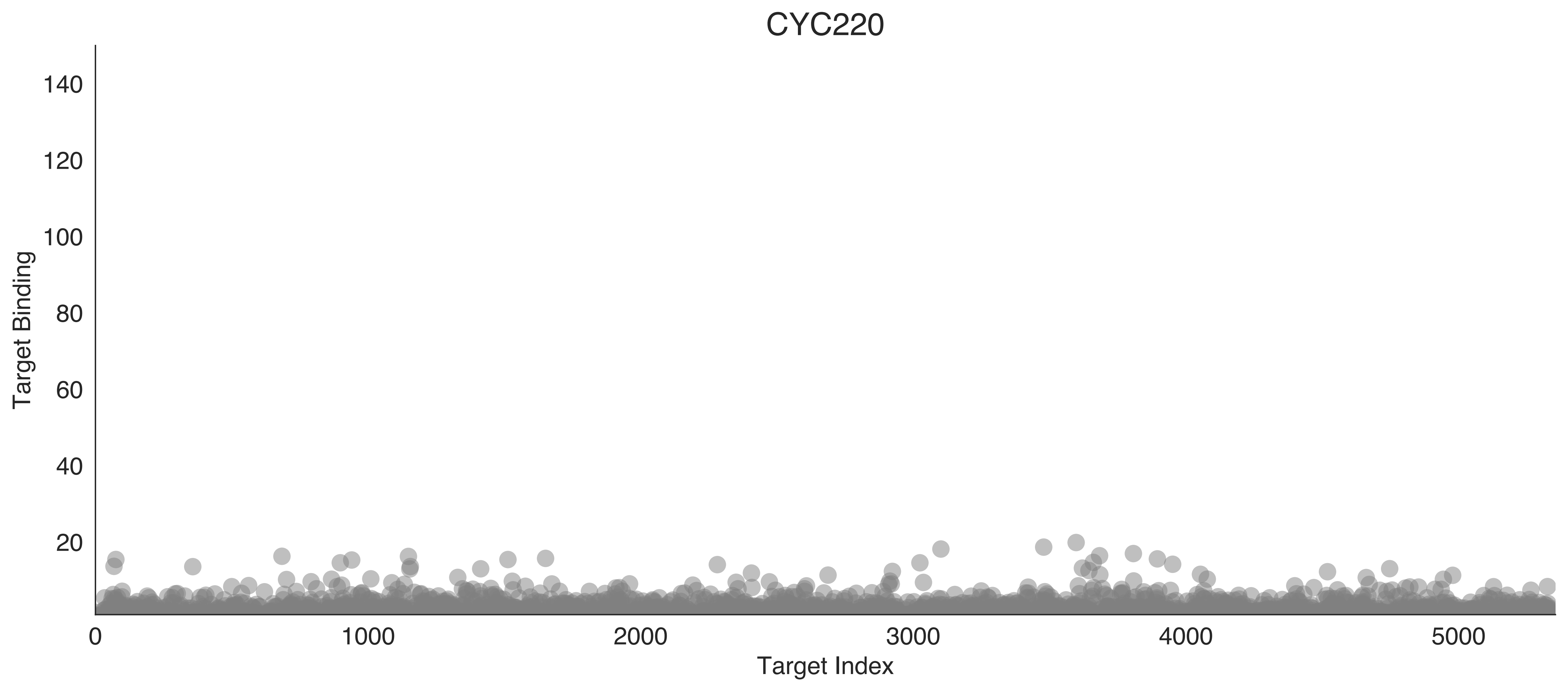
CYC220
Target Binding
140
120
100
80
60
40
20
0
1000
2000
3000
4000
5000
Target Index

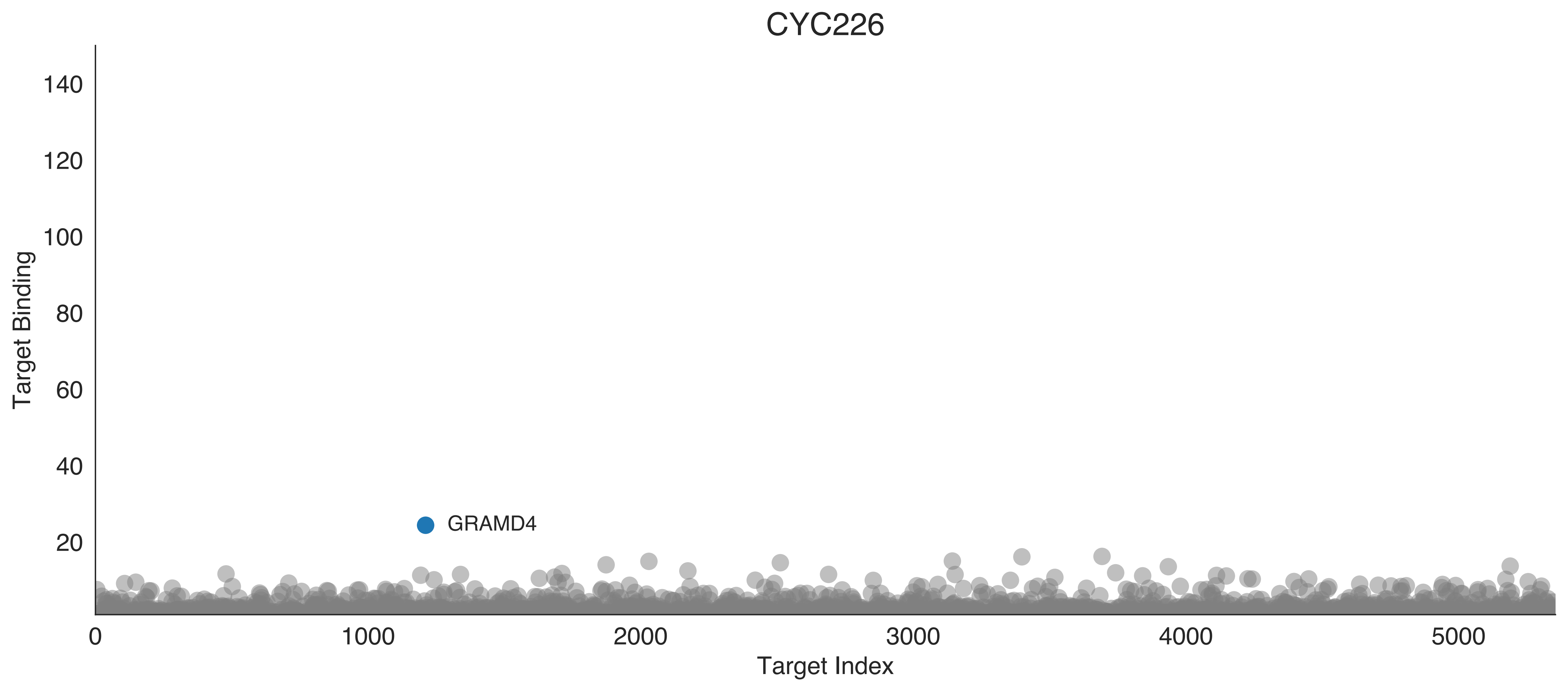
CYC226
Target Binding
140
120
100
80
60
40
20
GRAMD4
0
1000
2000
3000
4000
5000
Target Index

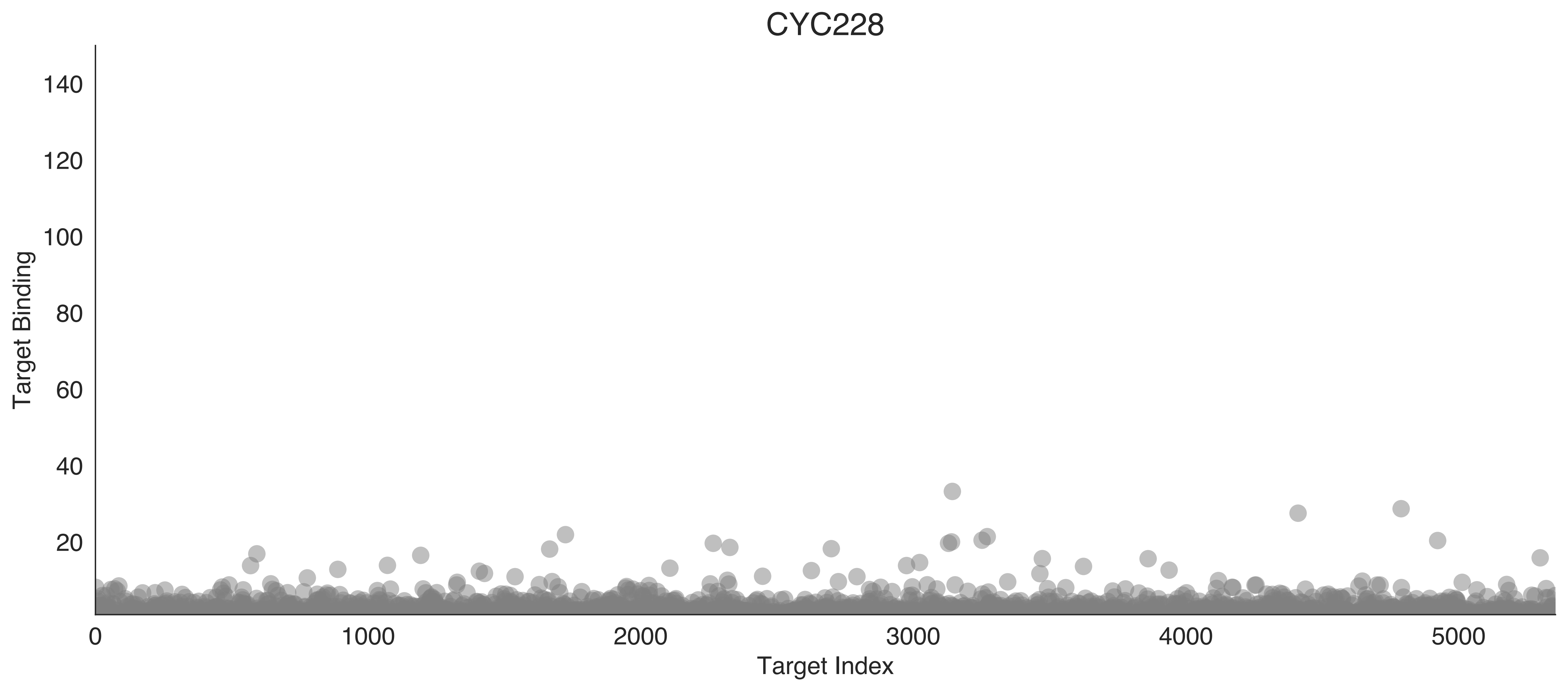
CYC228
Target Binding
140
120
100
80
60
40
20
0
1000
2000
3000
4000
5000
Target Index

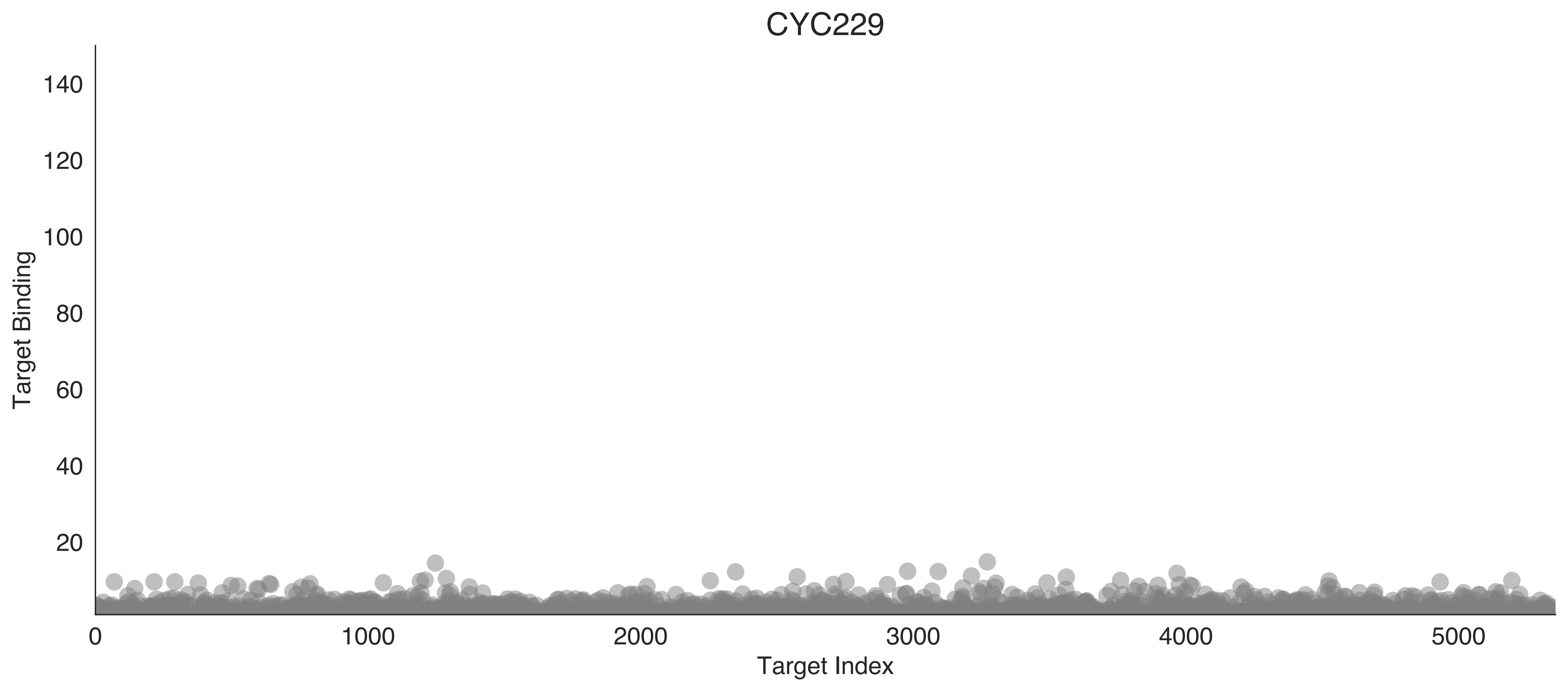

CYC229
Target Binding
140
120
100
80
60
40
20
0
1000
2000
3000
4000
5000
Target Index

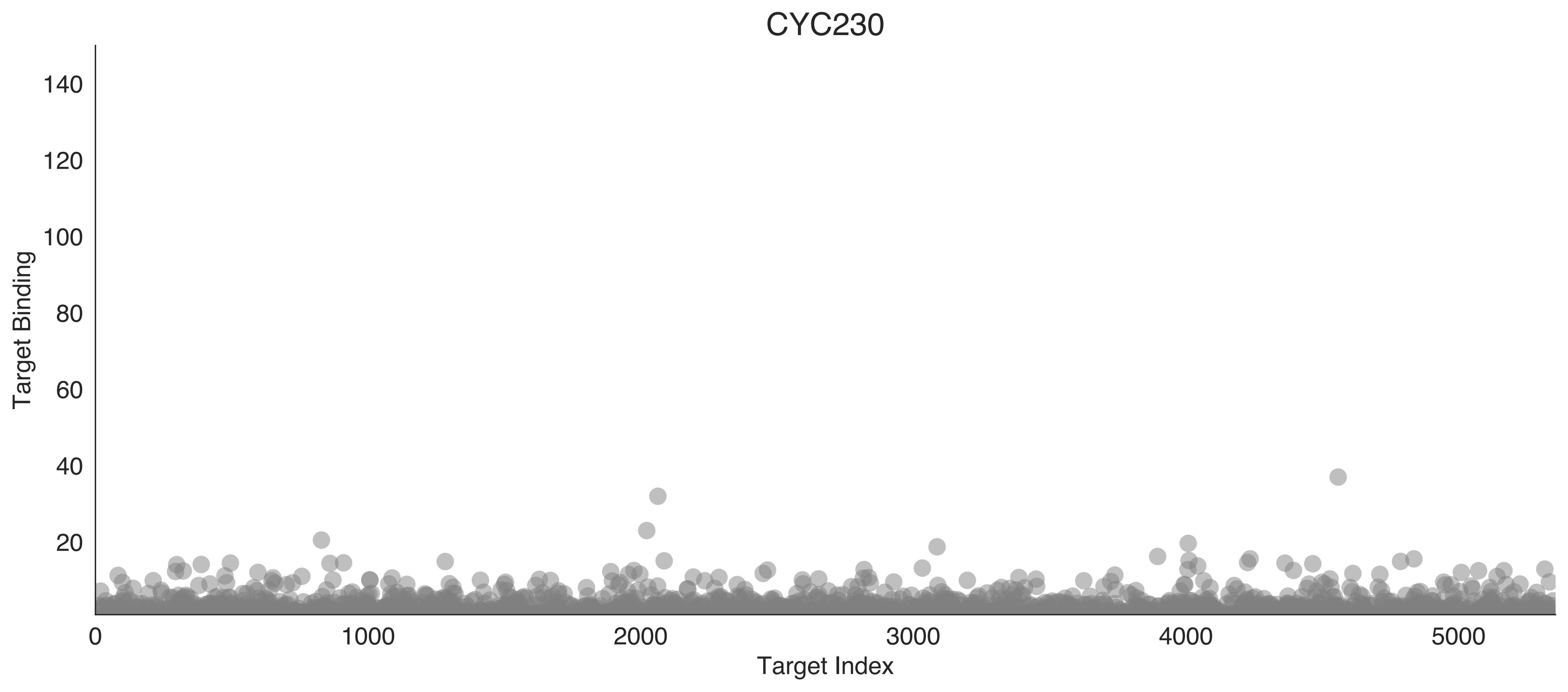

CYC230
Target Binding
Target Index
140
120
100
80
60
40
20
0
1000
2000
3000
4000
5000

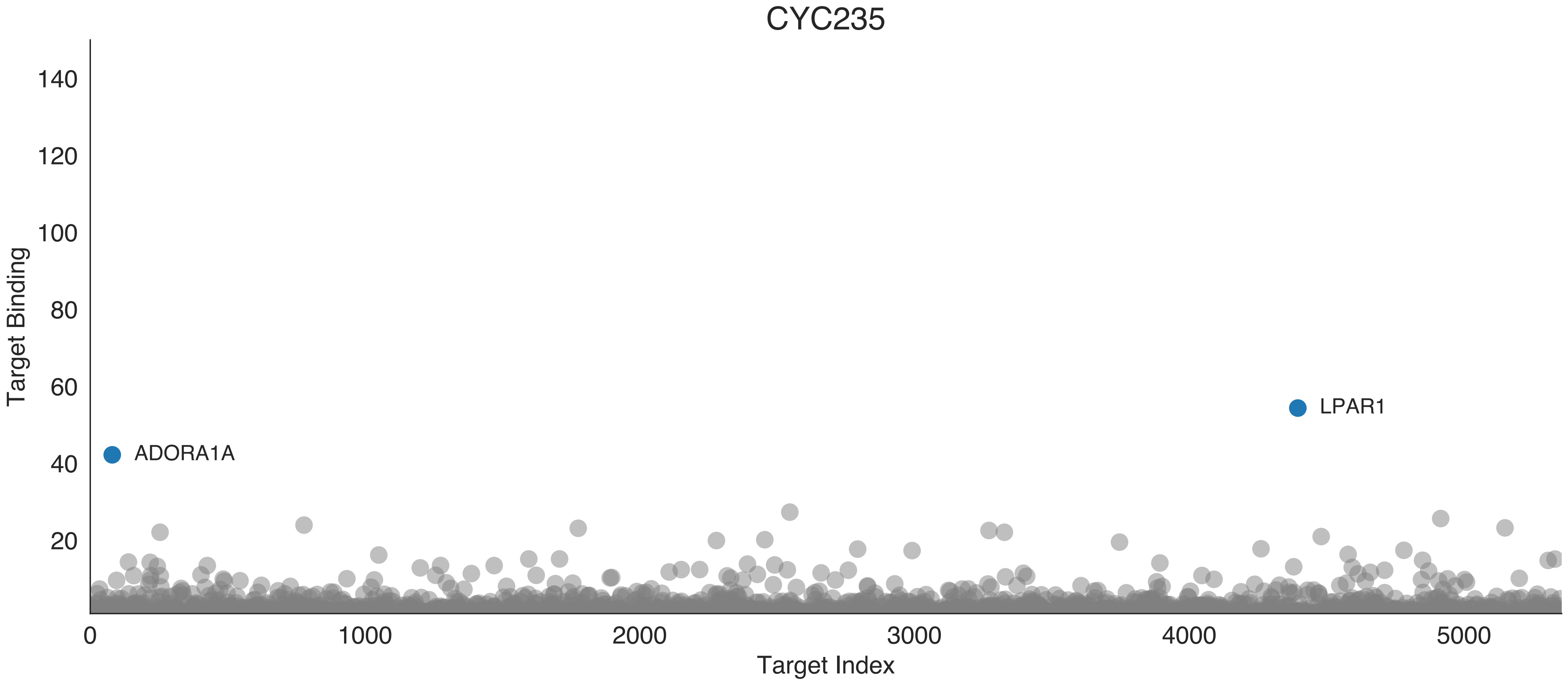
CYC235
Target Binding
140
120
100
80
60
40
20
0
1000
2000
3000
4000
5000
Target Index
ADORA1A
LPAR1

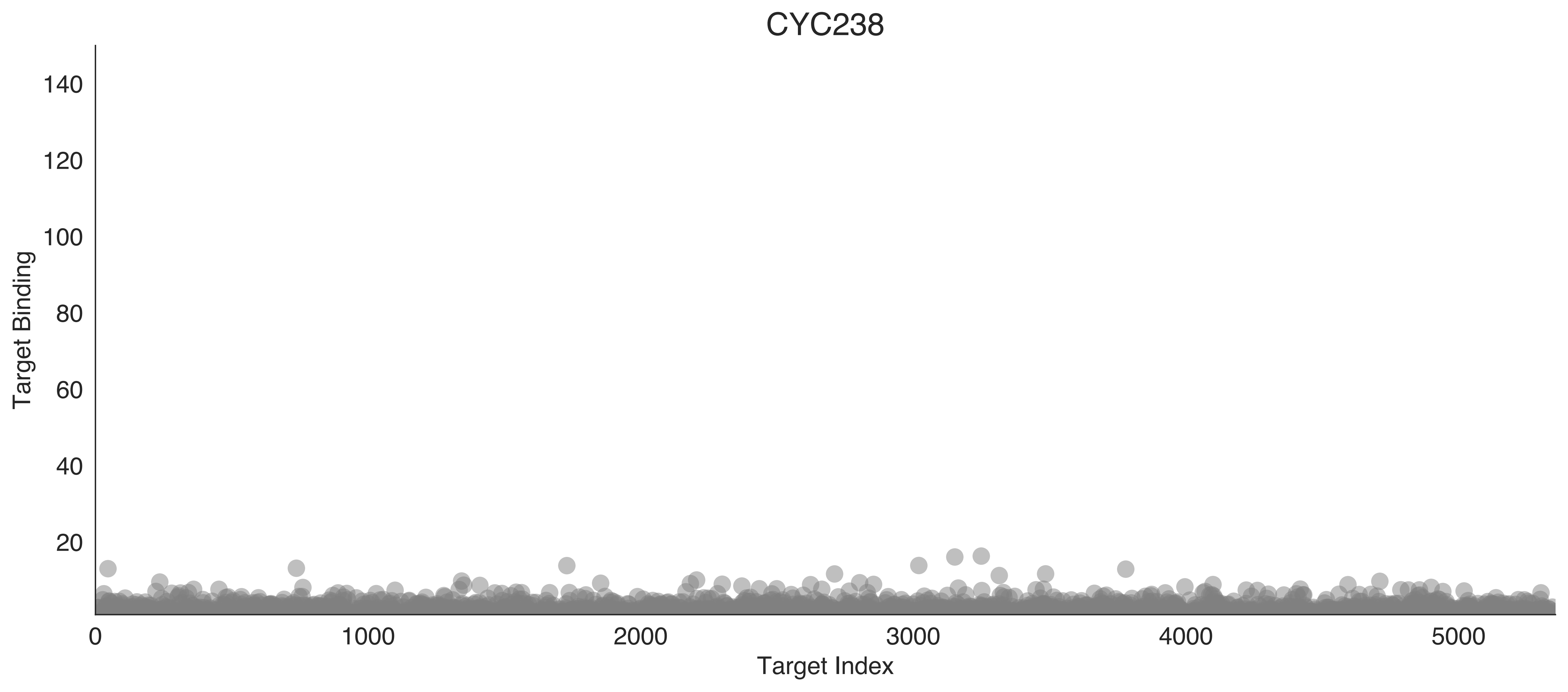
CYC238
Target Binding
Target Index
0
20
40
60
80
100
120
140
1000
2000
3000
4000
5000

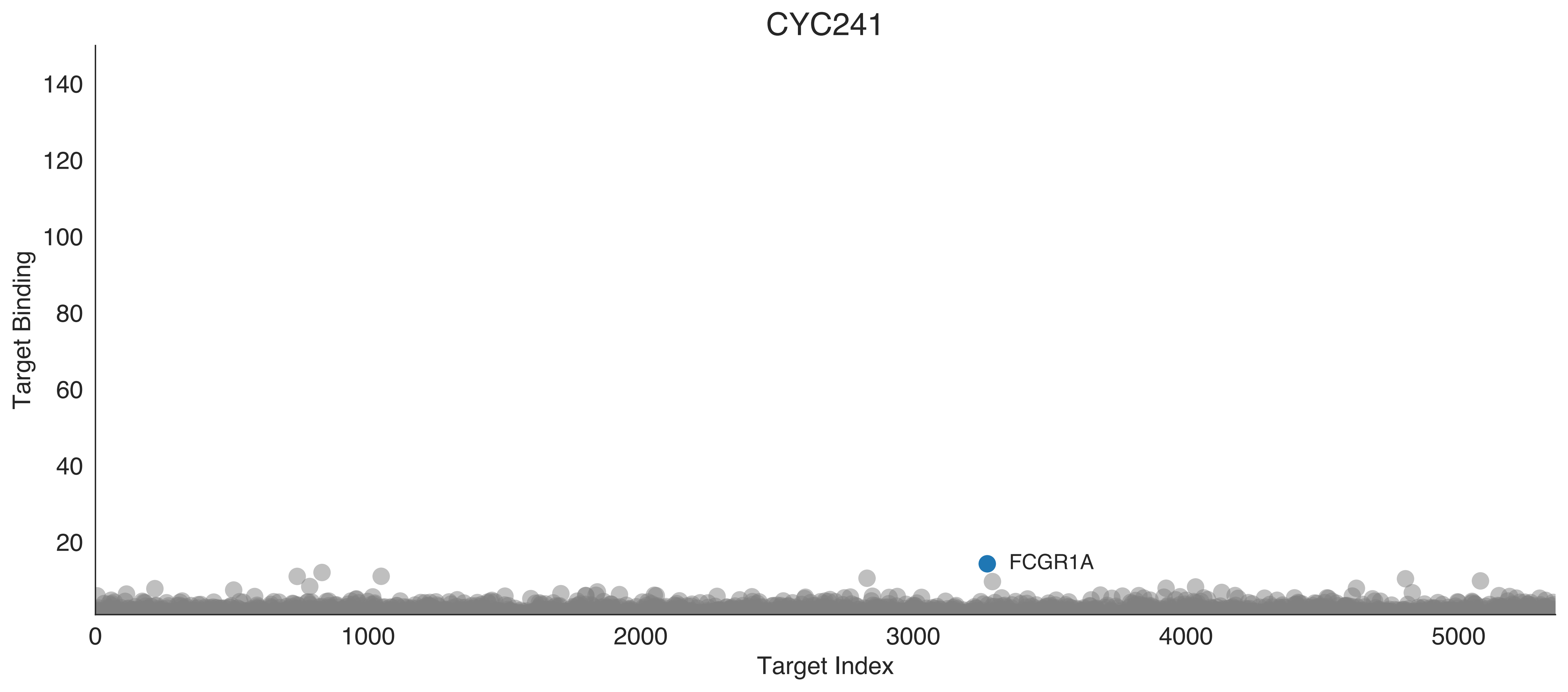
CYC241
Target Binding
Target Index
FCGR1A
140
120
100
80
60
40
20
0
1000
2000
3000
4000
5000

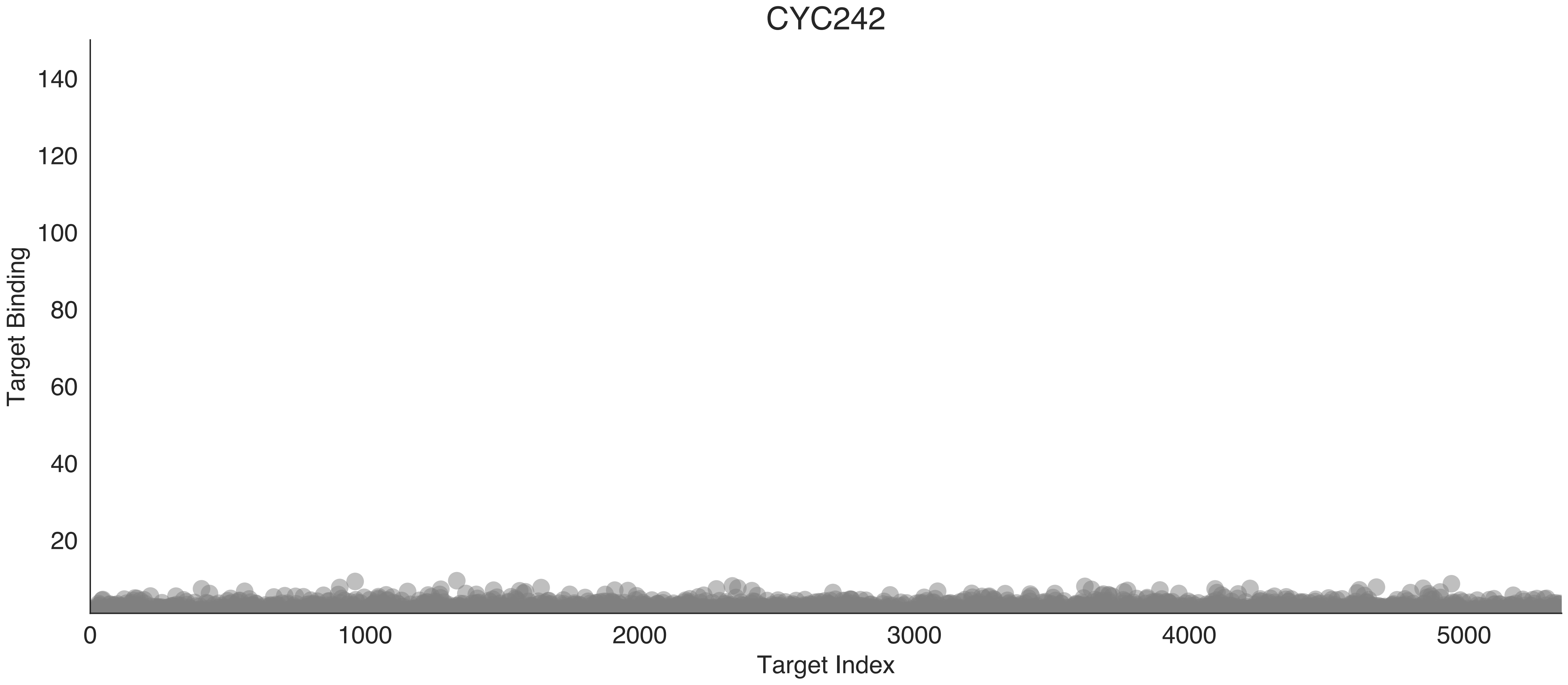

CYC242
Target Binding
140
120
100
80
60
40
20
0
1000
2000
3000
4000
5000
Target Index

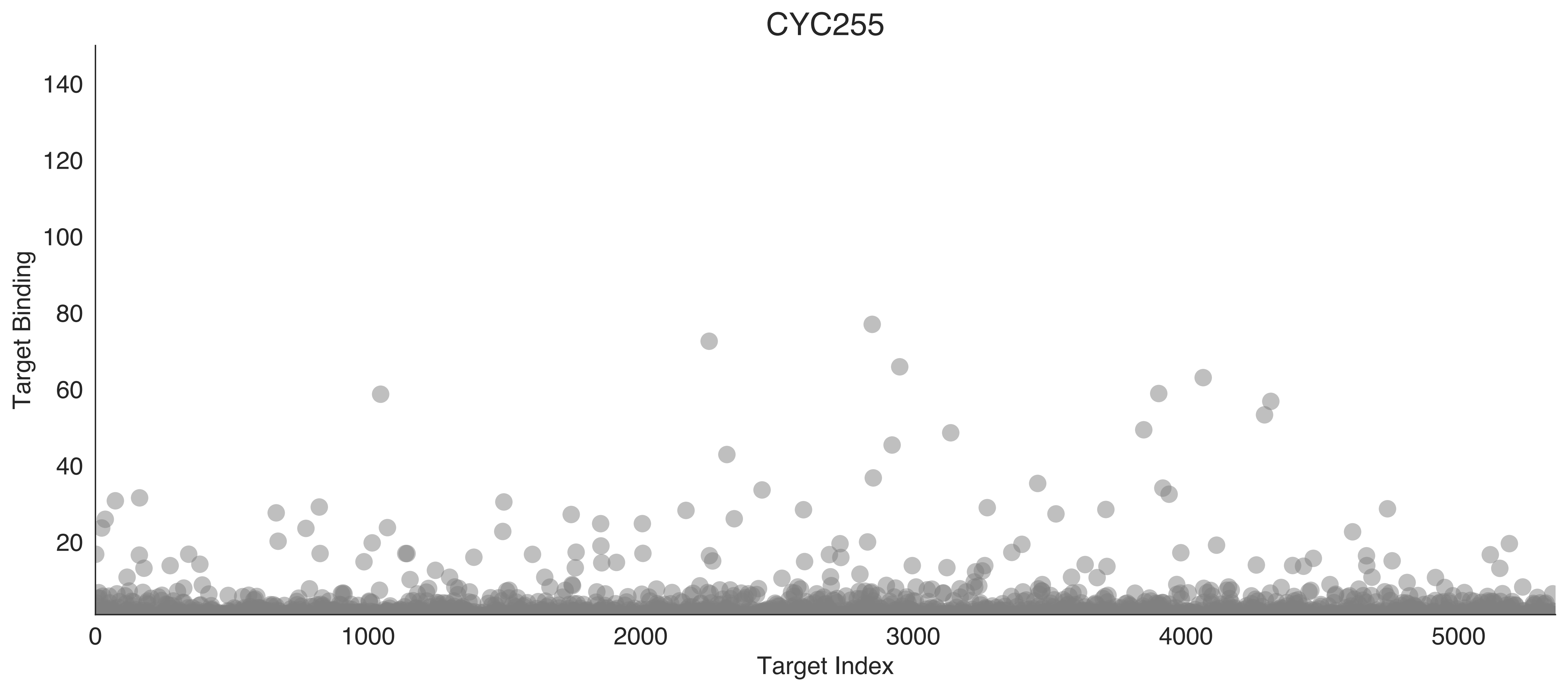

CYC255
Target Binding
Target Index
0
20
40
60
80
100
120
140
1000
2000
3000
4000
5000

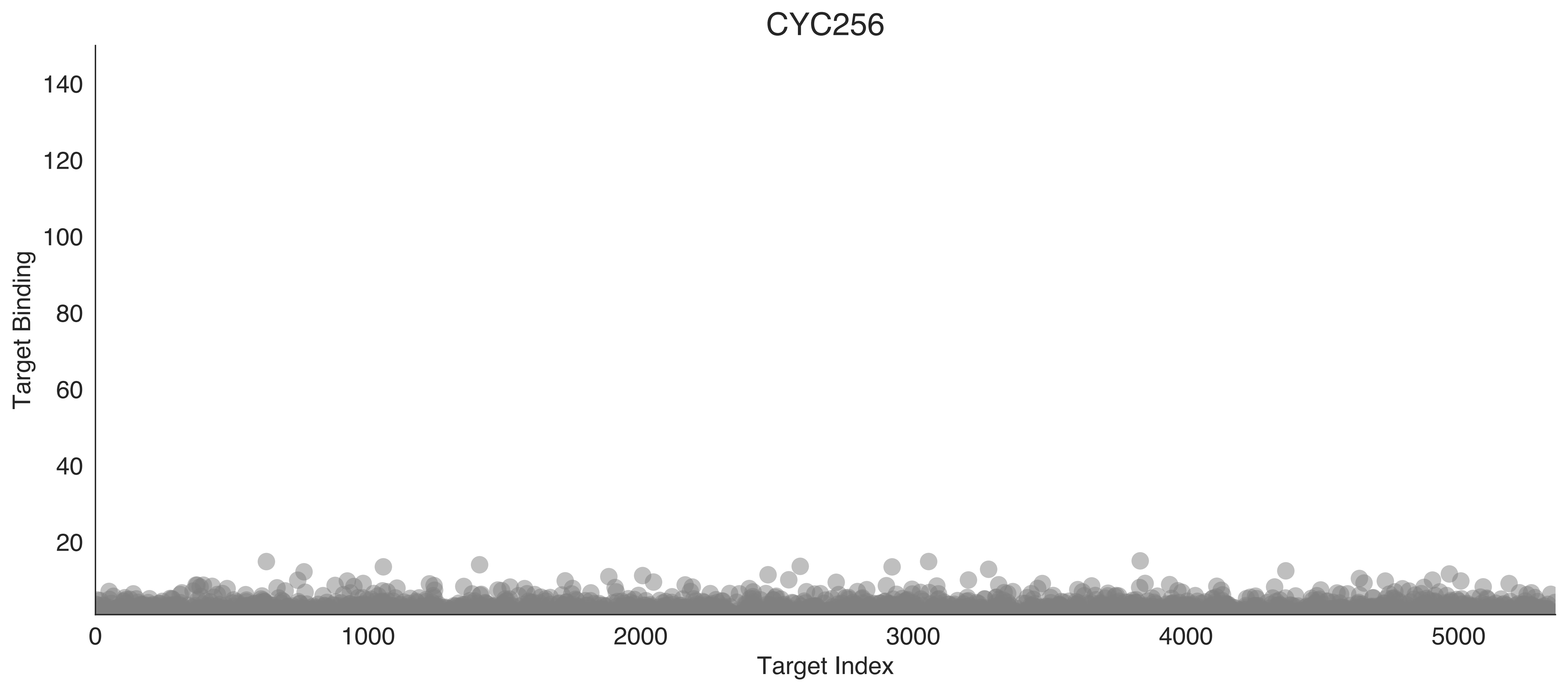
CYC256
Target Binding
Target Index
140
120
100
80
60
40
20
0
1000
2000
3000
4000
5000

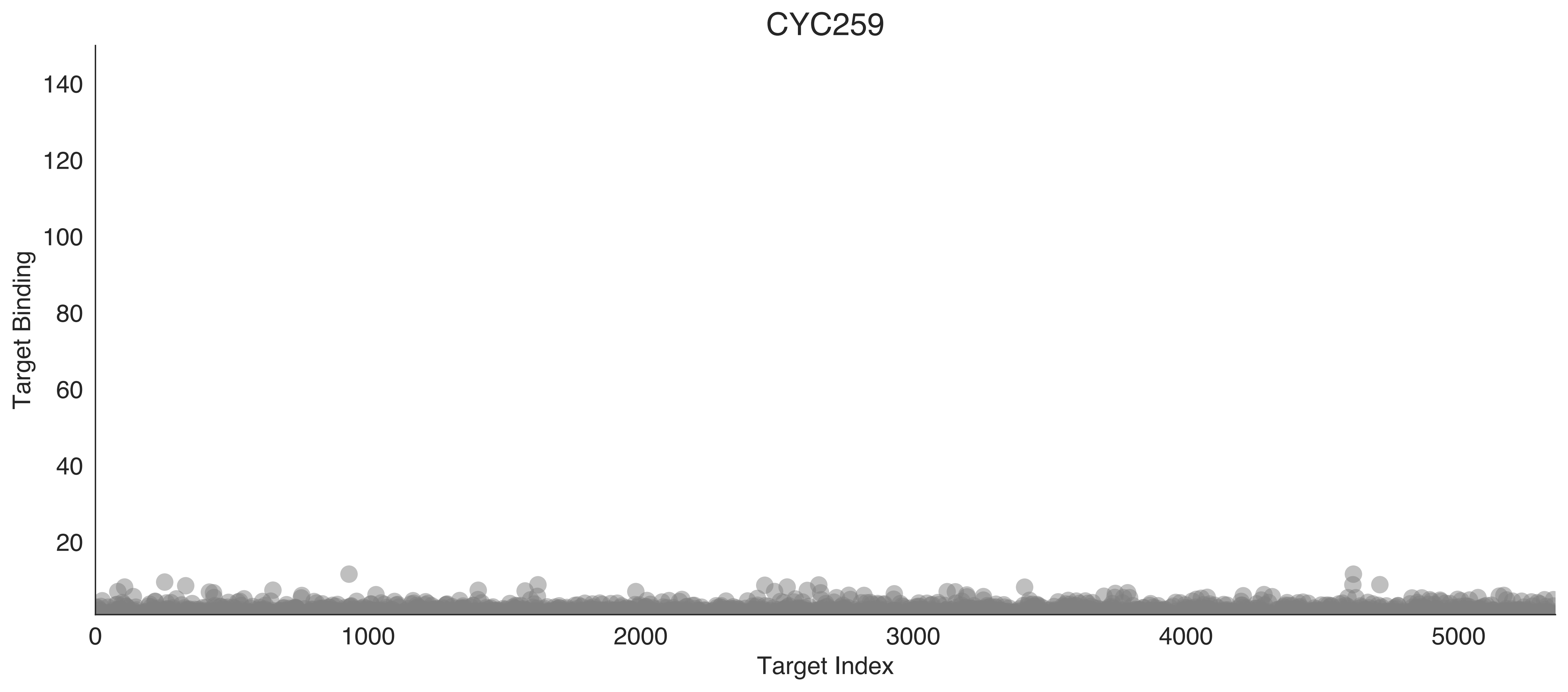
CYC259
Target Binding
Target Index
140
120
100
80
60
40
20
0
1000
2000
3000
4000
5000

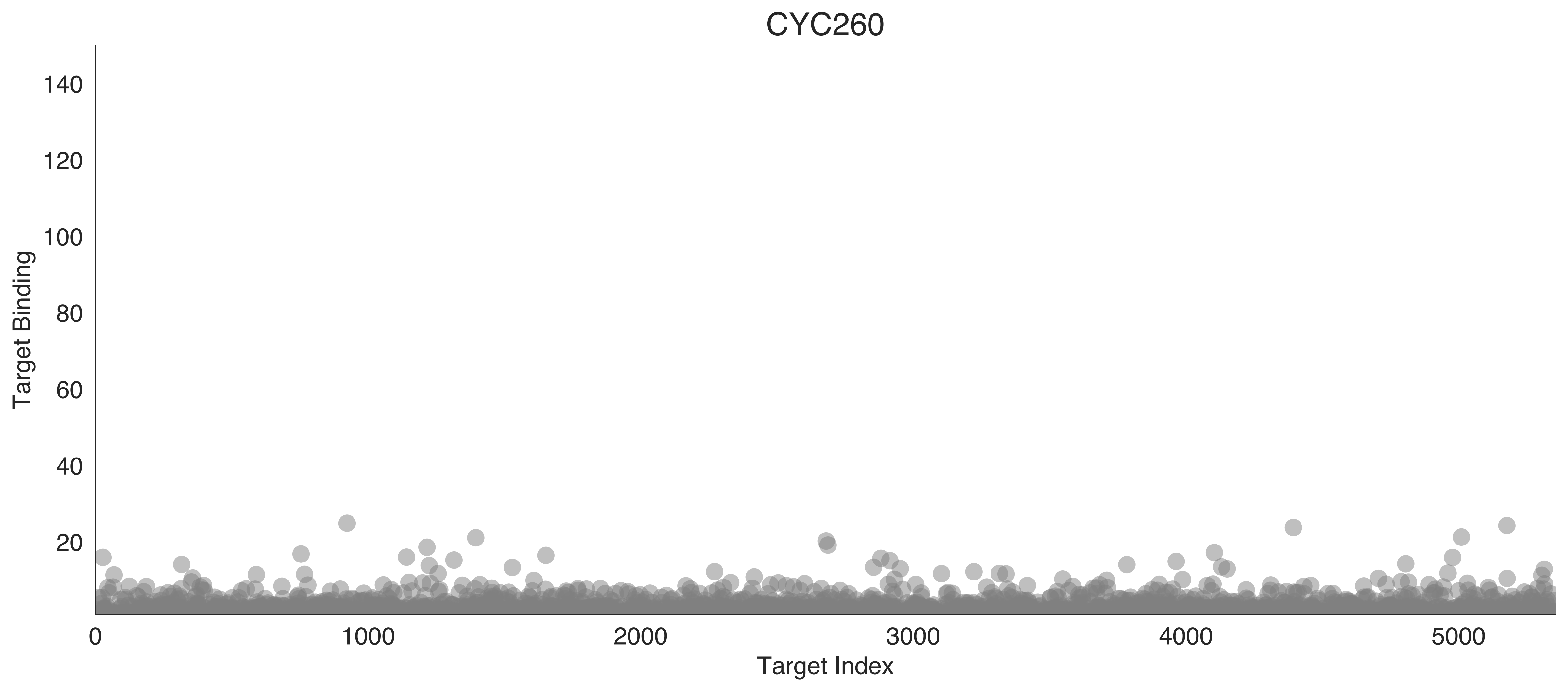
CYC260
Target Binding
140
120
100
80
60
40
20
0
1000
2000
3000
4000
5000
Target Index

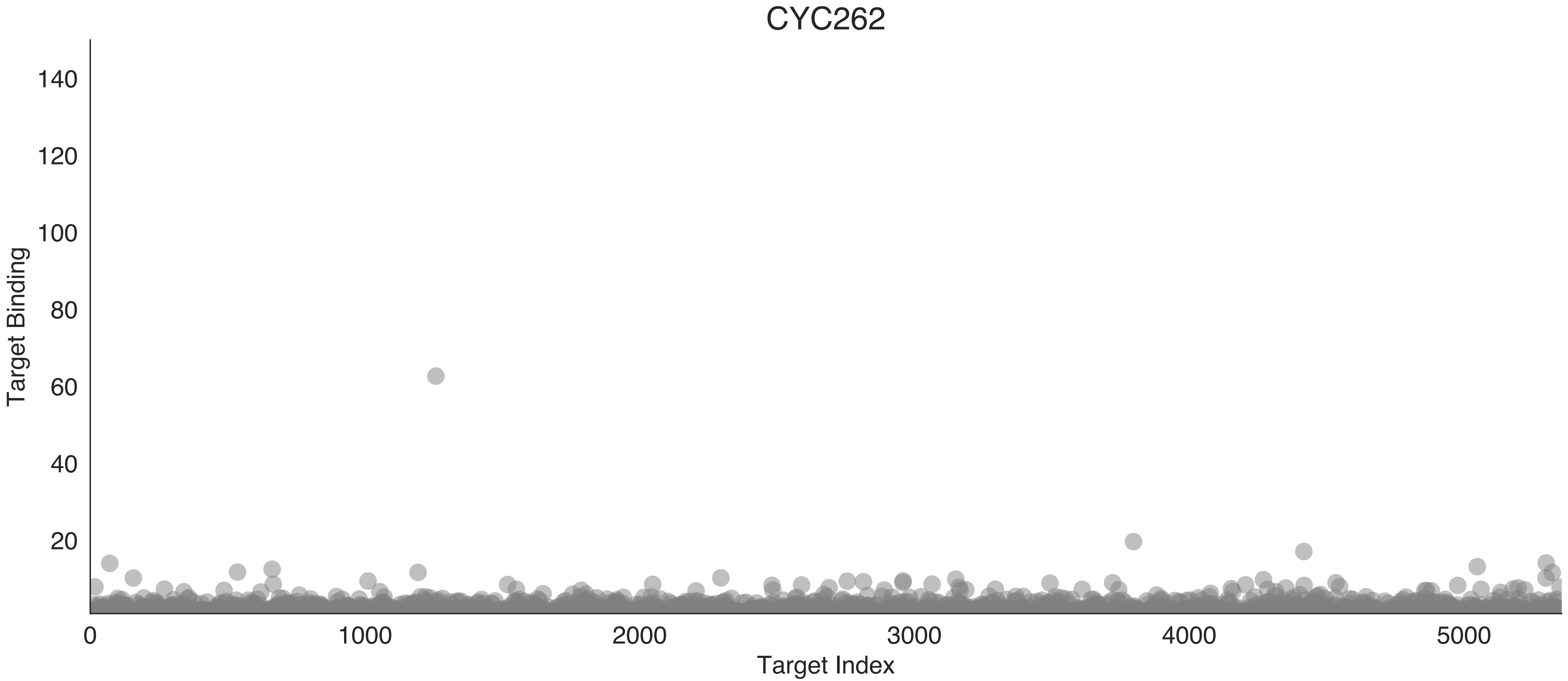
CYC262
Target Binding
140
120
100
80
60
40
20
0
1000
2000
3000
4000
5000
Target Index

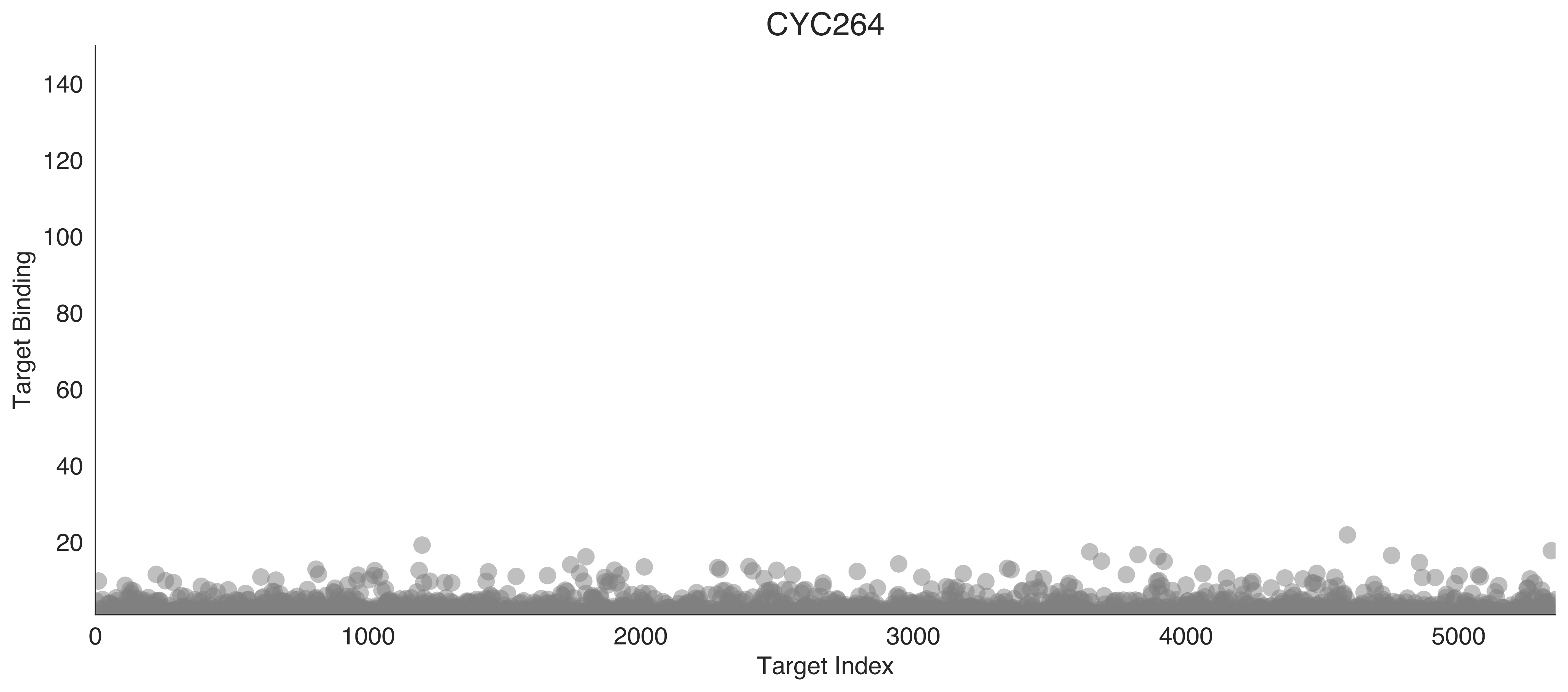
CYC264
Target Binding
Target Index
0
20
40
60
80
100
120
140
1000
2000
3000
4000
5000

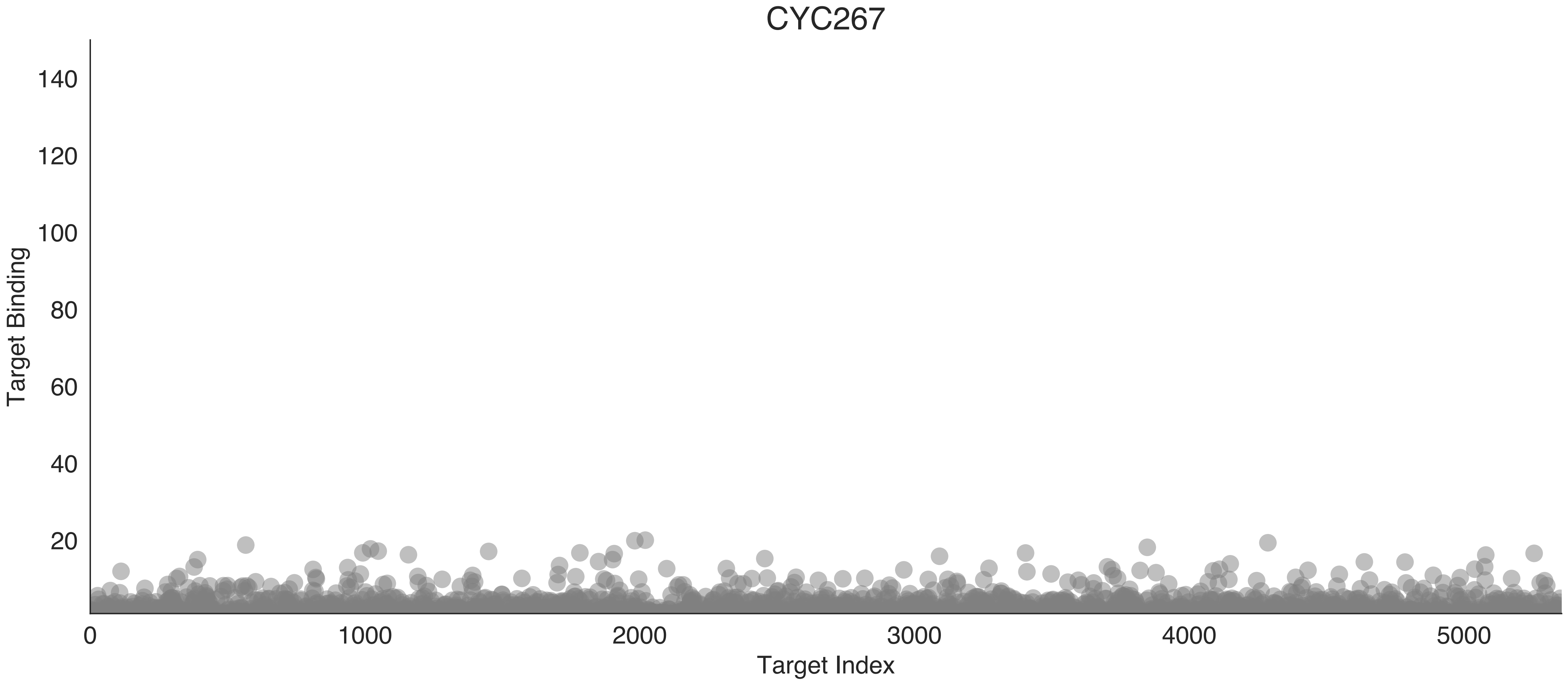

CYC267
Target Binding
140
120
100
80
60
40
20
0
1000
2000
3000
4000
5000
Target Index

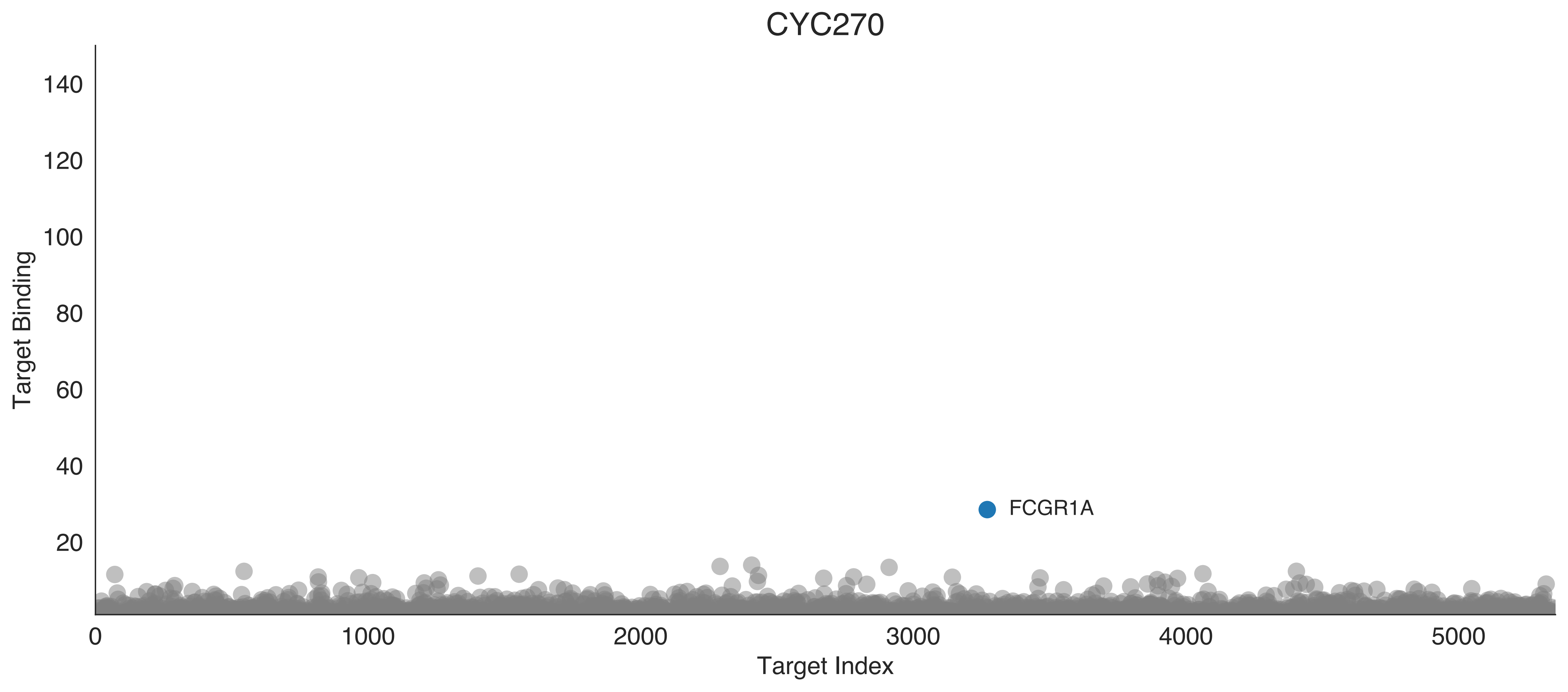
CYC270
Target Binding
Target Index
140
120
100
80
60
40
20
0
1000
2000
3000
4000
5000
FCGR1A

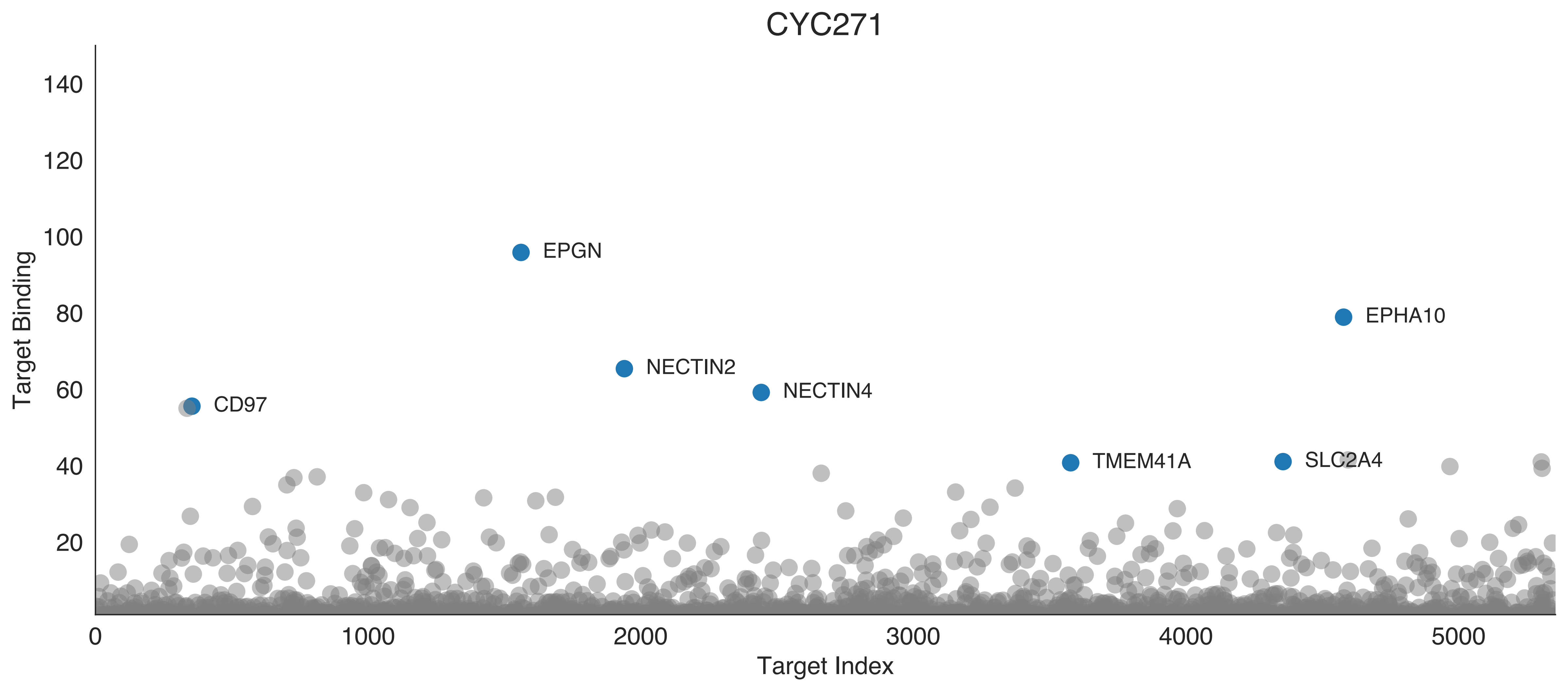

CYC271
Target Binding
Target Index
EPGN
EPHA10
NECTIN2
NECTIN4
CD97
TMEM41A
SLC2A4
0
20
40
60
80
100
120
140
1000
2000
3000
4000
5000

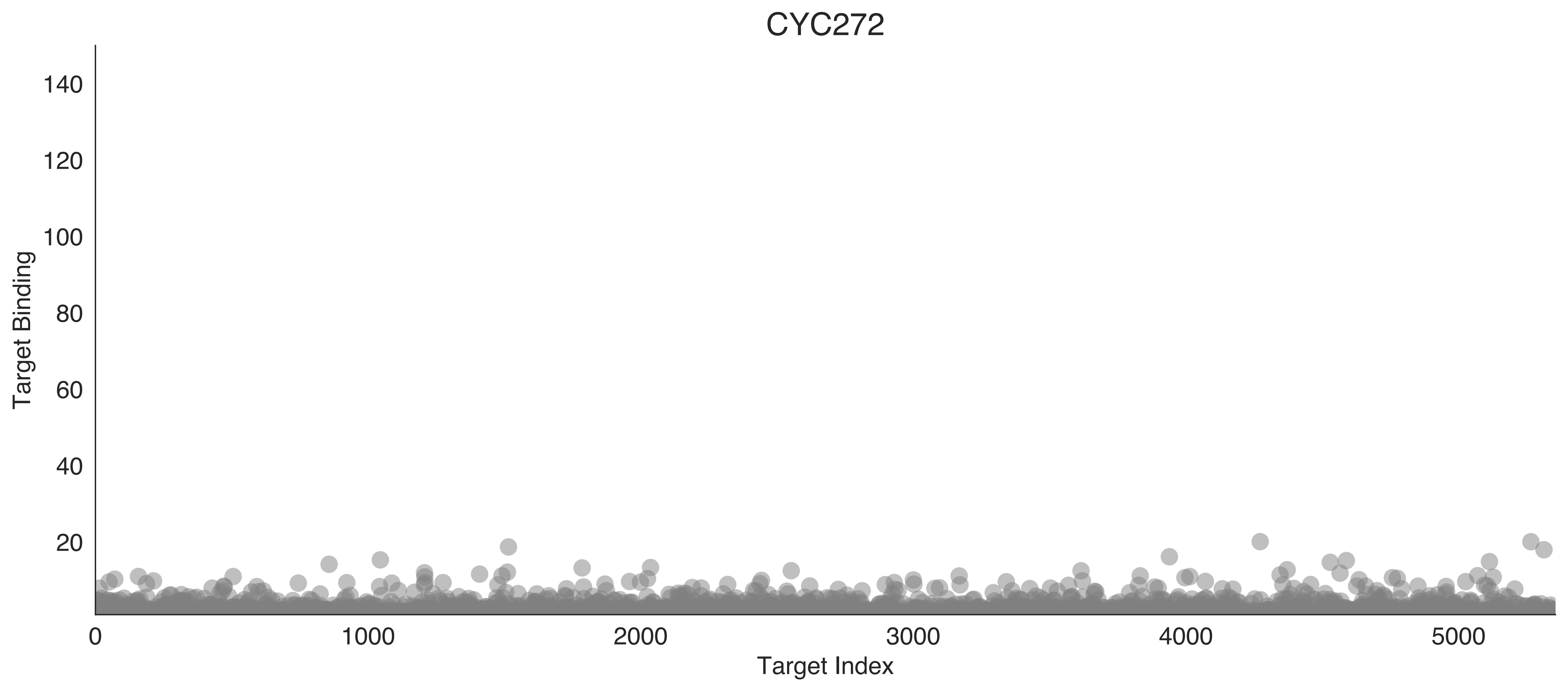

CYC272
Target Binding
Target Index
0
20
40
60
80
100
120
140
1000
2000
3000
4000
5000

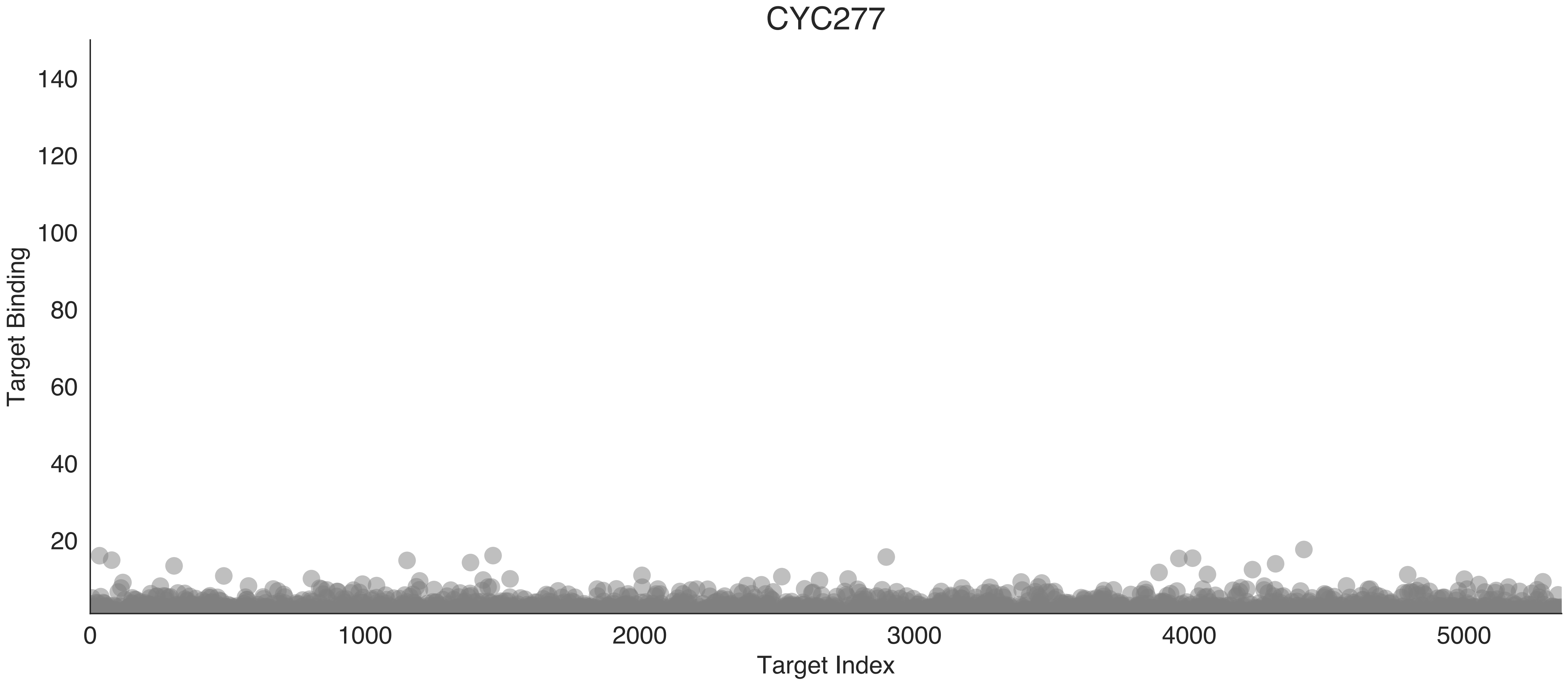

CYC277
Target Binding
140
120
100
80
60
40
20
0
1000
2000
3000
4000
5000
Target Index

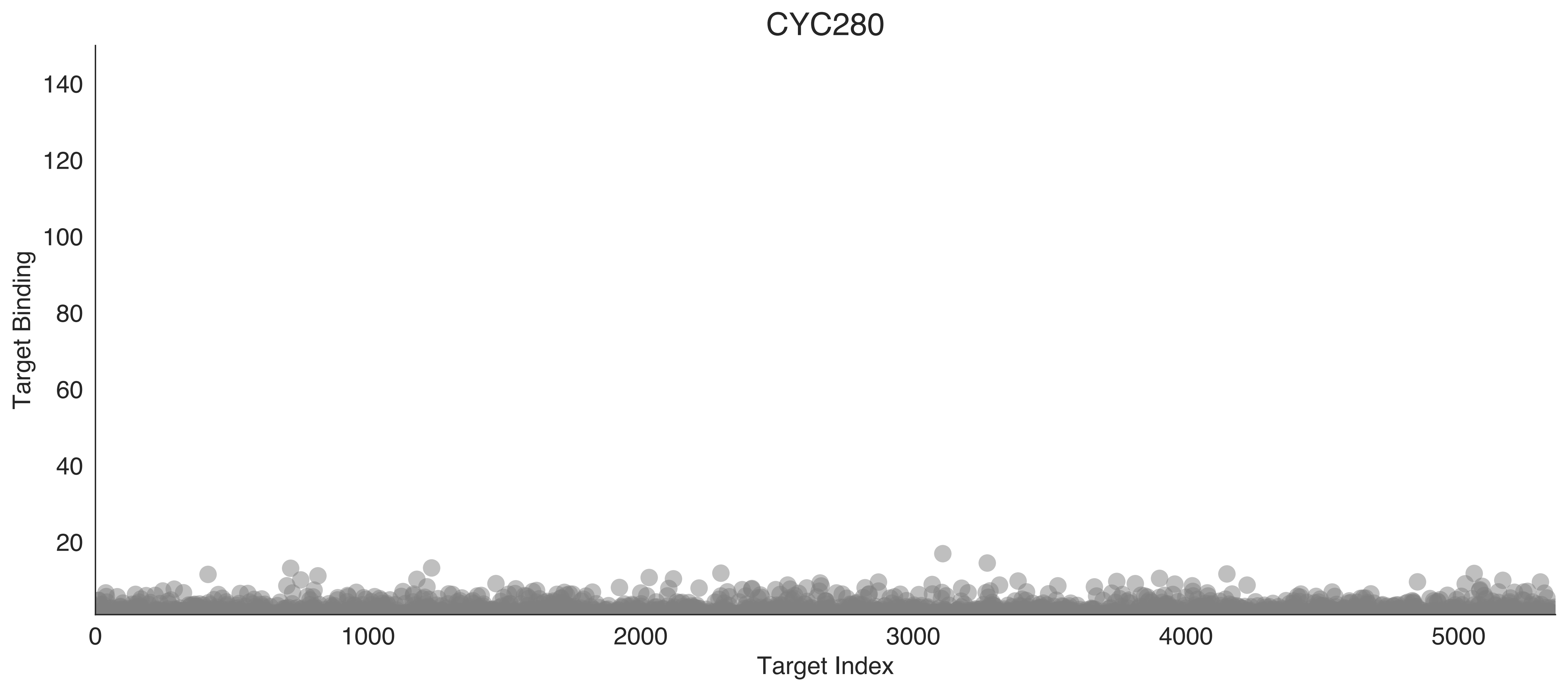
CYC280
Target Binding
140
120
100
80
60
40
20
0
1000
2000
3000
4000
5000
Target Index

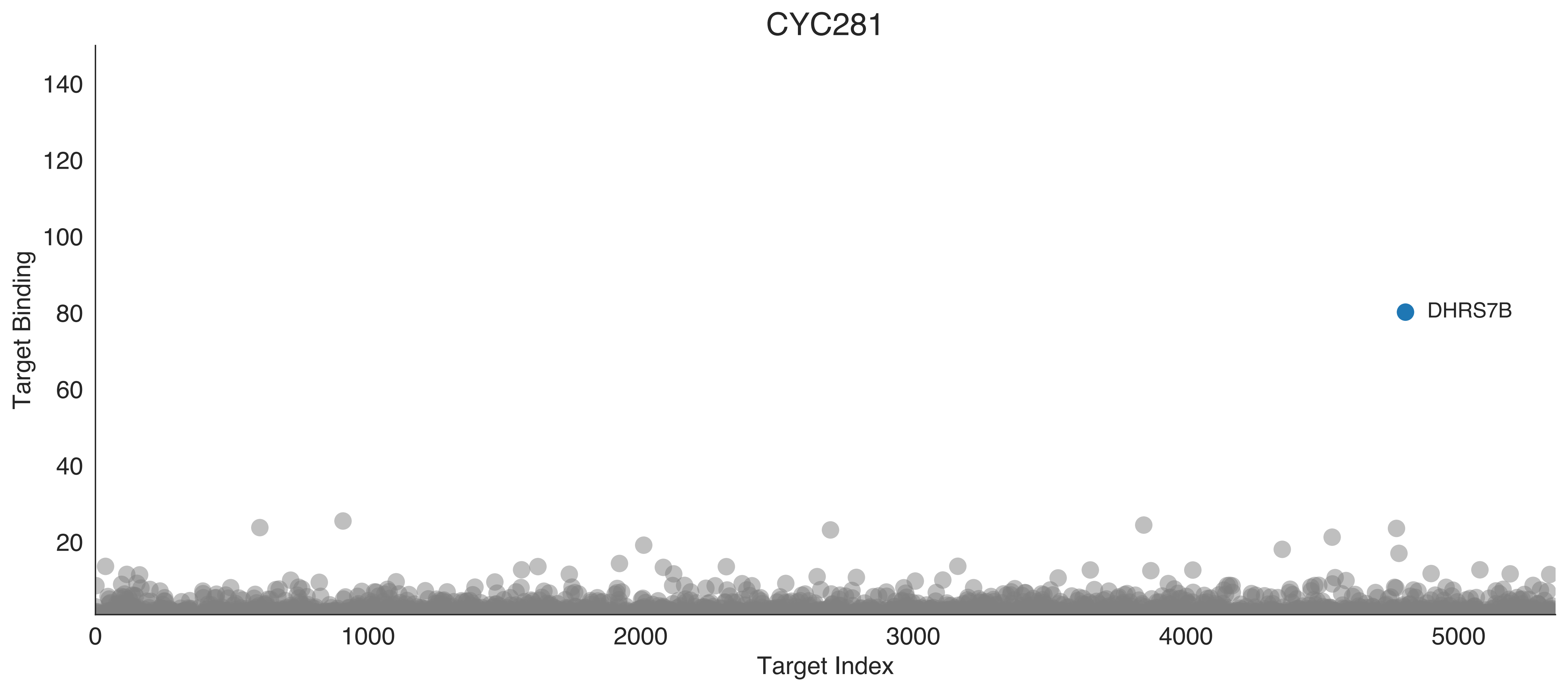

CYC281
Target Binding
140
120
100
80
60
40
20
DHRS7B
0
1000
2000
3000
4000
5000
Target Index
